# Pilot study: Trehalose-induced remodelling of the human microbiota affects *Clostridioides difficile* infection outcome in an *in vitro* colonic model

**DOI:** 10.1101/2021.02.16.431513

**Authors:** Anthony M. Buckley, Ines B. Moura, Norie Arai, William Spittal, Emma Clark, Yoshihiro Nishida, Hannah C. Harris, Karen Bentley, Georgina Davis, Dapeng Wang, Suparna Mitra, Takanobu Higashiyama, Mark H. Wilcox

## Abstract

Within the human intestinal tract, dietary, microbial- and host-derived compounds are used as signals by many pathogenic organisms, including *Clostridioides difficile*. Trehalose has been reported to enhance virulence of certain *C. difficile* ribotypes; however, such variants are widespread and not correlated with clinical outcomes for patients suffering from *C. difficile* infection (CDI). Here, we make preliminary observations to how to trehalose supplementation affects the microbiota in an *in vitro* model and show that trehalose can reduce the outgrowth of *C. difficile*, preventing simulated CDI. Three clinically reflective human gut models simulated the effects of sugar (trehalose or glucose) or saline ingestion on the microbiota. Models were instilled with sugar or saline and further exposed to *C. difficile* spores. The recovery of the microbiota following antibiotic treatment and CDI induction was monitored in each model. The human microbiota remodeled to utilise the bioavailable trehalose. Clindamycin induction caused simulated CDI in models supplemented with either glucose or saline; however, trehalose supplementation did not result in CDI, although limited spore germination did occur. The absence of CDI in trehalose model was associated with enhanced abundances of *Finegoldia, Faecalibacterium* and *Oscillospira*, and reduced abundances of *Klebsiella* and *Clostridium* spp., compared with the other models. Functional analysis of the microbiota in the trehalose model revealed differences in the metabolic pathways, such as amino acid metabolism, which could be attributed to prevention of CDI. Our data show that trehalose supplementation remodelled the microbiota, which prevented simulated CDI, potentially due to enhanced recovery of nutritionally competitive microbiota against *C. difficile*.

## Introduction

Trehalose is a disaccharide sugar consisting of two α-glucose monomers linked via 1,1-glycosidic bond and is present in a wide variety of organisms, such as bacteria, yeast, insects, plants, and animals. The structure of trehalose makes it highly resistant to acid hydrolysis, it is used as a high energy storage molecule for insect flight, and also as a dehydration or cryo-protectant in some microorganisms, plants and animals(1). This sugar is naturally found in foods, such as mushrooms and honey, but following the discovery of a cost-effective method of large-scale production(2) and regulatory approval as a food additive, trehalose is now also added to a range of processed food products (cereals, pasta, sweets and ice cream), cosmetics and some medicines(3,4). In the human digestive tract, trehalose is metabolised by host-produced trehalase enzymes located at the intestinal brush border, as well as microbial-produced trehalases. Many intestinal bacteria and yeasts produce trehalase enzymes, including *Bacillus* spp., *Escherichia coli, Blautia* spp., *Lactobacillaceae* and the nosocomial pathogen *Clostridioides difficile*(5–7).

*C. difficile* is the leading cause of antibiotic-associated diarrhoea(8), where antibiotic-induced microbiota depletion provides favourable conditions to the germination of *C. difficile* spores, which proliferate and produce toxins that cause disease. These toxins (TcdA and TcdB) are responsible for a range of symptoms from mild, self-limiting diarrhoea to pseudomembranous colitis, colonic perforation and death(8). Antibiotic treatment can fail to resolve the infection, with up to 30% of cases recurring after primary treatment(9). Depletion of the intestinal microbiota means *C. difficile* is exposed to several compounds normally metabolised by the healthy microbiota; some of these are used as either molecular signals or as a source of energy, such as the primary bile acids cholate and succinate(10,11). Collins *et al*. showed that ingested trehalose enhanced *C. difficile* virulence(12). Additionally, *C. difficile* ribotypes (RT) typically associated with CDI outbreaks, e.g. RT027 or RT017, have a mutated *treR* repressor gene, which overexpresses the trehalose metabolism gene (*treA*), whereas others, i.e. RT078, show enhanced trehalose uptake, through the presence of a novel phosphotransferase (PTS) system transporter(7). However, it is reported that such variants are common in *C. difficile* isolates and comparison of the clinical outcome of CDI patients to trehalose metabolic genotype of the isolated strain found no correlation between 30-day mortality and the trehalose metabolic genotype (13). Moreover, Saund *et al*. found no statistically significant association between the presence of trehalose utilisation variants in infecting *C. difficile* strains and the development of severe infection outcome(14).

Using a previously successful *in vitro* model developed to simulate human CDI in the large intestine(15), we measured microbial compositional changes following trehalose and glucose supplementation and the effect of sugar supplementation on *C. difficile* growth kinetics after antibiotic disruption of the microbiota. A schematic timeline of our experimental design is shown in **Figure 1**. Enumeration of *C. difficile* from the *in vitro* gut model when predicting treatment and antibiotic induction outcomes have been shown to be clinically reflective, and in some cases, more accurate than results from animal models. Antibiotics with a high propensity to induce CDI in patients also induce simulated CDI within the gut model(15–18). Conversely, antibiotics with a lower *in vitro* propensity to induce simulated CDI are now recognised to have low CDI risk(18,19). Crucially, the gut model has been used for *in vitro* evaluation of drug efficacy against simulated CDI at various stages of pre-clinical and clinical drug development. For fidaxomicin, data from animal and *in vitro* models correlated well with Phase III clinical trials(16,21,22), whereas for the toxin binding agent, tolevamer, results from the triple-stage gut model predicted clinical failure of this agent at Phase III, while animal model data did not(23–25). Following gut model investigations of cadazolid(26), surotomycin(27) and SMT19969(28) for treatment of CDI, these agents have progressed to Phase III clinical trials. A recent publication detailing efficacy of extended duration fidaxomicin therapy(29) led to the current Phase IIIB/IV trial investigating such dosing regimens in the clinic(30). In fact, data from gut model studies(19,31,32) have informed UK national antibiotic prescribing guidelines(33).

**Figure 1.**
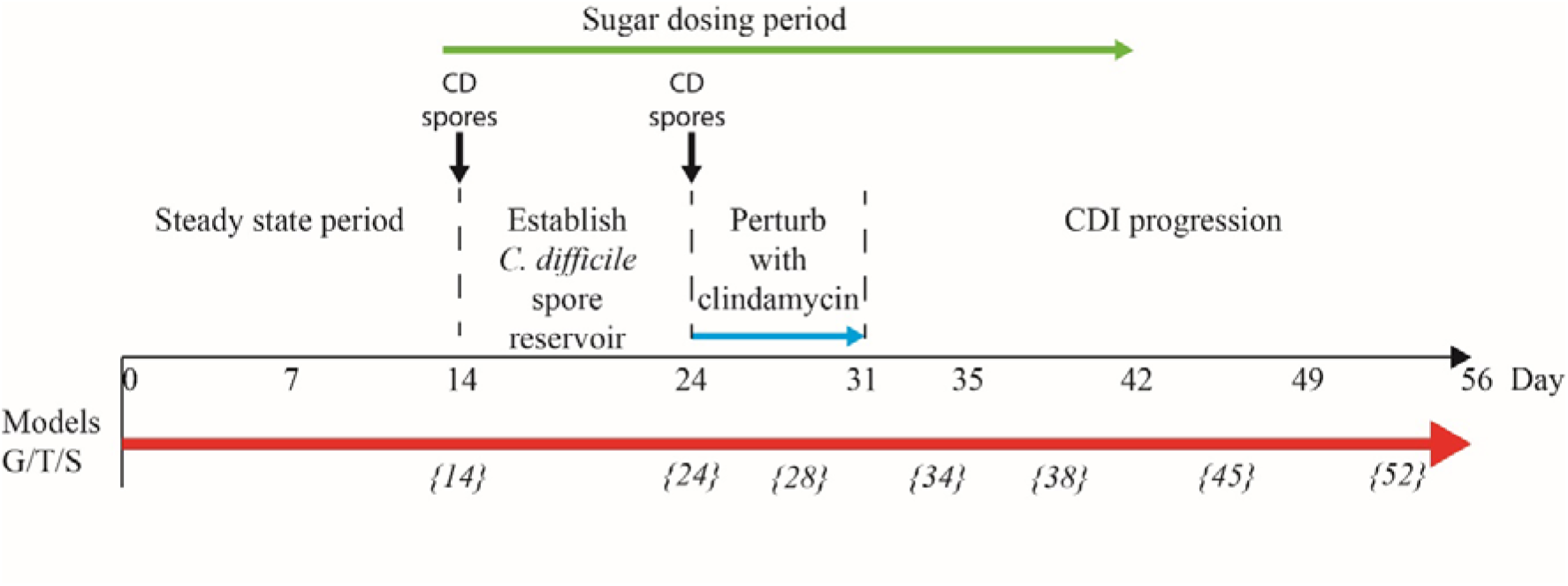
Schematic timeline of the *in vitro* triple vessel chemostat gut model and experimental design for each model. *C. difficile* spores (black lines) were instilled and commencement of sugar regimen [either trehalose (T), glucose (G) or saline (S) (green arrow)] before addition of antibiotics (blue arrow). Microbial populations were monitored post antibiotic for recovery and induction of CDI. Numbers in italicise brackets denotes the day that samples for DNA extraction were taken.

## Results

### Reconstitution of the human microbiota in the gut model

Each triple-staged gut model is seeded with a faecal slurry made using the pooled faecal matter from five healthy donors to capture a diverse representation of the human microbial populations. Firstly, we sought to characterise the microbial diversity captured from this slurry in each gut model. The donors used in this study showed distinct microbial profiles to each other (**Supplementary Table 1**), except for donors E and C, which clustered together, indicating similar microbiota profiles. Similarly, the faecal slurry used clustered distinctly from each donor but nested amongst the donors, suggesting the slurry contains the overall diversity from the donors; indeed, the slurry contained 36 bacterial families, more than each individual donor (**Figure 2A**). As determined by bacterial taxonomic analysis by 16S rRNA sequencing, the slurry was found to contain bacterial family members that were unique to a single donor, for example *Eubacteriaceae* and *Muribaculaceae* were only present in Donor C and the slurry (**Supplementary Table 1**). Once the microbial populations had stabilised in the three models, the predominant bacterial families were represented at similar levels to the slurry (**Figure 2BC**), although some differences were observed. Compared with the faecal slurry, *Bifidobacteriaceae* abundance increased and *Clostridiaceae* abundance decreased in all models at steady state, whereas *Coriobacteriaceae* showed increased abundance in models G and S and *Enterococcaceae* abundance increased in model T (**Figure 2C**). However, *Bifidobacteriaceae* did decrease in all models one week later, which was confirmed by direct enumeration.

**Figure 2.**
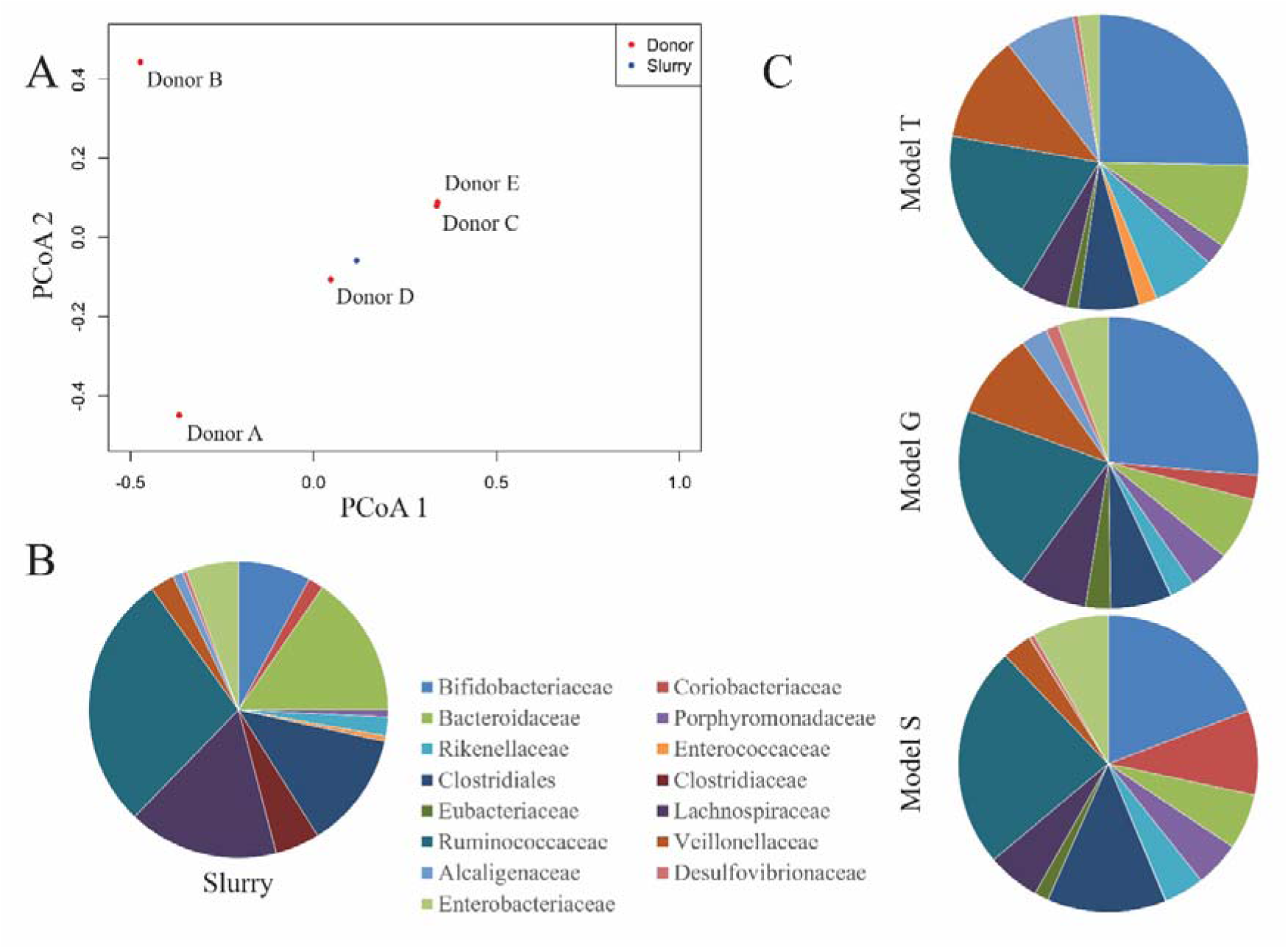
Taxonomic analysis of the faecal donors, pooled slurry and the steady state from the three individual models. Principal coordinate analysis of the five donors and the faecal slurry based on 16S rRNA sequencing (A). Bacterial family abundance (%) found within the pooled faecal slurry used to initiate all models (B) and at end of steady state period (experimental day 14) from the three models (G, T and S) (C).

### Trehalose induced changes to the gut model microbiota

Changes in the bacterial composition of the three models (T, G, and S) were monitored by 16S rRNA sequencing and taxonomic analysis (timepoints shown in **Figure 1**) and direct enumeration of selected bacterial populations (daily monitoring), to determine how trehalose dosing affected the gut microbiota, compared to glucose or saline. The trehalose dosing regimen in model T was designed to mimic the average trehalose content from meals consumed thrice daily by an adult, which can vary amongst adults and different countries(3,34). Trehalose was undetected in the saline (control) model S and was only detected periodically in vessel 1 (proximal colon, pH 5.5±0.2) of the model G dosed with glucose (0.01 mM), whereas trehalose supplementation in model T increased detectable trehalose concentrations, peaking at 8.1 mM in vessel 1 of model T prior to addition of antibiotics (**Figure 3**). Trehalose was largely undetectable in vessels 2 (medial colon, pH 6.2±0.2) and 3 (distal colon, pH 6.8±0.2) of each model, although trehalose was observed in vessel 2 up to three days after commencing dosing, peaking at 4.3 mM (**Supplementary Table 2**). As trehalose can be catabolised into two glucose molecules, model G was dosed with glucose at double the concentration of trehalose to determine if any effects of trehalose on the microbial populations was not due to the increased glucose availability (**Figure 3B**). Prior to antibiotic dosing, commencement of the glucose dosing regimen was associated with a peak glucose concentration of 16.1 mM, compared with 0.1 mM and 0.007 mM in models T and S, respectively. Whilst the microbial populations largely remained stable in the control (saline dosed) model, increased trehalose exposure was specifically associated with an increase in abundance of *Bacteroides uniformis* to almost 3 % of the total bacterial populations, identified by shotgun metagenomic sequencing. This increase was confirmed by selective enumeration of *Bacteroides* spp. and species identification by MALDI-TOF analysis. Similar abundance increases were observed in other microbial populations of model T, *Coprococcus* spp. (*C. catus, C. comes* and *C. eutactus*), *Blautia* spp. (*B. producta* and *B. obeum*), *Veillonella dispar* and *Lactobacillaceae* (*L. fermentum* and *L. johnsonii*) (**Figure 4A**). The enumerated recoveries of *Lactobacillus* spp. in model T showed a 1.2 log_10_ cfu/ml increase in response to trehalose instillation (**Supplementary Figure 1C**). However, the changes in microbiota did not result in germination of *C. difficile* spores. This increase in certain bacterial populations were mirrored by an increase in the number of Pfam domains associated with trehalose metabolism in model T. Trehalase (PF01204/EC:3.2.1.28) and trehalose phosphatase (PF02358/EC:3.1.3.12) Pfam domains increased 1.4- and 3.4-fold, respectively, in abundance upon commencement of trehalose dosing compared with pre-trehalose levels.

**Figure 3.**
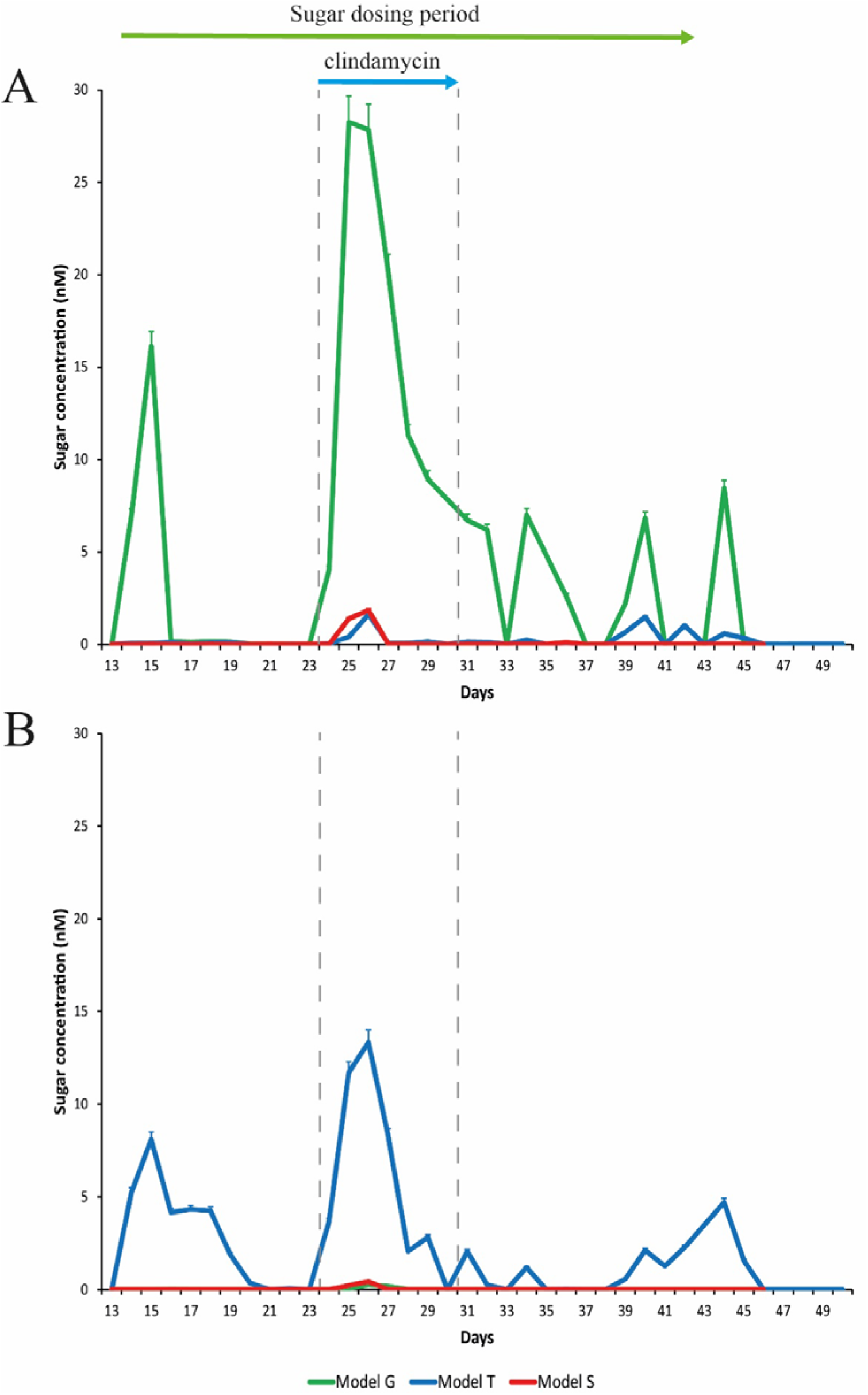
Detection of glucose (A) and trehalose (B) in the gut models. Sugar concentrations from model T (blue lines), G (green lines), and S (red lines) were detected by ion chromatography. Results are shown as mean ± S.D of three technical replicates. Sugar duration and clindamycin dosing are indicated by the top green and blue arrows, respectively.

**Figure 4.**
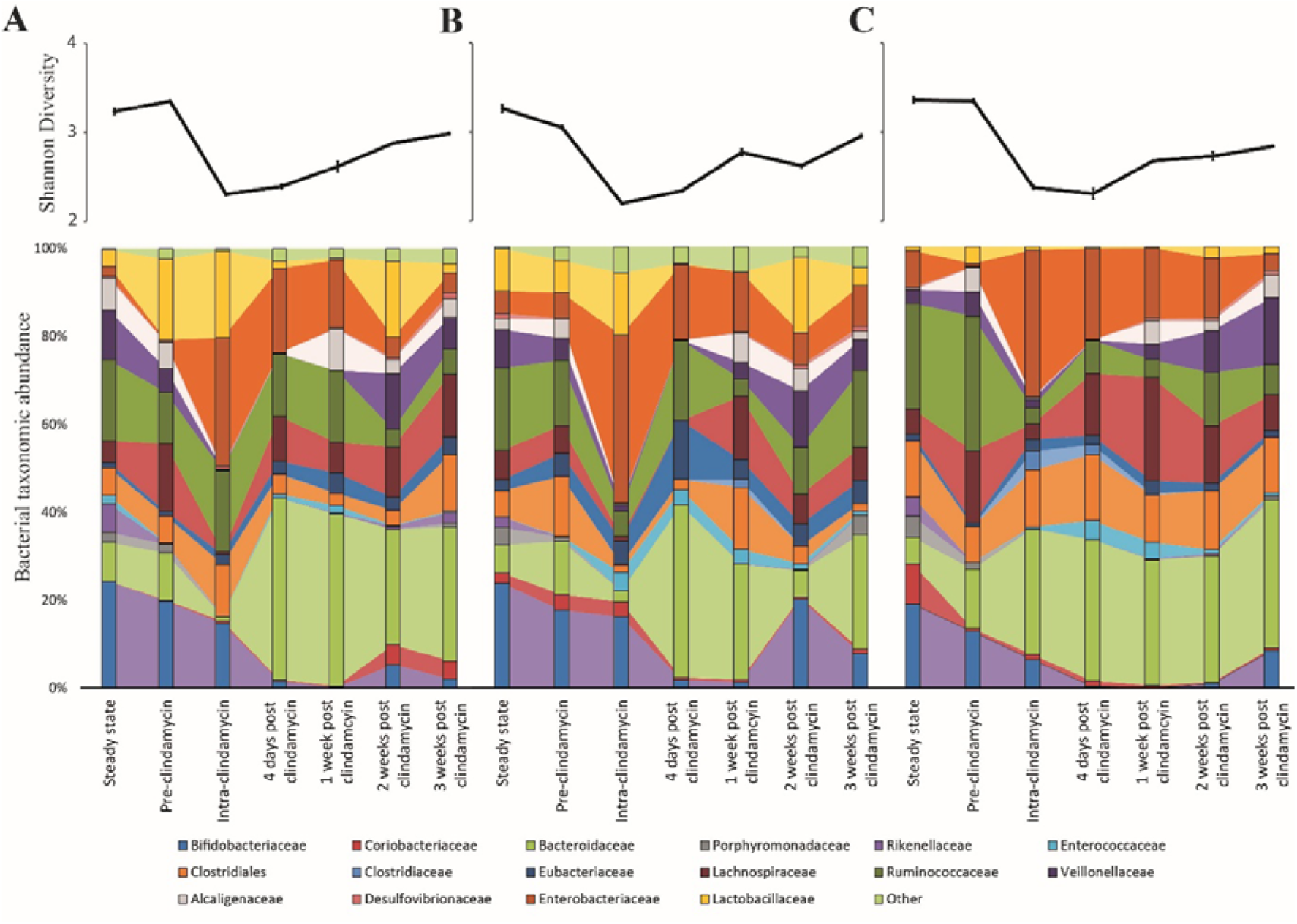
Microbiome changes throughout the model timeline after supplementation with either trehalose (model T; A), glucose (model G; B) or saline (model S; C). Upper line graphs represent diversity changes, as measured by Shannon Diversity Index, throughout the timeline. Results are shown as mean ± S.D of four technical replicates analysed by 16S rRNA sequencing. Lower stacked bar charts represent the bacterial taxonomical abundance changes in each model over time. Results are shown as mean ± S.D of four technical replicates analysed by 16S rRNA sequencing.

### Antibiotic induced dysbiosis increased trehalose bioavailability

CDI induction is reliant on microbial disruption, typically through antibiotic exposure. As we observed sugar-mediated microbiota reconfiguration in our models, we sought to determine how clindamycin induced(16) disruption affected the bioavailability of trehalose that could subsequently be utilised by *C. difficile*. Clindamycin levels in vessel 1 peaked at 93.1, 142.5 and 127.1 mgl, in models G, T and S, respectively. Upon clindamycin instillation, bacterial diversity significantly decreased (*p* < 0.001) from pre-clindamycin levels across all the models to a similar extent (**Figure 4**); however, by three weeks after cessation of clindamycin instillation, bacterial diversity had not recovered to pre-clindamycin levels. This decreased diversity in all models was characterised by significant (*p* ≤ 0.01) reductions to *Bifidobacteriaceae*, *Bacteroidaceae* (*B. gallinarum* and *B. intestinalis*), *Lachnospiraceae, Alcaligenaceae, Porphyromonadaceae* and *Veillonellaceae* populations, either during (intra-) or just after (four days post) clindamycin (**Figure 4**). However, *Enterobacteriaceae* (specifically *E. coli* and *K. pneumoniae*, as determined by MALDI-TOF analysis) and *Enterococcaceae* increased in relative abundance either during or just after antibiotic withdrawal, with an increase in bacterial populations confirmed by direct culture using selective agars (*Enterobacteriaceae* increased 2.4, 2.3, and 2.5 log_10_ cfu/ml; *Enterococcus* increased 1.8, 2.2, and 2.3 log_10_ cfu/ml, in models G, S and T, respectively, during this time) (**Supplementary Figure 1**). This pattern of clindamycin-induced microbial dysbiosis is synonymous with induction of CDI in our *in vitro* gut model, as previously seen (13,16).

Microbial dysbiosis in model T correlated with increased luminal concentrations of trehalose in all vessels (**Supplementary Table 2**); peak trehalose detection in vessels 1, 2 and 3 were 13.3, 6.7 and 2.5 mM, respectively (**Figure 3**). However, luminal concentrations of trehalose reduced to undetectable levels in vessels 2 and 3, and to approximately 2.1 mM in vessel 1, on the last day of clindamycin instillation (day 31). A similar pattern of increased trehalose bioavailability during clindamycin instillation, albeit at much lower levels, was observed in models G (vessels 1-2) and S (vessel 1-3), which, although were not dosed with trehalose throughout the experiment, had trehalose in the growth media. During clindamycin instillation, we detected greater bioavailability of glucose in all models at day 26, particularly in vessel 1 (**Figure 3**). Model G had the highest concentration, peaking at 28.3 mM and concentrations in models T and S peaked at 1.6 and 1.8, respectively. The initial reduction in microbial diversity caused by clindamycin, and the increased trehalose bioavailability, resulted in an increased abundance of microbial genomes harbouring Pfam domains for trehalose metabolism [trehalase (clan 0059) and trehalose phosphatase (clan 0137)]; these domains increased 4.4- and 12.8-fold, respectively, compared with the abundance of these domains pre-trehalose levels. One week after clindamycin, the abundance of these Pfam domains reduced to levels similar to pre-clindamycin.

### Trehalose induced microbiome changes prevents simulated CDI

We hypothesized that the differential pattern of clindamycin-induced microbiota disruption, seen between the three models, and the subsequent microbial recoveries would affect the progression of simulated CDI. Prior to clindamycin instillation, in both the saline supplemented model S (**Figure 5A**), and the glucose supplemented model G (**Figure 5B**), *C. difficile* populations remained in the form of spores from its addition to the models on day 14. However, *C. difficile* germination and outgrowth was detected one-week post clindamycin (day 38) with toxin detected from two days later until the end of the experiment, which is consistent with simulated CDI. Similarly, in the trehalose supplemented model, *C. difficile* spores remained quiescent until day 38, where we detected germination and limited outgrowth but, crucially, no toxin was detected throughout the experiment and the *C. difficile* levels decreased four days later to those consistent with the recovery of *C. difficile* spores (**Figure 5C**). We have previously reported this phenotypic observation in our gut model(13). Thus, we further investigated the comparative microbiota differences and metabolic abundances on this day between models S/G and model T related to *C. difficile* proliferation and onset of simulated CDI.

**Figure 5.**
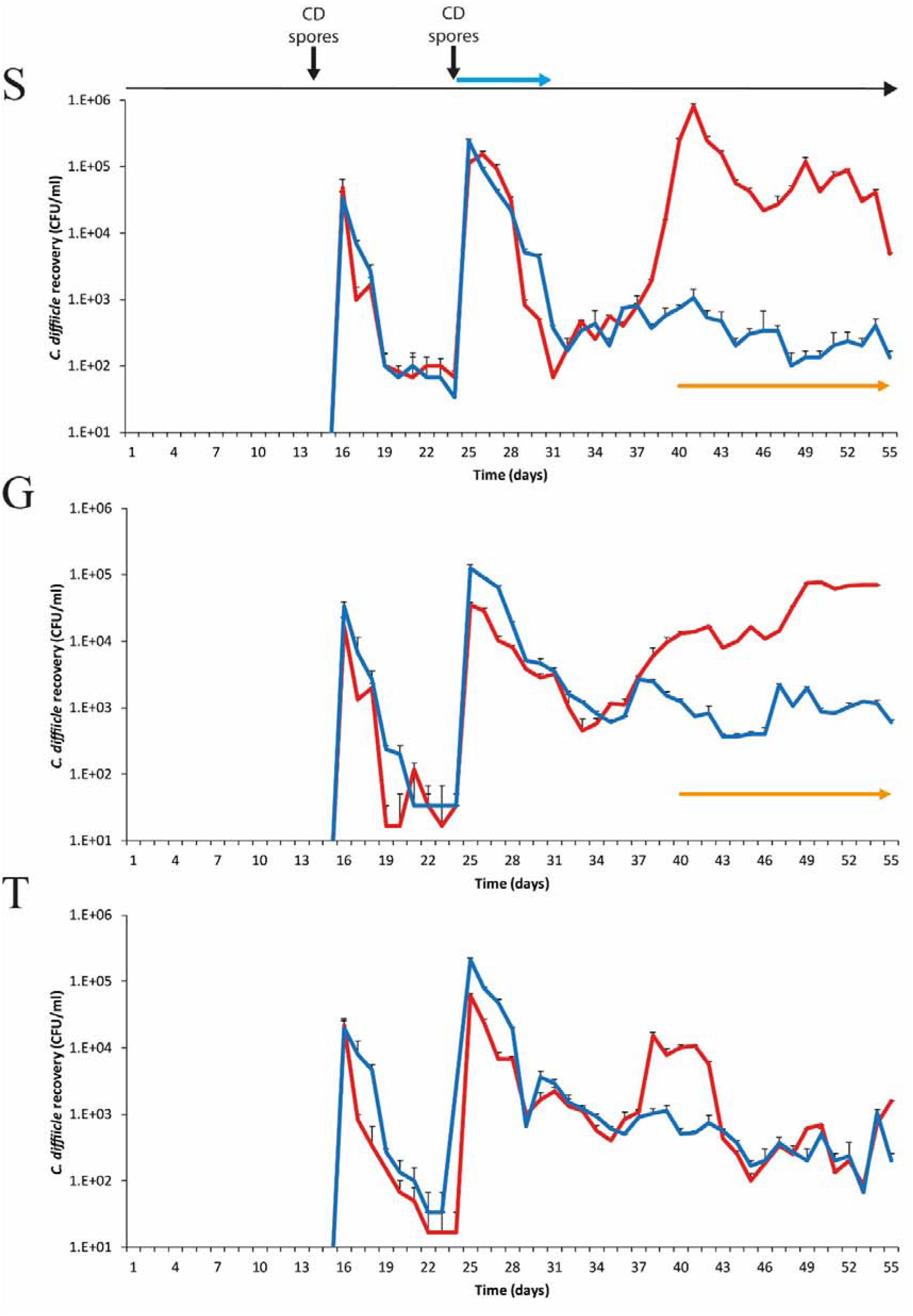
Recovery of *C. difficile* from model S, (A), model G, (B) and model T (C). Recovery of the total *C. difficile* populations (red lines) and the spore populations (blue lines) from vessel 3 are shown for each model. Spore germination and outgrowth was determined by a divergence between the red and blue lines. The detection of toxin is represented by orange arrows; there is no orange arrow for model T as no toxin was detected. Results expressed as mean ± SD from three technical replicates. CD spores denotes when the *C. difficile* spores were inoculated into the model. Blue arrow denotes when clindamycin was instilled into the models.

Metagenomic analysis of the microbiota between models S/G and model T on day 38 highlighted 25 significant differentially abundant bacterial communities between model T (no simulated CDI) and models S and G (induced simulated CDI) (**Table 1**). *Finegoldia magna, Blautia obeum, Faecalibacterium prausnitzii, Dorea formicigenerans, Lactobacillus rhamnosus* and *Oscillospira ruminantium* were significantly (*p*< 0.001) more abundant in model T compared with models G and S, whereas, *Phascolarctobacterium* spp., *Klebsiella* spp. (*K. pneumoniae* and *K. aerogenes*), *E. faecalis* and several *Clostridium* spp. [*C. symbiosum, C. clostridioforme, C. citroniae* and *Hungatella hathewayi* (formally known as *C. hathewayi*)] were significantly less abundant in model T. The increased abundance of *Lactobacillus* spp. in model T at day 38 compared with models S and G, was confirmed by direct enumeration, where *Lactobacillus* spp. levels in model T were 5.9 log_10_ cfu/mL, compared with 3.8 and 3.2 log_10_ cfu/mL for models S and G, respectively (**Supplementary Figure 1**). Similarly, at day 38, the recovery of *Clostridium* spp. and *Enterobacteriaceae* were higher in models S and G, compared with model T (**Supplementary Figure 1**). Analysing the metagenome function showed that components of the V/A-type ATPase were more abundant in models G and S, a conserved ATPase found only in *Enterococcus* spp.(35). Additionally, the high levels of *Clostridium* spp. (including former *Clostridium* species) found in models G and S was reflected in the increased abundance of those genetic elements involved in the sporulation cascade (*spo* genes).

**Table 1.** Differential abundance analysis of model T, compared to models G and S, at day 38.

| Population name<sup>a</sup> | Log<sub>2</sub> fold change<sup>b</sup> |  |
| --- | --- | --- |
|  | vs. Model G | vs. Model S |
| *s_Bacteroides uniformis* | -15.24 | -15.14 |
| *s_Faecalibacterium prausnitzii* | -15.07 | -14.97 |
| *s_Finegoldia magna* | -14.92 | -14.82 |
| *s_Blautia obeum* | -14.81 | -14.71 |
| *s_Dorea formicigenerans* | -14.57 | -14.47 |
| *s_Lactobacillus rhamnosus* | -12.35 | -12.25 |
| *s_Oscillibacter ruminantium* | -2.61 | -19.61 |
| *s_Hungatella hathewayi* | 18.22 | 16.8 |
| *s_Clostridium symbiosum* | 17.12 | 18.08 |
| *s_Klebsiella aerogenes* | 17.05 | 16.72 |
| *s_Blautia producta* | 15.51 | 16.06 |
| *s_Clostridium clostridioforme* | 15.30 | 16.43 |
| *s_Enterococcus faecalis* | 3.4 | 3.59 |
| *s_Klebsiella pneumoniae* | 3.18 | 2.27 |
| *s_Clostridium citroniae* | 3.05 | 2.57 |
| *g_Pantoea* | -14.91 | -14.81 |
| *g_Erysipelatoclostridium* | -14.83 | -14.73 |
| *g_Cronobacter* | -14.55 | -14.45 |
| *g_Eubacterium* | -14.45 | -14.36 |
| *g_Xanthomonas* | -14.38 | -14.28 |
| *g_Paraprevotella* | -14.34 | -14.24 |
| *g_Ochrobactrum* | -12.52 | -12.42 |
| *g_Phascolarctobacterium* | 20.06 | 19.12 |
| *g_Clostridium* | 15.98 | 16.47 |
| *f_Peptoniphilaceae* | -12.44 | -12.34 |

<sup>a</sup> Taxonomic level of microbial populations from whole genome sequencing data. s\_ species, g\_ genus or f\_ family.
<sup>b</sup> Log<sub>2</sub> fold change compared with model T, i.e. a positive number (red) indicates microbial population was significantly (FDR $p \leq 0.01$ ) less abundant in model T compared with models G and S, whilst a negative number (green) indicates microbial population was significantly (FDR $p \leq 0.01$ ) more abundant in model T.

Long-term supplementation of trehalose was associated with enhanced recovery of the Ruminococcaceae family, particularly the *Oscillospira*, *Erysipelotrichaceae* and *Anaeroplasmataceae* genera, 3-weeks post-clindamycin (**Figure 4A**). Inversely, *Bifidobacteriaceae* abundance in model T took longer to recover than the other models, where abundance remained at approximately 2 % compared with 7.2-7.6 % abundance observed in the other models. This was reflected in the bacterial recoveries where *Bifidobacterium* spp. levels recovered by days 13 and 9 (after clindamycin instillation) in models S and G, respectively, and day 19 in model T (**Figure 4**).

### Functional microbiome changes associated with CDI

In order to identify the microbial pathways active during CDI induction, a functional analysis of KEGG metabolic pathways was performed between model T, supplemented with trehalose, and models G and S, supplemented with glucose and saline (**Supplementary Table 3**). Comparison of the functional metabolic pathways between models was performed on day 38 (one-week post clindamycin), when CDI was detected in models G and S but not in model T. Model T showed differences to the metabolic landscape and cell surface components of the microbial species present, in comparison to models G and S. Whilst the abundance of genes involved in glucose/gluconeogenesis and the pentose phosphate pathway were similar across the models, other metabolic pathways that produced the central metabolites glyceraldehyde-3 phosphate (G3P) and D-glucose from different sugar sources, such lactose, galactose and *myo*-inositol, were more abundant in models G and S, whereas genes involved in trehalose metabolism were more abundant in model T (**Figure 6A**). Interestingly, several pathways to produce amino acids (glutamine and glutamate) or amino acid precursors (chorismate and D-ribose-1,5P) were abundant in models G and S. Alongside the increased abundance of the intracellular stress molecule guanosine pentaphosphate (ppGpp), this could indicate a depleted pool of free amino acids (**Supplementary Figure 2**). In contrast, we detected an increased abundance of genetic components involved in the conversion of L-glutamine to the short chain fatty acid, butyrate (**Figure 6A**).

**Figure 6.**
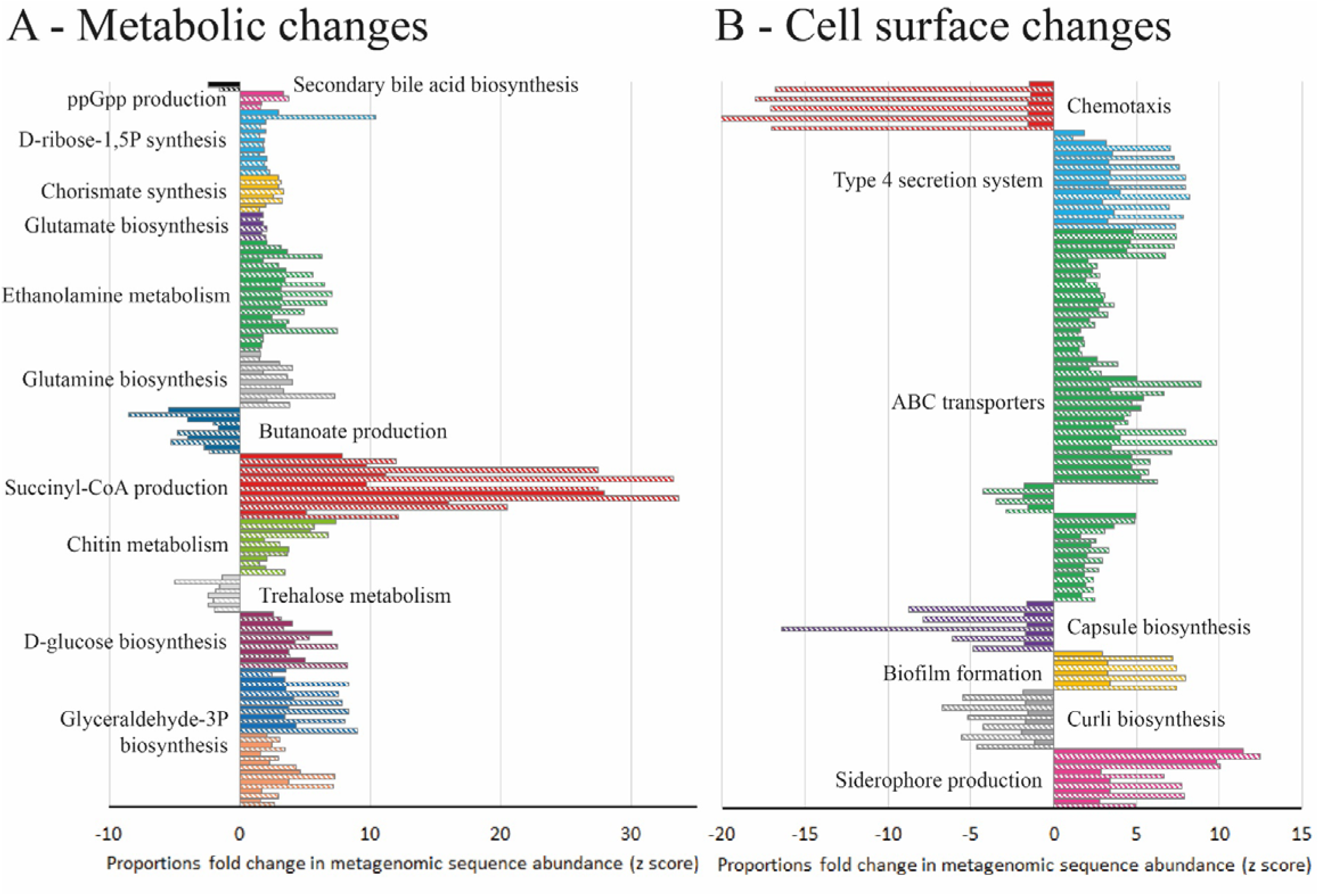
Differential abundance analysis (fold-change) of metabolic pathways in model T compared to models G (solid bars) and S (hatched bars) at day 38, based on metagenomics analysis. Abundance bars highlight the KEGG metabolic pathways (A) or KEGG cell surface components (B) that are significantly (*p*=<0.05) more abundant (>-1.5 fold-change) or less abundant (>1.5 fold-change) in model T. Pathways or components were only included where 80% of the genetic elements in that pathway/component showed an increased or decreased abundance from technical replicates of model T compared to both models G and S.

Differential abundance analysis of the shotgun metagenome sequences from these models suggests that the microbial species could have distinctive cell surface arrangements. In all three models, genetic elements associated with biofilm formation were present; however, these genetic elements were different between the models. Model T had increased abundance of genes involved with curli biosynthesis, important in Enterobacteriaceae biofilm formation, whereas, the genes involved in production of extracellular matrix (PGA) were more abundant in models G and S (**Figure 6B**). The presence of classical virulence factors were also different; genetic elements associated with siderophore production (yersiniabactin and bacitracin) and type 4 secretion systems (T4SS) were more abundant in models G and S, whilst genes involved in the production of polysaccharide capsule were less abundant. Additionally, ABC transporter systems were different between the models; sugar transport systems, for ribose, lactose, arabinogalactan and cellobiose, were more abundant in models G and S, whilst the general L-amino acid transporter was more abundant in model T (**Figure 6B**).

## Discussion

The consumption of trehalose has been proposed to contribute to the emergence and virulence of CDI outbreaks, specifically in those RTs harbouring genetic trehalose metabolism variants(7,12). Some RT lineages, such as RT027, have a mutated repressor, *treR*, gene, whilst others, such as RT078, have acquired a putative *ptsT* transport system, which gives them a competitive advantage during pathogenesis(12). However, we and others have reported that these genetic variants are common in multiple *C. difficile* lineages not associated with clinical outbreaks and there was no association between trehalose metabolism variant and clinical outcome(13,14). Here we describe the effects of trehalose supplementation on the human microbiota and how the recovery of the microbiota can affect simulated CDI, using a clinically reflective gut model. This model has been used to successfully predict trial outcome at various stages of pre-clinical and clinical therapeutic development. However, due to the limited number of biological replicates (n=2), our conclusions are presented as preliminary findings. Numerous reports have highlighted key microbiota differences between humans and mice(36,37), which is potentially important given the contribution of the microbiota towards colonisation resistance against CDI. Indeed, mice bred in captivity, on the same diet, have a similar microbiota(38), which is far removed from the heterogeneity displayed within in a human population, and could result in different clinical outcomes(39). The major human microbial communities present in the donor faeces were represented in the faecal slurry used to seed each model, and we subsequently recaptured these communities in each of our human ‘gut models’. Three independent models were run in parallel, supplemented with either trehalose, glucose or saline. As trehalose can be metabolised into two glucose molecules, this experimental approach was used to delineate the effects of trehalose rather than its breakdown product.

Trehalose supplementation immediately increased the trehalose concentration in vessels 1 and 2; however, despite continued trehalose supplementation, the levels decreased to undetectable levels, suggesting microbial trehalose metabolism. This drop in trehalose concentration was specifically associated with an increased abundance of trehalose metabolising Pfam domains and a corresponding remodelling of the microbiota to include higher abundancies of *B. uniformis, B. producta* and *V. dispar*. Many strains of these bacterial species are known to harbour *treA*-like genes, a trehalose-6-phosphatase that metabolises trehalose into glucose and glucose-6-phosphate monomers(40–42). Trehalose metabolism appears not to be restricted to these bacterial species, indeed, the UniProt protein sequence repository suggests trehalose metabolising genes are ubiquitous amongst bacterial species(6). Further evidence of the role of the intestinal microbiota to metabolise trehalose is shown during clindamycin exposure. Clindamycin is a broad-spectrum antibiotic that has previously been shown to induce CDI in our gut model(16). Clindamycin instillation decreased the microbial diversity in all models, causing simulated intestinal dysbiosis, which correlated with increased bioavailability of trehalose. The levels of trehalose returned to undetectable levels in vessels 2 and 3 prior to completion of the clindamycin regimen; these vessels simulate the medial and distal colon, which is most conducive for CDI(15). We used a highly sensitive method to quantify trehalose/glucose concentrations, with a limit of detection at 0.5μM. Collins *et al*. reported *C. difficile* trehalose metabolism variants that could utilise 50μM-25mM concentrations of trehalose more efficiently, compared with wildtype trehalose metabolism variants(12). We hypothesised that, although *C. difficile* trehalose metabolism variants could have a competitive advantage over wildtype isolates, this competitive advantage is diminished in the presence of the human microbiota, as the intestinal microbiota is an important source of trehalose metabolism, potentially reducing the bioavailability of trehalose during CDI. Further evidence for this is shown using degradation-resistant trehalose analogues, such as lactotrehalose, which show enhanced faecal bioavailability and, additionally, does not induce CDI in a mouse model(43). The Ribotype 027 strain used in our studies possesses a *treR* mutation; it would be interesting to determine the competitiveness of a strain harbouring a putative *ptsT* transport system, which gives them a competitive advantage during pathogenesis(12).

Instillation of clindamycin did result in induction of simulated CDI in models supplemented with either glucose or saline; however, trehalose supplementation did not result in simulated CDI ((13) and **Figure 5**). Differential taxonomic abundance between model T and models G/S identified bacterial species that were either more (n=7) or less (n=8) abundant in model T; although these data are from one independent biological repeat of each condition. Of the bacterial species associated with protection against CDI, *B. uniformis* was the dominant *Bacteroides* species identified in model T (4% of total reads) and is capable of metabolising trehalose. Through metabolite cross-feeding, *B. uniformis* can support the growth of *F. prausnitzii*(44), and vice versa, which enhances the growth of both species. *F. prausnitzii* is known to produce butyrate and formate, which have been shown to reduce inflammation during infection(45), but increased levels of *F. prausnitzii* were associated with recovery from CDI in patients and *in vivo* models(46,47). Strains of *B. obeum* can also utilise trehalose, and crucially, have been shown to reduce virulence gene expression, through quorum sensing, during *Vibrio cholerae* infection(48); toxin synthesis by *C. difficile* can be regulated by different quorum sensing(49,50). The metabolic interconnectivity between these microbial species could represent a consortium that competes for nutrients utilised by *C. difficile* after spore germination (as hypothesised in **Supplementary Figure 3**). Furthermore, the increased abundance of these bacterial species associated with CDI prevention in model T are less abundant in the faeces from CDI patients and have been attributed with protective effects in putative microbiome restoratives(51–53). Moreover, carbohydrate-induced microbiota outgrowth, and subsequent metabolic products, were associated with a decrease in *C. difficile* fitness(54), similar to our *in vitro* findings. Aside from acting as a potential carbon source, intracellular trehalose has unique properties that could protect cells from the effects of antibiotics, such as a membrane osmoprotectant and a protein stabilising molecule(1,55).

Interestingly, 4 different *Clostridium* spp. were associated with CDI induction in model’s G/S. A similar observation was noted by Khanna *et al*.(46) and Daquigan *et al*.(56), where several *Clostridium* spp. were associated with increased *C. difficile* abundance in CDI and recurrent CDI patients. Additionally, Girinathan *et al*., showed the presence of *Clostridium sardiniense* enhanced CDI in an *in vivo* model of disease(57). Conversely, the authors observed that *C. bifermentans*, an amino acid fermenter, prevented CDI-induced death in an animal model of infection. As *C. difficile* utilises amino acids during growth expansion as both a carbon source and an energy source, via Stickland reactions(58,59), microorganisms that compete for bioavailable amino acids, such as *C. bifermentans*, could prevent the rapid proliferation of *C. difficile* during CDI(57). Indeed, the CDI-linked functional profile of the microbiota favoured the production of amino acids (or amino acid precursors) from sugars, potentially due to a reduced pool of bioavailable amino acids after being used by *C. difficile* during pathogen expansion. This was mirrored in the functions of the cell surface ABC transporters where there was an increased abundance of sugar transporters in models G and S, potentially associated with *C. difficile* proliferation niche. The identified metabolic pathways and putative metabolic products associated with CDI induction in our gut models, namely G3P/D-ribose-5P/succinyl-CoA/inosine, have been shown to be utilised by *C. difficile* during *in vivo* growth in a murine model of infection(60).

By studying the effect of trehalose at a systems level, we outline plasticity of the human microbiome to adapt to utilise this carbon source, providing competition with *C. difficile*. The absence of enhanced CDI associated with trehalose in this model system, and the mechanisms explained, could help explain why clinical outcome was not associated with genetic markers for enhanced trehalose metabolism. Contrary to enhancing CDI, trehalose utilisation was preliminary linked with a consortium of cross-feeding microbial species that may be antagonistic to the growth of *C. difficile*, which can form the rationale for the basis of microbial therapies used for treating CDI infections (**Supplementary Figure 3**). Further *in vitro* and *in vivo* studies are required to confirm these preliminary data.

## Materials and methods

### Gut model set up

Three triple-stage *in vitro* gut models were run in parallel as previously described(61). Briefly, each model consists of three chemostat vessels arranged in a weir cascade system, top-fed with a complex growth medium at a controlled rate (D=0.015 h^−1^). All three vessels were continuously stirred, anaerobically maintained at 37 °C and regulated to reflect *in vivo* intestinal conditions. The three vessels are representative of the colon, reflecting increased pH and decreased nutrient availability from the proximal colon (vessel 1, pH 5.5±0.2), through the medial colon (vessel 2, pH 6.2±0.2) to the distal colon (vessel 3, pH 6.8±0.2). A faecal slurry (10% w/v) was made by diluting pooled human faeces from healthy elderly volunteers (n = five; >59 years of age; CDI negative; no prior three-month history of antibiotic exposure) with pre-reduced PBS. Donors over the age of 59 were used to represent the at-risk group for developing CDI. This slurry, approximately 500 ml per model, was used to seed the vessels for each gut model at the start of the experiment.

### Experimental timeline

For each model, the microbial populations were allowed to equilibrate for two weeks to reach ‘steady state’ without intervention; consistent stable microbial recoveries over 5 days from all microbial populations tested was used to determine ‘steady state’. A single one mL aliquot of *C. difficile* RT027 strain 210 spores (approximately 10^7^ cfu/ml) were inoculated into vessel 1 of each model and sugar supplementation commenced (**Figure 1**). Models were dosed with either glucose (n=1 model; model G; 1120 mM dosed thrice daily for 35 days), trehalose (n=1 model; model T; 560 mM thrice daily for 35 days), or phosphate buffered saline (n=1 model; model S; thrice daily for 35 days) (**Figure 1 – green arrow**). The dosing regimens were inoculated in vessel 1, and sufficient to achieve final trehalose and glucose concentrations in vessel 1 of each model of 10 mM and 20 mM, respectively, i.e. consistent with levels observed in humans consuming trehalose(3,34). A second dose of *C. difficile* strain 210 spores were inoculated as previously described, followed by a clindamycin regimen (33.9 mg/l four times daily for seven days) to induce simulated CDI(16). The first *C. difficile* spore dose was added to ensure that the microbial populations within the model had equilibrated and were able to prevent *C. difficile* spore germination; an effect called colonisation resistance. This ensured that the effects of any downstream manipulations of the microbiota were due to those manipulations rather than incomplete colonisation resistance.

### *Preparation of Ribotype-027* C. difficile *strain 210 spores*

Similar to other strains of ribotype-027, *C. difficile* 210 strain has a mutated *treR* repressor gene, as reported in(12). *C. difficile* spores for gut model inoculation were prepared as previously described(62). Briefly, *C. difficile* 210 was grown in BHI broth anaerobically at 37 °C for six days and removed from the incubator and incubated aerobically at room temperature overnight to further induce sporulation. Growth was harvested by centrifugation and incubated with PBS supplemented with 10 mg/ml lysozyme at 37 °C overnight. Samples were separated using a sucrose gradient and spores were treated with PBS supplemented with 20 ng/mL protease K and 200 nM EDTA. Spores were separated using a sucrose gradient and washed with PBS twice before a final resuspension in 30 ml. These were enumerated and diluted to approximately 1×10^7^ spores/ml for use in the models.

### Bacterial enumeration using selective agars

Vessel 3 of models T, S and G was sampled daily for culture profiling of total bacteria, lactose-fermenting Enterobacteriaceae, *Clostridium* spp., *Lactobacillus* spp., *Bifidobacterium* spp., *Enterococcus* spp., and *Bacteroides* spp., and *C. difficile* populations as described in **Supplementary Methods** and **Supplementary Table 4**. Colonies from these plates were identified to a species level by MALDI-TOF analysis.

### Cytotoxin assay

A VERO cell-based assay was used to approximate the level of toxin present in the gut model. This assay is a semi-quantitative measure of toxin activity within the gut models as many factors can affect the action of the toxin, e.g. protein levels, which were not normalised across each time point & sample. Samples from vessels 1, 2 & 3 were centrifuged and stored at 4 °C before testing. Samples were serially diluted 1:10 and co-incubated with a confluent monolayer of VERO cells and incubated for 48 hr at 37 °C, 5 % CO_2_. Toxin positivity was indicated by >50 % cell rounding, while the confluent cell monolayer was unaffected in toxin negative samples. Presence of *C. difficile* toxin was confirmed by neutralisation of neat sample with *C. sordellii* antitoxin.

### Antibiotic bioassay

The concentration of clindamycin in each vessel was determined by bioassays as previously described(16). Briefly, indicator organism Kocuria rhizophila (ATCC 9341) was inoculated into Wilkins-Chalgren agar and aseptically transferred into 245 x 245 mm agar plates. These plates were allowed to set, and nine mm wells were made using a cork borer. A calibration series of the antibiotic was added to each plate and samples loaded into the wells. Plates were incubated overnight, aerobically at 37 °C. Zone diameters were measured using callipers and concentration curves plotted from squared zone diameters and unknown concentrations from vessel supernatants determined. Assays were performed in triplicate.

### Trehalose and glucose concentration

Daily gut model samples from all vessels of each model were assessed by ion chromatography for trehalose and glucose concentrations. Samples were tested in a blind fashion. Gut model samples were centrifuged at 15,000 rpm for 10 mins, and the supernatant was further centrifuged using millipore centrifugal filter units with a 50 kDa and 3kDa cut off to remove proteins and bigger macromolecules. Samples were diluted 1:10 in ultrapure water and analysed by ion chromatography. 25 μl of sample was injected on a Dionex ICS-5000 plus ion chromatography system (Thermo fisher Scientific, USA). The separation was carried out on a Dionex CarboPac PA1 Analytical Column (4 x 250 mm) with a Dionex CarboPac PA1 Guard Column (4 x 50 mm). The column temperature was set at 30°C. Elution was performed using three elutants; A: ultrapure water, B: 300 mM sodium hydroxide and C:300 mM sodium hydroxide with 1500 mM sodium acetate, with a flow rate of one ml/min. The linear gradient condition is shown in **Supplementary Table 5**. The analytes were detected by using an electrochemical detector.

### DNA extraction, bacterial 16S rRNA library preparation and sequencing

The bacterial 16S rRNA sampling regimen from vessel 3 of all three models are shown in **Figure 1**. Samples were taken on the day but prior to the first sugar dose, first clindamycin dose, in the middle of clindamycin dosing, 2 days after last clindamycin dose, i.e. after washout of clindamycin from the system, and weekly thereafter. Four one ml aliquots, representing four technical replicates, from vessel 3 of each model were pelleted, the supernatant discarded, and the DNA extracted from the pellet using FastDNA™ SPIN kit for soil (MP Biomedicals™) following manufacturers’ instructions. DNA was stored at −80 °C until used for downstream analysis. DNA quality and double-stranded quantities were determined using the picogreeen absorption method. Bacterial 16S rRNA V4 fragments were PCR amplified using NEBNext Q5 Hot Start HiFi PCR master mix (NEB, U.K.) with universal 16S rRNA V4 primers [564F (TCGTCGGCAGCGTCAGATGTGTATAAGAGACAG-AYTGGGYDTAAAGNG) and 806R (GTCTCGTGGGCTCGGAGATGTGTATAAGAGACAG-TACNVGGGTATCTAATCC)] with Illumina adaptor sequence overhangs included using the PCR cycle [denaturation (95 °C x3 min for 1 cycle), amplification (95 °C x30 sec, 50 °C x30 sec, 72 °C x30 sec, for 28 cycles) and final elongation (72 °C x5 min for 1 cycle)]. PCR products were cleaned using AxyPrep Magnetic beads (Axygen, U.K.) and the 16S rRNA fragments checked using gel electrophoresis on an Agilent 2200 TapeStation system (Agilent Genomics, U.K.) before running a PCR for addition of the index sequences [denaturation (95 °C x3 min for 1 cycle), amplification (95 °C x30 sec, 55 °C x30 sec, 72 °C x30 sec, for 8 cycles) and final elongation (72 °C x5 min for 1 cycle)]. The fragments were cleaned, quantified (as before), normalised and samples sequenced using MiSeq sequencer (Illumina) with 250 bp paired-end reads. Library preparation and sequencing was done at the University of Leeds sequencing facility.

### Taxonomic analysis on 16S rRNA gene sequences

Demultiplexed FASTQ files of 16S rRNA sequences were trimmed of adapter sequences using cutadapt(63), and samples were filtered based on the number of reads across all samples to provide similar coverage. We followed the standard operating procedure from the MOTHUR package (v.1.41.3)(64). The paired reads were joined together and assembled into the contigs, and quality controlled based on the parameters such as maxambig=0, minlength=177 and maxlength=237. Unique sequences were aligned against a tailor-made reference generated from SILVA SEED database (version 132) and further filtered according to their start and end positions in the alignments. In order to reduce the possible redundancy to a minimum, the identical and very similar sequences (within 2bp mismatch) were merged. The chimeric sequences were discarded based on the built-in VSEARCH method(65). OTUs (operational taxonomic units) were identified by clustering (0.5 UniFrac distance) the sequences and were assigned the consensus taxonomy information with label=0.03. Low abundant reads (<50 reads per sample, which accounted for <0.04% of the total reads) were removed before further analysis; however, this precludes the contribution of low abundance taxonomic families in our analysis whilst ensuring accuracy from sequencing artifact. Taxonomic analysis is represented as mean percent abundance from four technical replicates. To determine if a microbial community was differentially abundant between the technical replicates of the three models on a given day a differential abundance analysis was performed and a microbial community was considered significantly different when the false discovery rate (FDR) *p* value was ≤ 0.01 and a −1.5 ≤ Log_2_ fold change ≥ 1.5 of the populations in model T in comparison with models G and S was observed. The same cut off values were used to determine the difference in abundance of the microbial populations of the slurry compared to the models T, G, and S at day 14. For the calculations of bacterial diversity, Shannon diversity index was computed for all samples and the distance matrix based on thetayc approach was used for Principal Coordinates (PCoA) analysis and visualization for each group of samples, based on four technical replicates. A Mann-Whitney U test was performed to determine statistical significance between the pre-clindamycin Shannon diversity index reference and the other timepoints for each model.

### Functional analysis of the microbiota

All samples collected from model T throughout the experiment (time points shown in Figure 1) and samples from model S and G on day 38 (1-week post clindamycin), were selected for shotgun metagenome sequencing. Four technical replicates were investigated for each time point. Extracted DNA was diluted to 500 ng and sheared to 200-300 bp using an E220 focused ultrasonicator (Covaris, U.K.). NEBNext Ultra DNA Library prep kit for Illumina was used for adaptor ligation and to PCR enrichment following manufacturer’s instructions. Libraries were sequenced using Illumina HiSeq 3000 sequencer (University of Leeds). FASTX-Toolkit (version 0.0.13) was used to trim the first 10 bp from the sequence reads. MEGAN UE (ultimate edition v6.18.0) was used to functionally annotate and compare the sequencing reads using inbuilt programme tools(66). Briefly, paired-end reads were aligned on to NCBI-nr database (version 14.12.19; nr.gr size: 52.5 Gb) using DIAMOND(67) and Meganizer was used to perform functional analysis of the DIAMOND input files using either an up-to-date representation of KEGG or Pfam-A database (version 32). KEGG orthologous groups (Ko groups) were mapped to enzymes that appear in metabolic pathways. Functional analysis using KEGG assigned 39-43% of the total reads to a Ko group. An abundance table of functional categories for each sample was generated and a differential proportion-based abundance analysis, based on z test(68), was performed. A functional annotation was considered significantly different when the false discovery rate (FDR) *p* value was ≤ 0.05 and a −1.5 ≤ Log2 fold change ≥ 1.5 of the populations, based on z-score, in model T in comparison with models G and S. Variations in Ko groups (metabolic pathways and surface components) were considered significant when the fold-change of at least 80 % of the genetic components in that pathway were either more or less abundant (as described above). The abundance of trehalase-specific Pfam terms, trehalase (clan 0059) and trehalose phosphatase (clan 0137), throughout all timepoints of model T were compared, and fold change abundance calculated.

### Taxonomic profiling of whole genome sequences

Taxonomic profiling from the whole genome paired sequences generated above was performed using CLC Genomics Workbench (version 12.0.3), with the CLC Microbial Genomics Module (version 4.8). Trimmed sequences were imported into the CLC software and each sequence read individually mapped to the fully curated (as of June 2019) microbial reference database, based on selected references from GenBank and RefSeq, using the default settings in the *Taxonomic Profiling* tool. Abundance tables were merged, and the *Differential Abundance Analysis* tool used to determine the abundance differential in taxa, compared to model T, based on a Log2 fold-change <-2 or >2 and a Wald test false discovery rate (FDR) *p* value <0.001.

## Acknowledgements

We thank Sharie Shearman for technical assistance with the gut model, and Dr Carr and Dr Raynor for preparing the sequencing libraries and performing the sequencing.

## Author Contributions

A.M.B., T.H. and M.H.W. conceived and designed the experimental studies. A.M.B., I.B.M., N.A., W.S., E.C., Y.N., H.C.H. and K.B. conducted the experiments. I.B.M., S.M., D.W. and A.M.B., analysed the experimental data. A.M.B., I.B.M., N.A. and M.H.W. wrote the manuscript with additional input from T.H., G.D., D.W., S.M. and I.B.M. All authors approved the manuscript.

## Ethics statement

The collection and use of human faeces in our gut model has been approved by the School of Medicine Research Ethics Committee, University of Leeds (MREC 15-070 – Investigation of the Interplay between Commensal Intestinal Organisms and Pathogenic Bacteria). Participants were provided with a ‘Participant Information Sheet’ (PIS) detailing a lay summary of the *in vitro* gut model and the scientific work they were contributing to by providing a faecal donation. Within this PIS, it is explained that by providing the sample, the participant is giving informed consent for that sample to be used in the gut model.

## References

1. Elbein AD, Pan YT, Pastuszak I, Carroll D. New insights on trehalose: a multifunctional molecule. Glycobiology. 2003;13(4):17–27.

2. Maruta K, Nakada T, Kubota M, Chaen H, Kurimoto M, Tsujisaka Y, et al. Formation of Trehalose from Maltooligosaccharides by a Novel Enzymatic System. Biosci Biotechnol Biochem. 1995;8451(10):1829–34.

3. Food Standards Australia New Zealand (FSANZ). Final assessment report application A453: trehalose as a novel food. FOOD Stand Aust NEW Zeal. 2003;(May).

4. Pinto-bonilla JC, Olmo-jimeno A, Llovet-osuna F, Hernández -E. A randomized crossover study comparing trehalose/hyaluronate eyedrops and standard treatment: patient satisfaction in the treatment of dry eye syndrome. Ther Clin Risk Manag. 2015;11:595–603.

5. Arguelles JC. Physiological roles of trehalose in bacteria and yeasts: a comparative analysis. Arch Microbiol. 2000;174(4):217–24.

6. Bateman A, Martin MJ, O’Donovan C, Magrane M, Alpi E, Antunes R, et al. UniProt: The universal protein knowledgebase. Nucleic Acids Res. 2017;45(D1):D158–69.

7. Collins J, Danhof H, Britton RA. The role of trehalose in the global spread of epidemic *Clostridium difficile*. Gut Microbes. 2018;00(00):1–6.

8. Rupnik M, Wilcox MH, Gerding DN. *Clostridium difficile* infection: new developments in epidemiology and pathogenesis. Nat Rev Microbiol. 2009;7(7):526–36.

9. Marsh, J.W., Arora, R., Schlackman, J.L., Shutt, K.A., Curry, S.R., Harrison LH. Association of Relapse of *Clostridium difficile* Disease with BI/NAP1/027. J Clin Microbiol. 2012;50(12):4078–82.

10. Sorg J a., Sonenshein AL. Bile salts and glycine as cogerminants for *Clostridium difficile* spores. J Bacteriol. 2008;190(7):2505–12.

11. Ferreyra JA, Wu KJ, Hryckowian AJ, Bouley DM, Weimer BC, Sonnenburg JL. Gut microbiota-produced succinate promotes *C. difficile* infection after antibiotic treatment or motility disturbance. Cell Host Microbe. 2014;16(6):770–7.

12. Collins J, Robinson C, Danhof H, Knetsch CW, Van Leeuwen HC, Lawley TD, et al. Dietary trehalose enhances virulence of epidemic *Clostridium difficile*. Nature. 2018;553(7688):291–4.

13. Eyre DW, Didelot X, Buckley AM, Freeman J, Moura IB, Crook DW, et al. *Clostridium difficile* trehalose metabolism variants are common and not associated with adverse patient outcomes when variably present in the same lineage. EBioMedicine. 2019;

14. Saund K, Rao K, Young VB, Snitkin ES. Genetic Determinants of Trehalose Utilization Are Not Associated With Severe *Clostridium difficile* Infection Outcome. Open Forum Infect Dis. 2020;1–4.

15. Freeman J, Neill FJO, Wilcox MH. Effects of cefotaxime and desacetylcefotaxime upon *Clostridium difficile* proliferation and toxin production in a triple-stage chemostat model of the human gut. J Antimicrob Chemother. 2003;52:96–102.

16. Chilton CH, Crowther GS, Freeman J, Todhunter SL, Nicholson S, Longshaw CM, et al. Successful treatment of simulated *Clostridium difficile* infection in a human gut model by fidaxomicin first line and after vancomycin or metronidazole failure. J Antimicrob Chemother. 2014;69:451–62.

17. Crowther GS, Baines SD, Todhunter SL, Freeman J, Chilton CH, Wilcox MH. Evaluation of NVB302 versus vancomycin activity in an *in vitro* human gut model of *Clostridium difficile* infection. J Antimicrob Chemother. 2013;68(1):168–76.

18. Saxton K, Baines SD, Freeman J, O’Connor R, Wilcox MH. Effects of exposure of *Clostridium difficile* PCR ribotypes 027 and 001 to fluoroquinolones in a human gut model. Antimicrob Agents Chemother. 2009;53(2):412–20.

19. Baines SD, Freeman J, Wilcox MH. Effects of piperacillin/tazobactam on *Clostridium difficile* growth and toxin production in a human gut model. J Antimicrob Chemother. 2005;55(6):974–82.

20. Baines SD, Saxton K, Freeman J, Wilcox MH. Tigecycline does not induce proliferation or cytotoxin production by epidemic *Clostridium difficile* strains in a human gut model. J Antimicrob Chemother. 2006;58(5):1062–5.

21. Swanson RN, Hardy DJ, Shipkowitz NL, Hanson CW, Ramer NC, Fernandes PB, et al. *In vitro* and *in vivo* evaluation of tiacumicins B and C against *Clostridium difficile*. Antimicrob Agents Chemother. 1991;35(6):1108–11.

22. Crook DW, Sarah Walker A, Kean Y, Weiss K, Cornely OA, Miller MA, et al. Fidaxomicin versus vancomycin for *Clostridium difficile* infection: Meta-analysis of pivotal randomized controlled trials. Clin Infect Dis. 2012;55(SUPPL.2):93–103.

23. Kurtz CB, Cannon EP, Brezzani A, Pitruzzello M, Dinardo C, Rinard E, et al. GT160-246, a toxin binding polymer for treatment of *Clostridium difficile* colitis. Antimicrob Agents Chemother. 2001;45(8):2340–7.

24. Johnson S, Louie TJ, Gerding DN, Cornely OA, Chasan-Taber S, Fitts D, et al. Vancomycin, metronidazole, or tolevamer for *Clostridium difficile* infection: Results from two multinational, randomized, controlled trials. Clin Infect Dis. 2014;59(3):345–54.

25. Baines SD, Freeman J, Wilcox MH. Tolevamer is not efficacious in the neutralization of cytotoxin in a human gut model of *Clostridium difficile* infection. Antimicrob Agents Chemother. 2009;53(5):2202–4.

26. Chilton CH, Crowther GS, Baines SD, Todhunter SL, Freeman J, Locher HH, et al. *In vitro* activity of cadazolid against clinically relevant *Clostridium difficile* isolates and in an *in vitro* gut model of *C. *difficile* infection*. J Antimicrob Chemother. 2014;69(3):697–705.

27. Chilton CH, Crowther GS, Todhunter SL, Nicholson S, Freeman J, Chesnel L, et al. Efficacy of surotomycin in an *in vitro* gut model of *Clostridium difficile* infection. J Antimicrob Chemother. 2014;69(9):2426–33.

28. Baines SD, Crowther GS, Freeman J, Todhunter S, Vickers R, Wilcox MH. SMT19969 as a treatment for *Clostridium difficile* infection: An assessment of antimicrobial activity using conventional susceptibility testing and an *in vitro* gut model. J Antimicrob Chemother. 2015;70(1):182–9.

29. Chilton CH, Crowther GS, Todhunter SL, Ashwin H, Longshaw CM, Karas A, et al. Efficacy of alternative fidaxomicin dosing regimens for treatment of simulated *Clostridium difficile* infection in an *in vitro* human gut model. J Antimicrob Chemother. 2015;70(9):2598–607.

30. Clinical trials [Internet]. Available from: https://clinicaltrials.gov/ct2/show/NCT02254967

31. Baines SD, Chilton CH, Crowther GS, Todhunter SL, Freeman J, Wilcox MH. Evaluation of antimicrobial activity of ceftaroline against *Clostridium difficile* and propensity to induce *C. *difficile* infection in an *in vitro* human gut model*. J Antimicrob Chemother. 2013;68(8):1842–9.

32. Chilton CH, Freeman J, Crowther GS, Todhunter SL, Nicholson S, Wilcox MH. Co-amoxiclav induces proliferation and cytotoxin production of *Clostridium difficile* ribotype 027 in a human gut model. J Antimicrob Chemother. 2012;67(4):951–4.

33. Public Health England. Management of infection guidance for primary care for consultation and local adaptation About Public Health England. 2016;1–75.

34. Abbott PJ, Chen J. WHO food additives series 46: Trehalose [Internet]. Available from: http://www.inchem.org/documents/jecfa/jecmono/v46je05.htm

35. Murata T, Yamato I, Kakinuma Y. Structure and Mechanism of Vacuolar Na+ Translocating ATPase From *Enterococcus hirae*. J Bioenerg Biomembr. 2005;37(6):411–3.

36. Ley RE, Backhed F, Turnbaugh P, Lozupone CA, Knight RD, Gordon JI. Obesity alters gut microbial ecology. Proc Natl Acad Sci. 2005;102(31):11070–5.

37. Nguyen TLA, Vieira-Silva S, Liston A, Raes J. How informative is the mouse for human gut microbiota research? Dis Model Mech. 2015;8:1–16.

38. Hufeldt MR, Nielsen DS, Vogensen FK, Midtvedt T, Hansen AK. Variation in the Gut Microbiota of Laboratory Mice Is Related to Both Genetic and Environmental Factors. Comp Med. 2010;60(5):336–42.

39. Selber-hnativ S, Rukundo B, Ahmadi M, Akoubi H, Baird A, Begum F, et al. Human Gut Microbiota: Toward an Ecology of Disease. 2017;8(July).

40. Rogosa M. The genus Veillonella. J Bacteriol. 1964;87(1):162–70.

41. Whatmore A, Chudek J, Reed R. The effects of osmotic upshock on the intracellular solute pools of *Bacillus subtilis*. J Gen Microbiol. 1990;136:2527–35.

42. Liu C, Finegold SM, Song Y, Lawson PA. Reclassification of *Clostridium coccoides, Ruminococcus hansenii, Ruminococcus hydrogenotrophicus, Ruminococcus luti, Ruminococcus productus* and *Ruminococcus schinkii* as *Blautia coccoides* gen. *nov., comb. nov., *Blautia hansenii**. Int J Syst Evol Microbiol. 2008;58:1896–902.

43. Zhang Y, DeBosch BJ. Microbial and metabolic impacts of trehalose and trehalose analogues. Gut Microbes. 2020 Sep 2;11(5):1475–82.

44. Chung W, Walker AW, Louis P, Parkhill J, Vermeiren J, Bosscher D, et al. Modulation of the human gut microbiota by dietary fibres occurs at the species level. BMC Biol. 2016;14:3.

45. Fachi JL, Felipe JDS, Farias A dos S, Varga-weisz P, Vinolo MAR. Butyrate Protects Mice from *Clostridium difficile*-Induced Colitis through an HIF-1-Dependent Mechanism. Cell Rep. 2019;27:750–61.

46. Khanna S, Montassier E, Schmidt B, Patel R, Knights D, Pardi DS, et al. Gut microbiome predictors of treatment response and recurrence in primary *Clostridium difficile* infection. Aliment Pharmacol Ther. 2016;44:715–27.

47. Roychowdhury S, Cadnum J, Glueck B, Obrenovich M, Donskey C, Cresci GAM, et al. *Faecalibacterium prausnitzii* and a Prebiotic Protect Intestinal Health in a Mouse Model of Antibiotic and *Clostridium difficile* Exposure. JPEN J Parenter Enter Nutr. 2018;42(7):1156–67.

48. Hsiao A, Ahmed AMS, Subramanian S, Griffin NW, Drewry LL, Jr WAP, et al. Members of the human gut microbiota involved in recovery from *Vibrio cholerae* infection. Nature. 2014;515:423.

49. Lee ASY, Song KP. LuxS/autoinducer-2 quorum sensing molecule regulates transcriptional virulence gene expression in *Clostridium difficile*. Biochem Biophys Res Commun. 2005;335(3):659–66.

50. Darkoh C, DuPont HL, Norris SJ, Kaplan HB. Toxin Synthesis by *Clostridium difficile* Is Regulated through Quorum Signaling. MBio. 2015;6(2):e02569–14.

51. Reeves AE, Theriot CM, Bergin IL, Huffnagle GB, Schloss PD, Young VB. The interplay between microbiome dynamics and pathogen dynamics in a murine model of *Clostridium difficile* infection. Gut Microbes. 2011;2(3):145–58.

52. Petrof EO, Gloor GB, Vanner SJ, Weese SJ, Carter D, Daigneault MC, et al. Stool substitute transplant therapy for the eradication of *Clostridium difficile* infection: ‘RePOOPulating’ the gut. Microbiome. 2013;1:1–12.

53. Milani C, Ticinesi A, Gerritsen J, Nouvenne A, Andrea Lugli G, Mancabelli L, et al. Gut microbiota composition and *Clostridium difficile* infection in hospitalized elderly individuals: A metagenomic study. Sci Rep. 2016;6(March):1–12.

54. Hryckowian AJ, Treuren W Van, Smits SA, Davis NM, Gardner JO, Bouley DM, et al. Microbiota-accessible carbohydrates suppress *Clostridium difficile* infection in a murine model. Nat Microbiol. 2018;

55. Jain NK, Roy I. Effect of trehalose on protein structure. Protein Sci. 2009;18:24–36.

56. Daquigan N, Seekatz AM, Greathouse KL, Young VB, White JR. High-resolution profiling of the gut microbiome reveals the extent of *Clostridium difficile* burden. npj Biofilms Microbiomes. 2017;35:1–8.

57. Girinathan B, Dibenedetto N, Worley J, Peltier J, Lavin R, Delaney D, et al. The mechanisms of *in vivo* commensal control of *Clostridioides difficile* virulence. BioRxiv. 2020;

58. Bouillaut L, Self WT, Sonenshein AL. Proline-dependent regulation of *Clostridium difficile* stickland metabolism. J Bacteriol. 2013;195(4):844–54.

59. Hofmann JD, Otto A, Berges M, Biedendieck R, Michel A, Becher D, et al. Metabolic Reprogramming of *Clostridioides difficile* During the Stationary Phase With the Induction of Toxin Production. Front Microbiol. 2018;9:1970.

60. Jenior ML, Leslie JL, Young VB, Schloss PD. *Clostridium difficile* Colonizes Alternative Nutrient Niches during Infection across Distinct Murine Gut Microbiomes. mSystems. 2017;2(4).

61. Moura IB, Buckley AM, Ewin D, Shearman S, Clark E, Wilcox MH, et al. Omadacycline Gut Microbiome Exposure Does Not Induce *Clostridium difficile* Proliferation or Toxin Production in a model that simulates the proximal, medial, and distal human colon. Antimicrob Agents Chemother. 2019;63(2):1–11.

62. Buckley AM, Spencer J, Candlish D, Irvine JJ, Douce GR. Infection of hamsters with the UK *Clostridium difficile* ribotype 027 outbreak strain R20291. J Med Microbiol. 2011;60(8):1174–80.

63. Martin M. Cutadapt removes adapter sequences from high-throughput sequencing reads. EMBnet.journal. 2011;17(1):10.

64. Schloss PD, Westcott SL, Ryabin T, Hall JR, Hartmann M, Hollister EB, et al. Introducing mothur: Open-source, platform-independent, community-supported software for describing and comparing microbial communities. Appl Environ Microbiol. 2009;75(23):7537–41.

65. Rognes T, Flouri T, Nichols B, Quince C, Mahé F. VSEARCH: a versatile open source tool for metagenomics. PeerJ. 2016;4:e2584.

66. Huson DH, Beier S, Flade I, Górska A, El-Hadidi M, Mitra S, et al. MEGAN Community Edition - Interactive Exploration and Analysis of Large-Scale Microbiome Sequencing Data. PLoS Comput Biol. 2016;12(6):1–12.

67. Buchfink B, Xie C, Huson DH. Fast and sensitive protein alignment using DIAMOND. Nat Methods. 2014;12(1):59–60.

68. Kal AJ, Zonneveld AJ Van, Benes V, Berg M Van Den, Koerkamp MG, Albermann K, et al. Dynamics of Gene Expression Revealed by Comparison of Serial Analysis of Gene Expression Different Carbon Sources. Mol Biol Cell. 1999;10(June):1859–72.

