## Supplementary Table 2 for "Pilot study: Trehalose-induced remodelling of the human microbiota affects *Clostridioides difficile* infection outcome in an *in vitro* colonic model"

| Experimental period | Day | Trehalose concentration (mM) | | | | | | | | | Glucose concentration (mM) | | | | | | | | |
|---|---|---|---|---|---|---|---|---|---|---|---|---|---|---|---|---|---|---|---|
| | | Model G | | | Model T | | | Model S | | | Model G | | | Model T | | | Model S | | |
| | | V1 | V2 | V3 | V1 | V2 | V3 | V1 | V2 | V3 | V1 | V2 | V3 | V1 | V2 | V3 | V1 | V2 | V3 |
| CD spore reservoir | 14 | ND | ND | ND | 5.2446 | 0.4353 | ND | ND | 0.0003 | ND | 6.98 | 0.0011 | 0.0021 | 0.0498 | 0.0253 | 0.0007 | 0.001 | 0.0007 | 0.0006 |
| | 15 | ND | ND | ND | 8.0957 | ND | ND | ND | ND | ND | 16.1177 | 0.0011 | 0.0017 | 0.0567 | 0.0003 | 0.0019 | 0.0017 | 0.0014 | 0.0066 |
| | 16 | 0.0128 | ND | 0.0004 | 4.1587 | 4.3267 | ND | ND | ND | ND | 0.1465 | 0.0067 | 0.0187 | 0.1018 | ND | ND | ND | ND | ND |
| | 17 | 0.0073 | ND | 0.0005 | 0.3459 | 0.0006 | ND | ND | ND | 0.0004 | 0.1272 | 0.0042 | 0.0029 | 0.0185 | 0.0018 | 0.0048 | 0.001 | 0.0009 | 0.0012 |
| | 18 | ND | ND | ND | 4.2475 | ND | ND | ND | ND | ND | 0.1478 | 0.0033 | ND | 0.0839 | 0.0018 | 0.0017 | 0.007 | 0.0021 | 0.0012 |
| | 19 | ND | ND | ND | 1.8968 | ND | ND | ND | ND | ND | 0.1343 | 0.0076 | 0.0014 | 0.1077 | 0.0012 | 0.0013 | 0.0009 | 0.0013 | 0.0009 |
| | 20 | ND | ND | ND | 0.3459 | ND | ND | ND | ND | ND | 0.0189 | 0.0108 | 0.0051 | ND | 0.0017 | 0.0024 | 0.0013 | 0.0014 | 0.001 |
| | 21 | ND | ND | ND | ND | ND | ND | ND | ND | ND | 0.0147 | 0.0034 | 0.0017 | ND | 0.001 | 0.0017 | 0.0015 | 0.0013 | 0.001 |
| | 22 | ND | ND | ND | 0.0502 | ND | ND | ND | ND | ND | 0.011 | 0.0015 | 0.0014 | 0.0025 | 0.0015 | 0.0011 | 0.0009 | 0.0014 | 0.0011 |
| | 23 | ND | ND | 0.0004 | ND | ND | ND | ND | ND | ND | 0.0009 | ND | 0.0235 | 0.0006 | 0.0012 | 0.0007 | 0.0015 | 0.001 | 0.0014 |
| Clindamycin | 24 | ND | ND | ND | 3.6687 | ND | ND | ND | ND | ND | 4.0395 | ND | 0.005 | 0.03 | 0.0015 | 0.0008 | 0.0014 | 0.0015 | 0.0013 |
| | 25 | 0.0583 | ND | ND | 11.7116 | 0.8146 | ND | 0.2221 | ND | ND | 28.252 | 0.3285 | ND | 0.4015 | 0.0163 | 0.0011 | 1.3918 | 0.0166 | 0.0016 |
| | 26 | 0.2536 | 0.007 | ND | 13.3425 | 6.6753 | 0.9993 | 0.4172 | 0.2168 | 0.021 | 27.8272 | 2.548 | 0.0147 | 1.6282 | 0.1154 | 0.0395 | 1.8408 | 0.5932 | 0.1239 |
| | 27 | 0.1788 | ND | ND | 8.2611 | 5.8113 | 2.5191 | ND | ND | ND | 20.1018 | 0.0954 | 0.0141 | 0.0552 | 0.1604 | 0.2399 | 0.0065 | 0.0027 | 0.0015 |
| | 28 | 0.0091 | 0.0127 | ND | 2.0596 | ND | ND | ND | ND | ND | 11.3207 | 0.0092 | 0.0051 | 0.059 | 0.0075 | 0.0024 | 0.0019 | 0.0013 | 0.001 |
| | 29 | 0.0048 | ND | ND | 2.8235 | ND | ND | ND | ND | ND | 8.9286 | 0.0046 | 0.008 | 0.1393 | 0.0042 | 0.0026 | 0.0013 | ND | ND |
| | 30 | ND | ND | ND | ND | ND | ND | ND | ND | ND | ND | ND | ND | 0.0016 | 0.0008 | 0.0009 | 0.0012 | 0.0047 | 0.0038 |
| | 31 | ND | ND | ND | 2.054 | ND | ND | ND | ND | ND | 6.7163 | 0.0032 | 0.001 | 0.1317 | 0.0016 | 0.0015 | 0.0049 | ND | ND |
| Simulated CDI | 32 | ND | ND | ND | 0.2552 | ND | ND | ND | ND | ND | 6.1944 | 0.0022 | 0.0007 | 0.1165 | 0.0049 | 0.0019 | 0.0051 | ND | ND |
| | 33 | ND | ND | ND | ND | ND | ND | ND | ND | ND | 0.0014 | 0.0021 | 0.0009 | 0.0046 | 0.002 | 0.0012 | 0.0078 | 0.0008 | ND |
| | 34 | ND | ND | ND | 1.1969 | ND | ND | ND | ND | ND | 7.0011 | 0.0021 | 0.001 | 0.2422 | 0.0042 | 0.0014 | 0.0053 | 0.0012 | ND |
| | 35 | ND | ND | ND | ND | ND | ND | ND | ND | ND | 0.0008 | 0.0137 | 0.0033 | ND | 0.0145 | ND | 0.0046 | ND | ND |
| | 36 | ND | ND | ND | 0.0112 | ND | ND | ND | ND | ND | 2.6103 | 0.0334 | 0.0168 | 0.0843 | 0.0652 | 0.0018 | 0.1078 | 0.001 | ND |
| | 37 | ND | ND | ND | ND | ND | ND | ND | ND | ND | 0.0009 | 0.0445 | 0.0116 | 0.0006 | 0.0129 | 0.0312 | ND | 0.001 | ND |
| | 38 | ND | ND | ND | ND | ND | ND | ND | ND | ND | 0.0017 | 0.0023 | 0.0013 | 0.001 | 0.0025 | 0.0009 | 0.0077 | 0.0018 | 0.0019 |
| | 39 | ND | ND | ND | 0.5673 | ND | ND | ND | ND | ND | 2.2232 | ND | 0.0009 | 0.6585 | 0.0007 | 0.0006 | 0.0089 | 0.0008 | 0.0009 |
| | 40 | ND | ND | ND | 2.1222 | ND | ND | ND | ND | ND | 6.8236 | 0.0019 | 0.0019 | 1.4805 | ND | ND | 0.0032 | 0.0011 | 0.0009 |
| | 41 | ND | ND | ND | 1.2728 | ND | ND | ND | ND | ND | 0.0031 | 0.0058 | ND | 0.0011 | 1.6883 | 0.0009 | 0.0081 | 0.0011 | 0.0018 |
| | 42 | ND | ND | ND | 2.2694 | ND | ND | ND | ND | ND | 0.0026 | 0.0058 | 0.0007 | 1.0114 | 0.0012 | 0.0019 | 0.0081 | 0.0009 | 0.0012 |
| | 43 | ND | ND | ND | ND | ND | ND | ND | ND | ND | 0.0065 | 0.0156 | 0.0022 | 0.0016 | 0.0011 | 0.0016 | 0.0035 | 0.0009 | 0.001 |
| | 44 | ND | ND | ND | 4.6967 | ND | ND | ND | ND | ND | 8.447 | 0.0145 | 0.0031 | 0.5768 | 0.0196 | 0.002 | 0.0044 | 0.001 | 0.0014 |
| | 45 | ND | ND | ND | 1.5605 | ND | ND | ND | ND | ND | 0.0102 | 0.0175 | 0.0087 | 0.337 | 0.0016 | 0.0022 | 0.0016 | 0.0013 | 0.0014 |
| | 46 | ND | ND | ND | ND | ND | ND | ND | ND | ND | 0.0046 | 0.0099 | 0.0027 | 0.0029 | 0.0126 | 0.0014 | 0.0013 | 0.0008 | 0.0007 |
| | 47 | ND | ND | ND | ND | ND | ND | | | | ND | ND | ND | ND | ND | ND | | | |
| | 48 | ND | ND | ND | ND | ND | ND | | | | 0.0022 | 0.0015 | 0.0013 | 0.0041 | 0.0008 | 0.0015 | | | |
| | 49 | ND | ND | ND | ND | ND | ND | | | | 0.0016 | 0.0013 | 0.0009 | 0.003 | 0.0011 | 0.001 | | | |
| | 50 | ND | ND | ND | ND | ND | ND | | | | 0.0027 | 0.0018 | 0.0016 | 0.0077 | 0.0009 | 0.0016 | | | |
