## Supplementary material for "Pilot study: Trehalose-induced remodelling of the human microbiota affects *Clostridioides difficile* infection outcome in an *in vitro* colonic model"

**Supplementary Table 1.** Comparison of bacterial families from the donors and the faecal slurry

| Bacterial family | Abundance (%) | | | | | |
|---|---|---|---|---|---|---|
| | Slurry | Donor A | Donor B | Donor C | Donor D | Donor E |
| Methanobacteriaceae | 0.042 | 0.041 | 0.035 | 0.111 | 0.003 | 0.000 |
| Actinomycetaceae | 0.006 | 0.000 | 0.001 | 0.017 | 0.000 | 0.007 |
| Bifidobacteriaceae | 6.525 | 5.504 | 5.701 | 4.932 | 0.024 | 15.795 |
| Coriobacteriaceae | 1.387 | 1.744 | 1.527 | 0.292 | 0.041 | 3.713 |
| Bacteroidales | 7.509 | 11.808 | 11.848 | 0.000 | 0.000 | 0.001 |
| Barnesiellaceae | 0.050 | 0.037 | 0.035 | 0.031 | 0.000 | 0.098 |
| Odoribacteraceae | 0.033 | 0.009 | 0.012 | 0.099 | 0.000 | 0.057 |
| Paraprevotellaceae | 0.955 | 1.531 | 1.283 | 0.000 | 0.000 | 0.000 |
| Bacteroidaceae | 12.905 | 7.875 | 8.530 | 12.540 | 20.431 | 13.921 |
| Porphyromonadaceae | 0.660 | 0.423 | 0.471 | 0.664 | 0.050 | 1.653 |
| Prevotellaceae | 0.693 | 1.105 | 0.953 | 0.000 | 0.000 | 0.000 |
| Rikenellaceae | 1.689 | 2.437 | 1.939 | 5.256 | 0.473 | 0.205 |
| Muribaculaceae | 0.006 | 0.000 | 0.000 | 0.144 | 0.000 | 0.000 |
| Lactobacillaceae | 1.510 | 0.003 | 0.004 | 0.003 | 9.777 | 0.000 |
| Streptococcaceae | 0.108 | 0.067 | 0.075 | 0.274 | 0.067 | 0.204 |
| Turicibacteraceae | 0.152 | 0.170 | 0.202 | 0.074 | 0.000 | 0.001 |
| Clostridiales | 10.576 | 13.343 | 14.081 | 18.764 | 6.648 | 11.586 |
| Christensenellaceae | 0.942 | 1.379 | 1.459 | 2.986 | 0.000 | 0.001 |
| Clostridiaceae | 3.998 | 4.984 | 4.872 | 4.056 | 0.338 | 0.912 |
| Eubacteriaceae | 0.012 | 0.000 | 0.000 | 0.086 | 0.000 | 0.000 |
| Lachnospiraceae | 13.547 | 10.156 | 10.459 | 1.789 | 20.529 | 18.324 |
| Peptococcaceae | 0.265 | 0.487 | 0.508 | 0.000 | 0.000 | 0.000 |
| Peptostreptococcaceae | 0.037 | 0.049 | 0.032 | 0.048 | 0.000 | 0.001 |
| Ruminococcaceae | 23.302 | 29.027 | 27.642 | 24.854 | 17.570 | 26.237 |
| Veillonellaceae | 2.158 | 2.847 | 2.822 | 0.888 | 2.216 | 1.655 |
| Erysipelotrichaceae | 1.422 | 1.251 | 1.392 | 0.331 | 1.850 | 2.492 |
| Victivallaceae | 0.443 | 1.002 | 1.344 | 0.000 | 0.000 | 0.001 |
| Unknown Family - RF32 | 0.482 | 0.720 | 0.523 | 3.605 | 0.373 | 0.027 |
| Alcaligenaceae | 0.912 | 0.597 | 0.631 | 0.703 | 1.100 | 2.532 |
| Desulfovibrionaceae | 0.370 | 0.669 | 0.749 | 0.096 | 0.024 | 0.445 |
| Enterobacteriaceae | 4.729 | 0.039 | 0.068 | 0.309 | 16.911 | 0.041 |
| Pasteurellaceae | 0.017 | 0.061 | 0.065 | 0.115 | 0.001 | 0.032 |
| Anaeroplasmataceae | 0.161 | 0.396 | 0.520 | 0.000 | 0.000 | 0.000 |
| Unknown Family - RF39 | 0.012 | 0.020 | 0.018 | 0.000 | 0.000 | 0.000 |
| Unknown Family - ML615J-28 | 0.105 | 0.000 | 0.000 | 3.686 | 0.000 | 0.006 |
| Verrucomicrobiaceae | 2.283 | 0.220 | 0.200 | 13.247 | 1.574 | 0.051 |

**Supplementary Table 2.** Glucose and trehalose concentrations from model G (glucose supplemented), model T (trehalose supplemented), and model S (saline supplemented) (vessels 1-3 for each model). File name: Supplemental Table 2 sugar concentrations.

**Supplementary Table 3.** Day 38 KEGG pathway analysis from model G (glucose supplemented), model T (trehalose supplemented), and model S (saline supplemented) (vessel 3 for each model). Data shown are KEGG pathway assigned reads from four technical replicates for each model. File name: Supplementary Table 3 KEGG pathway.

**Supplementary Figure 1.** Recovery of bacterial populations from vessel 3 of model S, (S), model G, (G) and model T (T). The bacterial populations enumerated were, total obligate anaerobic bacteria (black lines), lactose-fermenting *Enterobacteriaceae* (red lines), *Clostridium* spp. (green lines), *Lactobacillus* spp. (blue lines), and *C. difficile* total counts (indigo lines). Results expressed as mean ± SD of three technical replicates. Horizontal blue arrow represents the period of clindamycin dosing to all gut models.

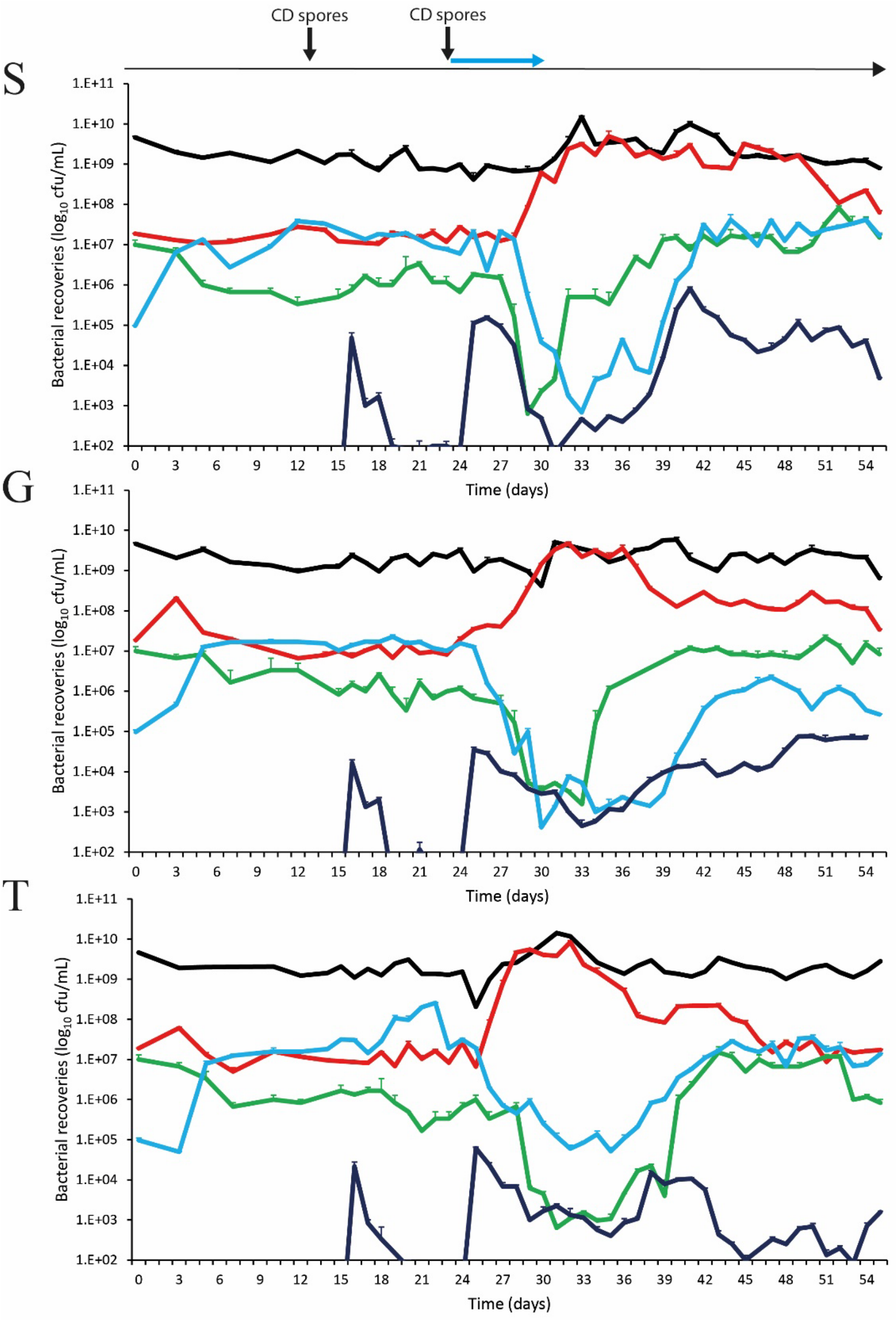

**Supplementary Figure 2.** Biosynthesis of amino acids (KEGG pathway Ko01230), with metabolic compounds produced from pathways that are most abundant in both model G and model S (CDI induction model) shown in red.

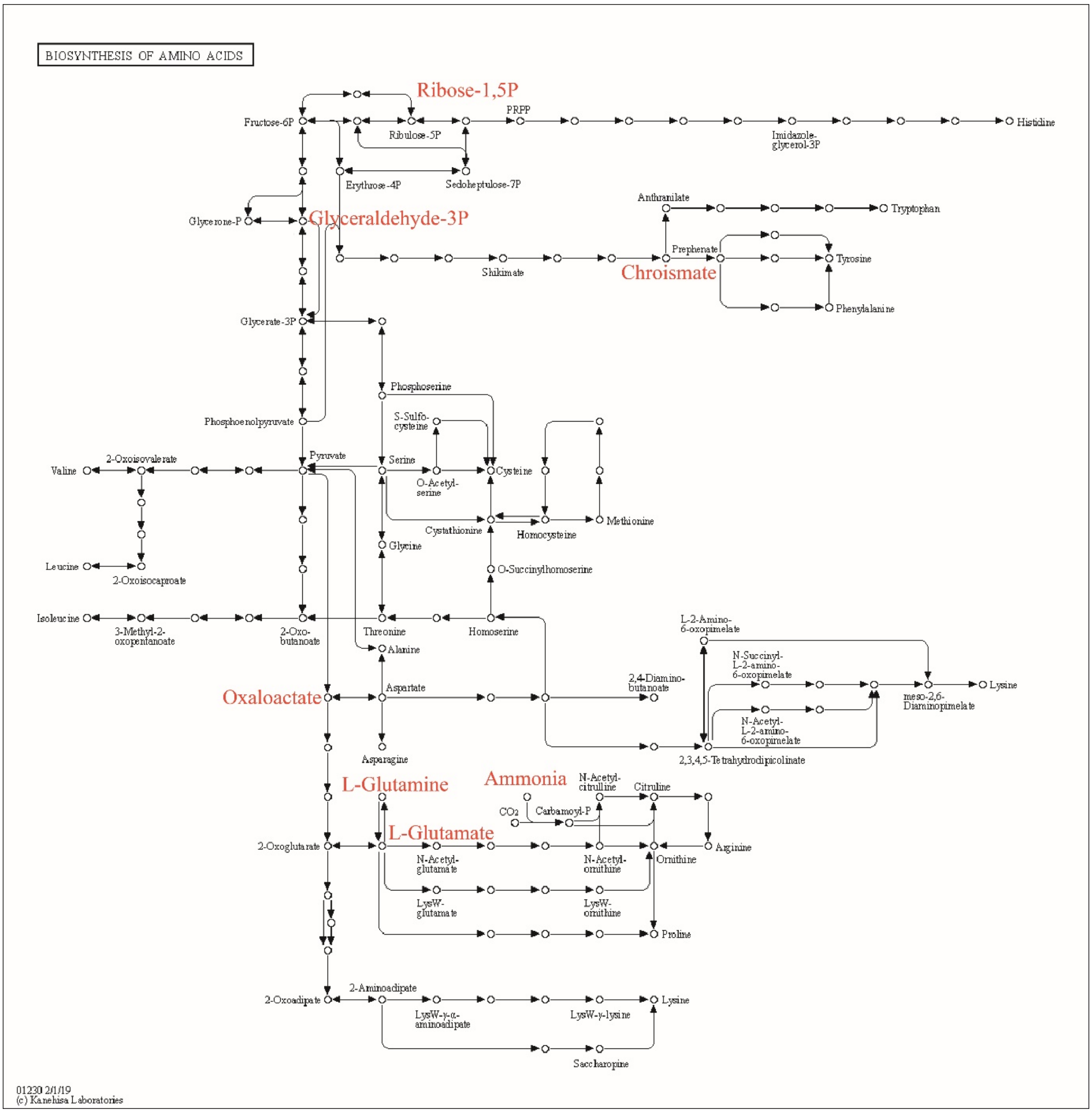

**Supplementary Figure 3.** Putative mechanisms of trehalose-induced protection against *C. difficile* infection.

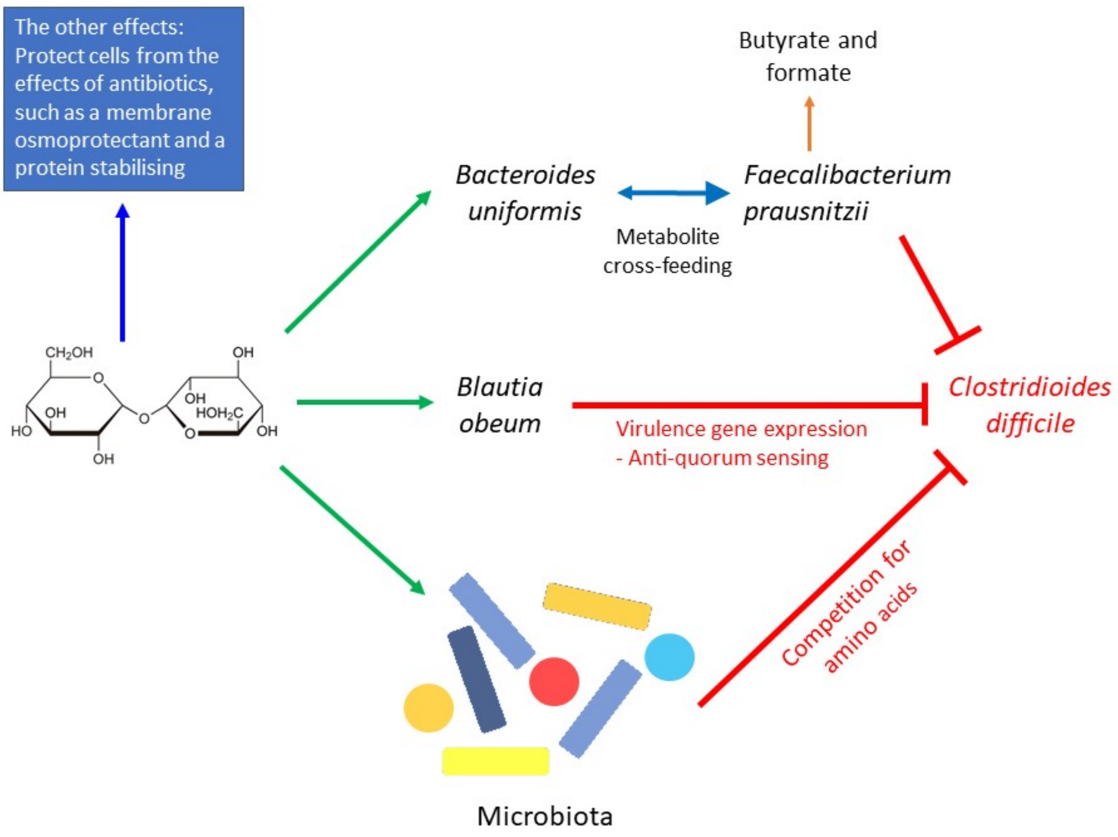

### Supplementary methods

*Enumeration of endogenous bacteria*

Models were sampled for culture profiling of key intestinal microbiota populations using selective and non-selective agars described in **Supplementary Table 4**. Populations of total bacteria, *Clostridium* spp. lactose-fermenting Enterobacteriaceae, *Enterococcus* spp., *Bacteroides* spp., *Bifidobacterium* spp., *Lactobacillus* spp., *C. difficile* total viable cells and *C. difficile* spores. *C. difficile* spores were isolated by treating 0.5 mL of gut model fluid with 0.5 mL of 96% ethanol. The samples were incubated at room temperature for 1 h, serially diluted to 10–3 in peptone water, and 20 µL of each sample dilution was plated in triplicate onto supplemented Braziers CCEY agar (**Supplementary Table 3**). Plates were incubated anaerobically for 48 h and distinctive colonies were enumerated and identified based on colony morphology and Matrix-Assisted Laser Desorption Ionization-Time of Flight (MALDI-TOF) identification. Each bacterial population was measured in triplicate (three technical replicates of a single biological replicate) in vessels 2 and 3. The limit of detection for either total viable counts or spores were 1.2 or l.5, respectively, $\log_{10}$ cfu/mL.

**Supplementary Table 4.** Target populations and agar composition for bacterial enumeration.

| Target populations | Agar | Supplements | Incubation (temp, environment) |
|---|---|---|---|
| Total anaerobes and total *Clostridium* spp. | Fastidious anaerobe agar | 5% horse blood | 37ºC, anerobic |
| *Bifidobacterium* spp. | 42.5 g/L Columbia agar, and 5 g/L agar technical | 0.5 g/L cysteine HCl, 5 g/L glucose | 37ºC, anerobic |
| *Bacteroides spp.* | Bacteroides bile aesculin agar | 5mg/L haemin, 10 µL/L vitamin K, 7.5 mg/L vancomycin, 1 mg/L penicillin, 75 mg/L kanamycin and 10 mg/L colistin | 37ºC, anerobic |
| *Lactobacillus* spp. | 52.2 g/L MRS broth and 20 g/L agar technical | 0.5 g/L cysteine hydrocloride, 20 mg/L vancomycin | 37ºC, anerobic |
| Total facultative anaerobes | Nutrient agar | N/A | 37ºC, aerobic |
| Lactose fermenting Enterobacteriaceae | MaConkey's agar | N/A | 37ºC, aerobic |
| *Enterococcus* spp. | *Kanamycin* aesculin *azide agar* | 10 mg/L nalidixic acid, 10 mg/L aztreonam, and 20 mg/L kanamycin | 37ºC, aerobic |
| Total spores (following alcohol shock for 1 hour) | Fastidious anaerobe agar | 5% horse blood | 37ºC, anerobic |
| *C. difficile* total viable cells | Braziers CCEY agar | D-cycloserine (250mg/L) cefoxitin (8mg/L), 5 mg/L lysozyme, and 20 mL/L lysed horse blood | 37ºC, anerobic |
| *C. difficile* spores | Braziers CCEY agar | 5 mg/L lysozyme, and 2% lysed horse blood | 37ºC, anerobic |

**Supplementary Table 5.** Ion chromatography conditions used to detect glucose and trehalose

Eluent Buffer

A: Water, B; 300 mM NaOH, C; 300 mM NaOH/1,500 mM sodium acetate

| Time (min) | Flow (ml/min) | A% | B% | C% | |
|---|---|---|---|---|---|
| -10 | 1 | 86.8 | 13.2 | 0.0 | |
| 0 | 1 | 86.8 | 13.2 | 0.0 | |
| 15 | 1 | 86.8 | 13.2 | 0.0 | Linear gradient |
| 25 | 1 | 67.0 | 0.0 | 33.0 | |
| 30 | 1 | 67.0 | 0.0 | 33.0 | |
