## Supplementary Table 3 for "Pilot study: Trehalose-induced remodelling of the human microbiota affects *Clostridioides difficile* infection outcome in an *in vitro* colonic model"

| | Saline model | | | | Trehalose model | | | | Glucose model | | | |
|---|---|---|---|---|---|---|---|---|---|---|---|---|
| #Datasets | SA D38 | SB D38 | SC D38 | SD D38 | TA D38 | TB D38 | TC D38 | TD D38 | GA D38 | GB D38 | GC D38 | GD D38 |
| KEGG | 9253574 | 9230269 | 9114938 | 9229998 | 8411070 | 8468258 | 8365147 | 8417468 | 8612545 | 8645575 | 8550853 | 8644083 |
| Metabolism | 3324452 | 3323200 | 3279165 | 3319148 | 3197007 | 3210979 | 3186753 | 3196959 | 3255493 | 3256410 | 3228229 | 3257346 |
| Carbohydrate metabolism | 1110504 | 1115680 | 1106843 | 1118697 | 1179116 | 1186477 | 1181704 | 1176146 | 1229890 | 1238874 | 1228443 | 1236169 |
| ko00010 Glycolysis / Gluconeogenesis | 173787 | 174835 | 172940 | 174648 | 172844 | 175917 | 173790 | 173529 | 184557 | 186962 | 184294 | 185290 |
| K00001 alcohol dehydrogenase [EC:1.1.1.1] | 1456 | 1456 | 1473 | 1402 | 1185 | 1259 | 1208 | 1141 | 1839 | 1925 | 1870 | 1902 |
| K00002 alcohol dehydrogenase (NADP+) [EC:1.1.1.2] | 174 | 169 | 149 | 148 | 43 | 37 | 52 | 65 | 306 | 308 | 336 | 335 |
| K00016 L-lactate dehydrogenase [EC:1.1.1.27] | 1548 | 1452 | 1512 | 1681 | 610 | 687 | 665 | 615 | 1685 | 1888 | 1825 | 1772 |
| K00114 alcohol dehydrogenase (cytochrome c) [EC:1.1.2.8] | 2545 | 2673 | 2492 | 2486 | 1293 | 1237 | 1225 | 1497 | 867 | 813 | 764 | 751 |
| K00121 S-(hydroxymethyl)glutathione dehydrogenase / alcohol dehydrogenase [EC:1.1.1.284 1.1.1.1] | 2258 | 2350 | 2107 | 2222 | 1410 | 1419 | 1333 | 1335 | 1558 | 1506 | 1505 | 1497 |
| K00128 aldehyde dehydrogenase (NAD+) [EC:1.2.1.3] | 5102 | 5084 | 4807 | 4818 | 3106 | 2867 | 2785 | 3017 | 1916 | 1754 | 1686 | 1825 |
| K00129 aldehyde dehydrogenase (NAD(P)+) [EC:1.2.1.5] | 16 | 16 | 10 | 10 | 14 | 12 | 15 | 10 | 3 | 4 | 6 | 4 |
| K00131 glyceraldehyde-3-phosphate dehydrogenase (NADP+) [EC:1.2.1.9] | 2124 | 2160 | 1918 | 2096 | 1063 | 1014 | 920 | 1156 | 658 | 722 | 700 | 712 |
| K00134 glyceraldehyde 3-phosphate dehydrogenase [EC:1.2.1.12] | 6507 | 6759 | 6631 | 6623 | 5652 | 5731 | 5693 | 5624 | 5233 | 5374 | 5245 | 5450 |
| K00138 aldehyde dehydrogenase [EC:1.2.1.-] | 4815 | 4669 | 4149 | 4632 | 2677 | 2513 | 2491 | 2640 | 2411 | 2391 | 2308 | 2483 |
| K00149 aldehyde dehydrogenase family 9 member A1 [EC:1.2.1.47 1.2.1.3] | 1 | 1 | 0 | 0 | 0 | 0 | 0 | 0 | 0 | 0 | 0 | 0 |
| K00150 glyceraldehyde-3-phosphate dehydrogenase (NAD(P)) [EC:1.2.1.59] | 0 | 2 | 2 | 1 | 0 | 3 | 2 | 3 | 4 | 12 | 6 | 13 |
| K00161 pyruvate dehydrogenase E1 component alpha subunit [EC:1.2.4.1] | 1703 | 1663 | 1418 | 1432 | 800 | 804 | 841 | 919 | 537 | 508 | 532 | 580 |
| K00162 pyruvate dehydrogenase E1 component beta subunit [EC:1.2.4.1] | 1648 | 1616 | 1373 | 1463 | 827 | 781 | 798 | 906 | 569 | 462 | 495 | 495 |
| K00163 pyruvate dehydrogenase E1 component [EC:1.2.4.1] | 3440 | 3384 | 3516 | 3516 | 2237 | 2394 | 2149 | 2183 | 2815 | 3102 | 2675 | 2964 |
| K00169 pyruvate ferredoxin oxidoreductase alpha subunit [EC:1.2.7.1] | 216 | 258 | 255 | 233 | 319 | 373 | 370 | 340 | 472 | 523 | 516 | 565 |
| K00170 pyruvate ferredoxin oxidoreductase beta subunit [EC:1.2.7.1] | 87 | 91 | 115 | 86 | 271 | 291 | 310 | 317 | 194 | 186 | 215 | 227 |
| K00171 pyruvate ferredoxin oxidoreductase delta subunit [EC:1.2.7.1] | 5 | 16 | 4 | 7 | 66 | 59 | 58 | 50 | 24 | 29 | 38 | 34 |
| K00172 pyruvate ferredoxin oxidoreductase gamma subunit [EC:1.2.7.1] | 92 | 139 | 139 | 122 | 156 | 171 | 181 | 166 | 270 | 280 | 293 | 294 |
| K00174 2-oxoglutarate/2-oxoacid ferredoxin oxidoreductase subunit alpha [EC:1.2.7.3 1.2.7.11] | 2742 | 2915 | 3097 | 2864 | 6669 | 6734 | 6879 | 6802 | 5579 | 5516 | 5922 | 5513 |
| K00175 2-oxoglutarate/2-oxoacid ferredoxin oxidoreductase subunit beta [EC:1.2.7.3 1.2.7.11] | 1687 | 1853 | 2015 | 1903 | 4182 | 4181 | 4270 | 4209 | 3675 | 3591 | 3674 | 3509 |
| K00382 dihydrolipoamide dehydrogenase [EC:1.8.1.4] | 9170 | 9150 | 8930 | 9118 | 9722 | 9692 | 9339 | 9543 | 8930 | 8583 | 8861 | 8866 |
| K00627 pyruvate dehydrogenase E2 component (dihydrolipoamide acetyltransferase) [EC:2.3.1.12] | 5358 | 5056 | 4803 | 4842 | 2778 | 2862 | 2887 | 3162 | 3520 | 3384 | 3306 | 3636 |
| K00844 hexokinase [EC:2.7.1.1] | 664 | 727 | 715 | 699 | 2558 | 2621 | 2670 | 2708 | 1967 | 1805 | 2044 | 1925 |
| K00845 glucokinase [EC:2.7.1.2] | 4977 | 5043 | 5103 | 5062 | 7634 | 7716 | 7883 | 7923 | 7174 | 7422 | 7334 | 7363 |
| K00850 6-phosphofructokinase 1 [EC:2.7.1.11] | 2702 | 2797 | 2789 | 2795 | 3416 | 3618 | 3575 | 3393 | 3769 | 3840 | 3773 | 3818 |
| K00873 pyruvate kinase [EC:2.7.1.40] | 7147 | 7129 | 7132 | 6836 | 6764 | 6674 | 6742 | 6952 | 7200 | 6971 | 7122 | 6903 |
| K00886 polyphosphate glucokinase [EC:2.7.1.63] | 2 | 2 | 2 | 3 | 2 | 0 | 0 | 0 | 8 | 3 | 13 | 9 |
| K00895 diphosphate-dependent phosphofructokinase [EC:2.7.1.90] | 937 | 1130 | 1188 | 1108 | 3502 | 3491 | 3555 | 3299 | 2447 | 2397 | 2481 | 2285 |
| K00918 ADP-dependent phosphofructokinase/glucokinase [EC:2.7.1.146 2.7.1.147] | 8 | 4 | 1 | 2 | 13 | 16 | 3 | 9 | 31 | 33 | 26 | 20 |
| K00927 phosphoglycerate kinase [EC:2.7.2.3] | 4518 | 4471 | 4418 | 4463 | 4657 | 4921 | 4595 | 4509 | 4638 | 4537 | 4454 | 4548 |
| K01085 glucose-1-phosphatase [EC:3.1.3.10] | 761 | 684 | 690 | 750 | 471 | 485 | 500 | 421 | 969 | 1082 | 956 | 998 |
| K01086 fructose-1,6-bisphosphatase I / sedoheptulose-1,7-bisphosphatase [EC:3.1.3.11 3.1.3.37] | 1 | 1 | 0 | 1 | 2 | 0 | 0 | 0 | 0 | 0 | 2 | 1 |
| K01222 6-phospho-beta-glucosidase [EC:3.2.1.86] | 2574 | 2700 | 2428 | 2620 | 978 | 1072 | 1015 | 963 | 4405 | 4885 | 4587 | 4583 |
| K01223 6-phospho-beta-glucosidase [EC:3.2.1.86] | 8415 | 8540 | 8261 | 8631 | 5678 | 6061 | 5781 | 5438 | 9521 | 10202 | 9329 | 10037 |
| K01596 phosphoenolpyruvate carboxykinase (GTP) [EC:4.1.1.32] | 36 | 22 | 30 | 21 | 43 | 23 | 42 | 31 | 45 | 20 | 34 | 33 |
| K01610 phosphoenolpyruvate carboxykinase (ATP) [EC:4.1.1.49] | 6524 | 6393 | 6680 | 6622 | 6704 | 6697 | 6816 | 6676 | 7149 | 7323 | 7398 | 7118 |
| K01622 fructose 1,6-bisphosphate aldolase/phosphatase [EC:4.1.2.13 3.1.3.11] | 2 | 0 | 0 | 0 | 1 | 4 | 0 | 0 | 2 | 13 | 3 | 7 |
| K01623 fructose-bisphosphate aldolase, class I [EC:4.1.2.13] | 167 | 167 | 157 | 154 | 32 | 55 | 55 | 37 | 349 | 303 | 338 | 327 |
| K01624 fructose-bisphosphate aldolase, class II [EC:4.1.2.13] | 6833 | 6723 | 6594 | 6717 | 5602 | 5734 | 5627 | 5650 | 6507 | 7002 | 6741 | 6911 |
| K01689 enolase [EC:4.2.1.11] | 4355 | 4359 | 4446 | 4327 | 5171 | 5542 | 5258 | 5229 | 4613 | 4407 | 4462 | 4639 |
| K01785 aldose 1-epimerase [EC:5.1.3.3] | 3982 | 4387 | 4311 | 4359 | 8224 | 8264 | 8357 | 8134 | 7455 | 7509 | 7425 | 7157 |
| K01792 glucose-6-phosphate 1-epimerase [EC:5.1.3.15] | 2168 | 2177 | 2047 | 2169 | 1293 | 1220 | 1206 | 1401 | 1221 | 1232 | 1200 | 1285 |
| K01803 triosephosphate isomerase (TIM) [EC:5.3.1.1] | 3258 | 3528 | 3483 | 3510 | 3358 | 3562 | 3401 | 3364 | 3850 | 3893 | 3906 | 3869 |
| K01810 glucose-6-phosphate isomerase [EC:5.3.1.9] | 5895 | 5831 | 6010 | 5964 | 5767 | 5779 | 5575 | 5595 | 5661 | 5601 | 5546 | 5744 |
| K01834 2,3-bisphosphoglycerate-dependent phosphoglycerate mutase [EC:5.4.2.11] | 1383 | 1365 | 1451 | 1352 | 2251 | 2339 | 2237 | 2282 | 2061 | 2089 | 2085 | 1930 |
| K01835 phosphoglucomutase [EC:5.4.2.2] | 3966 | 3823 | 3976 | 4107 | 4928 | 5010 | 5112 | 4791 | 5614 | 5460 | 5373 | 5604 |
| K01895 acetyl-CoA synthetase [EC:6.2.1.1] | 7843 | 7868 | 7754 | 7877 | 7723 | 7669 | 7743 | 7792 | 6178 | 6178 | 6273 | 6256 |
| K01905 acetate---CoA ligase (ADP-forming) subunit alpha [EC:6.2.1.13] | 13 | 20 | 12 | 8 | 0 | 0 | 0 | 0 | 0 | 3 | 0 | 0 |
| K02446 fructose-1,6-bisphosphatase II [EC:3.1.3.11] | 726 | 766 | 654 | 713 | 479 | 511 | 581 | 482 | 1109 | 1037 | 1015 | 1030 |
| K02752 PTS system, beta-glucoside (arbutin/salicin/cellobiose)-specific IIB component [EC:2.7.1.-] | 926 | 959 | 888 | 953 | 611 | 676 | 593 | 607 | 1247 | 1241 | 1232 | 1296 |
| K02777 PTS system, sugar-specific IIA component [EC:2.7.1.-] | 594 | 593 | 591 | 628 | 237 | 312 | 232 | 257 | 595 | 690 | 650 | 643 |
| K02778 PTS system, glucose-specific IIB component [EC:2.7.1.199] | 135 | 148 | 151 | 171 | 98 | 119 | 110 | 112 | 192 | 195 | 170 | 180 |
| K02779 PTS system, glucose-specific IIC component | 901 | 892 | 919 | 939 | 549 | 632 | 604 | 624 | 1247 | 1329 | 1351 | 1264 |
| K02790 PTS system, maltose/glucose-specific IIB component [EC:2.7.1.199 2.7.1.208] | 1360 | 1290 | 1245 | 1307 | 1098 | 1058 | 1100 | 1159 | 1403 | 1413 | 1351 | 1547 |
| K02791 PTS system, maltose/glucose-specific IIC component | 0 | 0 | 0 | 0 | 0 | 1 | 0 | 0 | 0 | 0 | 0 | 0 |
| K03737 pyruvate-ferredoxin/flavodoxin oxidoreductase [EC:1.2.7.1 1.2.7.-] | 7665 | 7761 | 7814 | 8173 | 8670 | 9001 | 8957 | 8852 | 11378 | 12075 | 11511 | 11442 |
| K03841 fructose-1,6-bisphosphatase I [EC:3.1.3.11] | 2218 | 2283 | 2214 | 2256 | 1383 | 1537 | 1396 | 1412 | 1620 | 1579 | 1627 | 1562 |
| K04022 alcohol dehydrogenase | 497 | 448 | 424 | 491 | 151 | 116 | 141 | 148 | 1108 | 973 | 1036 | 1071 |
| K04041 fructose-1,6-bisphosphatase III [EC:3.1.3.11] | 2186 | 2438 | 2585 | 2428 | 3966 | 4037 | 4107 | 3867 | 3809 | 3871 | 3716 | 3695 |
| K04072 acetaldehyde dehydrogenase / alcohol dehydrogenase [EC:1.2.1.10 1.1.1.1] | 2747 | 2670 | 2860 | 2783 | 1272 | 1265 | 1220 | 1306 | 3039 | 3274 | 3244 | 3201 |
| K06859 glucose-6-phosphate isomerase, archaeal [EC:5.3.1.9] | 8 | 10 | 14 | 10 | 8 | 9 | 12 | 4 | 39 | 56 | 39 | 53 |
| K10705 glyceraldehyde-3-phosphate dehydrogenase, spermatogenic [EC:1.2.1.12] | 1 | 1 | 0 | 0 | 0 | 0 | 0 | 0 | 0 | 0 | 0 | 0 |
| K11532 fructose-1,6-bisphosphatase II / sedoheptulose-1,7-bisphosphatase [EC:3.1.3.11 3.1.3.37] | 0 | 0 | 0 | 1 | 3 | 0 | 1 | 0 | 0 | 6 | 5 | 1 |
| K11645 fructose-bisphosphate aldolase, class I [EC:4.1.2.13] | 1631 | 1712 | 1832 | 1694 | 2982 | 3093 | 3088 | 2932 | 3033 | 3160 | 3008 | 3063 |
| K13953 alcohol dehydrogenase, propanol-preferring [EC:1.1.1.1] | 2060 | 2096 | 1985 | 1950 | 996 | 973 | 1021 | 1096 | 1236 | 1286 | 1235 | 1182 |
| K13954 alcohol dehydrogenase [EC:1.1.1.1] | 1899 | 1895 | 2188 | 2184 | 1270 | 1379 | 1287 | 1201 | 753 | 881 | 795 | 743 |
| K14085 aldehyde dehydrogenase family 7 member A1 [EC:1.2.1.31 1.2.1.8 1.2.1.3] | 0 | 0 | 0 | 0 | 0 | 1 | 0 | 0 | 0 | 0 | 0 | 0 |
| K15633 2,3-bisphosphoglycerate-independent phosphoglycerate mutase [EC:5.4.2.12] | 5655 | 5458 | 5484 | 5488 | 6065 | 6093 | 5980 | 5951 | 5807 | 5664 | 5764 | 5463 |
| K15634 probable phosphoglycerate mutase [EC:5.4.2.12] | 581 | 573 | 548 | 581 | 486 | 515 | 441 | 490 | 569 | 634 | 616 | 651 |
| K15635 2,3-bisphosphoglycerate-independent phosphoglycerate mutase [EC:5.4.2.12] | 1007 | 1018 | 1124 | 1052 | 2695 | 2757 | 2840 | 2675 | 2410 | 2236 | 2357 | 2276 |
| K15778 phosphomannomutase / phosphoglucomutase [EC:5.4.2.8 5.4.2.2] | 4256 | 4121 | 4159 | 4126 | 2399 | 2347 | 2312 | 2521 | 1575 | 1409 | 1453 | 1439 |
| K16306 fructose-bisphosphate aldolase / 2-amino-3,7-dideoxy-D-threo-hept-6-ulosonate synthase [EC:4.1.2.13 2.2.1.10] | 2 | 0 | 0 | 1 | 0 | 0 | 3 | 0 | 7 | 6 | 7 | 12 |
| K16370 6-phosphofructokinase 2 [EC:2.7.1.11] | 570 | 526 | 436 | 508 | 372 | 357 | 384 | 374 | 714 | 786 | 704 | 752 |
| K20118 PTS system, glucose-specific IIC component | 399 | 375 | 347 | 366 | 460 | 538 | 469 | 488 | 279 | 298 | 251 | 310 |
| K20866 glucose-1-phosphatase [EC:3.1.3.10] | 783 | 829 | 759 | 788 | 1713 | 1782 | 1660 | 1720 | 1473 | 1526 | 1432 | 1526 |
| K21071 ATP-dependent phosphofructokinase / diphosphate-dependent phosphofructokinase [EC:2.7.1.11 2.7.1.90] | 2786 | 2764 | 2797 | 2845 | 2883 | 2973 | 2874 | 2780 | 3195 | 3405 | 3320 | 3209 |
| K22224 acetate---CoA ligase (ADP-forming) subunit beta [EC:6.2.1.13] | 369 | 386 | 329 | 350 | 146 | 141 | 158 | 174 | 821 | 861 | 722 | 882 |
| ko00020 Citrate cycle (TCA cycle) | 120604 | 120221 | 119660 | 120947 | 124566 | 126571 | 125497 | 125566 | 128898 | 130185 | 129169 | 129342 |
| K00024 malate dehydrogenase [EC:1.1.1.37] | 1872 | 2021 | 2084 | 2002 | 4954 | 4766 | 4938 | 4822 | 4052 | 3976 | 3945 | 3824 |
| K00030 isocitrate dehydrogenase (NAD+) [EC:1.1.1.41] | 441 | 430 | 420 | 544 | 171 | 197 | 152 | 213 | 1001 | 1063 | 1071 | 1111 |
| K00031 isocitrate dehydrogenase [EC:1.1.1.42] | 5104 | 4863 | 4952 | 5034 | 4691 | 4819 | 4747 | 4729 | 4219 | 4124 | 4277 | 4186 |
| K00116 malate dehydrogenase (quinone) [EC:1.1.5.4] | 3904 | 3931 | 3863 | 4028 | 2189 | 2100 | 2060 | 2043 | 3112 | 3383 | 3160 | 3334 |
| K00164 2-oxoglutarate dehydrogenase E1 component [EC:1.2.4.2] | 2459 | 2319 | 2297 | 2442 | 1596 | 1686 | 1614 | 1642 | 2667 | 2841 | 2690 | 2800 |
| K00176 2-oxoglutarate ferredoxin oxidoreductase subunit delta [EC:1.2.7.3] | 264 | 316 | 255 | 299 | 517 | 557 | 459 | 401 | 544 | 478 | 556 | 539 |
| K00177 2-oxoglutarate ferredoxin oxidoreductase subunit gamma [EC:1.2.7.3] | 705 | 614 | 742 | 715 | 1048 | 1068 | 1133 | 1059 | 1123 | 1156 | 1161 | 1127 |
| K00235 succinate dehydrogenase (ubiquinone) iron-sulfur subunit [EC:1.3.5.1] | 0 | 0 | 0 | 0 | 0 | 1 | 0 | 0 | 1 | 0 | 0 | 0 |
| K00239 succinate dehydrogenase / fumarate reductase, flavoprotein subunit [EC:1.3.5.1 1.3.5.4] | 5317 | 5288 | 5481 | 5592 | 6988 | 7192 | 7109 | 7166 | 5851 | 5810 | 6036 | 5806 |
| K00240 succinate dehydrogenase / fumarate reductase, iron-sulfur subunit [EC:1.3.5.1 1.3.5.4] | 1948 | 2036 | 2067 | 2049 | 2675 | 2789 | 2897 | 2818 | 2257 | 2280 | 2269 | 2155 |
| K00241 succinate dehydrogenase / fumarate reductase, cytochrome b subunit | 1286 | 1222 | 1323 | 1293 | 2195 | 2131 | 2232 | 2159 | 1616 | 1653 | 1732 | 1656 |
| K00242 succinate dehydrogenase / fumarate reductase, membrane anchor subunit | 748 | 751 | 689 | 714 | 439 | 476 | 448 | 477 | 516 | 487 | 423 | 450 |
| K00244 fumarate reductase flavoprotein subunit [EC:1.3.5.4] | 4725 | 4730 | 4595 | 4828 | 4230 | 4751 | 4466 | 4583 | 6176 | 6587 | 6494 | 6696 |
| K00245 fumarate reductase iron-sulfur subunit [EC:1.3.5.4] | 501 | 530 | 527 | 547 | 429 | 434 | 487 | 422 | 699 | 758 | 745 | 784 |
| K00246 fumarate reductase subunit C | 287 | 314 | 312 | 316 | 274 | 286 | 235 | 261 | 400 | 435 | 418 | 445 |
| K00247 fumarate reductase subunit D | 234 | 245 | 224 | 226 | 146 | 158 | 131 | 158 | 365 | 341 | 340 | 356 |
| K00658 2-oxoglutarate dehydrogenase E2 component (dihydrolipoamide succinyltransferase) [EC:2.3.1.61] | 2943 | 3053 | 3004 | 3038 | 3621 | 3672 | 4004 | 4135 | 3136 | 3004 | 3021 | 3045 |
| K01647 citrate synthase [EC:2.3.3.1] | 4779 | 4725 | 4806 | 4753 | 4915 | 5021 | 4772 | 4786 | 5561 | 5716 | 5546 | 5551 |
| K01676 fumarate hydratase, class I [EC:4.2.1.2] | 5087 | 4968 | 4925 | 5045 | 6133 | 6182 | 6121 | 5997 | 5773 | 5797 | 5742 | 5683 |
| K01677 fumarate hydratase subunit alpha [EC:4.2.1.2] | 709 | 653 | 674 | 627 | 207 | 247 | 265 | 196 | 796 | 938 | 859 | 813 |
| K01678 fumarate hydratase subunit beta [EC:4.2.1.2] | 344 | 363 | 355 | 326 | 82 | 105 | 101 | 93 | 490 | 491 | 488 | 471 |
| K01679 fumarate hydratase, class II [EC:4.2.1.2] | 5167 | 5090 | 4879 | 4879 | 3118 | 3214 | 2912 | 3048 | 2480 | 2401 | 2358 | 2434 |
| K01681 aconitate hydratase [EC:4.2.1.3] | 9263 | 9138 | 8926 | 9323 | 8552 | 8488 | 8559 | 8683 | 11053 | 11240 | 11064 | 11185 |
| K01682 aconitate hydratase 2 / 2-methylisocitrate dehydratase [EC:4.2.1.3 4.2.1.99] | 6721 | 6802 | 6535 | 6544 | 4264 | 4417 | 4153 | 4069 | 5429 | 5283 | 5132 | 5471 |
| K01899 succinyl-CoA synthetase alpha subunit [EC:6.2.1.4 6.2.1.5] | 0 | 0 | 0 | 0 | 0 | 0 | 0 | 0 | 2 | 0 | 0 | 0 |
| K01900 succinyl-CoA synthetase beta subunit [EC:6.2.1.4 6.2.1.5] | 0 | 0 | 0 | 1 | 0 | 2 | 0 | 0 | 0 | 1 | 0 | 0 |
| K01902 succinyl-CoA synthetase alpha subunit [EC:6.2.1.5] | 2076 | 2121 | 2090 | 2143 | 2536 | 2402 | 2647 | 2449 | 2211 | 2212 | 2202 | 2257 |
| K01903 succinyl-CoA synthetase beta subunit [EC:6.2.1.5] | 3039 | 3001 | 3200 | 3071 | 3574 | 3533 | 3402 | 3266 | 3089 | 2990 | 3077 | 2994 |
| K01958 pyruvate carboxylase [EC:6.4.1.1] | 1059 | 899 | 885 | 920 | 548 | 578 | 622 | 649 | 1191 | 1363 | 1166 | 1431 |
| K01959 pyruvate carboxylase subunit A [EC:6.4.1.1] | 1857 | 1848 | 1833 | 1852 | 1040 | 951 | 958 | 1083 | 624 | 636 | 567 | 627 |
| K01960 pyruvate carboxylase subunit B [EC:6.4.1.1] | 5119 | 5279 | 5293 | 5058 | 8746 | 9129 | 8849 | 8842 | 6079 | 6007 | 6104 | 5734 |
| K05942 citrate (Re)-synthase [EC:2.3.3.3] | 162 | 161 | 153 | 160 | 40 | 45 | 57 | 37 | 270 | 307 | 244 | 308 |
| K18118 succinyl-CoA:acetate CoA-transferase [EC:2.8.3.18] | 2111 | 2193 | 2082 | 2176 | 1204 | 1111 | 1070 | 1124 | 958 | 845 | 812 | 793 |
| ko00030 Pentose phosphate pathway | 112743 | 112570 | 111715 | 113078 | 99656 | 101420 | 100478 | 99284 | 111300 | 112739 | 110524 | 112052 |
| K00032 phosphogluconate 2-dehydrogenase [EC:1.1.1.43] | 1364 | 1367 | 1230 | 1302 | 742 | 709 | 659 | 718 | 385 | 295 | 329 | 377 |
| K00033 6-phosphogluconate dehydrogenase [EC:1.1.1.44 1.1.1.343] | 2399 | 2455 | 2533 | 2601 | 3729 | 3701 | 3782 | 3586 | 3841 | 4250 | 3990 | 4053 |

| | | | | | | | | | | | | |
|---|---|---|---|---|---|---|---|---|---|---|---|---|
| K00034 glucose 1-dehydrogenase [EC:1.1.1.47] | 53 | 53 | 57 | 64 | 19 | 16 | 11 | 12 | 56 | 61 | 77 | 63 |
| K00036 glucose-6-phosphate 1-dehydrogenase [EC:1.1.1.49 1.1.1.363] | 6901 | 6838 | 6884 | 6843 | 6039 | 6066 | 6001 | 6174 | 5414 | 5131 | 5143 | 5096 |
| K00090 glyoxylate/hydroxypyruvate/2-ketogluconate reductase [EC:1.1.1.79 1.1.1.81 1.1.1.215] | 1733 | 1799 | 1552 | 1736 | 945 | 1062 | 973 | 1020 | 1230 | 1236 | 1198 | 1246 |
| K00117 quinoprotein glucose dehydrogenase [EC:1.1.5.2] | 5019 | 4864 | 4718 | 4740 | 3115 | 3347 | 3146 | 3189 | 3047 | 2972 | 2886 | 3127 |
| K00615 transketolase [EC:2.2.1.1] | 13517 | 13414 | 13461 | 13442 | 9604 | 9691 | 9778 | 9303 | 14442 | 14290 | 14105 | 14499 |
| K00616 transaldolase [EC:2.2.1.2] | 2686 | 2810 | 2779 | 2906 | 2831 | 2905 | 3012 | 2917 | 3855 | 3909 | 3806 | 3842 |
| K00851 gluconokinase [EC:2.7.1.12] | 1170 | 1269 | 1332 | 1296 | 801 | 740 | 808 | 797 | 965 | 957 | 924 | 948 |
| K00852 ribokinase [EC:2.7.1.15] | 4636 | 4221 | 4473 | 4477 | 3765 | 3703 | 3574 | 3590 | 4678 | 4839 | 4681 | 4857 |
| K00874 2-dehydro-3-deoxygluconokinase [EC:2.7.1.45] | 3277 | 3126 | 3415 | 3411 | 3608 | 3636 | 3623 | 3626 | 4248 | 4381 | 4480 | 4464 |
| K00948 ribose-phosphate pyrophosphokinase [EC:2.7.6.1] | 4083 | 4114 | 4036 | 4141 | 3554 | 3612 | 3629 | 3693 | 3819 | 3769 | 3811 | 3656 |
| K01053 gluconolactonase [EC:3.1.1.17] | 465 | 424 | 429 | 401 | 251 | 335 | 295 | 320 | 137 | 141 | 138 | 131 |
| K01057 6-phosphogluconolactonase [EC:3.1.1.31] | 1528 | 1496 | 1478 | 1452 | 1993 | 1981 | 1980 | 2071 | 1380 | 1230 | 1305 | 1215 |
| K01619 deoxyribose-phosphate aldolase [EC:4.1.2.4] | 2692 | 2736 | 2571 | 2624 | 2905 | 2980 | 3027 | 2905 | 3748 | 3807 | 3818 | 3705 |
| K01621 xylulose-5-phosphate/fructose-6-phosphate phosphoketolase [EC:4.1.2.9 4.1.2.22] | 154 | 128 | 146 | 120 | 267 | 277 | 259 | 236 | 34 | 79 | 72 | 63 |
| K01625 2-dehydro-3-deoxyphosphogluconate aldolase / (4S)-4-hydroxy-2-oxoglutarate aldolase [EC:4.1.2.14 4.1.3.42] | 3200 | 3039 | 2905 | 2986 | 2555 | 2647 | 2684 | 2680 | 2467 | 2661 | 2564 | 2611 |
| K01690 phosphogluconate dehydratase [EC:4.2.1.12] | 1462 | 1511 | 1427 | 1471 | 908 | 931 | 985 | 990 | 1487 | 1581 | 1470 | 1530 |
| K01783 ribulose-phosphate 3-epimerase [EC:5.1.3.1] | 2481 | 2589 | 2754 | 2803 | 2789 | 2882 | 2859 | 2620 | 2624 | 2620 | 2596 | 2576 |
| K01807 ribose 5-phosphate isomerase A [EC:5.3.1.6] | 1841 | 1888 | 1900 | 1896 | 2265 | 2129 | 2181 | 2110 | 2164 | 2173 | 2267 | 2216 |
| K01808 ribose 5-phosphate isomerase B [EC:5.3.1.6] | 1050 | 1039 | 1018 | 987 | 1233 | 1295 | 1262 | 1241 | 1392 | 1475 | 1373 | 1455 |
| K01839 phosphopentomutase [EC:5.4.2.7] | 1312 | 1341 | 1452 | 1472 | 666 | 636 | 671 | 617 | 1827 | 2075 | 1955 | 2113 |
| K03738 aldehyde:ferredoxin oxidoreductase [EC:1.2.7.5] | 212 | 255 | 244 | 232 | 154 | 150 | 151 | 119 | 574 | 691 | 564 | 608 |
| K05308 gluconate/galactonate dehydratase [EC:4.2.1.140] | 14 | 12 | 23 | 17 | 4 | 5 | 3 | 4 | 33 | 25 | 30 | 28 |
| K05774 ribose 1,5-bisphosphokinase [EC:2.7.4.23] | 1071 | 1081 | 986 | 960 | 533 | 515 | 468 | 569 | 685 | 740 | 723 | 796 |
| K06151 gluconate 2-dehydrogenase alpha chain [EC:1.1.99.3] | 2832 | 2866 | 2492 | 2813 | 1274 | 1403 | 1317 | 1487 | 1621 | 1595 | 1532 | 1532 |
| K06152 gluconate 2-dehydrogenase gamma chain [EC:1.1.99.3] | 1222 | 1216 | 1129 | 1190 | 596 | 559 | 491 | 570 | 890 | 923 | 859 | 917 |
| K07404 6-phosphogluconolactonase [EC:3.1.1.31] | 1754 | 1878 | 1811 | 1850 | 624 | 756 | 633 | 563 | 1879 | 2100 | 1948 | 1970 |
| K08093 3-hexulose-6-phosphate synthase [EC:4.1.2.43] | 3 | 4 | 3 | 2 | 2 | 1 | 0 | 2 | 13 | 16 | 13 | 12 |
| K08094 6-phospho-3-hexuloisomerase [EC:5.3.1.27] | 29 | 35 | 19 | 25 | 11 | 5 | 9 | 1 | 70 | 64 | 70 | 68 |
| K11441 dehydrogluconokinase [EC:2.7.1.13] | 1184 | 1196 | 1076 | 1135 | 532 | 575 | 550 | 631 | 344 | 304 | 298 | 361 |
| K11529 glycerate 2-kinase [EC:2.7.1.165] | 2766 | 2585 | 2489 | 2539 | 1699 | 1718 | 1667 | 1694 | 908 | 854 | 858 | 882 |
| K13812 bifunctional enzyme Fae/Hps [EC:4.2.1.147 4.1.2.43] | 2 | 0 | 0 | 0 | 0 | 0 | 2 | 2 | 6 | 4 | 2 | 8 |
| K13831 3-hexulose-6-phosphate synthase / 6-phospho-3-hexuloisomerase [EC:4.1.2.43 5.3.1.27] | 1 | 1 | 1 | 0 | 0 | 0 | 1 | 0 | 9 | 1 | 3 | 3 |
| K17463 2-dehydro-3-deoxy-phosphogluconate aldolase [EC:4.1.2.14] | 0 | 0 | 2 | 0 | 0 | 0 | 3 | 0 | 0 | 0 | 2 | 7 |
| K17464 PTS system, D-glucosaminate-specific IIA component [EC:2.7.1.203] | 28 | 32 | 26 | 21 | 8 | 12 | 4 | 2 | 45 | 74 | 49 | 51 |
| K17465 PTS system, D-glucosaminate-specific IIB component [EC:2.7.1.203] | 20 | 31 | 30 | 34 | 1 | 14 | 9 | 14 | 64 | 77 | 70 | 68 |
| K17466 PTS system, D-glucosaminate-specific IIC component | 76 | 63 | 53 | 59 | 18 | 12 | 5 | 18 | 110 | 84 | 88 | 90 |
| K17467 PTS system, D-glucosaminate-specific IID component | 56 | 71 | 67 | 64 | 8 | 10 | 9 | 16 | 98 | 106 | 112 | 111 |
| K17468 D-glucosaminate-6-phosphate ammonia-lyase [EC:4.3.1.29] | 0 | 0 | 2 | 0 | 2 | 0 | 2 | 0 | 1 | 0 | 1 | 2 |
| K18022 glyceraldehyde dehydrogenase small subunit [EC:1.2.99.8] | 0 | 0 | 0 | 0 | 1 | 0 | 0 | 0 | 0 | 0 | 0 | 0 |
| K19813 glucose dehydrogenase [EC:1.1.5.9] | 2 | 2 | 0 | 1 | 0 | 0 | 0 | 0 | 0 | 0 | 0 | 0 |
| K22229 2-ketogluconate reductase [EC:1.1.1.215] | 4 | 2 | 3 | 3 | 3 | 1 | 4 | 3 | 0 | 0 | 0 | 0 |
| K22345 glucosaminate ammonia-lyase [EC:4.3.1.9] | 1426 | 1463 | 1385 | 1436 | 801 | 818 | 771 | 872 | 571 | 463 | 437 | 450 |
| ko00040 Pentose and glucuronate interconversions | 67575 | 67509 | 68186 | 69040 | 87252 | 88003 | 88499 | 86556 | 94585 | 95082 | 95426 | 95212 |
| K00007 D-arabinitol 4-dehydrogenase [EC:1.1.1.11] | 720 | 748 | 707 | 732 | 525 | 557 | 506 | 561 | 863 | 963 | 910 | 905 |
| K00008 L-iditol 2-dehydrogenase [EC:1.1.1.14] | 1003 | 1060 | 1121 | 1055 | 1601 | 1652 | 1652 | 1568 | 2118 | 2078 | 2171 | 2130 |
| K00011 aldehyde reductase [EC:1.1.1.21] | 0 | 1 | 1 | 1 | 2 | 0 | 0 | 0 | 1 | 0 | 0 | 1 |
| K00012 UDPglucose 6-dehydrogenase [EC:1.1.1.22] | 4860 | 4927 | 4712 | 4734 | 4380 | 4256 | 4495 | 4275 | 4567 | 4348 | 4457 | 4574 |
| K00039 ribitol 2-dehydrogenase [EC:1.1.1.56] | 299 | 269 | 291 | 290 | 90 | 85 | 81 | 68 | 626 | 583 | 502 | 623 |
| K00040 fructuronate reductase [EC:1.1.1.57] | 2486 | 2472 | 2378 | 2464 | 1347 | 1297 | 1290 | 1280 | 3215 | 3389 | 3118 | 3429 |
| K00041 tagaturonate reductase [EC:1.1.1.58] | 2083 | 2071 | 2201 | 2086 | 3393 | 3504 | 3370 | 3429 | 3525 | 3390 | 3570 | 3383 |
| K00065 2-dehydro-3-deoxy-D-gluconate 5-dehydrogenase [EC:1.1.1.127] | 1115 | 1035 | 1014 | 1182 | 628 | 690 | 676 | 657 | 1474 | 1531 | 1418 | 1577 |
| K00078 dihydrodiol dehydrogenase / D-xylose 1-dehydrogenase (NADP) [EC:1.3.1.20 1.1.1.179] | 3 | 5 | 2 | 3 | 0 | 0 | 0 | 0 | 6 | 4 | 8 | 7 |
| K00848 rhamnulokinase [EC:2.7.1.5] | 3459 | 3558 | 3642 | 3705 | 6682 | 6534 | 6926 | 6491 | 6511 | 6449 | 6858 | 6456 |
| K00853 L-ribulokinase [EC:2.7.1.16] | 1299 | 1247 | 1142 | 1256 | 837 | 875 | 798 | 826 | 1743 | 1891 | 1807 | 1757 |
| K00854 xylulokinase [EC:2.7.1.17] | 4970 | 4712 | 4793 | 4953 | 5452 | 5612 | 5553 | 5484 | 6650 | 6832 | 6708 | 6951 |
| K00875 D-ribulokinase [EC:2.7.1.47] | 580 | 677 | 551 | 582 | 190 | 194 | 150 | 159 | 1142 | 1331 | 1145 | 1285 |
| K00880 L-xylulokinase [EC:2.7.1.53] | 823 | 804 | 818 | 881 | 776 | 776 | 759 | 779 | 1397 | 1476 | 1339 | 1455 |
| K00963 UTP--glucose-1-phosphate uridylyltransferase [EC:2.7.7.9] | 1849 | 1761 | 1751 | 1800 | 1380 | 1480 | 1322 | 1405 | 1673 | 1922 | 1817 | 1890 |
| K01051 pectinesterase [EC:3.1.1.11] | 3926 | 4011 | 4473 | 4185 | 11896 | 11951 | 11827 | 11802 | 9605 | 8792 | 9556 | 9076 |
| K01184 polygalacturonase [EC:3.2.1.15] | 289 | 304 | 279 | 275 | 409 | 449 | 442 | 414 | 0 | 0 | 0 | 1 |
| K01195 beta-glucuronidase [EC:3.2.1.31] | 2965 | 2979 | 3153 | 3092 | 7622 | 7568 | 7800 | 7477 | 7097 | 6746 | 7115 | 6902 |
| K01629 rhamnulose-1-phosphate aldolase [EC:4.1.2.19] | 1607 | 1667 | 1649 | 1640 | 2389 | 2398 | 2490 | 2204 | 2563 | 2676 | 2734 | 2633 |
| K01685 altronate hydrolase [EC:4.2.1.7] | 2431 | 2566 | 2555 | 2492 | 3502 | 3676 | 3728 | 3688 | 3753 | 3776 | 3898 | 4004 |
| K01686 mannonate dehydratase [EC:4.2.1.8] | 3849 | 3979 | 4004 | 3948 | 4450 | 4710 | 4738 | 4386 | 5266 | 5541 | 5429 | 5326 |
| K01728 pectate lyase [EC:4.2.2.2] | 3 | 0 | 0 | 1 | 0 | 0 | 5 | 0 | 2 | 0 | 2 | 4 |
| K01730 oligogalacturonide lyase [EC:4.2.2.6] | 631 | 689 | 615 | 699 | 365 | 326 | 336 | 314 | 1045 | 1122 | 1034 | 1116 |
| K01804 L-arabinose isomerase [EC:5.3.1.4] | 2549 | 2511 | 2697 | 2657 | 3997 | 3737 | 4147 | 3917 | 4192 | 4436 | 4335 | 4125 |
| K01805 xylose isomerase [EC:5.3.1.5] | 2294 | 2252 | 2320 | 2376 | 3355 | 3399 | 3445 | 3341 | 3967 | 4059 | 4083 | 4057 |
| K01812 glucuronate isomerase [EC:5.3.1.12] | 2614 | 2561 | 2772 | 2718 | 3536 | 3561 | 3659 | 3543 | 3609 | 3502 | 3594 | 3642 |
| K01815 4-deoxy-L-threo-5-hexosulose-uronate ketol-isomerase [EC:5.3.1.17] | 1960 | 1995 | 2170 | 2032 | 4663 | 4455 | 4441 | 4525 | 3514 | 3740 | 3703 | 3442 |
| K01820 L-rhamnose isomerase / sugar isomerase [EC:5.3.1.14 5.3.1.-] | 0 | 2 | 2 | 2 | 3 | 2 | 0 | 0 | 0 | 5 | 3 | 1 |
| K03077 L-ribulose-5-phosphate 4-epimerase [EC:5.1.3.4] | 2082 | 2170 | 2151 | 2159 | 2252 | 2448 | 2505 | 2392 | 2848 | 2789 | 2768 | 2669 |
| K03078 3-dehydro-L-gulonate-6-phosphate decarboxylase [EC:4.1.1.85] | 667 | 600 | 631 | 634 | 551 | 568 | 551 | 578 | 697 | 633 | 632 | 718 |
| K03079 L-ribulose-5-phosphate 3-epimerase [EC:5.1.3.22] | 1003 | 944 | 889 | 996 | 678 | 845 | 773 | 719 | 973 | 1027 | 1059 | 1008 |
| K03331 L-xylulose reductase [EC:1.1.1.10] | 0 | 0 | 0 | 1 | 0 | 0 | 0 | 0 | 0 | 0 | 0 | 1 |
| K05351 D-xylulose reductase [EC:1.1.1.9] | 4 | 1 | 2 | 5 | 6 | 1 | 3 | 3 | 2 | 10 | 5 | 6 |
| K05352 ribitol-5-phosphate 2-dehydrogenase (NADP+) [EC:1.1.1.405] | 1 | 5 | 0 | 2 | 0 | 1 | 2 | 7 | 4 | 11 | 6 | 9 |
| K08092 3-dehydro-L-gulonate 2-dehydrogenase [EC:1.1.1.130] | 292 | 283 | 295 | 327 | 451 | 465 | 435 | 407 | 114 | 162 | 122 | 132 |
| K08322 L-gulonate 5-dehydrogenase [EC:1.1.1.380] | 691 | 720 | 614 | 708 | 379 | 374 | 414 | 420 | 1084 | 1083 | 1004 | 1101 |
| K08323 mannonate dehydratase [EC:4.2.1.8] | 762 | 763 | 769 | 788 | 754 | 820 | 784 | 726 | 675 | 660 | 663 | 673 |
| K09988 D-lyxose ketol-isomerase [EC:5.3.1.15] | 514 | 517 | 477 | 597 | 469 | 522 | 523 | 499 | 618 | 720 | 608 | 718 |
| K13877 2,5-dioxopentanoate dehydrogenase [EC:1.2.1.26] | 4060 | 3708 | 3521 | 3761 | 2044 | 1876 | 1757 | 2170 | 1255 | 1019 | 1003 | 1098 |
| K14274 xylonolactonase [EC:3.1.1.-] | 565 | 558 | 598 | 604 | 372 | 363 | 342 | 419 | 739 | 781 | 786 | 817 |
| K16849 altronate dehydratase small subunit [EC:4.2.1.7] | 73 | 66 | 68 | 71 | 35 | 43 | 55 | 38 | 119 | 114 | 106 | 115 |
| K16850 altronate dehydratase large subunit [EC:4.2.1.7] | 649 | 677 | 635 | 681 | 321 | 321 | 308 | 286 | 985 | 1029 | 909 | 1014 |
| K17743 D-xylose reductase [EC:1.1.1.307] | 0 | 0 | 1 | 1 | 3 | 1 | 3 | 0 | 0 | 1 | 2 | 0 |
| K17818 D-arabinitol dehydrogenase (NADP+) [EC:1.1.1.287] | 2 | 1 | 2 | 3 | 2 | 2 | 3 | 0 | 4 | 2 | 3 | 5 |
| K18106 D-galacturonate reductase [EC:1.1.1.-] | 0 | 0 | 0 | 0 | 0 | 0 | 0 | 0 | 0 | 0 | 0 | 1 |
| K19588 2,5-dioxopentanoate dehydrogenase [EC:1.2.1.26] | 1 | 2 | 1 | 1 | 7 | 3 | 2 | 3 | 4 | 0 | 2 | 1 |
| K21030 D-ribitol-5-phosphate cytidylyltransferase [EC:2.7.7.40] | 4 | 2 | 2 | 1 | 0 | 0 | 1 | 0 | 1 | 12 | 3 | 18 |
| K21031 D-ribitol-5-phosphate cytidylyltransferase [EC:2.7.7.40] | 0 | 0 | 0 | 0 | 0 | 0 | 0 | 0 | 0 | 0 | 0 | 1 |
| K21619 tagaturonate epimerase [EC:5.1.2.7] | 39 | 40 | 38 | 26 | 2 | 0 | 0 | 0 | 97 | 85 | 76 | 82 |
| K21681 ribitol-5-phosphate 2-dehydrogenase (NADP+) / D-ribitol-5-phosphate cytidylyltransferase [EC:1.1.1.405 2.7.7.40] | 152 | 160 | 196 | 144 | 246 | 292 | 236 | 272 | 24 | 22 | 24 | 36 |
| K22185 D-xylose 1-dehydrogenase [EC:1.1.1.175] | 153 | 173 | 166 | 163 | 230 | 227 | 210 | 213 | 0 | 3 | 0 | 0 |
| K22186 xylonate dehydratase [EC:4.2.1.82] | 3 | 0 | 1 | 0 | 2 | 0 | 0 | 1 | 0 | 2 | 0 | 1 |
| K22187 alpha-ketoglutaric semialdehyde dehydrogenase [EC:1.2.1.-] | 3 | 0 | 3 | 1 | 5 | 0 | 2 | 2 | 2 | 4 | 2 | 7 |
| K22396 xylonate dehydratase [EC:4.2.1.82] | 873 | 919 | 772 | 881 | 1390 | 1450 | 1302 | 1371 | 2 | 6 | 11 | 9 |
| K22397 2-dehydro-3-deoxy-D-pentonate aldolase [EC:4.1.2.28] | 785 | 768 | 823 | 849 | 636 | 630 | 628 | 668 | 404 | 415 | 402 | 381 |
| K22933 galacturan 1,4-alpha-galacturonidase [EC:3.2.1.67] | 809 | 758 | 769 | 788 | 89 | 65 | 81 | 62 | 679 | 723 | 751 | 712 |
| K22935 D-ribulokinase [EC:2.7.1.47] | 55 | 48 | 35 | 47 | 5 | 1 | 0 | 3 | 85 | 118 | 111 | 98 |
| K22994 pectate disaccharide-lyase [EC:4.2.2.9] | 11 | 5 | 11 | 9 | 21 | 22 | 32 | 19 | 116 | 141 | 122 | 118 |
| ko00051 Fructose and mannose metabolism | 99507 | 100904 | 100066 | 100728 | 112467 | 113728 | 113153 | 110641 | 116005 | 117468 | 116532 | 116869 |
| K00009 mannitol-1-phosphate 5-dehydrogenase [EC:1.1.1.17] | 798 | 742 | 791 | 706 | 575 | 597 | 613 | 514 | 990 | 1087 | 996 | 1123 |
| K00045 mannitol 2-dehydrogenase [EC:1.1.1.67] | 2522 | 2627 | 2248 | 2329 | 3277 | 3234 | 3157 | 3215 | 2298 | 2077 | 2273 | 2148 |
| K00064 D-threo-aldose 1-dehydrogenase [EC:1.1.1.122] | 0 | 0 | 0 | 1 | 0 | 0 | 3 | 2 | 3 | 8 | 6 | 4 |
| K00066 GDP-mannose 6-dehydrogenase [EC:1.1.1.132] | 1877 | 1974 | 1877 | 1895 | 1191 | 1266 | 1246 | 1235 | 677 | 665 | 558 | 615 |
| K00068 sorbitol-6-phosphate 2-dehydrogenase [EC:1.1.1.140] | 736 | 823 | 820 | 770 | 547 | 688 | 608 | 582 | 753 | 750 | 794 | 795 |
| K00847 fructokinase [EC:2.7.1.4] | 5390 | 5227 | 5576 | 5413 | 7448 | 7276 | 7231 | 7228 | 6704 | 6628 | 7026 | 6951 |
| K00863 triose/dihydroxyacetone kinase / FAD-AMP lyase (cyclizing) [EC:2.7.1.28 2.7.1.29 4.6.1.15] | 1025 | 1087 | 1046 | 1046 | 693 | 734 | 712 | 666 | 1413 | 1313 | 1298 | 1436 |
| K00879 L-fuculokinase [EC:2.7.1.51] | 606 | 653 | 525 | 619 | 168 | 241 | 195 | 179 | 1253 | 1237 | 1196 | 1307 |
| K00881 allose kinase [EC:2.7.1.55] | 60 | 40 | 43 | 39 | 9 | 8 | 4 | 10 | 139 | 140 | 180 | 182 |
| K00882 1-phosphofructokinase [EC:2.7.1.56] | 1167 | 1077 | 1030 | 1092 | 545 | 532 | 565 | 511 | 1224 | 1285 | 1223 | 1436 |
| K00900 6-phosphofructo-2-kinase [EC:2.7.1.105] | 0 | 0 | 0 | 0 | 0 | 0 | 0 | 0 | 0 | 0 | 0 | 1 |
| K00966 mannose-1-phosphate guanylyltransferase [EC:2.7.7.13] | 14 | 19 | 16 | 21 | 0 | 0 | 0 | 0 | 24 | 38 | 28 | 42 |
| K00971 mannose-1-phosphate guanylyltransferase [EC:2.7.7.13] | 2438 | 2518 | 2612 | 2591 | 5333 | 5223 | 5491 | 5253 | 4634 | 4505 | 4610 | 4544 |
| K01218 mannan endo-1,4-beta-mannosidase [EC:3.2.1.78] | 489 | 505 | 496 | 509 | 185 | 210 | 185 | 232 | 326 | 345 | 360 | 355 |
| K01628 L-fuculose-phosphate aldolase [EC:4.1.2.17] | 2122 | 2068 | 2136 | 2074 | 1043 | 1137 | 1077 | 1150 | 1606 | 1665 | 1552 | 1667 |
| K01711 GDPmannose 4,6-dehydratase [EC:4.2.1.47] | 2114 | 2221 | 2320 | 2252 | 4540 | 4418 | 4456 | 4333 | 3096 | 3220 | 3169 | 3159 |
| K01729 poly(beta-D-mannuronate) lyase [EC:4.2.2.3] | 515 | 510 | 490 | 467 | 414 | 430 | 416 | 453 | 107 | 79 | 87 | 91 |
| K01795 mannuronan 5-epimerase [EC:5.1.3.37] | 1317 | 1300 | 1245 | 1245 | 660 | 631 | 605 | 663 | 406 | 376 | 352 | 417 |
| K01809 mannose-6-phosphate isomerase [EC:5.3.1.8] | 1973 | 1891 | 1983 | 1891 | 2930 | 3000 | 2881 | 2686 | 2662 | 2548 | 2657 | 2641 |

| | | | | | | | | | | | | |
|---|---|---|---|---|---|---|---|---|---|---|---|---|
| K01813 L-rhamnose isomerase [EC:5.3.1.14] | 2188 | 2321 | 2379 | 2329 | 3516 | 3678 | 3678 | 3581 | 3820 | 3984 | 3793 | 3794 |
| K01818 L-fucose/D-arabinose isomerase [EC:5.3.1.25 5.3.1.3] | 2097 | 2196 | 2258 | 2223 | 4023 | 3915 | 3965 | 3908 | 4625 | 4663 | 4665 | 4529 |
| K01840 phosphomannomutase [EC:5.4.2.8] | 1620 | 1746 | 1779 | 1789 | 3556 | 3505 | 3505 | 3544 | 3582 | 3481 | 3538 | 3321 |
| K02377 GDP-L-fucose synthase [EC:1.1.1.271] | 1022 | 1100 | 1029 | 1149 | 2584 | 2716 | 2653 | 2603 | 1655 | 1680 | 1508 | 1550 |
| K02768 PTS system, fructose-specific IIA component [EC:2.7.1.202] | 418 | 440 | 417 | 468 | 278 | 270 | 224 | 233 | 477 | 528 | 475 | 496 |
| K02769 PTS system, fructose-specific IIB component [EC:2.7.1.202] | 206 | 199 | 182 | 187 | 131 | 167 | 151 | 171 | 266 | 267 | 288 | 321 |
| K02770 PTS system, fructose-specific IIC component | 3676 | 3765 | 3503 | 3644 | 2009 | 1940 | 1789 | 1935 | 3411 | 3654 | 3318 | 3616 |
| K02771 PTS system, fructose-specific IID component | 0 | 0 | 0 | 0 | 0 | 0 | 0 | 0 | 0 | 0 | 0 | 1 |
| K02781 PTS system, glucitol/sorbitol-specific IIA component [EC:2.7.1.198] | 475 | 445 | 456 | 580 | 375 | 418 | 327 | 330 | 658 | 651 | 630 | 620 |
| K02783 PTS system, glucitol/sorbitol-specific IIC component | 956 | 938 | 905 | 941 | 736 | 721 | 712 | 640 | 1332 | 1359 | 1278 | 1367 |
| K02793 PTS system, mannose-specific IIA component [EC:2.7.1.191] | 188 | 157 | 147 | 152 | 31 | 26 | 61 | 17 | 105 | 127 | 138 | 114 |
| K02794 PTS system, mannose-specific IIB component [EC:2.7.1.191] | 967 | 997 | 871 | 958 | 417 | 470 | 400 | 418 | 1011 | 1114 | 1051 | 1071 |
| K02795 PTS system, mannose-specific IIC component | 1385 | 1343 | 1357 | 1420 | 488 | 505 | 471 | 487 | 1183 | 1299 | 1239 | 1259 |
| K02796 PTS system, mannose-specific IID component | 1417 | 1411 | 1444 | 1429 | 532 | 516 | 610 | 507 | 1261 | 1471 | 1362 | 1372 |
| K02798 PTS system, mannitol-specific IIA component [EC:2.7.1.197] | 65 | 59 | 51 | 64 | 24 | 23 | 27 | 16 | 102 | 111 | 93 | 90 |
| K02800 PTS system, mannitol-specific IIC component | 1552 | 1523 | 1422 | 1553 | 991 | 1062 | 993 | 1032 | 1889 | 2113 | 1955 | 2045 |
| K02812 PTS system, sorbose-specific IIA component [EC:2.7.1.206] | 87 | 79 | 93 | 74 | 66 | 70 | 73 | 58 | 25 | 36 | 31 | 26 |
| K02813 PTS system, sorbose-specific IIB component [EC:2.7.1.206] | 149 | 180 | 195 | 182 | 159 | 157 | 156 | 171 | 33 | 43 | 28 | 44 |
| K02814 PTS system, sorbose-specific IIC component | 190 | 195 | 245 | 203 | 251 | 265 | 268 | 269 | 45 | 67 | 62 | 67 |
| K02815 PTS system, sorbose-specific IID component | 267 | 287 | 269 | 286 | 292 | 296 | 294 | 292 | 45 | 66 | 45 | 54 |
| K03332 fructan beta-fructosidase [EC:3.2.1.80] | 1635 | 1913 | 1849 | 1781 | 5225 | 5214 | 5283 | 4912 | 4049 | 3933 | 4033 | 3799 |
| K05305 fucokinase [EC:2.7.1.52] | 104 | 100 | 121 | 106 | 37 | 69 | 92 | 49 | 204 | 203 | 182 | 229 |
| K07026 mannosyl-3-phosphoglycerate phosphatase [EC:3.1.3.70] | 160 | 158 | 163 | 173 | 178 | 299 | 257 | 251 | 26 | 46 | 19 | 34 |
| K07046 L-fuconolactonase [EC:3.1.1.-] | 9 | 2 | 4 | 7 | 0 | 2 | 0 | 1 | 0 | 0 | 2 | 0 |
| K08261 D-sorbitol dehydrogenase (acceptor) [EC:1.1.99.21] | 1 | 0 | 1 | 0 | 3 | 0 | 0 | 0 | 0 | 0 | 0 | 0 |
| K12660 2-dehydro-3-deoxy-L-rhamnonate aldolase [EC:4.1.2.53] | 530 | 518 | 454 | 518 | 335 | 313 | 322 | 342 | 638 | 605 | 622 | 668 |
| K12661 L-rhamnonate dehydratase [EC:4.2.1.90] | 699 | 741 | 709 | 746 | 482 | 581 | 485 | 465 | 1029 | 1064 | 1002 | 1100 |
| K15856 GDP-4-dehydro-6-deoxy-D-mannose reductase [EC:1.1.1.281] | 1859 | 1744 | 1660 | 1739 | 975 | 916 | 867 | 891 | 626 | 524 | 603 | 669 |
| K16011 mannose-1-phosphate guanylyltransferase / mannose-6-phosphate isomerase [EC:2.7.7.13 5.3.1.8] | 6543 | 6460 | 6085 | 6326 | 3518 | 3457 | 3448 | 3537 | 2144 | 2039 | 1943 | 1989 |
| K16881 mannose-1-phosphate guanylyltransferase / phosphomannomutase [EC:2.7.7.13 5.4.2.8] | 11 | 18 | 11 | 9 | 2 | 0 | 2 | 0 | 10 | 2 | 3 | 4 |
| K17195 D-allulose-6-phosphate 3-epimerase [EC:5.1.3.-] | 338 | 340 | 326 | 366 | 76 | 78 | 105 | 70 | 600 | 638 | 562 | 641 |
| K17742 sorbose reductase [EC:1.1.1.289] | 1 | 0 | 0 | 0 | 0 | 0 | 0 | 0 | 1 | 0 | 0 | 0 |
| K18333 L-fucose dehydrogenase | 20 | 9 | 11 | 13 | 3 | 2 | 0 | 4 | 13 | 11 | 5 | 7 |
| K18334 L-fuconate dehydratase [EC:4.2.1.68] | 3 | 2 | 3 | 2 | 1 | 0 | 7 | 6 | 3 | 10 | 15 | 4 |
| K18335 2-keto-3-deoxy-L-fuconate dehydrogenase [EC:1.1.1.-] | 3 | 5 | 0 | 1 | 0 | 5 | 3 | 0 | 2 | 1 | 1 | 3 |
| K18337 L-rhamnose 1-dehydrogenase [EC:1.1.1.378 1.1.1.377 1.1.1.173] | 2 | 7 | 1 | 1 | 0 | 0 | 0 | 0 | 0 | 1 | 0 | 0 |
| K19290 mannuronan synthase [EC:2.4.1.33] | 2230 | 2184 | 2287 | 2302 | 1302 | 1216 | 1162 | 1247 | 729 | 695 | 708 | 761 |
| K19291 mannuronan synthase [EC:2.4.1.33] | 1684 | 1554 | 1525 | 1517 | 929 | 827 | 777 | 888 | 507 | 556 | 458 | 511 |
| K19355 mannan endo-1,4-beta-mannosidase [EC:3.2.1.78] | 80 | 69 | 69 | 53 | 200 | 180 | 188 | 194 | 13 | 37 | 45 | 37 |
| K19956 L-sorbose 1-phosphate reductase [EC:1.1.1.-] | 434 | 405 | 465 | 418 | 456 | 447 | 443 | 435 | 202 | 202 | 190 | 197 |
| K21883 2-dehydro-3-deoxy-L-rhamnonate dehydrogenase (NAD+) [EC:1.1.1.401] | 1 | 2 | 3 | 5 | 0 | 0 | 0 | 0 | 1 | 4 | 2 | 1 |
| ko00052 Galactose metabolism | 98623 | 101417 | 103752 | 102272 | 155223 | 156456 | 157136 | 153939 | 154648 | 155519 | 155464 | 154988 |
| K00035 D-galactose 1-dehydrogenase [EC:1.1.1.48] | 1 | 0 | 0 | 0 | 1 | 2 | 0 | 2 | 2 | 0 | 0 | 2 |
| K00094 galactitol-1-phosphate 5-dehydrogenase [EC:1.1.1.251] | 5 | 3 | 3 | 2 | 10 | 13 | 10 | 6 | 47 | 55 | 38 | 43 |
| K00849 galactokinase [EC:2.7.1.6] | 2393 | 2603 | 2469 | 2460 | 3289 | 3311 | 3448 | 3314 | 3443 | 3599 | 3314 | 3607 |
| K00883 2-dehydro-3-deoxygalactonokinase [EC:2.7.1.58] | 995 | 924 | 945 | 984 | 527 | 566 | 582 | 540 | 1590 | 1511 | 1424 | 1533 |
| K00917 tagatose 6-phosphate kinase [EC:2.7.1.144] | 119 | 130 | 154 | 136 | 60 | 57 | 45 | 53 | 212 | 271 | 218 | 255 |
| K00965 UDPglucose--hexose-1-phosphate uridylyltransferase [EC:2.7.7.12] | 1321 | 1413 | 1373 | 1488 | 1030 | 1083 | 983 | 977 | 1529 | 1441 | 1494 | 1511 |
| K01182 oligo-1,6-glucosidase [EC:3.2.1.10] | 1373 | 1325 | 1385 | 1366 | 323 | 374 | 351 | 332 | 1526 | 1619 | 1547 | 1621 |
| K01187 alpha-glucosidase [EC:3.2.1.20] | 9858 | 10342 | 11056 | 10380 | 22900 | 23140 | 23139 | 22711 | 19666 | 19818 | 20303 | 19484 |
| K01190 beta-galactosidase [EC:3.2.1.23] | 22473 | 23862 | 24933 | 23509 | 54266 | 54176 | 54438 | 53231 | 40247 | 38418 | 39959 | 38794 |
| K01193 beta-fructofuranosidase [EC:3.2.1.26] | 4207 | 4334 | 4229 | 4392 | 2703 | 2867 | 2703 | 2603 | 5125 | 5236 | 5272 | 5441 |
| K01220 6-phospho-beta-galactosidase [EC:3.2.1.85] | 82 | 76 | 79 | 98 | 30 | 66 | 34 | 56 | 127 | 151 | 148 | 138 |
| K01229 lactase-phlorizin hydrolase [EC:3.2.1.108 3.2.1.62] | 1 | 0 | 0 | 0 | 0 | 0 | 0 | 0 | 0 | 0 | 0 | 1 |
| K01631 2-dehydro-3-deoxyphosphogalactonate aldolase [EC:4.1.2.21] | 593 | 623 | 600 | 749 | 336 | 346 | 340 | 374 | 1104 | 1112 | 1031 | 1021 |
| K01635 tagatose 1,6-diphosphate aldolase [EC:4.1.2.40] | 154 | 158 | 174 | 170 | 46 | 77 | 55 | 66 | 176 | 239 | 218 | 219 |
| K01684 galactonate dehydratase [EC:4.2.1.6] | 1720 | 1614 | 1721 | 1715 | 567 | 596 | 626 | 642 | 1664 | 1806 | 1670 | 1831 |
| K01784 UDP-glucose 4-epimerase [EC:5.1.3.2] | 4262 | 4362 | 4301 | 4297 | 4811 | 4755 | 4699 | 4702 | 5126 | 5274 | 5242 | 5493 |
| K01819 galactose-6-phosphate isomerase [EC:5.3.1.26] | 108 | 107 | 87 | 109 | 35 | 54 | 50 | 42 | 135 | 165 | 143 | 119 |
| K01854 UDP-galactopyranose mutase [EC:5.4.99.9] | 1271 | 1340 | 1336 | 1376 | 1278 | 1176 | 1317 | 1308 | 2606 | 2722 | 2528 | 2666 |
| K02079 N-acetylgalactosamine-6-phosphate deacetylase [EC:3.5.1.25] | 13 | 2 | 2 | 5 | 14 | 19 | 18 | 11 | 67 | 91 | 55 | 62 |
| K02080 galactosamine-6-phosphate isomerase [EC:5.3.1.-] | 185 | 143 | 150 | 173 | 256 | 282 | 252 | 227 | 72 | 67 | 71 | 58 |
| K02744 PTS system, N-acetylgalactosamine-specific IIA component [EC:2.7.1.-] | 326 | 308 | 394 | 383 | 197 | 217 | 199 | 206 | 167 | 205 | 184 | 219 |
| K02745 PTS system, N-acetylgalactosamine-specific IIB component [EC:2.7.1.-] | 41 | 55 | 48 | 61 | 13 | 15 | 16 | 9 | 90 | 53 | 78 | 60 |
| K02746 PTS system, N-acetylgalactosamine-specific IIC component | 83 | 91 | 90 | 84 | 34 | 34 | 24 | 29 | 135 | 167 | 136 | 108 |
| K02747 PTS system, N-acetylgalactosamine-specific IID component | 121 | 105 | 119 | 152 | 21 | 35 | 27 | 36 | 115 | 206 | 155 | 189 |
| K02773 PTS system, galactitol-specific IIA component [EC:2.7.1.200] | 67 | 68 | 86 | 57 | 27 | 30 | 29 | 26 | 95 | 132 | 122 | 148 |
| K02774 PTS system, galactitol-specific IIB component [EC:2.7.1.200] | 69 | 46 | 40 | 49 | 24 | 16 | 11 | 7 | 101 | 120 | 77 | 95 |
| K02775 PTS system, galactitol-specific IIC component | 1338 | 1472 | 1557 | 1487 | 3198 | 3233 | 3307 | 3170 | 3232 | 3324 | 3385 | 3450 |
| K02786 PTS system, lactose-specific IIA component [EC:2.7.1.207] | 24 | 27 | 23 | 21 | 8 | 20 | 6 | 18 | 39 | 48 | 25 | 28 |
| K02788 PTS system, lactose-specific IIC component | 119 | 104 | 138 | 96 | 73 | 79 | 69 | 45 | 138 | 179 | 169 | 139 |
| K06617 raffinose synthase [EC:2.4.1.82] | 0 | 0 | 1 | 0 | 0 | 2 | 0 | 0 | 0 | 0 | 0 | 0 |
| K07406 alpha-galactosidase [EC:3.2.1.22] | 2047 | 2029 | 1906 | 1946 | 490 | 530 | 489 | 431 | 3502 | 3637 | 3365 | 3697 |
| K07407 alpha-galactosidase [EC:3.2.1.22] | 8234 | 8579 | 9005 | 8696 | 14632 | 14741 | 14912 | 14439 | 15668 | 15810 | 15673 | 15470 |
| K08302 tagatose 1,6-diphosphate aldolase GatY/KbaY [EC:4.1.2.40] | 213 | 228 | 215 | 240 | 341 | 377 | 342 | 304 | 193 | 197 | 225 | 209 |
| K10984 PTS system, galactosamine-specific IIB component [EC:2.7.1.-] | 108 | 102 | 95 | 116 | 219 | 204 | 153 | 141 | 18 | 21 | 15 | 14 |
| K10985 PTS system, galactosamine-specific IIC component | 149 | 164 | 159 | 188 | 261 | 273 | 260 | 258 | 30 | 39 | 46 | 42 |
| K10986 PTS system, galactosamine-specific IID component | 210 | 160 | 162 | 192 | 269 | 301 | 268 | 279 | 64 | 57 | 49 | 47 |
| K12111 evolved beta-galactosidase subunit alpha [EC:3.2.1.23] | 582 | 521 | 562 | 576 | 131 | 102 | 102 | 102 | 772 | 834 | 781 | 836 |
| K12112 evolved beta-galactosidase subunit beta | 3 | 2 | 0 | 0 | 2 | 8 | 2 | 6 | 30 | 32 | 19 | 30 |
| K12308 beta-galactosidase [EC:3.2.1.23] | 7522 | 7550 | 7621 | 7746 | 8363 | 8310 | 8731 | 8624 | 12368 | 12998 | 12754 | 12824 |
| K12309 beta-galactosidase [EC:3.2.1.23] | 4 | 5 | 1 | 4 | 9 | 5 | 7 | 9 | 4 | 10 | 4 | 5 |
| K16371 D-tagatose-1,6-bisphosphate aldolase subunit GatZ/KbaZ | 272 | 291 | 266 | 251 | 551 | 519 | 524 | 488 | 133 | 147 | 150 | 157 |
| K17716 UDP-glucose 4-epimerase [EC:5.1.3.2] | 189 | 185 | 206 | 196 | 74 | 50 | 73 | 75 | 109 | 115 | 98 | 122 |
| K21620 galactitol 2-dehydrogenase [EC:1.1.1.16] | 2 | 0 | 0 | 2 | 0 | 0 | 0 | 5 | 2 | 2 | 2 | 1 |
| K21621 tagatose kinase [EC:2.7.1.101] | 0 | 2 | 0 | 0 | 4 | 2 | 3 | 2 | 2 | 0 | 2 | 7 |
| K21622 D-tagatose 6-phosphate 4-epimerase [EC:5.1.3.40] | 0 | 0 | 0 | 1 | 0 | 2 | 0 | 0 | 7 | 8 | 2 | 6 |
| ko00053 Ascorbate and aldarate metabolism | 31565 | 31134 | 30516 | 31214 | 27069 | 27246 | 27067 | 26966 | 31810 | 31724 | 31617 | 31764 |
| K01630 2-dehydro-3-deoxyglucarate aldolase [EC:4.1.2.20] | 736 | 719 | 667 | 770 | 367 | 336 | 352 | 361 | 824 | 912 | 887 | 885 |
| K01706 glucarate dehydratase [EC:4.2.1.40] | 1226 | 1190 | 1186 | 1252 | 1009 | 1030 | 1081 | 1072 | 1205 | 1300 | 1197 | 1233 |
| K01708 galactarate dehydratase [EC:4.2.1.42] | 874 | 873 | 852 | 928 | 708 | 701 | 656 | 681 | 1135 | 1122 | 1122 | 1141 |
| K02821 PTS system, ascorbate-specific IIA component [EC:2.7.1.194] | 1082 | 1013 | 1010 | 1057 | 648 | 578 | 543 | 500 | 1559 | 1706 | 1630 | 1771 |
| K02822 PTS system, ascorbate-specific IIB component [EC:2.7.1.194] | 841 | 776 | 849 | 845 | 373 | 353 | 375 | 424 | 1206 | 1330 | 1279 | 1242 |
| K03475 PTS system, ascorbate-specific IIC component | 3851 | 3813 | 3771 | 4014 | 2213 | 2255 | 2258 | 2055 | 5352 | 5462 | 5478 | 5488 |
| K03476 L-ascorbate 6-phosphate lactonase [EC:3.1.1.-] | 1146 | 1098 | 1060 | 1093 | 576 | 686 | 626 | 581 | 1334 | 1401 | 1441 | 1423 |
| K13874 L-arabinonolactonase [EC:3.1.1.15] | 273 | 364 | 297 | 312 | 80 | 79 | 61 | 64 | 648 | 720 | 642 | 652 |
| K13875 L-arabonate dehydrase [EC:4.2.1.25] | 18 | 20 | 16 | 14 | 3 | 1 | 2 | 3 | 39 | 30 | 26 | 27 |
| K13876 2-keto-3-deoxy-L-arabinonate dehydratase [EC:4.2.1.43] | 2 | 0 | 1 | 1 | 0 | 0 | 0 | 0 | 1 | 0 | 0 | 0 |
| K18981 uronate dehydrogenase [EC:1.1.1.203] | 0 | 2 | 0 | 0 | 0 | 0 | 0 | 2 | 2 | 2 | 0 | 1 |
| K18983 D-galactarolactone cycloisomerase [EC:5.5.1.27] | 43 | 54 | 40 | 52 | 4 | 4 | 8 | 6 | 112 | 95 | 96 | 131 |
| K20023 L-talarate/galactarate dehydratase [EC:4.2.1.156 4.2.1.42] | 267 | 261 | 234 | 257 | 337 | 353 | 321 | 300 | 69 | 77 | 83 | 84 |
| K23003 L-galactose dehydrogenase [EC:1.1.1.-] | 525 | 626 | 650 | 539 | 1906 | 1942 | 2073 | 2001 | 1449 | 1418 | 1577 | 1337 |
| K23004 L-galactono-1,5-lactonase [EC:3.1.1.-] | 579 | 635 | 763 | 574 | 2148 | 2193 | 2113 | 2164 | 1469 | 1420 | 1558 | 1325 |
| K23005 L-galactonate 5-dehydrogenase [EC:1.1.1.414] | 520 | 524 | 630 | 584 | 1904 | 1934 | 1901 | 1779 | 1429 | 1309 | 1307 | 1326 |
| K23007 L-galactonate 5-dehydrogenase [EC:1.1.1.414] | 226 | 219 | 236 | 209 | 297 | 361 | 337 | 311 | 66 | 63 | 82 | 83 |
| ko00500 Starch and sucrose metabolism | 170892 | 171950 | 172710 | 172783 | 219449 | 219930 | 220950 | 217325 | 233873 | 236153 | 235102 | 235064 |
| K00688 glycogen phosphorylase [EC:2.4.1.1] | 6185 | 6421 | 6444 | 6237 | 8504 | 8623 | 8431 | 8153 | 8608 | 8888 | 8959 | 8717 |
| K00690 sucrose phosphorylase [EC:2.4.1.7] | 318 | 290 | 349 | 318 | 70 | 58 | 62 | 42 | 474 | 518 | 449 | 504 |
| K00691 maltose phosphorylase [EC:2.4.1.8] | 177 | 178 | 171 | 213 | 51 | 47 | 50 | 34 | 128 | 225 | 142 | 214 |
| K00693 glycogen synthase [EC:2.4.1.11] | 0 | 0 | 0 | 0 | 2 | 0 | 0 | 0 | 0 | 0 | 0 | 0 |
| K00694 cellulose synthase (UDP-forming) [EC:2.4.1.12] | 3540 | 3546 | 3353 | 3629 | 2892 | 3025 | 2801 | 2835 | 4078 | 4291 | 3986 | 4262 |
| K00695 sucrose synthase [EC:2.4.1.13] | 0 | 0 | 0 | 0 | 0 | 0 | 0 | 0 | 0 | 0 | 2 | 0 |
| K00697 trehalose 6-phosphate synthase [EC:2.4.1.15 2.4.1.347] | 845 | 780 | 787 | 782 | 480 | 596 | 596 | 623 | 1152 | 1207 | 1084 | 1102 |
| K00700 1,4-alpha-glucan branching enzyme [EC:2.4.1.18] | 6462 | 6110 | 6258 | 6152 | 6080 | 6244 | 6255 | 6228 | 6869 | 7003 | 6728 | 6764 |
| K00702 cellobiose phosphorylase [EC:2.4.1.20] | 156 | 127 | 174 | 122 | 116 | 95 | 91 | 80 | 260 | 265 | 275 | 255 |
| K00703 starch synthase [EC:2.4.1.21] | 4177 | 3965 | 3914 | 4062 | 3609 | 3683 | 3581 | 3668 | 3659 | 3757 | 3570 | 3649 |
| K00705 4-alpha-glucanotransferase [EC:2.4.1.25] | 6389 | 6350 | 6405 | 6342 | 8163 | 7974 | 8232 | 8221 | 7412 | 7256 | 7207 | 7418 |
| K00975 glucose-1-phosphate adenylyltransferase [EC:2.7.7.27] | 2543 | 2459 | 2519 | 2552 | 1142 | 1250 | 1124 | 1157 | 2789 | 2953 | 2792 | 2914 |
| K00978 glucose-1-phosphate cytidylyltransferase [EC:2.7.7.33] | 781 | 832 | 884 | 837 | 1424 | 1425 | 1572 | 1430 | 1212 | 1160 | 1222 | 1103 |
| K01087 trehalose 6-phosphate phosphatase [EC:3.1.3.12] | 442 | 436 | 403 | 429 | 315 | 293 | 257 | 286 | 538 | 525 | 506 | 589 |
| K01176 alpha-amylase [EC:3.2.1.1] | 2391 | 2467 | 2547 | 2628 | 2064 | 2245 | 2070 | 2042 | 3137 | 3223 | 3102 | 3241 |

| | | | | | | | | | | | | |
|---|---|---|---|---|---|---|---|---|---|---|---|---|
| K01178 glucoamylase [EC:3.2.1.3] | 0 | 0 | 1 | 0 | 0 | 0 | 0 | 0 | 0 | 0 | 0 | 0 |
| K01179 endoglucanase [EC:3.2.1.4] | 1698 | 2008 | 2107 | 1985 | 6744 | 6610 | 6838 | 6589 | 5575 | 5269 | 5430 | 5180 |
| K01188 beta-glucosidase [EC:3.2.1.21] | 31 | 48 | 33 | 35 | 9 | 3 | 3 | 4 | 26 | 29 | 19 | 28 |
| K01194 alpha,alpha-trehalase [EC:3.2.1.28] | 3907 | 4151 | 3820 | 3903 | 2458 | 2568 | 2478 | 2616 | 3352 | 3528 | 3326 | 3468 |
| K01200 pullulanase [EC:3.2.1.41] | 2006 | 2118 | 2078 | 2080 | 2306 | 2459 | 2337 | 2359 | 3962 | 4010 | 3907 | 3955 |
| K01208 cyclomaltodextrinase / maltogenic alpha-amylase / neopullulanase [EC:3.2.1.54 3.2.1.133 3.2.1.135] | 361 | 346 | 396 | 350 | 71 | 92 | 70 | 77 | 554 | 635 | 570 | 608 |
| K01210 glucan 1,3-beta-glucosidase [EC:3.2.1.58] | 6 | 6 | 5 | 3 | 1 | 2 | 3 | 2 | 4 | 3 | 3 | 7 |
| K01212 levanase [EC:3.2.1.65] | 0 | 0 | 0 | 0 | 0 | 0 | 1 | 0 | 0 | 1 | 0 | 0 |
| K01214 isoamylase [EC:3.2.1.68] | 4004 | 3849 | 3641 | 3699 | 1975 | 2083 | 1914 | 2073 | 1735 | 1791 | 1654 | 1817 |
| K01226 trehalose-6-phosphate hydrolase [EC:3.2.1.93] | 1652 | 1610 | 1632 | 1648 | 947 | 967 | 877 | 879 | 2013 | 2040 | 1944 | 2059 |
| K01232 maltose-6'-phosphate glucosidase [EC:3.2.1.122] | 2048 | 2031 | 1797 | 2031 | 1528 | 1483 | 1565 | 1524 | 2044 | 2179 | 2078 | 2183 |
| K01236 maltooligosyltrehalose trehalohydrolase [EC:3.2.1.141] | 2885 | 2732 | 2464 | 2676 | 1644 | 1663 | 1595 | 1732 | 744 | 659 | 656 | 745 |
| K01513 ectonucleotide pyrophosphatase/phosphodiesterase family member 1/3 [EC:3.1.4.1 3.6.1.9] | 0 | 0 | 0 | 0 | 0 | 0 | 0 | 0 | 0 | 0 | 1 | 0 |
| K01838 beta-phosphoglucomutase [EC:5.4.2.6] | 782 | 846 | 733 | 784 | 154 | 181 | 196 | 184 | 580 | 608 | 597 | 629 |
| K02438 glycogen debranching enzyme [EC:3.2.1.196] | 1336 | 1305 | 1140 | 1208 | 943 | 956 | 986 | 939 | 1660 | 1805 | 1765 | 1798 |
| K02749 PTS system, alpha-glucoside-specific IIB component [EC:2.7.1.208 2.7.1.-] | 2357 | 2285 | 2216 | 2415 | 1167 | 1151 | 1115 | 1170 | 3660 | 4004 | 3822 | 4095 |
| K02759 PTS system, cellobiose-specific IIA component [EC:2.7.1.196 2.7.1.205] | 1007 | 1010 | 927 | 988 | 583 | 640 | 633 | 576 | 1404 | 1376 | 1376 | 1398 |
| K02760 PTS system, cellobiose-specific IIB component [EC:2.7.1.196 2.7.1.205] | 1217 | 1210 | 1151 | 1232 | 751 | 773 | 803 | 742 | 1416 | 1525 | 1441 | 1475 |
| K02761 PTS system, cellobiose-specific IIC component | 6568 | 6660 | 6655 | 6953 | 4127 | 4212 | 4058 | 3900 | 8486 | 8871 | 8450 | 8760 |
| K02810 PTS system, sucrose-specific IIC component | 2680 | 2451 | 2447 | 2534 | 1331 | 1372 | 1352 | 1234 | 2901 | 2866 | 2848 | 3027 |
| K02818 PTS system, trehalose-specific IIB component [EC:2.7.1.201] | 1718 | 1473 | 1572 | 1570 | 828 | 767 | 844 | 833 | 1852 | 1918 | 1822 | 1887 |
| K05341 amylosucrase [EC:2.4.1.4] | 328 | 322 | 378 | 343 | 44 | 45 | 61 | 42 | 357 | 342 | 371 | 318 |
| K05342 alpha,alpha-trehalose phosphorylase [EC:2.4.1.64] | 826 | 744 | 848 | 842 | 199 | 237 | 216 | 232 | 622 | 683 | 700 | 673 |
| K05343 maltose alpha-D-glucosyltransferase / alpha-amylase [EC:5.4.99.16 3.2.1.1] | 3055 | 3110 | 2805 | 2903 | 1416 | 1289 | 1298 | 1444 | 1916 | 2019 | 1916 | 1962 |
| K05349 beta-glucosidase [EC:3.2.1.21] | 26943 | 28311 | 29760 | 28470 | 68652 | 67441 | 69998 | 67356 | 59832 | 59055 | 60751 | 58564 |
| K05350 beta-glucosidase [EC:3.2.1.21] | 1972 | 1871 | 1862 | 1961 | 683 | 711 | 675 | 688 | 2175 | 2281 | 2252 | 2281 |
| K05988 dextranase [EC:3.2.1.11] | 1048 | 1062 | 1135 | 1068 | 3734 | 3636 | 3611 | 3786 | 3030 | 2873 | 3026 | 2833 |
| K06044 (1->4)-alpha-D-glucan 1-alpha-D-glucosylmutase [EC:5.4.99.15] | 4090 | 4131 | 3544 | 3668 | 2454 | 2351 | 2211 | 2619 | 1105 | 1019 | 893 | 1005 |
| K06896 maltose 6'-phosphate phosphatase [EC:3.1.3.90] | 70 | 79 | 81 | 87 | 14 | 16 | 17 | 23 | 73 | 67 | 77 | 57 |
| K07405 alpha-amylase [EC:3.2.1.1] | 815 | 892 | 860 | 804 | 2768 | 2632 | 2844 | 2642 | 1973 | 1834 | 1955 | 1873 |
| K16055 trehalose 6-phosphate synthase/phosphatase [EC:2.4.1.15 3.1.3.12] | 287 | 273 | 277 | 259 | 343 | 312 | 363 | 313 | 66 | 79 | 52 | 75 |
| K16147 starch synthase (maltosyl-transferring) [EC:2.4.99.16] | 2802 | 2615 | 2712 | 2708 | 1399 | 1405 | 1264 | 1435 | 842 | 785 | 780 | 849 |
| K16148 alpha-maltose-1-phosphate synthase [EC:2.4.1.342] | 6 | 1 | 2 | 1 | 0 | 4 | 3 | 0 | 2 | 24 | 7 | 20 |
| K16149 1,4-alpha-glucan branching enzyme [EC:2.4.1.18] | 1 | 0 | 0 | 0 | 0 | 0 | 0 | 0 | 0 | 0 | 0 | 0 |
| K16150 glycogen synthase [EC:2.4.1.11] | 0 | 0 | 0 | 0 | 0 | 0 | 0 | 1 | 0 | 0 | 0 | 0 |
| K16153 glycogen phosphorylase/synthase [EC:2.4.1.1 2.4.1.11] | 44 | 53 | 33 | 48 | 35 | 46 | 41 | 36 | 30 | 80 | 47 | 53 |
| K20108 PTS system, maltose-specific IIC component | 6 | 15 | 10 | 8 | 5 | 2 | 3 | 3 | 1 | 3 | 6 | 3 |
| K20811 inulosucrase [EC:2.4.1.9] | 0 | 0 | 0 | 0 | 1 | 0 | 3 | 1 | 0 | 2 | 2 | 1 |
| K21574 glucan 1,4-alpha-glucosidase [EC:3.2.1.3] | 2542 | 2701 | 2877 | 2705 | 9107 | 9067 | 9020 | 8791 | 6904 | 6443 | 6855 | 6309 |
| K22451 4-alpha-glucanotransferase [EC:2.4.1.25] | 2 | 3 | 4 | 6 | 77 | 73 | 94 | 43 | 13 | 10 | 8 | 12 |
| ko00520 Amino sugar and nucleotide sugar metabolism | 169556 | 172176 | 173086 | 173341 | 208423 | 209681 | 210430 | 207887 | 198209 | 199143 | 198548 | 199329 |
| K00075 UDP-N-acetylmuramate dehydrogenase [EC:1.3.1.98] | 2926 | 2947 | 3085 | 2872 | 3297 | 3293 | 3261 | 3156 | 3185 | 3292 | 3266 | 3238 |
| K00523 CDP-4-dehydro-6-deoxyglucose reductase, E3 [EC:1.17.1.1] | 1709 | 1616 | 1575 | 1618 | 918 | 951 | 974 | 920 | 534 | 497 | 546 | 496 |
| K00698 chitin synthase [EC:2.4.1.16] | 0 | 0 | 0 | 0 | 0 | 0 | 7 | 0 | 0 | 1 | 0 | 1 |
| K00790 UDP-N-acetylglucosamine 1-carboxyvinyltransferase [EC:2.5.1.7] | 4554 | 4585 | 4568 | 4572 | 4698 | 4801 | 4764 | 4706 | 5044 | 5245 | 5026 | 5183 |
| K00820 glutamine---fructose-6-phosphate transaminase (isomerizing) [EC:2.6.1.16] | 7697 | 8048 | 7856 | 7956 | 7582 | 7462 | 7512 | 7471 | 6392 | 6814 | 6788 | 6830 |
| K00884 N-acetylglucosamine kinase [EC:2.7.1.59] | 483 | 484 | 519 | 542 | 375 | 381 | 354 | 369 | 739 | 658 | 756 | 771 |
| K00885 N-acylmannosamine kinase [EC:2.7.1.60] | 191 | 220 | 185 | 202 | 270 | 358 | 278 | 317 | 64 | 64 | 55 | 50 |
| K00983 N-acylneuraminate cytidylyltransferase [EC:2.7.7.43] | 10 | 18 | 16 | 13 | 6 | 10 | 12 | 3 | 63 | 65 | 46 | 79 |
| K00992 N-acetyl-alpha-D-muramate 1-phosphate uridylyltransferase [EC:2.7.7.99] | 1339 | 1249 | 1230 | 1295 | 721 | 691 | 745 | 780 | 413 | 410 | 429 | 394 |
| K01183 chitinase [EC:3.2.1.14] | 693 | 751 | 706 | 781 | 349 | 363 | 372 | 266 | 1116 | 1231 | 1108 | 1242 |
| K01198 xylan 1,4-beta-xylosidase [EC:3.2.1.37] | 2949 | 2927 | 3148 | 3183 | 4585 | 4524 | 4509 | 4465 | 4836 | 4724 | 4819 | 5043 |
| K01207 beta-N-acetylhexosaminidase [EC:3.2.1.52] | 3873 | 3574 | 3672 | 3646 | 1612 | 1688 | 1524 | 1610 | 2495 | 2586 | 2467 | 2596 |
| K01209 alpha-L-arabinofuranosidase [EC:3.2.1.55] | 5134 | 5239 | 5499 | 5377 | 9252 | 9419 | 9774 | 9159 | 9916 | 10085 | 9932 | 9737 |
| K01443 N-acetylglucosamine-6-phosphate deacetylase [EC:3.5.1.25] | 5714 | 5811 | 5921 | 5787 | 8813 | 9010 | 9128 | 8941 | 7356 | 7456 | 7653 | 7274 |
| K01452 chitin deacetylase [EC:3.5.1.41] | 1 | 1 | 2 | 2 | 0 | 3 | 0 | 0 | 1 | 0 | 1 | 0 |
| K01639 N-acetylneuraminate lyase [EC:4.1.3.3] | 1266 | 1367 | 1353 | 1351 | 2186 | 2157 | 2082 | 2110 | 1842 | 1946 | 2011 | 1995 |
| K01654 N-acetylneuraminate synthase [EC:2.5.1.56] | 81 | 79 | 104 | 112 | 33 | 27 | 13 | 18 | 69 | 93 | 93 | 87 |
| K01709 CDP-glucose 4,6-dehydratase [EC:4.2.1.45] | 892 | 903 | 934 | 926 | 1446 | 1528 | 1459 | 1458 | 1188 | 1176 | 1313 | 1096 |
| K01787 N-acylglucosamine 2-epimerase [EC:5.1.3.8] | 1056 | 1133 | 1271 | 1118 | 3503 | 3522 | 3513 | 3443 | 2641 | 2483 | 2686 | 2408 |
| K01788 N-acylglucosamine-6-phosphate 2-epimerase [EC:5.1.3.9] | 890 | 827 | 836 | 870 | 444 | 426 | 382 | 430 | 1173 | 1453 | 1247 | 1316 |
| K01791 UDP-N-acetylglucosamine 2-epimerase (non-hydrolysing) [EC:5.1.3.14] | 2210 | 2334 | 2350 | 2282 | 2948 | 3012 | 2970 | 2871 | 3013 | 3133 | 3061 | 3076 |
| K01836 phosphoacetylglucosamine mutase [EC:5.4.2.3] | 0 | 0 | 0 | 0 | 0 | 0 | 0 | 0 | 0 | 0 | 2 | 0 |
| K02472 UDP-N-acetyl-D-mannosaminuronic acid dehydrogenase [EC:1.1.1.336] | 1073 | 1115 | 1046 | 1080 | 705 | 793 | 713 | 714 | 1127 | 1238 | 1106 | 1263 |
| K02473 UDP-N-acetylglucosamine 4-epimerase [EC:5.1.3.7] | 65 | 75 | 86 | 87 | 28 | 45 | 20 | 49 | 139 | 182 | 135 | 155 |
| K02474 UDP-N-acetyl-D-galactosamine dehydrogenase [EC:1.1.1.-] | 121 | 152 | 114 | 120 | 51 | 49 | 53 | 58 | 168 | 227 | 149 | 192 |
| K02564 glucosamine-6-phosphate deaminase [EC:3.5.99.6] | 3836 | 3987 | 4068 | 4057 | 8456 | 8355 | 8630 | 8362 | 7035 | 6951 | 6819 | 6862 |
| K02802 PTS system, N-acetylglucosamine-specific IIA component [EC:2.7.1.193] | 0 | 2 | 0 | 0 | 0 | 0 | 0 | 0 | 0 | 0 | 1 | 0 |
| K02803 PTS system, N-acetylglucosamine-specific IIB component [EC:2.7.1.193] | 4402 | 4420 | 4027 | 4273 | 2292 | 2198 | 2162 | 2385 | 2835 | 2911 | 2768 | 2985 |
| K03431 phosphoglucosamine mutase [EC:5.4.2.10] | 4357 | 4643 | 4514 | 4594 | 2419 | 2499 | 2259 | 2366 | 3789 | 3992 | 3756 | 4056 |
| K04042 bifunctional UDP-N-acetylglucosamine pyrophosphorylase / Glucosamine-1-phosphate N-acetyltransferase [EC:2.7.7.23 2.3.1.157] | 3633 | 3675 | 3504 | 3826 | 2398 | 2339 | 2224 | 2193 | 2478 | 2600 | 2445 | 2548 |
| K07102 N-acetylmuramate 1-kinase [EC:2.7.1.221] | 1837 | 1852 | 1700 | 1876 | 1147 | 1061 | 1063 | 1155 | 597 | 670 | 627 | 629 |
| K07106 N-acetylmuramic acid 6-phosphate etherase [EC:4.2.1.126] | 1936 | 1889 | 2005 | 1945 | 2656 | 2685 | 2780 | 2736 | 2901 | 2921 | 2923 | 2903 |
| K07806 UDP-4-amino-4-deoxy-L-arabinose-oxoglutarate aminotransferase [EC:2.6.1.87] | 2602 | 2532 | 2364 | 2549 | 1401 | 1376 | 1402 | 1624 | 1672 | 1653 | 1566 | 1572 |
| K08068 UDP-N-acetylglucosamine 2-epimerase (hydrolysing) [EC:3.2.1.183] | 0 | 0 | 0 | 0 | 0 | 0 | 0 | 0 | 0 | 2 | 0 | 0 |
| K08678 UDP-glucuronate decarboxylase [EC:4.1.1.35] | 524 | 623 | 646 | 574 | 1740 | 1900 | 1787 | 1817 | 1261 | 1197 | 1217 | 1287 |
| K08679 UDP-glucuronate 4-epimerase [EC:5.1.3.6] | 703 | 653 | 615 | 699 | 420 | 457 | 451 | 424 | 755 | 766 | 696 | 799 |
| K10011 UDP-4-amino-4-deoxy-L-arabinose formyltransferase / UDP-glucuronic acid dehydrogenase (UDP-4-keto-hexauronic acid decarboxylating) [EC:2.1.2.1 | 3065 | 3156 | 2686 | 2974 | 1666 | 1872 | 1710 | 1840 | 2274 | 2258 | 2252 | 2307 |
| K10012 undecaprenyl-phosphate 4-deoxy-4-formamido-L-arabinose transferase [EC:2.4.2.53] | 1848 | 1900 | 1714 | 1763 | 1129 | 1069 | 1079 | 1080 | 1451 | 1494 | 1295 | 1530 |
| K11192 PTS system, N-acetylmuramic acid-specific IIC component | 578 | 496 | 401 | 500 | 168 | 155 | 135 | 156 | 1044 | 1076 | 955 | 1034 |
| K12373 hexosaminidase [EC:3.2.1.52] | 12233 | 13626 | 14836 | 13716 | 42749 | 42813 | 43874 | 42787 | 31383 | 30037 | 31681 | 30023 |
| K12448 UDP-arabinose 4-epimerase [EC:5.1.3.5] | 0 | 0 | 2 | 0 | 0 | 0 | 0 | 0 | 0 | 4 | 1 | 0 | 2 |
| K12452 CDP-4-dehydro-6-deoxyglucose reductase, E1 [EC:1.17.1.1] | 955 | 974 | 1132 | 1109 | 1970 | 1963 | 2147 | 2043 | 1681 | 1617 | 1592 | 1694 |
| K12454 CDP-paratose 2-epimerase [EC:5.1.3.10] | 77 | 48 | 81 | 62 | 14 | 5 | 10 | 14 | 2 | 0 | 0 | 0 |
| K13010 perosamine synthetase [EC:2.6.1.102] | 0 | 3 | 3 | 5 | 1 | 2 | 4 | 3 | 4 | 5 | 6 | 4 |
| K13014 undecaprenyl phosphate-alpha-L-ara4FN deformylase [EC:3.5.1.-] | 1766 | 1864 | 1676 | 1751 | 987 | 1013 | 944 | 1143 | 1224 | 1254 | 1140 | 1219 |
| K13015 UDP-N-acetyl-D-glucosamine dehydrogenase [EC:1.1.1.136] | 1928 | 1892 | 2074 | 1907 | 1066 | 1204 | 1053 | 1100 | 644 | 618 | 575 | 630 |
| K13016 UDP-N-acetyl-2-amino-2-deoxyglucuronate dehydrogenase [EC:1.1.1.335] | 1411 | 1353 | 1464 | 1495 | 812 | 850 | 777 | 772 | 438 | 463 | 449 | 400 |
| K13017 UDP-2-acetamido-2-deoxy-ribo-hexuluronate aminotransferase [EC:2.6.1.98] | 1491 | 1555 | 1699 | 1597 | 866 | 913 | 847 | 958 | 502 | 479 | 465 | 444 |
| K13018 UDP-2-acetamido-3-amino-2,3-dideoxy-glucuronate N-acetyltransferase [EC:2.3.1.201] | 884 | 880 | 910 | 956 | 511 | 536 | 443 | 416 | 309 | 284 | 253 | 251 |
| K13019 UDP-GlcNAc3NAcA epimerase [EC:5.1.3.23] | 1488 | 1506 | 1662 | 1586 | 873 | 871 | 898 | 880 | 476 | 429 | 581 | 443 |
| K13020 UDP-N-acetyl-2-amino-2-deoxyglucuronate dehydrogenase [EC:1.1.1.335] | 1 | 0 | 0 | 1 | 0 | 0 | 0 | 0 | 0 | 0 | 0 | 0 |
| K13381 bifunctional chitinase/lysozyme [EC:3.2.1.14 3.2.1.17] | 6 | 5 | 9 | 8 | 8 | 5 | 12 | 5 | 142 | 179 | 128 | 156 |
| K15855 exo-1,4-beta-D-glucosaminidase [EC:3.2.1.165] | 0 | 0 | 0 | 0 | 0 | 2 | 0 | 0 | 0 | 0 | 0 | 0 |
| K15894 UDP-N-acetylglucosamine 4,6-dehydratase [EC:4.2.1.115] | 1 | 0 | 0 | 1 | 0 | 0 | 0 | 0 | 1 | 3 | 0 | 3 |
| K15895 UDP-4-amino-4,6-dideoxy-L-N-acetyl-beta-L-altrosamine transaminase [EC:2.6.1.92] | 0 | 0 | 0 | 0 | 0 | 0 | 0 | 0 | 1 | 1 | 0 | 0 |
| K15898 pseudaminic acid synthase [EC:2.5.1.97] | 76 | 66 | 53 | 76 | 12 | 9 | 12 | 16 | 116 | 128 | 107 | 136 |
| K15899 pseudaminic acid cytidylyltransferase [EC:2.7.7.81] | 13 | 15 | 15 | 15 | 7 | 4 | 5 | 3 | 27 | 21 | 23 | 24 |
| K15913 UDP-N-acetylbacillosamine N-acetyltransferase [EC:2.3.1.203] | 0 | 0 | 0 | 0 | 0 | 0 | 0 | 0 | 0 | 0 | 2 | 0 |
| K15920 xylan 1,4-beta-xylosidase [EC:3.2.1.37] | 2 | 1 | 0 | 1 | 0 | 0 | 0 | 0 | 1 | 0 | 1 | 0 |
| K15921 arabinoxylan arabinofuranohydrolase [EC:3.2.1.55] | 71 | 81 | 85 | 88 | 311 | 289 | 315 | 278 | 202 | 189 | 232 | 217 |
| K18429 GDP/UDP-N,N'-diacetylbacillosamine 2-epimerase (hydrolysing) [EC:3.2.1.184] | 89 | 87 | 95 | 84 | 24 | 35 | 29 | 26 | 25 | 29 | 20 | 38 |
| K18430 N,N'-diacetyllegionaminate synthase [EC:2.5.1.101] | 111 | 131 | 114 | 122 | 46 | 40 | 52 | 35 | 21 | 29 | 24 | 28 |
| K18431 CMP-N,N'-diacetyllegionaminic acid synthase [EC:2.7.7.82] | 57 | 42 | 76 | 44 | 13 | 19 | 40 | 18 | 21 | 24 | 19 | 22 |
| K18675 N,N'-diacetylchitobiose phosphorylase [EC:2.4.1.280] | 766 | 758 | 739 | 702 | 132 | 148 | 154 | 109 | 850 | 939 | 955 | 943 |
| K18676 glucosamine kinase [EC:2.7.1.8] | 11 | 18 | 15 | 13 | 0 | 0 | 5 | 2 | 8 | 13 | 13 | 7 |
| K21749 N-acylneuraminate/3-deoxy-D-glycero-D-galacto-nononate cytidylyltransferase [EC:2.7.7.43 2.7.7.92] | 0 | 0 | 0 | 0 | 0 | 0 | 0 | 0 | 0 | 2 | 0 | 1 |
| K22292 N-acetyl-D-muramate 6-phosphate phosphatase [EC:3.1.3.105] | 323 | 325 | 355 | 318 | 179 | 190 | 182 | 214 | 133 | 91 | 114 | 119 |
| ko00620 Pyruvate metabolism | 214233 | 214477 | 211102 | 214083 | 196205 | 198334 | 196154 | 197638 | 211431 | 213018 | 211213 | 213044 |
| K00027 malate dehydrogenase (oxaloacetate-decarboxylating) [EC:1.1.1.38] | 2930 | 2877 | 2871 | 2983 | 1923 | 2090 | 1951 | 1870 | 2048 | 2395 | 2230 | 2361 |
| K00029 malate dehydrogenase (oxaloacetate-decarboxylating)(NADP+) [EC:1.1.1.40] | 5209 | 5189 | 5154 | 5318 | 7027 | 7403 | 7149 | 7200 | 6273 | 6175 | 6137 | 6120 |
| K00049 glyoxylate/hydroxypyruvate reductase [EC:1.1.1.79 1.1.1.81] | 1 | 0 | 0 | 0 | 0 | 0 | 1 | 1 | 2 | 4 | 2 | 4 |
| K00051 malate dehydrogenase (NADP+) [EC:1.1.1.82] | 0 | 0 | 0 | 0 | 0 | 0 | 0 | 0 | 0 | 0 | 0 | 1 |
| K00101 L-lactate dehydrogenase (cytochrome) [EC:1.1.2.3] | 4494 | 4640 | 4165 | 4450 | 2519 | 2404 | 2267 | 2439 | 2929 | 3113 | 2871 | 3004 |
| K00102 D-lactate dehydrogenase (cytochrome) [EC:1.1.2.4] | 28 | 19 | 20 | 25 | 9 | 29 | 24 | 32 | 50 | 50 | 60 | 65 |
| K00132 acetaldehyde dehydrogenase (acetylating) [EC:1.2.1.10] | 521 | 502 | 477 | 475 | 101 | 101 | 99 | 77 | 281 | 355 | 272 | 316 |
| K00156 pyruvate dehydrogenase (quinone) [EC:1.2.5.1] | 6126 | 6183 | 6034 | 6165 | 6661 | 6490 | 6576 | 6476 | 5418 | 5462 | 5494 | 5374 |
| K00158 pyruvate oxidase [EC:1.2.3.3] | 5 | 6 | 3 | 5 | 17 | 11 | 12 | 6 | 5 | 10 | 8 | 5 |
| K00467 lactate 2-monooxygenase [EC:1.13.12.4] | 1 | 0 | 0 | 0 | 0 | 0 | 0 | 0 | 0 | 0 | 0 | 0 |
| K00625 phosphate acetyltransferase [EC:2.3.1.8] | 826 | 908 | 947 | 963 | 2238 | 2351 | 2492 | 2363 | 1822 | 1739 | 1902 | 1771 |
| K00626 acetyl-CoA C-acetyltransferase [EC:2.3.1.9] | 12594 | 12790 | 11954 | 12454 | 6888 | 6591 | 6385 | 6965 | 6821 | 6853 | 6638 | 6879 |

| | | | | | | | | | | | | |
|---|---|---|---|---|---|---|---|---|---|---|---|---|
| K00656 formate C-acetyltransferase [EC:2.3.1.54] | 7945 | 7917 | 8001 | 8345 | 7622 | 7705 | 7755 | 7440 | 11462 | 11561 | 11251 | 11563 |
| K00925 acetate kinase [EC:2.7.2.1] | 3296 | 3363 | 3477 | 3447 | 3947 | 4034 | 3901 | 3969 | 4148 | 4181 | 4090 | 4204 |
| K01003 oxaloacetate decarboxylase [EC:4.1.1.112] | 335 | 364 | 345 | 411 | 187 | 173 | 160 | 228 | 106 | 139 | 139 | 130 |
| K01006 pyruvate, orthophosphate dikinase [EC:2.7.9.1] | 3379 | 3242 | 3490 | 3196 | 5671 | 5800 | 5897 | 5735 | 5078 | 5121 | 5307 | 4888 |
| K01007 pyruvate, water dikinase [EC:2.7.9.2] | 5017 | 4873 | 5022 | 4913 | 2926 | 3187 | 2887 | 3093 | 3465 | 3464 | 3380 | 3573 |
| K01026 propionate CoA-transferase [EC:2.8.3.1] | 722 | 833 | 699 | 788 | 190 | 225 | 214 | 205 | 1436 | 1406 | 1544 | 1542 |
| K01067 acetyl-CoA hydrolase [EC:3.1.2.1] | 0 | 0 | 0 | 0 | 1 | 0 | 0 | 0 | 0 | 0 | 0 | 0 |
| K01069 hydroxyacylglutathione hydrolase [EC:3.1.2.6] | 3441 | 3472 | 3483 | 3508 | 3006 | 3075 | 3125 | 3021 | 2876 | 2791 | 2947 | 2861 |
| K01512 acylphosphatase [EC:3.6.1.7] | 739 | 755 | 725 | 761 | 403 | 405 | 428 | 396 | 548 | 515 | 531 | 561 |
| K01571 oxaloacetate decarboxylase (Na+ extruding) subunit alpha [EC:7.2.4.2] | 1987 | 1940 | 2047 | 1978 | 651 | 667 | 644 | 615 | 3462 | 3549 | 3473 | 3659 |
| K01573 oxaloacetate decarboxylase (Na+ extruding) subunit gamma | 143 | 144 | 156 | 159 | 462 | 434 | 440 | 461 | 356 | 315 | 350 | 349 |
| K01595 phosphoenolpyruvate carboxylase [EC:4.1.1.31] | 5733 | 5478 | 5363 | 5511 | 3393 | 3386 | 3216 | 3462 | 3684 | 3584 | 3501 | 3668 |
| K01638 malate synthase [EC:2.3.3.9] | 4633 | 4764 | 4455 | 4608 | 2739 | 2892 | 2747 | 2841 | 2776 | 2824 | 2661 | 2731 |
| K01649 2-isopropylmalate synthase [EC:2.3.3.13] | 5799 | 5824 | 5830 | 5944 | 5937 | 5857 | 5723 | 5763 | 6806 | 6975 | 6960 | 7000 |
| K01759 lactoylglutathione lyase [EC:4.4.1.5] | 3219 | 3155 | 3203 | 3128 | 2364 | 2303 | 2311 | 2320 | 2190 | 2142 | 2280 | 2265 |
| K01961 acetyl-CoA carboxylase, biotin carboxylase subunit [EC:6.4.1.2 6.3.4.14] | 6545 | 6692 | 6582 | 6701 | 8752 | 8665 | 9021 | 9035 | 7342 | 7278 | 7398 | 7276 |
| K01962 acetyl-CoA carboxylase carboxyl transferase subunit alpha [EC:6.4.1.2 2.1.3.15] | 1838 | 1852 | 1945 | 1848 | 1172 | 1258 | 1174 | 1308 | 1327 | 1443 | 1394 | 1391 |
| K01963 acetyl-CoA carboxylase carboxyl transferase subunit beta [EC:6.4.1.2 2.1.3.15] | 3022 | 2941 | 3063 | 2958 | 1607 | 1574 | 1475 | 1488 | 2072 | 2130 | 1947 | 2156 |
| K02160 acetyl-CoA carboxylase biotin carboxyl carrier protein | 666 | 678 | 660 | 627 | 438 | 423 | 505 | 462 | 592 | 635 | 563 | 626 |
| K02594 homocitrate synthase NifV [EC:2.3.3.14] | 274 | 274 | 209 | 220 | 317 | 353 | 317 | 319 | 12 | 14 | 13 | 13 |
| K03777 D-lactate dehydrogenase (quinone) [EC:1.1.5.12] | 1052 | 1006 | 934 | 1000 | 631 | 727 | 666 | 664 | 1369 | 1570 | 1424 | 1429 |
| K03778 D-lactate dehydrogenase [EC:1.1.1.28] | 2892 | 3138 | 2939 | 2981 | 3557 | 3360 | 3398 | 3298 | 3266 | 3112 | 3226 | 3184 |
| K04020 phosphotransacetylase | 373 | 387 | 341 | 420 | 113 | 142 | 130 | 131 | 823 | 823 | 843 | 868 |
| K04021 aldehyde dehydrogenase | 567 | 554 | 452 | 567 | 122 | 158 | 131 | 162 | 1096 | 1108 | 1107 | 1202 |
| K04073 acetaldehyde dehydrogenase [EC:1.2.1.10] | 315 | 280 | 266 | 314 | 71 | 85 | 72 | 75 | 569 | 558 | 533 | 568 |
| K05523 D-lactate dehydratase / protein deglycase [EC:4.2.1.130 3.5.1.124] | 1199 | 1261 | 1149 | 1187 | 683 | 667 | 583 | 658 | 468 | 396 | 394 | 381 |
| K07248 lactaldehyde dehydrogenase / glycolaldehyde dehydrogenase [EC:1.2.1.22 1.2.1.21] | 962 | 946 | 970 | 902 | 715 | 838 | 795 | 787 | 1141 | 999 | 1092 | 1161 |
| K10977 methanogen homocitrate synthase [EC:2.3.3.14 2.3.3.-] | 0 | 2 | 0 | 0 | 0 | 2 | 2 | 3 | 5 | 7 | 3 | 20 |
| K11262 acetyl-CoA carboxylase / biotin carboxylase 1 [EC:6.4.1.2 6.3.4.14 2.1.3.15] | 0 | 1 | 0 | 0 | 0 | 0 | 0 | 0 | 0 | 0 | 0 | 0 |
| K11263 acetyl-CoA/propionyl-CoA carboxylase, biotin carboxylase, biotin carboxyl carrier protein [EC:6.4.1.2 6.4.1.3 6.3.4.14] | 4 | 0 | 7 | 5 | 8 | 5 | 5 | 11 | 14 | 19 | 19 | 18 |
| K12972 glyoxylate/hydroxypyruvate reductase [EC:1.1.1.79 1.1.1.81] | 1137 | 1005 | 1024 | 1038 | 669 | 730 | 672 | 733 | 1340 | 1421 | 1401 | 1441 |
| K13788 phosphate acetyltransferase [EC:2.3.1.8] | 3822 | 3778 | 3660 | 3827 | 2226 | 2252 | 2136 | 2331 | 2613 | 2530 | 2488 | 2628 |
| K15024 putative phosphotransacetylase [EC:2.3.1.8] | 102 | 134 | 114 | 122 | 32 | 30 | 30 | 23 | 161 | 144 | 144 | 175 |
| K19266 lactaldehyde dehydrogenase [EC:1.2.1.22] | 0 | 0 | 3 | 0 | 2 | 2 | 0 | 1 | 1 | 10 | 8 | 19 |
| K20509 carboxybiotin decarboxylase [EC:7.2.4.1] | 5192 | 5339 | 5507 | 5349 | 8011 | 8264 | 8525 | 7967 | 9880 | 9486 | 9603 | 9806 |
| K22212 malolactic enzyme [EC:4.1.1.101] | 9 | 5 | 19 | 13 | 40 | 28 | 14 | 16 | 22 | 33 | 32 | 24 |
| K22373 lactate racemase [EC:5.1.2.1] | 121 | 147 | 151 | 104 | 108 | 137 | 123 | 126 | 155 | 190 | 156 | 177 |
| K23257 methylglyoxal/glyoxal reductase [EC:1.1.1.283 1.1.1.-] | 351 | 360 | 359 | 412 | 62 | 67 | 87 | 69 | 289 | 293 | 351 | 318 |
| ko00630 Glyoxylate and dicarboxylate metabolism | 166949 | 165626 | 161838 | 164554 | 151801 | 150936 | 149939 | 152568 | 143537 | 142741 | 141973 | 143116 |
| K00015 glyoxylate reductase [EC:1.1.1.26] | 157 | 160 | 147 | 137 | 30 | 48 | 32 | 35 | 128 | 146 | 125 | 117 |
| K00018 glycerate dehydrogenase [EC:1.1.1.29] | 2829 | 2934 | 2829 | 2928 | 3384 | 3389 | 3173 | 3221 | 2443 | 2232 | 2343 | 2242 |
| K00023 acetoacetyl-CoA reductase [EC:1.1.1.36] | 4 | 2 | 1 | 5 | 0 | 0 | 0 | 1 | 5 | 0 | 0 | 0 |
| K00042 2-hydroxy-3-oxopropionate reductase [EC:1.1.1.60] | 1782 | 1805 | 1742 | 1906 | 1057 | 1064 | 1010 | 1079 | 1389 | 1437 | 1272 | 1425 |
| K00048 lactaldehyde reductase [EC:1.1.1.77] | 1622 | 1672 | 1676 | 1709 | 2771 | 2693 | 2959 | 2897 | 3218 | 3155 | 3161 | 3284 |
| K00050 hydroxypyruvate reductase [EC:1.1.1.81] | 19 | 14 | 16 | 19 | 26 | 30 | 29 | 29 | 5 | 9 | 3 | 3 |
| K00104 glycolate oxidase [EC:1.1.3.15] | 2825 | 2710 | 2707 | 2630 | 1540 | 1395 | 1317 | 1572 | 1182 | 1297 | 1172 | 1204 |
| K00123 formate dehydrogenase major subunit [EC:1.17.1.9] | 3153 | 3196 | 3192 | 3230 | 2283 | 2466 | 2353 | 2415 | 3640 | 3732 | 3600 | 3756 |
| K00124 formate dehydrogenase iron-sulfur subunit | 2179 | 2244 | 2012 | 2167 | 1251 | 1283 | 1233 | 1462 | 1365 | 1382 | 1275 | 1336 |
| K00127 formate dehydrogenase subunit gamma | 1306 | 1270 | 1185 | 1191 | 835 | 894 | 803 | 877 | 947 | 891 | 989 | 1022 |
| K00281 glycine dehydrogenase [EC:1.4.4.2] | 11909 | 11722 | 11036 | 11585 | 10117 | 10076 | 9784 | 10444 | 8371 | 8044 | 7883 | 8286 |
| K00282 glycine dehydrogenase subunit 1 [EC:1.4.4.2] | 261 | 244 | 285 | 300 | 150 | 171 | 153 | 151 | 244 | 291 | 207 | 261 |
| K00283 glycine dehydrogenase subunit 2 [EC:1.4.4.2] | 502 | 502 | 578 | 500 | 186 | 208 | 238 | 206 | 421 | 447 | 410 | 401 |
| K00284 glutamate synthase (ferredoxin) [EC:1.4.7.1] | 0 | 1 | 0 | 1 | 1 | 0 | 1 | 0 | 0 | 0 | 1 | 2 |
| K00600 glycine hydroxymethyltransferase [EC:2.1.2.1] | 7337 | 7324 | 7059 | 7016 | 5975 | 5980 | 5914 | 5982 | 5305 | 5019 | 4966 | 5089 |
| K00605 aminomethyltransferase [EC:2.1.2.10] | 5198 | 4861 | 4793 | 4843 | 4648 | 4471 | 4657 | 4688 | 3830 | 3582 | 3702 | 3787 |
| K00830 alanine-glyoxylate transaminase / serine-glyoxylate transaminase / serine-pyruvate transaminase [EC:2.6.1.44 2.6.1.45 2.6.1.51] | 1 | 9 | 3 | 7 | 0 | 4 | 0 | 2 | 7 | 14 | 3 | 7 |
| K00865 glycerate 2-kinase [EC:2.7.1.165] | 3393 | 3614 | 3388 | 3486 | 3549 | 3779 | 3714 | 3650 | 4913 | 4975 | 4969 | 4881 |
| K01091 phosphoglycolate phosphatase [EC:3.1.3.18] | 5374 | 5387 | 5416 | 5395 | 5363 | 5279 | 5455 | 5452 | 4357 | 4435 | 4476 | 4410 |
| K01432 arylformamidase [EC:3.5.1.9] | 0 | 2 | 1 | 1 | 0 | 4 | 2 | 1 | 0 | 2 | 2 | 1 |
| K01433 formyltetrahydrofolate deformylase [EC:3.5.1.10] | 3475 | 3469 | 3512 | 3572 | 3545 | 3496 | 3489 | 3359 | 2886 | 2707 | 2745 | 2634 |
| K01455 formamidase [EC:3.5.1.49] | 56 | 39 | 52 | 47 | 3 | 5 | 3 | 0 | 112 | 91 | 98 | 107 |
| K01458 N-formylglutamate deformylase [EC:3.5.1.68] | 424 | 424 | 448 | 427 | 371 | 395 | 409 | 354 | 86 | 102 | 87 | 71 |
| K01569 oxalate decarboxylase [EC:4.1.1.2] | 1 | 0 | 0 | 0 | 0 | 0 | 0 | 0 | 0 | 0 | 0 | 0 |
| K01577 oxalyl-CoA decarboxylase [EC:4.1.1.8] | 8 | 11 | 11 | 3 | 28 | 23 | 26 | 22 | 80 | 104 | 109 | 97 |
| K01601 ribulose-bisphosphate carboxylase large chain [EC:4.1.1.39] | 8 | 10 | 10 | 13 | 0 | 4 | 4 | 2 | 3 | 5 | 8 | 9 |
| K01608 tartronate-semialdehyde synthase [EC:4.1.1.47] | 175 | 199 | 195 | 201 | 117 | 120 | 120 | 119 | 288 | 306 | 312 | 265 |
| K01637 isocitrate lyase [EC:4.1.3.1] | 2456 | 2370 | 2381 | 2529 | 1549 | 1591 | 1423 | 1487 | 1664 | 1720 | 1641 | 1694 |
| K01816 hydroxypyruvate isomerase [EC:5.3.1.22] | 828 | 684 | 797 | 794 | 343 | 352 | 389 | 379 | 388 | 451 | 452 | 453 |
| K01846 methylaspartate mutase sigma subunit [EC:5.4.99.1] | 20 | 23 | 9 | 15 | 2 | 0 | 7 | 3 | 38 | 43 | 41 | 27 |
| K01847 methylmalonyl-CoA mutase [EC:5.4.99.2] | 2870 | 2877 | 3145 | 3015 | 8943 | 9073 | 9289 | 8726 | 6520 | 6283 | 6668 | 6373 |
| K01848 methylmalonyl-CoA mutase, N-terminal domain [EC:5.4.99.2] | 103 | 124 | 113 | 115 | 10 | 2 | 10 | 4 | 242 | 296 | 269 | 261 |
| K01849 methylmalonyl-CoA mutase, C-terminal domain [EC:5.4.99.2] | 19 | 37 | 23 | 26 | 0 | 0 | 0 | 2 | 48 | 51 | 41 | 50 |
| K01915 glutamine synthetase [EC:6.3.1.2] | 22068 | 21588 | 21624 | 21420 | 21354 | 20720 | 21133 | 21547 | 16407 | 16493 | 16608 | 16343 |
| K01965 propionyl-CoA carboxylase alpha chain [EC:6.4.1.3] | 2 | 2 | 0 | 1 | 2 | 4 | 0 | 1 | 0 | 5 | 5 | 7 |
| K01966 propionyl-CoA carboxylase beta chain [EC:6.4.1.3 2.1.3.15] | 1894 | 1885 | 2156 | 2030 | 6363 | 6046 | 6237 | 6211 | 4383 | 4390 | 4578 | 4345 |
| K02437 glycine cleavage system H protein | 1982 | 1966 | 1784 | 1905 | 1825 | 1775 | 1737 | 1836 | 1417 | 1442 | 1516 | 1506 |
| K03418 N,N-dimethylformamidase [EC:3.5.1.56] | 0 | 0 | 0 | 0 | 0 | 0 | 0 | 0 | 0 | 2 | 2 | 5 |
| K03779 L(+)-tartrate dehydratase alpha subunit [EC:4.2.1.32] | 609 | 563 | 623 | 564 | 195 | 206 | 191 | 208 | 886 | 1000 | 931 | 905 |
| K03780 L(+)-tartrate dehydratase beta subunit [EC:4.2.1.32] | 499 | 521 | 487 | 475 | 199 | 193 | 200 | 198 | 706 | 806 | 727 | 755 |
| K03781 catalase [EC:1.11.1.6] | 7175 | 7002 | 7010 | 7190 | 3948 | 3758 | 3777 | 3844 | 4126 | 4085 | 3953 | 4001 |
| K04835 methylaspartate ammonia-lyase [EC:4.3.1.2] | 52 | 44 | 40 | 40 | 3 | 4 | 0 | 2 | 105 | 95 | 109 | 103 |
| K05606 methylmalonyl-CoA/ethylmalonyl-CoA epimerase [EC:5.1.99.1] | 287 | 276 | 345 | 291 | 832 | 861 | 851 | 888 | 681 | 579 | 632 | 656 |
| K07130 arylformamidase [EC:3.5.1.9] | 1003 | 934 | 847 | 867 | 537 | 475 | 426 | 542 | 339 | 339 | 350 | 383 |
| K07246 tartrate dehydrogenase/decarboxylase / D-malate dehydrogenase [EC:1.1.1.93 4.1.1.73 1.1.1.83] | 467 | 470 | 431 | 461 | 154 | 167 | 178 | 166 | 821 | 840 | 801 | 868 |
| K11472 glycolate oxidase FAD binding subunit | 2049 | 1906 | 1633 | 1820 | 1001 | 976 | 865 | 1061 | 660 | 655 | 584 | 638 |
| K11473 glycolate oxidase iron-sulfur subunit | 1792 | 1803 | 1602 | 1702 | 949 | 900 | 898 | 1001 | 701 | 639 | 632 | 633 |
| K11517 (S)-2-hydroxy-acid oxidase [EC:1.1.3.15] | 0 | 0 | 2 | 0 | 0 | 0 | 0 | 0 | 0 | 0 | 0 | 0 |
| K14067 malate-CoA ligase subunit beta [EC:6.2.1.9] | 0 | 0 | 0 | 0 | 0 | 0 | 0 | 0 | 0 | 1 | 0 | 0 |
| K15893 glycerate dehydrogenase [EC:1.1.1.29] | 0 | 0 | 3 | 0 | 0 | 0 | 0 | 0 | 2 | 5 | 3 | 4 |
| K15919 glyoxylate/hydroxypyruvate reductase [EC:1.1.1.79 1.1.1.81] | 6 | 1 | 2 | 0 | 2 | 5 | 3 | 1 | 3 | 5 | 8 | 0 |
| K17865 3-hydroxybutyryl-CoA dehydratase [EC:4.2.1.55] | 14 | 16 | 10 | 9 | 0 | 2 | 0 | 2 | 121 | 81 | 95 | 81 |
| K18123 4-hydroxy-2-oxoglutarate aldolase [EC:4.1.3.16] | 4 | 1 | 2 | 2 | 0 | 0 | 1 | 0 | 2 | 8 | 3 | 3 |
| K19268 methylaspartate mutase epsilon subunit [EC:5.4.99.1] | 49 | 52 | 55 | 47 | 4 | 6 | 7 | 2 | 116 | 143 | 116 | 155 |
| K19312 acetyl-CoA/propionyl-CoA carboxylase carboxyl transferase subunit [EC:6.4.1.2 6.4.1.3 2.1.3.15] | 17 | 12 | 9 | 14 | 0 | 2 | 3 | 1 | 15 | 16 | 22 | 20 |
| K22223 phosphoglycolate phosphatase [EC:3.1.3.18] | 8 | 12 | 19 | 17 | 8 | 4 | 3 | 9 | 11 | 11 | 12 | 16 |
| K22982 hydroxypyruvate reductase [EC:1.1.1.81] | 0 | 0 | 1 | 1 | 0 | 1 | 0 | 0 | 8 | 5 | 2 | 8 |
| ko00640 Propanoate metabolism | 135970 | 136336 | 133365 | 136367 | 121875 | 120804 | 120364 | 121682 | 126428 | 126127 | 125023 | 127665 |
| K00005 glycerol dehydrogenase [EC:1.1.1.6] | 1910 | 1939 | 2001 | 2111 | 1263 | 1249 | 1224 | 1189 | 2681 | 2744 | 2660 | 2733 |
| K00086 1,3-propanediol dehydrogenase [EC:1.1.1.202] | 518 | 506 | 473 | 532 | 141 | 150 | 138 | 109 | 1006 | 984 | 967 | 964 |
| K00140 malonate-semialdehyde dehydrogenase (acetylating) / methylmalonate-semialdehyde dehydrogenase [EC:1.2.1.18 1.2.1.27] | 6337 | 6070 | 6133 | 6269 | 3295 | 3105 | 3090 | 3356 | 3449 | 3543 | 3356 | 3453 |
| K00166 2-oxoisovalerate dehydrogenase E1 component alpha subunit [EC:1.2.4.4] | 1634 | 1549 | 1476 | 1553 | 856 | 722 | 674 | 865 | 489 | 529 | 491 | 461 |
| K00167 2-oxoisovalerate dehydrogenase E1 component beta subunit [EC:1.2.4.4] | 1569 | 1482 | 1369 | 1523 | 754 | 670 | 681 | 873 | 509 | 493 | 421 | 478 |
| K00232 acyl-CoA oxidase [EC:1.3.3.6] | 0 | 0 | 0 | 0 | 0 | 0 | 0 | 0 | 0 | 0 | 3 | 0 |
| K00248 butyryl-CoA dehydrogenase [EC:1.3.8.1] | 2181 | 2107 | 1995 | 2105 | 1256 | 1107 | 1142 | 1159 | 1076 | 1071 | 960 | 1072 |
| K00822 beta-alanine--pyruvate transaminase [EC:2.6.1.18] | 4661 | 4602 | 4448 | 4407 | 2607 | 2372 | 2306 | 2470 | 1425 | 1440 | 1256 | 1422 |
| K00823 4-aminobutyrate aminotransferase [EC:2.6.1.19] | 796 | 809 | 810 | 773 | 558 | 572 | 560 | 509 | 903 | 1031 | 887 | 1109 |
| K00932 propionate kinase [EC:2.7.2.15] | 495 | 410 | 364 | 515 | 149 | 130 | 143 | 151 | 929 | 1046 | 970 | 1026 |
| K01034 acetate CoA/acetoacetate CoA-transferase alpha subunit [EC:2.8.3.8 2.8.3.9] | 5 | 4 | 7 | 3 | 6 | 4 | 8 | 10 | 27 | 49 | 39 | 50 |
| K01035 acetate CoA/acetoacetate CoA-transferase beta subunit [EC:2.8.3.8 2.8.3.9] | 25 | 28 | 26 | 17 | 1 | 7 | 6 | 10 | 47 | 58 | 56 | 71 |
| K01574 acetoacetate decarboxylase [EC:4.1.1.4] | 5 | 0 | 1 | 3 | 0 | 2 | 1 | 1 | 6 | 5 | 6 | 1 |
| K01578 malonyl-CoA decarboxylase [EC:4.1.1.9] | 1 | 0 | 0 | 0 | 0 | 2 | 0 | 0 | 0 | 0 | 0 | 0 |
| K01659 2-methylcitrate synthase [EC:2.3.3.5] | 1830 | 1831 | 1579 | 1644 | 922 | 925 | 899 | 1037 | 615 | 553 | 641 | 634 |
| K01692 enoyl-CoA hydratase [EC:4.2.1.17] | 2564 | 2527 | 2395 | 2447 | 1388 | 1298 | 1221 | 1333 | 1259 | 1171 | 1202 | 1224 |
| K01699 propanediol dehydratase large subunit [EC:4.2.1.28] | 1066 | 1114 | 1062 | 1073 | 641 | 776 | 709 | 662 | 1396 | 1484 | 1426 | 1608 |
| K01720 2-methylcitrate dehydratase [EC:4.2.1.79] | 2277 | 2286 | 2125 | 2145 | 1242 | 1197 | 1176 | 1242 | 784 | 708 | 696 | 790 |
| K01734 methylglyoxal synthase [EC:4.2.3.3] | 1150 | 1399 | 1222 | 1243 | 1539 | 1518 | 1570 | 1388 | 1740 | 1838 | 1854 | 1746 |
| K01782 3-hydroxyacyl-CoA dehydrogenase / enoyl-CoA hydratase / 3-hydroxybutyryl-CoA epimerase [EC:1.1.1.35 4.2.1.17 5.1.2.3] | 4154 | 4016 | 3994 | 4206 | 2376 | 2402 | 2240 | 2417 | 2705 | 2627 | 2473 | 2733 |
| K01825 3-hydroxyacyl-CoA dehydrogenase / enoyl-CoA hydratase / 3-hydroxybutyryl-CoA epimerase / enoyl-CoA isomerase [EC:1.1.1.35 4.2.1.17 5.1.2.3 5.3.3 | 4786 | 4649 | 4709 | 4546 | 2924 | 2929 | 2807 | 2978 | 2916 | 2903 | 2954 | 3089 |
| K01908 propionyl-CoA synthetase [EC:6.2.1.17] | 2515 | 2413 | 2227 | 2350 | 1265 | 1273 | 1247 | 1381 | 933 | 940 | 832 | 871 |
| K03416 methylmalonyl-CoA carboxyltransferase 5S subunit [EC:2.1.3.1] | 0 | 1 | 1 | 1 | 1 | 0 | 0 | 0 | 1 | 0 | 0 | 0 |
| K03417 methylisocitrate lyase [EC:4.1.3.30] | 1310 | 1195 | 1095 | 1202 | 658 | 641 | 591 | 669 | 458 | 404 | 413 | 404 |

| | | | | | | | | | | | | |
|---|---|---|---|---|---|---|---|---|---|---|---|---|
| K05605 3-hydroxyisobutyryl-CoA hydrolase [EC:3.1.2.4] | 0 | 2 | 0 | 0 | 1 | 1 | 0 | 0 | 0 | 0 | 0 | 0 |
| K07250 4-aminobutyrate aminotransferase / (S)-3-amino-2-methylpropionate transaminase / 5-aminovalerate transaminase [EC:2.6.1.19 2.6.1.22 2.6.1.48] | 734 | 766 | 717 | 757 | 193 | 186 | 210 | 197 | 1689 | 1785 | 1611 | 1652 |
| K07511 enoyl-CoA hydratase [EC:4.2.1.17] | 0 | 0 | 2 | 1 | 0 | 0 | 0 | 0 | 1 | 0 | 0 | 0 |
| K08325 NADP-dependent alcohol dehydrogenase [EC:1.1.-.-] | 1535 | 1559 | 1482 | 1559 | 2777 | 2916 | 2841 | 2880 | 2702 | 2590 | 2641 | 2548 |
| K09699 2-oxoisovalerate dehydrogenase E2 component (dihydrolipoyl transacylase) [EC:2.3.1.168] | 1217 | 1181 | 1058 | 1257 | 643 | 558 | 501 | 681 | 395 | 376 | 383 | 397 |
| K09788 2-methylaconitate isomerase [EC:5.3.3.-] | 1552 | 1548 | 1442 | 1392 | 775 | 686 | 824 | 896 | 492 | 507 | 444 | 429 |
| K11264 methylmalonyl-CoA decarboxylase [EC:4.1.1.-] | 0 | 0 | 0 | 0 | 11 | 8 | 3 | 3 | 2 | 3 | 0 | 0 |
| K11381 2-oxoisovalerate dehydrogenase E1 component [EC:1.2.4.4] | 1194 | 1302 | 1407 | 1331 | 4001 | 4056 | 4138 | 4040 | 2733 | 2770 | 2799 | 2600 |
| K13919 propanediol dehydratase medium subunit [EC:4.2.1.28] | 405 | 439 | 389 | 432 | 313 | 271 | 310 | 247 | 557 | 489 | 603 | 598 |
| K13920 propanediol dehydratase small subunit [EC:4.2.1.28] | 340 | 343 | 300 | 329 | 200 | 192 | 220 | 217 | 403 | 405 | 415 | 494 |
| K13921 1-propanol dehydrogenase | 728 | 733 | 727 | 778 | 435 | 485 | 432 | 457 | 958 | 906 | 970 | 955 |
| K13922 propionaldehyde dehydrogenase [EC:1.2.1.87] | 1002 | 1096 | 1023 | 1077 | 639 | 658 | 626 | 596 | 1389 | 1314 | 1349 | 1412 |
| K13923 phosphate propanoyltransferase [EC:2.3.1.222] | 321 | 407 | 274 | 364 | 239 | 305 | 236 | 267 | 447 | 483 | 438 | 482 |
| K14729 multifunctional beta-oxidation protein [EC:4.2.1.- 1.1.1.-] | 1 | 0 | 0 | 0 | 0 | 2 | 0 | 0 | 0 | 0 | 0 | 0 |
| K17489 methylmalonyl-CoA carboxyltransferase 12S subunit [EC:2.1.3.1] | 17 | 9 | 7 | 12 | 1 | 0 | 2 | 2 | 7 | 15 | 18 | 28 |
| K18369 alcohol dehydrogenase [EC:1.1.1.-] | 865 | 917 | 1009 | 960 | 1432 | 1404 | 1330 | 1328 | 1336 | 1315 | 1299 | 1395 |
| K18471 methylglyoxal reductase [EC:1.1.1.-] | 482 | 478 | 461 | 465 | 390 | 400 | 390 | 359 | 744 | 798 | 696 | 730 |
| K19697 propionate kinase [EC:2.7.2.15] | 716 | 720 | 688 | 802 | 430 | 432 | 449 | 467 | 928 | 971 | 890 | 1050 |
| K19709 acetate CoA-transferase [EC:2.8.3.8] | 85 | 78 | 77 | 59 | 118 | 133 | 115 | 135 | 205 | 172 | 191 | 216 |
| K19745 acrylyl-CoA reductase (NADPH) [EC:1.3.1.-] | 469 | 513 | 420 | 496 | 375 | 393 | 353 | 390 | 607 | 696 | 633 | 708 |
| K20143 acrylyl-CoA reductase (NADH) [EC:1.3.1.95] | 3 | 2 | 0 | 0 | 0 | 0 | 2 | 0 | 19 | 15 | 10 | 14 |
| K20455 2-methylcitrate dehydratase (2-methyl-trans-aconitate forming) [EC:4.2.1.117] | 2716 | 2810 | 2478 | 2598 | 1542 | 1344 | 1370 | 1497 | 836 | 783 | 709 | 748 |
| K20626 lactoyl-CoA dehydratase subunit alpha [EC:4.2.1.54] | 9 | 6 | 12 | 15 | 2 | 0 | 0 | 0 | 35 | 45 | 39 | 57 |
| K20627 lactoyl-CoA dehydratase subunit beta [EC:4.2.1.54] | 9 | 5 | 7 | 10 | 6 | 7 | 8 | 5 | 53 | 51 | 51 | 44 |
| K22214 propionyl-CoA:succinyl-CoA transferase [EC:2.8.3.-] | 0 | 0 | 0 | 0 | 14 | 9 | 6 | 23 | 9 | 4 | 3 | 1 |
| K23259 isopropanol dehydrogenase (NADP+) [EC:1.1.1.80] | 55 | 49 | 63 | 66 | 6 | 9 | 10 | 6 | 16 | 9 | 9 | 7 |
| ko00650 Butanoate metabolism | 135646 | 135907 | 133457 | 136759 | 114541 | 115903 | 114379 | 115720 | 128075 | 130074 | 127806 | 130251 |
| K00004 (R,R)-butanediol dehydrogenase / meso-butanediol dehydrogenase / diacetyl reductase [EC:1.1.1.4 1.1.1.- 1.1.1.303] | 3444 | 3483 | 3113 | 3297 | 1824 | 1759 | 1697 | 1874 | 1076 | 1003 | 1054 | 1109 |
| K00019 3-hydroxybutyrate dehydrogenase [EC:1.1.1.30] | 1498 | 1522 | 1433 | 1481 | 830 | 811 | 726 | 800 | 793 | 912 | 898 | 970 |
| K00022 3-hydroxyacyl-CoA dehydrogenase [EC:1.1.1.35] | 4 | 0 | 0 | 0 | 0 | 0 | 0 | 1 | 0 | 0 | 0 | 0 |
| K00043 4-hydroxybutyrate dehydrogenase [EC:1.1.1.61] | 0 | 1 | 0 | 0 | 0 | 0 | 0 | 0 | 0 | 1 | 1 | 0 |
| K00074 3-hydroxybutyryl-CoA dehydrogenase [EC:1.1.1.157] | 5128 | 5060 | 4541 | 4899 | 2552 | 2575 | 2367 | 2719 | 3579 | 3720 | 3337 | 3574 |
| K00100 butanol dehydrogenase [EC:1.1.1.-] | 1738 | 1689 | 1622 | 1580 | 834 | 913 | 861 | 940 | 597 | 586 | 619 | 681 |
| K00135 succinate-semialdehyde dehydrogenase / glutarate-semialdehyde dehydrogenase [EC:1.2.1.16 1.2.1.79 1.2.1.20] | 6810 | 6633 | 6518 | 6842 | 3726 | 3547 | 3634 | 3664 | 4741 | 4836 | 4859 | 5036 |
| K00139 succinate-semialdehyde dehydrogenase [EC:1.2.1.24] | 1 | 0 | 0 | 1 | 0 | 0 | 0 | 0 | 0 | 0 | 0 | 0 |
| K00209 enoyl-[acyl-carrier protein] reductase / trans-2-enoyl-CoA reductase (NAD+) [EC:1.3.1.9 1.3.1.44] | 1825 | 1887 | 1679 | 1778 | 986 | 959 | 898 | 1029 | 630 | 663 | 627 | 641 |
| K00634 phosphate butyryltransferase [EC:2.3.1.19] | 490 | 477 | 584 | 546 | 907 | 821 | 877 | 760 | 674 | 710 | 743 | 709 |
| K00929 butyrate kinase [EC:2.7.2.7] | 1473 | 1531 | 1662 | 1468 | 2622 | 2513 | 2600 | 2609 | 2351 | 2331 | 2333 | 2340 |
| K01027 3-oxoacid CoA-transferase [EC:2.8.3.5] | 3 | 5 | 5 | 3 | 9 | 11 | 8 | 7 | 6 | 10 | 10 | 6 |
| K01028 3-oxoacid CoA-transferase subunit A [EC:2.8.3.5] | 535 | 564 | 506 | 593 | 365 | 405 | 378 | 431 | 568 | 607 | 549 | 584 |
| K01029 3-oxoacid CoA-transferase subunit B [EC:2.8.3.5] | 562 | 526 | 450 | 542 | 313 | 320 | 347 | 333 | 470 | 568 | 497 | 534 |
| K01039 glutaconate CoA-transferase, subunit A [EC:2.8.3.12] | 639 | 643 | 562 | 667 | 357 | 371 | 367 | 379 | 216 | 174 | 199 | 208 |
| K01040 glutaconate CoA-transferase, subunit B [EC:2.8.3.12] | 1338 | 1295 | 1227 | 1187 | 814 | 755 | 702 | 720 | 438 | 431 | 358 | 417 |
| K01575 acetolactate decarboxylase [EC:4.1.1.5] | 426 | 473 | 451 | 504 | 286 | 330 | 364 | 277 | 640 | 634 | 594 | 666 |
| K01580 glutamate decarboxylase [EC:4.1.1.15] | 1039 | 1071 | 1083 | 1066 | 3318 | 3331 | 3323 | 3374 | 2576 | 2482 | 2624 | 2516 |
| K01615 glutaconyl-CoA decarboxylase subunit alpha [EC:7.2.4.5] | 25 | 35 | 36 | 30 | 67 | 79 | 76 | 89 | 26 | 37 | 40 | 36 |
| K01640 hydroxymethylglutaryl-CoA lyase [EC:4.1.3.4] | 1325 | 1249 | 1153 | 1244 | 662 | 657 | 638 | 652 | 447 | 425 | 397 | 373 |
| K01641 hydroxymethylglutaryl-CoA synthase [EC:2.3.3.10] | 56 | 93 | 87 | 56 | 19 | 21 | 27 | 20 | 70 | 113 | 81 | 89 |
| K01652 acetolactate synthase I/II/III large subunit [EC:2.2.1.6] | 16148 | 16395 | 15882 | 16536 | 11375 | 11742 | 11289 | 11370 | 13807 | 14064 | 13638 | 14040 |
| K01653 acetolactate synthase I/III small subunit [EC:2.2.1.6] | 2139 | 2147 | 2225 | 2167 | 2163 | 2234 | 2198 | 2149 | 2281 | 2397 | 2291 | 2373 |
| K01715 enoyl-CoA hydratase [EC:4.2.1.17] | 178 | 202 | 190 | 225 | 132 | 114 | 114 | 119 | 260 | 357 | 355 | 361 |
| K01799 maleate isomerase [EC:5.2.1.1] | 3 | 2 | 2 | 3 | 0 | 0 | 0 | 0 | 3 | 0 | 2 | 1 |
| K01907 acetoacetyl-CoA synthetase [EC:6.2.1.16] | 2504 | 2541 | 2409 | 2385 | 1287 | 1314 | 1136 | 1379 | 796 | 745 | 688 | 726 |
| K03366 meso-butanediol dehydrogenase / (S,S)-butanediol dehydrogenase / diacetyl reductase [EC:1.1.1.- 1.1.1.76 1.1.1.304] | 868 | 834 | 869 | 857 | 771 | 899 | 840 | 778 | 576 | 616 | 578 | 692 |
| K03821 polyhydroxyalkanoate synthase subunit PhaC [EC:2.3.1.-] | 5497 | 5392 | 5402 | 5410 | 2994 | 2931 | 2851 | 3101 | 1824 | 1632 | 1603 | 1702 |
| K07516 3-hydroxyacyl-CoA dehydrogenase [EC:1.1.1.35] | 11 | 3 | 5 | 1 | 16 | 13 | 24 | 20 | 1 | 4 | 2 | 7 |
| K07518 hydroxybutyrate-dimer hydrolase [EC:3.1.1.22] | 0 | 0 | 0 | 0 | 0 | 0 | 0 | 0 | 0 | 2 | 0 | 0 |
| K08318 4-hydroxybutyrate dehydrogenase / sulfolactaldehyde 3-reductase [EC:1.1.1.61 1.1.1.373] | 263 | 248 | 227 | 235 | 348 | 395 | 364 | 352 | 66 | 67 | 98 | 79 |
| K08324 succinate-semialdehyde dehydrogenase [EC:1.2.1.16 1.2.1.24] | 815 | 685 | 639 | 827 | 553 | 589 | 521 | 529 | 1000 | 1118 | 985 | 1134 |
| K11258 acetolactate synthase II small subunit [EC:2.2.1.6] | 179 | 151 | 146 | 175 | 122 | 133 | 119 | 115 | 220 | 198 | 173 | 230 |
| K14268 5-aminovalerate/4-aminobutyrate aminotransferase [EC:2.6.1.48 2.6.1.19] | 2343 | 2184 | 2110 | 2276 | 1241 | 1257 | 1264 | 1354 | 679 | 688 | 645 | 610 |
| K14534 4-hydroxybutyryl-CoA dehydratase / vinylacetyl-CoA-Delta-isomerase [EC:4.2.1.120 5.3.3.3] | 780 | 723 | 818 | 827 | 127 | 121 | 133 | 109 | 1441 | 1486 | 1568 | 1619 |
| K16871 4-aminobutyrate---pyruvate transaminase [EC:2.6.1.96] | 14 | 17 | 16 | 27 | 0 | 2 | 2 | 0 | 12 | 12 | 15 | 8 |
| K18009 meso-butanediol dehydrogenase / (S,S)-butanediol dehydrogenase / diacetyl reductase [EC:1.1.1.- 1.1.1.76 1.1.1.304] | 1 | 2 | 2 | 3 | 2 | 4 | 3 | 3 | 2 | 1 | 1 | 0 |
| K18119 succinate-semialdehyde dehydrogenase [EC:1.2.1.76] | 3 | 12 | 2 | 8 | 1 | 0 | 2 | 0 | 0 | 3 | 0 | 7 |
| K18120 4-hydroxybutyrate dehydrogenase [EC:1.1.1.61] | 81 | 53 | 56 | 56 | 19 | 20 | 15 | 16 | 17 | 23 | 38 | 34 |
| K18122 4-hydroxybutyrate CoA-transferase [EC:2.8.3.-] | 65 | 72 | 72 | 74 | 35 | 49 | 29 | 36 | 39 | 55 | 47 | 45 |
| K23351 glutaconyl-CoA/methylmalonyl-CoA decarboxylase subunit gamma | 417 | 444 | 456 | 432 | 963 | 903 | 899 | 908 | 721 | 843 | 788 | 670 |
| K23352 glutaconyl-CoA/methylmalonyl-CoA decarboxylase subunit delta | 198 | 251 | 239 | 213 | 49 | 49 | 43 | 53 | 175 | 249 | 223 | 201 |
| ko00660 C5-Branched dibasic acid metabolism | 37781 | 38036 | 37586 | 38769 | 35226 | 35582 | 34868 | 34361 | 36217 | 36500 | 35721 | 36542 |
| K00052 3-isopropylmalate dehydrogenase [EC:1.1.1.85] | 2570 | 2661 | 2578 | 2686 | 3458 | 3326 | 3436 | 3182 | 3374 | 3477 | 3304 | 3389 |
| K01703 3-isopropylmalate/(R)-2-methylmalate dehydratase large subunit [EC:4.2.1.33 4.2.1.35] | 4521 | 4354 | 4339 | 4552 | 4913 | 5049 | 4994 | 4747 | 4882 | 4859 | 4710 | 4855 |
| K01704 3-isopropylmalate/(R)-2-methylmalate dehydratase small subunit [EC:4.2.1.33 4.2.1.35] | 1899 | 1885 | 1875 | 1955 | 2206 | 2229 | 2088 | 2223 | 1894 | 2033 | 2112 | 1963 |
| K09011 (R)-citramalate synthase [EC:2.3.1.182] | 751 | 807 | 800 | 766 | 2452 | 2524 | 2365 | 2523 | 1770 | 1703 | 1705 | 1687 |
| K10218 4-hydroxy-4-methyl-2-oxoglutarate aldolase [EC:4.1.3.17] | 301 | 329 | 292 | 358 | 106 | 95 | 87 | 89 | 633 | 605 | 623 | 651 |
| K18288 itaconate CoA-transferase [EC:2.8.3.-] | 3 | 2 | 1 | 0 | 0 | 0 | 1 | 1 | 3 | 4 | 3 | 6 |
| K18289 itaconate CoA-transferase [EC:2.8.3.- 2.8.3.22] | 1841 | 1785 | 1804 | 1878 | 1022 | 952 | 875 | 974 | 537 | 481 | 526 | 574 |
| K18291 itaconyl-CoA hydratase / mesaconyl-C4 CoA hydratase [EC:4.2.1.56 4.2.1.-] | 1174 | 1149 | 1161 | 1190 | 619 | 628 | 637 | 631 | 392 | 346 | 323 | 373 |
| K18292 (S)-citramalyl-CoA lyase [EC:4.1.3.25] | 593 | 657 | 638 | 686 | 385 | 395 | 352 | 357 | 225 | 216 | 174 | 199 |
| K22004 trans-aconitate decarbocylase [EC:4.1.1.-] | 0 | 0 | 0 | 0 | 0 | 0 | 0 | 1 | 0 | 0 | 0 | 0 |
| ko00562 Inositol phosphate metabolism | 25821 | 26006 | 25548 | 26338 | 17182 | 17350 | 16758 | 17402 | 27334 | 28147 | 27205 | 28180 |
| K00010 myo-inositol 2-dehydrogenase / D-chiro-inositol 1-dehydrogenase [EC:1.1.1.18 1.1.1.369] | 1408 | 1350 | 1304 | 1378 | 282 | 287 | 299 | 281 | 2542 | 2713 | 2549 | 2702 |
| K00889 1-phosphatidylinositol-4-phosphate 5-kinase [EC:2.7.1.68] | 19 | 18 | 24 | 22 | 75 | 67 | 0 | 0 | 47 | 57 | 69 | 43 |
| K01092 myo-inositol-1(or 4)-monophosphatase [EC:3.1.3.25] | 2188 | 2392 | 2374 | 2397 | 2796 | 2862 | 2851 | 2860 | 3783 | 3738 | 3833 | 3876 |
| K01093 4-phytase / acid phosphatase [EC:3.1.3.26 3.1.3.2] | 432 | 537 | 445 | 505 | 153 | 125 | 115 | 109 | 1025 | 1067 | 985 | 1022 |
| K01114 phospholipase C [EC:3.1.4.3] | 6513 | 6572 | 6098 | 6453 | 3388 | 3540 | 3352 | 3634 | 2143 | 1954 | 1808 | 2070 |
| K01771 1-phosphatidylinositol phosphodiesterase [EC:4.6.1.13] | 2 | 0 | 2 | 1 | 1 | 5 | 11 | 17 | 19 | 11 | 12 | 7 |
| K01858 myo-inositol-1-phosphate synthase [EC:5.5.1.4] | 804 | 815 | 981 | 800 | 2615 | 2637 | 2548 | 2656 | 1886 | 1832 | 1970 | 1759 |
| K03335 inosose dehydratase [EC:4.2.1.44] | 1403 | 1322 | 1315 | 1416 | 355 | 330 | 291 | 352 | 2619 | 2923 | 2752 | 2996 |
| K03336 3D-(3,5/4)-trihydroxycyclohexane-1,2-dione acylhydrolase (decyclizing) [EC:3.7.1.22] | 1345 | 1287 | 1237 | 1319 | 292 | 276 | 260 | 258 | 2242 | 2401 | 2196 | 2405 |
| K03337 5-deoxy-glucuronate isomerase [EC:5.3.1.30] | 506 | 465 | 474 | 531 | 123 | 107 | 117 | 89 | 838 | 901 | 826 | 916 |
| K03338 5-dehydro-2-deoxygluconokinase [EC:2.7.1.92] | 1003 | 1010 | 1025 | 1075 | 268 | 279 | 257 | 240 | 1908 | 2041 | 1992 | 2085 |
| K03339 6-phospho-5-dehydro-2-deoxy-D-gluconate aldolase [EC:4.1.2.29] | 2 | 0 | 0 | 0 | 5 | 2 | 1 | 0 | 0 | 0 | 0 | 0 |
| K06606 2-keto-myo-inositol isomerase [EC:5.3.99.11] | 319 | 371 | 357 | 386 | 88 | 96 | 92 | 99 | 792 | 870 | 739 | 791 |
| K13711 phosphatidylinositol 4-kinase type 2 [EC:2.7.1.67] | 0 | 0 | 0 | 0 | 0 | 0 | 0 | 0 | 0 | 1 | 0 | 0 |
| K16043 scyllo-inositol 2-dehydrogenase (NAD+) [EC:1.1.1.370] | 1 | 1 | 3 | 1 | 0 | 0 | 0 | 0 | 12 | 7 | 10 | 12 |
| K16044 scyllo-inositol 2-dehydrogenase (NADP+) [EC:1.1.1.371] | 0 | 0 | 2 | 0 | 7 | 5 | 9 | 3 | 3 | 0 | 0 | 0 |
| K19517 1D-myo-inositol 3-kinase [EC:2.7.1.64] | 0 | 0 | 0 | 0 | 0 | 0 | 1 | 0 | 2 | 0 | 0 | 0 |
| K20279 synaptojanin [EC:3.1.3.36] | 0 | 0 | 0 | 0 | 0 | 0 | 0 | 0 | 0 | 0 | 2 | 0 |
| K22230 scyllo-inositol 2-dehydrogenase (NADP+) [EC:1.1.1.-] | 274 | 263 | 287 | 270 | 78 | 63 | 59 | 79 | 162 | 180 | 183 | 171 |
| K22232 3-dehydro-scyllo-inosose hydrolase [EC:3.7.1.-] | 4 | 3 | 2 | 2 | 3 | 2 | 4 | 5 | 9 | 10 | 12 | 1 |
| K22233 5-keto-L-gluconate epimerase [EC:5.1.-.-] | 3 | 2 | 2 | 3 | 0 | 0 | 0 | 0 | 3 | 5 | 5 | 2 |
| Energy metabolism | 609856 | 609815 | 605232 | 611512 | 560444 | 566647 | 558081 | 560060 | 560446 | 560068 | 556434 | 559554 |
| ko00190 Oxidative phosphorylation | 145690 | 146078 | 147707 | 148635 | 137554 | 140073 | 137205 | 137298 | 132458 | 132786 | 132346 | 132247 |
| K00330 NADH-quinone oxidoreductase subunit A [EC:7.1.1.2] | 1433 | 1445 | 1509 | 1515 | 1399 | 1425 | 1357 | 1446 | 1177 | 1089 | 1165 | 1146 |
| K00331 NADH-quinone oxidoreductase subunit B [EC:7.1.1.2] | 1446 | 1366 | 1571 | 1441 | 1930 | 2135 | 2012 | 1793 | 1664 | 1547 | 1612 | 1580 |
| K00332 NADH-quinone oxidoreductase subunit C [EC:7.1.1.2] | 3 | 0 | 2 | 1 | 0 | 0 | 0 | 3 | 0 | 5 | 4 | 5 |
| K00333 NADH-quinone oxidoreductase subunit D [EC:7.1.1.2] | 4 | 5 | 5 | 9 | 6 | 0 | 3 | 3 | 13 | 8 | 6 | 12 |
| K00334 NADH-quinone oxidoreductase subunit E [EC:7.1.1.2] | 1153 | 1088 | 1127 | 1214 | 624 | 648 | 609 | 596 | 889 | 1012 | 974 | 1041 |
| K00335 NADH-quinone oxidoreductase subunit F [EC:7.1.1.2] | 4128 | 4299 | 4308 | 4210 | 1920 | 1947 | 1939 | 2061 | 3183 | 3392 | 3218 | 3386 |
| K00336 NADH-quinone oxidoreductase subunit G [EC:7.1.1.2] | 6013 | 6018 | 5860 | 6234 | 3570 | 3483 | 3245 | 3546 | 4424 | 4570 | 4417 | 4514 |
| K00337 NADH-quinone oxidoreductase subunit H [EC:7.1.1.2] | 2556 | 2557 | 2583 | 2712 | 3415 | 3447 | 3372 | 3358 | 2974 | 2834 | 2864 | 2736 |
| K00338 NADH-quinone oxidoreductase subunit I [EC:7.1.1.2] | 1383 | 1360 | 1410 | 1480 | 1713 | 1763 | 1582 | 1600 | 1482 | 1406 | 1451 | 1448 |
| K00339 NADH-quinone oxidoreductase subunit J [EC:7.1.1.2] | 1086 | 1116 | 1091 | 1124 | 1593 | 1603 | 1651 | 1537 | 1471 | 1293 | 1388 | 1387 |
| K00340 NADH-quinone oxidoreductase subunit K [EC:7.1.1.2] | 695 | 695 | 677 | 775 | 930 | 936 | 940 | 1005 | 826 | 775 | 802 | 743 |
| K00341 NADH-quinone oxidoreductase subunit L [EC:7.1.1.2] | 4976 | 4931 | 5089 | 5096 | 6336 | 6697 | 6358 | 6523 | 5528 | 5094 | 5373 | 5320 |
| K00342 NADH-quinone oxidoreductase subunit M [EC:7.1.1.2] | 3813 | 4033 | 4058 | 3920 | 4784 | 4774 | 4475 | 4797 | 4009 | 4056 | 4065 | 3896 |
| K00343 NADH-quinone oxidoreductase subunit N [EC:7.1.1.2] | 3847 | 3828 | 3915 | 3895 | 4945 | 4961 | 4795 | 4726 | 4168 | 4039 | 4024 | 3935 |
| K00356 NADH dehydrogenase [EC:1.6.99.3] | 1 | 2 | 1 | 0 | 0 | 0 | 2 | 3 | 4 | 1 | 2 | 4 |
| K00404 cytochrome c oxidase cbb3-type subunit I [EC:1.9.3.1] | 7457 | 7450 | 7363 | 7317 | 4004 | 3861 | 3840 | 4044 | 2358 | 2173 | 2352 | 2334 |
| K00405 cytochrome c oxidase cbb3-type subunit II | 987 | 1037 | 987 | 1061 | 601 | 503 | 561 | 567 | 378 | 343 | 328 | 352 |

| | | | | | | | | | | | | |
|---|---|---|---|---|---|---|---|---|---|---|---|---|
| K00406 cytochrome c oxidase cbb3-type subunit III | 1334 | 1269 | 1349 | 1174 | 697 | 688 | 604 | 624 | 429 | 413 | 352 | 375 |
| K00407 cytochrome c oxidase cbb3-type subunit IV | 513 | 481 | 484 | 504 | 268 | 283 | 262 | 264 | 164 | 156 | 165 | 146 |
| K00410 ubiquinol-cytochrome c reductase cytochrome b/c1 subunit | 0 | 0 | 0 | 0 | 0 | 1 | 0 | 0 | 0 | 0 | 0 | 0 |
| K00411 ubiquinol-cytochrome c reductase iron-sulfur subunit [EC:7.1.1.8] | 810 | 772 | 828 | 843 | 548 | 545 | 478 | 471 | 310 | 295 | 276 | 259 |
| K00412 ubiquinol-cytochrome c reductase cytochrome b subunit | 1861 | 1865 | 1926 | 1879 | 1034 | 1152 | 1052 | 1024 | 620 | 583 | 604 | 575 |
| K00413 ubiquinol-cytochrome c reductase cytochrome c1 subunit | 997 | 1054 | 1090 | 1087 | 591 | 600 | 575 | 557 | 331 | 378 | 288 | 312 |
| K00424 cytochrome bd-I ubiquinol oxidase subunit X [EC:7.1.1.7] | 81 | 50 | 56 | 68 | 53 | 50 | 63 | 43 | 85 | 105 | 98 | 86 |
| K00425 cytochrome bd ubiquinol oxidase subunit I [EC:7.1.1.7] | 5480 | 5553 | 5511 | 5510 | 5784 | 5813 | 5934 | 5746 | 7072 | 7205 | 7038 | 7319 |
| K00426 cytochrome bd ubiquinol oxidase subunit II [EC:7.1.1.7] | 2959 | 2867 | 2991 | 3017 | 3849 | 3798 | 3878 | 3585 | 4715 | 4781 | 4544 | 4698 |
| K00937 polyphosphate kinase [EC:2.7.4.1] | 6997 | 7138 | 7219 | 7038 | 6878 | 6880 | 7000 | 6953 | 6127 | 5929 | 6021 | 6240 |
| K01507 inorganic pyrophosphatase [EC:3.6.1.1] | 1496 | 1529 | 1406 | 1505 | 849 | 861 | 863 | 864 | 1326 | 1449 | 1351 | 1374 |
| K01535 H+-transporting ATPase [EC:7.1.2.1] | 1 | 5 | 0 | 1 | 0 | 0 | 0 | 0 | 0 | 0 | 0 | 0 |
| K02107 V/A-type H+/Na+-transporting ATPase subunit G/H | 21 | 20 | 16 | 22 | 5 | 8 | 3 | 1 | 21 | 30 | 24 | 23 |
| K02108 F-type H+-transporting ATPase subunit a | 3450 | 3448 | 3579 | 3489 | 3547 | 3695 | 3655 | 3531 | 3459 | 3383 | 3469 | 3219 |
| K02109 F-type H+-transporting ATPase subunit b | 1309 | 1405 | 1420 | 1390 | 1419 | 1309 | 1334 | 1293 | 1368 | 1222 | 1300 | 1215 |
| K02110 F-type H+-transporting ATPase subunit c | 912 | 941 | 961 | 977 | 765 | 765 | 746 | 710 | 810 | 801 | 820 | 785 |
| K02111 F-type H+/Na+-transporting ATPase subunit alpha [EC:7.1.2.2 7.2.2.1] | 6594 | 6558 | 6528 | 6717 | 5992 | 5937 | 6007 | 6005 | 6024 | 5856 | 6208 | 6156 |
| K02112 F-type H+/Na+-transporting ATPase subunit beta [EC:7.1.2.2 7.2.2.1] | 5890 | 5915 | 6324 | 6109 | 5721 | 5799 | 5765 | 5756 | 5414 | 5731 | 5557 | 5672 |
| K02113 F-type H+-transporting ATPase subunit delta | 2032 | 2047 | 2064 | 2153 | 2016 | 2074 | 2171 | 1973 | 1919 | 2025 | 1985 | 1931 |
| K02114 F-type H+-transporting ATPase subunit epsilon | 1484 | 1561 | 1664 | 1533 | 1374 | 1311 | 1162 | 1234 | 1277 | 1285 | 1287 | 1235 |
| K02115 F-type H+-transporting ATPase subunit gamma | 3407 | 3367 | 3468 | 3463 | 3059 | 3303 | 3183 | 3135 | 3288 | 3359 | 3264 | 3258 |
| K02117 V/A-type H+/Na+-transporting ATPase subunit A [EC:7.1.2.2 7.2.2.1] | 2616 | 2519 | 2787 | 2696 | 3612 | 3909 | 3931 | 3644 | 3812 | 3941 | 3968 | 3900 |
| K02118 V/A-type H+/Na+-transporting ATPase subunit B | 2186 | 2117 | 2181 | 2127 | 3166 | 3183 | 3224 | 3030 | 3171 | 3295 | 3139 | 3056 |
| K02119 V/A-type H+/Na+-transporting ATPase subunit C | 849 | 753 | 836 | 813 | 203 | 208 | 248 | 200 | 712 | 759 | 663 | 800 |
| K02120 V/A-type H+/Na+-transporting ATPase subunit D | 976 | 993 | 980 | 946 | 1307 | 1235 | 1259 | 1374 | 1341 | 1334 | 1450 | 1248 |
| K02121 V/A-type H+/Na+-transporting ATPase subunit E | 340 | 425 | 433 | 411 | 998 | 1052 | 1048 | 1038 | 733 | 780 | 854 | 810 |
| K02122 V/A-type H+/Na+-transporting ATPase subunit F | 315 | 315 | 287 | 303 | 85 | 81 | 95 | 76 | 321 | 272 | 298 | 284 |
| K02123 V/A-type H+/Na+-transporting ATPase subunit I | 2506 | 2549 | 2727 | 2650 | 4058 | 4216 | 4044 | 3975 | 3810 | 3912 | 3938 | 3944 |
| K02124 V/A-type H+/Na+-transporting ATPase subunit K | 599 | 600 | 591 | 594 | 1049 | 1056 | 1043 | 999 | 890 | 932 | 978 | 915 |
| K02126 F-type H+-transporting ATPase subunit a | 0 | 2 | 0 | 0 | 0 | 0 | 0 | 0 | 0 | 0 | 0 | 0 |
| K02128 F-type H+-transporting ATPase subunit c | 0 | 2 | 0 | 0 | 0 | 0 | 0 | 0 | 0 | 0 | 0 | 0 |
| K02132 F-type H+-transporting ATPase subunit alpha | 3 | 3 | 11 | 6 | 1 | 2 | 1 | 0 | 12 | 6 | 15 | 5 |
| K02133 F-type H+-transporting ATPase subunit beta [EC:7.1.2.2] | 2 | 1 | 0 | 1 | 3 | 3 | 3 | 1 | 0 | 1 | 3 | 2 |
| K02145 V-type H+-transporting ATPase subunit A [EC:7.1.2.2] | 0 | 0 | 0 | 0 | 0 | 1 | 0 | 0 | 0 | 0 | 0 | 0 |
| K02147 V-type H+-transporting ATPase subunit B | 0 | 0 | 0 | 1 | 0 | 0 | 0 | 0 | 0 | 0 | 0 | 0 |
| K02256 cytochrome c oxidase subunit 1 [EC:1.9.3.1] | 0 | 0 | 0 | 0 | 0 | 1 | 3 | 0 | 2 | 0 | 0 | 0 |
| K02257 heme o synthase [EC:2.5.1.141] | 2834 | 2634 | 2610 | 2714 | 1595 | 1628 | 1539 | 1675 | 1489 | 1512 | 1339 | 1572 |
| K02258 cytochrome c oxidase assembly protein subunit 11 | 819 | 864 | 905 | 885 | 450 | 447 | 414 | 486 | 280 | 316 | 276 | 261 |
| K02259 cytochrome c oxidase assembly protein subunit 15 | 1687 | 1642 | 1668 | 1596 | 903 | 885 | 761 | 802 | 541 | 555 | 555 | 480 |
| K02262 cytochrome c oxidase subunit 3 | 0 | 0 | 0 | 0 | 1 | 0 | 0 | 0 | 0 | 0 | 0 | 0 |
| K02274 cytochrome c oxidase subunit I [EC:1.9.3.1] | 2751 | 2830 | 2831 | 2868 | 1674 | 1633 | 1516 | 1621 | 1028 | 994 | 907 | 909 |
| K02275 cytochrome c oxidase subunit II [EC:1.9.3.1] | 1468 | 1566 | 1355 | 1422 | 842 | 777 | 760 | 861 | 545 | 460 | 434 | 477 |
| K02276 cytochrome c oxidase subunit III [EC:1.9.3.1] | 1618 | 1508 | 1462 | 1537 | 806 | 874 | 765 | 859 | 529 | 459 | 504 | 503 |
| K02277 cytochrome c oxidase subunit IV [EC:1.9.3.1] | 1 | 0 | 0 | 0 | 0 | 1 | 0 | 0 | 0 | 0 | 0 | 0 |
| K02297 cytochrome o ubiquinol oxidase subunit II [EC:7.1.1.3] | 1168 | 1118 | 1171 | 1138 | 730 | 721 | 740 | 782 | 976 | 1086 | 992 | 1022 |
| K02298 cytochrome o ubiquinol oxidase subunit I [EC:7.1.1.3] | 4065 | 3975 | 4072 | 4192 | 2380 | 2492 | 2400 | 2439 | 2624 | 2635 | 2616 | 2544 |
| K02299 cytochrome o ubiquinol oxidase subunit III | 1231 | 1290 | 1277 | 1318 | 910 | 803 | 705 | 787 | 844 | 858 | 780 | 840 |
| K02300 cytochrome o ubiquinol oxidase subunit IV | 510 | 518 | 534 | 542 | 288 | 363 | 352 | 334 | 356 | 401 | 347 | 375 |
| K02826 cytochrome aa3-600 menaquinol oxidase subunit II [EC:7.1.1.5] | 0 | 0 | 0 | 0 | 0 | 0 | 0 | 2 | 0 | 0 | 0 | 0 |
| K02827 cytochrome aa3-600 menaquinol oxidase subunit I [EC:7.1.1.5] | 1 | 0 | 0 | 0 | 0 | 0 | 0 | 0 | 0 | 0 | 1 | 0 |
| K02828 cytochrome aa3-600 menaquinol oxidase subunit III [EC:7.1.1.5] | 1 | 0 | 0 | 1 | 0 | 0 | 0 | 1 | 0 | 0 | 0 | 0 |
| K02829 cytochrome aa3-600 menaquinol oxidase subunit IV [EC:7.1.1.5] | 0 | 0 | 0 | 1 | 0 | 0 | 0 | 1 | 0 | 0 | 0 | 0 |
| K03878 NADH-ubiquinone oxidoreductase chain 1 [EC:7.1.1.2] | 0 | 0 | 2 | 0 | 2 | 0 | 0 | 0 | 0 | 0 | 0 | 0 |
| K03881 NADH-ubiquinone oxidoreductase chain 4 [EC:7.1.1.2] | 0 | 0 | 0 | 0 | 0 | 0 | 0 | 0 | 0 | 0 | 1 | 0 |
| K03883 NADH-ubiquinone oxidoreductase chain 5 [EC:7.1.1.2] | 0 | 0 | 0 | 0 | 0 | 2 | 3 | 0 | 2 | 1 | 0 | 0 |
| K03885 NADH dehydrogenase [EC:1.6.99.3] | 5318 | 5498 | 5154 | 5339 | 5243 | 5237 | 5092 | 5163 | 4225 | 4045 | 4222 | 4154 |
| K03888 menaquinol-cytochrome c reductase cytochrome b/c subunit | 2 | 0 | 1 | 0 | 0 | 0 | 0 | 0 | 0 | 0 | 0 | 1 |
| K03934 NADH dehydrogenase (ubiquinone) Fe-S protein 1 [EC:7.1.1.2 1.6.99.3] | 0 | 0 | 0 | 0 | 0 | 0 | 0 | 0 | 0 | 0 | 0 | 3 |
| K05576 NAD(P)H-quinone oxidoreductase subunit 4L [EC:7.1.1.2] | 0 | 0 | 0 | 0 | 1 | 1 | 0 | 0 | 0 | 1 | 0 | 1 |
| K05577 NAD(P)H-quinone oxidoreductase subunit 5 [EC:7.1.1.2] | 0 | 1 | 0 | 0 | 0 | 0 | 0 | 0 | 0 | 0 | 0 | 0 |
| K05587 bidirectional [NiFe] hydrogenase diaphorase subunit [EC:7.1.1.2] | 0 | 0 | 0 | 1 | 1 | 0 | 0 | 0 | 0 | 0 | 1 | 1 |
| K05588 bidirectional [NiFe] hydrogenase diaphorase subunit [EC:7.1.1.2] | 1 | 2 | 1 | 1 | 0 | 0 | 0 | 0 | 0 | 0 | 0 | 0 |
| K06019 pyrophosphatase PpaX [EC:3.6.1.1] | 23 | 31 | 19 | 22 | 6 | 3 | 2 | 0 | 37 | 41 | 52 | 42 |
| K13378 NADH-quinone oxidoreductase subunit C/D [EC:7.1.1.2] | 4220 | 4253 | 4392 | 4509 | 5441 | 5591 | 5568 | 5468 | 4731 | 4770 | 4478 | 4604 |
| K15862 cytochrome c oxidase cbb3-type subunit I/II [EC:1.9.3.1] | 2 | 0 | 1 | 0 | 1 | 0 | 2 | 0 | 1 | 4 | 2 | 1 |
| K15986 manganese-dependent inorganic pyrophosphatase [EC:3.6.1.1] | 1275 | 1326 | 1365 | 1326 | 707 | 765 | 793 | 718 | 907 | 1137 | 995 | 1072 |
| K22468 polyphosphate kinase [EC:2.7.4.1] | 2648 | 2550 | 2396 | 2627 | 1493 | 1630 | 1413 | 1538 | 1347 | 1302 | 1383 | 1352 |
| K22501 cytochrome bd-II ubiquinol oxidase subunit AppX [EC:7.1.1.7] | 0 | 2 | 0 | 0 | 3 | 0 | 0 | 3 | 4 | 3 | 2 | 4 |
| ko00195 Photosynthesis | 25078 | 25242 | 26008 | 25831 | 23893 | 24193 | 24023 | 23637 | 23559 | 23662 | 23890 | 23471 |
| ko00710 Carbon fixation in photosynthetic organisms | 70353 | 70765 | 71165 | 70998 | 72378 | 73942 | 73301 | 71665 | 77711 | 78355 | 77439 | 77675 |
| K00855 phosphoribulokinase [EC:2.7.1.19] | 539 | 523 | 533 | 525 | 354 | 428 | 394 | 378 | 657 | 756 | 667 | 764 |
| K11214 sedoheptulokinase [EC:2.7.1.14] | 1 | 0 | 2 | 2 | 0 | 0 | 0 | 0 | 7 | 3 | 0 | 4 |
| K14454 aspartate aminotransferase, cytoplasmic [EC:2.6.1.1] | 0 | 1 | 0 | 0 | 0 | 0 | 0 | 0 | 0 | 0 | 0 | 0 |
| K14455 aspartate aminotransferase, mitochondrial [EC:2.6.1.1] | 0 | 0 | 0 | 0 | 1 | 0 | 2 | 0 | 0 | 1 | 0 | 0 |
| ko00720 Carbon fixation pathways in prokaryotes | 143254 | 143346 | 143547 | 144074 | 155127 | 157183 | 156797 | 156520 | 153867 | 154439 | 154047 | 153805 |
| K00194 acetyl-CoA decarbonylase/synthase, CODH/ACS complex subunit delta [EC:2.1.1.245] | 24 | 36 | 29 | 54 | 19 | 31 | 29 | 36 | 90 | 111 | 90 | 114 |
| K00196 anaerobic carbon-monoxide dehydrogenase iron sulfur subunit | 87 | 78 | 68 | 86 | 37 | 41 | 119 | 128 | 64 | 68 | 76 | 76 |
| K00197 acetyl-CoA decarbonylase/synthase, CODH/ACS complex subunit gamma [EC:2.1.1.245] | 76 | 49 | 43 | 42 | 59 | 42 | 35 | 53 | 127 | 177 | 130 | 151 |
| K00198 anaerobic carbon-monoxide dehydrogenase catalytic subunit [EC:1.2.7.4] | 317 | 304 | 321 | 328 | 101 | 93 | 91 | 80 | 609 | 649 | 581 | 565 |
| K00297 methylenetetrahydrofolate reductase (NADPH) [EC:1.5.1.20] | 3363 | 3383 | 3519 | 3382 | 3530 | 3539 | 3691 | 3747 | 3378 | 3132 | 3176 | 3155 |
| K01491 methylenetetrahydrofolate dehydrogenase (NADP+) / methenyltetrahydrofolate cyclohydrolase [EC:1.5.1.5 3.5.4.9] | 2495 | 2524 | 2501 | 2444 | 2967 | 2892 | 2862 | 2850 | 2795 | 2679 | 2741 | 2709 |
| K01938 formate--tetrahydrofolate ligase [EC:6.3.4.3] | 1952 | 1977 | 2051 | 1957 | 3553 | 3585 | 3732 | 3511 | 3430 | 3406 | 3202 | 3300 |
| K05299 formate dehydrogenase (NADP+) alpha subunit [EC:1.17.1.10] | 9 | 9 | 11 | 8 | 4 | 6 | 6 | 8 | 16 | 19 | 13 | 13 |
| K14138 acetyl-CoA synthase [EC:2.3.1.169] | 101 | 83 | 87 | 102 | 52 | 52 | 56 | 48 | 228 | 288 | 230 | 258 |
| K14470 2-methylfumaryl-CoA isomerase [EC:5.4.1.3] | 0 | 0 | 0 | 1 | 0 | 0 | 0 | 0 | 1 | 0 | 1 | 0 |
| K15016 enoyl-CoA hydratase / 3-hydroxyacyl-CoA dehydrogenase [EC:4.2.1.17 1.1.1.35] | 5 | 2 | 2 | 1 | 1 | 1 | 1 | 0 | 2 | 5 | 6 | 6 |
| K15019 3-hydroxypropionyl-coenzyme A dehydratase [EC:4.2.1.116] | 0 | 2 | 2 | 7 | 0 | 3 | 2 | 1 | 1 | 3 | 2 | 1 |
| K15020 acryloyl-coenzyme A reductase [EC:1.3.1.84] | 0 | 0 | 0 | 0 | 0 | 0 | 0 | 1 | 0 | 0 | 0 | 0 |
| K15022 formate dehydrogenase (NADP+) beta subunit [EC:1.17.1.10] | 76 | 72 | 79 | 79 | 119 | 91 | 95 | 96 | 106 | 130 | 127 | 129 |
| K15023 5-methyltetrahydrofolate corrinoid/iron sulfur protein methyltransferase [EC:2.1.1.258] | 43 | 29 | 44 | 35 | 35 | 36 | 29 | 28 | 90 | 91 | 102 | 87 |
| K15039 3-hydroxypropionate dehydrogenase (NADP+) [EC:1.1.1.298] | 4 | 2 | 2 | 0 | 0 | 0 | 0 | 0 | 0 | 0 | 1 | 0 |
| K18209 fumarate reductase (CoM/CoB) subunit A [EC:1.3.4.1] | 34 | 34 | 26 | 39 | 3 | 9 | 5 | 9 | 63 | 54 | 39 | 60 |
| K18210 fumarate reductase (CoM/CoB) subunit B [EC:1.3.4.1] | 0 | 3 | 2 | 0 | 3 | 2 | 3 | 0 | 7 | 9 | 3 | 23 |
| K18556 NADH-dependent fumarate reductase subunit A [EC:1.3.1.6] | 0 | 0 | 0 | 0 | 0 | 0 | 0 | 1 | 0 | 0 | 0 | 0 |
| K18559 NADH-dependent fumarate reductase subunit D | 0 | 0 | 0 | 0 | 0 | 0 | 0 | 2 | 0 | 0 | 0 | 0 |
| ko00680 Methane metabolism | 103508 | 103110 | 102439 | 103040 | 99764 | 102035 | 100002 | 99858 | 104726 | 104681 | 103420 | 104820 |
| K00058 D-3-phosphoglycerate dehydrogenase / 2-oxoglutarate reductase [EC:1.1.1.95 1.1.1.399] | 7254 | 6707 | 6997 | 6996 | 5036 | 5169 | 4978 | 4913 | 7449 | 7651 | 7393 | 7411 |
| K00125 formate dehydrogenase (coenzyme F420) beta subunit [EC:1.17.98.3 1.8.98.6] | 0 | 2 | 0 | 5 | 0 | 0 | 2 | 0 | 13 | 10 | 13 | 12 |
| K00148 glutathione-independent formaldehyde dehydrogenase [EC:1.2.1.46] | 2130 | 2040 | 2041 | 2117 | 1154 | 1053 | 1063 | 1161 | 693 | 627 | 630 | 710 |
| K00200 formylmethanofuran dehydrogenase subunit A [EC:1.2.7.12] | 2 | 1 | 3 | 1 | 0 | 0 | 2 | 0 | 8 | 17 | 13 | 16 |
| K00201 formylmethanofuran dehydrogenase subunit B [EC:1.2.7.12] | 0 | 4 | 4 | 1 | 0 | 4 | 2 | 0 | 4 | 15 | 2 | 12 |
| K00202 formylmethanofuran dehydrogenase subunit C [EC:1.2.7.12] | 2 | 2 | 3 | 0 | 0 | 0 | 2 | 2 | 4 | 4 | 2 | 10 |
| K00203 formylmethanofuran dehydrogenase subunit D [EC:1.2.7.12] | 0 | 0 | 0 | 0 | 0 | 0 | 0 | 0 | 2 | 2 | 3 | 1 |
| K00204 4Fe-4S ferredoxin | 0 | 2 | 0 | 0 | 0 | 0 | 2 | 1 | 2 | 0 | 3 | 9 |
| K00205 4Fe-4S ferredoxin | 0 | 1 | 2 | 0 | 2 | 6 | 1 | 0 | 7 | 19 | 11 | 15 |
| K00317 dimethylamine/trimethylamine dehydrogenase [EC:1.5.8.1 1.5.8.2] | 2 | 2 | 4 | 2 | 2 | 1 | 3 | 2 | 0 | 7 | 5 | 7 |
| K00319 methylenetetrahydromethanopterin dehydrogenase [EC:1.5.98.1] | 0 | 2 | 2 | 1 | 0 | 0 | 0 | 0 | 1 | 5 | 3 | 7 |
| K00320 5,10-methylenetetrahydromethanopterin reductase [EC:1.5.98.2] | 0 | 0 | 0 | 1 | 1 | 0 | 0 | 0 | 7 | 12 | 2 | 4 |
| K00399 methyl-coenzyme M reductase alpha subunit [EC:2.8.4.1] | 0 | 2 | 2 | 0 | 5 | 0 | 0 | 0 | 4 | 12 | 2 | 11 |
| K00400 methyl coenzyme M reductase system, component A2 | 3 | 5 | 0 | 3 | 4 | 11 | 0 | 2 | 13 | 25 | 13 | 21 |
| K00401 methyl-coenzyme M reductase beta subunit [EC:2.8.4.1] | 0 | 2 | 0 | 1 | 0 | 2 | 2 | 2 | 5 | 11 | 9 | 9 |
| K00402 methyl-coenzyme M reductase gamma subunit [EC:2.8.4.1] | 0 | 0 | 0 | 0 | 0 | 0 | 3 | 0 | 5 | 4 | 1 | 4 |
| K00440 coenzyme F420 hydrogenase subunit alpha [EC:1.12.98.1] | 3 | 0 | 2 | 1 | 0 | 0 | 0 | 3 | 4 | 12 | 12 | 2 |
| K00441 coenzyme F420 hydrogenase subunit beta [EC:1.12.98.1] | 3 | 0 | 0 | 1 | 2 | 1 | 1 | 2 | 8 | 21 | 12 | 24 |
| K00442 coenzyme F420 hydrogenase subunit delta | 0 | 0 | 0 | 0 | 0 | 0 | 0 | 0 | 2 | 2 | 0 | 1 |
| K00443 coenzyme F420 hydrogenase subunit gamma [EC:1.12.98.1] | 0 | 0 | 2 | 3 | 0 | 1 | 1 | 2 | 0 | 4 | 8 | 7 |
| K00577 tetrahydromethanopterin S-methyltransferase subunit A [EC:2.1.1.86] | 0 | 0 | 0 | 0 | 0 | 0 | 0 | 3 | 4 | 6 | 1 | 17 |
| K00578 tetrahydromethanopterin S-methyltransferase subunit B [EC:2.1.1.86] | 0 | 0 | 1 | 1 | 0 | 0 | 0 | 0 | 2 | 5 | 1 | 3 |
| K00579 tetrahydromethanopterin S-methyltransferase subunit C [EC:2.1.1.86] | 2 | 0 | 1 | 0 | 2 | 0 | 0 | 0 | 3 | 3 | 4 | 6 |
| K00580 tetrahydromethanopterin S-methyltransferase subunit D [EC:2.1.1.86] | 3 | 0 | 0 | 0 | 0 | 0 | 1 | 1 | 4 | 6 | 2 | 8 |
| K00581 tetrahydromethanopterin S-methyltransferase subunit E [EC:2.1.1.86] | 0 | 3 | 0 | 0 | 0 | 0 | 0 | 2 | 5 | 3 | 2 | 6 |

| | | | | | | | | | | | | |
|---|---|---|---|---|---|---|---|---|---|---|---|---|
| K00582 tetrahydromethanopterin S-methyltransferase subunit F [EC:2.1.1.86] | 0 | 0 | 1 | 0 | 0 | 0 | 0 | 1 | 0 | 0 | 0 | 1 |
| K00583 tetrahydromethanopterin S-methyltransferase subunit G [EC:2.1.1.86] | 0 | 1 | 0 | 0 | 1 | 1 | 0 | 0 | 0 | 0 | 0 | 0 |
| K00584 tetrahydromethanopterin S-methyltransferase subunit H [EC:2.1.1.86] | 0 | 0 | 0 | 3 | 0 | 2 | 0 | 2 | 12 | 16 | 5 | 22 |
| K00672 formylmethanofuran--tetrahydromethanopterin N-formyltransferase [EC:2.3.1.101] | 3 | 2 | 1 | 0 | 2 | 2 | 0 | 2 | 4 | 16 | 6 | 12 |
| K00831 phosphoserine aminotransferase [EC:2.6.1.52] | 2303 | 2293 | 2385 | 2285 | 3216 | 3259 | 3311 | 3247 | 3226 | 3008 | 3200 | 3103 |
| K01070 S-formylglutathione hydrolase [EC:3.1.2.12] | 1734 | 1674 | 1659 | 1699 | 992 | 1073 | 941 | 996 | 1185 | 1169 | 1119 | 1169 |
| K01079 phosphoserine phosphatase [EC:3.1.3.3] | 2516 | 2498 | 2413 | 2455 | 3399 | 3604 | 3451 | 3294 | 2869 | 2831 | 2918 | 2902 |
| K01499 methenyltetrahydromethanopterin cyclohydrolase [EC:3.5.4.27] | 8 | 7 | 8 | 17 | 0 | 0 | 3 | 2 | 10 | 13 | 5 | 12 |
| K02203 phosphoserine / homoserine phosphotransferase [EC:3.1.3.3 2.7.1.39] | 866 | 946 | 883 | 906 | 523 | 449 | 510 | 428 | 274 | 244 | 309 | 263 |
| K03388 heterodisulfide reductase subunit A2 [EC:1.8.7.3 1.8.98.4 1.8.98.5 1.8.98.6] | 104 | 119 | 124 | 150 | 53 | 38 | 39 | 35 | 244 | 265 | 193 | 247 |
| K03389 heterodisulfide reductase subunit B2 [EC:1.8.7.3 1.8.98.4 1.8.98.5 1.8.98.6] | 57 | 49 | 38 | 61 | 44 | 29 | 29 | 38 | 108 | 100 | 104 | 107 |
| K03390 heterodisulfide reductase subunit C2 [EC:1.8.7.3 1.8.98.4 1.8.98.5 1.8.98.6] | 34 | 19 | 21 | 31 | 18 | 21 | 17 | 21 | 56 | 70 | 61 | 61 |
| K03421 methyl-coenzyme M reductase subunit C | 0 | 0 | 0 | 0 | 2 | 0 | 0 | 2 | 2 | 4 | 8 | 6 |
| K03422 methyl-coenzyme M reductase subunit D | 0 | 0 | 0 | 0 | 0 | 0 | 0 | 2 | 0 | 8 | 0 | 3 |
| K03532 trimethylamine-N-oxide reductase (cytochrome c), cytochrome c-type subunit TorC | 14 | 26 | 26 | 31 | 38 | 46 | 30 | 35 | 103 | 82 | 101 | 84 |
| K03533 TorA specific chaperone | 4 | 9 | 2 | 3 | 8 | 6 | 5 | 10 | 47 | 36 | 51 | 28 |
| K04480 methanol---5-hydroxybenzimidazolylcobamide Co-methyltransferase [EC:2.1.1.90] | 10 | 7 | 25 | 19 | 24 | 32 | 18 | 14 | 47 | 50 | 46 | 73 |
| K05884 L-2-hydroxycarboxylate dehydrogenase (NAD+) [EC:1.1.1.337] | 36 | 47 | 30 | 28 | 6 | 2 | 2 | 1 | 87 | 90 | 88 | 106 |
| K05979 2-phosphosulfolactate phosphatase [EC:3.1.3.71] | 13 | 6 | 12 | 21 | 1 | 5 | 1 | 2 | 15 | 27 | 24 | 26 |
| K06034 sulfopyruvate decarboxylase subunit alpha [EC:4.1.1.79] | 0 | 0 | 0 | 0 | 0 | 0 | 2 | 0 | 4 | 2 | 5 | 9 |
| K06914 tyramine---L-glutamate ligase [EC:6.3.4.24] | 0 | 0 | 0 | 0 | 0 | 1 | 0 | 0 | 2 | 9 | 3 | 7 |
| K07072 (4-(4-[2-(gamma-L-glutamylamino)ethyl]phenoxymethyl)furan-2-yl)methanamine synthase [EC:2.5.1.131] | 2 | 0 | 0 | 0 | 0 | 5 | 0 | 2 | 3 | 5 | 2 | 6 |
| K07144 5-(aminomethyl)-3-furanmethanol phosphate kinase [EC:2.7.4.31] | 0 | 0 | 0 | 2 | 0 | 0 | 0 | 0 | 4 | 2 | 3 | 4 |
| K07811 trimethylamine-N-oxide reductase (cytochrome c) [EC:1.7.2.3] | 9 | 11 | 15 | 9 | 38 | 48 | 19 | 49 | 121 | 155 | 166 | 150 |
| K07812 trimethylamine-N-oxide reductase (cytochrome c) [EC:1.7.2.3] | 251 | 262 | 291 | 207 | 443 | 446 | 453 | 484 | 524 | 470 | 464 | 507 |
| K07821 trimethylamine-N-oxide reductase (cytochrome c), cytochrome c-type subunit TorY | 2 | 8 | 2 | 3 | 12 | 14 | 17 | 12 | 52 | 43 | 37 | 42 |
| K08097 phosphosulfolactate synthase [EC:4.4.1.19] | 0 | 0 | 0 | 0 | 0 | 0 | 0 | 0 | 2 | 6 | 2 | 4 |
| K08264 heterodisulfide reductase subunit D [EC:1.8.98.1] | 3 | 2 | 0 | 3 | 2 | 0 | 3 | 0 | 4 | 9 | 1 | 8 |
| K08685 quinohemoprotein amine dehydrogenase [EC:1.4.9.1] | 1 | 0 | 0 | 2 | 0 | 0 | 0 | 0 | 0 | 0 | 0 | 0 |
| K09733 (5-formylfuran-3-yl)methyl phosphate synthase [EC:4.2.3.153] | 2 | 3 | 0 | 0 | 0 | 0 | 2 | 0 | 0 | 9 | 3 | 4 |
| K10978 methanogen homoisocitrate dehydrogenase [EC:1.1.1.87 1.1.1.-] | 0 | 0 | 0 | 0 | 2 | 4 | 0 | 0 | 0 | 9 | 8 | 6 |
| K11212 LPPG:FO 2-phospho-L-lactate transferase [EC:2.7.8.28] | 0 | 1 | 5 | 0 | 2 | 4 | 2 | 0 | 8 | 4 | 4 | 1 |
| K11260 4Fe-4S ferredoxin | 0 | 0 | 1 | 0 | 0 | 0 | 2 | 0 | 0 | 1 | 1 | 1 |
| K11261 formylmethanofuran dehydrogenase subunit E [EC:1.2.7.12] | 127 | 123 | 125 | 149 | 54 | 45 | 47 | 52 | 211 | 259 | 234 | 247 |
| K11780 7,8-didemethyl-8-hydroxy-5-deazariboflavin synthase [EC:4.3.1.32] | 0 | 0 | 2 | 0 | 3 | 0 | 4 | 2 | 4 | 6 | 4 | 3 |
| K11781 5-amino-6-(D-ribitylamino)uracil---L-tyrosine 4-hydroxyphenyl transferase [EC:2.5.1.147] | 3 | 3 | 0 | 0 | 7 | 0 | 0 | 0 | 5 | 13 | 3 | 8 |
| K12234 coenzyme F420-0:L-glutamate ligase / coenzyme F420-1:gamma-L-glutamate ligase [EC:6.3.2.31 6.3.2.34] | 0 | 1 | 0 | 0 | 3 | 0 | 3 | 0 | 1 | 2 | 1 | 9 |
| K13039 sulfopyruvate decarboxylase subunit beta [EC:4.1.1.79] | 0 | 0 | 0 | 1 | 0 | 0 | 0 | 0 | 2 | 9 | 3 | 1 |
| K13942 5,10-methenyltetrahydromethanopterin hydrogenase [EC:1.12.98.2] | 2 | 2 | 0 | 0 | 3 | 0 | 0 | 2 | 0 | 6 | 3 | 11 |
| K14080 [methyl-Co(III) methanol-specific corrinoid protein]:coenzyme M methyltransferase [EC:2.1.1.246] | 8 | 20 | 11 | 9 | 11 | 10 | 8 | 4 | 16 | 24 | 21 | 31 |
| K14081 methanol corrinoid protein | 0 | 0 | 1 | 0 | 0 | 2 | 2 | 5 | 4 | 9 | 3 | 9 |
| K14083 trimethylamine---corrinoid protein Co-methyltransferase [EC:2.1.1.250] | 822 | 808 | 820 | 875 | 723 | 693 | 718 | 659 | 1915 | 2087 | 1829 | 2205 |
| K14084 trimethylamine corrinoid protein | 2 | 2 | 1 | 6 | 0 | 0 | 2 | 0 | 1 | 2 | 2 | 2 |
| K14126 F420-non-reducing hydrogenase large subunit [EC:1.12.99.- 1.8.98.5] | 3 | 6 | 0 | 3 | 0 | 4 | 5 | 4 | 4 | 2 | 0 | 28 |
| K14127 F420-non-reducing hydrogenase iron-sulfur subunit [EC:1.12.99.- 1.8.98.5 1.8.98.6] | 0 | 0 | 0 | 0 | 0 | 1 | 0 | 0 | 6 | 4 | 0 | 4 |
| K14128 F420-non-reducing hydrogenase small subunit [EC:1.12.99.- 1.8.98.5] | 0 | 0 | 1 | 0 | 0 | 0 | 2 | 1 | 2 | 8 | 3 | 20 |
| K14941 2-phospho-L-lactate guanylyltransferase [EC:2.7.7.68] | 2 | 0 | 1 | 1 | 0 | 0 | 0 | 0 | 2 | 1 | 3 | 7 |
| K16176 methylamine---corrinoid protein Co-methyltransferase [EC:2.1.1.248] | 10 | 12 | 12 | 8 | 3 | 5 | 2 | 2 | 5 | 4 | 9 | 4 |
| K16178 dimethylamine---corrinoid protein Co-methyltransferase [EC:2.1.1.249] | 0 | 0 | 0 | 0 | 0 | 0 | 0 | 0 | 0 | 0 | 0 | 1 |
| K16179 dimethylamine corrinoid protein | 0 | 0 | 0 | 1 | 0 | 0 | 1 | 0 | 0 | 0 | 0 | 1 |
| K16792 methanogen homoaconitase large subunit [EC:4.2.1.114] | 1 | 0 | 4 | 4 | 1 | 3 | 3 | 0 | 4 | 7 | 9 | 12 |
| K16793 methanogen homoaconitase small subunit [EC:4.2.1.114] | 1 | 0 | 0 | 0 | 0 | 0 | 1 | 0 | 1 | 2 | 1 | 5 |
| K18933 tyrosine decarboxylase / aspartate 1-decarboxylase [EC:4.1.1.25 4.1.1.11] | 0 | 0 | 0 | 3 | 2 | 1 | 2 | 2 | 3 | 0 | 8 | 5 |
| K19793 (5-formylfuran-3-yl)methyl phosphate transaminase [EC:2.6.1.108] | 0 | 0 | 1 | 0 | 0 | 1 | 0 | 0 | 0 | 0 | 0 | 0 |
| K22081 methylamine---glutamate N-methyltransferase subunit A [EC:2.1.1.21] | 0 | 0 | 0 | 0 | 0 | 0 | 0 | 2 | 0 | 0 | 0 | 0 |
| K22083 methylamine---glutamate N-methyltransferase subunit C [EC:2.1.1.21] | 1 | 0 | 0 | 0 | 0 | 0 | 0 | 3 | 0 | 0 | 0 | 0 |
| K22084 methylglutamate dehydrogenase subunit A [EC:1.5.99.5] | 0 | 0 | 0 | 1 | 0 | 1 | 0 | 0 | 0 | 1 | 0 | 0 |
| K22305 phosphoserine phosphatase [EC:3.1.3.3] | 4 | 2 | 3 | 3 | 3 | 1 | 2 | 3 | 6 | 7 | 10 | 11 |
| K22480 heterodisulfide reductase subunit A1 [EC:1.8.7.3] | 8 | 3 | 3 | 1 | 0 | 0 | 2 | 0 | 13 | 15 | 3 | 15 |
| K22481 heterodisulfide reductase subunit B1 [EC:1.8.7.3] | 2 | 0 | 0 | 1 | 0 | 1 | 0 | 2 | 6 | 7 | 3 | 10 |
| K22482 heterodisulfide reductase subunit C1 [EC:1.8.7.3] | 0 | 1 | 0 | 1 | 0 | 1 | 0 | 0 | 3 | 4 | 0 | 1 |
| K22516 formate dehydrogenase (coenzyme F420) alpha subunit [EC:1.17.98.3 1.8.98.6] | 36 | 26 | 20 | 31 | 14 | 8 | 19 | 9 | 22 | 35 | 29 | 26 |
| ko00910 Nitrogen metabolism | 120034 | 119552 | 117160 | 118852 | 102402 | 101623 | 100410 | 101796 | 98543 | 97125 | 97001 | 97654 |
| K00260 glutamate dehydrogenase [EC:1.4.1.2] | 59 | 72 | 65 | 44 | 29 | 27 | 25 | 23 | 92 | 109 | 114 | 93 |
| K00261 glutamate dehydrogenase (NAD(P)+) [EC:1.4.1.3] | 584 | 540 | 564 | 646 | 147 | 173 | 164 | 144 | 1125 | 1169 | 1191 | 1171 |
| K00262 glutamate dehydrogenase (NADP+) [EC:1.4.1.4] | 4671 | 4827 | 5022 | 4918 | 7000 | 7211 | 7056 | 7039 | 6549 | 6261 | 6492 | 6096 |
| K00264 glutamate synthase (NADH) [EC:1.4.1.14] | 8 | 14 | 19 | 11 | 11 | 10 | 19 | 9 | 22 | 11 | 21 | 14 |
| K00265 glutamate synthase (NADPH) large chain [EC:1.4.1.13] | 14239 | 14566 | 14524 | 14318 | 15161 | 14791 | 14306 | 14093 | 14526 | 14253 | 14248 | 14084 |
| K00266 glutamate synthase (NADPH) small chain [EC:1.4.1.13] | 7387 | 7375 | 7496 | 7262 | 9781 | 9848 | 10089 | 9830 | 9306 | 9188 | 9350 | 9228 |
| K00360 assimilatory nitrate reductase electron transfer subunit [EC:1.7.99.-] | 3 | 0 | 0 | 0 | 0 | 1 | 0 | 0 | 0 | 0 | 0 | 0 |
| K00362 nitrite reductase (NADH) large subunit [EC:1.7.1.15] | 7321 | 7187 | 6973 | 7380 | 4370 | 4329 | 4308 | 4460 | 6239 | 6149 | 6048 | 6145 |
| K00363 nitrite reductase (NADH) small subunit [EC:1.7.1.15] | 730 | 770 | 719 | 766 | 457 | 492 | 452 | 427 | 517 | 526 | 448 | 476 |
| K00366 ferredoxin-nitrite reductase [EC:1.7.7.1] | 30 | 42 | 26 | 40 | 12 | 3 | 11 | 5 | 19 | 26 | 24 | 29 |
| K00368 nitrite reductase (NO-forming) [EC:1.7.2.1] | 0 | 0 | 0 | 2 | 1 | 2 | 1 | 2 | 4 | 16 | 3 | 5 |
| K00370 nitrate reductase / nitrite oxidoreductase, alpha subunit [EC:1.7.5.1 1.7.99.-] | 9684 | 9631 | 9278 | 9660 | 5346 | 5224 | 4896 | 5196 | 7670 | 7529 | 7284 | 7779 |
| K00371 nitrate reductase / nitrite oxidoreductase, beta subunit [EC:1.7.5.1 1.7.99.-] | 2332 | 2280 | 2264 | 2393 | 1238 | 1296 | 1185 | 1290 | 2727 | 2691 | 2562 | 2621 |
| K00372 assimilatory nitrate reductase catalytic subunit [EC:1.7.99.-] | 5354 | 5390 | 5194 | 5361 | 3092 | 3005 | 2854 | 2930 | 3085 | 3080 | 2888 | 3275 |
| K00374 nitrate reductase gamma subunit [EC:1.7.5.1 1.7.99.-] | 1527 | 1484 | 1539 | 1529 | 920 | 870 | 839 | 857 | 1255 | 1177 | 1231 | 1213 |
| K00376 nitrous-oxide reductase [EC:1.7.2.4] | 1875 | 1864 | 1726 | 1846 | 980 | 1035 | 970 | 1056 | 578 | 579 | 588 | 540 |
| K00459 nitronate monooxygenase [EC:1.13.12.16] | 5347 | 5300 | 5010 | 5282 | 2775 | 2694 | 2695 | 2877 | 2322 | 2312 | 2341 | 2508 |
| K00531 nitrogenase delta subunit [EC:1.18.6.1] | 3 | 0 | 2 | 1 | 1 | 1 | 0 | 4 | 2 | 5 | 0 | 2 |
| K00926 carbamate kinase [EC:2.7.2.2] | 1632 | 1635 | 1708 | 1774 | 978 | 970 | 941 | 925 | 1466 | 1519 | 1470 | 1404 |
| K01501 nitrilase [EC:3.5.5.1] | 21 | 22 | 22 | 21 | 17 | 12 | 5 | 10 | 39 | 45 | 44 | 36 |
| K01673 carbonic anhydrase [EC:4.2.1.1] | 4600 | 4718 | 4518 | 4496 | 3490 | 3383 | 3441 | 3485 | 3333 | 3341 | 3278 | 3471 |
| K01674 carbonic anhydrase [EC:4.2.1.1] | 25 | 27 | 33 | 33 | 14 | 5 | 6 | 10 | 31 | 50 | 41 | 21 |
| K01725 cyanate lyase [EC:4.2.1.104] | 420 | 419 | 364 | 384 | 208 | 246 | 210 | 255 | 110 | 111 | 107 | 136 |
| K01948 carbamoyl-phosphate synthase (ammonia) [EC:6.3.4.16] | 1 | 0 | 0 | 0 | 0 | 1 | 0 | 0 | 0 | 0 | 0 | 0 |
| K02305 nitric oxide reductase subunit C | 682 | 676 | 629 | 675 | 332 | 371 | 336 | 389 | 241 | 209 | 215 | 189 |
| K02567 periplasmic nitrate reductase NapA [EC:1.7.99.-] | 3951 | 4109 | 3678 | 3776 | 2245 | 2189 | 2197 | 2421 | 1612 | 1601 | 1529 | 1539 |
| K02568 cytochrome c-type protein NapB | 659 | 618 | 619 | 583 | 323 | 356 | 339 | 319 | 220 | 219 | 210 | 214 |
| K02575 MFS transporter, NNP family, nitrate/nitrite transporter | 5611 | 5652 | 5390 | 5528 | 2960 | 3034 | 2762 | 2960 | 3469 | 3354 | 3279 | 3647 |
| K02586 nitrogenase molybdenum-iron protein alpha chain [EC:1.18.6.1] | 336 | 316 | 283 | 348 | 347 | 427 | 416 | 424 | 78 | 100 | 101 | 95 |
| K02588 nitrogenase iron protein NifH | 289 | 262 | 266 | 309 | 316 | 344 | 334 | 326 | 137 | 183 | 169 | 163 |
| K02591 nitrogenase molybdenum-iron protein beta chain [EC:1.18.6.1] | 315 | 315 | 305 | 292 | 471 | 456 | 447 | 438 | 18 | 42 | 27 | 41 |
| K03385 nitrite reductase (cytochrome c-552) [EC:1.7.2.2] | 1591 | 1725 | 1843 | 1715 | 5713 | 5810 | 5969 | 5551 | 4412 | 4137 | 4603 | 4216 |
| K04561 nitric oxide reductase subunit B [EC:1.7.2.5] | 2180 | 2103 | 1969 | 2058 | 1250 | 1118 | 1042 | 1223 | 724 | 676 | 616 | 701 |
| K05601 hydroxylamine reductase [EC:1.7.99.1] | 2761 | 2650 | 2685 | 2897 | 4045 | 4002 | 4128 | 3953 | 4184 | 4047 | 4092 | 4162 |
| K15371 glutamate dehydrogenase [EC:1.4.1.2] | 7394 | 7227 | 6792 | 7016 | 3804 | 3820 | 3548 | 3904 | 2316 | 2180 | 2149 | 2180 |
| K15576 nitrate/nitrite transport system substrate-binding protein | 777 | 655 | 646 | 707 | 419 | 412 | 419 | 450 | 1017 | 968 | 973 | 1050 |
| K15577 nitrate/nitrite transport system permease protein | 384 | 438 | 418 | 415 | 287 | 277 | 321 | 337 | 510 | 594 | 591 | 600 |
| K15578 nitrate/nitrite transport system ATP-binding protein [EC:3.6.3.-] | 477 | 442 | 394 | 445 | 284 | 358 | 304 | 264 | 520 | 557 | 553 | 611 |
| K15864 nitrite reductase (NO-forming) / hydroxylamine reductase [EC:1.7.2.1 1.7.99.1] | 2360 | 2254 | 2132 | 2155 | 1234 | 1232 | 1176 | 1326 | 759 | 712 | 630 | 750 |
| K15876 cytochrome c nitrite reductase small subunit | 290 | 319 | 369 | 298 | 986 | 1063 | 1062 | 1037 | 790 | 815 | 784 | 697 |
| ko00920 Sulfur metabolism | 96207 | 96252 | 92776 | 95736 | 67620 | 67770 | 65884 | 68472 | 68196 | 68496 | 67501 | 68763 |
| K00299 FMN reductase [EC:1.5.1.38] | 1645 | 1659 | 1532 | 1701 | 898 | 914 | 843 | 982 | 789 | 875 | 765 | 915 |
| K00380 sulfite reductase (NADPH) flavoprotein alpha-component [EC:1.8.1.2] | 5044 | 4900 | 4798 | 4852 | 2923 | 2808 | 2932 | 2771 | 2580 | 2662 | 2748 | 2818 |
| K00381 sulfite reductase (NADPH) hemoprotein beta-component [EC:1.8.1.2] | 6118 | 6141 | 5914 | 6135 | 3497 | 3591 | 3390 | 3681 | 3065 | 2912 | 2792 | 2922 |
| K00385 anaerobic sulfite reductase subunit C | 563 | 491 | 552 | 542 | 98 | 120 | 141 | 113 | 436 | 463 | 521 | 475 |
| K00390 phosphoadenosine phosphosulfate reductase [EC:1.8.4.8 1.8.4.10] | 1562 | 1515 | 1571 | 1588 | 575 | 610 | 560 | 570 | 1002 | 1192 | 1134 | 1214 |
| K00394 adenylylsulfate reductase, subunit A [EC:1.8.99.2] | 24 | 14 | 16 | 14 | 5 | 9 | 7 | 6 | 21 | 36 | 26 | 29 |
| K00395 adenylylsulfate reductase, subunit B [EC:1.8.99.2] | 4 | 13 | 2 | 9 | 9 | 11 | 5 | 9 | 3 | 9 | 0 | 4 |
| K00640 serine O-acetyltransferase [EC:2.3.1.30] | 3960 | 3994 | 3982 | 4081 | 4079 | 4016 | 4022 | 3925 | 3366 | 3741 | 3521 | 3468 |
| K00641 homoserine O-acetyltransferase/O-succinyltransferase [EC:2.3.1.31 2.3.1.46] | 2008 | 1968 | 1997 | 1939 | 1195 | 1153 | 1046 | 1201 | 703 | 632 | 644 | 623 |
| K00651 homoserine O-succinyltransferase/O-acetyltransferase [EC:2.3.1.46 2.3.1.31] | 1767 | 1758 | 1818 | 1816 | 2681 | 2773 | 2688 | 2704 | 2670 | 2718 | 2753 | 2761 |
| K00860 adenylylsulfate kinase [EC:2.7.1.25] | 725 | 772 | 758 | 778 | 1408 | 1355 | 1567 | 1466 | 1322 | 1429 | 1381 | 1414 |
| K00955 bifunctional enzyme CysN/CysC [EC:2.7.7.4 2.7.1.25] | 2782 | 2672 | 2860 | 2862 | 1714 | 1659 | 1620 | 1664 | 991 | 876 | 916 | 840 |
| K00956 sulfate adenylyltransferase subunit 1 [EC:2.7.7.4] | 1789 | 1797 | 1856 | 1898 | 3663 | 3894 | 3870 | 3734 | 3343 | 3321 | 3303 | 3223 |
| K00957 sulfate adenylyltransferase subunit 2 [EC:2.7.7.4] | 1611 | 1619 | 1611 | 1618 | 2522 | 2665 | 2560 | 2458 | 2268 | 2038 | 2169 | 2153 |
| K00958 sulfate adenylyltransferase [EC:2.7.7.4] | 5 | 9 | 9 | 7 | 2 | 4 | 3 | 3 | 9 | 14 | 8 | 22 |
| K01011 thiosulfate/3-mercaptopyruvate sulfurtransferase [EC:2.8.1.1 2.8.1.2] | 2402 | 2361 | 2238 | 2334 | 1421 | 1280 | 1297 | 1442 | 1247 | 1241 | 1223 | 1251 |
| K01082 3'(2'), 5'-bisphosphate nucleotidase [EC:3.1.3.7] | 2530 | 2409 | 2464 | 2419 | 2850 | 2841 | 2836 | 2810 | 2174 | 2084 | 2295 | 2239 |

| | | | | | | | | | | | | |
|---|---|---|---|---|---|---|---|---|---|---|---|---|
| K01738 cysteine synthase [EC:2.5.1.47] | 5828 | 5923 | 5827 | 5924 | 6532 | 6628 | 6576 | 6497 | 6452 | 6559 | 6224 | 6254 |
| K01739 cystathionine gamma-synthase [EC:2.5.1.48] | 774 | 733 | 744 | 871 | 557 | 639 | 570 | 495 | 1056 | 1074 | 1219 | 1158 |
| K02045 sulfate/thiosulfate transport system ATP-binding protein [EC:7.3.2.3] | 2029 | 1857 | 1833 | 1972 | 1270 | 1194 | 1245 | 1187 | 1482 | 1409 | 1431 | 1375 |
| K02046 sulfate/thiosulfate transport system permease protein | 1860 | 1919 | 1712 | 1923 | 1118 | 1130 | 1126 | 1156 | 1145 | 1123 | 1214 | 1115 |
| K02047 sulfate/thiosulfate transport system permease protein | 1419 | 1475 | 1404 | 1437 | 902 | 884 | 909 | 929 | 1071 | 1095 | 1007 | 1031 |
| K02048 sulfate/thiosulfate transport system substrate-binding protein | 1531 | 1579 | 1521 | 1574 | 1018 | 978 | 870 | 1008 | 1170 | 1180 | 1148 | 1151 |
| K02439 thiosulfate sulfurtransferase [EC:2.8.1.1] | 727 | 741 | 741 | 780 | 497 | 496 | 417 | 473 | 421 | 449 | 517 | 507 |
| K03119 taurine dioxygenase [EC:1.14.11.17] | 2826 | 2871 | 2720 | 2625 | 1552 | 1438 | 1320 | 1472 | 1485 | 1438 | 1385 | 1494 |
| K04091 alkanesulfonate monooxygenase [EC:1.14.14.5] | 6864 | 6964 | 6398 | 6640 | 3271 | 3206 | 2913 | 3432 | 4432 | 4178 | 4191 | 4399 |
| K05907 adenylyl-sulfate reductase (glutathione) [EC:1.8.4.9] | 0 | 1 | 0 | 0 | 0 | 0 | 0 | 0 | 0 | 0 | 0 | 0 |
| K06881 bifunctional oligoribonuclease and PAP phosphatase NrnA [EC:3.1.3.7 3.1.13.3] | 1176 | 1280 | 1336 | 1269 | 2344 | 2524 | 2387 | 2379 | 2009 | 2022 | 2079 | 2008 |
| K07306 anaerobic dimethyl sulfoxide reductase subunit A [EC:1.8.5.3] | 1341 | 1417 | 1470 | 1467 | 1024 | 1062 | 1010 | 974 | 1886 | 2099 | 1944 | 2014 |
| K07307 anaerobic dimethyl sulfoxide reductase subunit B | 709 | 712 | 628 | 621 | 426 | 482 | 495 | 509 | 909 | 1046 | 959 | 1030 |
| K07308 anaerobic dimethyl sulfoxide reductase subunit C | 505 | 551 | 481 | 557 | 337 | 363 | 373 | 287 | 748 | 812 | 706 | 767 |
| K08352 thiosulfate reductase / polysulfide reductase chain A [EC:1.8.5.5] | 71 | 62 | 52 | 48 | 36 | 34 | 22 | 36 | 95 | 68 | 84 | 105 |
| K08353 thiosulfate reductase electron transport protein | 3 | 0 | 0 | 1 | 0 | 2 | 0 | 0 | 0 | 0 | 0 | 0 |
| K08354 thiosulfate reductase cytochrome b subunit | 4 | 2 | 5 | 1 | 7 | 22 | 8 | 14 | 40 | 53 | 34 | 33 |
| K08357 tetrathionate reductase subunit A | 14 | 22 | 8 | 20 | 4 | 17 | 5 | 14 | 14 | 32 | 27 | 45 |
| K08358 tetrathionate reductase subunit B | 4 | 0 | 4 | 3 | 5 | 0 | 0 | 0 | 10 | 12 | 1 | 4 |
| K08359 tetrathionate reductase subunit C | 0 | 0 | 1 | 1 | 2 | 2 | 2 | 2 | 1 | 6 | 3 | 2 |
| K10764 O-succinylhomoserine sulfhydrylase [EC:2.5.1.-] | 1782 | 1803 | 1799 | 1710 | 900 | 953 | 926 | 1075 | 592 | 547 | 536 | 528 |
| K10831 taurine transport system ATP-binding protein [EC:7.6.2.7] | 1730 | 1629 | 1524 | 1597 | 743 | 805 | 720 | 770 | 1103 | 950 | 966 | 1068 |
| K11180 dissimilatory sulfite reductase alpha subunit [EC:1.8.99.5] | 8 | 21 | 15 | 13 | 6 | 5 | 5 | 4 | 11 | 16 | 9 | 7 |
| K11181 dissimilatory sulfite reductase beta subunit [EC:1.8.99.5] | 16 | 5 | 19 | 6 | 4 | 3 | 6 | 7 | 4 | 35 | 9 | 15 |
| K15551 taurine transport system substrate-binding protein | 1978 | 1940 | 1804 | 1930 | 1142 | 1208 | 1080 | 1150 | 1166 | 1223 | 1114 | 1285 |
| K15552 taurine transport system permease protein | 1927 | 1906 | 1773 | 1774 | 1065 | 1098 | 1007 | 1061 | 1207 | 1220 | 1226 | 1221 |
| K15553 sulfonate transport system substrate-binding protein | 9591 | 9871 | 9258 | 9576 | 4932 | 4712 | 4690 | 5068 | 5048 | 4838 | 4846 | 5025 |
| K15554 sulfonate transport system permease protein | 4942 | 4799 | 4470 | 4804 | 2505 | 2319 | 2292 | 2630 | 2186 | 2104 | 1983 | 2178 |
| K15555 sulfonate transport system ATP-binding protein [EC:3.6.3.-] | 4238 | 4267 | 3970 | 4307 | 2165 | 2222 | 1919 | 2348 | 1861 | 1976 | 1874 | 1853 |
| K16937 thiosulfate dehydrogenase [quinone] large subunit [EC:1.8.5.2] | 5 | 4 | 7 | 6 | 0 | 0 | 2 | 0 | 1 | 2 | 0 | 1 |
| K16950 anaerobic sulfite reductase subunit A | 480 | 399 | 490 | 510 | 100 | 108 | 101 | 129 | 363 | 445 | 427 | 427 |
| K16951 anaerobic sulfite reductase subunit B | 376 | 385 | 455 | 411 | 88 | 93 | 103 | 76 | 309 | 371 | 326 | 332 |
| K17069 O-acetylhomoserine/O-acetylserine sulfhydrylase [EC:2.5.1.49 2.5.1.47] | 0 | 2 | 0 | 0 | 0 | 0 | 0 | 1 | 0 | 0 | 0 | 0 |
| K17217 cystathionine gamma-lyase / homocysteine desulfhydrase [EC:4.4.1.1 4.4.1.2] | 0 | 0 | 1 | 0 | 0 | 0 | 0 | 0 | 0 | 0 | 1 | 1 |
| K17218 sulfide:quinone oxidoreductase [EC:1.8.5.4] | 3466 | 3442 | 3097 | 3222 | 1692 | 1618 | 1590 | 1787 | 1106 | 1004 | 960 | 1028 |
| K17222 L-cysteine S-thiosulfotransferase [EC:2.8.5.2] | 0 | 0 | 0 | 0 | 0 | 0 | 0 | 0 | 2 | 0 | 0 | 0 |
| K17226 sulfur-oxidizing protein SoxY | 2 | 0 | 0 | 0 | 0 | 0 | 0 | 0 | 0 | 0 | 0 | 0 |
| K17228 dimethylsulfone monooxygenase [EC:1.14.14.35] | 536 | 566 | 474 | 540 | 268 | 293 | 259 | 332 | 186 | 165 | 169 | 162 |
| K20034 3-(methylthio)propionyl---CoA ligase [EC:6.2.1.44] | 597 | 593 | 554 | 653 | 170 | 156 | 156 | 151 | 1277 | 1254 | 1263 | 1289 |
| K20035 3-(methylthio)propanoyl-CoA dehydrogenase [EC:1.3.8.-] | 2 | 0 | 1 | 1 | 0 | 0 | 0 | 0 | 1 | 1 | 1 | 0 |
| K21307 sulfite dehydrogenase (quinone) subunit SoeA [EC:1.8.5.6] | 10 | 1 | 2 | 9 | 2 | 1 | 3 | 3 | 4 | 11 | 13 | 11 |
| K21308 sulfite dehydrogenase (quinone) subunit SoeB | 1 | 0 | 0 | 0 | 0 | 0 | 0 | 0 | 0 | 0 | 0 | 0 |
| K23163 sulfate/thiosulfate transport system substrate-binding protein | 2296 | 2418 | 2200 | 2339 | 1391 | 1371 | 1387 | 1477 | 1354 | 1436 | 1406 | 1468 |
| K23304 serine O-acetyltransferase [EC:2.3.1.30] | 0 | 0 | 0 | 1 | 5 | 1 | 3 | 0 | 0 | 0 | 0 | 1 |
| Lipid metabolism | 309133 | 311310 | 306979 | 308645 | 306871 | 305504 | 305084 | 306841 | 292771 | 289500 | 288010 | 289817 |
| ko00061 Fatty acid biosynthesis | 66452 | 66872 | 66378 | 66508 | 70556 | 69105 | 70239 | 70803 | 62022 | 61518 | 61559 | 61627 |
| K00059 3-oxoacyl-[acyl-carrier protein] reductase [EC:1.1.1.100] | 13605 | 13158 | 13121 | 13004 | 9680 | 9267 | 9323 | 9691 | 8537 | 8520 | 8456 | 8451 |
| K00208 enoyl-[acyl-carrier protein] reductase I [EC:1.3.1.9 1.3.1.10] | 1438 | 1495 | 1521 | 1533 | 2368 | 2390 | 2378 | 2253 | 2194 | 2014 | 2126 | 2133 |
| K00645 [acyl-carrier-protein] S-malonyltransferase [EC:2.3.1.39] | 3126 | 3021 | 3156 | 3243 | 2939 | 2989 | 2957 | 2925 | 3173 | 3232 | 3216 | 3225 |
| K00647 3-oxoacyl-[acyl-carrier-protein] synthase I [EC:2.3.1.41] | 4745 | 4733 | 4459 | 4601 | 3014 | 2927 | 2781 | 3133 | 2225 | 2219 | 2169 | 2295 |
| K00648 3-oxoacyl-[acyl-carrier-protein] synthase III [EC:2.3.1.180] | 3715 | 4111 | 4025 | 3925 | 5668 | 5615 | 5633 | 5569 | 4836 | 4801 | 4796 | 4751 |
| K00667 fatty acid synthase subunit alpha, fungi type [EC:2.3.1.86] | 0 | 2 | 0 | 0 | 0 | 0 | 0 | 0 | 0 | 0 | 0 | 0 |
| K01071 medium-chain acyl-[acyl-carrier-protein] hydrolase [EC:3.1.2.21] | 65 | 58 | 52 | 57 | 24 | 20 | 30 | 21 | 53 | 90 | 50 | 57 |
| K01716 3-hydroxyacyl-[acyl-carrier protein] dehydratase / trans-2-decenoyl-[acyl-carrier protein] isomerase [EC:4.2.1.59 5.3.3.14] | 750 | 917 | 808 | 819 | 537 | 511 | 492 | 553 | 599 | 602 | 634 | 596 |
| K01897 long-chain acyl-CoA synthetase [EC:6.2.1.3] | 8770 | 9406 | 9278 | 9093 | 17228 | 16696 | 17565 | 17395 | 13676 | 13087 | 13358 | 13238 |
| K02078 acyl carrier protein | 1597 | 1533 | 1518 | 1673 | 1524 | 1501 | 1511 | 1628 | 1542 | 1495 | 1572 | 1570 |
| K02371 enoyl-[acyl-carrier protein] reductase II [EC:1.3.1.9] | 1555 | 1614 | 1626 | 1652 | 2122 | 2055 | 2065 | 2065 | 2538 | 2576 | 2560 | 2511 |
| K02372 3-hydroxyacyl-[acyl-carrier-protein] dehydratase [EC:4.2.1.59] | 1529 | 1441 | 1523 | 1471 | 819 | 796 | 844 | 837 | 1074 | 1132 | 1061 | 1174 |
| K03921 acyl-[acyl-carrier-protein] desaturase [EC:1.14.19.2 1.14.19.11 1.14.19.26] | 0 | 2 | 0 | 0 | 0 | 0 | 0 | 0 | 0 | 0 | 2 | 0 |
| K09458 3-oxoacyl-[acyl-carrier-protein] synthase II [EC:2.3.1.179] | 7077 | 6999 | 7052 | 7012 | 6574 | 6571 | 6635 | 6363 | 6355 | 6449 | 6461 | 6317 |
| K10780 enoyl-[acyl-carrier protein] reductase III [EC:1.3.1.104] | 2 | 2 | 1 | 1 | 0 | 0 | 1 | 0 | 3 | 1 | 3 | 1 |
| K11533 fatty acid synthase, bacteria type [EC:2.3.1.-] | 11 | 8 | 7 | 15 | 8 | 10 | 12 | 14 | 39 | 49 | 53 | 101 |
| K13370 17beta-estradiol 17-dehydrogenase / 3alpha(17beta)-hydroxysteroid dehydrogenase (NAD+) / 3-oxoacyl-[acyl-carrier protein] reductase alpha subuni | 1 | 1 | 0 | 1 | 0 | 1 | 0 | 0 | 0 | 0 | 0 | 0 |
| K16363 UDP-3-O-[3-hydroxymyristoyl] N-acetylglucosamine deacetylase / 3-hydroxyacyl-[acyl-carrier-protein] dehydratase [EC:3.5.1.108 4.2.1.59] | 974 | 927 | 938 | 919 | 2968 | 3011 | 2989 | 2986 | 2053 | 1991 | 2036 | 1939 |
| K18473 acetoacetyl-[acyl-carrier protein] synthase [EC:2.3.1.180] | 3589 | 3386 | 3346 | 3566 | 2120 | 1860 | 1943 | 2037 | 1148 | 1092 | 1056 | 1157 |
| K18474 trans-2-decenoyl-[acyl-carrier protein] isomerase [EC:5.3.3.14] | 3 | 7 | 10 | 5 | 0 | 0 | 2 | 0 | 0 | 0 | 2 | 3 |
| K18660 malonyl-CoA/methylmalonyl-CoA synthetase [EC:6.2.1.-] | 0 | 0 | 0 | 0 | 0 | 1 | 0 | 0 | 0 | 0 | 0 | 0 |
| K22540 3-hydroxyacyl-thioester dehydratase, animal type [EC:4.2.1.-] | 0 | 0 | 1 | 1 | 0 | 0 | 0 | 0 | 0 | 0 | 0 | 0 |
| ko00062 Fatty acid elongation | 5 | 0 | 4 | 1 | 2 | 0 | 0 | 3 | 1 | 0 | 0 | 0 |
| K01068 acyl-coenzyme A thioesterase 1/2/4 [EC:3.1.2.2] | 0 | 0 | 0 | 0 | 2 | 0 | 0 | 0 | 0 | 0 | 0 | 0 |
| K01074 palmitoyl-protein thioesterase [EC:3.1.2.22] | 0 | 0 | 1 | 0 | 0 | 0 | 0 | 0 | 0 | 0 | 0 | 0 |
| K07508 acetyl-CoA acyltransferase 2 [EC:2.3.1.16] | 1 | 0 | 1 | 0 | 0 | 0 | 0 | 2 | 0 | 0 | 0 | 0 |
| ko00071 Fatty acid degradation | 83216 | 82836 | 80666 | 81489 | 59791 | 58151 | 57555 | 59765 | 53800 | 53036 | 52240 | 53654 |
| K00249 acyl-CoA dehydrogenase [EC:1.3.8.7] | 12040 | 11683 | 11576 | 11551 | 6589 | 6315 | 6025 | 6736 | 3910 | 3629 | 3652 | 3657 |
| K00252 glutaryl-CoA dehydrogenase [EC:1.3.8.6] | 1884 | 1899 | 1710 | 1822 | 1064 | 957 | 991 | 992 | 606 | 580 | 507 | 596 |
| K00496 alkane 1-monooxygenase [EC:1.14.15.3] | 3112 | 2999 | 3072 | 3017 | 1802 | 1663 | 1528 | 1644 | 984 | 1003 | 897 | 988 |
| K00529 3-phenylpropionate/trans-cinnamate dioxygenase ferredoxin reductase component [EC:1.18.1.3] | 127 | 149 | 118 | 99 | 20 | 5 | 13 | 8 | 4 | 7 | 5 | 5 |
| K00632 acetyl-CoA acyltransferase [EC:2.3.1.16] | 6771 | 6698 | 6402 | 6636 | 4031 | 3892 | 3923 | 3925 | 3815 | 3724 | 3658 | 3839 |
| K01909 long-chain-fatty-acid--[acyl-carrier-protein] ligase [EC:6.2.1.20] | 0 | 0 | 1 | 1 | 0 | 0 | 0 | 0 | 0 | 0 | 0 | 1 |
| K05297 rubredoxin---NAD+ reductase [EC:1.18.1.1] | 2218 | 2122 | 2033 | 1965 | 1193 | 1119 | 1081 | 1201 | 704 | 613 | 622 | 710 |
| K05939 acyl-[acyl-carrier-protein]-phospholipid O-acyltransferase / long-chain-fatty-acid--[acyl-carrier-protein] ligase [EC:2.3.1.40 6.2.1.20] | 1432 | 1359 | 1316 | 1323 | 896 | 1075 | 946 | 966 | 1758 | 1863 | 1890 | 1888 |
| K06445 acyl-CoA dehydrogenase [EC:1.3.99.-] | 5037 | 4873 | 4680 | 4861 | 2879 | 2925 | 2805 | 2925 | 3217 | 3266 | 3076 | 3375 |
| K09478 short/branched chain acyl-CoA dehydrogenase [EC:1.3.99.12] | 6 | 2 | 5 | 1 | 1 | 0 | 2 | 1 | 4 | 7 | 8 | 3 |
| K09479 very long chain acyl-CoA dehydrogenase [EC:1.3.8.9] | 0 | 1 | 0 | 1 | 1 | 1 | 3 | 1 | 2 | 1 | 0 | 0 |
| K13239 Delta3-Delta2-enoyl-CoA isomerase [EC:5.3.3.8] | 1 | 0 | 0 | 0 | 0 | 0 | 0 | 0 | 0 | 0 | 0 | 0 |
| K13767 enoyl-CoA hydratase [EC:4.2.1.17] | 0 | 1 | 0 | 0 | 0 | 0 | 0 | 0 | 0 | 1 | 0 | 0 |
| ko00072 Synthesis and degradation of ketone bodies | 16578 | 16749 | 15589 | 16376 | 9086 | 8818 | 8510 | 9209 | 9181 | 9493 | 9076 | 9436 |
| ko00100 Steroid biosynthesis | 0 | 1 | 0 | 2 | 0 | 1 | 0 | 0 | 2 | 1 | 3 | 8 |
| K12298 bile salt-stimulated lipase [EC:3.1.1.3 3.1.1.13] | 0 | 1 | 0 | 2 | 0 | 1 | 0 | 0 | 2 | 1 | 3 | 8 |
| ko00120 Primary bile acid biosynthesis | 2512 | 2686 | 2823 | 2640 | 6374 | 6506 | 6651 | 6256 | 5338 | 5106 | 5378 | 5153 |
| K01442 choloylglycine hydrolase [EC:3.5.1.24] | 2512 | 2685 | 2822 | 2640 | 6374 | 6506 | 6651 | 6256 | 5338 | 5105 | 5377 | 5153 |
| K01796 alpha-methylacyl-CoA racemase [EC:5.1.99.4] | 0 | 1 | 1 | 0 | 0 | 0 | 0 | 0 | 0 | 1 | 1 | 0 |
| ko00121 Secondary bile acid biosynthesis | 2784 | 2957 | 3101 | 2906 | 6442 | 6575 | 6715 | 6334 | 5557 | 5342 | 5614 | 5384 |
| K00076 7-alpha-hydroxysteroid dehydrogenase [EC:1.1.1.159] | 5 | 0 | 3 | 3 | 3 | 4 | 8 | 15 | 30 | 40 | 36 | 36 |
| K15868 bile acid-coenzyme A ligase [EC:6.2.1.7] | 15 | 7 | 2 | 5 | 5 | 5 | 7 | 12 | 24 | 21 | 32 | 23 |
| K22604 3alpha-hydroxycholanate dehydrogenase (NAD+) [EC:1.1.1.52] | 2 | 0 | 3 | 3 | 1 | 1 | 0 | 0 | 0 | 0 | 1 | 0 |
| K22605 3alpha-hydroxycholanate dehydrogenase (NADP+) [EC:1.1.1.392] | 0 | 0 | 1 | 0 | 0 | 0 | 1 | 0 | 1 | 0 | 1 | 1 |
| K22606 3beta-hydroxycholanate 3-dehydrogenase (NAD+) [EC:1.1.1.391] | 250 | 265 | 270 | 255 | 59 | 59 | 48 | 51 | 164 | 176 | 167 | 171 |
| ko00140 Steroid hormone biosynthesis | 6731 | 7112 | 6987 | 6939 | 13699 | 13841 | 14055 | 13839 | 12228 | 11460 | 11859 | 11357 |
| K00038 3alpha(or 20beta)-hydroxysteroid dehydrogenase [EC:1.1.1.53] | 8 | 17 | 23 | 9 | 1 | 0 | 3 | 0 | 11 | 6 | 8 | 18 |
| K01130 arylsulfatase [EC:3.1.6.1] | 6304 | 6570 | 6426 | 6461 | 12061 | 12191 | 12306 | 12246 | 11033 | 10408 | 10703 | 10185 |
| K01131 steryl-sulfatase [EC:3.1.6.2] | 0 | 0 | 0 | 0 | 0 | 0 | 0 | 0 | 1 | 10 | 4 | 7 | 7 |
| K05296 3(or 17)beta-hydroxysteroid dehydrogenase [EC:1.1.1.51] | 1 | 0 | 0 | 1 | 0 | 0 | 0 | 0 | 1 | 0 | 1 | 1 |
| K12343 3-oxo-5-alpha-steroid 4-dehydrogenase 1 [EC:1.3.1.22] | 417 | 524 | 538 | 467 | 1637 | 1648 | 1746 | 1592 | 1173 | 1042 | 1140 | 1146 |
| K15680 corticosteroid 11-beta-dehydrogenase isozyme 1 [EC:1.1.1.146] | 0 | 0 | 0 | 0 | 0 | 1 | 0 | 0 | 0 | 0 | 0 | 0 |
| ko00561 Glycerolipid metabolism | 55557 | 55846 | 54836 | 55553 | 43485 | 43921 | 43349 | 43586 | 56522 | 57228 | 55353 | 56920 |
| K00631 glycerol-3-phosphate O-acyltransferase [EC:2.3.1.15] | 5380 | 5546 | 5306 | 5526 | 3300 | 3295 | 3243 | 3293 | 3417 | 3501 | 3446 | 3469 |
| K00655 1-acyl-sn-glycerol-3-phosphate acyltransferase [EC:2.3.1.51] | 3000 | 2920 | 3125 | 2866 | 2710 | 2794 | 2705 | 2866 | 2810 | 2835 | 2743 | 2793 |
| K00864 glycerol kinase [EC:2.7.1.30] | 9915 | 9952 | 9554 | 9759 | 5312 | 5490 | 5238 | 5655 | 6381 | 6677 | 6150 | 6601 |
| K00901 diacylglycerol kinase (ATP) [EC:2.7.1.107] | 908 | 784 | 836 | 879 | 567 | 572 | 545 | 542 | 537 | 527 | 451 | 557 |
| K01046 triacylglycerol lipase [EC:3.1.1.3] | 2846 | 2685 | 2656 | 2739 | 1445 | 1388 | 1389 | 1448 | 986 | 881 | 817 | 915 |
| K01054 acylglycerol lipase [EC:3.1.1.23] | 1 | 0 | 0 | 1 | 0 | 1 | 0 | 1 | 0 | 0 | 0 | 1 |
| K03429 processive 1,2-diacylglycerol beta-glucosyltransferase [EC:2.4.1.315] | 208 | 204 | 187 | 182 | 72 | 83 | 47 | 64 | 220 | 278 | 276 | 246 |
| K03621 phosphate acyltransferase [EC:2.3.1.274] | 2248 | 2183 | 2244 | 2195 | 1169 | 1108 | 1233 | 1215 | 1595 | 1681 | 1531 | 1654 |
| K05878 phosphoenolpyruvate---glycerone phosphotransferase subunit DhaK [EC:2.7.1.121] | 1017 | 991 | 957 | 1107 | 653 | 644 | 639 | 585 | 1679 | 1702 | 1597 | 1680 |
| K05879 phosphoenolpyruvate---glycerone phosphotransferase subunit DhaL [EC:2.7.1.121] | 687 | 632 | 560 | 605 | 356 | 390 | 390 | 374 | 913 | 867 | 854 | 918 |
| K05881 phosphoenolpyruvate---glycerone phosphotransferase subunit DhaM [EC:2.7.1.121] | 890 | 966 | 907 | 1002 | 586 | 611 | 553 | 594 | 1311 | 1258 | 1326 | 1362 |
| K06117 glycerol-1-phosphatase [EC:3.1.3.21] | 1 | 4 | 0 | 3 | 0 | 5 | 5 | 3 | 12 | 10 | 8 | 11 |
| K06119 sulfoquinovosyltransferase [EC:2.4.1.-] | 0 | 0 | 0 | 0 | 0 | 0 | 0 | 0 | 0 | 1 | 1 | 1 |
| K06120 glycerol dehydratase large subunit [EC:4.2.1.30] | 661 | 741 | 600 | 728 | 193 | 190 | 186 | 223 | 1333 | 1405 | 1329 | 1349 |
| K06121 glycerol dehydratase medium subunit [EC:4.2.1.30] | 244 | 251 | 298 | 285 | 60 | 70 | 64 | 60 | 510 | 502 | 543 | 555 |

| | | | | | | | | | | | | |
|---|---|---|---|---|---|---|---|---|---|---|---|---|
| K06122 glycerol dehydratase small subunit [EC:4.2.1.30] | 176 | 177 | 213 | 184 | 66 | 47 | 47 | 52 | 389 | 357 | 292 | 337 |
| K07029 diacylglycerol kinase (ATP) [EC:2.7.1.107] | 59 | 62 | 43 | 72 | 15 | 28 | 24 | 17 | 43 | 102 | 66 | 76 |
| K08591 acyl phosphate:glycerol-3-phosphate acyltransferase [EC:2.3.1.275] | 1652 | 1663 | 1529 | 1593 | 919 | 867 | 870 | 897 | 1300 | 1325 | 1257 | 1283 |
| K13677 1,2-diacylglycerol-3-alpha-glucose alpha-1,2-glucosyltransferase [EC:2.4.1.208] | 134 | 157 | 152 | 140 | 85 | 92 | 72 | 61 | 193 | 231 | 221 | 211 |
| K18097 glycerol 2-dehydrogenase (NADP+) [EC:1.1.1.156] | 0 | 0 | 0 | 1 | 2 | 0 | 0 | 0 | 1 | 1 | 1 | 0 |
| K19002 1,2-diacylglycerol 3-alpha-glucosyltransferase [EC:2.4.1.337] | 226 | 222 | 285 | 229 | 304 | 382 | 336 | 307 | 453 | 525 | 465 | 507 |
| K19003 1,2-diacylglycerol 3-beta-glucosyltransferase [EC:2.4.1.336] | 1 | 0 | 4 | 1 | 0 | 0 | 1 | 0 | 5 | 3 | 3 | 1 |
| K19005 lipoteichoic acid synthase [EC:2.7.8.20] | 130 | 106 | 113 | 129 | 50 | 57 | 66 | 69 | 107 | 174 | 155 | 154 |
| K22251 glycerol dehydrogenase [EC:1.1.1.6] | 3 | 5 | 2 | 2 | 3 | 0 | 3 | 0 | 11 | 5 | 6 | 7 |
| ko00564 Glycerophospholipid metabolism | 77188 | 77094 | 75885 | 77077 | 65895 | 66463 | 65676 | 66179 | 70002 | 69934 | 68518 | 69693 |
| K00057 glycerol-3-phosphate dehydrogenase (NAD(P)+) [EC:1.1.1.94] | 3404 | 3354 | 3433 | 3401 | 3649 | 3771 | 3650 | 3735 | 3691 | 3369 | 3504 | 3480 |
| K00096 glycerol-1-phosphate dehydrogenase [NAD(P)+] [EC:1.1.1.261] | 257 | 242 | 316 | 296 | 182 | 199 | 191 | 170 | 632 | 631 | 602 | 611 |
| K00105 alpha-glycerophosphate oxidase [EC:1.1.3.21] | 87 | 88 | 106 | 112 | 24 | 37 | 32 | 52 | 95 | 126 | 98 | 116 |
| K00111 glycerol-3-phosphate dehydrogenase [EC:1.1.5.3] | 9039 | 8963 | 8559 | 9138 | 5136 | 4938 | 4829 | 4954 | 5979 | 5996 | 5881 | 6057 |
| K00112 glycerol-3-phosphate dehydrogenase subunit B [EC:1.1.5.3] | 779 | 745 | 664 | 789 | 534 | 581 | 549 | 538 | 1036 | 1121 | 1045 | 1140 |
| K00113 glycerol-3-phosphate dehydrogenase subunit C | 643 | 703 | 606 | 692 | 408 | 534 | 501 | 497 | 934 | 934 | 836 | 1016 |
| K00570 phosphatidylethanolamine/phosphatidyl-N-methylethanolamine N-methyltransferase [EC:2.1.1.17 2.1.1.71] | 3 | 0 | 0 | 1 | 0 | 3 | 0 | 0 | 0 | 2 | 0 | 1 |
| K00968 choline-phosphate cytidylyltransferase [EC:2.7.7.15] | 2 | 0 | 0 | 0 | 3 | 7 | 5 | 2 | 7 | 9 | 4 | 11 |
| K00980 glycerol-3-phosphate cytidylyltransferase [EC:2.7.7.39] | 128 | 112 | 121 | 117 | 122 | 135 | 154 | 134 | 69 | 72 | 65 | 66 |
| K00981 phosphatidate cytidylyltransferase [EC:2.7.7.41] | 4288 | 4498 | 4412 | 4303 | 5619 | 5594 | 5414 | 5341 | 4725 | 4684 | 4710 | 4563 |
| K00995 CDP-diacylglycerol---glycerol-3-phosphate 3-phosphatidyltransferase [EC:2.7.8.5] | 2497 | 2587 | 2647 | 2492 | 2480 | 2396 | 2450 | 2518 | 2057 | 2024 | 2017 | 2075 |
| K00998 CDP-diacylglycerol---serine O-phosphatidyltransferase [EC:2.7.8.8] | 792 | 787 | 797 | 785 | 497 | 629 | 570 | 514 | 1092 | 1170 | 1179 | 1107 |
| K01004 phosphatidylcholine synthase [EC:2.7.8.24] | 1093 | 1075 | 1019 | 1055 | 610 | 611 | 575 | 605 | 338 | 366 | 320 | 332 |
| K01048 lysophospholipase [EC:3.1.1.5] | 730 | 681 | 699 | 776 | 552 | 529 | 541 | 444 | 975 | 1046 | 1005 | 1073 |
| K01058 phospholipase A1/A2 [EC:3.1.1.32 3.1.1.4] | 506 | 600 | 573 | 624 | 373 | 403 | 400 | 382 | 768 | 767 | 775 | 839 |
| K01095 phosphatidylglycerophosphatase A [EC:3.1.3.27] | 1286 | 1315 | 1254 | 1325 | 1485 | 1451 | 1476 | 1513 | 1355 | 1531 | 1346 | 1275 |
| K01096 phosphatidylglycerophosphatase B [EC:3.1.3.27 3.1.3.81 3.1.3.4 3.6.1.27] | 476 | 475 | 467 | 485 | 269 | 341 | 291 | 302 | 786 | 832 | 708 | 714 |
| K01115 phospholipase D1/2 [EC:3.1.4.4] | 0 | 0 | 0 | 0 | 0 | 0 | 0 | 0 | 0 | 0 | 1 | 0 |
| K01126 glycerophosphoryl diester phosphodiesterase [EC:3.1.4.46] | 10729 | 10609 | 10677 | 10476 | 13418 | 13053 | 13563 | 13347 | 12072 | 11885 | 11814 | 11854 |
| K01521 CDP-diacylglycerol pyrophosphatase [EC:3.6.1.26] | 463 | 456 | 462 | 501 | 338 | 379 | 306 | 375 | 677 | 704 | 626 | 679 |
| K01613 phosphatidylserine decarboxylase [EC:4.1.1.65] | 2101 | 2214 | 2222 | 2138 | 2511 | 2566 | 2603 | 2499 | 2540 | 2529 | 2457 | 2424 |
| K03735 ethanolamine ammonia-lyase large subunit [EC:4.3.1.7] | 3617 | 3521 | 3472 | 3614 | 1959 | 1924 | 1872 | 1928 | 3009 | 2967 | 2743 | 2784 |
| K03736 ethanolamine ammonia-lyase small subunit [EC:4.3.1.7] | 2089 | 1998 | 1899 | 1919 | 1039 | 1008 | 986 | 1051 | 1785 | 1810 | 1786 | 1879 |
| K04019 ethanolamine utilization protein EutA | 632 | 660 | 637 | 693 | 170 | 176 | 178 | 152 | 1209 | 1256 | 1267 | 1408 |
| K06131 cardiolipin synthase A/B [EC:2.7.8.-] | 6648 | 6775 | 6634 | 6800 | 7975 | 8246 | 8057 | 7929 | 8213 | 8151 | 8305 | 8149 |
| K06132 cardiolipin synthase C [EC:2.7.8.-] | 3218 | 3215 | 3112 | 3166 | 1911 | 1907 | 1917 | 2208 | 1262 | 1269 | 1212 | 1301 |
| K08744 cardiolipin synthase (CMP-forming) [EC:2.7.8.41] | 244 | 256 | 272 | 248 | 112 | 117 | 97 | 133 | 323 | 273 | 309 | 293 |
| K13333 lysophospholipase [EC:3.1.1.5] | 0 | 0 | 0 | 0 | 0 | 0 | 0 | 0 | 2 | 0 | 0 | 0 |
| K13535 cardiolipin-specific phospholipase [EC:3.1.1.-] | 0 | 0 | 0 | 0 | 0 | 1 | 0 | 0 | 0 | 0 | 0 | 1 |
| K13622 S-adenosylmethionine-diacylglycerol 3-amino-3-carboxypropyl transferase | 0 | 0 | 3 | 0 | 0 | 4 | 0 | 0 | 0 | 4 | 2 | 3 |
| K13623 S-adenosylmethionine-diacylgycerolhomoserine-N-methlytransferase | 0 | 0 | 0 | 0 | 0 | 0 | 0 | 2 | 0 | 2 | 0 | 1 |
| K17103 CDP-diacylglycerol---serine O-phosphatidyltransferase [EC:2.7.8.8] | 2051 | 1856 | 2184 | 2020 | 2455 | 2446 | 2492 | 2366 | 1806 | 1746 | 1710 | 1702 |
| K17104 phosphoglycerol geranylgeranyltransferase [EC:2.5.1.41] | 0 | 0 | 0 | 1 | 2 | 0 | 0 | 0 | 0 | 1 | 2 | 11 |
| K17105 geranylgeranylglycerol-phosphate geranylgeranyltransferase [EC:2.5.1.42] | 0 | 0 | 0 | 1 | 0 | 0 | 0 | 0 | 8 | 6 | 1 | 9 |
| K17830 digeranylgeranylglycerophospholipid reductase [EC:1.3.1.101 1.3.7.11] | 1 | 0 | 0 | 1 | 3 | 0 | 3 | 0 | 4 | 6 | 6 | 3 |
| K18695 glycerophosphocholine phosphodiesterase GPCPD1 [EC:3.1.4.2] | 0 | 1 | 0 | 0 | 0 | 0 | 0 | 0 | 1 | 0 | 0 | 0 |
| K18697 phosphatidylglycerophosphatase C [EC:3.1.3.27] | 441 | 400 | 385 | 395 | 262 | 302 | 289 | 273 | 542 | 527 | 517 | 572 |
| K19664 CDP-2,3-bis-(O-geranylgeranyl)-sn-glycerol synthase [EC:2.7.7.67] | 0 | 2 | 0 | 1 | 2 | 2 | 0 | 0 | 0 | 8 | 4 | 5 |
| K19665 archaetidylserine synthase [EC:2.7.8.38] | 0 | 0 | 0 | 0 | 0 | 2 | 0 | 0 | 2 | 3 | 0 | 3 |
| ko00565 Ether lipid metabolism | 9206 | 9409 | 8708 | 9131 | 4871 | 4989 | 4837 | 5257 | 3592 | 3364 | 3216 | 3687 |
| K00803 alkyldihydroxyacetonephosphate synthase [EC:2.5.1.26] | 2186 | 2237 | 2037 | 2054 | 1110 | 1046 | 1085 | 1241 | 681 | 643 | 632 | 778 |
| K22387 lysophospholipase D [EC:3.1.4.39] | 1 | 0 | 0 | 0 | 0 | 0 | 0 | 0 | 0 | 0 | 0 | 0 |
| ko00600 Sphingolipid metabolism | 43452 | 45719 | 47280 | 45329 | 92594 | 92965 | 93944 | 91652 | 79267 | 76794 | 78525 | 76530 |
| K01134 arylsulfatase A [EC:3.1.6.8] | 4 | 2 | 2 | 1 | 1 | 0 | 1 | 2 | 3 | 0 | 0 | 0 |
| K01186 sialidase-1 [EC:3.2.1.18] | 2176 | 2319 | 2579 | 2298 | 7677 | 7901 | 8247 | 7864 | 6136 | 5760 | 6039 | 5759 |
| K01201 glucosylceramidase [EC:3.2.1.45] | 1191 | 1264 | 1341 | 1326 | 2858 | 2834 | 3006 | 2823 | 2322 | 2407 | 2435 | 2307 |
| K01202 galactosylceramidase [EC:3.2.1.46] | 0 | 0 | 0 | 0 | 0 | 0 | 0 | 0 | 0 | 3 | 3 | 2 |
| K04708 3-dehydrosphinganine reductase [EC:1.1.1.102] | 2 | 2 | 2 | 2 | 0 | 0 | 0 | 0 | 3 | 5 | 0 | 1 |
| K12349 neutral ceramidase [EC:3.5.1.23] | 1013 | 1084 | 1084 | 1083 | 591 | 584 | 534 | 594 | 349 | 336 | 342 | 310 |
| K17108 non-lysosomal glucosylceramidase [EC:3.2.1.45] | 4 | 3 | 1 | 3 | 9 | 3 | 4 | 13 | 0 | 0 | 2 | 0 |
| ko00590 Arachidonic acid metabolism | 6272 | 6577 | 6237 | 6248 | 4543 | 4560 | 4500 | 4880 | 4094 | 3858 | 4112 | 3981 |
| K00432 glutathione peroxidase [EC:1.11.1.9] | 3214 | 3293 | 3268 | 3173 | 2836 | 2907 | 2880 | 3011 | 2566 | 2388 | 2579 | 2357 |
| K18592 gamma-glutamyltranspeptidase / glutathione hydrolase / leukotriene-C4 hydrolase [EC:2.3.2.2 3.4.19.13 3.4.19.14] | 0 | 0 | 0 | 0 | 0 | 0 | 0 | 0 | 0 | 1 | 0 | 1 |
| K19246 arachidonate 15-lipoxygenase [EC:1.13.11.33] | 2551 | 2684 | 2396 | 2450 | 1334 | 1250 | 1220 | 1487 | 760 | 702 | 757 | 783 |
| K23486 9,11-endoperoxide prostaglandin H2 reductase [EC:1.1.1.-] | 1 | 0 | 0 | 1 | 0 | 0 | 0 | 0 | 0 | 0 | 1 | 1 |
| ko00591 Linoleic acid metabolism | 1600 | 1779 | 1452 | 1725 | 975 | 935 | 954 | 1024 | 1128 | 1105 | 1087 | 1195 |
| K00508 linoleoyl-CoA desaturase [EC:1.14.19.3] | 1094 | 1179 | 879 | 1101 | 602 | 532 | 554 | 642 | 360 | 338 | 312 | 356 |
| ko00592 alpha-Linolenic acid metabolism | 7277 | 7298 | 6975 | 7260 | 4404 | 4295 | 4323 | 4307 | 4583 | 4492 | 4436 | 4678 |
| K10526 OPC-8:0 CoA ligase 1 [EC:6.2.1.-] | 0 | 0 | 0 | 0 | 0 | 0 | 0 | 0 | 0 | 1 | 0 | 0 |
| ko01040 Biosynthesis of unsaturated fatty acids | 3612 | 3527 | 3206 | 3417 | 2253 | 2179 | 2077 | 2167 | 2306 | 2323 | 2265 | 2300 |
| K00507 stearoyl-CoA desaturase (Delta-9 desaturase) [EC:1.14.19.1] | 1 | 0 | 0 | 2 | 0 | 0 | 0 | 0 | 0 | 0 | 0 | 0 |
| K10804 acyl-CoA thioesterase I [EC:3.1.2.- 3.1.2.2 3.1.1.2 3.1.1.5] | 1295 | 1250 | 1073 | 1161 | 804 | 759 | 722 | 798 | 772 | 799 | 764 | 813 |
| K10805 acyl-CoA thioesterase II [EC:3.1.2.-] | 1421 | 1451 | 1311 | 1448 | 932 | 889 | 814 | 879 | 1019 | 1060 | 980 | 987 |
| K10806 acyl-CoA thioesterase YciA [EC:3.1.2.-] | 895 | 824 | 822 | 806 | 515 | 531 | 541 | 490 | 515 | 464 | 516 | 500 |
| Nucleotide metabolism | 313861 | 313109 | 312215 | 313931 | 302565 | 306153 | 303551 | 302813 | 312106 | 311970 | 308504 | 310587 |
| ko00230 Purine metabolism | 232710 | 232578 | 231060 | 232794 | 215359 | 217957 | 216249 | 216099 | 224942 | 224048 | 221457 | 223084 |
| K00073 ureidoglycolate dehydrogenase (NAD+) [EC:1.1.1.350] | 105 | 97 | 90 | 114 | 33 | 47 | 30 | 28 | 136 | 156 | 163 | 144 |
| K00087 xanthine dehydrogenase molybdenum-binding subunit [EC:1.17.1.4] | 1397 | 1235 | 1202 | 1161 | 450 | 511 | 579 | 427 | 1154 | 1155 | 1139 | 1178 |
| K00088 IMP dehydrogenase [EC:1.1.1.205] | 6113 | 6256 | 6328 | 6186 | 5917 | 5929 | 5688 | 5877 | 6696 | 6633 | 6656 | 6447 |
| K00106 xanthine dehydrogenase/oxidase [EC:1.17.1.4 1.17.3.2] | 0 | 2 | 0 | 0 | 0 | 4 | 0 | 0 | 0 | 0 | 0 | 0 |
| K00364 GMP reductase [EC:1.7.1.7] | 851 | 748 | 793 | 795 | 546 | 649 | 569 | 568 | 967 | 1054 | 1017 | 1132 |
| K00525 ribonucleoside-diphosphate reductase alpha chain [EC:1.17.4.1] | 13702 | 13778 | 13670 | 13792 | 12280 | 12707 | 12530 | 12324 | 11834 | 11674 | 11872 | 11631 |
| K00526 ribonucleoside-diphosphate reductase beta chain [EC:1.17.4.1] | 3364 | 3435 | 3494 | 3409 | 2172 | 2083 | 2073 | 2205 | 2985 | 3126 | 2896 | 2972 |
| K00527 ribonucleoside-triphosphate reductase (thioredoxin) [EC:1.17.4.2] | 0 | 4 | 5 | 0 | 0 | 0 | 0 | 0 | 0 | 0 | 3 | 0 |
| K00602 phosphoribosylaminoimidazolecarboxamide formyltransferase / IMP cyclohydrolase [EC:2.1.2.3 3.5.4.10] | 6501 | 6459 | 6298 | 6439 | 8668 | 8594 | 8740 | 8669 | 7719 | 7319 | 7267 | 7176 |
| K00759 adenine phosphoribosyltransferase [EC:2.4.2.7] | 1881 | 1730 | 1859 | 1756 | 2044 | 2048 | 2018 | 2092 | 2126 | 2020 | 2115 | 2097 |
| K00760 hypoxanthine phosphoribosyltransferase [EC:2.4.2.8] | 1754 | 1724 | 1747 | 1742 | 1867 | 2028 | 1899 | 1948 | 1907 | 1905 | 1916 | 1783 |
| K00764 amidophosphoribosyltransferase [EC:2.4.2.14] | 6666 | 6934 | 6999 | 6655 | 8994 | 8899 | 9306 | 8904 | 8218 | 8390 | 8157 | 7922 |
| K00769 xanthine phosphoribosyltransferase [EC:2.4.2.22] | 366 | 263 | 330 | 309 | 200 | 232 | 228 | 201 | 386 | 457 | 453 | 450 |
| K00839 (S)-ureidoglycine---glyoxylate transaminase [EC:2.6.1.112] | 778 | 856 | 701 | 773 | 489 | 545 | 501 | 550 | 924 | 1048 | 932 | 1000 |
| K00856 adenosine kinase [EC:2.7.1.20] | 21 | 23 | 33 | 23 | 28 | 27 | 26 | 32 | 29 | 36 | 41 | 25 |
| K00892 inosine kinase [EC:2.7.1.73] | 762 | 853 | 789 | 733 | 556 | 557 | 524 | 551 | 1097 | 1159 | 1098 | 1121 |
| K00939 adenylate kinase [EC:2.7.4.3] | 2278 | 2392 | 2287 | 2309 | 2317 | 2314 | 2309 | 2307 | 2295 | 2302 | 2241 | 2242 |
| K00940 nucleoside-diphosphate kinase [EC:2.7.4.6] | 1293 | 1297 | 1283 | 1331 | 1669 | 1584 | 1622 | 1657 | 1383 | 1281 | 1352 | 1408 |
| K00942 guanylate kinase [EC:2.7.4.8] | 2224 | 2361 | 2259 | 2284 | 2343 | 2319 | 2419 | 2373 | 2354 | 2243 | 2421 | 2309 |
| K00951 GTP pyrophosphokinase [EC:2.7.6.5] | 8335 | 8795 | 8948 | 8781 | 12489 | 12533 | 12563 | 12401 | 10402 | 10539 | 10661 | 10155 |
| K01081 5'-nucleotidase [EC:3.1.3.5] | 601 | 692 | 723 | 572 | 2066 | 1866 | 1899 | 1819 | 1548 | 1413 | 1592 | 1529 |
| K01119 2',3'-cyclic-nucleotide 2'-phosphodiesterase / 3'-nucleotidase [EC:3.1.4.16 3.1.3.6] | 2063 | 2122 | 2135 | 2070 | 2992 | 3075 | 2865 | 2958 | 3265 | 3362 | 3009 | 3238 |
| K01129 dGTPase [EC:3.1.5.1] | 5928 | 5682 | 5651 | 5698 | 5441 | 5507 | 5437 | 5429 | 5223 | 5168 | 5054 | 5183 |
| K01139 GTP diphosphokinase / guanosine-3',5'-bis(diphosphate) 3'-diphosphatase [EC:2.7.6.5 3.1.7.2] | 2661 | 2580 | 2551 | 2683 | 1714 | 1888 | 1727 | 1802 | 2428 | 2485 | 2366 | 2480 |
| K01239 purine nucleosidase [EC:3.2.2.1] | 2083 | 2000 | 1842 | 1966 | 1081 | 1028 | 996 | 1081 | 976 | 960 | 833 | 871 |
| K01241 AMP nucleosidase [EC:3.2.2.4] | 3632 | 3717 | 3515 | 3590 | 3442 | 3374 | 3221 | 3261 | 2988 | 2834 | 2780 | 2887 |
| K01427 urease [EC:3.5.1.5] | 0 | 1 | 2 | 1 | 0 | 0 | 0 | 1 | 0 | 0 | 0 | 0 |
| K01428 urease subunit alpha [EC:3.5.1.5] | 3987 | 3782 | 3507 | 3779 | 1882 | 1699 | 1728 | 1859 | 2884 | 2892 | 2701 | 2946 |
| K01429 urease subunit beta [EC:3.5.1.5] | 660 | 709 | 630 | 611 | 318 | 310 | 294 | 348 | 402 | 401 | 419 | 446 |
| K01430 urease subunit gamma [EC:3.5.1.5] | 584 | 622 | 555 | 574 | 257 | 295 | 268 | 293 | 439 | 443 | 452 | 474 |
| K01466 allantoinase [EC:3.5.2.5] | 334 | 292 | 304 | 325 | 149 | 192 | 156 | 172 | 324 | 335 | 318 | 327 |
| K01477 allantoicase [EC:3.5.3.4] | 1509 | 1516 | 1417 | 1468 | 825 | 762 | 787 | 820 | 494 | 407 | 431 | 428 |
| K01483 ureidoglycolate lyase [EC:4.3.2.3] | 777 | 794 | 715 | 718 | 451 | 412 | 390 | 371 | 234 | 227 | 249 | 252 |
| K01486 adenine deaminase [EC:3.5.4.2] | 2414 | 2266 | 2294 | 2344 | 624 | 587 | 612 | 596 | 2638 | 3008 | 2783 | 3082 |
| K01487 guanine deaminase [EC:3.5.4.3] | 4282 | 4148 | 4110 | 4335 | 2042 | 2144 | 2015 | 2100 | 2466 | 2586 | 2269 | 2508 |
| K01488 adenosine deaminase [EC:3.5.4.4] | 1348 | 1350 | 1330 | 1429 | 549 | 561 | 616 | 517 | 1394 | 1573 | 1511 | 1534 |
| K01515 ADP-ribose pyrophosphatase [EC:3.6.1.13] | 1708 | 1705 | 1696 | 1753 | 942 | 862 | 1051 | 1030 | 1202 | 1228 | 1128 | 1284 |
| K01518 bis(5'-nucleosidyl)-tetraphosphatase [EC:3.6.1.17] | 0 | 0 | 0 | 0 | 0 | 0 | 0 | 1 | 2 | 0 | 1 | 0 |
| K01524 exopolyphosphatase / guanosine-5'-triphosphate,3'-diphosphate pyrophosphatase [EC:3.6.1.11 3.6.1.40] | 5769 | 5686 | 5729 | 5637 | 3508 | 3685 | 3614 | 3602 | 4826 | 4859 | 4863 | 4873 |
| K01525 bis(5'-nucleosyl)-tetraphosphatase (symmetrical) [EC:3.6.1.41] | 1844 | 1661 | 1791 | 1880 | 1138 | 1130 | 1117 | 1163 | 1277 | 1188 | 1217 | 1218 |
| K01588 5-(carboxyamino)imidazole ribonucleotide mutase [EC:5.4.99.18] | 1580 | 1637 | 1637 | 1564 | 1756 | 1871 | 1753 | 1800 | 1900 | 1711 | 1818 | 1767 |
| K01589 5-(carboxyamino)imidazole ribonucleotide synthase [EC:6.3.4.18] | 1984 | 1984 | 1810 | 1889 | 1216 | 1312 | 1160 | 1313 | 1568 | 1437 | 1575 | 1488 |
| K01756 adenylosuccinate lyase [EC:4.3.2.2] | 4686 | 4570 | 4836 | 4919 | 5238 | 5319 | 5155 | 5028 | 5016 | 5066 | 4726 | 4872 |
| K01768 adenylate cyclase [EC:4.6.1.1] | 1713 | 1620 | 1470 | 1628 | 874 | 800 | 827 | 846 | 577 | 476 | 507 | 508 |
| K01923 phosphoribosylaminoimidazole-succinocarboxamide synthase [EC:6.3.2.6] | 2751 | 2713 | 2651 | 2728 | 3209 | 3193 | 3283 | 3222 | 3071 | 2993 | 2966 | 2975 |

| | | | | | | | | | | | | |
|---|---|---|---|---|---|---|---|---|---|---|---|---|
| K01933 phosphoribosylformylglycinamidine cyclo-ligase [EC:6.3.3.1] | 3845 | 3888 | 3914 | 4006 | 4236 | 4459 | 4139 | 4206 | 4268 | 4279 | 4371 | 4262 |
| K01939 adenylosuccinate synthase [EC:6.3.4.4] | 4749 | 4755 | 4893 | 4816 | 5048 | 5100 | 4981 | 4927 | 4913 | 4993 | 4954 | 5049 |
| K01945 phosphoribosylamine---glycine ligase [EC:6.3.4.13] | 4766 | 4817 | 4698 | 4624 | 5075 | 5203 | 5160 | 5270 | 4978 | 4924 | 5054 | 4804 |
| K01951 GMP synthase (glutamine-hydrolysing) [EC:6.3.5.2] | 7368 | 7456 | 7759 | 7400 | 9383 | 9471 | 9336 | 9310 | 8978 | 8745 | 8786 | 8640 |
| K01952 phosphoribosylformylglycinamidine synthase [EC:6.3.5.3] | 12215 | 12259 | 12273 | 12342 | 13001 | 13269 | 13457 | 13621 | 12170 | 11877 | 11837 | 12247 |
| K02083 allantoate deiminase [EC:3.5.3.9] | 841 | 833 | 793 | 829 | 517 | 602 | 526 | 504 | 970 | 1110 | 1022 | 1052 |
| K02428 XTP/dITP diphosphohydrolase [EC:3.6.1.66] | 2249 | 2296 | 2426 | 2300 | 3759 | 3755 | 3788 | 3673 | 3398 | 3276 | 3362 | 3300 |
| K03651 3',5'-cyclic-AMP phosphodiesterase [EC:3.1.4.53] | 2371 | 2281 | 2089 | 2282 | 1228 | 1218 | 1255 | 1392 | 1983 | 1897 | 1829 | 1988 |
| K03783 purine-nucleoside phosphorylase [EC:2.4.2.1] | 973 | 1038 | 1075 | 993 | 1973 | 2101 | 2051 | 1976 | 1723 | 1673 | 1671 | 1749 |
| K03784 purine-nucleoside phosphorylase [EC:2.4.2.1] | 868 | 985 | 869 | 1002 | 591 | 631 | 509 | 517 | 965 | 1076 | 1039 | 1019 |
| K03787 5'-nucleotidase [EC:3.1.3.5] | 1497 | 1639 | 1662 | 1567 | 2344 | 2396 | 2515 | 2301 | 2187 | 2128 | 2139 | 2191 |
| K03815 xanthosine phosphorylase [EC:2.4.2.-] | 3 | 0 | 3 | 0 | 3 | 12 | 9 | 16 | 34 | 65 | 66 | 42 |
| K03816 xanthine phosphoribosyltransferase [EC:2.4.2.22] | 1731 | 1816 | 1755 | 1746 | 1795 | 1838 | 1924 | 1799 | 1565 | 1552 | 1527 | 1418 |
| K04765 nucleoside triphosphate diphosphatase [EC:3.6.1.9] | 703 | 695 | 699 | 766 | 494 | 503 | 476 | 458 | 704 | 742 | 643 | 837 |
| K05851 adenylate cyclase, class 1 [EC:4.6.1.1] | 6311 | 6262 | 5805 | 6160 | 3757 | 3673 | 3565 | 3734 | 3676 | 3624 | 3452 | 3589 |
| K05873 adenylate cyclase, class 2 [EC:4.6.1.1] | 0 | 5 | 5 | 7 | 8 | 2 | 3 | 3 | 0 | 9 | 13 | 7 |
| K06863 5-formaminoimidazole-4-carboxamide-1-(beta)-D-ribofuranosyl 5'-monophosphate synthetase [EC:6.3.4.23] | 0 | 0 | 2 | 1 | 0 | 2 | 0 | 0 | 7 | 9 | 6 | 15 |
| K06928 nucleoside-triphosphatase [EC:3.6.1.15] | 1 | 0 | 0 | 0 | 12 | 18 | 15 | 20 | 3 | 0 | 3 | 9 |
| K06966 pyrimidine/purine-5'-nucleotide nucleosidase [EC:3.2.2.10 3.2.2.-] | 929 | 1029 | 974 | 976 | 650 | 657 | 641 | 707 | 1261 | 1242 | 1230 | 1346 |
| K07127 5-hydroxyisourate hydrolase [EC:3.5.2.17] | 944 | 944 | 851 | 901 | 487 | 517 | 469 | 476 | 744 | 850 | 729 | 834 |
| K07816 putative GTP pyrophosphokinase [EC:2.7.6.5] | 1089 | 1086 | 1222 | 1117 | 315 | 332 | 293 | 286 | 1116 | 1201 | 1183 | 1185 |
| K08289 phosphoribosylglycinamide formyltransferase 2 [EC:2.1.2.2] | 2128 | 2104 | 2047 | 2107 | 2336 | 2350 | 2356 | 2228 | 2495 | 2475 | 2450 | 2507 |
| K08312 ADP-ribose diphosphatase [EC:3.6.1.-] | 1128 | 1004 | 1100 | 1154 | 689 | 729 | 692 | 625 | 735 | 648 | 753 | 665 |
| K08693 2',3'-cyclic-nucleotide 2'-phosphodiesterase / 3'-nucleotidase / 5'-nucleotidase [EC:3.1.4.16 3.1.3.6 3.1.3.5] | 0 | 0 | 1 | 0 | 0 | 0 | 0 | 0 | 1 | 0 | 3 | 1 |
| K08723 5'-nucleotidase [EC:3.1.3.5] | 433 | 488 | 433 | 460 | 303 | 312 | 318 | 312 | 546 | 649 | 526 | 623 |
| K09913 purine/pyrimidine-nucleoside phosphorylase [EC:2.4.2.1 2.4.2.2] | 554 | 642 | 623 | 593 | 368 | 320 | 340 | 368 | 407 | 420 | 382 | 442 |
| K10213 ribosylpyrimidine nucleosidase [EC:3.2.2.8] | 164 | 168 | 154 | 159 | 231 | 247 | 223 | 253 | 31 | 61 | 59 | 58 |
| K10353 deoxyadenosine kinase [EC:2.7.1.76] | 0 | 0 | 0 | 0 | 0 | 3 | 0 | 2 | 1 | 0 | 0 | 0 |
| K10807 ribonucleoside-diphosphate reductase subunit M1 [EC:1.17.4.1] | 0 | 0 | 0 | 0 | 0 | 0 | 0 | 0 | 0 | 0 | 0 | 3 |
| K11175 phosphoribosylglycinamide formyltransferase 1 [EC:2.1.2.2] | 2241 | 2257 | 2294 | 2276 | 2476 | 2529 | 2566 | 2446 | 2400 | 2427 | 2297 | 2273 |
| K11176 IMP cyclohydrolase [EC:3.5.4.10] | 0 | 5 | 0 | 0 | 2 | 0 | 0 | 0 | 0 | 3 | 5 | 2 |
| K11177 xanthine dehydrogenase YagR molybdenum-binding subunit [EC:1.17.1.4] | 3204 | 3053 | 3060 | 3008 | 1685 | 1584 | 1622 | 1687 | 901 | 845 | 966 | 942 |
| K11178 xanthine dehydrogenase YagS FAD-binding subunit [EC:1.17.1.4] | 1319 | 1325 | 1181 | 1242 | 672 | 634 | 663 | 649 | 331 | 319 | 313 | 399 |
| K11751 5'-nucleotidase / UDP-sugar diphosphatase [EC:3.1.3.5 3.6.1.45] | 2851 | 2774 | 2825 | 2851 | 988 | 1083 | 967 | 938 | 3132 | 3277 | 3245 | 3338 |
| K11788 phosphoribosylamine--glycine ligase / phosphoribosylformylglycinamidine cyclo-ligase [EC:6.3.4.13 6.3.3.1] | 1 | 0 | 1 | 1 | 0 | 1 | 0 | 2 | 2 | 0 | 3 | 1 |
| K13479 xanthine dehydrogenase FAD-binding subunit [EC:1.17.1.4] | 465 | 435 | 428 | 459 | 110 | 134 | 128 | 144 | 372 | 464 | 405 | 434 |
| K13480 xanthine dehydrogenase iron-sulfur-binding subunit | 2 | 8 | 7 | 3 | 8 | 5 | 11 | 11 | 25 | 25 | 24 | 38 |
| K13481 xanthine dehydrogenase small subunit [EC:1.17.1.4] | 1804 | 1887 | 1875 | 1885 | 1019 | 1047 | 978 | 1081 | 639 | 556 | 516 | 612 |
| K13482 xanthine dehydrogenase large subunit [EC:1.17.1.4] | 3438 | 3542 | 3398 | 3534 | 1833 | 1840 | 1816 | 1983 | 1123 | 1089 | 953 | 1095 |
| K13483 xanthine dehydrogenase YagT iron-sulfur-binding subunit | 642 | 707 | 590 | 631 | 309 | 372 | 330 | 380 | 205 | 184 | 192 | 210 |
| K14048 urease subunit gamma/beta [EC:3.5.1.5] | 13 | 23 | 15 | 16 | 4 | 5 | 3 | 1 | 33 | 34 | 24 | 49 |
| K14642 golgi apyrase [EC:3.6.1.5] | 0 | 0 | 0 | 0 | 0 | 0 | 0 | 0 | 0 | 0 | 2 | 0 |
| K14977 (S)-ureidoglycine aminohydrolase [EC:3.5.3.26] | 1534 | 1319 | 1269 | 1377 | 755 | 776 | 692 | 841 | 600 | 585 | 497 | 589 |
| K15518 deoxyguanosine kinase [EC:2.7.1.113] | 2 | 0 | 0 | 0 | 0 | 0 | 0 | 0 | 0 | 0 | 0 | 0 |
| K15519 deoxyadenosine/deoxycytidine kinase [EC:2.7.1.76 2.7.1.74] | 2 | 0 | 0 | 0 | 0 | 0 | 0 | 0 | 0 | 0 | 0 | 0 |
| K15780 bifunctional protein TilS/HprT [EC:6.3.4.19 2.4.2.8] | 0 | 0 | 1 | 0 | 0 | 0 | 0 | 0 | 0 | 0 | 2 | 1 |
| K16839 FAD-dependent urate hydroxylase [EC:1.14.13.113] | 691 | 717 | 599 | 666 | 450 | 454 | 427 | 468 | 930 | 996 | 884 | 972 |
| K16840 2-oxo-4-hydroxy-4-carboxy-5-ureidoimidazoline decarboxylase [EC:4.1.1.97] | 267 | 263 | 238 | 276 | 166 | 184 | 214 | 185 | 377 | 372 | 404 | 410 |
| K16841 allantoin racemase [EC:5.1.99.3] | 414 | 428 | 422 | 481 | 278 | 302 | 333 | 264 | 630 | 586 | 564 | 625 |
| K16842 allantoinase [EC:3.5.2.5] | 544 | 544 | 491 | 562 | 394 | 364 | 375 | 359 | 593 | 656 | 579 | 641 |
| K16856 ureidoglycolate lyase [EC:4.3.2.3] | 0 | 0 | 0 | 1 | 0 | 0 | 0 | 0 | 0 | 0 | 0 | 0 |
| K18151 ureidoglycolate amidohydrolase [EC:3.5.1.116] | 7 | 2 | 5 | 5 | 0 | 1 | 1 | 0 | 3 | 2 | 2 | 0 |
| K18532 adenylate kinase [EC:2.7.4.3] | 0 | 0 | 0 | 0 | 2 | 0 | 0 | 0 | 2 | 4 | 0 | 10 |
| K19572 adenosine deaminase CECR1 [EC:3.5.4.4] | 0 | 0 | 0 | 1 | 0 | 0 | 0 | 0 | 1 | 0 | 0 | 1 |
| K19710 ATP adenylyltransferase [EC:2.7.7.53] | 6 | 5 | 3 | 3 | 0 | 4 | 0 | 1 | 4 | 13 | 4 | 9 |
| K20881 5'-nucleotidase [EC:3.1.3.5] | 1623 | 1681 | 1635 | 1586 | 972 | 1035 | 963 | 1047 | 1004 | 1011 | 951 | 1006 |
| K21053 adenine deaminase [EC:3.5.4.2] | 1663 | 1684 | 1419 | 1586 | 849 | 816 | 805 | 890 | 525 | 463 | 465 | 487 |
| K21636 ribonucleoside-triphosphate reductase (formate) [EC:1.1.98.6] | 7675 | 7450 | 7766 | 7789 | 9242 | 9496 | 9482 | 9090 | 9724 | 9824 | 9930 | 9508 |
| K22026 nucleoside kinase [EC:2.7.1.73 2.7.1.213 2.7.1.-] | 2 | 0 | 0 | 0 | 2 | 2 | 0 | 1 | 4 | 4 | 4 | 8 |
| K22601 oxamate carbamoyltransferase [EC:2.1.3.5] | 686 | 686 | 724 | 748 | 505 | 550 | 498 | 612 | 824 | 843 | 807 | 909 |
| K22602 oxamate amidohydrolase [EC:3.5.1.126] | 956 | 959 | 899 | 896 | 608 | 652 | 654 | 621 | 1278 | 1195 | 1158 | 1245 |
| K23264 phosphoribosylformylglycinamidine synthase subunit PurS [EC:6.3.5.3] | 14 | 16 | 15 | 16 | 2 | 5 | 4 | 2 | 19 | 35 | 19 | 15 |
| K23265 phosphoribosylformylglycinamidine synthase subunit PurQ / glutaminase [EC:6.3.5.3 3.5.1.2] | 36 | 43 | 47 | 40 | 16 | 6 | 19 | 15 | 50 | 70 | 53 | 51 |
| K23269 phosphoribosylformylglycinamidine synthase subunit PurL [EC:6.3.5.3] | 115 | 126 | 122 | 149 | 30 | 40 | 29 | 16 | 133 | 160 | 137 | 158 |
| K23270 phosphoribosylformylglycinamidine synthase subunit PurSL [EC:6.3.5.3] | 36 | 26 | 33 | 26 | 5 | 8 | 6 | 11 | 31 | 51 | 33 | 31 |
| ko00240 Pyrimidine metabolism | 120448 | 120448 | 121181 | 121053 | 126543 | 128294 | 126776 | 125646 | 129870 | 130885 | 129597 | 130410 |
| K00226 dihydroorotate dehydrogenase (fumarate) [EC:1.3.98.1] | 564 | 580 | 641 | 647 | 1926 | 1835 | 1959 | 1870 | 1445 | 1374 | 1317 | 1250 |
| K00254 dihydroorotate dehydrogenase [EC:1.3.5.2] | 1807 | 1666 | 1607 | 1751 | 1165 | 1114 | 1060 | 1044 | 1148 | 1259 | 1176 | 1240 |
| K00560 thymidylate synthase [EC:2.1.1.45] | 2636 | 2575 | 2688 | 2634 | 2925 | 2856 | 2896 | 2814 | 2874 | 2808 | 2929 | 2870 |
| K00609 aspartate carbamoyltransferase catalytic subunit [EC:2.1.3.2] | 2939 | 2896 | 2850 | 2874 | 3021 | 3172 | 3208 | 3139 | 2955 | 2907 | 3075 | 3068 |
| K00610 aspartate carbamoyltransferase regulatory subunit | 769 | 735 | 804 | 794 | 1095 | 1202 | 1096 | 1110 | 1211 | 1227 | 1249 | 1201 |
| K00756 pyrimidine-nucleoside phosphorylase [EC:2.4.2.2] | 463 | 442 | 446 | 439 | 81 | 93 | 84 | 97 | 330 | 387 | 360 | 349 |
| K00757 uridine phosphorylase [EC:2.4.2.3] | 2183 | 2103 | 2314 | 2113 | 2573 | 2578 | 2680 | 2605 | 2910 | 3000 | 2945 | 2912 |
| K00758 thymidine phosphorylase [EC:2.4.2.4] | 813 | 893 | 836 | 891 | 609 | 653 | 576 | 566 | 1222 | 1237 | 1100 | 1315 |
| K00761 uracil phosphoribosyltransferase [EC:2.4.2.9] | 2157 | 2183 | 2153 | 2242 | 2502 | 2393 | 2458 | 2481 | 2619 | 2616 | 2505 | 2579 |
| K00762 orotate phosphoribosyltransferase [EC:2.4.2.10] | 2979 | 2878 | 3209 | 3023 | 2765 | 2761 | 2728 | 2734 | 3093 | 3056 | 3101 | 3005 |
| K00857 thymidine kinase [EC:2.7.1.21] | 820 | 816 | 911 | 865 | 1564 | 1631 | 1663 | 1644 | 1519 | 1596 | 1547 | 1449 |
| K00876 uridine kinase [EC:2.7.1.48] | 2775 | 2805 | 2927 | 2935 | 5792 | 5796 | 5919 | 5533 | 5055 | 5071 | 5253 | 5061 |
| K00943 dTMP kinase [EC:2.7.4.9] | 773 | 786 | 699 | 754 | 480 | 520 | 570 | 498 | 781 | 814 | 742 | 735 |
| K00945 CMP/dCMP kinase [EC:2.7.4.25] | 1662 | 1789 | 1866 | 1810 | 2403 | 2393 | 2354 | 2287 | 2291 | 2369 | 2486 | 2314 |
| K01240 uridine nucleosidase [EC:3.2.2.3] | 0 | 0 | 0 | 1 | 0 | 0 | 1 | 2 | 3 | 0 | 0 | 0 |
| K01464 dihydropyrimidinase [EC:3.5.2.2] | 3509 | 3257 | 3061 | 3231 | 1774 | 1730 | 1630 | 1808 | 1484 | 1525 | 1433 | 1370 |
| K01465 dihydroorotase [EC:3.5.2.3] | 8609 | 8435 | 8482 | 8595 | 6462 | 6377 | 6196 | 6464 | 6230 | 5985 | 6074 | 6215 |
| K01485 cytosine/creatinine deaminase [EC:3.5.4.1 3.5.4.21] | 3946 | 4137 | 3722 | 3907 | 2167 | 2340 | 2245 | 2205 | 3905 | 4103 | 3980 | 3957 |
| K01489 cytidine deaminase [EC:3.5.4.5] | 1112 | 1096 | 1205 | 1186 | 1343 | 1273 | 1280 | 1303 | 1687 | 1758 | 1738 | 1697 |
| K01493 dCMP deaminase [EC:3.5.4.12] | 724 | 744 | 762 | 708 | 1123 | 1211 | 1214 | 1074 | 1234 | 1026 | 1076 | 1087 |
| K01494 dCTP deaminase [EC:3.5.4.13] | 1440 | 1367 | 1399 | 1417 | 768 | 772 | 734 | 740 | 1105 | 1108 | 1076 | 1173 |
| K01520 dUTP pyrophosphatase [EC:3.6.1.23] | 1968 | 1979 | 1980 | 1941 | 2141 | 2086 | 2055 | 2182 | 1868 | 1930 | 1921 | 1929 |
| K01591 orotidine-5'-phosphate decarboxylase [EC:4.1.1.23] | 2609 | 2494 | 2565 | 2615 | 2938 | 2865 | 2790 | 2940 | 2768 | 2611 | 2621 | 2746 |
| K01937 CTP synthase [EC:6.3.4.2] | 3440 | 3517 | 3557 | 3395 | 4528 | 4932 | 4744 | 4575 | 4789 | 4799 | 4742 | 4839 |
| K01954 carbamoyl-phosphate synthase [EC:6.3.5.5] | 0 | 0 | 1 | 0 | 0 | 0 | 0 | 0 | 0 | 0 | 0 | 0 |
| K01955 carbamoyl-phosphate synthase large subunit [EC:6.3.5.5] | 11446 | 11724 | 11976 | 11789 | 17411 | 17349 | 17323 | 17253 | 16167 | 16086 | 16146 | 15808 |
| K01956 carbamoyl-phosphate synthase small subunit [EC:6.3.5.5] | 3413 | 3507 | 3550 | 3397 | 3829 | 3872 | 3800 | 3724 | 3568 | 3644 | 3572 | 3711 |
| K02823 dihydroorotate dehydrogenase electron transfer subunit | 887 | 920 | 995 | 870 | 1506 | 1532 | 1662 | 1477 | 1386 | 1501 | 1370 | 1441 |
| K02825 pyrimidine operon attenuation protein / uracil phosphoribosyltransferase [EC:2.4.2.9] | 564 | 545 | 597 | 536 | 246 | 277 | 272 | 337 | 261 | 219 | 205 | 282 |
| K03365 cytosine/creatinine deaminase [EC:3.5.4.1 3.5.4.21] | 0 | 0 | 0 | 0 | 0 | 3 | 0 | 0 | 3 | 6 | 0 | 4 |
| K03465 thymidylate synthase (FAD) [EC:2.1.1.148] | 47 | 50 | 51 | 71 | 33 | 43 | 61 | 53 | 114 | 134 | 146 | 121 |
| K06016 beta-ureidopropionase / N-carbamoyl-L-amino-acid hydrolase [EC:3.5.1.6 3.5.1.87] | 2461 | 2229 | 2262 | 2343 | 1629 | 1721 | 1528 | 1566 | 1372 | 1452 | 1340 | 1460 |
| K08722 5'-deoxynucleotidase [EC:3.1.3.89] | 482 | 411 | 368 | 422 | 247 | 258 | 298 | 281 | 472 | 552 | 512 | 542 |
| K08728 nucleoside deoxyribosyltransferase [EC:2.4.2.6] | 0 | 0 | 0 | 0 | 2 | 2 | 0 | 1 | 0 | 0 | 0 | 0 |
| K09018 pyrimidine oxygenase [EC:1.14.99.46] | 567 | 569 | 544 | 586 | 390 | 477 | 416 | 440 | 784 | 774 | 712 | 796 |
| K09019 3-hydroxypropanoate dehydrogenase [EC:1.1.1.-] | 300 | 301 | 275 | 311 | 223 | 228 | 222 | 255 | 399 | 473 | 448 | 440 |
| K09020 ureidoacrylate peracid hydrolase [EC:3.5.1.110] | 370 | 410 | 410 | 405 | 263 | 281 | 313 | 279 | 546 | 558 | 513 | 507 |
| K09021 aminoacrylate peracid reductase | 191 | 216 | 174 | 201 | 127 | 151 | 153 | 130 | 262 | 263 | 263 | 252 |
| K09023 aminoacrylate hydrolase [EC:3.5.1.-] | 491 | 454 | 410 | 515 | 269 | 327 | 332 | 341 | 667 | 628 | 604 | 740 |
| K09024 flavin reductase [EC:1.5.1.-] | 300 | 315 | 319 | 285 | 214 | 242 | 213 | 216 | 460 | 468 | 390 | 458 |
| K09903 uridylate kinase [EC:2.7.4.22] | 2613 | 2606 | 2717 | 2626 | 2881 | 2951 | 2902 | 2792 | 2506 | 2754 | 2585 | 2703 |
| K11540 carbamoyl-phosphate synthase / aspartate carbamoyltransferase / dihydroorotase [EC:6.3.5.5 2.1.3.2 3.5.2.3] | 1 | 2 | 2 | 0 | 0 | 2 | 3 | 0 | 2 | 1 | 0 | 1 |
| K11541 carbamoyl-phosphate synthase / aspartate carbamoyltransferase [EC:6.3.5.5 2.1.3.2] | 1 | 2 | 1 | 0 | 1 | 2 | 1 | 3 | 1 | 0 | 0 | 1 |
| K13421 uridine monophosphate synthetase [EC:2.4.2.10 4.1.1.23] | 1 | 0 | 2 | 0 | 0 | 2 | 2 | 0 | 0 | 0 | 0 | 0 |
| K16066 3-hydroxy acid dehydrogenase / malonic semialdehyde reductase [EC:1.1.1.381 1.1.1.-] | 507 | 368 | 384 | 421 | 278 | 309 | 305 | 295 | 597 | 643 | 588 | 695 |
| K16328 pseudouridine kinase [EC:2.7.1.83] | 515 | 438 | 415 | 422 | 678 | 683 | 650 | 688 | 174 | 157 | 168 | 163 |
| K16329 pseudouridylate synthase [EC:4.2.1.70] | 271 | 234 | 215 | 241 | 321 | 346 | 310 | 334 | 99 | 113 | 123 | 93 |
| K17722 dihydropyrimidine dehydrogenase (NAD+) subunit PreT [EC:1.3.1.1] | 1823 | 1764 | 1655 | 1692 | 1294 | 1289 | 1292 | 1320 | 510 | 547 | 514 | 514 |
| K17723 dihydropyrimidine dehydrogenase (NAD+) subunit PreA [EC:1.3.1.1] | 2259 | 2260 | 2049 | 2121 | 1488 | 1395 | 1201 | 1539 | 618 | 659 | 641 | 665 |
| K17828 dihydroorotate dehydrogenase (NAD+) catalytic subunit [EC:1.3.1.14] | 944 | 1033 | 1099 | 1115 | 1731 | 1871 | 1903 | 1675 | 1670 | 1724 | 1761 | 1745 |
| Amino acid metabolism | 928212 | 920354 | 906433 | 918771 | 830701 | 829189 | 823081 | 834247 | 824292 | 823098 | 814008 | 821853 |
| ko00250 Alanine, aspartate and glutamate metabolism | 165048 | 165483 | 165153 | 164301 | 182305 | 181946 | 182325 | 181734 | 169127 | 169019 | 168291 | 167352 |
| K00259 alanine dehydrogenase [EC:1.4.1.1] | 721 | 851 | 830 | 738 | 2287 | 2337 | 2369 | 2165 | 1856 | 1597 | 1732 | 1630 |
| K00278 L-aspartate oxidase [EC:1.4.3.16] | 4449 | 4501 | 4437 | 4428 | 5809 | 5871 | 5924 | 5696 | 5188 | 5096 | 4897 | 4862 |
| K00294 1-pyrroline-5-carboxylate dehydrogenase [EC:1.2.1.88] | 21 | 14 | 18 | 16 | 3 | 0 | 0 | 0 | 3 | 10 | 4 | 3 |

| | | | | | | | | | | | | |
|---|---|---|---|---|---|---|---|---|---|---|---|---|
| K00812 aspartate aminotransferase [EC:2.6.1.1] | 1589 | 1772 | 1943 | 1689 | 5232 | 5097 | 5317 | 5193 | 3844 | 3733 | 3875 | 3655 |
| K00813 aspartate aminotransferase [EC:2.6.1.1] | 671 | 674 | 656 | 736 | 477 | 493 | 430 | 444 | 884 | 942 | 872 | 1056 |
| K01424 L-asparaginase [EC:3.5.1.1] | 3849 | 3972 | 3973 | 4000 | 7585 | 7501 | 7727 | 7685 | 7429 | 7388 | 7394 | 7118 |
| K01425 glutaminase [EC:3.5.1.2] | 1423 | 1361 | 1424 | 1428 | 2703 | 2855 | 3083 | 2739 | 2523 | 2349 | 2628 | 2450 |
| K01744 aspartate ammonia-lyase [EC:4.3.1.1] | 4427 | 4526 | 4316 | 4271 | 5404 | 5342 | 5430 | 5568 | 5061 | 4961 | 4983 | 4932 |
| K01755 argininosuccinate lyase [EC:4.3.2.1] | 5369 | 5397 | 5524 | 5321 | 5867 | 6017 | 6011 | 6088 | 5538 | 5625 | 5496 | 5635 |
| K01779 aspartate racemase [EC:5.1.1.13] | 530 | 565 | 483 | 574 | 426 | 482 | 409 | 398 | 374 | 434 | 408 | 412 |
| K01914 aspartate--ammonia ligase [EC:6.3.1.1] | 1802 | 1845 | 1862 | 1810 | 2827 | 2924 | 3004 | 2780 | 2999 | 3036 | 3042 | 2906 |
| K01940 argininosuccinate synthase [EC:6.3.4.5] | 4155 | 4382 | 4200 | 3990 | 4692 | 4578 | 4660 | 4906 | 4226 | 4227 | 4107 | 4251 |
| K01953 asparagine synthase (glutamine-hydrolysing) [EC:6.3.5.4] | 11561 | 11240 | 10660 | 10963 | 8426 | 8356 | 8249 | 8602 | 7075 | 6942 | 6910 | 6985 |
| K05597 glutamin-(asparagin-)ase [EC:3.5.1.38] | 209 | 199 | 178 | 188 | 110 | 92 | 104 | 115 | 64 | 62 | 56 | 71 |
| K09758 aspartate 4-decarboxylase [EC:4.1.1.12] | 940 | 1089 | 1072 | 1119 | 2774 | 2888 | 2855 | 2967 | 2094 | 2022 | 2219 | 1939 |
| K11358 aspartate aminotransferase [EC:2.6.1.1] | 502 | 502 | 518 | 516 | 147 | 125 | 149 | 145 | 447 | 472 | 411 | 468 |
| K13051 L-asparaginase / beta-aspartyl-peptidase [EC:3.5.1.1 3.4.19.5] | 657 | 610 | 607 | 647 | 478 | 581 | 556 | 459 | 737 | 781 | 709 | 770 |
| K13566 omega-amidase [EC:3.5.1.3] | 2149 | 2128 | 2191 | 2119 | 2595 | 2652 | 2570 | 2537 | 2126 | 2238 | 2171 | 2142 |
| K13821 RHH-type transcriptional regulator, proline utilization regulon repressor / proline dehydrogenase / delta 1-pyrroline-5-carboxylate dehydrogenase [E( | 6056 | 5732 | 5737 | 5696 | 3788 | 3676 | 3667 | 3932 | 4224 | 4224 | 3917 | 4216 |
| K14260 alanine-synthesizing transaminase [EC:2.6.1.66 2.6.1.2] | 2511 | 2457 | 2615 | 2520 | 1531 | 1566 | 1519 | 1504 | 1910 | 1937 | 1854 | 1927 |
| K14681 argininosuccinate lyase / amino-acid N-acetyltransferase [EC:4.3.2.1 2.3.1.1] | 8 | 8 | 12 | 9 | 10 | 12 | 15 | 11 | 16 | 11 | 14 | 12 |
| K19244 alanine dehydrogenase [EC:1.4.1.1] | 0 | 3 | 0 | 2 | 0 | 0 | 1 | 2 | 4 | 1 | 2 | 10 |
| K22457 asparagine---oxo-acid transaminase [EC:2.6.1.14] | 82 | 55 | 80 | 70 | 22 | 27 | 16 | 6 | 62 | 102 | 68 | 107 |
| ko00260 Glycine, serine and threonine metabolism | 158044 | 156295 | 152537 | 155504 | 140898 | 141299 | 139962 | 141766 | 141694 | 139878 | 139509 | 140126 |
| K00003 homoserine dehydrogenase [EC:1.1.1.3] | 2777 | 2707 | 2779 | 2703 | 1433 | 1334 | 1351 | 1461 | 1522 | 1514 | 1523 | 1526 |
| K00060 threonine 3-dehydrogenase [EC:1.1.1.103] | 1000 | 949 | 1044 | 1073 | 490 | 592 | 496 | 493 | 1565 | 1590 | 1511 | 1580 |
| K00108 choline dehydrogenase [EC:1.1.99.1] | 6688 | 6362 | 6082 | 6470 | 3340 | 3240 | 3512 | 3591 | 3390 | 3448 | 3156 | 3252 |
| K00130 betaine-aldehyde dehydrogenase [EC:1.2.1.8] | 3519 | 3673 | 3393 | 3561 | 1726 | 1657 | 1553 | 1708 | 2545 | 2587 | 2481 | 2467 |
| K00133 aspartate-semialdehyde dehydrogenase [EC:1.2.1.11] | 3907 | 4075 | 3992 | 4083 | 3987 | 4038 | 4111 | 3983 | 4784 | 4641 | 4814 | 4672 |
| K00274 monoamine oxidase [EC:1.4.3.4] | 2541 | 2664 | 2610 | 2588 | 1445 | 1409 | 1307 | 1544 | 864 | 874 | 791 | 749 |
| K00276 primary-amine oxidase [EC:1.4.3.21] | 799 | 774 | 720 | 792 | 209 | 222 | 195 | 209 | 1545 | 1642 | 1487 | 1568 |
| K00302 sarcosine oxidase, subunit alpha [EC:1.5.3.1] | 5148 | 5221 | 4686 | 4775 | 2694 | 2506 | 2568 | 2926 | 1705 | 1576 | 1489 | 1702 |
| K00303 sarcosine oxidase, subunit beta [EC:1.5.3.1] | 1941 | 1835 | 1589 | 1816 | 968 | 904 | 927 | 992 | 623 | 552 | 526 | 603 |
| K00304 sarcosine oxidase, subunit delta [EC:1.5.3.1] | 519 | 561 | 507 | 502 | 340 | 321 | 268 | 322 | 173 | 193 | 188 | 214 |
| K00305 sarcosine oxidase, subunit gamma [EC:1.5.3.1] | 6 | 13 | 2 | 3 | 3 | 3 | 1 | 5 | 4 | 4 | 3 | 1 |
| K00314 sarcosine dehydrogenase [EC:1.5.8.3] | 0 | 0 | 0 | 0 | 0 | 0 | 0 | 0 | 0 | 1 | 0 | 0 |
| K00613 glycine amidinotransferase [EC:2.1.4.1] | 11 | 8 | 5 | 11 | 12 | 9 | 7 | 12 | 19 | 30 | 26 | 31 |
| K00639 glycine C-acetyltransferase [EC:2.3.1.29] | 1585 | 1696 | 1665 | 1695 | 2989 | 3283 | 3250 | 3087 | 2927 | 2976 | 2973 | 2938 |
| K00643 5-aminolevulinate synthase [EC:2.3.1.37] | 1 | 0 | 0 | 1 | 2 | 1 | 3 | 0 | 2 | 1 | 3 | 3 |
| K00836 diaminobutyrate-2-oxoglutarate transaminase [EC:2.6.1.76] | 1899 | 1881 | 1861 | 1888 | 1116 | 1101 | 1039 | 1063 | 1493 | 1590 | 1377 | 1548 |
| K00872 homoserine kinase [EC:2.7.1.39] | 690 | 706 | 656 | 708 | 424 | 458 | 492 | 439 | 852 | 883 | 918 | 982 |
| K00928 aspartate kinase [EC:2.7.2.4] | 5433 | 5415 | 5737 | 5526 | 7655 | 7537 | 7771 | 7396 | 6513 | 6555 | 6530 | 6417 |
| K01620 threonine aldolase [EC:4.1.2.48] | 5121 | 5056 | 4889 | 5075 | 4321 | 4190 | 4266 | 4486 | 4014 | 3819 | 3782 | 3979 |
| K01695 tryptophan synthase alpha chain [EC:4.2.1.20] | 2398 | 2529 | 2414 | 2500 | 2507 | 2534 | 2459 | 2570 | 2450 | 2351 | 2350 | 2428 |
| K01696 tryptophan synthase beta chain [EC:4.2.1.20] | 3407 | 3488 | 3374 | 3542 | 3727 | 3666 | 3808 | 3760 | 3718 | 3670 | 3836 | 3655 |
| K01697 cystathionine beta-synthase [EC:4.2.1.22] | 2036 | 2021 | 1964 | 2117 | 1134 | 1258 | 1185 | 1214 | 1517 | 1575 | 1531 | 1593 |
| K01733 threonine synthase [EC:4.2.3.1] | 5015 | 4928 | 5092 | 5061 | 5326 | 5341 | 5175 | 5358 | 5075 | 4936 | 5149 | 5069 |
| K01752 L-serine dehydratase [EC:4.3.1.17] | 6980 | 6897 | 6631 | 6587 | 6102 | 6057 | 6175 | 6262 | 6318 | 6343 | 6198 | 6353 |
| K01753 D-serine dehydratase [EC:4.3.1.18] | 3125 | 3132 | 2796 | 3041 | 1806 | 1728 | 1710 | 1741 | 2098 | 2015 | 2003 | 1984 |
| K01754 threonine dehydratase [EC:4.3.1.19] | 6866 | 6744 | 6331 | 6812 | 3613 | 3717 | 3565 | 3712 | 4149 | 4296 | 4212 | 4377 |
| K01758 cystathionine gamma-lyase [EC:4.4.1.1] | 2729 | 2548 | 2406 | 2518 | 1539 | 1646 | 1410 | 1619 | 1603 | 1519 | 1478 | 1509 |
| K02204 homoserine kinase type II [EC:2.7.1.39] | 1420 | 1487 | 1433 | 1311 | 809 | 724 | 781 | 800 | 479 | 379 | 451 | 469 |
| K06001 tryptophan synthase beta chain [EC:4.2.1.20] | 1743 | 1712 | 1843 | 1772 | 3103 | 2922 | 3130 | 3111 | 2950 | 2971 | 3086 | 2895 |
| K06720 L-ectoine synthase [EC:4.2.1.108] | 0 | 0 | 0 | 0 | 2 | 1 | 1 | 0 | 0 | 2 | 0 | 1 |
| K11440 choline dehydrogenase [EC:1.1.1.1] | 1 | 0 | 0 | 0 | 0 | 1 | 1 | 0 | 0 | 0 | 0 | 0 |
| K12524 bifunctional aspartokinase / homoserine dehydrogenase 1 [EC:2.7.2.4 1.1.1.3] | 3160 | 3193 | 3211 | 3233 | 6630 | 6536 | 6608 | 6626 | 5999 | 5925 | 6009 | 5745 |
| K12525 bifunctional aspartokinase / homoserine dehydrogenase 2 [EC:2.7.2.4 1.1.1.3] | 1610 | 1664 | 1653 | 1589 | 1158 | 1176 | 1126 | 1080 | 2170 | 2228 | 2181 | 2255 |
| K13290 serine---pyruvate transaminase [EC:2.6.1.51] | 1 | 0 | 1 | 0 | 2 | 0 | 0 | 0 | 0 | 0 | 0 | 0 |
| K13745 L-2,4-diaminobutyrate decarboxylase [EC:4.1.1.86] | 783 | 741 | 729 | 765 | 481 | 520 | 533 | 537 | 1070 | 1082 | 965 | 1120 |
| K15783 ectoine hydrolase [EC:3.5.4.44] | 2 | 0 | 0 | 1 | 3 | 1 | 1 | 0 | 0 | 0 | 2 | 2 |
| K15784 N2-acetyl-L-2,4-diaminobutanoate deacetylase [EC:3.5.1.125] | 0 | 0 | 2 | 1 | 0 | 0 | 2 | 0 | 5 | 6 | 2 | 0 |
| K15785 L-2,4-diaminobutyrate transaminase [EC:2.6.1.76] | 17 | 12 | 14 | 19 | 0 | 2 | 3 | 0 | 8 | 13 | 11 | 9 |
| K15786 aspartate-semialdehyde dehydrogenase [EC:1.2.1.-] | 1 | 0 | 0 | 0 | 0 | 2 | 0 | 0 | 0 | 0 | 2 | 0 |
| K20498 D-serine ammonia-lyase [EC:4.3.1.18] | 2 | 2 | 0 | 0 | 1 | 0 | 1 | 0 | 0 | 0 | 0 | 1 |
| ko00270 Cysteine and methionine metabolism | 136148 | 134863 | 135435 | 136507 | 136745 | 137420 | 136655 | 137130 | 140832 | 141208 | 139817 | 139860 |
| K00547 homocysteine S-methyltransferase [EC:2.1.1.10] | 663 | 682 | 600 | 663 | 454 | 423 | 423 | 455 | 884 | 861 | 741 | 856 |
| K00548 5-methyltetrahydrofolate--homocysteine methyltransferase [EC:2.1.1.13] | 8752 | 8490 | 8722 | 8756 | 9047 | 9115 | 9238 | 9191 | 9750 | 9455 | 9498 | 9473 |
| K00549 5-methyltetrahydropteroyltriglutamate--homocysteine methyltransferase [EC:2.1.1.14] | 4885 | 4929 | 4876 | 5003 | 3146 | 3387 | 3120 | 3223 | 4303 | 4520 | 4335 | 4511 |
| K00558 DNA (cytosine-5)-methyltransferase 1 [EC:2.1.1.37] | 3000 | 3049 | 3137 | 3037 | 5781 | 6034 | 6013 | 6006 | 5676 | 5447 | 5471 | 5544 |
| K00772 5'-methylthioadenosine phosphorylase [EC:2.4.2.28] | 100 | 104 | 114 | 114 | 72 | 78 | 59 | 84 | 141 | 140 | 132 | 147 |
| K00789 S-adenosylmethionine synthetase [EC:2.5.1.6] | 4569 | 4549 | 4585 | 4822 | 4605 | 4545 | 4364 | 4428 | 4753 | 4870 | 4666 | 4677 |
| K00797 spermidine synthase [EC:2.5.1.16] | 3573 | 3591 | 3385 | 3518 | 2135 | 2097 | 1887 | 2094 | 1797 | 1759 | 1669 | 1721 |
| K00816 kynurenine---oxoglutarate transaminase / cysteine-S-conjugate beta-lyase / glutamine---phenylpyruvate transaminase [EC:2.6.1.7 4.4.1.13 2.6.1.64] | 0 | 0 | 0 | 0 | 0 | 0 | 0 | 0 | 0 | 0 | 1 | 0 |
| K00826 branched-chain amino acid aminotransferase [EC:2.6.1.42] | 3364 | 3225 | 3321 | 3308 | 4020 | 3688 | 3853 | 3828 | 3706 | 3768 | 3705 | 3625 |
| K00832 aromatic-amino-acid transaminase [EC:2.6.1.57] | 4599 | 4472 | 4392 | 4485 | 2742 | 2763 | 2799 | 2815 | 2957 | 2867 | 2938 | 3036 |
| K00838 aromatic amino acid aminotransferase I / 2-aminoadipate transaminase [EC:2.6.1.57 2.6.1.39 2.6.1.27 2.6.1.5] | 0 | 0 | 0 | 0 | 0 | 0 | 0 | 0 | 0 | 1 | 0 | 0 |
| K00899 5-methylthioribose kinase [EC:2.7.1.100] | 794 | 757 | 688 | 831 | 637 | 707 | 698 | 621 | 1106 | 1180 | 1034 | 1210 |
| K01243 adenosylhomocysteine nucleosidase [EC:3.2.2.9] | 1316 | 1216 | 1237 | 1342 | 1725 | 1655 | 1725 | 1685 | 1752 | 1740 | 1923 | 1808 |
| K01251 adenosylhomocysteinase [EC:3.3.1.1] | 2101 | 2254 | 2347 | 2234 | 3792 | 3988 | 3703 | 3764 | 3716 | 3771 | 3663 | 3584 |
| K01505 1-aminocyclopropane-1-carboxylate deaminase [EC:3.5.99.7] | 1155 | 1089 | 1002 | 1080 | 600 | 570 | 579 | 665 | 322 | 352 | 341 | 321 |
| K01611 S-adenosylmethionine decarboxylase [EC:4.1.1.50] | 2392 | 2220 | 2317 | 2268 | 1395 | 1415 | 1280 | 1362 | 1324 | 1416 | 1348 | 1333 |
| K01740 O-acetylhomoserine (thiol)-lyase [EC:2.5.1.49] | 4902 | 4900 | 5249 | 4953 | 6668 | 6795 | 7074 | 6918 | 5724 | 5624 | 5536 | 5354 |
| K01760 cysteine-S-conjugate beta-lyase [EC:4.4.1.13] | 1260 | 1241 | 1256 | 1308 | 802 | 879 | 874 | 852 | 2225 | 2238 | 2184 | 2242 |
| K01761 methionine-gamma-lyase [EC:4.4.1.11] | 33 | 17 | 26 | 19 | 94 | 134 | 111 | 106 | 267 | 255 | 222 | 233 |
| K01919 glutamate--cysteine ligase [EC:6.3.2.2] | 3641 | 3578 | 3483 | 3575 | 2210 | 2153 | 2038 | 2248 | 2446 | 2478 | 2536 | 2573 |
| K01920 glutathione synthase [EC:6.3.2.3] | 2012 | 2093 | 1998 | 2020 | 1250 | 1273 | 1269 | 1275 | 1310 | 1230 | 1180 | 1358 |
| K05396 D-cysteine desulfhydrase [EC:4.4.1.15] | 689 | 621 | 625 | 710 | 509 | 588 | 511 | 532 | 883 | 985 | 912 | 1012 |
| K07173 S-ribosylhomocysteine lyase [EC:4.4.1.21] | 720 | 706 | 722 | 737 | 534 | 536 | 522 | 530 | 602 | 658 | 600 | 702 |
| K08963 methylthioribose-1-phosphate isomerase [EC:5.3.1.23] | 1792 | 1744 | 1840 | 1799 | 1142 | 1180 | 1178 | 1162 | 1544 | 1551 | 1516 | 1635 |
| K08964 methylthioribulose-1-phosphate dehydratase [EC:4.2.1.109] | 1178 | 1136 | 1117 | 1183 | 781 | 678 | 662 | 732 | 687 | 744 | 683 | 730 |
| K08965 2,3-diketo-5-methylthiopentyl-1-phosphate enolase [EC:5.3.2.5] | 374 | 317 | 340 | 310 | 50 | 63 | 67 | 58 | 161 | 190 | 204 | 192 |
| K08966 2-hydroxy-3-keto-5-methylthiopentenyl-1-phosphate phosphatase [EC:3.1.3.87] | 1 | 0 | 2 | 0 | 0 | 0 | 0 | 0 | 0 | 0 | 0 | 0 |
| K08967 1,2-dihydroxy-3-keto-5-methylthiopentene dioxygenase [EC:1.13.11.53 1.13.11.54] | 635 | 712 | 620 | 726 | 513 | 555 | 453 | 503 | 551 | 513 | 469 | 544 |
| K08968 L-methionine (R)-S-oxide reductase [EC:1.8.4.14] | 1228 | 1262 | 1199 | 1180 | 1435 | 1560 | 1534 | 1530 | 1376 | 1356 | 1292 | 1275 |
| K08969 aminotransferase [EC:2.6.1.-] | 648 | 646 | 625 | 656 | 559 | 485 | 496 | 458 | 833 | 835 | 838 | 900 |
| K09880 enolase-phosphatase E1 [EC:3.1.3.77] | 1454 | 1325 | 1269 | 1358 | 791 | 764 | 830 | 786 | 887 | 799 | 867 | 850 |
| K12339 S-sulfo-L-cysteine synthase (O-acetyl-L-serine-dependent) [EC:2.5.1.144] | 1955 | 1957 | 1928 | 2010 | 1245 | 1233 | 1132 | 1160 | 1331 | 1200 | 1243 | 1314 |
| K12960 5-methylthioadenosine/S-adenosylhomocysteine deaminase [EC:3.5.4.31 3.5.4.28] | 2497 | 2335 | 2302 | 2298 | 1275 | 1307 | 1195 | 1285 | 1037 | 1055 | 1076 | 984 |
| K13060 acyl homoserine lactone synthase [EC:2.3.1.184] | 189 | 230 | 226 | 204 | 97 | 129 | 101 | 157 | 56 | 43 | 58 | 65 |
| K13061 acyl homoserine lactone synthase [EC:2.3.1.184] | 685 | 695 | 709 | 688 | 391 | 356 | 427 | 388 | 251 | 215 | 217 | 238 |
| K14155 cysteine-S-conjugate beta-lyase [EC:4.4.1.13] | 3799 | 3885 | 4001 | 3971 | 5788 | 5893 | 6057 | 5942 | 5877 | 6169 | 6249 | 6073 |
| K16843 (S)-sulfolactate dehydrogenase [EC:1.1.1.310] | 9 | 8 | 4 | 5 | 1 | 1 | 6 | 3 | 2 | 2 | 4 | 5 |
| K16844 (2R)-3-sulfolactate dehydrogenase (NADP+) [EC:1.1.1.338] | 7 | 4 | 3 | 5 | 0 | 0 | 0 | 0 | 0 | 3 | 0 | 0 |
| K16845 (2R)-sulfolactate sulfo-lyase subunit alpha [EC:4.4.1.24] | 3 | 8 | 2 | 5 | 4 | 1 | 4 | 5 | 8 | 14 | 13 | 8 |
| K16846 (2R)-sulfolactate sulfo-lyase subunit beta [EC:4.4.1.24] | 34 | 35 | 30 | 24 | 14 | 13 | 11 | 3 | 34 | 48 | 23 | 39 |
| K17216 cystathionine beta-synthase (O-acetyl-L-serine) [EC:2.5.1.134] | 0 | 1 | 0 | 0 | 0 | 0 | 0 | 0 | 0 | 0 | 0 | 1 |
| K17398 DNA (cytosine-5)-methyltransferase 3A [EC:2.1.1.37] | 16 | 10 | 13 | 15 | 42 | 45 | 34 | 35 | 5 | 8 | 11 | 4 |
| K17462 putative AdoMet-dependent methyltransferase [EC:2.1.1.-] | 45 | 33 | 37 | 36 | 14 | 7 | 7 | 10 | 29 | 20 | 29 | 22 |
| K17950 L-cysteate sulfo-lyase [EC:4.4.1.25] | 250 | 252 | 202 | 247 | 409 | 400 | 342 | 362 | 0 | 8 | 1 | 1 |
| K19696 5'-methylthioinosine phosphorylase [EC:2.4.2.44] | 997 | 912 | 938 | 948 | 521 | 418 | 469 | 542 | 326 | 314 | 265 | 317 |
| ko00280 Valine, leucine and isoleucine degradation | 100118 | 98754 | 96172 | 98193 | 78402 | 75896 | 75111 | 78377 | 66083 | 64843 | 64138 | 65077 |
| K00020 3-hydroxyisobutyrate dehydrogenase [EC:1.1.1.31] | 3819 | 3771 | 3646 | 3751 | 1823 | 1757 | 1864 | 2024 | 2096 | 1992 | 1959 | 1982 |
| K00186 2-oxoisovalerate ferredoxin oxidoreductase, alpha subunit [EC:1.2.7.7] | 2 | 2 | 0 | 3 | 0 | 0 | 2 | 0 | 3 | 6 | 3 | 7 |
| K00187 2-oxoisovalerate ferredoxin oxidoreductase, beta subunit [EC:1.2.7.7] | 0 | 0 | 1 | 1 | 1 | 2 | 0 | 0 | 7 | 10 | 7 | 15 |
| K00188 2-oxoisovalerate ferredoxin oxidoreductase, delta subunit [EC:1.2.7.7] | 0 | 0 | 0 | 0 | 0 | 0 | 0 | 0 | 0 | 2 | 2 | 1 |
| K00253 isovaleryl-CoA dehydrogenase [EC:1.3.8.4] | 1163 | 1258 | 1138 | 1176 | 657 | 655 | 580 | 679 | 442 | 387 | 371 | 343 |
| K00263 leucine dehydrogenase [EC:1.4.1.9] | 14 | 17 | 16 | 21 | 17 | 15 | 25 | 12 | 21 | 28 | 18 | 44 |
| K01968 3-methylcrotonyl-CoA carboxylase alpha subunit [EC:6.4.1.4] | 2713 | 2605 | 2290 | 2618 | 1305 | 1278 | 1257 | 1370 | 804 | 668 | 618 | 726 |
| K01969 3-methylcrotonyl-CoA carboxylase beta subunit [EC:6.4.1.4] | 2139 | 2018 | 1971 | 2011 | 1132 | 1014 | 1128 | 1238 | 697 | 609 | 587 | 637 |
| K08683 3-hydroxyacyl-CoA dehydrogenase / 3-hydroxy-2-methylbutyryl-CoA dehydrogenase [EC:1.1.1.35 1.1.1.178] | 0 | 0 | 0 | 1 | 0 | 0 | 0 | 0 | 0 | 0 | 0 | 0 |
| K11410 short/branched chain acyl-CoA dehydrogenase [EC:1.3.99.12] | 0 | 0 | 0 | 0 | 0 | 0 | 0 | 0 | 1 | 2 | 0 | 0 |
| K13766 methylglutaconyl-CoA hydratase [EC:4.2.1.18] | 3 | 4 | 3 | 3 | 3 | 1 | 4 | 1 | 4 | 6 | 3 | 4 |
| K18661 malonyl-CoA/methylmalonyl-CoA synthetase [EC:6.2.1.-] | 0 | 2 | 0 | 0 | 0 | 0 | 0 | 0 | 0 | 0 | 0 | 1 |
| ko00290 Valine, leucine and isoleucine biosynthesis | 60133 | 59525 | 59186 | 60762 | 53405 | 53901 | 52691 | 52867 | 59166 | 60533 | 59088 | 60512 |

| | | | | | | | | | | | | |
|---|---|---|---|---|---|---|---|---|---|---|---|---|
| K00053 ketol-acid reductoisomerase [EC:1.1.1.86] | 3866 | 3730 | 3880 | 3924 | 4061 | 4120 | 3962 | 4071 | 4363 | 4431 | 4412 | 4490 |
| K00835 valine--pyruvate aminotransferase [EC:2.6.1.66] | 817 | 806 | 807 | 777 | 591 | 582 | 644 | 578 | 1124 | 1048 | 956 | 1128 |
| K01687 dihydroxy-acid dehydratase [EC:4.2.1.9] | 8688 | 8322 | 8541 | 8619 | 6945 | 7119 | 6911 | 7090 | 8859 | 9319 | 9037 | 9384 |
| K01702 3-isopropylmalate dehydratase [EC:4.2.1.33] | 1 | 0 | 0 | 0 | 0 | 0 | 0 | 0 | 0 | 0 | 1 | 0 |
| K21360 3-isopropylmalate/methylthioalkylmalate dehydrogenase [EC:1.1.1.85 1.1.1.-] | 0 | 0 | 0 | 0 | 1 | 0 | 0 | 0 | 0 | 0 | 0 | 0 |
| ko00300 Lysine biosynthesis | 57096 | 56875 | 57438 | 57773 | 66099 | 66061 | 66541 | 65933 | 65460 | 66424 | 65814 | 65405 |
| K00215 4-hydroxy-tetrahydrodipicolinate reductase [EC:1.17.1.8] | 2573 | 2480 | 2670 | 2483 | 3013 | 2926 | 2920 | 2925 | 2740 | 2726 | 2703 | 2757 |
| K00290 saccharopine dehydrogenase (NAD+, L-lysine forming) [EC:1.5.1.7] | 886 | 888 | 1022 | 899 | 2611 | 2589 | 2683 | 2681 | 1898 | 1907 | 1951 | 1821 |
| K00293 saccharopine dehydrogenase (NADP+, L-glutamate forming) [EC:1.5.1.10] | 0 | 0 | 2 | 0 | 0 | 0 | 0 | 0 | 0 | 0 | 0 | 1 |
| K00674 2,3,4,5-tetrahydropyridine-2,6-dicarboxylate N-succinyltransferase [EC:2.3.1.117] | 1486 | 1409 | 1370 | 1525 | 1589 | 1562 | 1515 | 1477 | 1603 | 1540 | 1563 | 1547 |
| K00821 acetylornithine/N-succinyldiaminopimelate aminotransferase [EC:2.6.1.11 2.6.1.17] | 3969 | 3790 | 4033 | 4230 | 4166 | 4199 | 4155 | 4092 | 4137 | 4117 | 4016 | 4109 |
| K00841 aminotransferase [EC:2.6.1.-] | 51 | 51 | 76 | 68 | 23 | 32 | 26 | 24 | 73 | 82 | 67 | 65 |
| K01439 succinyl-diaminopimelate desuccinylase [EC:3.5.1.18] | 4200 | 4116 | 3902 | 4232 | 1963 | 1900 | 1906 | 1850 | 3574 | 3814 | 3592 | 3793 |
| K01586 diaminopimelate decarboxylase [EC:4.1.1.20] | 3521 | 3526 | 3366 | 3564 | 3677 | 3768 | 3646 | 3791 | 4079 | 4198 | 4044 | 3920 |
| K01714 4-hydroxy-tetrahydrodipicolinate synthase [EC:4.3.3.7] | 7653 | 7602 | 7412 | 7583 | 5481 | 5507 | 5501 | 5630 | 5832 | 6314 | 6024 | 6051 |
| K01778 diaminopimelate epimerase [EC:5.1.1.7] | 3391 | 3264 | 3244 | 3354 | 2935 | 2940 | 2978 | 2942 | 2879 | 3109 | 2908 | 3027 |
| K01928 UDP-N-acetylmuramoyl-L-alanyl-D-glutamate--2,6-diaminopimelate ligase [EC:6.3.2.13] | 3073 | 3120 | 3241 | 3076 | 4560 | 4751 | 4650 | 4488 | 4568 | 4635 | 4478 | 4797 |
| K01929 UDP-N-acetylmuramoyl-tripeptide--D-alanyl-D-alanine ligase [EC:6.3.2.10] | 3391 | 3468 | 3536 | 3528 | 4813 | 5042 | 5129 | 4962 | 4166 | 4428 | 4240 | 4184 |
| K03340 diaminopimelate dehydrogenase [EC:1.4.1.16] | 1012 | 1010 | 1112 | 1043 | 1656 | 1695 | 1878 | 1730 | 1551 | 1637 | 1856 | 1678 |
| K05822 tetrahydrodipicolinate N-acetyltransferase [EC:2.3.1.89] | 70 | 50 | 64 | 63 | 76 | 76 | 33 | 34 | 78 | 73 | 66 | 74 |
| K05823 N-acetyldiaminopimelate deacetylase [EC:3.5.1.47] | 54 | 66 | 77 | 72 | 8 | 6 | 8 | 12 | 49 | 73 | 47 | 74 |
| K05824 homoisocitrate dehydrogenase [EC:1.1.1.87] | 3 | 0 | 0 | 1 | 0 | 0 | 0 | 0 | 9 | 13 | 13 | 9 |
| K05825 2-aminoadipate transaminase [EC:2.6.1.-] | 954 | 1032 | 1020 | 946 | 2515 | 2474 | 2406 | 2580 | 2427 | 2339 | 2483 | 2353 |
| K05827 [lysine-biosynthesis-protein LysW]---L-2-aminoadipate ligase [EC:6.3.2.43] | 0 | 0 | 0 | 0 | 0 | 0 | 0 | 0 | 2 | 3 | 0 | 0 |
| K05828 LysW-gamma-L-alpha-aminoadipate/LysW-L-glutamate kinase [EC:2.7.2.-] | 0 | 0 | 0 | 0 | 0 | 2 | 0 | 0 | 0 | 0 | 0 | 0 |
| K05829 LysW-gamma-L-alpha-aminoadipyl-6-phosphate/LysW-L-glutamyl-5-phosphate reductase [EC:1.2.1.-] | 2 | 2 | 0 | 0 | 0 | 4 | 3 | 0 | 1 | 2 | 3 | 1 |
| K05830 LysW-gamma-L-lysine/LysW-L-ornithine aminotransferase [EC:2.6.1.-] | 0 | 0 | 0 | 0 | 0 | 0 | 0 | 0 | 0 | 3 | 0 | 1 |
| K05831 [amino group carrier protein]-lysine/ornithine hydrolase [EC:3.5.1.130 3.5.1.132] | 0 | 0 | 2 | 0 | 0 | 0 | 0 | 0 | 2 | 0 | 0 | 1 |
| K10206 LL-diaminopimelate aminotransferase [EC:2.6.1.83] | 2008 | 2082 | 2134 | 2131 | 5095 | 5006 | 5048 | 5036 | 4180 | 3948 | 4045 | 3924 |
| K14267 N-succinyldiaminopimelate aminotransferase [EC:2.6.1.17] | 1897 | 1850 | 1768 | 1823 | 1042 | 942 | 1071 | 1109 | 606 | 564 | 626 | 546 |
| K15792 murE/murF fusion protein [EC:6.3.2.13 6.3.2.10] | 7 | 5 | 5 | 3 | 3 | 9 | 6 | 12 | 6 | 7 | 6 | 9 |
| K16180 methylornithine synthase [EC:5.4.99.58] | 4 | 5 | 2 | 5 | 1 | 0 | 3 | 3 | 0 | 0 | 2 | 3 |
| K16181 pyrrolysine biosynthesis protein PylC | 0 | 2 | 2 | 4 | 0 | 0 | 0 | 1 | 2 | 0 | 0 | 1 |
| K16182 pyrrolysine biosynthesis protein PylD | 0 | 0 | 0 | 1 | 0 | 0 | 0 | 2 | 0 | 0 | 0 | 0 |
| K17450 homoaconitase [EC:4.2.1.-] | 2 | 1 | 2 | 1 | 6 | 1 | 3 | 3 | 0 | 3 | 3 | 1 |
| ko00310 Lysine degradation | 64790 | 64535 | 62440 | 64129 | 48507 | 47720 | 47077 | 48598 | 50967 | 50225 | 49737 | 51131 |
| K00137 aminobutyraldehyde dehydrogenase [EC:1.2.1.19] | 2653 | 2680 | 2673 | 2777 | 1576 | 1569 | 1579 | 1506 | 1804 | 1643 | 1648 | 1625 |
| K00468 lysine 2-monooxygenase [EC:1.13.12.2] | 0 | 0 | 0 | 0 | 0 | 0 | 0 | 0 | 0 | 0 | 0 | 1 |
| K00824 D-alanine transaminase [EC:2.6.1.21] | 303 | 263 | 271 | 266 | 243 | 286 | 267 | 233 | 133 | 123 | 116 | 127 |
| K01506 5-aminopentanamidase [EC:3.5.1.30] | 6 | 9 | 12 | 13 | 3 | 4 | 3 | 4 | 0 | 0 | 0 | 0 |
| K01582 lysine decarboxylase [EC:4.1.1.18] | 2277 | 2322 | 2086 | 2428 | 1315 | 1493 | 1354 | 1251 | 3888 | 3932 | 3770 | 3918 |
| K01843 lysine 2,3-aminomutase [EC:5.4.3.2] | 1370 | 1425 | 1543 | 1341 | 4447 | 4328 | 4465 | 4320 | 2925 | 2854 | 3205 | 2965 |
| K01844 beta-lysine 5,6-aminomutase alpha subunit [EC:5.4.3.3] | 0 | 0 | 0 | 0 | 2 | 4 | 3 | 0 | 0 | 6 | 3 | 1 |
| K03894 N2-citryl-N6-acetyl-N6-hydroxylysine synthase [EC:6.3.2.38] | 0 | 2 | 3 | 0 | 3 | 2 | 3 | 5 | 3 | 6 | 3 | 6 |
| K03895 aerobactin synthase [EC:6.3.2.39] | 0 | 0 | 5 | 3 | 2 | 5 | 2 | 0 | 6 | 4 | 6 | 5 |
| K03896 acetyl CoA:N6-hydroxylysine acetyl transferase [EC:2.3.1.102] | 0 | 0 | 0 | 0 | 1 | 3 | 5 | 2 | 0 | 4 | 1 | 0 |
| K03897 lysine N6-hydroxylase [EC:1.14.13.59] | 0 | 0 | 1 | 0 | 3 | 0 | 0 | 4 | 6 | 14 | 3 | 9 |
| K09251 putrescine aminotransferase [EC:2.6.1.82] | 952 | 892 | 828 | 920 | 657 | 722 | 611 | 649 | 1272 | 1295 | 1186 | 1340 |
| K13609 delta1-piperideine-2-carboxylate reductase [EC:1.5.1.21] | 2009 | 1931 | 1872 | 1844 | 971 | 938 | 812 | 909 | 1270 | 1315 | 1210 | 1291 |
| K15736 L-2-hydroxyglutarate oxidase [EC:1.1.5.-] | 446 | 514 | 435 | 490 | 148 | 132 | 135 | 175 | 1074 | 1036 | 947 | 1114 |
| K15737 glutarate hydroxylase [EC:1.14.11.-] | 375 | 389 | 386 | 378 | 167 | 135 | 153 | 133 | 908 | 841 | 830 | 877 |
| K18011 beta-lysine 5,6-aminomutase beta subunit [EC:5.4.3.3] | 0 | 0 | 0 | 1 | 2 | 0 | 1 | 2 | 4 | 0 | 0 | 1 |
| K18012 L-erythro-3,5-diaminohexanoate dehydrogenase [EC:1.4.1.11] | 0 | 0 | 0 | 0 | 3 | 3 | 2 | 1 | 4 | 2 | 4 | 1 |
| K18013 3-keto-5-aminohexanoate cleavage enzyme [EC:2.3.1.247] | 33 | 33 | 34 | 33 | 10 | 5 | 6 | 8 | 52 | 54 | 42 | 64 |
| K18014 3-aminobutyryl-CoA ammonia-lyase [EC:4.3.1.14] | 0 | 0 | 0 | 0 | 1 | 4 | 1 | 2 | 2 | 0 | 3 | 2 |
| K21672 2,4-diaminopentanoate dehydrogenase [EC:1.4.1.12 1.4.1.26] | 352 | 395 | 378 | 380 | 86 | 88 | 98 | 97 | 245 | 298 | 257 | 286 |
| K23385 D-ornithine/D-lysine decarboxylase [EC:4.1.1.116] | 9 | 8 | 6 | 11 | 0 | 0 | 0 | 0 | 0 | 0 | 0 | 0 |
| ko00220 Arginine biosynthesis | 88438 | 87351 | 86788 | 87091 | 82048 | 81614 | 81599 | 82787 | 73934 | 73370 | 73133 | 73303 |
| K00145 N-acetyl-gamma-glutamyl-phosphate reductase [EC:1.2.1.38] | 3863 | 3704 | 3705 | 3740 | 4147 | 4166 | 4168 | 4274 | 3582 | 3664 | 3889 | 3713 |
| K00611 ornithine carbamoyltransferase [EC:2.1.3.3] | 4632 | 4603 | 4223 | 4517 | 2496 | 2427 | 2461 | 2531 | 2931 | 3050 | 2917 | 2933 |
| K00619 amino-acid N-acetyltransferase [EC:2.3.1.1] | 17 | 31 | 19 | 21 | 4 | 5 | 8 | 2 | 33 | 40 | 26 | 44 |
| K00620 glutamate N-acetyltransferase / amino-acid N-acetyltransferase [EC:2.3.1.35 2.3.1.1] | 2449 | 2332 | 2459 | 2494 | 1268 | 1313 | 1207 | 1291 | 1292 | 1226 | 1303 | 1266 |
| K00818 acetylornithine aminotransferase [EC:2.6.1.11] | 0 | 3 | 1 | 1 | 3 | 1 | 3 | 1 | 2 | 1 | 1 | 0 |
| K00930 acetylglutamate kinase [EC:2.7.2.8] | 2976 | 3045 | 2974 | 2812 | 3129 | 3150 | 3037 | 3011 | 3040 | 2852 | 2813 | 2924 |
| K01438 acetylornithine deacetylase [EC:3.5.1.16] | 5644 | 5532 | 5365 | 5417 | 5429 | 5466 | 5415 | 5537 | 4210 | 4031 | 3917 | 3952 |
| K01457 allophanate hydrolase [EC:3.5.1.54] | 401 | 333 | 316 | 417 | 532 | 607 | 527 | 569 | 0 | 0 | 2 | 0 |
| K01476 arginase [EC:3.5.3.1] | 330 | 351 | 343 | 337 | 207 | 222 | 250 | 209 | 640 | 626 | 603 | 652 |
| K01478 arginine deiminase [EC:3.5.3.6] | 2289 | 2248 | 2325 | 2369 | 1391 | 1330 | 1307 | 1307 | 979 | 930 | 902 | 1032 |
| K01941 urea carboxylase [EC:6.3.4.6] | 776 | 742 | 706 | 764 | 1071 | 1087 | 1083 | 1123 | 1 | 3 | 4 | 4 |
| K14677 aminoacylase [EC:3.5.1.14] | 1 | 0 | 3 | 1 | 0 | 1 | 1 | 0 | 0 | 1 | 0 | 2 |
| K14682 amino-acid N-acetyltransferase [EC:2.3.1.1] | 2813 | 2674 | 2591 | 2609 | 1617 | 1542 | 1486 | 1567 | 1753 | 1775 | 1615 | 1774 |
| K22477 N-acetylglutamate synthase [EC:2.3.1.1] | 150 | 139 | 122 | 126 | 29 | 20 | 19 | 29 | 116 | 119 | 136 | 117 |
| ko00330 Arginine and proline metabolism | 122379 | 121549 | 116445 | 119568 | 85355 | 84913 | 82894 | 86883 | 84914 | 83852 | 82967 | 84481 |
| K00147 glutamate-5-semialdehyde dehydrogenase [EC:1.2.1.41] | 3111 | 3201 | 3251 | 3186 | 4225 | 4097 | 4268 | 4323 | 4220 | 4145 | 4361 | 4216 |
| K00286 pyrroline-5-carboxylate reductase [EC:1.5.1.2] | 1964 | 1999 | 1944 | 1992 | 2477 | 2654 | 2443 | 2434 | 2616 | 2680 | 2738 | 2650 |
| K00316 spermidine dehydrogenase [EC:1.5.99.6] | 2957 | 3020 | 2903 | 2924 | 1666 | 1437 | 1342 | 1524 | 947 | 856 | 817 | 896 |
| K00318 proline dehydrogenase [EC:1.5.5.2] | 8 | 14 | 3 | 7 | 2 | 0 | 2 | 0 | 2 | 0 | 0 | 0 |
| K00472 prolyl 4-hydroxylase [EC:1.14.11.2] | 1 | 0 | 0 | 0 | 0 | 0 | 0 | 0 | 0 | 0 | 0 | 0 |
| K00657 diamine N-acetyltransferase [EC:2.3.1.57] | 1602 | 1692 | 1729 | 1763 | 1758 | 1840 | 1728 | 1747 | 1676 | 1784 | 1627 | 1624 |
| K00673 arginine N-succinyltransferase [EC:2.3.1.109] | 2463 | 2353 | 2273 | 2345 | 1256 | 1318 | 1306 | 1349 | 1934 | 2052 | 1914 | 2026 |
| K00819 ornithine--oxo-acid transaminase [EC:2.6.1.13] | 137 | 149 | 175 | 156 | 203 | 235 | 227 | 179 | 14 | 35 | 27 | 27 |
| K00840 succinylornithine aminotransferase [EC:2.6.1.81] | 3157 | 3003 | 3022 | 3106 | 1624 | 1658 | 1585 | 1761 | 2566 | 2339 | 2474 | 2452 |
| K00931 glutamate 5-kinase [EC:2.7.2.11] | 3625 | 3469 | 3519 | 3619 | 4005 | 3881 | 4175 | 4081 | 3738 | 3631 | 3897 | 3850 |
| K01259 proline iminopeptidase [EC:3.4.11.5] | 1733 | 1695 | 1748 | 1711 | 930 | 972 | 885 | 975 | 570 | 524 | 558 | 501 |
| K01426 amidase [EC:3.5.1.4] | 9167 | 9061 | 8492 | 8895 | 4978 | 4956 | 4644 | 5208 | 3233 | 3275 | 3176 | 3276 |
| K01470 creatinine amidohydrolase [EC:3.5.2.10] | 974 | 931 | 903 | 893 | 289 | 331 | 358 | 338 | 1348 | 1320 | 1252 | 1370 |
| K01473 N-methylhydantoinase A [EC:3.5.2.14] | 47 | 39 | 65 | 46 | 2 | 2 | 3 | 1 | 19 | 19 | 15 | 16 |
| K01474 N-methylhydantoinase B [EC:3.5.2.14] | 53 | 34 | 31 | 36 | 2 | 2 | 1 | 0 | 15 | 29 | 5 | 23 |
| K01480 agmatinase [EC:3.5.3.11] | 847 | 956 | 885 | 946 | 538 | 692 | 580 | 530 | 1294 | 1352 | 1290 | 1334 |
| K01484 succinylarginine dihydrolase [EC:3.5.3.23] | 2102 | 2175 | 1935 | 2222 | 1077 | 1177 | 1082 | 1133 | 2202 | 2309 | 2288 | 2375 |
| K01581 ornithine decarboxylase [EC:4.1.1.17] | 2931 | 2878 | 2973 | 3051 | 2098 | 2211 | 2148 | 2164 | 2776 | 2756 | 2679 | 2860 |
| K01584 arginine decarboxylase [EC:4.1.1.19] | 3205 | 3099 | 3012 | 3142 | 1650 | 1642 | 1540 | 1617 | 1275 | 1131 | 1263 | 1244 |
| K01585 arginine decarboxylase [EC:4.1.1.19] | 6227 | 6173 | 6159 | 6006 | 7212 | 7223 | 7308 | 7277 | 6418 | 6347 | 6530 | 6231 |
| K01750 ornithine cyclodeaminase [EC:4.3.1.12] | 1342 | 1356 | 1182 | 1287 | 642 | 632 | 602 | 731 | 477 | 474 | 422 | 509 |
| K01777 proline racemase [EC:5.1.1.4] | 66 | 71 | 80 | 66 | 84 | 90 | 93 | 109 | 111 | 106 | 112 | 121 |
| K02626 arginine decarboxylase [EC:4.1.1.19] | 8 | 5 | 9 | 7 | 0 | 2 | 0 | 0 | 1 | 4 | 2 | 0 |
| K05526 succinylglutamate desuccinylase [EC:3.5.1.96] | 1654 | 1639 | 1477 | 1561 | 817 | 806 | 804 | 774 | 1575 | 1724 | 1677 | 1790 |
| K06447 succinylglutamic semialdehyde dehydrogenase [EC:1.2.1.71] | 3752 | 3713 | 3505 | 3659 | 1914 | 1861 | 1884 | 1970 | 2955 | 2850 | 2897 | 3053 |
| K08660 cytosolic nonspecific dipeptidase [EC:3.4.13.18] | 1 | 0 | 0 | 0 | 0 | 0 | 0 | 0 | 0 | 0 | 0 | 0 |
| K09123 cis-L-3-hydroxyproline dehydratase [EC:4.2.1.171] | 1994 | 2004 | 1767 | 1902 | 963 | 933 | 925 | 1109 | 635 | 561 | 561 | 634 |
| K09470 gamma-glutamylputrescine synthase [EC:6.3.1.11] | 693 | 806 | 738 | 769 | 505 | 496 | 556 | 487 | 1067 | 1126 | 1094 | 1052 |
| K09471 gamma-glutamylputrescine oxidase [EC:1.4.3.-] | 6055 | 5978 | 5587 | 5777 | 2773 | 2866 | 2730 | 2892 | 3493 | 3458 | 3348 | 3370 |
| K09472 4-(gamma-glutamylamino)butanal dehydrogenase [EC:1.2.1.99] | 2648 | 2561 | 2434 | 2563 | 1299 | 1292 | 1154 | 1420 | 1722 | 1843 | 1677 | 1762 |
| K09473 gamma-glutamyl-gamma-aminobutyrate hydrolase [EC:3.5.1.94] | 287 | 283 | 305 | 278 | 69 | 76 | 87 | 78 | 589 | 684 | 658 | 629 |
| K10536 agmatine deiminase [EC:3.5.3.12] | 5492 | 5545 | 5061 | 5239 | 3165 | 3096 | 2936 | 3313 | 1783 | 1650 | 1584 | 1724 |
| K10793 D-proline reductase (dithiol) PrdA [EC:1.21.4.1] | 11 | 22 | 16 | 25 | 1 | 2 | 0 | 1 | 11 | 0 | 3 | 1 |
| K10794 D-proline reductase (dithiol) PrdB [EC:1.21.4.1] | 7 | 8 | 5 | 7 | 0 | 0 | 0 | 0 | 0 | 0 | 0 | 0 |
| K12251 N-carbamoylputrescine amidase [EC:3.5.1.53] | 3617 | 3497 | 3424 | 3406 | 3464 | 3402 | 3406 | 3615 | 2163 | 1835 | 1981 | 2032 |
| K12252 arginine:pyruvate transaminase [EC:2.6.1.84] | 2127 | 2046 | 1890 | 1988 | 1088 | 1051 | 958 | 1180 | 680 | 647 | 625 | 686 |
| K12253 5-guanidino-2-oxopentanoate decarboxylase [EC:4.1.1.75] | 2613 | 2606 | 2318 | 2444 | 1402 | 1225 | 1218 | 1324 | 857 | 754 | 780 | 796 |
| K12254 4-guanidinobutyraldehyde dehydrogenase / NAD-dependent aldehyde dehydrogenase [EC:1.2.1.54 1.2.1.-] | 2719 | 2717 | 2534 | 2661 | 1508 | 1569 | 1444 | 1547 | 887 | 816 | 865 | 853 |
| K12255 guanidinobutyrase [EC:3.5.3.7] | 1698 | 1767 | 1657 | 1755 | 765 | 794 | 701 | 772 | 1187 | 1137 | 1193 | 1104 |
| K12256 putrescine---pyruvate transaminase [EC:2.6.1.113] | 229 | 220 | 242 | 224 | 120 | 108 | 100 | 128 | 69 | 85 | 74 | 71 |
| K12658 4-hydroxyproline epimerase [EC:5.1.1.8] | 1522 | 1490 | 1187 | 1321 | 778 | 757 | 612 | 713 | 488 | 368 | 376 | 399 |
| K13747 carboxynorspermidine decarboxylase [EC:4.1.1.96] | 718 | 772 | 904 | 826 | 2351 | 2266 | 2464 | 2186 | 1701 | 1699 | 1737 | 1594 |
| K21061 D-hydroxyproline dehydrogenase subunit beta [EC:1.5.99.-] | 962 | 887 | 738 | 861 | 438 | 409 | 382 | 447 | 262 | 246 | 242 | 299 |
| K21062 1-pyrroline-4-hydroxy-2-carboxylate deaminase [EC:3.5.4.22] | 1407 | 1364 | 1276 | 1314 | 692 | 700 | 694 | 778 | 481 | 408 | 335 | 417 |
| K22548 trans-L-3-hydroxyproline dehydratase [EC:4.2.1.77] | 1301 | 1265 | 1169 | 1247 | 734 | 643 | 586 | 774 | 429 | 339 | 356 | 391 |
| K22549 D-hydroxyproline dehydrogenase subunit alpha [EC:1.5.99.-] | 1149 | 1075 | 842 | 1018 | 512 | 462 | 416 | 622 | 311 | 301 | 284 | 329 |
| K22550 D-hydroxyproline dehydrogenase subunit gamma | 335 | 315 | 228 | 286 | 155 | 134 | 129 | 155 | 78 | 81 | 96 | 87 |
| K23121 trans-4-hydroxy-L-proline dehydratase [EC:4.2.1.172] | 662 | 614 | 635 | 586 | 447 | 494 | 506 | 475 | 713 | 755 | 685 | 756 |

| | | | | | | | | | | | | |
|---|---|---|---|---|---|---|---|---|---|---|---|---|
| ko00340 Histidine metabolism | 60479 | 59535 | 59990 | 59197 | 62077 | 61787 | 62134 | 62424 | 52445 | 51474 | 51374 | 51745 |
| K00013 histidinol dehydrogenase [EC:1.1.1.23] | 2886 | 2775 | 2704 | 2896 | 2688 | 2738 | 2759 | 2880 | 2762 | 2862 | 2734 | 2838 |
| K00603 glutamate formiminotransferase / 5-formyltetrahydrofolate cyclo-ligase [EC:2.1.2.5 6.3.3.2] | 671 | 669 | 766 | 679 | 2041 | 2184 | 2185 | 2182 | 1583 | 1536 | 1581 | 1478 |
| K00765 ATP phosphoribosyltransferase [EC:2.4.2.17] | 2507 | 2343 | 2433 | 2365 | 2773 | 2844 | 2825 | 2874 | 2649 | 2733 | 2589 | 2667 |
| K00817 histidinol-phosphate aminotransferase [EC:2.6.1.9] | 7709 | 7663 | 7838 | 7675 | 8330 | 8147 | 8236 | 8130 | 6354 | 6206 | 6285 | 6231 |
| K01089 imidazoleglycerol-phosphate dehydratase / histidinol-phosphatase [EC:4.2.1.19 3.1.3.15] | 1369 | 1390 | 1460 | 1370 | 2724 | 2567 | 2822 | 2592 | 2418 | 2451 | 2545 | 2539 |
| K01468 imidazolonepropionase [EC:3.5.2.7] | 3878 | 3650 | 3709 | 3931 | 4276 | 4223 | 4403 | 4414 | 3847 | 3604 | 3650 | 3629 |
| K01479 formiminoglutamase [EC:3.5.3.8] | 1815 | 1699 | 1721 | 1699 | 903 | 855 | 814 | 811 | 1166 | 1054 | 1114 | 1260 |
| K01496 phosphoribosyl-AMP cyclohydrolase [EC:3.5.4.19] | 723 | 760 | 748 | 719 | 408 | 394 | 457 | 428 | 294 | 270 | 241 | 325 |
| K01523 phosphoribosyl-ATP pyrophosphohydrolase [EC:3.6.1.31] | 598 | 593 | 565 | 579 | 343 | 336 | 334 | 403 | 232 | 244 | 201 | 214 |
| K01590 histidine decarboxylase [EC:4.1.1.22] | 12 | 5 | 0 | 2 | 3 | 2 | 3 | 0 | 17 | 17 | 13 | 12 |
| K01663 imidazole glycerol-phosphate synthase [EC:4.3.2.10] | 0 | 0 | 1 | 0 | 0 | 0 | 0 | 0 | 0 | 0 | 0 | 0 |
| K01693 imidazoleglycerol-phosphate dehydratase [EC:4.2.1.19] | 1305 | 1271 | 1255 | 1142 | 681 | 631 | 662 | 735 | 615 | 617 | 555 | 619 |
| K01712 urocanate hydratase [EC:4.2.1.49] | 5722 | 5474 | 5473 | 5440 | 6589 | 6727 | 6833 | 6651 | 5795 | 5866 | 5642 | 5721 |
| K01745 histidine ammonia-lyase [EC:4.3.1.3] | 8324 | 8421 | 8336 | 8264 | 9886 | 9874 | 9767 | 9966 | 8177 | 7973 | 8036 | 8016 |
| K01814 phosphoribosylformimino-5-aminoimidazole carboxamide ribotide isomerase [EC:5.3.1.16] | 2363 | 2406 | 2548 | 2385 | 2522 | 2483 | 2418 | 2545 | 2494 | 2317 | 2438 | 2459 |
| K02500 imidazole glycerol-phosphate synthase subunit HisF [EC:4.3.2.10] | 3360 | 3295 | 3402 | 3348 | 3250 | 3334 | 3384 | 3172 | 2792 | 2729 | 2798 | 2790 |
| K02501 imidazole glycerol-phosphate synthase subunit HisH [EC:4.3.2.10] | 2696 | 2584 | 2685 | 2615 | 2605 | 2544 | 2557 | 2564 | 2305 | 2162 | 2172 | 2305 |
| K02502 ATP phosphoribosyltransferase regulatory subunit | 1479 | 1342 | 1329 | 1282 | 698 | 649 | 613 | 686 | 740 | 829 | 793 | 784 |
| K04486 histidinol-phosphatase (PHP family) [EC:3.1.3.15] | 3182 | 3250 | 3462 | 3204 | 4102 | 4281 | 4268 | 4165 | 3607 | 3425 | 3470 | 3391 |
| K05602 histidinol-phosphatase [EC:3.1.3.15] | 0 | 0 | 0 | 1 | 0 | 0 | 0 | 0 | 0 | 0 | 0 | 1 |
| K05603 formimidoylglutamate deiminase [EC:3.5.3.13] | 691 | 671 | 572 | 655 | 591 | 583 | 542 | 611 | 102 | 132 | 123 | 112 |
| K11755 phosphoribosyl-ATP pyrophosphohydrolase / phosphoribosyl-AMP cyclohydrolase [EC:3.6.1.31 3.5.4.19] | 1087 | 1062 | 1086 | 1079 | 1699 | 1681 | 1701 | 1658 | 1595 | 1669 | 1775 | 1661 |
| K13990 glutamate formiminotransferase / formiminotetrahydrofolate cyclodeaminase [EC:2.1.2.5 4.3.1.4] | 5 | 2 | 3 | 2 | 7 | 9 | 10 | 12 | 0 | 1 | 3 | 2 |
| K14152 phosphoribosyl-ATP pyrophosphohydrolase / phosphoribosyl-AMP cyclohydrolase / histidinol dehydrogenase [EC:3.6.1.31 3.5.4.19 1.1.1.23] | 0 | 0 | 0 | 0 | 0 | 0 | 0 | 1 | 0 | 0 | 0 | 0 |
| K17363 urocanate reductase [EC:1.3.99.33] | 12 | 21 | 19 | 22 | 22 | 17 | 25 | 19 | 32 | 43 | 46 | 42 |
| ko00350 Tyrosine metabolism | 63187 | 61811 | 60724 | 62184 | 42265 | 41713 | 40887 | 42573 | 46100 | 46102 | 45123 | 46338 |
| K00055 aryl-alcohol dehydrogenase [EC:1.1.1.90] | 406 | 379 | 402 | 465 | 104 | 135 | 95 | 139 | 881 | 891 | 833 | 844 |
| K00151 5-carboxymethyl-2-hydroxymuconic-semialdehyde dehydrogenase [EC:1.2.1.60] | 2534 | 2420 | 2364 | 2407 | 1194 | 1183 | 1010 | 1175 | 1775 | 1746 | 1655 | 1830 |
| K00450 gentisate 1,2-dioxygenase [EC:1.13.11.4] | 1483 | 1452 | 1354 | 1397 | 726 | 707 | 746 | 830 | 472 | 464 | 416 | 396 |
| K00451 homogentisate 1,2-dioxygenase [EC:1.13.11.5] | 393 | 434 | 343 | 340 | 189 | 174 | 192 | 215 | 136 | 125 | 109 | 134 |
| K00455 3,4-dihydroxyphenylacetate 2,3-dioxygenase [EC:1.13.11.15] | 1658 | 1562 | 1439 | 1461 | 765 | 724 | 742 | 793 | 1088 | 1163 | 1061 | 1141 |
| K00457 4-hydroxyphenylpyruvate dioxygenase [EC:1.13.11.27] | 5276 | 5060 | 4895 | 4938 | 2674 | 2608 | 2324 | 2809 | 2760 | 2716 | 2748 | 2897 |
| K00483 4-hydroxyphenylacetate 3-monooxygenase [EC:1.14.14.9] | 3388 | 3198 | 3008 | 3277 | 1796 | 1609 | 1542 | 1705 | 2360 | 2156 | 1982 | 2181 |
| K00484 flavin reductase (NADH) [EC:1.5.1.36] | 584 | 571 | 486 | 500 | 234 | 242 | 273 | 281 | 515 | 534 | 479 | 541 |
| K01555 fumarylacetoacetase [EC:3.7.1.2] | 1681 | 1512 | 1501 | 1683 | 902 | 848 | 762 | 854 | 524 | 449 | 449 | 457 |
| K01557 acylpyruvate hydrolase [EC:3.7.1.5] | 1 | 0 | 0 | 0 | 0 | 0 | 0 | 0 | 0 | 0 | 0 | 0 |
| K01593 aromatic-L-amino-acid/L-tryptophan decarboxylase [EC:4.1.1.28 4.1.1.105] | 0 | 1 | 0 | 0 | 0 | 0 | 0 | 0 | 0 | 0 | 0 | 0 |
| K01668 tyrosine phenol-lyase [EC:4.1.99.2] | 306 | 280 | 273 | 283 | 359 | 405 | 407 | 360 | 13 | 14 | 5 | 8 |
| K01800 maleylacetoacetate isomerase [EC:5.2.1.2] | 1700 | 1716 | 1539 | 1668 | 919 | 833 | 803 | 929 | 556 | 464 | 459 | 513 |
| K01826 5-carboxymethyl-2-hydroxymuconate isomerase [EC:5.3.3.10] | 1071 | 1058 | 949 | 1044 | 495 | 546 | 491 | 537 | 587 | 609 | 574 | 636 |
| K02509 2-oxo-hept-3-ene-1,7-dioate hydratase [EC:4.2.1.-] | 1720 | 1620 | 1518 | 1636 | 853 | 839 | 774 | 958 | 1117 | 1075 | 1054 | 1123 |
| K02510 4-hydroxy-2-oxoheptanedioate aldolase [EC:4.1.2.52] | 2182 | 2003 | 1932 | 2165 | 1061 | 1147 | 1103 | 1118 | 1196 | 1210 | 1194 | 1248 |
| K05921 5-oxopent-3-ene-1,2,5-tricarboxylate decarboxylase / 2-hydroxyhepta-2,4-diene-1,7-dioate isomerase [EC:4.1.1.68 5.3.3.-] | 2889 | 2653 | 2638 | 2757 | 1384 | 1475 | 1328 | 1487 | 1862 | 1784 | 1707 | 1825 |
| K10219 2-hydroxy-4-carboxymuconate semialdehyde hemiacetal dehydrogenase [EC:1.1.1.312] | 8 | 5 | 7 | 11 | 0 | 0 | 0 | 0 | 4 | 6 | 4 | 4 |
| K16165 fumarylpyruvate hydrolase [EC:3.7.1.20] | 150 | 140 | 146 | 142 | 111 | 89 | 110 | 116 | 64 | 67 | 80 | 76 |
| K16901 anthranilate 3-monooxygenase (FAD) / 4-hydroxyphenylacetate 3-monooxygenase [EC:1.14.14.8 1.14.14.9] | 1 | 0 | 0 | 0 | 0 | 0 | 0 | 2 | 0 | 0 | 0 | 0 |
| K22330 tyrosine decarboxylase [EC:4.1.1.25] | 97 | 104 | 100 | 124 | 38 | 35 | 28 | 30 | 107 | 159 | 119 | 133 |
| K23109 aromatic 2-oxoacid reductase [EC:1.1.1.110] | 3 | 5 | 12 | 9 | 3 | 3 | 2 | 0 | 16 | 20 | 14 | 18 |
| ko00360 Phenylalanine metabolism | 67599 | 67361 | 65044 | 66281 | 47773 | 47165 | 46342 | 48668 | 56908 | 57652 | 56287 | 58293 |
| K00146 phenylacetaldehyde dehydrogenase [EC:1.2.1.39] | 3074 | 3228 | 2751 | 2991 | 1383 | 1316 | 1355 | 1467 | 2754 | 2743 | 2608 | 2749 |
| K00285 D-amino-acid dehydrogenase [EC:1.4.5.1] | 5374 | 5202 | 5358 | 5270 | 3033 | 2959 | 2812 | 3147 | 3184 | 3145 | 3203 | 3362 |
| K00500 phenylalanine-4-hydroxylase [EC:1.14.16.1] | 1235 | 1169 | 1118 | 1207 | 678 | 690 | 622 | 661 | 367 | 382 | 315 | 350 |
| K01451 hippurate hydrolase [EC:3.5.1.32] | 4297 | 4273 | 4089 | 4137 | 2111 | 1923 | 1897 | 2168 | 2794 | 2838 | 2619 | 2805 |
| K01666 4-hydroxy 2-oxovalerate aldolase [EC:4.1.3.39] | 424 | 414 | 433 | 447 | 96 | 102 | 116 | 85 | 812 | 819 | 761 | 789 |
| K01912 phenylacetate-CoA ligase [EC:6.2.1.30] | 2593 | 2821 | 2790 | 2674 | 5548 | 5595 | 5892 | 5643 | 4880 | 4983 | 4988 | 5098 |
| K02554 2-keto-4-pentenoate hydratase [EC:4.2.1.80] | 413 | 442 | 449 | 407 | 324 | 419 | 357 | 333 | 583 | 580 | 637 | 651 |
| K02609 ring-1,2-phenylacetyl-CoA epoxidase subunit PaaA [EC:1.14.13.149] | 388 | 358 | 346 | 401 | 95 | 79 | 94 | 109 | 744 | 692 | 696 | 750 |
| K02610 ring-1,2-phenylacetyl-CoA epoxidase subunit PaaB | 128 | 119 | 100 | 103 | 46 | 26 | 29 | 39 | 223 | 216 | 216 | 217 |
| K02611 ring-1,2-phenylacetyl-CoA epoxidase subunit PaaC [EC:1.14.13.149] | 270 | 274 | 276 | 307 | 86 | 86 | 66 | 87 | 512 | 585 | 558 | 657 |
| K02612 ring-1,2-phenylacetyl-CoA epoxidase subunit PaaD | 174 | 190 | 195 | 191 | 43 | 52 | 53 | 54 | 322 | 421 | 430 | 413 |
| K02613 ring-1,2-phenylacetyl-CoA epoxidase subunit PaaE | 310 | 428 | 410 | 390 | 101 | 99 | 113 | 102 | 726 | 826 | 860 | 895 |
| K02614 acyl-CoA thioesterase [EC:3.1.2.-] | 1143 | 1167 | 1155 | 1101 | 1313 | 1300 | 1305 | 1427 | 1301 | 1313 | 1249 | 1267 |
| K02615 3-oxo-5,6-didehydrosuberyl-CoA/3-oxoadipyl-CoA thiolase [EC:2.3.1.223 2.3.1.174] | 430 | 473 | 382 | 441 | 134 | 106 | 114 | 133 | 885 | 900 | 899 | 923 |
| K02618 oxepin-CoA hydrolase / 3-oxo-5,6-dehydrosuberyl-CoA semialdehyde dehydrogenase [EC:3.3.2.12 1.2.1.91] | 671 | 634 | 647 | 693 | 194 | 211 | 207 | 184 | 1352 | 1424 | 1402 | 1494 |
| K03782 catalase-peroxidase [EC:1.11.1.21] | 1258 | 1181 | 1142 | 1263 | 789 | 896 | 772 | 806 | 1641 | 1801 | 1708 | 1710 |
| K05708 3-phenylpropionate/trans-cinnamate dioxygenase subunit alpha [EC:1.14.12.19] | 0 | 0 | 0 | 0 | 11 | 3 | 8 | 4 | 4 | 1 | 3 | 1 |
| K05709 3-phenylpropionate/trans-cinnamate dioxygenase subunit beta [EC:1.14.12.19] | 0 | 0 | 0 | 0 | 2 | 2 | 3 | 3 | 0 | 1 | 1 | 1 |
| K05710 3-phenylpropionate/trans-cinnamate dioxygenase ferredoxin component | 0 | 1 | 0 | 0 | 3 | 4 | 4 | 3 | 1 | 0 | 2 | 4 |
| K05711 2,3-dihydroxy-2,3-dihydrophenylpropionate dehydrogenase [EC:1.3.1.87] | 0 | 0 | 0 | 0 | 10 | 10 | 4 | 5 | 4 | 2 | 6 | 2 |
| K05712 3-(3-hydroxy-phenyl)propionate hydroxylase [EC:1.14.13.127] | 601 | 533 | 477 | 546 | 157 | 190 | 148 | 178 | 1041 | 1126 | 1068 | 1154 |
| K05713 2,3-dihydroxyphenylpropionate 1,2-dioxygenase [EC:1.13.11.16] | 352 | 356 | 276 | 304 | 101 | 81 | 98 | 84 | 683 | 757 | 658 | 745 |
| K05714 2-hydroxy-6-oxonona-2,4-dienedioate hydrolase [EC:3.7.1.14] | 283 | 293 | 266 | 314 | 98 | 81 | 84 | 100 | 648 | 690 | 614 | 702 |
| K10797 2-enoate reductase [EC:1.3.1.31] | 1240 | 1260 | 1193 | 1190 | 199 | 187 | 213 | 189 | 1244 | 1330 | 1284 | 1369 |
| K15866 2-(1,2-epoxy-1,2-dihydrophenyl)acetyl-CoA isomerase [EC:5.3.3.18] | 1225 | 1220 | 1112 | 1080 | 588 | 529 | 521 | 643 | 866 | 851 | 866 | 872 |
| K18356 phenylglyoxylate dehydrogenase beta subunit [EC:1.2.1.58] | 1 | 2 | 0 | 1 | 0 | 0 | 0 | 0 | 5 | 3 | 7 | 13 |
| ko00380 Tryptophan metabolism | 67727 | 67568 | 65289 | 66791 | 46993 | 46039 | 45017 | 47573 | 42448 | 41918 | 41277 | 42338 |
| K00452 3-hydroxyanthranilate 3,4-dioxygenase [EC:1.13.11.6] | 1 | 0 | 0 | 0 | 0 | 2 | 0 | 0 | 0 | 0 | 1 | 0 |
| K00453 tryptophan 2,3-dioxygenase [EC:1.13.11.11] | 1198 | 1272 | 1328 | 1245 | 689 | 730 | 729 | 786 | 355 | 366 | 356 | 383 |
| K01556 kynureninase [EC:3.7.1.3] | 450 | 470 | 445 | 444 | 238 | 240 | 210 | 217 | 133 | 163 | 127 | 138 |
| K01667 tryptophanase [EC:4.1.99.1] | 992 | 1068 | 1125 | 1117 | 3054 | 2955 | 3140 | 3031 | 1964 | 1901 | 2035 | 1842 |
| K03392 aminocarboxymuconate-semialdehyde decarboxylase [EC:4.1.1.45] | 14 | 9 | 16 | 22 | 3 | 0 | 2 | 2 | 4 | 0 | 3 | 9 |
| K04103 indolepyruvate decarboxylase [EC:4.1.1.74] | 693 | 702 | 727 | 670 | 203 | 190 | 229 | 198 | 1443 | 1568 | 1374 | 1535 |
| K10217 aminomuconate-semialdehyde/2-hydroxymuconate-6-semialdehyde dehydrogenase [EC:1.2.1.32 1.2.1.85] | 1 | 0 | 0 | 0 | 0 | 0 | 0 | 0 | 0 | 0 | 0 | 0 |
| K14264 kynurenine aminotransferase [EC:2.6.1.7] | 0 | 0 | 0 | 0 | 0 | 0 | 0 | 1 | 0 | 0 | 0 | 0 |
| K20807 nitrile hydratase subunit beta [EC:4.2.1.84] | 0 | 0 | 0 | 0 | 0 | 0 | 0 | 0 | 1 | 0 | 0 | 0 |
| K21801 indoleacetamide hydrolase [EC:3.5.1.-] | 1 | 0 | 0 | 0 | 0 | 0 | 0 | 0 | 0 | 0 | 0 | 0 |
| K23384 indoleacetate decarboxylase [EC:4.1.1.115] | 6 | 6 | 7 | 9 | 0 | 0 | 0 | 0 | 0 | 2 | 1 | 0 |
| ko00400 Phenylalanine, tyrosine and tryptophan biosynthesis | 82297 | 82245 | 82864 | 82360 | 80409 | 79695 | 80181 | 79849 | 78406 | 78029 | 77696 | 78041 |
| K00014 shikimate dehydrogenase [EC:1.1.1.25] | 4001 | 3925 | 4135 | 4063 | 3512 | 3324 | 3485 | 3433 | 4683 | 4655 | 4415 | 4703 |
| K00210 prephenate dehydrogenase [EC:1.3.1.12] | 484 | 593 | 611 | 550 | 1720 | 1747 | 1773 | 1642 | 1302 | 1179 | 1262 | 1227 |
| K00220 cyclohexadieny/prephenate dehydrogenase [EC:1.3.1.43 1.3.1.12] | 2 | 0 | 1 | 1 | 3 | 0 | 2 | 3 | 2 | 3 | 4 | 1 |
| K00766 anthranilate phosphoribosyltransferase [EC:2.4.2.18] | 2809 | 2797 | 2832 | 2778 | 3137 | 3063 | 3089 | 3018 | 2440 | 2233 | 2427 | 2257 |
| K00800 3-phosphoshikimate 1-carboxyvinyltransferase [EC:2.5.1.19] | 5576 | 5694 | 5740 | 5617 | 5392 | 5341 | 5479 | 5349 | 4614 | 4702 | 4753 | 4710 |
| K00891 shikimate kinase [EC:2.7.1.71] | 2326 | 2469 | 2468 | 2371 | 2276 | 2358 | 2300 | 2480 | 2640 | 2612 | 2515 | 2510 |
| K01609 indole-3-glycerol phosphate synthase [EC:4.1.1.48] | 2147 | 2194 | 2150 | 2113 | 2385 | 2239 | 2417 | 2510 | 1794 | 1763 | 1912 | 1820 |
| K01626 3-deoxy-7-phosphoheptulonate synthase [EC:2.5.1.54] | 8495 | 8301 | 8246 | 8294 | 4738 | 4501 | 4268 | 4534 | 5044 | 5246 | 4836 | 5059 |
| K01656 anthranilate synthase / indole-3-glycerol phosphate synthase [EC:4.1.3.27 4.1.1.48] | 0 | 0 | 0 | 0 | 0 | 1 | 0 | 0 | 0 | 0 | 0 | 0 |
| K01657 anthranilate synthase component I [EC:4.1.3.27] | 5953 | 6024 | 6120 | 5943 | 5535 | 5595 | 5523 | 5413 | 5197 | 4925 | 4909 | 4982 |
| K01658 anthranilate synthase component II [EC:4.1.3.27] | 2717 | 2705 | 2636 | 2690 | 2255 | 2308 | 2218 | 2346 | 1660 | 1627 | 1614 | 1567 |
| K01713 cyclohexadienyl dehydratase [EC:4.2.1.51 4.2.1.91] | 1488 | 1454 | 1382 | 1433 | 714 | 631 | 634 | 695 | 1043 | 1112 | 1022 | 1021 |
| K01735 3-dehydroquinate synthase [EC:4.2.3.4] | 3884 | 3778 | 3727 | 3933 | 4281 | 4278 | 4111 | 4072 | 3840 | 3881 | 3949 | 3808 |
| K01736 chorismate synthase [EC:4.2.3.5] | 3778 | 3757 | 3785 | 3685 | 3912 | 4003 | 4219 | 3944 | 3782 | 3981 | 3958 | 4017 |
| K01817 phosphoribosylanthranilate isomerase [EC:5.3.1.24] | 1294 | 1299 | 1349 | 1259 | 1575 | 1536 | 1605 | 1633 | 1362 | 1326 | 1407 | 1352 |
| K03785 3-dehydroquinate dehydratase I [EC:4.2.1.10] | 616 | 615 | 628 | 653 | 476 | 487 | 533 | 458 | 804 | 989 | 865 | 930 |
| K03786 3-dehydroquinate dehydratase II [EC:4.2.1.10] | 2215 | 2284 | 2358 | 2375 | 2111 | 2065 | 2051 | 2081 | 1753 | 1788 | 1743 | 1775 |
| K03856 3-deoxy-7-phosphoheptulonate synthase [EC:2.5.1.54] | 646 | 605 | 667 | 619 | 206 | 224 | 246 | 213 | 744 | 785 | 659 | 743 |
| K04092 chorismate mutase [EC:5.4.99.5] | 1 | 0 | 2 | 1 | 3 | 1 | 0 | 0 | 4 | 10 | 6 | 9 |
| K04093 chorismate mutase [EC:5.4.99.5] | 920 | 980 | 844 | 933 | 538 | 522 | 534 | 504 | 314 | 277 | 257 | 310 |
| K04516 chorismate mutase [EC:5.4.99.5] | 676 | 727 | 806 | 726 | 2316 | 2326 | 2246 | 2317 | 1689 | 1525 | 1738 | 1578 |
| K04517 prephenate dehydrogenase [EC:1.3.1.12] | 631 | 620 | 627 | 618 | 205 | 211 | 172 | 182 | 615 | 734 | 637 | 621 |
| K04518 prephenate dehydratase [EC:4.2.1.51] | 560 | 658 | 751 | 655 | 1974 | 1931 | 2037 | 1926 | 1422 | 1411 | 1429 | 1473 |
| K05358 quinate dehydrogenase (quinone) [EC:1.1.5.8] | 1340 | 1300 | 1259 | 1230 | 802 | 901 | 823 | 811 | 1722 | 1761 | 1619 | 1750 |
| K05359 arogenate/prephenate dehydratase [EC:4.2.1.91 4.2.1.51] | 0 | 0 | 0 | 0 | 0 | 0 | 0 | 0 | 2 | 2 | 0 | 4 |
| K05887 quinate/shikimate dehydrogenase [EC:1.1.1.282] | 36 | 54 | 49 | 36 | 89 | 70 | 89 | 65 | 98 | 88 | 98 | 116 |
| K06208 chorismate mutase [EC:5.4.99.5] | 0 | 1 | 0 | 2 | 0 | 0 | 0 | 0 | 6 | 12 | 5 | 5 |
| K06209 chorismate mutase [EC:5.4.99.5] | 268 | 237 | 261 | 275 | 60 | 85 | 85 | 89 | 226 | 248 | 262 | 289 |
| K11646 3-dehydroquinate synthase II [EC:1.4.1.24] | 15 | 16 | 12 | 19 | 16 | 2 | 5 | 6 | 21 | 25 | 10 | 27 |
| K13497 anthranilate synthase/phosphoribosyltransferase [EC:4.1.3.27 2.4.2.18] | 969 | 943 | 912 | 966 | 572 | 623 | 600 | 602 | 1301 | 1384 | 1442 | 1367 |
| K13498 indole-3-glycerol phosphate synthase / phosphoribosylanthranilate isomerase [EC:4.1.1.48 5.3.1.24] | 811 | 744 | 756 | 813 | 546 | 540 | 595 | 545 | 1107 | 1076 | 971 | 1128 |
| K13503 anthranilate synthase [EC:4.1.3.27] | 0 | 0 | 0 | 0 | 2 | 7 | 1 | 2 | 2 | 6 | 3 | 0 |

| | | | | | | | | | | | | |
|---|---|---|---|---|---|---|---|---|---|---|---|---|
| K13829 shikimate kinase / 3-dehydroquinate synthase [EC:2.7.1.71 4.2.3.4] | 10 | 15 | 8 | 15 | 26 | 14 | 21 | 19 | 25 | 25 | 24 | 32 |
| K13832 3-dehydroquinate dehydratase / shikimate dehydrogenase [EC:4.2.1.10 1.1.1.25] | 0 | 2 | 2 | 1 | 2 | 2 | 3 | 0 | 1 | 0 | 0 | 1 |
| K13853 3-deoxy-7-phosphoheptulonate synthase / chorismate mutase [EC:2.5.1.54 5.4.99.5] | 0 | 0 | 2 | 2 | 0 | 0 | 1 | 0 | 0 | 1 | 2 | 1 |
| K14170 chorismate mutase / prephenate dehydratase [EC:5.4.99.5 4.2.1.51] | 3001 | 2781 | 2992 | 2825 | 1598 | 1754 | 1517 | 1626 | 2205 | 2038 | 2000 | 2037 |
| K14187 chorismate mutase / prephenate dehydrogenase [EC:5.4.99.5 1.3.1.12] | 772 | 691 | 649 | 743 | 487 | 568 | 544 | 502 | 964 | 996 | 968 | 1025 |
| K15652 3-dehydroshikimate dehydratase [EC:4.2.1.118] | 0 | 0 | 1 | 0 | 1 | 0 | 0 | 0 | 0 | 0 | 0 | 0 |
| K17749 3-benzylmalate isomerase [EC:4.2.1.-] | 1 | 0 | 0 | 0 | 0 | 0 | 0 | 0 | 0 | 0 | 0 | 0 |
| K17750 3-benzylmalate dehydrogenase [EC:1.1.1.-] | 0 | 0 | 0 | 0 | 0 | 0 | 0 | 0 | 0 | 1 | 0 | 0 |
| Metabolism of other amino acids | 285297 | 284883 | 280899 | 284452 | 282862 | 283029 | 281654 | 282129 | 282463 | 280216 | 279210 | 280375 |
| ko00410 beta-Alanine metabolism | 54483 | 53513 | 52036 | 52952 | 36094 | 34911 | 33537 | 35631 | 33584 | 33314 | 32046 | 33284 |
| K01579 aspartate 1-decarboxylase [EC:4.1.1.11] | 968 | 1037 | 1032 | 990 | 1278 | 1240 | 1193 | 1165 | 1013 | 1069 | 1076 | 1027 |
| K01918 pantoate--beta-alanine ligase [EC:6.3.2.1] | 2715 | 2495 | 2689 | 2623 | 3123 | 3153 | 3021 | 3109 | 2737 | 2592 | 2593 | 2551 |
| K09722 4-phosphopantoate---beta-alanine ligase [EC:6.3.2.36] | 1 | 0 | 0 | 1 | 1 | 0 | 0 | 0 | 2 | 2 | 2 | 3 |
| K15372 taurine---2-oxoglutarate transaminase [EC:2.6.1.55] | 852 | 915 | 766 | 875 | 512 | 591 | 496 | 535 | 1206 | 1209 | 1140 | 1240 |
| ko00430 Taurine and hypotaurine metabolism | 30793 | 30662 | 29791 | 30251 | 25650 | 25797 | 25113 | 26040 | 23162 | 22413 | 22532 | 23041 |
| K00681 gamma-glutamyltranspeptidase / glutathione hydrolase [EC:2.3.2.2 3.4.19.13] | 9341 | 8938 | 8847 | 9001 | 5414 | 5263 | 5126 | 5548 | 4207 | 4148 | 4127 | 4293 |
| K03851 taurine-pyruvate aminotransferase [EC:2.6.1.77] | 10 | 6 | 16 | 9 | 3 | 5 | 0 | 2 | 10 | 7 | 4 | 4 |
| K03852 sulfoacetaldehyde acetyltransferase [EC:2.3.3.15] | 8 | 18 | 6 | 7 | 7 | 5 | 2 | 6 | 23 | 20 | 20 | 39 |
| K15373 sulfoacetaldehyde reductase [EC:1.1.1.313] | 497 | 507 | 468 | 509 | 281 | 313 | 344 | 323 | 643 | 627 | 611 | 763 |
| ko00440 Phosphonate and phosphinate metabolism | 17247 | 17191 | 16246 | 16839 | 10181 | 9988 | 9611 | 10350 | 14425 | 14444 | 13863 | 14440 |
| K01841 phosphoenolpyruvate phosphomutase [EC:5.4.2.9] | 132 | 132 | 142 | 146 | 74 | 64 | 66 | 63 | 83 | 85 | 108 | 71 |
| K03430 2-aminoethylphosphonate-pyruvate transaminase [EC:2.6.1.37] | 2348 | 2353 | 2331 | 2360 | 1232 | 1341 | 1253 | 1410 | 1969 | 2037 | 1902 | 1914 |
| K03823 phosphinothricin acetyltransferase [EC:2.3.1.183] | 2141 | 2223 | 2057 | 2159 | 1490 | 1407 | 1447 | 1485 | 1689 | 1680 | 1599 | 1732 |
| K05306 phosphonoacetaldehyde hydrolase [EC:3.11.1.1] | 1963 | 1815 | 1677 | 1838 | 1005 | 1000 | 1026 | 1046 | 1320 | 1424 | 1312 | 1387 |
| K05780 alpha-D-ribose 1-methylphosphonate 5-triphosphate synthase subunit PhnL [EC:2.7.8.37] | 1001 | 964 | 912 | 976 | 425 | 424 | 385 | 456 | 1000 | 962 | 919 | 1019 |
| K06162 alpha-D-ribose 1-methylphosphonate 5-triphosphate diphosphatase [EC:3.6.1.63] | 1845 | 1853 | 1663 | 1690 | 824 | 774 | 704 | 795 | 1543 | 1395 | 1414 | 1547 |
| K06163 alpha-D-ribose 1-methylphosphonate 5-phosphate C-P lyase [EC:4.7.1.1] | 1651 | 1657 | 1628 | 1612 | 786 | 811 | 741 | 844 | 1254 | 1251 | 1214 | 1243 |
| K06164 alpha-D-ribose 1-methylphosphonate 5-triphosphate synthase subunit PhnI [EC:2.7.8.37] | 1761 | 1656 | 1558 | 1683 | 822 | 723 | 665 | 923 | 1422 | 1451 | 1302 | 1410 |
| K06165 alpha-D-ribose 1-methylphosphonate 5-triphosphate synthase subunit PhnH [EC:2.7.8.37] | 1260 | 1315 | 1096 | 1263 | 589 | 523 | 511 | 602 | 898 | 891 | 788 | 858 |
| K06166 alpha-D-ribose 1-methylphosphonate 5-triphosphate synthase subunit PhnG [EC:2.7.8.37] | 963 | 900 | 835 | 889 | 488 | 372 | 421 | 398 | 706 | 692 | 695 | 688 |
| K06167 phosphoribosyl 1,2-cyclic phosphate phosphodiesterase [EC:3.1.4.55] | 2000 | 2144 | 2173 | 2030 | 2270 | 2357 | 2183 | 2149 | 2220 | 2221 | 2220 | 2209 |
| K09459 phosphonopyruvate decarboxylase [EC:4.1.1.82] | 72 | 66 | 50 | 64 | 35 | 30 | 41 | 27 | 109 | 107 | 124 | 122 |
| K09994 (aminoalkyl)phosphonate N-acetyltransferase [EC:2.3.1.280] | 12 | 25 | 16 | 18 | 3 | 6 | 7 | 6 | 43 | 44 | 61 | 84 |
| K19670 phosphonoacetate hydrolase [EC:3.11.1.2] | 72 | 65 | 64 | 82 | 127 | 138 | 146 | 128 | 105 | 92 | 114 | 89 |
| K20859 phosphoribosyl 1,2-cyclic phosphate 1,2-diphosphodiesterase [EC:3.1.4.57] | 24 | 23 | 44 | 29 | 8 | 11 | 10 | 16 | 57 | 103 | 87 | 56 |
| ko00450 Selenocompound metabolism | 49483 | 49507 | 49234 | 50116 | 48072 | 49879 | 48981 | 48250 | 55905 | 55802 | 55947 | 55855 |
| K00384 thioredoxin reductase (NADPH) [EC:1.8.1.9] | 5048 | 5103 | 5254 | 5193 | 4075 | 4326 | 4004 | 3898 | 5206 | 5283 | 5317 | 5310 |
| K01008 selenide, water dikinase [EC:2.7.9.3] | 2726 | 2788 | 2603 | 2646 | 1346 | 1332 | 1280 | 1444 | 1713 | 1789 | 1707 | 1815 |
| K01042 L-seryl-tRNA(Ser) seleniumtransferase [EC:2.9.1.1] | 2558 | 2671 | 2455 | 2469 | 1542 | 1562 | 1633 | 1584 | 2588 | 2805 | 2590 | 2854 |
| K01874 methionyl-tRNA synthetase [EC:6.1.1.10] | 6412 | 6714 | 6299 | 6516 | 7488 | 7743 | 7705 | 7311 | 7785 | 7475 | 7715 | 7530 |
| K07309 Tat-targeted selenate reductase subunit YnfE [EC:1.97.1.9] | 1255 | 1258 | 1157 | 1358 | 893 | 967 | 820 | 875 | 1950 | 1859 | 1894 | 1844 |
| K07310 Tat-targeted selenate reductase subunit YnfF [EC:1.97.1.9] | 5 | 11 | 11 | 8 | 34 | 39 | 29 | 33 | 114 | 144 | 137 | 114 |
| K11717 cysteine desulfurase / selenocysteine lyase [EC:2.8.1.7 4.4.1.16] | 1734 | 1738 | 1834 | 1809 | 3381 | 3485 | 3574 | 3359 | 3340 | 3274 | 3425 | 3343 |
| K12527 putative selenate reductase [EC:1.97.1.9] | 1310 | 1273 | 1237 | 1279 | 392 | 470 | 448 | 416 | 1301 | 1482 | 1357 | 1361 |
| K12528 putative selenate reductase molybdopterin-binding subunit | 16 | 10 | 17 | 6 | 38 | 34 | 44 | 33 | 173 | 170 | 187 | 189 |
| K12529 putative selenate reductase FAD-binding subunit | 0 | 1 | 0 | 1 | 9 | 6 | 11 | 10 | 43 | 41 | 37 | 58 |
| ko00460 Cyanoamino acid metabolism | 54789 | 55753 | 56554 | 55832 | 93081 | 91752 | 94310 | 92020 | 83690 | 82659 | 84193 | 81963 |
| K10814 hydrogen cyanide synthase HcnA [EC:1.4.99.5] | 397 | 408 | 399 | 379 | 236 | 188 | 190 | 240 | 150 | 137 | 112 | 102 |
| K10815 hydrogen cyanide synthase HcnB [EC:1.4.99.5] | 652 | 653 | 577 | 703 | 333 | 293 | 316 | 354 | 190 | 155 | 193 | 197 |
| K10816 hydrogen cyanide synthase HcnC [EC:1.4.99.5] | 1731 | 1712 | 1501 | 1742 | 869 | 850 | 793 | 913 | 489 | 493 | 486 | 471 |
| K18282 cyanide dihydratase [EC:3.5.5.-] | 0 | 0 | 0 | 0 | 0 | 0 | 0 | 1 | 0 | 0 | 0 | 0 |
| ko00471 D-Glutamine and D-glutamate metabolism | 15224 | 15081 | 14944 | 15213 | 16826 | 17335 | 17417 | 17037 | 17085 | 16882 | 17014 | 17054 |
| K01776 glutamate racemase [EC:5.1.1.3] | 2921 | 2983 | 2975 | 3007 | 3151 | 3357 | 3239 | 3144 | 3002 | 3005 | 3000 | 2980 |
| K01924 UDP-N-acetylmuramate--alanine ligase [EC:6.3.2.8] | 5304 | 5105 | 5096 | 5223 | 5477 | 5627 | 5569 | 5659 | 5370 | 5179 | 5158 | 5376 |
| K01925 UDP-N-acetylmuramoylalanine--D-glutamate ligase [EC:6.3.2.9] | 4613 | 4697 | 4526 | 4574 | 5196 | 5177 | 5211 | 5184 | 4866 | 4958 | 4854 | 4875 |
| K22210 D-glutamate cyclase [EC:4.2.1.48] | 1 | 0 | 0 | 1 | 0 | 0 | 0 | 0 | 0 | 0 | 0 | 1 |
| K23169 UDP-N-acetylmuramoyl-L-alanine---L-glutamate ligase [EC:6.3.2.53] | 75 | 73 | 68 | 64 | 8 | 19 | 15 | 23 | 37 | 52 | 40 | 44 |
| K23170 UDP-N-acetyl-alpha-D-muramoyl-L-alanyl-L-glutamate epimerase [EC:5.1.1.23] | 58 | 80 | 66 | 42 | 18 | 29 | 13 | 14 | 48 | 38 | 34 | 35 |
| ko00472 D-Arginine and D-ornithine metabolism | 4026 | 4096 | 3930 | 3926 | 2283 | 2259 | 2158 | 2272 | 1573 | 1428 | 1390 | 1514 |
| K17898 D-ornithine 4,5-aminomutase subunit beta [EC:5.4.3.5] | 1 | 0 | 0 | 0 | 0 | 0 | 2 | 3 | 1 | 4 | 5 | 2 |
| K17899 D-ornithine 4,5-aminomutase subunit alpha [EC:5.4.3.5] | 0 | 3 | 0 | 0 | 0 | 0 | 2 | 0 | 0 | 0 | 0 | 3 |
| K19744 L-arginine dehydrogenase [EC:1.4.1.25] | 1557 | 1535 | 1412 | 1442 | 918 | 860 | 777 | 839 | 478 | 449 | 421 | 452 |
| K19746 D-arginine dehydrogenase [EC:1.4.99.6] | 1785 | 1847 | 1822 | 1796 | 996 | 968 | 983 | 1063 | 689 | 529 | 547 | 592 |
| K21898 ornithine racemase [EC:5.1.1.12] | 19 | 45 | 41 | 31 | 40 | 57 | 29 | 37 | 27 | 25 | 44 | 52 |
| ko00473 D-Alanine metabolism | 11584 | 11376 | 11284 | 11249 | 12835 | 12757 | 12829 | 12622 | 12103 | 12115 | 11999 | 11890 |
| K01775 alanine racemase [EC:5.1.1.1] | 5618 | 5449 | 5415 | 5381 | 7500 | 7409 | 7367 | 7298 | 6646 | 6678 | 6636 | 6487 |
| K01921 D-alanine-D-alanine ligase [EC:6.3.2.4] | 5117 | 5141 | 5072 | 5038 | 4567 | 4518 | 4615 | 4569 | 5003 | 5021 | 4923 | 4991 |
| K03367 D-alanine--poly(phosphoribitol) ligase subunit 1 [EC:6.1.1.13] | 469 | 453 | 456 | 482 | 466 | 463 | 498 | 443 | 268 | 260 | 267 | 241 |
| K14188 D-alanine--poly(phosphoribitol) ligase subunit 2 [EC:6.1.1.13] | 77 | 70 | 70 | 82 | 59 | 81 | 82 | 79 | 53 | 33 | 57 | 44 |
| ko00480 Glutathione metabolism | 72117 | 71420 | 70079 | 71801 | 54876 | 55182 | 53923 | 55239 | 55062 | 55030 | 54029 | 55526 |
| K00383 glutathione reductase (NADPH) [EC:1.8.1.7] | 2927 | 2810 | 2641 | 2678 | 1547 | 1643 | 1704 | 1649 | 1776 | 1850 | 1640 | 1831 |
| K00799 glutathione S-transferase [EC:2.5.1.18] | 8066 | 7801 | 7767 | 8129 | 4589 | 4538 | 4351 | 4600 | 5020 | 4797 | 4675 | 4936 |
| K01255 leucyl aminopeptidase [EC:3.4.11.1] | 3148 | 3053 | 2887 | 3141 | 1693 | 1832 | 1609 | 1785 | 2297 | 2556 | 2342 | 2495 |
| K01256 aminopeptidase N [EC:3.4.11.2] | 5582 | 5558 | 5449 | 5600 | 3753 | 3630 | 3708 | 3565 | 3451 | 3498 | 3494 | 3525 |
| K01270 dipeptidase D [EC:3.4.13.-] | 3016 | 3181 | 3164 | 3080 | 6994 | 7072 | 7013 | 6803 | 6070 | 6048 | 5922 | 6206 |
| K01460 glutathionylspermidine amidase/synthetase [EC:3.5.1.78 6.3.1.8] | 664 | 693 | 768 | 780 | 250 | 183 | 231 | 200 | 1538 | 1638 | 1592 | 1483 |
| K01469 5-oxoprolinase (ATP-hydrolysing) [EC:3.5.2.9] | 2 | 2 | 2 | 3 | 0 | 5 | 3 | 0 | 4 | 10 | 4 | 8 |
| K06048 glutamate---cysteine ligase / carboxylate-amine ligase [EC:6.3.2.2 6.3.-.-] | 2188 | 2244 | 2138 | 2295 | 1250 | 1269 | 1182 | 1253 | 1345 | 1489 | 1400 | 1502 |
| K07160 5-oxoprolinase (ATP-hydrolysing) subunit A [EC:3.5.2.9] | 3965 | 4053 | 3697 | 3950 | 2222 | 2221 | 2098 | 2406 | 1956 | 1786 | 1667 | 1790 |
| K07232 glutathione-specific gamma-glutamylcyclotransferase [EC:4.3.2.7] | 366 | 390 | 365 | 427 | 289 | 320 | 260 | 252 | 503 | 529 | 532 | 539 |
| K07751 PepB aminopeptidase [EC:3.4.11.23] | 867 | 819 | 775 | 760 | 482 | 577 | 542 | 548 | 1031 | 1114 | 1060 | 1178 |
| K15428 Cys-Gly metallodipeptidase DUG1 [EC:3.4.13.-] | 2 | 2 | 2 | 0 | 0 | 0 | 0 | 0 | 0 | 0 | 0 | 0 |
| K23123 5-oxoprolinase (ATP-hydrolysing) subunit B [EC:3.5.2.9] | 355 | 358 | 351 | 363 | 278 | 288 | 294 | 269 | 408 | 444 | 461 | 485 |
| K23124 5-oxoprolinase (ATP-hydrolysing) subunit C [EC:3.5.2.9] | 489 | 562 | 520 | 477 | 385 | 405 | 391 | 362 | 702 | 711 | 731 | 670 |
| Glycan biosynthesis and metabolism | 241570 | 246088 | 250980 | 246569 | 364846 | 367201 | 367403 | 360665 | 333664 | 329566 | 333903 | 329667 |
| ko00510 N-Glycan biosynthesis | 791 | 837 | 897 | 859 | 1733 | 1750 | 1884 | 1744 | 1186 | 1137 | 1310 | 1226 |
| K00721 dolichol-phosphate mannosyltransferase [EC:2.4.1.83] | 763 | 807 | 855 | 829 | 1728 | 1742 | 1879 | 1741 | 1150 | 1063 | 1255 | 1170 |
| K00737 beta-1,4-mannosyl-glycoprotein beta-1,4-N-acetylglucosaminyltransferase [EC:2.4.1.144] | 8 | 9 | 13 | 9 | 0 | 4 | 1 | 1 | 13 | 15 | 10 | 12 |
| K01001 UDP-N-acetylglucosamine--dolichyl-phosphate N-acetylglucosaminephosphotransferase [EC:2.7.8.15] | 0 | 0 | 2 | 0 | 2 | 0 | 0 | 1 | 4 | 11 | 5 | 7 |
| K07151 dolichyl-diphosphooligosaccharide---protein glycosyltransferase [EC:2.4.99.18] | 20 | 21 | 27 | 21 | 3 | 4 | 4 | 1 | 19 | 48 | 40 | 37 |
| ko00513 Various types of N-glycan biosynthesis | 12253 | 13647 | 14863 | 13737 | 42752 | 42817 | 43878 | 42788 | 31402 | 30085 | 31721 | 30060 |
| ko00515 Mannose type O-glycan biosynthesis | 53 | 56 | 40 | 57 | 46 | 51 | 59 | 46 | 76 | 58 | 79 | 113 |
| K00728 dolichyl-phosphate-mannose-protein mannosyltransferase [EC:2.4.1.109] | 53 | 56 | 40 | 57 | 46 | 51 | 59 | 46 | 76 | 58 | 79 | 112 |
| ko00514 Other types of O-glycan biosynthesis | 53 | 56 | 40 | 57 | 46 | 51 | 59 | 46 | 76 | 58 | 79 | 112 |
| ko00531 Glycosaminoglycan degradation | 21940 | 23170 | 25148 | 23636 | 62104 | 61784 | 63099 | 61476 | 47710 | 45944 | 48090 | 46046 |
| K01135 arylsulfatase B [EC:3.1.6.12] | 0 | 0 | 0 | 1 | 0 | 0 | 0 | 0 | 2 | 0 | 1 | 1 |
| K01137 N-acetylglucosamine-6-sulfatase [EC:3.1.6.14] | 2 | 4 | 2 | 0 | 2 | 1 | 0 | 0 | 0 | 0 | 0 | 0 |
| K01197 hyaluronoglucosaminidase [EC:3.2.1.35] | 255 | 265 | 275 | 308 | 396 | 310 | 390 | 345 | 22 | 50 | 29 | 63 |
| K01205 alpha-N-acetylglucosaminidase [EC:3.2.1.50] | 2600 | 2712 | 3198 | 2864 | 9707 | 9394 | 9500 | 9246 | 6704 | 6511 | 6790 | 6454 |
| K01217 L-iduronidase [EC:3.2.1.76] | 0 | 1 | 5 | 1 | 0 | 0 | 0 | 0 | 1 | 0 | 0 | 1 |
| K01565 N-sulfoglucosamine sulfohydrolase [EC:3.10.1.1] | 8 | 4 | 6 | 4 | 7 | 5 | 4 | 2 | 2 | 4 | 3 | 1 |
| ko00563 Glycosylphosphatidylinositol (GPI)-anchor biosynthesis | 0 | 1 | 0 | 0 | 0 | 0 | 0 | 0 | 0 | 0 | 0 | 0 |
| K03434 N-acetylglucosaminylphosphatidylinositol deacetylase [EC:3.5.1.89] | 0 | 1 | 0 | 0 | 0 | 0 | 0 | 0 | 0 | 0 | 0 | 0 |
| ko00601 Glycosphingolipid biosynthesis - lacto and neolacto series | 3 | 1 | 0 | 1 | 0 | 0 | 0 | 0 | 0 | 0 | 1 | 0 |
| K00709 histo-blood group ABO system transferase [EC:2.4.1.40 2.4.1.37] | 3 | 1 | 0 | 1 | 0 | 0 | 0 | 0 | 0 | 0 | 1 | 0 |
| ko00603 Glycosphingolipid biosynthesis - globo and isoglobo series | 22516 | 24234 | 25747 | 24358 | 57871 | 58084 | 59275 | 57657 | 50553 | 49484 | 50719 | 49190 |
| K00722 globoside alpha-N-acetylgalactosaminyltransferase [EC:2.4.1.88] | 1 | 0 | 0 | 0 | 0 | 0 | 0 | 0 | 0 | 0 | 0 | 0 |
| K01204 alpha-N-acetylgalactosaminidase [EC:3.2.1.49] | 1 | 0 | 0 | 0 | 0 | 0 | 0 | 0 | 0 | 0 | 0 | 0 |
| ko00604 Glycosphingolipid biosynthesis - ganglio series | 12237 | 13631 | 14837 | 13720 | 42758 | 42818 | 43881 | 42796 | 31387 | 30047 | 31685 | 30028 |
| ko00540 Lipopolysaccharide biosynthesis | 66881 | 66967 | 66581 | 67463 | 71571 | 71940 | 68813 | 69446 | 65795 | 65943 | 65880 | 66690 |
| K00677 UDP-N-acetylglucosamine acyltransferase [EC:2.3.1.129] | 2770 | 2740 | 2830 | 2703 | 4653 | 4531 | 3883 | 3869 | 3556 | 3626 | 3787 | 3728 |
| K00713 UDP-glucose:(glucosyl)LPS alpha-1,2-glucosyltransferase [EC:2.4.1.-] | 0 | 2 | 0 | 0 | 0 | 0 | 0 | 0 | 3 | 1 | 0 | 2 |
| K00748 lipid-A-disaccharide synthase [EC:2.4.1.182] | 3125 | 3234 | 3284 | 3270 | 4118 | 4108 | 4159 | 4172 | 3297 | 3140 | 3304 | 3184 |
| K00912 tetraacyldisaccharide 4'-kinase [EC:2.7.1.130] | 2414 | 2506 | 2585 | 2674 | 3674 | 3722 | 3750 | 3709 | 3173 | 3032 | 3022 | 3115 |
| K00979 3-deoxy-manno-octulosonate cytidylyltransferase (CMP-KDO synthetase) [EC:2.7.7.38] | 1974 | 1930 | 1876 | 2088 | 2453 | 2364 | 2353 | 2372 | 2043 | 2041 | 2070 | 2009 |
| K01627 2-dehydro-3-deoxyphosphooctonate aldolase (KDO 8-P synthase) [EC:2.5.1.55] | 2071 | 2202 | 2086 | 2038 | 2779 | 2741 | 2641 | 2791 | 2147 | 2132 | 2182 | 2170 |
| K02517 Kdo2-lipid IVA lauroyltransferase/acyltransferase [EC:2.3.1.241 2.3.1.-] | 3808 | 3961 | 4091 | 4021 | 3871 | 4034 | 3783 | 3886 | 3632 | 3574 | 3706 | 3707 |
| K02527 3-deoxy-D-manno-octulosonic-acid transferase [EC:2.4.99.12 2.4.99.13 2.4.99.14 2.4.99.15] | 3225 | 3236 | 3198 | 3155 | 4292 | 4216 | 4254 | 4338 | 3576 | 3638 | 3666 | 3697 |
| K02535 UDP-3-O-[3-hydroxymyristoyl] N-acetylglucosamine deacetylase [EC:3.5.1.108] | 1773 | 1928 | 1896 | 1957 | 1209 | 1309 | 1214 | 1203 | 1270 | 1455 | 1291 | 1318 |
| K02536 UDP-3-O-[3-hydroxymyristoyl] glucosamine N-acyltransferase [EC:2.3.1.191] | 3561 | 3477 | 3705 | 3580 | 5749 | 5688 | 4056 | 3973 | 4542 | 4399 | 4505 | 4457 |
| K02560 lauroyl-Kdo2-lipid IVA myristoyltransferase [EC:2.3.1.243] | 556 | 540 | 481 | 527 | 416 | 329 | 421 | 427 | 728 | 750 | 646 | 770 |

| | | | | | | | | | | | | |
|---|---|---|---|---|---|---|---|---|---|---|---|---|
| K02840 UDP-D-galactose:(glucosyl)LPS alpha-1,6-D-galactosyltransferase [EC:2.4.1.-] | 0 | 1 | 2 | 0 | 2 | 1 | 3 | 5 | 1 | 2 | 8 | 2 |
| K02841 heptosyltransferase I [EC:2.4.-.-] | 1136 | 1156 | 1031 | 1117 | 787 | 813 | 819 | 709 | 1056 | 1108 | 1020 | 1084 |
| K02843 heptosyltransferase II [EC:2.4.-.-] | 1417 | 1459 | 1494 | 1453 | 2864 | 2763 | 2854 | 2843 | 2798 | 2936 | 3055 | 2937 |
| K02844 UDP-glucose:(heptosyl)LPS alpha-1,3-glucosyltransferase [EC:2.4.1.-] | 2594 | 2631 | 2620 | 2715 | 1550 | 1600 | 1572 | 1548 | 1569 | 1694 | 1541 | 1768 |
| K02847 O-antigen ligase [EC:2.4.1.-] | 3 | 3 | 6 | 3 | 11 | 20 | 22 | 13 | 93 | 79 | 95 | 76 |
| K02848 heptose I phosphotransferase [EC:2.7.1.-] | 1272 | 1369 | 1271 | 1324 | 796 | 712 | 722 | 712 | 483 | 457 | 433 | 465 |
| K02849 heptosyltransferase III [EC:2.4.-.-] | 670 | 660 | 670 | 723 | 514 | 528 | 483 | 512 | 935 | 995 | 911 | 977 |
| K02850 heptose II phosphotransferase [EC:2.7.1.-] | 3 | 5 | 2 | 0 | 9 | 8 | 15 | 7 | 30 | 30 | 29 | 46 |
| K03269 UDP-2,3-diacylglucosamine hydrolase [EC:3.6.1.54] | 1327 | 1451 | 1477 | 1400 | 2227 | 2327 | 2303 | 2302 | 2065 | 1922 | 2066 | 1982 |
| K03270 3-deoxy-D-manno-octulosonate 8-phosphate phosphatase (KDO 8-P phosphatase) [EC:3.1.3.45] | 1622 | 1568 | 1564 | 1644 | 1846 | 2004 | 1798 | 1865 | 1543 | 1647 | 1530 | 1508 |
| K03271 D-sedoheptulose 7-phosphate isomerase [EC:5.3.1.28] | 1990 | 2035 | 2006 | 2133 | 1773 | 1831 | 1810 | 1818 | 1959 | 1942 | 1886 | 1913 |
| K03272 D-beta-D-heptose 7-phosphate kinase / D-beta-D-heptose 1-phosphate adenosyltransferase [EC:2.7.1.167 2.7.7.70] | 3566 | 3596 | 3293 | 3507 | 2168 | 2069 | 1971 | 2076 | 2075 | 2174 | 2221 | 2334 |
| K03273 D-glycero-D-manno-heptose 1,7-bisphosphate phosphatase [EC:3.1.3.82 3.1.3.83] | 1994 | 1962 | 2018 | 1991 | 2343 | 2506 | 2489 | 2469 | 2153 | 2214 | 2190 | 2103 |
| K03274 ADP-L-glycero-D-manno-heptose 6-epimerase [EC:5.1.3.20] | 2039 | 1895 | 1910 | 1962 | 1281 | 1294 | 1168 | 1222 | 1447 | 1423 | 1403 | 1386 |
| K03275 UDP-glucose:(glucosyl)LPS alpha-1,3-glucosyltransferase [EC:2.4.1.-] | 4 | 4 | 5 | 9 | 16 | 9 | 18 | 8 | 51 | 38 | 56 | 51 |
| K03276 UDP-glucose/galactose:(glucosyl)LPS alpha-1,2-glucosyl/galactosyltransferase [EC:2.4.1.-] | 10 | 9 | 13 | 14 | 50 | 38 | 36 | 23 | 98 | 82 | 59 | 71 |
| K03277 heptosyltransferase IV [EC:2.4.-.-] | 0 | 1 | 0 | 0 | 0 | 2 | 0 | 0 | 0 | 0 | 0 | 0 |
| K03760 lipid A ethanolaminephosphotransferase [EC:2.7.8.43] | 3898 | 3769 | 3561 | 3734 | 2223 | 2275 | 2116 | 2246 | 2325 | 2238 | 2254 | 2280 |
| K06041 arabinose-5-phosphate isomerase [EC:5.3.1.13] | 1936 | 1913 | 1875 | 1927 | 2737 | 2829 | 2835 | 2656 | 2108 | 2325 | 2257 | 2307 |
| K07031 D-glycero-alpha-D-manno-heptose-7-phosphate kinase [EC:2.7.1.168] | 2056 | 2087 | 2180 | 2135 | 2988 | 2871 | 3189 | 3087 | 3007 | 3028 | 3033 | 3081 |
| K07264 4-amino-4-deoxy-L-arabinose transferase [EC:2.4.2.43] | 2759 | 2548 | 2371 | 2608 | 1493 | 1523 | 1410 | 1659 | 1853 | 1839 | 1702 | 1928 |
| K09778 Kdo2-lipid IVA 3' secondary acyltransferase [EC:2.3.1.-] | 3 | 3 | 5 | 7 | 0 | 0 | 3 | 2 | 1 | 0 | 2 | 1 |
| K12973 lipid IVA palmitoyltransferase [EC:2.3.1.251] | 327 | 330 | 358 | 339 | 287 | 318 | 314 | 313 | 478 | 599 | 525 | 532 |
| K12974 KDO2-lipid IV(A) palmitoleoyltransferase [EC:2.3.1.242] | 500 | 504 | 485 | 554 | 334 | 414 | 367 | 350 | 707 | 701 | 678 | 821 |
| K12975 KDO II ethanolaminephosphotransferase [EC:2.7.8.42] | 1181 | 1121 | 1104 | 1121 | 806 | 806 | 771 | 791 | 1578 | 1572 | 1557 | 1592 |
| K12976 lipid A 3-O-deacylase [EC:3.1.1.-] | 426 | 467 | 442 | 431 | 257 | 274 | 315 | 373 | 138 | 136 | 141 | 142 |
| K12979 beta-hydroxylase [EC:1.14.11.-] | 3162 | 3088 | 3045 | 3050 | 1659 | 1635 | 1596 | 1728 | 1543 | 1377 | 1480 | 1554 |
| K12981 KDO transferase III [EC:2.4.99.-] | 86 | 88 | 82 | 82 | 123 | 114 | 172 | 128 | 0 | 1 | 0 | 0 |
| K12983 UDP-glucose:(glucosyl)LPS beta-1,3-glucosyltransferase [EC:2.4.1.-] | 2 | 0 | 0 | 2 | 15 | 14 | 11 | 13 | 43 | 44 | 49 | 44 |
| K12985 (galactosyl)LPS 1,2-glucosyltransferase [EC:2.4.1.-] | 1 | 3 | 1 | 6 | 8 | 15 | 5 | 4 | 35 | 49 | 55 | 50 |
| K12986 1,5-rhamnosyltransferase [EC:2.4.1.-] | 0 | 0 | 0 | 0 | 3 | 0 | 2 | 0 | 0 | 0 | 0 | 1 |
| K15669 D-glycero-alpha-D-manno-heptose 1-phosphate guanylyltransferase [EC:2.7.7.71] | 672 | 663 | 749 | 639 | 1895 | 1754 | 1964 | 1889 | 1498 | 1401 | 1434 | 1430 |
| K19353 heptose-I-phosphate ethanolaminephosphotransferase [EC:2.7.8.-] | 3056 | 2983 | 3105 | 2989 | 1698 | 1800 | 1585 | 1716 | 1257 | 1230 | 1256 | 1260 |
| K19803 Kdo2-lipid A phosphotransferase [EC:2.7.4.29] | 829 | 826 | 764 | 838 | 499 | 599 | 443 | 486 | 723 | 765 | 630 | 745 |
| K21344 D-glycero-beta-D-manno-heptose-7-phosphate kinase [EC:2.7.1.167] | 56 | 63 | 62 | 47 | 88 | 84 | 86 | 93 | 78 | 87 | 77 | 81 |
| K21345 D-glycero-beta-D-manno-heptose 1-phosphate adenylyltransferase [EC:2.7.7.70] | 33 | 23 | 40 | 27 | 39 | 37 | 43 | 54 | 47 | 29 | 32 | 41 |
| K23067 beta-1,4-galactosyltransferase | 0 | 0 | 0 | 0 | 0 | 0 | 0 | 0 | 0 | 0 | 0 | 1 |
| ko00550 Peptidoglycan biosynthesis | 86848 | 87008 | 87124 | 86745 | 88584 | 90742 | 90751 | 89004 | 93249 | 94341 | 93030 | 93778 |
| K00687 penicillin-binding protein 2B | 110 | 135 | 106 | 126 | 39 | 39 | 32 | 21 | 116 | 185 | 110 | 144 |
| K00887 undecaprenol kinase [EC:2.7.1.66] | 25 | 23 | 29 | 15 | 3 | 5 | 8 | 8 | 31 | 35 | 39 | 29 |
| K01000 phospho-N-acetylmuramoyl-pentapeptide-transferase [EC:2.7.8.13] | 2617 | 2792 | 2880 | 2888 | 3912 | 4145 | 4297 | 3919 | 4056 | 4316 | 4115 | 3923 |
| K02563 UDP-N-acetylglucosamine--N-acetylmuramyl-(pentapeptide) pyrophosphoryl-undecaprenol N-acetylglucosamine transferase [EC:2.4.1.227] | 3791 | 3850 | 3717 | 3791 | 4322 | 4365 | 4445 | 4345 | 4045 | 3783 | 3953 | 3955 |
| K03587 cell division protein FtsI (penicillin-binding protein 3) [EC:3.4.16.4] | 7696 | 7695 | 7686 | 7682 | 8334 | 8378 | 8283 | 8272 | 8046 | 7716 | 7911 | 7807 |
| K03693 penicillin-binding protein 1B | 125 | 118 | 93 | 124 | 17 | 45 | 26 | 40 | 115 | 158 | 117 | 118 |
| K03814 monofunctional glycosyltransferase [EC:2.4.1.129] | 2004 | 2089 | 2011 | 2010 | 2209 | 2110 | 2163 | 2224 | 1960 | 1845 | 1838 | 1928 |
| K05362 UDP-N-acetylmuramoyl-L-alanyl-D-glutamate-L-lysine ligase [EC:6.3.2.7] | 80 | 98 | 104 | 99 | 19 | 17 | 23 | 22 | 102 | 136 | 82 | 94 |
| K05363 serine/alanine adding enzyme [EC:2.3.2.10] | 48 | 65 | 51 | 72 | 4 | 7 | 7 | 9 | 76 | 72 | 50 | 60 |
| K05364 penicillin-binding protein A | 396 | 381 | 382 | 369 | 78 | 77 | 117 | 86 | 310 | 408 | 358 | 395 |
| K05365 penicillin-binding protein 1B [EC:2.4.1.129 3.4.16.4] | 5137 | 5053 | 5024 | 5118 | 3198 | 3236 | 2919 | 3065 | 3079 | 3245 | 3213 | 3295 |
| K05366 penicillin-binding protein 1A [EC:2.4.1.129 3.4.16.4] | 8090 | 8285 | 8511 | 8221 | 8144 | 8350 | 8238 | 8210 | 8035 | 8209 | 8080 | 8270 |
| K05367 penicillin-binding protein 1C [EC:2.4.1.129] | 2517 | 2542 | 2480 | 2586 | 4923 | 5069 | 5341 | 4910 | 4851 | 4925 | 4855 | 4779 |
| K05515 penicillin-binding protein 2 [EC:3.4.16.4] | 6289 | 6351 | 6261 | 6331 | 6424 | 6962 | 6650 | 6492 | 7691 | 7628 | 7682 | 7322 |
| K06153 undecaprenyl-diphosphatase [EC:3.6.1.27] | 2935 | 3057 | 3042 | 2946 | 3187 | 3061 | 3096 | 3008 | 3367 | 3219 | 3317 | 3238 |
| K07009 lipid II isoglutaminyl synthase (glutamine-hydrolysing) [EC:6.3.5.13] | 92 | 118 | 91 | 92 | 87 | 76 | 73 | 90 | 145 | 161 | 170 | 154 |
| K07258 serine-type D-Ala-D-Ala carboxypeptidase (penicillin-binding protein 5/6) [EC:3.4.16.4] | 6654 | 6142 | 6430 | 6460 | 2947 | 3104 | 2973 | 2969 | 5581 | 5957 | 5625 | 5964 |
| K07259 serine-type D-Ala-D-Ala carboxypeptidase/endopeptidase (penicillin-binding protein 4) [EC:3.4.16.4 3.4.21.-] | 3295 | 3261 | 3325 | 3077 | 4556 | 4888 | 5044 | 4764 | 4140 | 3979 | 4147 | 4049 |
| K07260 zinc D-Ala-D-Ala carboxypeptidase [EC:3.4.17.14] | 295 | 265 | 307 | 294 | 80 | 88 | 86 | 85 | 312 | 364 | 325 | 337 |
| K08384 stage V sporulation protein D (sporulation-specific penicillin-binding protein) | 1553 | 1440 | 1465 | 1499 | 413 | 438 | 483 | 406 | 1472 | 1676 | 1562 | 1640 |
| K08724 penicillin-binding protein 2B | 3 | 3 | 3 | 4 | 7 | 15 | 19 | 10 | 9 | 5 | 5 | 11 |
| K11693 peptidoglycan pentaglycine glycine transferase (the first glycine) [EC:2.3.2.16] | 0 | 2 | 0 | 0 | 4 | 0 | 0 | 0 | 0 | 0 | 2 | 0 |
| K11694 peptidoglycan pentaglycine glycine transferase (the second and third glycine) [EC:2.3.2.17] | 1 | 0 | 0 | 0 | 0 | 2 | 0 | 3 | 1 | 2 | 0 | 3 |
| K12554 alanine adding enzyme [EC:2.3.2.-] | 76 | 102 | 90 | 50 | 16 | 15 | 18 | 11 | 81 | 87 | 87 | 99 |
| K12555 penicillin-binding protein 2A [EC:2.4.1.129 3.4.16.4] | 263 | 337 | 304 | 311 | 64 | 60 | 72 | 60 | 250 | 299 | 216 | 232 |
| K12556 penicillin-binding protein 2X | 107 | 115 | 107 | 117 | 36 | 39 | 35 | 22 | 131 | 140 | 137 | 128 |
| K18149 penicillin-binding protein | 106 | 97 | 88 | 98 | 15 | 8 | 17 | 11 | 85 | 94 | 79 | 119 |
| K18770 penicillin-binding protein 4 [EC:2.4.1.129 3.4.16.4] | 2 | 0 | 0 | 0 | 0 | 0 | 0 | 5 | 0 | 0 | 0 | 0 |
| K18866 zinc D-Ala-D-Ala dipeptidase/carboxypeptidase [EC:3.4.13.22 3.4.17.14] | 7 | 5 | 12 | 5 | 8 | 0 | 4 | 4 | 4 | 8 | 9 | 5 |
| K19302 undecaprenyl-diphosphatase [EC:3.6.1.27] | 3331 | 3310 | 3180 | 3247 | 2750 | 2751 | 2951 | 3076 | 2635 | 2592 | 2702 | 2670 |
| K21464 penicillin-binding protein 2D [EC:2.4.1.129 3.4.16.4] | 6 | 5 | 5 | 4 | 5 | 7 | 4 | 3 | 14 | 9 | 8 | 11 |
| K21465 penicillin-binding protein A | 0 | 0 | 2 | 0 | 3 | 4 | 3 | 2 | 3 | 7 | 0 | 0 |
| K21469 serine-type D-Ala-D-Ala carboxypeptidase [EC:3.4.16.4] | 3 | 0 | 0 | 0 | 13 | 10 | 10 | 9 | 10 | 9 | 7 | 15 |
| K23393 lipid II isoglutaminyl synthase (glutamine-hydrolysing) [EC:6.3.5.13] | 209 | 204 | 209 | 223 | 156 | 153 | 109 | 107 | 288 | 307 | 278 | 331 |
| ko00511 Other glycan degradation | 64353 | 68557 | 72676 | 68431 | 166618 | 166689 | 169506 | 165064 | 136142 | 130969 | 136413 | 130815 |
| K01191 alpha-mannosidase [EC:3.2.1.24] | 4577 | 4779 | 4906 | 4864 | 5245 | 5153 | 5480 | 5166 | 8407 | 8418 | 8678 | 8612 |
| K01192 beta-mannosidase [EC:3.2.1.25] | 5160 | 5322 | 5520 | 5290 | 9105 | 8987 | 9196 | 8860 | 11382 | 11333 | 11323 | 11053 |
| K01206 alpha-L-fucosidase [EC:3.2.1.51] | 5778 | 5816 | 6254 | 5966 | 12704 | 12642 | 12804 | 12621 | 11046 | 10391 | 10911 | 10345 |
| K01227 mannosyl-glycoprotein endo-beta-N-acetylglucosaminidase [EC:3.2.1.96] | 868 | 851 | 866 | 879 | 159 | 209 | 140 | 124 | 469 | 569 | 542 | 553 |
| K01444 N4-(beta-N-acetylglucosaminyl)-L-asparaginase [EC:3.5.1.26] | 2 | 0 | 0 | 0 | 0 | 0 | 0 | 0 | 0 | 0 | 2 | 1 |
| K15923 alpha-L-fucosidase 2 [EC:3.2.1.51] | 9302 | 10187 | 10877 | 10000 | 31704 | 31856 | 32206 | 31458 | 23944 | 22760 | 24037 | 22497 |
| ko00571 Lipoarabinomannan (LAM) biosynthesis | 1751 | 1744 | 1665 | 1680 | 1219 | 1166 | 1119 | 1207 | 693 | 621 | 662 | 751 |
| K08256 phosphatidyl-myo-inositol alpha-mannosyltransferase [EC:2.4.1.345] | 60 | 79 | 76 | 85 | 270 | 285 | 292 | 268 | 186 | 175 | 178 | 183 |
| K14335 alpha-1,6-mannosyltransferase [EC:2.4.1.-] | 1691 | 1663 | 1589 | 1595 | 949 | 881 | 827 | 939 | 507 | 446 | 484 | 568 |
| K22311 phosphatidylinositol dimannoside acyltransferase [EC:2.3.1.265] | 0 | 2 | 0 | 0 | 0 | 0 | 0 | 0 | 0 | 0 | 0 | 0 |
| ko00572 Arabinogalactan biosynthesis - Mycobacterium | 904 | 770 | 775 | 775 | 607 | 626 | 652 | 604 | 1030 | 1153 | 1085 | 1109 |
| K02851 UDP-GlcNAc:undecaprenyl-phosphate/decaprenyl-phosphate GlcNAc-1-phosphate transferase [EC:2.7.8.33 2.7.8.35] | 899 | 768 | 770 | 768 | 601 | 622 | 649 | 597 | 1015 | 1125 | 1085 | 1095 |
| K16650 galactofuranosylgalactofuranosylrhamnosyl-N-acetylglucosaminyl-diphospho-decaprenol beta-1,5/1,6-galactofuranosyltransferase [EC:2.4.1.288] | 0 | 0 | 0 | 0 | 0 | 1 | 3 | 0 | 6 | 17 | 0 | 10 |
| K22902 D-galactosaminyltransferase | 5 | 2 | 5 | 7 | 6 | 3 | 0 | 7 | 9 | 11 | 0 | 4 |
| Metabolism of cofactors and vitamins | 535572 | 534387 | 523670 | 530729 | 500589 | 500853 | 497817 | 500763 | 493153 | 490778 | 487504 | 492888 |
| ko00730 Thiamine metabolism | 50870 | 51286 | 51216 | 51131 | 57812 | 58113 | 58381 | 57578 | 55567 | 56271 | 55802 | 55628 |
| K00788 thiamine-phosphate pyrophosphorylase [EC:2.5.1.3] | 2332 | 2451 | 2410 | 2347 | 3140 | 3356 | 3377 | 3192 | 3054 | 3112 | 3124 | 3008 |
| K00877 hydroxymethylpyrimidine/phosphomethylpyrimidine kinase / thiaminase [EC:2.7.1.49 2.7.4.7 3.5.99.2] | 1 | 0 | 0 | 0 | 1 | 0 | 0 | 0 | 0 | 0 | 1 | 1 |
| K00878 hydroxyethylthiazole kinase [EC:2.7.1.50] | 1075 | 1055 | 979 | 1087 | 494 | 461 | 464 | 500 | 1125 | 1242 | 1289 | 1205 |
| K00941 hydroxymethylpyrimidine/phosphomethylpyrimidine kinase [EC:2.7.1.49 2.7.4.7] | 2533 | 2627 | 2533 | 2549 | 2677 | 2559 | 2537 | 2576 | 2655 | 2768 | 2610 | 2679 |
| K00946 thiamine-monophosphate kinase [EC:2.7.4.16] | 2743 | 2729 | 2744 | 2634 | 3546 | 3540 | 3659 | 3592 | 3121 | 3005 | 2964 | 3160 |
| K00949 thiamine pyrophosphokinase [EC:2.7.6.2] | 689 | 780 | 810 | 751 | 1511 | 1565 | 1519 | 1410 | 1380 | 1386 | 1431 | 1414 |
| K01077 alkaline phosphatase [EC:3.1.3.1] | 4527 | 4672 | 4930 | 4814 | 7494 | 7562 | 7672 | 7634 | 6192 | 6091 | 6417 | 6081 |
| K01662 1-deoxy-D-xylulose-5-phosphate synthase [EC:2.2.1.7] | 7630 | 7602 | 7659 | 7587 | 10318 | 10121 | 10353 | 10111 | 8777 | 8802 | 8735 | 8488 |
| K03147 phosphomethylpyrimidine synthase [EC:4.1.99.17] | 5511 | 5615 | 5403 | 5549 | 5776 | 6134 | 5935 | 5893 | 5663 | 5637 | 5592 | 5612 |
| K03148 sulfur carrier protein ThiS adenylyltransferase [EC:2.7.7.73] | 463 | 506 | 491 | 504 | 338 | 399 | 391 | 360 | 563 | 684 | 685 | 716 |
| K03149 thiazole synthase [EC:2.8.1.10] | 2400 | 2321 | 2316 | 2286 | 2682 | 2684 | 2706 | 2685 | 2312 | 2394 | 2266 | 2344 |
| K03150 2-iminoacetate synthase [EC:4.1.99.19] | 2820 | 2952 | 3123 | 2994 | 5164 | 5307 | 5410 | 5255 | 4832 | 5121 | 5029 | 4892 |
| K03151 tRNA uracil 4-sulfurtransferase [EC:2.8.1.4] | 3924 | 3873 | 3721 | 3831 | 2120 | 2105 | 2009 | 2050 | 2627 | 2657 | 2514 | 2650 |
| K03153 glycine oxidase [EC:1.4.3.19] | 1655 | 1667 | 1702 | 1643 | 995 | 933 | 915 | 883 | 548 | 490 | 545 | 558 |
| K03707 thiaminase (transcriptional activator TenA) [EC:3.5.99.2] | 642 | 662 | 727 | 669 | 163 | 142 | 163 | 147 | 1120 | 1119 | 1144 | 1151 |
| K04487 cysteine desulfurase [EC:2.8.1.7] | 4959 | 4725 | 4754 | 4819 | 2670 | 2858 | 2652 | 2803 | 3115 | 3420 | 3078 | 3237 |
| K06949 ribosome biogenesis GTPase / thiamine phosphate phosphatase [EC:3.6.1.- 3.1.3.100] | 4179 | 4204 | 4153 | 4278 | 6058 | 5742 | 5991 | 5834 | 5565 | 5440 | 5456 | 5486 |
| K07251 thiamine kinase [EC:2.7.1.89] | 502 | 449 | 466 | 472 | 328 | 307 | 296 | 323 | 615 | 599 | 669 | 689 |
| K10810 thiazole tautomerase (transcriptional regulator TenI) [EC:5.3.99.10] | 0 | 0 | 0 | 0 | 0 | 0 | 0 | 1 | 0 | 0 | 0 | 0 |
| K14153 hydroxymethylpyrimidine kinase / phosphomethylpyrimidine kinase / thiamine-phosphate diphosphorylase [EC:2.7.1.49 2.7.4.7 2.5.1.3] | 2 | 1 | 3 | 3 | 8 | 5 | 7 | 1 | 3 | 2 | 7 | 4 |
| K14154 thiamine-phosphate diphosphorylase / hydroxyethylthiazole kinase [EC:2.5.1.3 2.7.1.50] | 1 | 0 | 0 | 0 | 0 | 0 | 0 | 0 | 0 | 0 | 2 | 0 |
| K14394 low molecular weight phosphotyrosine protein phosphatase [EC:3.1.3.2 3.1.3.48] | 0 | 1 | 0 | 0 | 0 | 1 | 0 | 1 | 0 | 0 | 0 | 1 |
| K18278 pyrimidine precursor biosynthesis enzyme | 0 | 0 | 0 | 0 | 0 | 0 | 0 | 0 | 2 | 0 | 0 | 0 |
| K21219 hydroxymethylpyrimidine kinase / phosphomethylpyrimidine kinase / thiamine-phosphate diphosphorylase [EC:2.7.1.49 2.7.4.7 2.5.1.3] | 0 | 0 | 0 | 0 | 0 | 0 | 0 | 0 | 0 | 0 | 0 | 1 |
| K22699 sulfide-dependent adenosine diphosphate thiazole synthase [EC:2.4.2.59] | 3 | 2 | 5 | 5 | 0 | 0 | 1 | 0 | 0 | 0 | 0 | 0 |
| ko00740 Riboflavin metabolism | 21818 | 22168 | 20993 | 21472 | 21783 | 21538 | 21803 | 21674 | 20935 | 21071 | 21151 | 21578 |
| K00793 riboflavin synthase [EC:2.5.1.9] | 1809 | 1813 | 1763 | 1828 | 2157 | 2248 | 2266 | 2293 | 1585 | 1633 | 1582 | 1642 |
| K00794 6,7-dimethyl-8-ribityllumazine synthase [EC:2.5.1.78] | 1539 | 1505 | 1390 | 1321 | 1856 | 1851 | 1881 | 1822 | 1584 | 1441 | 1616 | 1517 |
| K01497 GTP cyclohydrolase II [EC:3.5.4.25] | 1432 | 1332 | 1287 | 1330 | 840 | 759 | 719 | 786 | 777 | 833 | 810 | 741 |
| K02858 3,4-dihydroxy 2-butanone 4-phosphate synthase [EC:4.1.99.12] | 429 | 403 | 392 | 446 | 288 | 326 | 331 | 302 | 592 | 610 | 543 | 659 |
| K03788 acid phosphatase (class B) [EC:3.1.3.2] | 467 | 433 | 415 | 481 | 331 | 300 | 294 | 288 | 574 | 538 | 611 | 627 |
| K04719 5,6-dimethylbenzimidazole synthase [EC:1.13.11.79] | 718 | 703 | 572 | 618 | 321 | 292 | 291 | 378 | 202 | 206 | 155 | 213 |

| | | | | | | | | | | | | |
|---|---|---|---|---|---|---|---|---|---|---|---|---|
| K05368 aquacobalamin reductase / NAD(P)H-flavin reductase [EC:1.16.1.3 1.5.1.41] | 494 | 491 | 431 | 458 | 324 | 363 | 321 | 358 | 616 | 705 | 678 | 647 |
| K07732 riboflavin kinase, archaea type [EC:2.7.1.161] | 0 | 0 | 1 | 0 | 0 | 0 | 0 | 0 | 0 | 0 | 0 | 4 |
| K08096 GTP cyclohydrolase IIa [EC:3.5.4.29] | 0 | 0 | 0 | 0 | 0 | 0 | 0 | 0 | 2 | 1 | 0 | 9 |
| K09474 acid phosphatase (class A) [EC:3.1.3.2] | 324 | 343 | 281 | 298 | 221 | 196 | 215 | 167 | 500 | 502 | 523 | 577 |
| K11752 diaminohydroxyphosphoribosylaminopyrimidine deaminase / 5-amino-6-(5-phosphoribosylamino)uracil reductase [EC:3.5.4.26 1.1.1.193] | 3137 | 3256 | 3132 | 3150 | 3768 | 3781 | 3814 | 3841 | 3253 | 3172 | 3285 | 3229 |
| K11753 riboflavin kinase / FMN adenylyltransferase [EC:2.7.1.26 2.7.7.2] | 3398 | 3497 | 3462 | 3572 | 3861 | 3778 | 3835 | 3685 | 3328 | 3489 | 3526 | 3620 |
| K14379 tartrate-resistant acid phosphatase type 5 [EC:3.1.3.2] | 0 | 0 | 2 | 0 | 0 | 0 | 0 | 0 | 0 | 1 | 0 | 0 |
| K14652 3,4-dihydroxy 2-butanone 4-phosphate synthase / GTP cyclohydrolase II [EC:4.1.99.12 3.5.4.25] | 2644 | 2748 | 2624 | 2532 | 4167 | 3995 | 4181 | 4032 | 2922 | 2656 | 2914 | 2777 |
| K14653 2-amino-5-formylamino-6-ribosylaminopyrimidin-4(3H)-one 5'-monophosphate deformylase [EC:3.5.1.102] | 0 | 1 | 0 | 0 | 3 | 0 | 2 | 0 | 2 | 7 | 0 | 2 |
| K14654 2,5-diamino-6-(ribosylamino)-4(3H)-pyrimidinone 5'-phosphate reductase [EC:1.1.1.302] | 98 | 102 | 120 | 106 | 19 | 23 | 9 | 12 | 56 | 68 | 76 | 77 |
| K14655 tRNA pseudouridine32 synthase / 2,5-diamino-6-(5-phospho-D-ribitylamino)-pyrimidin-4(3H)-one deaminase [EC:5.4.99.28] | 0 | 1 | 0 | 0 | 0 | 0 | 0 | 0 | 0 | 0 | 0 | 0 |
| K14656 FAD synthetase [EC:2.7.7.2] | 0 | 2 | 0 | 1 | 0 | 0 | 0 | 2 | 3 | 5 | 2 | 3 |
| K19285 FMN reductase (NADPH) [EC:1.5.1.38] | 284 | 258 | 313 | 245 | 57 | 61 | 59 | 33 | 152 | 233 | 198 | 174 |
| K19286 FMN reductase [NAD(P)H] [EC:1.5.1.39] | 0 | 2 | 9 | 4 | 0 | 4 | 3 | 1 | 4 | 4 | 5 | 5 |
| K20861 FMN hydrolase / 5-amino-6-(5-phospho-D-ribitylamino)uracil phosphatase [EC:3.1.3.102 3.1.3.104] | 253 | 283 | 235 | 273 | 74 | 89 | 90 | 74 | 574 | 606 | 520 | 603 |
| K20862 FMN hydrolase / 5-amino-6-(5-phospho-D-ribitylamino)uracil phosphatase [EC:3.1.3.102 3.1.3.104] | 1697 | 1768 | 1620 | 1671 | 963 | 1019 | 936 | 1056 | 994 | 1014 | 957 | 1087 |
| K20884 riboflavin kinase / FMN hydrolase [EC:2.7.1.26 3.1.3.102] | 0 | 1 | 0 | 1 | 2 | 1 | 1 | 2 | 3 | 3 | 0 | 3 |
| K21063 5-amino-6-(5-phospho-D-ribitylamino)uracil phosphatase [EC:3.1.3.104] | 388 | 408 | 421 | 395 | 1222 | 1162 | 1305 | 1153 | 830 | 797 | 854 | 840 |
| K21064 5-amino-6-(5-phospho-D-ribitylamino)uracil phosphatase [EC:3.1.3.104] | 46 | 48 | 58 | 35 | 20 | 6 | 19 | 14 | 49 | 71 | 64 | 42 |
| K22393 flavin reductase (NADH) subunit 1 [EC:1.5.1.36] | 0 | 2 | 2 | 1 | 2 | 0 | 0 | 2 | 2 | 0 | 2 | 0 |
| K22394 flavin reductase (NADH) subunit 2 [EC:1.5.1.36] | 0 | 0 | 0 | 0 | 2 | 2 | 0 | 0 | 2 | 0 | 0 | 1 |
| ko00750 Vitamin B6 metabolism | 23086 | 22667 | 23367 | 23030 | 24096 | 24317 | 24101 | 24190 | 23554 | 23284 | 23850 | 23711 |
| K00097 4-hydroxythreonine-4-phosphate dehydrogenase [EC:1.1.1.262] | 2960 | 2953 | 3132 | 2991 | 3372 | 3586 | 3568 | 3516 | 2813 | 2853 | 3013 | 2932 |
| K00275 pyridoxamine 5'-phosphate oxidase [EC:1.4.3.5] | 1726 | 1683 | 1734 | 1679 | 2065 | 2058 | 2103 | 2149 | 1777 | 1748 | 1757 | 1745 |
| K00868 pyridoxine kinase [EC:2.7.1.35] | 3578 | 3318 | 3436 | 3500 | 3294 | 3252 | 3229 | 3131 | 3631 | 3703 | 3660 | 3803 |
| K03472 D-erythrose 4-phosphate dehydrogenase [EC:1.2.1.72] | 2305 | 2249 | 2257 | 2171 | 1443 | 1364 | 1336 | 1300 | 1414 | 1362 | 1385 | 1490 |
| K03473 erythronate-4-phosphate dehydrogenase [EC:1.1.1.290] | 2646 | 2529 | 2568 | 2668 | 2464 | 2501 | 2379 | 2635 | 2134 | 2163 | 2225 | 2285 |
| K03474 pyridoxine 5-phosphate synthase [EC:2.6.99.2] | 1775 | 1969 | 1952 | 1812 | 2458 | 2493 | 2542 | 2416 | 2119 | 2156 | 2131 | 1970 |
| K05275 pyridoxine 4-dehydrogenase [EC:1.1.1.65] | 465 | 476 | 494 | 556 | 375 | 374 | 337 | 353 | 780 | 749 | 789 | 740 |
| K06215 pyridoxal 5'-phosphate synthase pdxS subunit [EC:4.3.3.6] | 220 | 174 | 229 | 199 | 57 | 62 | 88 | 55 | 377 | 384 | 351 | 362 |
| K08681 5'-phosphate synthase pdxT subunit [EC:4.3.3.6] | 93 | 95 | 88 | 108 | 26 | 27 | 33 | 30 | 208 | 222 | 190 | 212 |
| ko00760 Nicotinate and nicotinamide metabolism | 59558 | 60554 | 59167 | 60289 | 56365 | 56642 | 55916 | 56048 | 57758 | 57918 | 56796 | 57969 |
| K00322 NAD(P) transhydrogenase [EC:1.6.1.1] | 2781 | 2779 | 2718 | 2741 | 1775 | 1707 | 1608 | 1634 | 1688 | 2005 | 1783 | 1817 |
| K00324 H+-translocating NAD(P) transhydrogenase subunit alpha [EC:1.6.1.2 7.1.1.1] | 2888 | 2849 | 2754 | 2906 | 1638 | 1695 | 1518 | 1726 | 1814 | 2016 | 1922 | 1929 |
| K00325 H+-translocating NAD(P) transhydrogenase subunit beta [EC:1.6.1.2 7.1.1.1] | 2693 | 2723 | 2530 | 2680 | 1608 | 1625 | 1494 | 1679 | 1727 | 1767 | 1546 | 1640 |
| K00763 nicotinate phosphoribosyltransferase [EC:6.3.4.21] | 5586 | 5535 | 5523 | 5912 | 5298 | 5116 | 4970 | 5139 | 4930 | 4817 | 4804 | 4864 |
| K00767 nicotinate-nucleotide pyrophosphorylase (carboxylating) [EC:2.4.2.19] | 2535 | 2631 | 2535 | 2675 | 2884 | 3032 | 2916 | 2886 | 2527 | 2422 | 2480 | 2451 |
| K00858 NAD+ kinase [EC:2.7.1.23] | 2402 | 2491 | 2459 | 2402 | 2983 | 2853 | 2957 | 2879 | 2913 | 2792 | 2797 | 3023 |
| K00952 nicotinamide-nucleotide adenylyltransferase [EC:2.7.7.1] | 0 | 0 | 2 | 1 | 0 | 0 | 0 | 0 | 2 | 2 | 4 | 4 |
| K00969 nicotinate-nucleotide adenylyltransferase [EC:2.7.7.18] | 1937 | 1985 | 1914 | 1872 | 2119 | 2178 | 2093 | 2230 | 2280 | 1964 | 1991 | 2106 |
| K01440 nicotinamidase [EC:3.5.1.19] | 1 | 3 | 2 | 2 | 1 | 0 | 4 | 2 | 4 | 2 | 2 | 4 |
| K01916 NAD+ synthase [EC:6.3.1.5] | 1964 | 2007 | 1923 | 1863 | 1167 | 1176 | 1132 | 1231 | 1231 | 1249 | 1141 | 1332 |
| K01950 NAD+ synthase (glutamine-hydrolysing) [EC:6.3.5.1] | 1641 | 1831 | 1760 | 1693 | 2907 | 2996 | 3072 | 2914 | 2775 | 2845 | 2856 | 2671 |
| K03426 NAD+ diphosphatase [EC:3.6.1.22] | 2450 | 2564 | 2504 | 2504 | 2696 | 2686 | 2733 | 2785 | 2295 | 2350 | 2267 | 2416 |
| K03462 nicotinamide phosphoribosyltransferase [EC:2.4.2.12] | 316 | 300 | 294 | 274 | 392 | 430 | 469 | 434 | 8 | 21 | 7 | 5 |
| K03517 quinolinate synthase [EC:2.5.1.72] | 2648 | 2927 | 2791 | 2837 | 3454 | 3494 | 3466 | 3475 | 3065 | 2987 | 3012 | 2910 |
| K03742 nicotinamide-nucleotide amidase [EC:3.5.1.42] | 1172 | 1183 | 1295 | 1218 | 2183 | 2336 | 2352 | 2279 | 2190 | 2152 | 2240 | 2092 |
| K03743 nicotinamide-nucleotide amidase [EC:3.5.1.42] | 1064 | 1001 | 1037 | 979 | 684 | 754 | 695 | 749 | 654 | 601 | 662 | 682 |
| K06211 HTH-type transcriptional regulator, transcriptional repressor of NAD biosynthesis genes [EC:2.7.7.1 2.7.1.22] | 840 | 843 | 815 | 782 | 546 | 548 | 485 | 510 | 1189 | 1171 | 1161 | 1174 |
| K06989 aspartate dehydrogenase [EC:1.4.1.21] | 859 | 938 | 1007 | 903 | 2005 | 1840 | 2028 | 1959 | 1554 | 1555 | 1521 | 1463 |
| K08281 nicotinamidase/pyrazinamidase [EC:3.5.1.19 3.5.1.-] | 1235 | 1297 | 1163 | 1248 | 683 | 780 | 717 | 738 | 826 | 876 | 870 | 853 |
| K13522 bifunctional NMN adenylyltransferase/nudix hydrolase [EC:2.7.7.1 3.6.1.-] | 3 | 2 | 4 | 9 | 12 | 16 | 11 | 16 | 9 | 10 | 8 | 7 |
| K18029 nicotinate dehydrogenase subunit A [EC:1.17.2.1] | 749 | 786 | 699 | 827 | 402 | 387 | 388 | 397 | 265 | 216 | 232 | 231 |
| K18030 nicotinate dehydrogenase subunit B [EC:1.17.2.1] | 1 | 0 | 0 | 0 | 0 | 0 | 1 | 0 | 0 | 1 | 0 | 0 |
| K19647 2-hydroxymethylglutarate dehydrogenase [EC:1.1.1.291] | 3 | 0 | 1 | 2 | 0 | 2 | 0 | 0 | 4 | 5 | 1 | 5 |
| K20445 nicotinate dehydrogenase FAD-subunit [EC:1.17.1.5] | 10 | 5 | 9 | 15 | 3 | 1 | 3 | 2 | 7 | 6 | 6 | 4 |
| K20446 nicotinate dehydrogenase small FeS subunit [EC:1.17.1.5] | 3 | 2 | 1 | 3 | 0 | 1 | 0 | 0 | 5 | 3 | 2 | 1 |
| K20447 nicotinate dehydrogenase large molybdopterin subunit [EC:1.17.1.5] | 25 | 31 | 31 | 35 | 11 | 5 | 8 | 13 | 20 | 18 | 19 | 18 |
| K20448 nicotinate dehydrogenase medium molybdopterin subunit [EC:1.17.1.5] | 0 | 3 | 0 | 0 | 0 | 0 | 0 | 0 | 0 | 0 | 0 | 1 |
| K20449 6-hydroxynicotinate reductase [EC:1.3.7.1] | 0 | 0 | 0 | 1 | 0 | 0 | 0 | 0 | 3 | 5 | 2 | 1 |
| K20451 methylitaconate Delta-isomerase [EC:5.3.3.6] | 9 | 3 | 5 | 11 | 0 | 0 | 0 | 0 | 0 | 1 | 3 | 1 |
| K20452 dimethylmaleate hydratase large subunit [EC:4.2.1.85] | 20 | 17 | 21 | 21 | 13 | 17 | 21 | 21 | 42 | 61 | 50 | 51 |
| K20453 dimethylmaleate hydratase small subunit [EC:4.2.1.85] | 4 | 2 | 4 | 3 | 1 | 5 | 1 | 10 | 4 | 16 | 14 | 11 |
| K20454 2,3-dimethylmalate lyase [EC:4.1.3.32] | 10 | 3 | 6 | 6 | 0 | 0 | 0 | 0 | 6 | 3 | 7 | 5 |
| ko00770 Pantothenate and CoA biosynthesis | 63114 | 62017 | 61895 | 62925 | 55709 | 56100 | 54483 | 55734 | 58102 | 59151 | 58005 | 59172 |
| K00077 2-dehydropantoate 2-reductase [EC:1.1.1.169] | 3579 | 3561 | 3309 | 3444 | 3587 | 3491 | 3369 | 3663 | 3820 | 3883 | 3711 | 3763 |
| K00606 3-methyl-2-oxobutanoate hydroxymethyltransferase [EC:2.1.2.11] | 3045 | 2788 | 2804 | 2908 | 3043 | 2993 | 2992 | 3058 | 2371 | 2307 | 2438 | 2362 |
| K00859 dephospho-CoA kinase [EC:2.7.1.24] | 1675 | 1650 | 1809 | 1684 | 1928 | 2039 | 1977 | 2001 | 1939 | 1992 | 1988 | 1995 |
| K00867 type I pantothenate kinase [EC:2.7.1.33] | 670 | 646 | 648 | 681 | 465 | 498 | 492 | 476 | 902 | 945 | 829 | 888 |
| K00954 pantetheine-phosphate adenylyltransferase [EC:2.7.7.3] | 1663 | 1703 | 1758 | 1690 | 1844 | 1954 | 1803 | 1872 | 1711 | 1763 | 1769 | 1812 |
| K00997 holo-[acyl-carrier protein] synthase [EC:2.7.8.7] | 264 | 284 | 285 | 306 | 170 | 146 | 160 | 140 | 375 | 373 | 337 | 401 |
| K01598 phosphopantothenoylcysteine decarboxylase [EC:4.1.1.36] | 31 | 43 | 30 | 38 | 5 | 13 | 10 | 7 | 34 | 57 | 26 | 32 |
| K02201 pantetheine-phosphate adenylyltransferase [EC:2.7.7.3] | 0 | 1 | 2 | 0 | 0 | 3 | 1 | 0 | 2 | 0 | 2 | 3 |
| K03525 type III pantothenate kinase [EC:2.7.1.33] | 921 | 977 | 1016 | 985 | 1615 | 1725 | 1678 | 1703 | 1611 | 1536 | 1591 | 1657 |
| K06133 4'-phosphopantetheinyl transferase [EC:2.7.8.-] | 159 | 139 | 128 | 124 | 68 | 76 | 77 | 73 | 269 | 248 | 234 | 234 |
| K06982 pantoate kinase [EC:2.7.1.169] | 0 | 0 | 2 | 0 | 2 | 0 | 0 | 0 | 8 | 1 | 3 | 5 |
| K08682 acyl carrier protein phosphodiesterase [EC:3.1.4.14] | 454 | 438 | 392 | 407 | 255 | 293 | 263 | 221 | 622 | 670 | 617 | 678 |
| K09680 type II pantothenate kinase [EC:2.7.1.33] | 469 | 477 | 519 | 513 | 242 | 223 | 254 | 245 | 203 | 268 | 256 | 222 |
| K13038 phosphopantothenoylcysteine decarboxylase / phosphopantothenate---cysteine ligase [EC:4.1.1.36 6.3.2.5] | 3792 | 3774 | 3977 | 3930 | 4328 | 4285 | 4247 | 4235 | 3864 | 3742 | 3981 | 3962 |
| K21977 phosphopantothenate---cysteine ligase (CTP) [EC:6.3.2.5] | 30 | 58 | 36 | 59 | 17 | 14 | 13 | 16 | 64 | 53 | 56 | 44 |
| ko00780 Biotin metabolism | 52419 | 52001 | 51297 | 51644 | 50898 | 50582 | 50622 | 50821 | 47169 | 46759 | 46602 | 46867 |
| K00652 8-amino-7-oxononanoate synthase [EC:2.3.1.47] | 3864 | 3973 | 3828 | 4002 | 5987 | 6010 | 5968 | 6186 | 4780 | 4732 | 4738 | 4513 |
| K00833 adenosylmethionine---8-amino-7-oxononanoate aminotransferase [EC:2.6.1.62] | 4656 | 4722 | 4614 | 4617 | 6637 | 6775 | 6836 | 6557 | 5454 | 5287 | 5334 | 5309 |
| K01012 biotin synthase [EC:2.8.1.6] | 3621 | 3621 | 3651 | 3482 | 3961 | 3827 | 4014 | 3916 | 3594 | 3594 | 3557 | 3486 |
| K01906 6-carboxyhexanoate--CoA ligase [EC:6.2.1.14] | 36 | 31 | 43 | 36 | 3 | 2 | 3 | 0 | 47 | 51 | 50 | 66 |
| K01935 dethiobiotin synthetase [EC:6.3.3.3] | 2240 | 2175 | 1965 | 2068 | 2315 | 2356 | 2229 | 2328 | 2340 | 2337 | 2244 | 2432 |
| K02169 malonyl-CoA O-methyltransferase [EC:2.1.1.197] | 2220 | 2226 | 2058 | 2114 | 2486 | 2417 | 2372 | 2491 | 2141 | 2005 | 1976 | 2027 |
| K02170 pimeloyl-[acyl-carrier protein] methyl ester esterase [EC:3.1.1.85] | 1319 | 1318 | 1166 | 1214 | 739 | 786 | 761 | 776 | 899 | 892 | 897 | 987 |
| K03524 BirA family transcriptional regulator, biotin operon repressor / biotin---[acetyl-CoA-carboxylase] ligase [EC:6.3.4.15] | 2884 | 2920 | 3011 | 3065 | 2971 | 2878 | 3016 | 2861 | 2984 | 2990 | 3044 | 3041 |
| K08351 biotin/methionine sulfoxide reductase [EC:1.-.-.-] | 2765 | 2777 | 2770 | 2965 | 1999 | 2199 | 2088 | 2057 | 3616 | 3592 | 3542 | 3745 |
| K09789 pimeloyl-[acyl-carrier protein] methyl ester esterase [EC:3.1.1.85] | 420 | 412 | 515 | 460 | 1345 | 1381 | 1374 | 1372 | 929 | 945 | 947 | 891 |
| ko00785 Lipoic acid metabolism | 5132 | 5236 | 5166 | 5296 | 6687 | 6801 | 6843 | 6620 | 6268 | 6246 | 6287 | 6388 |
| K03644 lipoyl synthase [EC:2.8.1.8] | 2019 | 1993 | 1996 | 2054 | 2671 | 2719 | 2634 | 2661 | 2222 | 2341 | 2273 | 2342 |
| K03800 lipoate---protein ligase [EC:6.3.1.20] | 1521 | 1540 | 1446 | 1557 | 1963 | 1961 | 1997 | 1876 | 2337 | 2238 | 2273 | 2323 |
| K03801 lipoyl(octanoyl) transferase [EC:2.3.1.181] | 1592 | 1703 | 1724 | 1685 | 2053 | 2121 | 2212 | 2083 | 1709 | 1667 | 1741 | 1723 |
| ko00790 Folate biosynthesis | 58287 | 58335 | 57107 | 58486 | 56359 | 55992 | 56085 | 56472 | 50063 | 49360 | 49527 | 49777 |
| K00287 dihydrofolate reductase [EC:1.5.1.3] | 1870 | 1966 | 1797 | 1854 | 2007 | 1976 | 1869 | 1991 | 1901 | 1836 | 1768 | 1819 |
| K00796 dihydropteroate synthase [EC:2.5.1.15] | 2557 | 2549 | 2575 | 2525 | 3195 | 3197 | 3098 | 3016 | 2376 | 2229 | 2421 | 2288 |
| K00950 2-amino-4-hydroxy-6-hydroxymethyldihydropteridine diphosphokinase [EC:2.7.6.3] | 1869 | 1999 | 1975 | 2000 | 2617 | 2627 | 2654 | 2564 | 2174 | 2141 | 2140 | 2191 |
| K01113 alkaline phosphatase D [EC:3.1.3.1] | 2489 | 2529 | 2267 | 2456 | 1483 | 1253 | 1253 | 1372 | 763 | 775 | 716 | 816 |
| K01495 GTP cyclohydrolase IA [EC:3.5.4.16] | 2349 | 2352 | 2405 | 2392 | 2446 | 2325 | 2518 | 2410 | 2027 | 1996 | 2009 | 2010 |
| K01633 7,8-dihydroneopterin aldolase/epimerase/oxygenase [EC:4.1.2.25 5.1.99.8 1.13.11.81] | 1082 | 1174 | 1080 | 1178 | 1361 | 1246 | 1327 | 1260 | 1139 | 1204 | 1066 | 1125 |
| K01664 para-aminobenzoate synthetase component II [EC:2.6.1.85] | 396 | 402 | 425 | 402 | 248 | 307 | 336 | 254 | 511 | 543 | 550 | 595 |
| K01665 para-aminobenzoate synthetase component I [EC:2.6.1.85] | 2908 | 3036 | 3014 | 3122 | 3426 | 3505 | 3416 | 3479 | 3038 | 3042 | 2995 | 2953 |
| K01724 4a-hydroxytetrahydrobiopterin dehydratase [EC:4.2.1.96] | 540 | 489 | 493 | 470 | 278 | 261 | 249 | 271 | 155 | 150 | 119 | 143 |
| K01737 6-pyruvoyltetrahydropterin/6-carboxytetrahydropterin synthase [EC:4.2.3.12 4.1.2.50] | 975 | 1026 | 920 | 971 | 1213 | 1217 | 1267 | 1276 | 990 | 1045 | 987 | 992 |
| K01930 folylpolyglutamate synthase [EC:6.3.2.17] | 0 | 0 | 2 | 1 | 0 | 0 | 1 | 0 | 0 | 1 | 0 | 0 |
| K02619 4-amino-4-deoxychorismate lyase [EC:4.1.3.38] | 2023 | 2066 | 1963 | 2004 | 2076 | 2157 | 2160 | 2119 | 1902 | 1726 | 1869 | 1817 |
| K03635 molybdopterin synthase catalytic subunit [EC:2.8.1.12] | 1026 | 943 | 931 | 983 | 587 | 635 | 554 | 606 | 718 | 675 | 606 | 651 |
| K03637 cyclic pyranopterin monophosphate synthase [EC:4.6.1.17] | 1454 | 1302 | 1284 | 1376 | 690 | 796 | 750 | 777 | 884 | 893 | 856 | 875 |
| K03638 molybdopterin adenylyltransferase [EC:2.7.7.75] | 1785 | 1930 | 1651 | 1778 | 950 | 995 | 967 | 1055 | 896 | 905 | 900 | 1006 |
| K03639 GTP 3',8-cyclase [EC:4.1.99.22] | 4352 | 4054 | 4045 | 4159 | 2344 | 2304 | 2291 | 2440 | 2369 | 2216 | 2268 | 2430 |
| K03750 molybdopterin molybdotransferase [EC:2.10.1.1] | 4770 | 4504 | 4363 | 4442 | 2670 | 2749 | 2553 | 2789 | 2549 | 2829 | 2764 | 2924 |
| K03752 molybdenum cofactor guanylyltransferase [EC:2.7.7.77] | 1288 | 1276 | 1185 | 1245 | 691 | 762 | 741 | 713 | 821 | 799 | 775 | 822 |
| K03831 molybdopterin adenylyltransferase [EC:2.7.7.75] | 403 | 442 | 396 | 477 | 348 | 379 | 342 | 359 | 615 | 612 | 637 | 668 |
| K06879 7-cyano-7-deazaguanine reductase [EC:1.7.1.13] | 1563 | 1628 | 1546 | 1649 | 1008 | 987 | 1003 | 1015 | 1163 | 1150 | 1049 | 1202 |
| K06897 7,8-dihydropterin-6-yl-methyl-4-(beta-D-ribofuranosyl)aminobenzene 5'-phosphate synthase [EC:2.5.1.105] | 538 | 566 | 663 | 592 | 1360 | 1291 | 1391 | 1342 | 1345 | 1442 | 1357 | 1270 |
| K06920 7-cyano-7-deazaguanine synthase [EC:6.3.4.20] | 1614 | 1602 | 1727 | 1862 | 2139 | 2234 | 2088 | 2091 | 1981 | 1870 | 1820 | 1721 |
| K06984 beta-ribofuranosylaminobenzene 5'-phosphate synthase [EC:2.4.2.54] | 0 | 0 | 2 | 0 | 0 | 0 | 0 | 2 | 5 | 4 | 8 | 5 |
| K07141 molybdenum cofactor cytidylyltransferase [EC:2.7.7.76] | 1371 | 1336 | 1194 | 1399 | 651 | 631 | 634 | 596 | 632 | 663 | 652 | 599 |
| K07142 2-amino-4-hydroxy-6-hydroxymethyldihydropteridine diphosphokinase [EC:2.7.6.3] | 2 | 2 | 1 | 0 | 0 | 0 | 0 | 0 | 4 | 0 | 0 | 10 |

| | | | | | | | | | | | | |
|---|---|---|---|---|---|---|---|---|---|---|---|---|
| K07589 D-erythro-7,8-dihydroneopterin triphosphate epimerase [EC:5.1.99.7] | 710 | 719 | 711 | 738 | 412 | 411 | 452 | 487 | 460 | 460 | 431 | 422 |
| K08310 dihydroneopterin triphosphate diphosphatase [EC:3.6.1.67] | 267 | 215 | 243 | 272 | 207 | 200 | 185 | 191 | 345 | 336 | 349 | 334 |
| K09007 GTP cyclohydrolase IB [EC:3.5.4.16] | 1681 | 1672 | 1617 | 1648 | 929 | 927 | 955 | 995 | 582 | 581 | 526 | 602 |
| K09457 7-cyano-7-deazaguanine reductase [EC:1.7.1.13] | 280 | 306 | 328 | 315 | 969 | 913 | 1012 | 960 | 718 | 766 | 700 | 668 |
| K09739 dihydroneopterin aldolase [EC:4.1.2.25] | 0 | 0 | 2 | 0 | 1 | 0 | 0 | 0 | 1 | 3 | 1 | 4 |
| K10026 7-carboxy-7-deazaguanine synthase [EC:4.3.99.3] | 1826 | 1828 | 1887 | 1846 | 2044 | 2061 | 2148 | 2107 | 1737 | 1666 | 1679 | 1801 |
| K11754 dihydrofolate synthase / folylpolyglutamate synthase [EC:6.3.2.12 6.3.2.17] | 3561 | 3514 | 3520 | 3516 | 4296 | 4091 | 4236 | 4304 | 4116 | 3981 | 4245 | 4130 |
| K13818 molybdopterin-guanine dinucleotide biosynthesis protein [EC:2.7.7.77] | 49 | 35 | 55 | 46 | 30 | 41 | 35 | 36 | 16 | 30 | 15 | 20 |
| K13938 dihydromonapterin reductase / dihydrofolate reductase [EC:1.5.1.50 1.5.1.3] | 761 | 842 | 766 | 768 | 473 | 460 | 376 | 451 | 711 | 632 | 630 | 692 |
| K13939 dihydroneopterin aldolase / 2-amino-4-hydroxy-6-hydroxymethyldihydropteridine diphosphokinase / dihydropteroate synthase [EC:4.1.2.25 2.7.6.3 2 | 0 | 0 | 2 | 0 | 0 | 0 | 0 | 0 | 0 | 0 | 0 | 0 |
| K13940 dihydroneopterin aldolase / 2-amino-4-hydroxy-6-hydroxymethyldihydropteridine diphosphokinase [EC:4.1.2.25 2.7.6.3] | 87 | 107 | 104 | 111 | 23 | 41 | 24 | 24 | 154 | 181 | 153 | 200 |
| K13950 para-aminobenzoate synthetase [EC:2.6.1.85] | 0 | 0 | 2 | 1 | 0 | 0 | 0 | 0 | 0 | 0 | 1 | 1 |
| K15376 gephyrin [EC:2.10.1.1 2.7.7.75] | 1 | 0 | 3 | 3 | 1 | 1 | 4 | 2 | 0 | 2 | 2 | 3 |
| K17488 GTP cyclohydrolase IV [EC:3.5.4.39] | 0 | 2 | 0 | 1 | 2 | 5 | 0 | 0 | 1 | 9 | 3 | 8 |
| K18590 dihydrofolate reductase (trimethoprim resistance protein) [EC:1.5.1.3] | 0 | 0 | 0 | 0 | 0 | 1 | 0 | 0 | 0 | 0 | 0 | 1 |
| K18824 dihydropteroate synthase type 2 [EC:2.5.1.15] | 0 | 0 | 0 | 0 | 0 | 0 | 0 | 0 | 0 | 3 | 0 | 1 |
| K18853 dihydromethanopterin reductase (acceptor) [EC:1.5.99.15] | 2 | 0 | 3 | 0 | 0 | 0 | 0 | 0 | 0 | 2 | 2 | 6 |
| K18974 dihydropteroate synthase type 1 [EC:2.5.1.15] | 0 | 0 | 0 | 0 | 3 | 0 | 2 | 2 | 4 | 0 | 2 | 2 |
| K20967 GTP 3',8-cyclase / cyclic pyranopterin monophosphate synthase [EC:4.1.99.22 4.6.1.17] | 0 | 0 | 0 | 0 | 0 | 0 | 0 | 3 | 1 | 0 | 0 | 0 |
| ko00670 One carbon pool by folate | 54772 | 54409 | 54069 | 54069 | 59811 | 59720 | 60266 | 59574 | 56390 | 54279 | 54334 | 54214 |
| K00288 methylenetetrahydrofolate dehydrogenase (NADP+) / methenyltetrahydrofolate cyclohydrolase / formyltetrahydrofolate synthetase [EC:1.5.1.5 3.5.4 | 0 | 0 | 1 | 0 | 0 | 0 | 0 | 2 | 0 | 0 | 0 | 0 |
| K00604 methionyl-tRNA formyltransferase [EC:2.1.2.9] | 3337 | 3486 | 3389 | 3382 | 3518 | 3551 | 3628 | 3484 | 3402 | 3362 | 3462 | 3391 |
| K01500 methenyltetrahydrofolate cyclohydrolase [EC:3.5.4.9] | 20 | 9 | 9 | 7 | 1 | 3 | 0 | 3 | 11 | 14 | 20 | 15 |
| K01746 formiminotetrahydrofolate cyclodeaminase [EC:4.3.1.4] | 1 | 0 | 0 | 0 | 34 | 28 | 34 | 21 | 3 | 4 | 4 | 7 |
| K01934 5-formyltetrahydrofolate cyclo-ligase [EC:6.3.3.2] | 1982 | 1962 | 1802 | 1880 | 2027 | 2058 | 1962 | 1890 | 1803 | 1750 | 1746 | 1715 |
| K13403 methylenetetrahydrofolate dehydrogenase(NAD+) / 5,10-methenyltetrahydrofolate cyclohydrolase [EC:1.5.1.15 3.5.4.9] | 0 | 0 | 1 | 0 | 0 | 0 | 0 | 0 | 0 | 1 | 1 | 0 |
| ko00830 Retinol metabolism | 5941 | 6112 | 5761 | 5748 | 3891 | 3929 | 3897 | 3874 | 4653 | 4767 | 4640 | 4615 |
| K07249 retinal dehydrogenase [EC:1.2.1.36] | 0 | 1 | 0 | 0 | 0 | 0 | 0 | 0 | 0 | 1 | 0 | 0 |
| K09516 all-trans-retinol 13,14-reductase [EC:1.3.99.23] | 166 | 209 | 195 | 173 | 300 | 277 | 335 | 302 | 19 | 50 | 29 | 34 |
| K11153 retinol dehydrogenase 12 [EC:1.1.1.300] | 1 | 0 | 0 | 0 | 0 | 1 | 0 | 0 | 0 | 0 | 1 | 0 |
| K15734 all-trans-retinol dehydrogenase (NAD+) [EC:1.1.1.105] | 0 | 0 | 1 | 1 | 0 | 0 | 0 | 0 | 0 | 0 | 0 | 0 |
| ko00860 Porphyrin and chlorophyll metabolism | 107231 | 106167 | 101632 | 103492 | 85572 | 84623 | 83570 | 85907 | 87872 | 86755 | 86125 | 88181 |
| K00228 coproporphyrinogen III oxidase [EC:1.3.3.3] | 1705 | 1678 | 1563 | 1636 | 1004 | 1006 | 907 | 1004 | 1132 | 1089 | 1047 | 1205 |
| K00230 menaquinone-dependent protoporphyrinogen oxidase [EC:1.3.5.3] | 643 | 656 | 658 | 601 | 1344 | 1476 | 1412 | 1398 | 1393 | 1233 | 1347 | 1301 |
| K00231 protoporphyrinogen/coproporphyrinogen III oxidase [EC:1.3.3.4 1.3.3.15] | 21 | 19 | 9 | 23 | 3 | 4 | 2 | 1 | 5 | 6 | 2 | 3 |
| K00435 Fe-coproporphyrin III decarboxylase [EC:1.11.1.-] | 5 | 1 | 16 | 4 | 0 | 2 | 0 | 0 | 0 | 0 | 0 | 0 |
| K00595 precorrin-6Y C5,15-methyltransferase (decarboxylating) [EC:2.1.1.132] | 2523 | 2527 | 2392 | 2232 | 2508 | 2397 | 2435 | 2517 | 1992 | 1918 | 1940 | 1911 |
| K00768 nicotinate-nucleotide--dimethylbenzimidazole phosphoribosyltransferase [EC:2.4.2.21] | 3496 | 3344 | 3195 | 3212 | 3467 | 3553 | 3463 | 3463 | 4066 | 4046 | 4190 | 4181 |
| K00798 cob(I)alamin adenosyltransferase [EC:2.5.1.17] | 1525 | 1524 | 1519 | 1485 | 1838 | 1768 | 1950 | 1872 | 1274 | 1233 | 1348 | 1355 |
| K01599 uroporphyrinogen decarboxylase [EC:4.1.1.37] | 4728 | 4731 | 4722 | 4832 | 4010 | 3847 | 3962 | 3800 | 5051 | 5254 | 5075 | 5250 |
| K01698 porphobilinogen synthase [EC:4.2.1.24] | 1666 | 1702 | 1755 | 1713 | 1127 | 1159 | 1066 | 1119 | 1610 | 1740 | 1710 | 1633 |
| K01719 uroporphyrinogen-III synthase [EC:4.2.1.75] | 2077 | 2174 | 2045 | 2125 | 2745 | 2754 | 2751 | 2744 | 2278 | 2258 | 2234 | 2258 |
| K01749 hydroxymethylbilane synthase [EC:2.5.1.61] | 2741 | 2700 | 2590 | 2731 | 1548 | 1509 | 1568 | 1546 | 1858 | 1789 | 1711 | 1825 |
| K01772 protoporphyrin/coproporphyrin ferrochelatase [EC:4.99.1.1 4.99.1.9] | 2321 | 2326 | 2149 | 2417 | 1410 | 1385 | 1369 | 1402 | 1427 | 1399 | 1430 | 1451 |
| K01845 glutamate-1-semialdehyde 2,1-aminomutase [EC:5.4.3.8] | 4650 | 4500 | 4416 | 4575 | 2808 | 2739 | 2266 | 2515 | 2773 | 2796 | 2700 | 2837 |
| K01885 glutamyl-tRNA synthetase [EC:6.1.1.17] | 4654 | 4628 | 4821 | 4547 | 5461 | 5490 | 5648 | 5570 | 5240 | 5181 | 5266 | 5201 |
| K02188 cobalt-precorrin-5B (C1)-methyltransferase [EC:2.1.1.195] | 2691 | 2742 | 2441 | 2512 | 2670 | 2616 | 2549 | 2681 | 2672 | 2605 | 2617 | 2604 |
| K02189 cobalt-precorrin 5A hydrolase [EC:3.7.1.12] | 1400 | 1388 | 1294 | 1422 | 803 | 711 | 690 | 808 | 1327 | 1310 | 1346 | 1356 |
| K02190 sirohydrochlorin cobaltochelatase [EC:4.99.1.3] | 1394 | 1517 | 1392 | 1537 | 2321 | 2366 | 2446 | 2294 | 2292 | 2211 | 2297 | 2212 |
| K02191 cobalt-precorrin-6B (C15)-methyltransferase [EC:2.1.1.196] | 340 | 336 | 315 | 309 | 219 | 228 | 207 | 224 | 346 | 341 | 319 | 361 |
| K02224 cobyrinic acid a,c-diamide synthase [EC:6.3.5.9 6.3.5.11] | 3600 | 3658 | 3339 | 3466 | 3065 | 2838 | 2823 | 3001 | 3075 | 3041 | 2956 | 3034 |
| K02225 cobalamin biosynthetic protein CobC | 1352 | 1390 | 1102 | 1185 | 678 | 657 | 558 | 642 | 424 | 331 | 346 | 395 |
| K02226 alpha-ribazole phosphatase [EC:3.1.3.73] | 1656 | 1656 | 1589 | 1579 | 1915 | 1876 | 1886 | 1880 | 1761 | 1676 | 1748 | 1696 |
| K02227 adenosylcobinamide-phosphate synthase [EC:6.3.1.10] | 2266 | 2255 | 2046 | 2109 | 1745 | 1598 | 1468 | 1659 | 1636 | 1659 | 1666 | 1730 |
| K02229 precorrin-3B synthase [EC:1.14.13.83] | 1437 | 1389 | 1114 | 1231 | 608 | 611 | 628 | 761 | 414 | 366 | 340 | 416 |
| K02230 cobaltochelatase CobN [EC:6.6.1.2] | 10190 | 10207 | 9284 | 9685 | 5545 | 5371 | 5200 | 5913 | 3310 | 3008 | 2924 | 3061 |
| K02231 adenosylcobinamide kinase / adenosylcobinamide-phosphate guanylyltransferase [EC:2.7.1.156 2.7.7.62] | 1568 | 1572 | 1478 | 1543 | 1596 | 1572 | 1593 | 1530 | 1548 | 1489 | 1579 | 1625 |
| K02232 adenosylcobyric acid synthase [EC:6.3.5.10] | 3752 | 3683 | 3480 | 3543 | 2597 | 2595 | 2562 | 2602 | 3047 | 3109 | 2919 | 3026 |
| K02233 adenosylcobinamide-GDP ribazoletransferase [EC:2.7.8.26] | 2102 | 2147 | 2069 | 2043 | 2459 | 2477 | 2280 | 2476 | 2285 | 2246 | 2388 | 2309 |
| K02234 cobalamin biosynthesis protein CobW | 1692 | 1648 | 1506 | 1438 | 853 | 763 | 694 | 727 | 494 | 493 | 419 | 485 |
| K02302 uroporphyrin-III C-methyltransferase / precorrin-2 dehydrogenase / sirohydrochlorin ferrochelatase [EC:2.1.1.107 1.3.1.76 4.99.1.4] | 2627 | 2711 | 2438 | 2655 | 1564 | 1675 | 1647 | 1602 | 2539 | 2649 | 2605 | 2650 |
| K02303 uroporphyrin-III C-methyltransferase [EC:2.1.1.107] | 2540 | 2582 | 2452 | 2542 | 1472 | 1467 | 1417 | 1445 | 812 | 693 | 715 | 777 |
| K02304 precorrin-2 dehydrogenase / sirohydrochlorin ferrochelatase [EC:1.3.1.76 4.99.1.4] | 239 | 210 | 208 | 241 | 52 | 59 | 45 | 57 | 227 | 238 | 244 | 246 |
| K02492 glutamyl-tRNA reductase [EC:1.2.1.70] | 3206 | 3030 | 3076 | 2865 | 1904 | 1693 | 1830 | 1865 | 2167 | 2095 | 2028 | 2120 |
| K02495 oxygen-independent coproporphyrinogen III oxidase [EC:1.3.98.3] | 1933 | 1963 | 2004 | 1974 | 1293 | 1287 | 1341 | 1257 | 1671 | 1782 | 1621 | 1736 |
| K02496 uroporphyrin-III C-methyltransferase [EC:2.1.1.107] | 2751 | 2649 | 2568 | 2620 | 1636 | 1609 | 1527 | 1599 | 1570 | 1506 | 1495 | 1559 |
| K03394 precorrin-2/cobalt-factor-2 C20-methyltransferase [EC:2.1.1.130 2.1.1.151] | 1890 | 1885 | 1725 | 1834 | 978 | 956 | 1001 | 1056 | 1240 | 1266 | 1230 | 1234 |
| K03399 cobalt-precorrin-7 (C5)-methyltransferase [EC:2.1.1.289] | 450 | 366 | 350 | 360 | 244 | 256 | 210 | 208 | 542 | 522 | 497 | 540 |
| K03404 magnesium chelatase subunit D [EC:6.6.1.1] | 76 | 93 | 95 | 64 | 8 | 4 | 5 | 2 | 82 | 90 | 95 | 121 |
| K03405 magnesium chelatase subunit I [EC:6.6.1.1] | 356 | 357 | 324 | 316 | 159 | 176 | 190 | 211 | 165 | 136 | 135 | 172 |
| K03594 bacterioferritin [EC:1.16.3.1] | 2524 | 2510 | 2698 | 2466 | 2382 | 2384 | 2170 | 2433 | 1866 | 1864 | 1890 | 1780 |
| K04032 ethanolamine utilization cobalamin adenosyltransferase [EC:2.5.1.17] | 328 | 275 | 327 | 334 | 83 | 99 | 106 | 107 | 690 | 732 | 654 | 810 |
| K04034 anaerobic magnesium-protoporphyrin IX monomethyl ester cyclase [EC:1.21.98.3] | 3 | 1 | 3 | 1 | 1 | 0 | 0 | 0 | 5 | 9 | 5 | 6 |
| K04720 threonine-phosphate decarboxylase [EC:4.1.1.81] | 1353 | 1439 | 1470 | 1432 | 1879 | 1811 | 1959 | 1945 | 2207 | 2261 | 2235 | 2173 |
| K05895 precorrin-6A/cobalt-precorrin-6A reductase [EC:1.3.1.54 1.3.1.106] | 1412 | 1332 | 1208 | 1373 | 745 | 731 | 710 | 796 | 1056 | 1057 | 988 | 1052 |
| K05934 precorrin-3B C17-methyltransferase [EC:2.1.1.131] | 600 | 600 | 558 | 583 | 319 | 314 | 339 | 343 | 769 | 812 | 699 | 788 |
| K05936 precorrin-4/cobalt-precorrin-4 C11-methyltransferase [EC:2.1.1.133 2.1.1.271] | 1988 | 1775 | 1744 | 1822 | 954 | 955 | 888 | 930 | 1301 | 1257 | 1141 | 1277 |
| K06042 precorrin-8X/cobalt-precorrin-8 methylmutase [EC:5.4.99.61 5.4.99.60] | 1366 | 1374 | 1266 | 1262 | 562 | 610 | 627 | 659 | 988 | 882 | 972 | 1090 |
| K07145 heme oxygenase (staphylobilin-producing) [EC:1.14.99.48] | 1 | 0 | 0 | 0 | 0 | 0 | 0 | 0 | 0 | 0 | 0 | 0 |
| K07215 heme oxygenase (biliverdin-IX-beta and delta-forming) [EC:1.14.99.58] | 684 | 709 | 666 | 685 | 327 | 384 | 342 | 342 | 247 | 227 | 192 | 196 |
| K08260 adenosylcobinamide hydrolase [EC:3.5.1.90] | 1 | 2 | 2 | 2 | 1 | 0 | 0 | 0 | 4 | 2 | 11 | 6 |
| K09882 cobaltochelatase CobS [EC:6.6.1.2] | 34 | 46 | 44 | 54 | 33 | 20 | 31 | 43 | 69 | 66 | 45 | 74 |
| K09883 cobaltochelatase CobT [EC:6.6.1.2] | 1 | 0 | 0 | 0 | 2 | 2 | 0 | 0 | 5 | 3 | 3 | 4 |
| K13541 cobalt-precorrin 5A hydrolase / precorrin-3B C17-methyltransferase [EC:3.7.1.12 2.1.1.131] | 1939 | 1966 | 1599 | 1746 | 881 | 948 | 956 | 1098 | 599 | 540 | 495 | 528 |
| K13542 uroporphyrinogen III methyltransferase / synthase [EC:2.1.1.107 4.2.1.75] | 678 | 632 | 641 | 639 | 131 | 162 | 167 | 132 | 708 | 756 | 724 | 866 |
| K13543 uroporphyrinogen III methyltransferase / synthase [EC:2.1.1.107 4.2.1.75] | 34 | 39 | 44 | 45 | 42 | 30 | 51 | 67 | 57 | 51 | 57 | 50 |
| K13786 cob(II)yrinic acid a,c-diamide reductase [EC:1.16.8.1] | 2 | 0 | 0 | 0 | 0 | 0 | 0 | 2 | 3 | 0 | 2 | 1 |
| K16651 L-threonine kinase [EC:2.7.1.177] | 593 | 507 | 552 | 534 | 379 | 351 | 383 | 398 | 736 | 731 | 658 | 786 |
| K19221 cob(I)alamin adenosyltransferase [EC:2.5.1.17] | 940 | 858 | 927 | 976 | 430 | 440 | 437 | 449 | 910 | 920 | 893 | 966 |
| K19712 adenosylcobinamide-phosphate guanylyltransferase [EC:2.7.7.62] | 0 | 2 | 2 | 1 | 0 | 0 | 0 | 0 | 2 | 2 | 3 | 1 |
| K21479 cobalt-factor III methyltransferase [EC:2.1.1.272] | 1 | 0 | 0 | 0 | 0 | 0 | 0 | 5 | 0 | 0 | 0 | 0 |
| K21480 heme oxygenase (biliverdin-producing, ferredoxin) [EC:1.14.15.20] | 1 | 0 | 0 | 0 | 0 | 6 | 0 | 1 | 0 | 0 | 0 | 0 |
| K21610 Ni-sirohydrochlorin a,c-diamide reductive cyclase subunit CfbC [EC:6.3.3.7] | 0 | 0 | 0 | 0 | 0 | 0 | 0 | 0 | 0 | 2 | 0 | 12 |
| K21611 Ni-sirohydrochlorin a,c-diamide reductive cyclase subunit CfbD [EC:6.3.3.7] | 2 | 0 | 0 | 0 | 0 | 2 | 0 | 0 | 3 | 3 | 3 | 9 |
| K21612 coenzyme F430 synthetase [EC:6.4.1.9] | 3 | 2 | 1 | 1 | 0 | 2 | 2 | 0 | 5 | 13 | 3 | 7 |
| K22011 sirohydrochlorin cobalto/nickelchelatase [EC:4.99.1.3 4.99.1.11] | 0 | 0 | 0 | 0 | 0 | 0 | 3 | 0 | 6 | 6 | 5 | 10 |
| K22012 Ni-sirohydrochlorin a,c-diamide synthase [EC:6.3.5.12] | 0 | 0 | 0 | 0 | 0 | 0 | 0 | 0 | 0 | 0 | 0 | 1 |
| K22226 Fe-coproporphyrin III synthase | 18 | 10 | 14 | 18 | 12 | 6 | 8 | 9 | 22 | 31 | 27 | 26 |
| K22227 heme synthase | 480 | 396 | 463 | 447 | 306 | 371 | 371 | 385 | 124 | 201 | 198 | 181 |
| K22316 ribonuclease H / adenosylcobalamin/alpha-ribazole phosphatase [EC:3.1.26.4 3.1.3.73] | 1 | 2 | 2 | 4 | 0 | 0 | 0 | 0 | 0 | 4 | 3 | 1 |
| ko00130 Ubiquinone and other terpenoid-quinone biosynthesis | 45775 | 46182 | 44534 | 45669 | 37405 | 38116 | 37464 | 37982 | 38645 | 38412 | 38247 | 38384 |
| K00355 NAD(P)H dehydrogenase (quinone) [EC:1.6.5.2] | 2683 | 2813 | 2569 | 2767 | 1666 | 1603 | 1531 | 1798 | 1381 | 1306 | 1406 | 1253 |
| K00568 2-polyprenyl-6-hydroxyphenyl methylase / 3-demethylubiquinone-9 3-methyltransferase [EC:2.1.1.222 2.1.1.64] | 1445 | 1476 | 1336 | 1398 | 860 | 859 | 847 | 802 | 868 | 968 | 954 | 937 |
| K01075 4-hydroxybenzoyl-CoA thioesterase [EC:3.1.2.23] | 527 | 565 | 585 | 501 | 338 | 317 | 295 | 337 | 133 | 180 | 160 | 180 |
| K01661 naphthoate synthase [EC:4.1.3.36] | 1142 | 1140 | 1181 | 1180 | 2133 | 2077 | 2150 | 2162 | 2048 | 2051 | 2008 | 1901 |
| K01851 salicylate biosynthesis isochorismate synthase [EC:5.4.4.2] | 2193 | 2185 | 2061 | 1980 | 1178 | 1199 | 1080 | 1193 | 779 | 777 | 587 | 649 |
| K01904 4-coumarate--CoA ligase [EC:6.2.1.12] | 1 | 0 | 0 | 0 | 0 | 1 | 0 | 0 | 0 | 0 | 0 | 0 |
| K01911 O-succinylbenzoic acid---CoA ligase [EC:6.2.1.26] | 1651 | 1572 | 1666 | 1659 | 2798 | 2936 | 3019 | 2972 | 2962 | 2813 | 2866 | 2796 |
| K02361 isochorismate synthase [EC:5.4.4.2] | 1345 | 1495 | 1415 | 1408 | 2784 | 3006 | 2892 | 2783 | 2615 | 2573 | 2648 | 2626 |
| K02548 1,4-dihydroxy-2-naphthoate polyprenyltransferase [EC:2.5.1.74] | 1290 | 1336 | 1345 | 1315 | 2460 | 2458 | 2675 | 2391 | 2221 | 2330 | 2334 | 2267 |
| K02549 O-succinylbenzoate synthase [EC:4.2.1.113] | 736 | 716 | 638 | 736 | 429 | 446 | 419 | 443 | 910 | 981 | 924 | 960 |
| K02551 2-succinyl-5-enolpyruvyl-6-hydroxy-3-cyclohexene-1-carboxylate synthase [EC:2.2.1.9] | 2671 | 2693 | 2629 | 2639 | 4076 | 4220 | 4122 | 4083 | 4170 | 4050 | 4277 | 4098 |
| K02552 menaquinone-specific isochorismate synthase [EC:5.4.4.2] | 860 | 860 | 803 | 841 | 481 | 524 | 516 | 543 | 1116 | 1202 | 1125 | 1156 |
| K03179 4-hydroxybenzoate polyprenyltransferase [EC:2.5.1.39] | 2082 | 2081 | 2030 | 2079 | 1232 | 1303 | 1197 | 1260 | 1256 | 1232 | 1148 | 1187 |
| K03181 chorismate--pyruvate lyase [EC:4.1.3.40] | 1232 | 1303 | 1216 | 1406 | 749 | 834 | 776 | 899 | 901 | 851 | 829 | 810 |
| K03182 4-hydroxy-3-polyprenylbenzoate decarboxylase [EC:4.1.1.98] | 6185 | 6218 | 6215 | 6471 | 3644 | 3761 | 3422 | 3624 | 3197 | 3151 | 3203 | 3226 |
| K03183 demethylmenaquinone methyltransferase / 2-methoxy-6-polyprenyl-1,4-benzoquinol methylase [EC:2.1.1.163 2.1.1.201] | 2175 | 2244 | 2323 | 2195 | 2667 | 2669 | 2821 | 2595 | 2381 | 2414 | 2521 | 2380 |
| K03184 3-demethoxyubiquinol 3-hydroxylase [EC:1.14.99.60] | 786 | 769 | 723 | 714 | 483 | 443 | 488 | 486 | 1071 | 1096 | 981 | 1065 |
| K03185 2-octaprenyl-6-methoxyphenol hydroxylase [EC:1.14.13.-] | 2494 | 2664 | 2552 | 2625 | 1547 | 1583 | 1496 | 1534 | 1578 | 1548 | 1495 | 1661 |
| K03186 flavin prenyltransferase [EC:2.5.1.129] | 1990 | 1929 | 1674 | 1811 | 1097 | 1146 | 1155 | 1146 | 1469 | 1430 | 1460 | 1401 |
| K03809 NAD(P)H dehydrogenase (quinone) [EC:1.6.5.2] | 1252 | 1300 | 1317 | 1289 | 760 | 721 | 804 | 699 | 726 | 724 | 718 | 822 |

| | | | | | | | | | | | | |
|---|---|---|---|---|---|---|---|---|---|---|---|---|
| K06127 2-methoxy-6-polyprenyl-1,4-benzoquinol methylase [EC:2.1.1.201] | 1 | 0 | 0 | 0 | 2 | 0 | 1 | 3 | 0 | 0 | 0 | 0 |
| K06134 3-demethoxyubiquinol 3-hydroxylase [EC:1.14.99.60] | 770 | 762 | 781 | 739 | 454 | 457 | 470 | 411 | 257 | 249 | 259 | 219 |
| K08680 2-succinyl-6-hydroxy-2,4-cyclohexadiene-1-carboxylate synthase [EC:4.2.99.20] | 520 | 525 | 477 | 506 | 284 | 302 | 311 | 302 | 632 | 697 | 627 | 674 |
| K11782 chorismate dehydratase [EC:4.2.1.151] | 9 | 8 | 5 | 5 | 7 | 11 | 3 | 6 | 5 | 5 | 5 | 7 |
| K11783 futalosine hydrolase [EC:3.2.2.26] | 2 | 3 | 2 | 1 | 3 | 4 | 3 | 0 | 7 | 7 | 4 | 8 |
| K11784 cyclic dehypoxanthinyl futalosine synthase [EC:1.21.98.1] | 7 | 2 | 10 | 8 | 5 | 18 | 8 | 2 | 10 | 15 | 6 | 14 |
| K11785 1,4-dihydroxy-6-naphthoate synthase [EC:1.14.-.-] | 3 | 5 | 9 | 3 | 3 | 10 | 5 | 13 | 18 | 11 | 8 | 11 |
| K18285 aminodeoxyfutalosine synthase [EC:2.5.1.120] | 19 | 5 | 9 | 9 | 3 | 11 | 9 | 7 | 12 | 19 | 13 | 15 |
| K18286 aminodeoxyfutalosine deaminase [EC:3.5.4.40] | 1 | 2 | 2 | 1 | 0 | 0 | 0 | 0 | 0 | 0 | 0 | 0 |
| K18800 2-polyprenylphenol 6-hydroxylase [EC:1.14.13.240] | 3061 | 3005 | 2822 | 2994 | 1750 | 1682 | 1716 | 1760 | 1871 | 1709 | 1746 | 1833 |
| K19222 1,4-dihydroxy-2-naphthoyl-CoA hydrolase [EC:3.1.2.28] | 884 | 927 | 750 | 855 | 504 | 472 | 466 | 523 | 539 | 492 | 490 | 534 |
| K19267 NAD(P)H dehydrogenase (quinone) [EC:1.6.5.2] | 481 | 519 | 494 | 596 | 333 | 434 | 443 | 396 | 752 | 814 | 697 | 796 |
| K20810 aminodeoxyfutalosine deaminase [EC:3.5.4.40] | 1 | 0 | 0 | 0 | 1 | 1 | 0 | 0 | 0 | 0 | 0 | 1 |
| Metabolism of terpenoids and polyketides | 154125 | 153253 | 148955 | 151067 | 124182 | 123608 | 122819 | 122769 | 130332 | 129860 | 127856 | 130163 |
| ko00900 Terpenoid backbone biosynthesis | 46655 | 47022 | 46688 | 46804 | 47838 | 47169 | 47643 | 47051 | 44876 | 45329 | 45135 | 44698 |
| K00021 hydroxymethylglutaryl-CoA reductase (NADPH) [EC:1.1.1.34] | 0 | 0 | 0 | 0 | 0 | 0 | 0 | 0 | 2 | 10 | 3 | 9 |
| K00054 hydroxymethylglutaryl-CoA reductase [EC:1.1.1.88] | 63 | 55 | 59 | 51 | 8 | 18 | 5 | 7 | 64 | 75 | 75 | 66 |
| K00099 1-deoxy-D-xylulose-5-phosphate reductoisomerase [EC:1.1.1.267] | 3978 | 4155 | 4145 | 4129 | 4369 | 4365 | 4375 | 4273 | 4072 | 4161 | 4110 | 4079 |
| K00795 farnesyl diphosphate synthase [EC:2.5.1.1 2.5.1.10] | 584 | 611 | 556 | 619 | 505 | 469 | 444 | 417 | 816 | 909 | 835 | 873 |
| K00805 heptaprenyl diphosphate synthase [EC:2.5.1.30] | 351 | 403 | 389 | 348 | 107 | 85 | 69 | 98 | 330 | 370 | 427 | 397 |
| K00806 undecaprenyl diphosphate synthase [EC:2.5.1.31] | 2035 | 2016 | 2098 | 2092 | 2578 | 2639 | 2695 | 2553 | 2507 | 2686 | 2570 | 2686 |
| K00869 mevalonate kinase [EC:2.7.1.36] | 75 | 49 | 52 | 58 | 26 | 16 | 15 | 10 | 53 | 94 | 76 | 68 |
| K00919 4-diphosphocytidyl-2-C-methyl-D-erythritol kinase [EC:2.7.1.148] | 2604 | 2616 | 2691 | 2541 | 2838 | 2750 | 2710 | 2558 | 2734 | 2690 | 2845 | 2647 |
| K00938 phosphomevalonate kinase [EC:2.7.4.2] | 62 | 67 | 70 | 68 | 24 | 16 | 19 | 18 | 82 | 104 | 75 | 87 |
| K00991 2-C-methyl-D-erythritol 4-phosphate cytidylyltransferase [EC:2.7.7.60] | 2549 | 2397 | 2531 | 2609 | 2428 | 2371 | 2597 | 2526 | 2422 | 2523 | 2344 | 2422 |
| K01597 diphosphomevalonate decarboxylase [EC:4.1.1.33] | 49 | 79 | 58 | 61 | 17 | 10 | 13 | 16 | 52 | 78 | 73 | 73 |
| K01770 2-C-methyl-D-erythritol 2,4-cyclodiphosphate synthase [EC:4.6.1.12] | 1547 | 1699 | 1652 | 1685 | 1785 | 1777 | 1782 | 1705 | 1619 | 1705 | 1652 | 1719 |
| K01823 isopentenyl-diphosphate Delta-isomerase [EC:5.3.3.2] | 436 | 490 | 434 | 506 | 261 | 294 | 302 | 286 | 631 | 695 | 621 | 606 |
| K02523 octaprenyl-diphosphate synthase [EC:2.5.1.90] | 2606 | 2446 | 2653 | 2517 | 3389 | 3610 | 3615 | 3531 | 3009 | 2964 | 3020 | 2948 |
| K03526 (E)-4-hydroxy-3-methylbut-2-enyl-diphosphate synthase [EC:1.17.7.1 1.17.7.3] | 4251 | 4405 | 4447 | 4420 | 6126 | 6050 | 6161 | 5977 | 5224 | 4958 | 5337 | 4994 |
| K03527 4-hydroxy-3-methylbut-2-en-1-yl diphosphate reductase [EC:1.17.7.4] | 2847 | 2824 | 2873 | 2679 | 3256 | 3123 | 3159 | 3158 | 3313 | 3252 | 3240 | 3215 |
| K06013 STE24 endopeptidase [EC:3.4.24.84] | 9 | 4 | 2 | 3 | 9 | 9 | 11 | 7 | 4 | 5 | 10 | 13 |
| K06981 isopentenyl phosphate kinase [EC:2.7.4.26] | 0 | 3 | 3 | 0 | 1 | 0 | 0 | 1 | 2 | 6 | 8 | 4 |
| K12506 2-C-methyl-D-erythritol 4-phosphate cytidylyltransferase / 2-C-methyl-D-erythritol 2,4-cyclodiphosphate synthase [EC:2.7.7.60 4.6.1.12] | 63 | 60 | 63 | 70 | 62 | 72 | 67 | 99 | 90 | 77 | 85 | 77 |
| K13787 geranylgeranyl diphosphate synthase, type I [EC:2.5.1.1 2.5.1.10 2.5.1.29] | 11 | 9 | 11 | 16 | 10 | 7 | 7 | 6 | 31 | 20 | 21 | 23 |
| K13789 geranylgeranyl diphosphate synthase, type II [EC:2.5.1.1 2.5.1.10 2.5.1.29] | 2253 | 2148 | 2200 | 2235 | 2812 | 2752 | 2832 | 2709 | 2146 | 2176 | 2251 | 2226 |
| K15888 tritrans,polycis-undecaprenyl-diphosphate synthase [geranylgeranyl-diphosphate specific] [EC:2.5.1.89] | 1 | 1 | 0 | 0 | 2 | 2 | 0 | 0 | 5 | 3 | 3 | 10 |
| K21273 trans,polycis-polyprenyl diphosphate synthase [EC:2.5.1.88] | 1 | 0 | 1 | 0 | 0 | 1 | 0 | 0 | 0 | 0 | 0 | 0 |
| ko00902 Monoterpenoid biosynthesis | 1 | 0 | 0 | 0 | 0 | 0 | 0 | 0 | 0 | 1 | 0 | 0 |
| K15095 (+)-neomenthol dehydrogenase [EC:1.1.1.208] | 1 | 0 | 0 | 0 | 0 | 0 | 0 | 0 | 0 | 0 | 0 | 0 |
| K23232 8-hydroxygeraniol dehydrogenase [EC:1.1.1.324] | 0 | 0 | 0 | 0 | 0 | 0 | 0 | 0 | 0 | 1 | 0 | 0 |
| ko00909 Sesquiterpenoid and triterpenoid biosynthesis | 3 | 0 | 2 | 3 | 0 | 0 | 0 | 0 | 0 | 0 | 0 | 0 |
| K06045 squalene-hopene/tetraprenyl-beta-curcumene cyclase [EC:5.4.99.17 4.2.1.129] | 3 | 0 | 2 | 3 | 0 | 0 | 0 | 0 | 0 | 0 | 0 | 0 |
| ko00906 Carotenoid biosynthesis | 29 | 53 | 60 | 56 | 43 | 55 | 56 | 50 | 125 | 101 | 88 | 98 |
| K02291 15-cis-phytoene synthase [EC:2.5.1.32] | 10 | 16 | 12 | 25 | 14 | 20 | 18 | 20 | 42 | 37 | 26 | 34 |
| K09835 prolycopene isomerase [EC:5.2.1.13] | 1 | 0 | 2 | 2 | 0 | 0 | 0 | 0 | 1 | 1 | 3 | 1 |
| K10027 phytoene desaturase [EC:1.3.99.26 1.3.99.28 1.3.99.29 1.3.99.31] | 14 | 24 | 35 | 25 | 24 | 35 | 32 | 27 | 72 | 53 | 55 | 54 |
| K10210 diapolycopene oxygenase [EC:1.14.99.44] | 2 | 6 | 5 | 3 | 0 | 0 | 6 | 3 | 6 | 2 | 2 | 6 |
| K10211 4,4'-diaponeurosporenoate glycosyltransferase [EC:2.4.1.-] | 0 | 5 | 4 | 1 | 0 | 0 | 0 | 0 | 4 | 0 | 2 | 2 |
| K10212 glycosyl-4,4'-diaponeurosporenoate acyltransferase [EC:2.3.1.-] | 2 | 2 | 2 | 0 | 5 | 0 | 0 | 0 | 0 | 4 | 0 | 1 |
| K15746 beta-carotene 3-hydroxylase [EC:1.14.15.24] | 0 | 0 | 0 | 0 | 0 | 0 | 0 | 0 | 0 | 4 | 0 | 0 |
| ko00981 Insect hormone biosynthesis | 5102 | 5084 | 4807 | 4818 | 3106 | 2867 | 2785 | 3017 | 1916 | 1755 | 1686 | 1825 |
| K14938 cholesterol 7-desaturase [EC:1.14.19.21] | 0 | 0 | 0 | 0 | 0 | 0 | 0 | 0 | 0 | 1 | 0 | 0 |
| ko00908 Zeatin biosynthesis | 3123 | 3207 | 3387 | 3217 | 5160 | 5000 | 5206 | 4962 | 4225 | 4307 | 4377 | 4260 |
| K00791 tRNA dimethylallyltransferase [EC:2.5.1.75] | 3123 | 3207 | 3387 | 3217 | 5160 | 5000 | 5206 | 4962 | 4225 | 4307 | 4377 | 4260 |
| ko00903 Limonene and pinene degradation | 16698 | 16373 | 16025 | 16123 | 9818 | 9523 | 9105 | 9778 | 8914 | 8576 | 8466 | 9038 |
| K14731 epsilon-lactone hydrolase [EC:3.1.1.83] | 92 | 97 | 120 | 106 | 24 | 27 | 52 | 33 | 118 | 121 | 151 | 167 |
| ko00281 Geraniol degradation | 32077 | 31388 | 30074 | 30983 | 17949 | 17595 | 16844 | 17810 | 15019 | 14526 | 14201 | 14848 |
| K11731 citronellyl-CoA dehydrogenase [EC:1.3.99.-] | 1806 | 1704 | 1660 | 1740 | 939 | 903 | 941 | 923 | 511 | 506 | 498 | 480 |
| K13774 citronellol/citronellal dehydrogenase | 849 | 907 | 840 | 831 | 449 | 471 | 408 | 457 | 280 | 260 | 252 | 275 |
| K13775 citronellol/citronellal dehydrogenase | 1202 | 1233 | 1030 | 1152 | 659 | 603 | 543 | 642 | 388 | 373 | 421 | 352 |
| K13776 citronellyl-CoA synthetase [EC:6.2.1.-] | 2534 | 2508 | 2308 | 2469 | 1315 | 1348 | 1220 | 1343 | 794 | 764 | 691 | 764 |
| K13777 geranyl-CoA carboxylase alpha subunit [EC:6.4.1.5] | 2904 | 2884 | 2601 | 2663 | 1520 | 1451 | 1323 | 1417 | 892 | 864 | 787 | 827 |
| K13778 geranyl-CoA carboxylase beta subunit [EC:6.4.1.5] | 2360 | 2282 | 2336 | 2284 | 1279 | 1250 | 1202 | 1241 | 726 | 681 | 668 | 638 |
| K13779 isohexenylglutaconyl-CoA hydratase [EC:4.2.1.57] | 818 | 731 | 646 | 765 | 407 | 391 | 378 | 481 | 286 | 228 | 200 | 254 |
| ko01052 Type I polyketide structures | 0 | 1 | 1 | 0 | 1 | 1 | 2 | 5 | 7 | 7 | 8 | 2 |
| K16378 pipecolate-incorporating enzyme | 0 | 0 | 0 | 0 | 0 | 1 | 2 | 3 | 4 | 6 | 8 | 1 |
| K16384 polyene macrolide polyketide synthase | 0 | 0 | 0 | 0 | 1 | 0 | 0 | 1 | 2 | 0 | 0 | 1 |
| K16395 epothilone synthetase B | 0 | 1 | 1 | 0 | 0 | 0 | 0 | 0 | 1 | 1 | 0 | 0 |
| K16397 epothilone polyketide synthase D | 0 | 0 | 0 | 0 | 0 | 0 | 0 | 1 | 0 | 0 | 0 | 0 |
| ko01051 Biosynthesis of ansamycins | 13517 | 13414 | 13461 | 13442 | 9604 | 9691 | 9778 | 9303 | 14442 | 14290 | 14105 | 14499 |
| ko01059 Biosynthesis of enediyne antibiotics | 0 | 0 | 0 | 0 | 0 | 0 | 0 | 0 | 0 | 0 | 0 | 1 |
| K15320 6-methylsalicylic acid synthase [EC:2.3.1.165] | 0 | 0 | 0 | 0 | 0 | 0 | 0 | 0 | 0 | 0 | 0 | 1 |
| ko01056 Biosynthesis of type II polyketide backbone | 2 | 1 | 0 | 0 | 0 | 0 | 0 | 0 | 2 | 1 | 0 | 1 |
| K12420 ketoreductase [EC:1.1.1.-] | 2 | 1 | 0 | 0 | 0 | 0 | 0 | 0 | 2 | 1 | 0 | 1 |
| ko01057 Biosynthesis of type II polyketide products | 0 | 0 | 0 | 0 | 0 | 0 | 0 | 0 | 0 | 1 | 0 | 0 |
| K05556 ketoreductase RED1 [EC:1.1.1.-] | 0 | 0 | 0 | 0 | 0 | 0 | 0 | 0 | 0 | 1 | 0 | 0 |
| ko00253 Tetracycline biosynthesis | 857 | 898 | 977 | 897 | 3124 | 3092 | 3110 | 3055 | 2340 | 2086 | 2325 | 2189 |
| K18221 tetracycline 11a-monooxygenase, tetracycline resistance protein [EC:1.14.13.231] | 855 | 897 | 977 | 897 | 3124 | 3092 | 3110 | 3055 | 2338 | 2085 | 2325 | 2188 |
| ko00523 Polyketide sugar unit biosynthesis | 14902 | 15109 | 15013 | 14959 | 16606 | 17024 | 17232 | 16617 | 13995 | 14146 | 13887 | 13846 |
| K00067 dTDP-4-dehydrorhamnose reductase [EC:1.1.1.133] | 4197 | 4324 | 4281 | 4273 | 4572 | 4728 | 4685 | 4483 | 3463 | 3571 | 3423 | 3316 |
| K00973 glucose-1-phosphate thymidylyltransferase [EC:2.7.7.24] | 4335 | 4362 | 4461 | 4283 | 5425 | 5406 | 5669 | 5509 | 4932 | 4888 | 4972 | 4998 |
| K01710 dTDP-glucose 4,6-dehydratase [EC:4.2.1.46] | 4263 | 4310 | 4172 | 4262 | 4525 | 4735 | 4663 | 4518 | 3965 | 4125 | 3926 | 3887 |
| K01790 dTDP-4-dehydrorhamnose 3,5-epimerase [EC:5.1.3.13] | 2099 | 2103 | 2089 | 2123 | 2084 | 2149 | 2213 | 2106 | 1619 | 1546 | 1541 | 1629 |
| K13315 NDP-hexose C3-ketoreductase / dTDP-4-oxo-2-deoxy-alpha-D-pentos-2-ene 2,3-reductase [EC:1.1.1.-] | 6 | 5 | 6 | 10 | 0 | 0 | 0 | 0 | 3 | 4 | 3 | 5 |
| K13327 dTDP-3,4-didehydro-2,6-dideoxy-alpha-D-glucose 3-reductase [EC:1.1.1.384] | 0 | 0 | 0 | 3 | 0 | 0 | 0 | 0 | 3 | 0 | 2 | 0 |
| K13328 dTDP-4-dehydro-2,6-dideoxy-D-glucose 3-dehydratase [EC:4.2.1.164] | 0 | 0 | 0 | 0 | 0 | 0 | 0 | 1 | 0 | 0 | 0 | 0 |
| K16435 dTDP-4-dehydro-6-deoxy-alpha-D-glucopyranose 2,3-dehydratase [EC:4.2.1.159] | 0 | 3 | 2 | 3 | 0 | 4 | 0 | 0 | 3 | 5 | 8 | 3 |
| K16436 dTDP-3-amino-2,3,6-trideoxy-4-keto-D-glucose/dTDP-3-amino-3,4,6-trideoxy-alpha-D-glucose/dTDP-2,6-dideoxy-D-kanosamine transaminase [EC:2.6 | 0 | 2 | 0 | 2 | 0 | 2 | 0 | 0 | 0 | 0 | 0 | 1 |
| K23144 UDP-N-acetylglucosamine diphosphorylase / glucose-1-phosphate thymidylyltransferase / UDP-N-acetylgalactosamine diphosphorylase / glucosamine | 2 | 0 | 2 | 0 | 0 | 0 | 2 | 0 | 7 | 7 | 12 | 7 |
| ko01054 Nonribosomal peptide structures | 545 | 573 | 494 | 584 | 430 | 483 | 410 | 398 | 391 | 459 | 418 | 429 |
| K15654 surfactin family lipopeptide synthetase A | 1 | 1 | 1 | 0 | 0 | 1 | 0 | 0 | 2 | 1 | 0 | 3 |
| K15655 surfactin family lipopeptide synthetase B | 0 | 0 | 0 | 0 | 0 | 0 | 0 | 0 | 2 | 1 | 0 | 1 |
| K15656 surfactin family lipopeptide synthetase C | 3 | 0 | 0 | 1 | 0 | 0 | 0 | 0 | 0 | 0 | 0 | 0 |
| K15662 iturin family lipopeptide synthetase B | 2 | 2 | 2 | 3 | 1 | 0 | 0 | 0 | 1 | 5 | 3 | 3 |
| K15664 fengycin family lipopeptide synthetase A | 0 | 0 | 0 | 0 | 0 | 0 | 0 | 0 | 0 | 1 | 0 | 0 |
| K15665 fengycin family lipopeptide synthetase B | 0 | 1 | 0 | 1 | 0 | 0 | 0 | 0 | 1 | 2 | 1 | 1 |
| K15667 fengycin family lipopeptide synthetase D | 1 | 0 | 0 | 0 | 0 | 0 | 0 | 0 | 1 | 0 | 0 | 0 |
| K15668 fengycin family lipopeptide synthetase E | 0 | 0 | 0 | 0 | 0 | 0 | 0 | 0 | 0 | 0 | 0 | 1 |
| K16093 bacitracin synthase 1 | 1 | 2 | 1 | 1 | 1 | 0 | 0 | 0 | 4 | 5 | 1 | 3 |
| K16094 bacitracin synthase 2 | 3 | 0 | 2 | 2 | 2 | 0 | 1 | 0 | 1 | 1 | 0 | 1 |
| K16095 bacitracin synthase 3 | 3 | 2 | 2 | 2 | 0 | 0 | 0 | 0 | 4 | 3 | 2 | 3 |
| K16119 lichenysin synthetase A | 0 | 0 | 0 | 0 | 0 | 0 | 0 | 0 | 1 | 3 | 2 | 1 |
| K16122 tyrocidine synthetase I | 0 | 0 | 1 | 0 | 0 | 0 | 0 | 0 | 0 | 0 | 0 | 0 |
| K16124 tyrocidine synthetase III | 1 | 0 | 2 | 0 | 0 | 0 | 0 | 0 | 0 | 0 | 0 | 0 |
| K16129 microcystin synthetase protein McyE | 0 | 0 | 0 | 0 | 0 | 0 | 0 | 0 | 0 | 1 | 0 | 0 |
| K16133 microcystin synthetase protein McyI | 0 | 0 | 0 | 0 | 0 | 0 | 0 | 0 | 0 | 2 | 1 | 0 |
| ko01053 Biosynthesis of siderophore group nonribosomal peptides | 37222 | 36407 | 33871 | 35198 | 20296 | 20603 | 19701 | 20468 | 32877 | 32731 | 31475 | 33301 |
| K00216 2,3-dihydro-2,3-dihydroxybenzoate dehydrogenase [EC:1.3.1.28] | 458 | 465 | 427 | 532 | 313 | 313 | 384 | 333 | 630 | 625 | 643 | 696 |
| K01252 bifunctional isochorismate lyase / aryl carrier protein [EC:3.3.2.1 6.3.2.14] | 599 | 509 | 515 | 536 | 359 | 338 | 413 | 351 | 703 | 748 | 688 | 789 |
| K02362 enterobactin synthetase component D [EC:6.3.2.14 2.7.8.-] | 603 | 666 | 545 | 609 | 402 | 415 | 355 | 378 | 581 | 599 | 561 | 601 |
| K02363 2,3-dihydroxybenzoate-AMP ligase [EC:6.3.2.14 2.7.7.58] | 976 | 1012 | 803 | 910 | 602 | 722 | 697 | 582 | 1354 | 1379 | 1363 | 1418 |
| K02364 enterobactin synthetase component F [EC:6.3.2.14] | 2262 | 2096 | 1927 | 2212 | 1656 | 1688 | 1535 | 1523 | 2661 | 2674 | 2612 | 2816 |
| K04780 nonribosomal peptide synthetase DhbF | 0 | 0 | 0 | 0 | 1 | 0 | 0 | 0 | 0 | 0 | 0 | 0 |
| K04781 salicylate synthetase [EC:5.4.4.2 4.2.99.21] | 284 | 257 | 287 | 329 | 84 | 100 | 94 | 82 | 645 | 713 | 627 | 772 |
| K04782 isochorismate pyruvate lyase [EC:4.2.99.21] | 540 | 528 | 501 | 490 | 286 | 267 | 260 | 295 | 177 | 137 | 153 | 159 |
| K04783 yersiniabactin salicyl-AMP ligase [EC:6.3.2.-] | 5668 | 5505 | 5691 | 5563 | 3100 | 3069 | 2981 | 2944 | 3076 | 2930 | 2845 | 2787 |
| K04784 yersiniabactin nonribosomal peptide synthetase | 3308 | 3413 | 3295 | 3564 | 1120 | 1147 | 1135 | 1079 | 5606 | 5762 | 5451 | 5755 |
| K04785 yersiniabactin synthetase, thiazolinyl reductase component | 452 | 463 | 391 | 429 | 116 | 135 | 106 | 108 | 938 | 904 | 967 | 931 |
| K04786 yersiniabactin nonribosomal peptide/polyketide synthase | 3662 | 3508 | 3359 | 3641 | 969 | 1039 | 967 | 867 | 7355 | 7636 | 7382 | 7909 |
| K04787 mycobactin salicyl-AMP ligase [EC:6.3.2.-] | 0 | 0 | 0 | 0 | 0 | 0 | 0 | 1 | 1 | 0 | 0 | 0 |

| | | | | | | | | | | | | |
|---|---|---|---|---|---|---|---|---|---|---|---|---|
| K04788 mycobactin phenyloxazoline synthetase | 56 | 51 | 54 | 40 | 18 | 23 | 45 | 36 | 10 | 11 | 7 | 13 |
| K04792 mycobactin peptide synthetase MbtF | 0 | 0 | 0 | 0 | 0 | 0 | 0 | 0 | 0 | 0 | 0 | 1 |
| K12238 pyochelin biosynthesis protein PchD | 2623 | 2311 | 2267 | 2387 | 1293 | 1242 | 1170 | 1312 | 802 | 711 | 660 | 743 |
| K12239 dihydroaeruginoic acid synthetase | 2557 | 2576 | 2172 | 2273 | 1306 | 1227 | 1234 | 1445 | 850 | 675 | 692 | 790 |
| K12240 pyochelin synthetase | 7108 | 6939 | 6050 | 6098 | 3437 | 3387 | 3113 | 3701 | 2465 | 2153 | 2021 | 2271 |
| K12241 pyochelin biosynthetic protein PchG | 1668 | 1568 | 1308 | 1356 | 791 | 762 | 724 | 912 | 513 | 522 | 443 | 419 |
| ko01055 Biosynthesis of vancomycin group antibiotics | 4263 | 4315 | 4174 | 4267 | 4526 | 4742 | 4663 | 4518 | 3969 | 4130 | 3934 | 3891 |
| K16428 nonribosomal peptide synthetase CepA | 0 | 0 | 0 | 0 | 1 | 1 | 0 | 0 | 1 | 0 | 0 | 0 |
| Biosynthesis of other secondary metabolites | 190903 | 190852 | 192732 | 191796 | 233503 | 232484 | 235180 | 231851 | 217484 | 215139 | 217005 | 214577 |
| ko00940 Phenylpropanoid biosynthesis | 32196 | 33340 | 34471 | 33540 | 71230 | 70199 | 72603 | 70003 | 65143 | 64596 | 66190 | 63984 |
| K12355 coniferyl-aldehyde dehydrogenase [EC:1.2.1.68] | 0 | 0 | 0 | 0 | 0 | 1 | 0 | 1 | 0 | 0 | 0 | 0 |
| K20039 phenacrylate decarboxylase [EC:4.1.1.102] | 1 | 0 | 0 | 0 | 0 | 0 | 0 | 2 | 0 | 0 | 0 | 0 |
| ko00944 Flavone and flavonol biosynthesis | 2965 | 2979 | 3153 | 3092 | 7622 | 7568 | 7800 | 7477 | 7097 | 6746 | 7115 | 6902 |
| ko00901 Indole alkaloid biosynthesis | 0 | 1 | 0 | 0 | 0 | 0 | 0 | 0 | 0 | 0 | 0 | 0 |
| ko00950 Isoquinoline alkaloid biosynthesis | 10701 | 10860 | 10839 | 10806 | 10253 | 10109 | 10199 | 10350 | 10541 | 10532 | 10374 | 10532 |
| ko00960 Tropane, piperidine and pyridine alkaloid biosynthesis | 22018 | 22019 | 22162 | 22222 | 20537 | 20411 | 20343 | 20144 | 21829 | 21745 | 21503 | 21807 |
| K00808 homospermidine synthase [EC:2.5.1.44] | 1853 | 1903 | 2132 | 2044 | 1112 | 1128 | 1049 | 1047 | 632 | 622 | 648 | 573 |
| K08081 tropinone reductase I [EC:1.1.1.206] | 1 | 0 | 0 | 0 | 0 | 0 | 0 | 0 | 0 | 0 | 0 | 0 |
| K13607 cinnamoyl-CoA:phenyllactate CoA-transferase [EC:2.8.3.17] | 9 | 5 | 5 | 9 | 1 | 5 | 0 | 0 | 4 | 9 | 5 | 7 |
| K19064 lysine 6-dehydrogenase [EC:1.4.1.18] | 0 | 0 | 0 | 4 | 0 | 0 | 0 | 1 | 4 | 3 | 2 | 4 |
| ko00232 Caffeine metabolism | 0 | 2 | 0 | 0 | 0 | 4 | 0 | 0 | 0 | 0 | 0 | 0 |
| ko00965 Betalain biosynthesis | 1718 | 1703 | 1494 | 1617 | 930 | 975 | 830 | 922 | 1039 | 1035 | 1005 | 963 |
| K15777 4,5-DOPA dioxygenase extradiol [EC:1.13.11.-] | 1718 | 1702 | 1494 | 1617 | 930 | 975 | 830 | 922 | 1039 | 1035 | 1005 | 963 |
| ko00966 Glucosinolate biosynthesis | 4521 | 4354 | 4339 | 4552 | 4914 | 5049 | 4994 | 4747 | 4882 | 4859 | 4710 | 4855 |
| ko00311 Penicillin and cephalosporin biosynthesis | 8866 | 8634 | 8627 | 8540 | 7640 | 7633 | 7507 | 7599 | 5580 | 5068 | 5180 | 5221 |
| K01060 cephalosporin-C deacetylase [EC:3.1.1.41] | 124 | 150 | 136 | 154 | 88 | 95 | 70 | 91 | 66 | 62 | 64 | 63 |
| K01434 penicillin amidase [EC:3.5.1.11] | 7230 | 6914 | 6660 | 6777 | 3698 | 3620 | 3556 | 3726 | 2261 | 1970 | 1884 | 2099 |
| K17836 beta-lactamase class A [EC:3.5.2.6] | 1501 | 1561 | 1824 | 1604 | 3845 | 3906 | 3872 | 3772 | 3243 | 3030 | 3224 | 3054 |
| K19200 isopenicillin-N N-acyltransferase like protein | 11 | 9 | 7 | 5 | 9 | 12 | 9 | 10 | 10 | 6 | 8 | 5 |
| ko00332 Carbapenem biosynthesis | 6736 | 6670 | 6770 | 6805 | 8230 | 7978 | 8443 | 8404 | 7958 | 7776 | 8258 | 8066 |
| ko00261 Monobactam biosynthesis | 30837 | 30802 | 31277 | 31199 | 35977 | 36067 | 36221 | 35656 | 34760 | 34734 | 34737 | 34241 |
| K05375 MbtH protein | 314 | 276 | 266 | 317 | 152 | 125 | 131 | 157 | 111 | 96 | 80 | 106 |
| ko00521 Streptomycin biosynthesis | 33159 | 33370 | 33617 | 33510 | 39818 | 40498 | 40907 | 40356 | 38527 | 38516 | 38430 | 38505 |
| ko00524 Neomycin, kanamycin and gentamicin biosynthesis | 5641 | 5770 | 5818 | 5761 | 10192 | 10337 | 10553 | 10631 | 9141 | 9227 | 9378 | 9288 |
| ko00525 Acarbose and validamycin biosynthesis | 8600 | 8672 | 8635 | 8545 | 9951 | 10141 | 10334 | 10028 | 8904 | 9020 | 8913 | 8892 |
| K19974 2-epi-5-epi-valiolone 7-phosphate 2-epimerase [EC:5.1.3.35] | 0 | 0 | 0 | 0 | 0 | 0 | 0 | 1 | 0 | 0 | 0 | 0 |
| K20437 validoxylamine A 7'-phosphate phosphatase [EC:3.1.3.101] | 0 | 0 | 0 | 0 | 1 | 0 | 0 | 0 | 0 | 0 | 3 | 0 |
| ko00401 Novobiocin biosynthesis | 16959 | 16987 | 17235 | 17013 | 19343 | 19151 | 19422 | 19056 | 17369 | 17133 | 17252 | 17320 |
| ko00404 Staurosporine biosynthesis | 1 | 0 | 0 | 2 | 0 | 0 | 0 | 0 | 0 | 0 | 0 | 0 |
| K14266 tryptophan 7-halogenase [EC:1.14.19.9] | 1 | 0 | 0 | 2 | 0 | 0 | 0 | 0 | 0 | 0 | 0 | 0 |
| ko00405 Phenazine biosynthesis | 23666 | 23152 | 22957 | 22726 | 16136 | 15901 | 15484 | 16081 | 11436 | 10981 | 10783 | 10955 |
| K06998 trans-2,3-dihydro-3-hydroxyanthranilate isomerase [EC:5.3.3.17] | 208 | 153 | 160 | 144 | 257 | 276 | 265 | 225 | 4 | 7 | 5 | 3 |
| K13063 2-amino-4-deoxychorismate synthase [EC:2.6.1.86] | 2265 | 2135 | 2010 | 2043 | 1114 | 1129 | 1062 | 1184 | 653 | 703 | 650 | 646 |
| K17940 2-heptyl-3-hydroxy-4(1H)-quinolone synthase [EC:1.14.13.182] | 566 | 565 | 591 | 550 | 336 | 273 | 287 | 293 | 167 | 157 | 180 | 167 |
| K18000 anthranilate-CoA ligase [EC:6.2.1.32] | 2371 | 2396 | 2409 | 2415 | 1474 | 1327 | 1240 | 1360 | 770 | 690 | 666 | 807 |
| K18001 2-heptyl-4(1H)-quinolone synthase subunit PqsB [EC:2.3.1.230] | 1422 | 1379 | 1447 | 1388 | 807 | 804 | 765 | 850 | 463 | 410 | 445 | 413 |
| K18002 2-heptyl-4(1H)-quinolone synthase subunit PqsC [EC:2.3.1.230] | 1481 | 1476 | 1555 | 1402 | 864 | 810 | 828 | 821 | 478 | 463 | 453 | 426 |
| K18003 anthraniloyl-CoA anthraniloyltransferase [EC:2.3.1.262] | 1711 | 1526 | 1560 | 1526 | 918 | 868 | 870 | 885 | 515 | 514 | 449 | 474 |
| K20257 2-aminobenzoylacetyl-CoA thioesterase [EC:3.1.2.32] | 272 | 265 | 248 | 260 | 136 | 135 | 149 | 166 | 79 | 89 | 123 | 82 |
| K20260 phenazine biosynthesis protein | 1399 | 1332 | 1291 | 1318 | 722 | 720 | 624 | 819 | 407 | 464 | 384 | 441 |
| K20261 trans-2,3-dihydro-3-hydroxyanthranilic acid synthase [EC:3.3.2.15] | 812 | 892 | 774 | 807 | 448 | 423 | 459 | 434 | 297 | 243 | 217 | 277 |
| K20262 dihydrophenazinedicarboxylate synthase [EC:1.10.3.16] | 542 | 526 | 459 | 447 | 256 | 242 | 243 | 256 | 153 | 136 | 115 | 134 |
| K20940 5-methylphenazine-1-carboxylate 1-monooxygenase [EC:1.14.13.218] | 1944 | 1773 | 1695 | 1788 | 1012 | 988 | 948 | 1026 | 591 | 551 | 573 | 536 |
| K21103 phenazine-1-carboxylate N-methyltransferase [EC:2.1.1.327] | 3 | 5 | 2 | 5 | 2 | 3 | 3 | 3 | 2 | 2 | 0 | 0 |
| ko00333 Prodigiosin biosynthesis | 18169 | 17674 | 17798 | 17780 | 14987 | 14646 | 14658 | 14869 | 13904 | 13766 | 13798 | 13809 |
| ko00254 Aflatoxin biosynthesis | 0 | 1 | 0 | 0 | 0 | 0 | 0 | 0 | 0 | 0 | 0 | 0 |
| ko00998 Biosynthesis of secondary metabolites - other antibiotics | 1602 | 1539 | 1528 | 1680 | 1581 | 1563 | 1563 | 1682 | 1555 | 1506 | 1587 | 1594 |
| K19548 dihydroanticapsin dehydrogenase [EC:1.1.1.385] | 0 | 0 | 0 | 1 | 0 | 0 | 0 | 0 | 0 | 1 | 0 | 1 |
| K21721 N-[(2S)-2-amino-2-carboxyethyl]-L-glutamate dehydrogenase [EC:1.5.1.51] | 5 | 6 | 10 | 6 | 57 | 62 | 52 | 54 | 13 | 10 | 15 | 23 |
| ko00999 Biosynthesis of secondary metabolites - unclassified | 25 | 51 | 51 | 37 | 97 | 120 | 81 | 91 | 40 | 35 | 59 | 75 |
| Xenobiotics biodegradation and metabolism | 199531 | 197852 | 190839 | 196157 | 142449 | 142050 | 138282 | 143945 | 152281 | 150905 | 148117 | 152118 |
| ko00362 Benzoate degradation | 70270 | 69863 | 66855 | 68977 | 44700 | 44506 | 43305 | 45274 | 55403 | 54819 | 53560 | 56141 |
| K00448 protocatechuate 3,4-dioxygenase, alpha subunit [EC:1.13.11.3] | 737 | 762 | 660 | 790 | 327 | 316 | 308 | 372 | 628 | 607 | 537 | 626 |
| K00449 protocatechuate 3,4-dioxygenase, beta subunit [EC:1.13.11.3] | 721 | 633 | 618 | 649 | 313 | 290 | 285 | 278 | 653 | 651 | 622 | 668 |
| K00481 p-hydroxybenzoate 3-monooxygenase [EC:1.14.13.2] | 919 | 937 | 924 | 986 | 413 | 399 | 411 | 394 | 1075 | 1205 | 1125 | 1166 |
| K01031 3-oxoadipate CoA-transferase, alpha subunit [EC:2.8.3.6] | 227 | 233 | 224 | 237 | 62 | 72 | 78 | 76 | 483 | 467 | 480 | 498 |
| K01032 3-oxoadipate CoA-transferase, beta subunit [EC:2.8.3.6] | 232 | 203 | 203 | 236 | 52 | 54 | 50 | 62 | 490 | 484 | 465 | 471 |
| K01055 3-oxoadipate enol-lactonase [EC:3.1.1.24] | 2533 | 2579 | 2441 | 2533 | 1371 | 1420 | 1303 | 1544 | 1404 | 1278 | 1265 | 1360 |
| K01607 4-carboxymuconolactone decarboxylase [EC:4.1.1.44] | 2843 | 2945 | 2818 | 2956 | 3284 | 3509 | 3411 | 3366 | 4000 | 3821 | 3975 | 3897 |
| K01821 4-oxalocrotonate tautomerase [EC:5.3.2.6] | 286 | 247 | 234 | 251 | 150 | 177 | 206 | 194 | 206 | 221 | 217 | 220 |
| K01856 muconate cycloisomerase [EC:5.5.1.1] | 1916 | 1833 | 1716 | 1885 | 888 | 867 | 798 | 906 | 1370 | 1326 | 1303 | 1316 |
| K01857 3-carboxy-cis,cis-muconate cycloisomerase [EC:5.5.1.2] | 2839 | 2790 | 2531 | 2662 | 1457 | 1398 | 1324 | 1481 | 1532 | 1502 | 1398 | 1626 |
| K03381 catechol 1,2-dioxygenase [EC:1.13.11.1] | 1027 | 979 | 895 | 903 | 462 | 441 | 418 | 451 | 811 | 823 | 753 | 885 |
| K03464 muconolactone D-isomerase [EC:5.3.3.4] | 530 | 514 | 524 | 509 | 248 | 269 | 214 | 282 | 383 | 382 | 338 | 377 |
| K04108 4-hydroxybenzoyl-CoA reductase subunit alpha [EC:1.3.7.9] | 4 | 6 | 4 | 5 | 1 | 3 | 1 | 1 | 4 | 1 | 2 | 1 |
| K04116 cyclohexanecarboxylate-CoA ligase [EC:6.2.1.-] | 4 | 0 | 2 | 1 | 0 | 0 | 1 | 0 | 2 | 8 | 4 | 0 |
| K05549 benzoate/toluate 1,2-dioxygenase subunit alpha [EC:1.14.12.10 1.14.12.-] | 2308 | 2308 | 2172 | 2153 | 1056 | 1027 | 953 | 1069 | 1568 | 1568 | 1504 | 1603 |
| K05550 benzoate/toluate 1,2-dioxygenase subunit beta [EC:1.14.12.10 1.14.12.-] | 924 | 904 | 755 | 851 | 426 | 421 | 395 | 474 | 546 | 519 | 569 | 560 |
| K05783 dihydroxycyclohexadiene carboxylate dehydrogenase [EC:1.3.1.25 1.3.1.-] | 1165 | 1103 | 1003 | 1077 | 439 | 486 | 480 | 532 | 758 | 705 | 684 | 804 |
| K05784 benzoate/toluate 1,2-dioxygenase reductase component [EC:1.18.1.-] | 1819 | 1665 | 1768 | 1794 | 920 | 866 | 763 | 911 | 1109 | 1109 | 1124 | 1222 |
| K07104 catechol 2,3-dioxygenase [EC:1.13.11.2] | 98 | 104 | 107 | 110 | 21 | 23 | 21 | 18 | 64 | 131 | 102 | 116 |
| K07535 2-hydroxycyclohexanecarboxyl-CoA dehydrogenase [EC:1.1.1.-] | 1 | 2 | 2 | 1 | 0 | 0 | 0 | 0 | 9 | 1 | 0 | 2 |
| K07536 2-ketocyclohexanecarboxyl-CoA hydrolase [EC:3.1.2.-] | 0 | 0 | 0 | 0 | 0 | 0 | 0 | 0 | 0 | 1 | 0 | 0 |
| K07823 3-oxoadipyl-CoA thiolase [EC:2.3.1.174] | 2241 | 2106 | 2116 | 2080 | 1212 | 1157 | 1123 | 1234 | 728 | 638 | 661 | 731 |
| K14333 2,3-dihydroxybenzoate decarboxylase [EC:4.1.1.46] | 401 | 428 | 378 | 390 | 112 | 131 | 104 | 105 | 879 | 837 | 755 | 875 |
| K16173 glutaryl-CoA dehydrogenase (non-decarboxylating) [EC:1.3.99.32] | 1 | 0 | 0 | 0 | 0 | 0 | 0 | 0 | 0 | 0 | 0 | 0 |
| K16514 4-oxalomesaconate tautomerase [EC:5.3.2.8] | 464 | 485 | 481 | 501 | 127 | 142 | 141 | 143 | 852 | 918 | 870 | 929 |
| K16515 4-oxalomesaconate hydratase [EC:4.2.1.83] | 317 | 350 | 297 | 342 | 95 | 68 | 87 | 96 | 609 | 666 | 664 | 689 |
| K18365 4-hydroxy-2-oxovalerate/4-hydroxy-2-oxohexanoate aldolase [EC:4.1.3.39 4.1.3.43] | 0 | 0 | 0 | 0 | 0 | 0 | 0 | 0 | 1 | 0 | 0 | 0 |
| K19066 cyclohex-1-ene-1-carbonyl-CoA dehydrogenase [EC:1.3.8.10] | 0 | 0 | 0 | 0 | 0 | 0 | 0 | 0 | 1 | 3 | 0 | 1 |
| K19067 cyclohexane-1-carbonyl-CoA dehydrogenase [EC:1.3.8.11] | 0 | 0 | 2 | 2 | 0 | 0 | 2 | 0 | 0 | 1 | 0 | 1 |
| ko00627 Aminobenzoate degradation | 28705 | 28147 | 25968 | 27495 | 15633 | 15452 | 14431 | 15936 | 15093 | 14652 | 14286 | 15017 |
| K00141 benzaldehyde dehydrogenase (NAD) [EC:1.2.1.28] | 0 | 0 | 2 | 1 | 0 | 0 | 0 | 0 | 1 | 0 | 3 | 0 |
| K01101 4-nitrophenyl phosphatase [EC:3.1.3.41] | 48 | 62 | 44 | 49 | 163 | 144 | 130 | 153 | 48 | 72 | 71 | 61 |
| K01576 benzoylformate decarboxylase [EC:4.1.1.7] | 2777 | 2649 | 2356 | 2493 | 1388 | 1423 | 1230 | 1496 | 901 | 759 | 726 | 782 |
| K01612 vanillate/4-hydroxybenzoate decarboxylase subunit C [EC:4.1.1.- 4.1.1.61] | 721 | 697 | 620 | 755 | 537 | 578 | 458 | 516 | 773 | 856 | 812 | 837 |
| K01781 mandelate racemase [EC:5.1.2.2] | 0 | 0 | 2 | 2 | 2 | 0 | 0 | 0 | 2 | 4 | 0 | 1 |
| K03862 vanillate monooxygenase [EC:1.14.13.82] | 1886 | 1818 | 1655 | 1723 | 1021 | 995 | 953 | 1066 | 580 | 531 | 468 | 595 |
| K03863 vanillate monooxygenase ferredoxin subunit | 1631 | 1612 | 1389 | 1511 | 931 | 804 | 810 | 909 | 525 | 418 | 442 | 439 |
| K04099 gallate dioxygenase [EC:1.13.11.57] | 542 | 613 | 544 | 617 | 176 | 176 | 140 | 136 | 1130 | 1066 | 1034 | 1091 |
| K05599 anthranilate 1,2-dioxygenase (deaminating, decarboxylating) large subunit [EC:1.14.12.1] | 1917 | 1855 | 1782 | 2006 | 1033 | 994 | 872 | 1032 | 659 | 654 | 533 | 614 |
| K05600 anthranilate 1,2-dioxygenase (deaminating, decarboxylating) small subunit [EC:1.14.12.1] | 711 | 688 | 649 | 648 | 359 | 306 | 345 | 333 | 217 | 203 | 206 | 160 |
| K08295 2-aminobenzoate-CoA ligase [EC:6.2.1.32] | 0 | 2 | 0 | 0 | 0 | 0 | 0 | 3 | 2 | 3 | 2 | 4 |
| K09461 anthraniloyl-CoA monooxygenase [EC:1.14.13.40] | 3 | 3 | 2 | 2 | 1 | 0 | 2 | 1 | 5 | 13 | 5 | 5 |
| K11311 anthranilate 1,2-dioxygenase reductase component [EC:1.18.1.-] | 1479 | 1341 | 1249 | 1296 | 717 | 652 | 634 | 752 | 391 | 379 | 367 | 419 |
| K14581 naphthalene 1,2-dioxygenase ferredoxin reductase component [EC:1.18.1.7] | 2 | 0 | 0 | 1 | 2 | 0 | 0 | 0 | 0 | 2 | 3 | 4 |
| K15054 (S)-mandelate dehydrogenase [EC:1.1.99.31] | 219 | 226 | 220 | 206 | 299 | 370 | 293 | 315 | 0 | 0 | 2 | 1 |
| K15066 vanillate/3-O-methylgallate O-demethylase [EC:2.1.1.341] | 4 | 11 | 5 | 7 | 0 | 0 | 0 | 0 | 0 | 0 | 1 | 0 |
| K20712 3-(hydroxyamino)phenol mutase [EC:5.4.4.3] | 2 | 0 | 0 | 0 | 0 | 0 | 0 | 0 | 0 | 0 | 0 | 0 |
| K21759 vanillate/4-hydroxybenzoate decarboxylase subunit D [EC:4.1.1.- 4.1.1.61] | 131 | 174 | 132 | 165 | 82 | 120 | 114 | 107 | 217 | 195 | 198 | 206 |
| K21802 vanillin dehydrogenase [EC:1.2.1.67] | 10 | 9 | 7 | 15 | 0 | 2 | 2 | 1 | 2 | 2 | 3 | 4 |
| K22958 gallate decarboxylase subunit C [EC:4.1.1.59] | 1567 | 1502 | 1451 | 1528 | 790 | 784 | 745 | 755 | 1880 | 1888 | 1915 | 2041 |
| K22959 gallate decarboxylase subunit B | 46 | 42 | 50 | 48 | 8 | 5 | 7 | 6 | 39 | 37 | 25 | 34 |
| K22960 gallate decarboxylase subunit D | 0 | 0 | 0 | 1 | 3 | 4 | 9 | 1 | 5 | 11 | 10 | 4 |
| ko00364 Fluorobenzoate degradation | 11923 | 11475 | 10761 | 11216 | 5574 | 5506 | 5066 | 5738 | 8307 | 8265 | 7839 | 8534 |
| K01061 carboxymethylenebutenolidase [EC:3.1.1.45] | 2764 | 2683 | 2452 | 2553 | 1383 | 1398 | 1259 | 1395 | 2144 | 2215 | 1902 | 2144 |
| ko00625 Chloroalkane and chloroalkene degradation | 23344 | 23348 | 22868 | 23078 | 13900 | 13717 | 13373 | 14123 | 13057 | 13185 | 12792 | 12980 |
| K01560 2-haloacid dehalogenase [EC:3.8.1.2] | 1018 | 1018 | 910 | 991 | 470 | 427 | 459 | 529 | 448 | 416 | 417 | 474 |
| K01561 haloacetate dehalogenase [EC:3.8.1.3] | 1186 | 1173 | 1149 | 1175 | 609 | 610 | 575 | 648 | 473 | 373 | 349 | 394 |
| ko00361 Chlorocyclohexane and chlorobenzene degradation | 8845 | 8646 | 8012 | 8465 | 4252 | 4247 | 3967 | 4404 | 5607 | 5527 | 5098 | 5588 |
| K21607 tetrachlorobenzoquinone reductase [EC:1.1.1.404] | 836 | 856 | 783 | 848 | 419 | 481 | 437 | 457 | 297 | 243 | 272 | 259 |

| | | | | | | | | | | | | |
|---|---|---|---|---|---|---|---|---|---|---|---|---|
| ko00623 Toluene degradation | 6114 | 5879 | 5469 | 5807 | 2841 | 2841 | 2570 | 2893 | 5207 | 5255 | 4797 | 5189 |
| K07540 benzylsuccinate synthase [EC:4.1.99.11] | 0 | 5 | 2 | 0 | 3 | 0 | 0 | 0 | 0 | 0 | 3 | 0 |
| K07546 E-phenylitaconyl-CoA hydratase [EC:4.2.1.-] | 0 | 0 | 0 | 0 | 0 | 0 | 0 | 1 | 0 | 0 | 0 | 0 |
| K20199 4-hydroxybenzaldehyde dehydrogenase (NADP+) [EC:1.2.1.96] | 1 | 0 | 0 | 0 | 1 | 0 | 0 | 1 | 0 | 0 | 0 | 0 |
| ko00622 Xylene degradation | 9871 | 9674 | 9163 | 9556 | 4523 | 4632 | 4263 | 4765 | 7636 | 7565 | 7427 | 7904 |
| K10619 p-cumate 2,3-dioxygenase subunit alpha [EC:1.14.12.25] | 1712 | 1828 | 1572 | 1686 | 916 | 891 | 805 | 935 | 538 | 464 | 460 | 527 |
| ko00633 Nitrotoluene degradation | 6332 | 6311 | 6128 | 6155 | 4941 | 5183 | 5347 | 5250 | 4905 | 5077 | 4827 | 5081 |
| K06281 hydrogenase large subunit [EC:1.12.99.6] | 537 | 526 | 562 | 504 | 733 | 803 | 760 | 739 | 400 | 415 | 383 | 415 |
| K06282 hydrogenase small subunit [EC:1.12.99.6] | 357 | 315 | 329 | 370 | 491 | 441 | 472 | 475 | 245 | 255 | 265 | 314 |
| K10678 nitroreductase [EC:1.-.-.-] | 464 | 467 | 415 | 442 | 267 | 309 | 305 | 247 | 491 | 580 | 574 | 608 |
| K10679 nitroreductase / dihydropteridine reductase [EC:1.-.-.- 1.5.1.34] | 441 | 456 | 399 | 425 | 353 | 399 | 366 | 352 | 613 | 669 | 593 | 582 |
| K10680 N-ethylmaleimide reductase [EC:1.-.-.-] | 3703 | 3635 | 3487 | 3532 | 2135 | 2195 | 2304 | 2345 | 1508 | 1370 | 1272 | 1375 |
| ko00642 Ethylbenzene degradation | 6773 | 6698 | 6402 | 6637 | 4033 | 3892 | 3923 | 3925 | 3815 | 3726 | 3661 | 3843 |
| ko00643 Styrene degradation | 18851 | 18860 | 17262 | 18375 | 9778 | 9520 | 9073 | 10010 | 9489 | 9345 | 9140 | 9558 |
| K01502 aliphatic nitrilase [EC:3.5.5.7] | 0 | 1 | 0 | 0 | 0 | 0 | 0 | 0 | 4 | 6 | 2 | 9 |
| ko00791 Atrazine degradation | 6469 | 6237 | 5767 | 6209 | 4067 | 4011 | 3908 | 4196 | 3783 | 3807 | 3628 | 3940 |
| K03382 hydroxydechloroatrazine ethylaminohydrolase [EC:3.5.4.43] | 35 | 20 | 22 | 35 | 2 | 5 | 3 | 0 | 19 | 28 | 24 | 20 |
| K03383 cyanuric acid amidohydrolase [EC:3.5.2.15] | 3 | 0 | 0 | 0 | 0 | 0 | 0 | 0 | 0 | 0 | 0 | 0 |
| K19837 1-carboxybiuret hydrolase [EC:3.5.1.131] | 1 | 0 | 0 | 0 | 0 | 0 | 0 | 0 | 0 | 0 | 0 | 0 |
| K23359 biuret amidohydrolase [EC:3.5.1.84] | 9 | 5 | 14 | 12 | 1 | 3 | 2 | 2 | 5 | 6 | 2 | 1 |
| ko00930 Caprolactam degradation | 17149 | 16501 | 16484 | 16659 | 9697 | 9599 | 8993 | 9768 | 8999 | 8844 | 8740 | 9199 |
| K06446 acyl-CoA dehydrogenase [EC:1.3.99.-] | 1797 | 1620 | 1614 | 1787 | 889 | 907 | 798 | 977 | 573 | 569 | 589 | 532 |
| K17754 cyclopentanol dehydrogenase [EC:1.1.1.163] | 0 | 0 | 0 | 0 | 0 | 0 | 0 | 0 | 0 | 0 | 1 | 0 |
| K19962 6-oxohexanoate dehydrogenase [EC:1.2.1.63] | 1 | 0 | 0 | 0 | 0 | 1 | 0 | 0 | 0 | 0 | 0 | 0 |
| ko00363 Bisphenol degradation | 861 | 817 | 768 | 864 | 467 | 483 | 480 | 503 | 290 | 239 | 272 | 286 |
| K05913 2,4'-dihydroxyacetophenone dioxygenase [EC:1.13.11.41] | 861 | 817 | 768 | 864 | 467 | 483 | 480 | 503 | 290 | 239 | 272 | 286 |
| ko00621 Dioxin degradation | 1439 | 1383 | 1382 | 1420 | 641 | 783 | 751 | 687 | 2173 | 2181 | 2151 | 2229 |
| K00480 salicylate hydroxylase [EC:1.14.13.1] | 1 | 0 | 0 | 1 | 0 | 0 | 0 | 0 | 2 | 3 | 3 | 1 |
| ko00626 Naphthalene degradation | 10423 | 10467 | 10613 | 10543 | 6135 | 6295 | 6069 | 6079 | 8427 | 8877 | 8655 | 8530 |
| ko00624 Polycyclic aromatic hydrocarbon degradation | 1461 | 1395 | 1278 | 1441 | 642 | 606 | 593 | 650 | 1283 | 1263 | 1165 | 1299 |
| ko00365 Furfural degradation | 32 | 44 | 36 | 20 | 1 | 2 | 3 | 4 | 11 | 9 | 7 | 9 |
| K16873 5-(hydroxymethyl)furfural/furfural oxidase [EC:1.1.3.47 1.1.3.-] | 0 | 0 | 0 | 0 | 0 | 0 | 0 | 2 | 0 | 0 | 0 | 0 |
| K16874 2,5-furandicarboxylate decarboxylase 1 | 30 | 44 | 35 | 20 | 1 | 2 | 3 | 2 | 10 | 9 | 7 | 9 |
| K16875 2,5-furandicarboxylate decarboxylase 2 | 1 | 0 | 1 | 0 | 0 | 0 | 0 | 0 | 1 | 0 | 0 | 0 |
| K16880 2-oxoglutaroyl-CoA hydrolase | 1 | 0 | 0 | 0 | 0 | 0 | 0 | 0 | 0 | 0 | 0 | 0 |
| ko00984 Steroid degradation | 8 | 2 | 0 | 7 | 0 | 4 | 1 | 2 | 5 | 1 | 4 | 3 |
| K03333 cholesterol oxidase [EC:1.1.3.6] | 0 | 0 | 0 | 2 | 0 | 2 | 0 | 0 | 2 | 0 | 1 | 1 |
| K05898 3-oxosteroid 1-dehydrogenase [EC:1.3.99.4] | 3 | 1 | 0 | 1 | 0 | 0 | 1 | 2 | 2 | 1 | 2 | 1 |
| K16048 3-hydroxy-9,10-secoandrosta-1,3,5(10)-triene-9,17-dione monooxygenase reductase component [EC:1.5.1.-] | 1 | 1 | 0 | 0 | 0 | 1 | 0 | 0 | 0 | 0 | 0 | 0 |
| K16050 4,5:9,10-diseco-3-hydroxy-5,9,17-trioxoandrosta-1(10),2-diene-4-oate hydrolase [EC:3.7.1.17] | 2 | 0 | 0 | 3 | 0 | 0 | 0 | 0 | 0 | 0 | 0 | 0 |
| K18687 HIP---CoA ligase [EC:6.2.1.41] | 1 | 0 | 0 | 0 | 0 | 1 | 0 | 0 | 0 | 0 | 0 | 0 |
| ko00980 Metabolism of xenobiotics by cytochrome P450 | 13914 | 13780 | 13393 | 13756 | 8198 | 8210 | 7931 | 8189 | 9759 | 9639 | 9396 | 9631 |
| K15303 aflatoxin B1 aldehyde reductase | 55 | 56 | 49 | 40 | 4 | 8 | 3 | 7 | 97 | 117 | 97 | 103 |
| ko00982 Drug metabolism - cytochrome P450 | 16397 | 16383 | 15952 | 16301 | 9639 | 9610 | 9235 | 9726 | 10520 | 10392 | 10082 | 10270 |
| ko00983 Drug metabolism - other enzymes | 42944 | 42575 | 43326 | 43204 | 48183 | 48290 | 47711 | 47793 | 48778 | 48281 | 48403 | 47848 |
| K00569 thiopurine S-methyltransferase [EC:2.1.1.67] | 946 | 930 | 910 | 899 | 551 | 523 | 438 | 477 | 310 | 255 | 258 | 286 |
| K01044 carboxylesterase 1 [EC:3.1.1.1] | 0 | 0 | 1 | 0 | 0 | 0 | 0 | 0 | 0 | 1 | 0 | 0 |
| K03927 carboxylesterase 2 [EC:3.1.1.1 3.1.1.84 3.1.1.56] | 0 | 0 | 0 | 0 | 0 | 0 | 0 | 0 | 1 | 0 | 0 | 0 |
| Not included in regular maps | 3286 | 3287 | 3218 | 3416 | 1938 | 2038 | 1810 | 1973 | 1851 | 1965 | 1844 | 1886 |
| ko09113 Global maps only | 3286 | 3287 | 3218 | 3416 | 1938 | 2038 | 1810 | 1973 | 1851 | 1965 | 1844 | 1886 |
| K00528 ferredoxin/flavodoxin---NADP+ reductase [EC:1.18.1.2 1.19.1.1] | 3105 | 3135 | 3077 | 3251 | 1703 | 1779 | 1580 | 1759 | 1851 | 1965 | 1844 | 1883 |
| K08088 fimbrial protein FimY | 181 | 152 | 141 | 165 | 235 | 259 | 230 | 214 | 0 | 0 | 0 | 3 |
| Genetic Information Processing | 688494 | 689615 | 696901 | 695431 | 729254 | 736705 | 735480 | 727754 | 712116 | 714939 | 716173 | 713046 |
| Transcription | 27799 | 28114 | 28971 | 28527 | 34261 | 35071 | 34594 | 33772 | 33945 | 34478 | 34784 | 33403 |
| ko03020 RNA polymerase | 27778 | 28095 | 28948 | 28514 | 34250 | 35071 | 34590 | 33769 | 33902 | 34425 | 34744 | 33346 |
| K02999 DNA-directed RNA polymerase I subunit RPA1 [EC:2.7.7.6] | 0 | 0 | 0 | 0 | 0 | 0 | 0 | 0 | 2 | 0 | 0 | 0 |
| K03002 DNA-directed RNA polymerase I subunit RPA2 [EC:2.7.7.6] | 0 | 2 | 0 | 0 | 0 | 0 | 0 | 0 | 0 | 0 | 0 | 0 |
| K03011 DNA-directed RNA polymerase II subunit RPB3 | 0 | 2 | 0 | 0 | 0 | 0 | 0 | 0 | 0 | 0 | 0 | 0 |
| K03040 DNA-directed RNA polymerase subunit alpha [EC:2.7.7.6] | 3331 | 3411 | 3419 | 3372 | 3946 | 3945 | 3957 | 3979 | 3914 | 3962 | 3948 | 3902 |
| K03041 DNA-directed RNA polymerase subunit A' [EC:2.7.7.6] | 2 | 0 | 0 | 3 | 3 | 0 | 3 | 3 | 18 | 31 | 13 | 32 |
| K03042 DNA-directed RNA polymerase subunit A [EC:2.7.7.6] | 2 | 0 | 8 | 0 | 2 | 0 | 0 | 1 | 12 | 16 | 4 | 7 |
| K03043 DNA-directed RNA polymerase subunit beta [EC:2.7.7.6] | 10872 | 11269 | 11552 | 11363 | 14140 | 14675 | 14483 | 13922 | 13783 | 14035 | 14231 | 13540 |
| K03044 DNA-directed RNA polymerase subunit B' [EC:2.7.7.6] | 2 | 0 | 2 | 1 | 0 | 0 | 2 | 0 | 4 | 12 | 8 | 15 |
| K03045 DNA-directed RNA polymerase subunit B [EC:2.7.7.6] | 0 | 1 | 1 | 1 | 0 | 1 | 0 | 2 | 2 | 4 | 8 | 6 |
| K03046 DNA-directed RNA polymerase subunit beta' [EC:2.7.7.6] | 12740 | 12599 | 13223 | 12943 | 15703 | 15944 | 15695 | 15415 | 15625 | 15800 | 15959 | 15315 |
| K03047 DNA-directed RNA polymerase subunit D [EC:2.7.7.6] | 1 | 2 | 0 | 0 | 0 | 1 | 1 | 0 | 1 | 5 | 2 | 6 |
| K03048 DNA-directed RNA polymerase subunit delta | 20 | 27 | 11 | 23 | 4 | 5 | 5 | 7 | 24 | 13 | 20 | 27 |
| K03049 DNA-directed RNA polymerase subunit E' [EC:2.7.7.6] | 0 | 0 | 0 | 1 | 0 | 0 | 0 | 0 | 0 | 7 | 0 | 3 |
| K03050 DNA-directed RNA polymerase subunit E [EC:2.7.7.6] | 0 | 0 | 0 | 0 | 0 | 0 | 0 | 1 | 1 | 0 | 2 | 1 |
| K03051 DNA-directed RNA polymerase subunit F [EC:2.7.7.6] | 0 | 0 | 0 | 0 | 0 | 0 | 0 | 1 | 0 | 1 | 3 | 0 |
| K03053 DNA-directed RNA polymerase subunit H [EC:2.7.7.6] | 0 | 0 | 0 | 0 | 2 | 0 | 2 | 0 | 5 | 4 | 0 | 1 |
| K03055 DNA-directed RNA polymerase subunit K [EC:2.7.7.6] | 2 | 0 | 0 | 0 | 0 | 0 | 0 | 0 | 3 | 0 | 1 | 1 |
| K03056 DNA-directed RNA polymerase subunit L [EC:2.7.7.6] | 0 | 0 | 0 | 1 | 0 | 0 | 0 | 0 | 0 | 2 | 2 | 1 |
| K03058 DNA-directed RNA polymerase subunit N [EC:2.7.7.6] | 0 | 0 | 0 | 0 | 2 | 0 | 0 | 0 | 0 | 0 | 0 | 1 |
| K03059 DNA-directed RNA polymerase subunit P [EC:2.7.7.6] | 0 | 0 | 0 | 0 | 0 | 0 | 0 | 0 | 0 | 1 | 1 | 1 |
| K03060 DNA-directed RNA polymerase subunit omega [EC:2.7.7.6] | 805 | 781 | 730 | 805 | 447 | 499 | 442 | 437 | 508 | 531 | 538 | 482 |
| K13797 DNA-directed RNA polymerase subunit beta-beta' [EC:2.7.7.6] | 1 | 1 | 0 | 0 | 1 | 0 | 0 | 1 | 0 | 0 | 1 | 0 |
| K13798 DNA-directed RNA polymerase subunit B [EC:2.7.7.6] | 0 | 0 | 2 | 1 | 0 | 1 | 0 | 0 | 0 | 1 | 3 | 5 |
| ko03022 Basal transcription factors | 3 | 2 | 4 | 1 | 11 | 0 | 4 | 3 | 14 | 24 | 10 | 16 |
| K03120 transcription initiation factor TFIID TATA-box-binding protein | 0 | 2 | 0 | 0 | 2 | 0 | 0 | 0 | 3 | 4 | 3 | 3 |
| K03124 transcription initiation factor TFIIB | 0 | 0 | 2 | 1 | 3 | 0 | 0 | 0 | 5 | 7 | 3 | 6 |
| K03135 transcription initiation factor TFIID subunit 11 | 0 | 0 | 0 | 0 | 1 | 0 | 0 | 0 | 0 | 0 | 0 | 0 |
| K03136 transcription initiation factor TFIIE subunit alpha | 0 | 0 | 2 | 0 | 3 | 0 | 0 | 0 | 3 | 4 | 1 | 0 |
| K10844 DNA excision repair protein ERCC-2 [EC:3.6.4.12] | 3 | 0 | 0 | 0 | 2 | 0 | 4 | 3 | 3 | 9 | 3 | 7 |
| ko03040 Spliceosome | 18 | 17 | 19 | 12 | 0 | 0 | 0 | 0 | 29 | 29 | 30 | 41 |
| K03283 heat shock 70kDa protein 1/2/6/8 | 0 | 0 | 1 | 0 | 0 | 0 | 0 | 0 | 0 | 0 | 0 | 1 |
| K12733 peptidyl-prolyl cis-trans isomerase-like 1 [EC:5.2.1.8] | 18 | 13 | 17 | 12 | 0 | 0 | 0 | 0 | 29 | 27 | 30 | 39 |
| K12823 ATP-dependent RNA helicase DDX5/DBP2 [EC:3.6.4.13] | 0 | 1 | 1 | 0 | 0 | 0 | 0 | 0 | 0 | 2 | 0 | 1 |
| K12827 splicing factor 3A subunit 3 | 0 | 1 | 0 | 0 | 0 | 0 | 0 | 0 | 0 | 0 | 0 | 0 |
| K12856 pre-mRNA-processing factor 8 | 0 | 2 | 0 | 0 | 0 | 0 | 0 | 0 | 0 | 0 | 0 | 0 |
| Translation | 231419 | 233704 | 237905 | 236785 | 242947 | 245463 | 245107 | 240471 | 239930 | 242047 | 241466 | 242016 |
| ko03010 Ribosome | 78384 | 79586 | 82698 | 81135 | 88612 | 88928 | 90151 | 86988 | 84998 | 85396 | 85577 | 85767 |
| K02863 large subunit ribosomal protein L1 | 2408 | 2351 | 2592 | 2524 | 2685 | 2884 | 2814 | 2668 | 2571 | 2730 | 2743 | 2735 |
| K02864 large subunit ribosomal protein L10 | 1440 | 1481 | 1633 | 1602 | 1914 | 2024 | 1966 | 1858 | 1801 | 1851 | 1798 | 1729 |
| K02865 large subunit ribosomal protein L10Ae | 0 | 0 | 0 | 0 | 0 | 0 | 0 | 0 | 0 | 2 | 0 | 0 |
| K02866 large subunit ribosomal protein L10e | 0 | 0 | 0 | 0 | 0 | 0 | 0 | 0 | 2 | 6 | 3 | 3 |
| K02867 large subunit ribosomal protein L11 | 1591 | 1667 | 1685 | 1674 | 1764 | 1771 | 1747 | 1756 | 1766 | 1760 | 1759 | 1733 |
| K02869 large subunit ribosomal protein L12 | 0 | 0 | 0 | 0 | 0 | 0 | 0 | 1 | 0 | 0 | 0 | 1 |
| K02871 large subunit ribosomal protein L13 | 1662 | 1700 | 1614 | 1696 | 1833 | 1844 | 1712 | 1764 | 1728 | 1808 | 1830 | 1796 |
| K02874 large subunit ribosomal protein L14 | 1469 | 1320 | 1396 | 1436 | 1491 | 1581 | 1519 | 1445 | 1482 | 1457 | 1404 | 1498 |
| K02875 large subunit ribosomal protein L14e | 0 | 0 | 0 | 0 | 0 | 0 | 0 | 0 | 2 | 1 | 2 | 0 |
| K02876 large subunit ribosomal protein L15 | 1576 | 1651 | 1688 | 1715 | 1793 | 1799 | 1953 | 1904 | 1737 | 1842 | 1831 | 1831 |
| K02877 large subunit ribosomal protein L15e | 0 | 0 | 0 | 1 | 0 | 1 | 0 | 0 | 0 | 6 | 2 | 7 |
| K02878 large subunit ribosomal protein L16 | 1648 | 1558 | 1642 | 1643 | 1743 | 1865 | 1762 | 1839 | 1733 | 1706 | 1843 | 1807 |
| K02879 large subunit ribosomal protein L17 | 1628 | 1563 | 1778 | 1660 | 1737 | 1696 | 1762 | 1684 | 1631 | 1699 | 1630 | 1751 |
| K02881 large subunit ribosomal protein L18 | 1263 | 1294 | 1374 | 1288 | 1426 | 1545 | 1530 | 1405 | 1379 | 1348 | 1348 | 1351 |
| K02883 large subunit ribosomal protein L18e | 1 | 0 | 2 | 1 | 0 | 1 | 0 | 2 | 0 | 3 | 0 | 4 |
| K02884 large subunit ribosomal protein L19 | 1192 | 1116 | 1279 | 1160 | 1434 | 1347 | 1356 | 1299 | 1346 | 1282 | 1262 | 1338 |
| K02885 large subunit ribosomal protein L19e | 0 | 0 | 2 | 0 | 3 | 0 | 0 | 3 | 4 | 4 | 0 | 3 |
| K02886 large subunit ribosomal protein L2 | 2448 | 2431 | 2632 | 2452 | 3071 | 2911 | 3201 | 3021 | 3097 | 3168 | 3032 | 3127 |
| K02887 large subunit ribosomal protein L20 | 1118 | 1197 | 1213 | 1168 | 1210 | 1182 | 1240 | 1177 | 1136 | 1193 | 1261 | 1170 |
| K02888 large subunit ribosomal protein L21 | 1165 | 1264 | 1267 | 1222 | 1321 | 1258 | 1238 | 1174 | 1319 | 1301 | 1196 | 1257 |
| K02889 large subunit ribosomal protein L21e | 3 | 0 | 0 | 0 | 0 | 0 | 0 | 2 | 0 | 3 | 2 | 1 |
| K02890 large subunit ribosomal protein L22 | 1298 | 1351 | 1419 | 1459 | 1500 | 1558 | 1598 | 1465 | 1512 | 1601 | 1475 | 1395 |
| K02892 large subunit ribosomal protein L23 | 1159 | 1096 | 1155 | 1156 | 1172 | 1204 | 1229 | 1129 | 1205 | 1194 | 1273 | 1241 |
| K02895 large subunit ribosomal protein L24 | 1186 | 1113 | 1141 | 1166 | 1208 | 1300 | 1280 | 1233 | 1192 | 1251 | 1198 | 1132 |
| K02896 large subunit ribosomal protein L24e | 0 | 0 | 0 | 0 | 0 | 0 | 0 | 1 | 0 | 0 | 0 | 4 |
| K02897 large subunit ribosomal protein L25 | 1730 | 1746 | 1690 | 1746 | 2019 | 2068 | 1941 | 1977 | 1629 | 1638 | 1746 | 1725 |
| K02899 large subunit ribosomal protein L27 | 1075 | 1123 | 1068 | 1099 | 1078 | 1112 | 1105 | 1014 | 1057 | 1125 | 1033 | 1086 |
| K02902 large subunit ribosomal protein L28 | 807 | 845 | 802 | 790 | 950 | 911 | 991 | 840 | 834 | 797 | 827 | 827 |
| K02904 large subunit ribosomal protein L29 | 719 | 730 | 798 | 711 | 780 | 784 | 833 | 813 | 740 | 737 | 753 | 774 |

| | | | | | | | | | | | | |
|---|---|---|---|---|---|---|---|---|---|---|---|---|
| K02906 large subunit ribosomal protein L3 | 2422 | 2528 | 2666 | 2514 | 2605 | 2682 | 2652 | 2572 | 2428 | 2222 | 2407 | 2424 |
| K02907 large subunit ribosomal protein L30 | 720 | 677 | 723 | 687 | 723 | 736 | 728 | 763 | 704 | 736 | 693 | 751 |
| K02908 large subunit ribosomal protein L30e | 0 | 0 | 0 | 0 | 0 | 0 | 0 | 1 | 2 | 0 | 0 | 5 |
| K02909 large subunit ribosomal protein L31 | 1479 | 1511 | 1535 | 1459 | 1428 | 1386 | 1325 | 1296 | 1325 | 1355 | 1276 | 1246 |
| K02910 large subunit ribosomal protein L31e | 0 | 0 | 0 | 0 | 0 | 0 | 0 | 0 | 2 | 0 | 2 | 1 |
| K02911 large subunit ribosomal protein L32 | 615 | 620 | 672 | 679 | 736 | 785 | 777 | 747 | 762 | 662 | 739 | 767 |
| K02912 large subunit ribosomal protein L32e | 0 | 2 | 0 | 0 | 0 | 2 | 0 | 3 | 4 | 6 | 0 | 3 |
| K02913 large subunit ribosomal protein L33 | 577 | 649 | 656 | 666 | 636 | 660 | 692 | 608 | 555 | 628 | 653 | 641 |
| K02914 large subunit ribosomal protein L34 | 519 | 553 | 519 | 516 | 535 | 544 | 611 | 611 | 523 | 522 | 551 | 560 |
| K02915 large subunit ribosomal protein L34e | 2 | 0 | 0 | 0 | 0 | 0 | 0 | 0 | 2 | 0 | 0 | 1 |
| K02916 large subunit ribosomal protein L35 | 561 | 629 | 650 | 576 | 641 | 668 | 681 | 653 | 561 | 639 | 627 | 639 |
| K02919 large subunit ribosomal protein L36 | 876 | 781 | 783 | 888 | 689 | 704 | 714 | 680 | 649 | 654 | 690 | 696 |
| K02921 large subunit ribosomal protein L37Ae | 0 | 0 | 0 | 0 | 0 | 0 | 0 | 0 | 0 | 2 | 1 | 4 |
| K02922 large subunit ribosomal protein L37e | 1 | 0 | 0 | 1 | 0 | 0 | 0 | 0 | 0 | 0 | 0 | 0 |
| K02924 large subunit ribosomal protein L39e | 0 | 0 | 0 | 0 | 0 | 0 | 0 | 0 | 0 | 0 | 1 | 1 |
| K02925 large subunit ribosomal protein L3e | 0 | 0 | 2 | 0 | 0 | 0 | 0 | 0 | 0 | 0 | 0 | 0 |
| K02926 large subunit ribosomal protein L4 | 1580 | 1777 | 1733 | 1774 | 2221 | 2155 | 2306 | 2168 | 2291 | 2242 | 2276 | 2211 |
| K02927 large subunit ribosomal protein L40e | 0 | 0 | 0 | 1 | 0 | 0 | 0 | 0 | 0 | 0 | 0 | 0 |
| K02929 large subunit ribosomal protein L44e | 1 | 0 | 0 | 0 | 0 | 1 | 0 | 0 | 0 | 0 | 0 | 0 |
| K02930 large subunit ribosomal protein L4e | 0 | 2 | 0 | 0 | 1 | 2 | 0 | 0 | 4 | 13 | 2 | 5 |
| K02931 large subunit ribosomal protein L5 | 2074 | 2130 | 2145 | 2160 | 2287 | 2503 | 2277 | 2328 | 2283 | 2398 | 2251 | 2311 |
| K02933 large subunit ribosomal protein L6 | 1988 | 2022 | 2134 | 2056 | 2398 | 2359 | 2380 | 2238 | 2135 | 2095 | 2264 | 2030 |
| K02935 large subunit ribosomal protein L7/L12 | 1405 | 1452 | 1385 | 1414 | 1499 | 1579 | 1484 | 1286 | 1447 | 1424 | 1454 | 1451 |
| K02936 large subunit ribosomal protein L7Ae | 0 | 0 | 0 | 0 | 0 | 0 | 0 | 0 | 1 | 8 | 1 | 2 |
| K02939 large subunit ribosomal protein L9 | 1565 | 1528 | 1625 | 1664 | 1731 | 1765 | 1842 | 1787 | 1639 | 1731 | 1682 | 1704 |
| K02942 large subunit ribosomal protein LP1 | 0 | 1 | 0 | 0 | 0 | 0 | 0 | 0 | 0 | 0 | 0 | 0 |
| K02944 large subunit ribosomal protein LX | 1 | 0 | 0 | 1 | 0 | 0 | 0 | 0 | 0 | 0 | 0 | 4 |
| K02945 small subunit ribosomal protein S1 | 5297 | 5553 | 5571 | 5476 | 6499 | 6100 | 6274 | 6259 | 6058 | 5901 | 6041 | 5817 |
| K02946 small subunit ribosomal protein S10 | 1122 | 1208 | 1283 | 1176 | 1339 | 1210 | 1344 | 1192 | 1259 | 1272 | 1334 | 1320 |
| K02948 small subunit ribosomal protein S11 | 1380 | 1422 | 1511 | 1437 | 1624 | 1650 | 1636 | 1590 | 1580 | 1585 | 1512 | 1511 |
| K02950 small subunit ribosomal protein S12 | 1453 | 1585 | 1598 | 1651 | 1590 | 1616 | 1545 | 1608 | 1565 | 1594 | 1694 | 1688 |
| K02952 small subunit ribosomal protein S13 | 1414 | 1417 | 1475 | 1412 | 1529 | 1551 | 1600 | 1449 | 1437 | 1450 | 1426 | 1471 |
| K02954 small subunit ribosomal protein S14 | 1149 | 1177 | 1261 | 1209 | 1234 | 1272 | 1236 | 1221 | 1210 | 1203 | 1182 | 1189 |
| K02956 small subunit ribosomal protein S15 | 888 | 988 | 981 | 1024 | 1127 | 1017 | 1091 | 1126 | 1045 | 1049 | 997 | 1020 |
| K02957 small subunit ribosomal protein S15Ae | 0 | 0 | 0 | 0 | 0 | 0 | 0 | 0 | 0 | 0 | 0 | 1 |
| K02958 small subunit ribosomal protein S15e | 0 | 0 | 0 | 0 | 1 | 0 | 0 | 0 | 0 | 0 | 0 | 0 |
| K02959 small subunit ribosomal protein S16 | 985 | 973 | 1145 | 996 | 1233 | 1259 | 1265 | 1267 | 1088 | 973 | 1166 | 1189 |
| K02961 small subunit ribosomal protein S17 | 889 | 887 | 1090 | 982 | 1083 | 1119 | 1151 | 1021 | 1037 | 1097 | 1045 | 1062 |
| K02962 small subunit ribosomal protein S17e | 0 | 2 | 1 | 0 | 0 | 0 | 0 | 0 | 0 | 2 | 0 | 1 |
| K02963 small subunit ribosomal protein S18 | 947 | 1005 | 1069 | 1049 | 1040 | 1078 | 1139 | 1115 | 988 | 984 | 997 | 1097 |
| K02965 small subunit ribosomal protein S19 | 1047 | 1114 | 1156 | 1082 | 1191 | 1151 | 1111 | 1113 | 1245 | 1217 | 1150 | 1202 |
| K02966 small subunit ribosomal protein S19e | 1 | 0 | 0 | 3 | 0 | 0 | 0 | 0 | 0 | 5 | 2 | 1 |
| K02967 small subunit ribosomal protein S2 | 2361 | 2530 | 2496 | 2533 | 2993 | 2977 | 3183 | 2906 | 2881 | 2807 | 2899 | 2836 |
| K02968 small subunit ribosomal protein S20 | 861 | 849 | 910 | 796 | 1084 | 1025 | 1104 | 1033 | 813 | 915 | 949 | 925 |
| K02970 small subunit ribosomal protein S21 | 738 | 727 | 741 | 758 | 842 | 717 | 877 | 856 | 950 | 946 | 860 | 875 |
| K02974 small subunit ribosomal protein S24e | 0 | 0 | 0 | 0 | 0 | 0 | 0 | 0 | 0 | 0 | 0 | 1 |
| K02977 small subunit ribosomal protein S27Ae | 0 | 2 | 0 | 0 | 0 | 0 | 0 | 0 | 0 | 1 | 1 | 1 |
| K02978 small subunit ribosomal protein S27e | 0 | 0 | 0 | 0 | 0 | 2 | 0 | 0 | 0 | 0 | 0 | 0 |
| K02979 small subunit ribosomal protein S28e | 0 | 0 | 0 | 1 | 0 | 0 | 0 | 0 | 1 | 4 | 2 | 3 |
| K02982 small subunit ribosomal protein S3 | 2646 | 2621 | 2896 | 2758 | 2817 | 2979 | 2892 | 2890 | 2876 | 2768 | 2823 | 2881 |
| K02984 small subunit ribosomal protein S3Ae | 0 | 0 | 0 | 0 | 0 | 2 | 0 | 0 | 0 | 5 | 7 | 10 |
| K02986 small subunit ribosomal protein S4 | 2416 | 2453 | 2472 | 2458 | 2697 | 2637 | 2661 | 2736 | 2594 | 2622 | 2467 | 2552 |
| K02987 small subunit ribosomal protein S4e | 5 | 1 | 0 | 1 | 2 | 0 | 0 | 0 | 11 | 6 | 2 | 2 |
| K02988 small subunit ribosomal protein S5 | 1862 | 1807 | 2030 | 2022 | 2011 | 2083 | 2176 | 2003 | 1938 | 1903 | 1886 | 1967 |
| K02990 small subunit ribosomal protein S6 | 1218 | 1223 | 1222 | 1202 | 1237 | 1233 | 1292 | 1298 | 1183 | 1195 | 1283 | 1276 |
| K02991 small subunit ribosomal protein S6e | 2 | 0 | 0 | 0 | 0 | 0 | 0 | 0 | 0 | 4 | 0 | 3 |
| K02992 small subunit ribosomal protein S7 | 1658 | 1621 | 1658 | 1635 | 1876 | 1882 | 1883 | 1920 | 1931 | 1854 | 1953 | 1873 |
| K02994 small subunit ribosomal protein S8 | 1525 | 1527 | 1555 | 1578 | 1731 | 1655 | 1777 | 1640 | 1566 | 1660 | 1600 | 1602 |
| K02995 small subunit ribosomal protein S8e | 1 | 0 | 2 | 0 | 0 | 0 | 0 | 0 | 0 | 2 | 0 | 6 |
| K02996 small subunit ribosomal protein S9 | 1533 | 1417 | 1460 | 1452 | 1554 | 1528 | 1650 | 1521 | 1498 | 1496 | 1456 | 1573 |
| K07590 large subunit ribosomal protein L7A | 14 | 18 | 17 | 19 | 16 | 8 | 18 | 9 | 24 | 26 | 22 | 29 |
| ko00970 Aminoacyl-tRNA biosynthesis | 142167 | 143005 | 144509 | 144727 | 143932 | 146276 | 144845 | 143047 | 144532 | 146340 | 145552 | 145840 |
| K01866 tyrosyl-tRNA synthetase [EC:6.1.1.1] | 6742 | 6605 | 6727 | 6974 | 6388 | 6321 | 6459 | 6334 | 5664 | 5665 | 5753 | 5770 |
| K01867 tryptophanyl-tRNA synthetase [EC:6.1.1.2] | 3614 | 3622 | 3676 | 3711 | 3792 | 4023 | 4044 | 4030 | 4428 | 4742 | 4735 | 4608 |
| K01868 threonyl-tRNA synthetase [EC:6.1.1.3] | 6660 | 6655 | 7015 | 6994 | 6974 | 6942 | 6936 | 6609 | 7203 | 7560 | 7247 | 7437 |
| K01869 leucyl-tRNA synthetase [EC:6.1.1.4] | 9583 | 9582 | 9705 | 9565 | 10668 | 10848 | 10592 | 10552 | 10301 | 10537 | 10280 | 10063 |
| K01870 isoleucyl-tRNA synthetase [EC:6.1.1.5] | 10289 | 10379 | 10980 | 10790 | 11304 | 11532 | 11439 | 11340 | 10797 | 10641 | 10720 | 10544 |
| K01872 alanyl-tRNA synthetase [EC:6.1.1.7] | 9066 | 9219 | 9137 | 9110 | 9633 | 9766 | 9737 | 9506 | 9083 | 9105 | 9340 | 9507 |
| K01873 valyl-tRNA synthetase [EC:6.1.1.9] | 9179 | 8897 | 9059 | 9315 | 7199 | 7385 | 7188 | 7018 | 7897 | 7971 | 7694 | 8077 |
| K01875 seryl-tRNA synthetase [EC:6.1.1.11] | 4363 | 4409 | 4466 | 4443 | 4674 | 4740 | 4702 | 4674 | 4469 | 4824 | 4691 | 4769 |
| K01876 aspartyl-tRNA synthetase [EC:6.1.1.12] | 5602 | 5653 | 5659 | 5922 | 4487 | 4708 | 4525 | 4545 | 4629 | 4751 | 4821 | 4732 |
| K01878 glycyl-tRNA synthetase alpha chain [EC:6.1.1.14] | 2497 | 2510 | 2409 | 2488 | 1531 | 1499 | 1458 | 1493 | 1542 | 1541 | 1603 | 1523 |
| K01879 glycyl-tRNA synthetase beta chain [EC:6.1.1.14] | 5273 | 5390 | 5062 | 5278 | 3211 | 3377 | 3074 | 3328 | 3368 | 3520 | 3394 | 3562 |
| K01880 glycyl-tRNA synthetase [EC:6.1.1.14] | 2009 | 1953 | 2101 | 2054 | 3947 | 4053 | 3875 | 3831 | 3549 | 3258 | 3509 | 3352 |
| K01881 prolyl-tRNA synthetase [EC:6.1.1.15] | 5371 | 5276 | 5451 | 5584 | 4901 | 5020 | 4916 | 4819 | 5372 | 5524 | 5216 | 5467 |
| K01883 cysteinyl-tRNA synthetase [EC:6.1.1.16] | 4181 | 4270 | 4450 | 4306 | 5064 | 5226 | 4883 | 5016 | 4914 | 4708 | 4919 | 4899 |
| K01886 glutaminyl-tRNA synthetase [EC:6.1.1.18] | 5316 | 5378 | 5480 | 5444 | 5606 | 5793 | 5776 | 5569 | 5269 | 5325 | 5470 | 5239 |
| K01887 arginyl-tRNA synthetase [EC:6.1.1.19] | 6214 | 6256 | 6241 | 6249 | 7091 | 7075 | 6708 | 6915 | 6311 | 6389 | 6285 | 6315 |
| K01889 phenylalanyl-tRNA synthetase alpha chain [EC:6.1.1.20] | 3043 | 3223 | 3267 | 3119 | 3450 | 3369 | 3418 | 3400 | 3250 | 3406 | 3288 | 3315 |
| K01890 phenylalanyl-tRNA synthetase beta chain [EC:6.1.1.20] | 7083 | 6995 | 7352 | 7149 | 7961 | 7927 | 7809 | 7840 | 7446 | 7653 | 7654 | 7474 |
| K01892 histidyl-tRNA synthetase [EC:6.1.1.21] | 4269 | 4468 | 4612 | 4493 | 4784 | 4923 | 4876 | 4876 | 4404 | 4574 | 4406 | 4337 |
| K01893 asparaginyl-tRNA synthetase [EC:6.1.1.22] | 2827 | 2850 | 2953 | 2865 | 4053 | 4108 | 4264 | 3926 | 4302 | 4435 | 4302 | 4279 |
| K02433 aspartyl-tRNA(Asn)/glutamyl-tRNA(Gln) amidotransferase subunit A [EC:6.3.5.6 6.3.5.7] | 3038 | 2913 | 2942 | 2974 | 1487 | 1489 | 1511 | 1571 | 1644 | 1777 | 1588 | 1823 |
| K02434 aspartyl-tRNA(Asn)/glutamyl-tRNA(Gln) amidotransferase subunit B [EC:6.3.5.6 6.3.5.7] | 1873 | 1908 | 1765 | 1871 | 1007 | 978 | 1039 | 1096 | 1222 | 1275 | 1235 | 1285 |
| K02435 aspartyl-tRNA(Asn)/glutamyl-tRNA(Gln) amidotransferase subunit C [EC:6.3.5.6 6.3.5.7] | 572 | 584 | 589 | 576 | 335 | 312 | 324 | 307 | 320 | 347 | 330 | 354 |
| K03330 glutamyl-tRNA(Gln) amidotransferase subunit E [EC:6.3.5.7] | 2 | 2 | 3 | 0 | 0 | 4 | 3 | 2 | 18 | 16 | 19 | 14 |
| K04566 lysyl-tRNA synthetase, class I [EC:6.1.1.6] | 70 | 55 | 78 | 80 | 17 | 23 | 13 | 19 | 119 | 180 | 138 | 160 |
| K04567 lysyl-tRNA synthetase, class II [EC:6.1.1.6] | 5796 | 5870 | 5768 | 5831 | 6175 | 6320 | 6497 | 6310 | 7141 | 6987 | 7010 | 7076 |
| K09482 glutamyl-tRNA(Gln) amidotransferase subunit D [EC:6.3.5.7] | 2 | 0 | 0 | 0 | 0 | 1 | 0 | 0 | 13 | 14 | 13 | 8 |
| K09698 nondiscriminating glutamyl-tRNA synthetase [EC:6.1.1.24] | 171 | 178 | 159 | 173 | 57 | 59 | 57 | 53 | 235 | 263 | 279 | 268 |
| K09759 nondiscriminating aspartyl-tRNA synthetase [EC:6.1.1.23] | 501 | 406 | 439 | 455 | 126 | 109 | 108 | 119 | 607 | 529 | 580 | 606 |
| K11627 pyrrolysyl-tRNA synthetase [EC:6.1.1.26] | 0 | 0 | 0 | 0 | 0 | 0 | 0 | 0 | 0 | 0 | 0 | 1 |
| K14164 glycyl-tRNA synthetase [EC:6.1.1.14] | 0 | 0 | 0 | 0 | 1 | 0 | 0 | 0 | 0 | 0 | 0 | 0 |
| ko03013 RNA transport | 6412 | 6540 | 6301 | 6264 | 5983 | 5863 | 5904 | 5942 | 6316 | 6304 | 6250 | 6427 |
| K00784 ribonuclease Z [EC:3.1.26.11] | 1950 | 1994 | 1984 | 1970 | 2651 | 2610 | 2667 | 2549 | 3011 | 2983 | 2929 | 3015 |
| K00974 tRNA nucleotidyltransferase (CCA-adding enzyme) [EC:2.7.7.72 3.1.3.- 3.1.4.-] | 3378 | 3396 | 3247 | 3271 | 2040 | 1991 | 1921 | 2029 | 2262 | 2277 | 2269 | 2303 |
| K03113 translation initiation factor 1 | 1068 | 1131 | 1058 | 1015 | 1286 | 1257 | 1307 | 1356 | 1015 | 975 | 1021 | 1049 |
| K03231 elongation factor 1-alpha | 0 | 3 | 3 | 0 | 2 | 0 | 0 | 0 | 2 | 11 | 3 | 12 |
| K03236 translation initiation factor 1A | 0 | 0 | 1 | 0 | 0 | 0 | 0 | 0 | 2 | 1 | 3 | 5 |
| K03237 translation initiation factor 2 subunit 1 | 0 | 0 | 0 | 1 | 0 | 1 | 2 | 0 | 4 | 1 | 0 | 4 |
| K03238 translation initiation factor 2 subunit 2 | 0 | 0 | 0 | 0 | 0 | 2 | 0 | 1 | 3 | 1 | 3 | 3 |
| K03239 translation initiation factor eIF-2B subunit alpha | 8 | 9 | 5 | 4 | 1 | 0 | 2 | 2 | 2 | 4 | 4 | 5 |
| K03242 translation initiation factor 2 subunit 3 | 2 | 0 | 2 | 0 | 1 | 0 | 2 | 0 | 10 | 11 | 7 | 9 |
| K03243 translation initiation factor 5B | 2 | 2 | 1 | 1 | 2 | 0 | 0 | 2 | 1 | 16 | 3 | 12 |
| K03251 translation initiation factor 3 subunit D | 0 | 0 | 0 | 2 | 0 | 0 | 0 | 0 | 0 | 0 | 0 | 0 |
| K03537 ribonuclease P/MRP protein subunit POP5 [EC:3.1.26.5] | 0 | 0 | 0 | 0 | 0 | 0 | 0 | 0 | 0 | 4 | 0 | 1 |
| K03538 ribonuclease P protein subunit POP4 [EC:3.1.26.5] | 0 | 0 | 0 | 0 | 0 | 0 | 0 | 0 | 0 | 1 | 0 | 0 |
| K03539 ribonuclease P/MRP protein subunit RPP1 [EC:3.1.26.5] | 2 | 0 | 0 | 0 | 0 | 0 | 0 | 0 | 0 | 6 | 2 | 1 |
| K03540 ribonuclease P protein subunit RPR2 [EC:3.1.26.5] | 2 | 1 | 0 | 0 | 0 | 0 | 0 | 0 | 0 | 3 | 0 | 1 |
| K07562 nonsense-mediated mRNA decay protein 3 | 0 | 2 | 0 | 0 | 0 | 1 | 1 | 2 | 4 | 9 | 6 | 6 |
| K07936 GTP-binding nuclear protein Ran | 0 | 0 | 0 | 0 | 0 | 0 | 0 | 1 | 0 | 0 | 0 | 0 |
| K13126 polyadenylate-binding protein | 0 | 0 | 0 | 0 | 0 | 0 | 2 | 0 | 0 | 0 | 0 | 1 |
| K14288 exportin-T | 0 | 2 | 0 | 0 | 0 | 0 | 0 | 0 | 0 | 0 | 0 | 0 |
| K14293 importin subunit beta-1 | 0 | 0 | 0 | 0 | 0 | 0 | 0 | 0 | 0 | 1 | 0 | 0 |
| K14296 nuclear pore complex protein Nup153 | 0 | 0 | 0 | 0 | 0 | 1 | 0 | 0 | 0 | 0 | 0 | 0 |
| ko03015 mRNA surveillance pathway | 0 | 1 | 5 | 1 | 0 | 2 | 7 | 3 | 5 | 14 | 8 | 15 |
| K03265 peptide chain release factor subunit 1 | 0 | 0 | 1 | 1 | 0 | 0 | 2 | 3 | 5 | 13 | 3 | 9 |
| K06269 serine/threonine-protein phosphatase PP1 catalytic subunit [EC:3.1.3.16] | 0 | 0 | 0 | 0 | 0 | 2 | 0 | 0 | 0 | 0 | 0 | 1 |
| K06965 protein pelota | 0 | 0 | 2 | 0 | 0 | 0 | 3 | 0 | 0 | 1 | 3 | 4 |
| K14403 cleavage and polyadenylation specificity factor subunit 3 [EC:3.1.27.-] | 0 | 0 | 2 | 0 | 0 | 0 | 0 | 0 | 0 | 0 | 0 | 0 |
| K14408 cleavage stimulation factor subunit 3 | 0 | 0 | 0 | 0 | 0 | 0 | 0 | 0 | 0 | 0 | 2 | 0 |

| | | | | | | | | | | | | |
|---|---|---|---|---|---|---|---|---|---|---|---|---|
| K15544 RNA polymerase II subunit A C-terminal domain phosphatase SSU72 [EC:3.1.3.16] | 0 | 1 | 0 | 0 | 0 | 0 | 0 | 0 | 0 | 0 | 0 | 0 |
| ko03008 Ribosome biogenesis in eukaryotes | 4458 | 4574 | 4392 | 4658 | 4420 | 4395 | 4203 | 4494 | 4083 | 4013 | 4087 | 3976 |
| K03264 translation initiation factor 6 | 4 | 2 | 0 | 0 | 0 | 0 | 2 | 0 | 5 | 10 | 5 | 6 |
| K03685 ribonuclease III [EC:3.1.26.3] | 2267 | 2346 | 2291 | 2492 | 3097 | 3097 | 3018 | 3174 | 2861 | 2793 | 2859 | 2738 |
| K06943 nucleolar GTP-binding protein | 0 | 2 | 0 | 1 | 2 | 0 | 0 | 0 | 2 | 7 | 1 | 9 |
| K07178 RIO kinase 1 [EC:2.7.11.1] | 1020 | 1090 | 964 | 1024 | 567 | 572 | 522 | 600 | 361 | 339 | 346 | 318 |
| K11130 H/ACA ribonucleoprotein complex subunit 3 | 0 | 0 | 0 | 0 | 0 | 1 | 0 | 1 | 0 | 0 | 2 | 1 |
| K11131 H/ACA ribonucleoprotein complex subunit 4 [EC:5.4.99.-] | 0 | 0 | 0 | 0 | 0 | 0 | 2 | 0 | 0 | 14 | 5 | 9 |
| K13288 oligoribonuclease [EC:3.1.-.-] | 1165 | 1127 | 1135 | 1141 | 747 | 718 | 656 | 712 | 844 | 816 | 854 | 864 |
| K14536 ribosome assembly protein 1 [EC:3.6.5.-] | 0 | 0 | 0 | 0 | 2 | 0 | 0 | 0 | 0 | 0 | 2 | 0 |
| K14538 nuclear GTP-binding protein | 0 | 0 | 2 | 0 | 0 | 0 | 0 | 0 | 0 | 0 | 0 | 0 |
| K14550 U3 small nucleolar RNA-associated protein 10 | 0 | 0 | 0 | 0 | 3 | 0 | 0 | 0 | 0 | 0 | 0 | 0 |
| K14558 periodic tryptophan protein 2 | 0 | 0 | 0 | 0 | 0 | 1 | 0 | 0 | 0 | 0 | 0 | 0 |
| K14561 U3 small nucleolar ribonucleoprotein protein IMP4 | 0 | 0 | 0 | 0 | 0 | 0 | 0 | 0 | 0 | 4 | 0 | 3 |
| K14564 nucleolar protein 56 | 0 | 5 | 0 | 0 | 0 | 2 | 0 | 2 | 4 | 4 | 0 | 4 |
| K14572 midasin | 0 | 0 | 0 | 0 | 0 | 1 | 0 | 0 | 0 | 0 | 0 | 1 |
| K14574 ribosome maturation protein SDO1 | 0 | 0 | 0 | 0 | 0 | 2 | 2 | 2 | 0 | 2 | 5 | 5 |
| Folding, sorting and degradation | 174908 | 175003 | 177026 | 175944 | 193526 | 195858 | 195289 | 193074 | 181441 | 180503 | 180689 | 179546 |
| ko03060 Protein export | 53866 | 54160 | 54942 | 54304 | 61801 | 62256 | 62358 | 61735 | 57576 | 56944 | 56998 | 55828 |
| K03070 preprotein translocase subunit SecA [EC:7.4.2.8] | 10636 | 10483 | 10919 | 10637 | 11748 | 12054 | 12206 | 12189 | 11170 | 11026 | 10733 | 10698 |
| K03071 preprotein translocase subunit SecB | 1175 | 1157 | 1179 | 1120 | 702 | 770 | 702 | 752 | 698 | 794 | 706 | 760 |
| K03072 preprotein translocase subunit SecD | 4397 | 4244 | 4335 | 4240 | 2718 | 2657 | 2590 | 2699 | 2659 | 2863 | 2592 | 2701 |
| K03073 preprotein translocase subunit SecE | 1019 | 1096 | 1075 | 1013 | 1008 | 1037 | 1073 | 929 | 913 | 986 | 980 | 942 |
| K03074 preprotein translocase subunit SecF | 2190 | 2071 | 2004 | 2046 | 1228 | 1245 | 1263 | 1258 | 1487 | 1553 | 1511 | 1455 |
| K03075 preprotein translocase subunit SecG | 1123 | 1199 | 1200 | 1236 | 1332 | 1245 | 1054 | 1023 | 1139 | 1098 | 1089 | 1151 |
| K03076 preprotein translocase subunit SecY | 3699 | 3992 | 4003 | 4120 | 4596 | 4977 | 4909 | 4691 | 4737 | 4881 | 4778 | 4751 |
| K03100 signal peptidase I [EC:3.4.21.89] | 5225 | 5239 | 5296 | 5203 | 8286 | 8407 | 8553 | 8605 | 7011 | 6772 | 7071 | 6586 |
| K03101 signal peptidase II [EC:3.4.23.36] | 2010 | 2059 | 2083 | 2088 | 2159 | 2103 | 2138 | 2050 | 1992 | 1913 | 2007 | 1894 |
| K03105 signal recognition particle subunit SRP19 | 0 | 0 | 0 | 2 | 2 | 0 | 1 | 0 | 0 | 0 | 0 | 4 |
| K03106 signal recognition particle subunit SRP54 [EC:3.6.5.4] | 4524 | 4669 | 4614 | 4627 | 4994 | 4932 | 4883 | 4805 | 4624 | 4591 | 4572 | 4428 |
| K03107 signal recognition particle subunit SRP68 | 0 | 0 | 2 | 0 | 0 | 0 | 0 | 0 | 0 | 0 | 0 | 0 |
| K03110 fused signal recognition particle receptor | 3300 | 3317 | 3191 | 3120 | 3400 | 3219 | 3550 | 3550 | 3295 | 3072 | 3238 | 3093 |
| K03116 sec-independent protein translocase protein TatA | 693 | 696 | 691 | 670 | 638 | 648 | 615 | 694 | 604 | 582 | 595 | 563 |
| K03117 sec-independent protein translocase protein TatB | 764 | 676 | 724 | 677 | 504 | 557 | 489 | 572 | 548 | 474 | 499 | 490 |
| K03118 sec-independent protein translocase protein TatC | 2545 | 2535 | 2586 | 2472 | 3282 | 3200 | 3104 | 3217 | 2643 | 2539 | 2553 | 2516 |
| K03210 preprotein translocase subunit YajC | 1080 | 1137 | 1098 | 1124 | 1096 | 1105 | 994 | 1023 | 1057 | 1035 | 1019 | 1118 |
| K03217 YidC/Oxa1 family membrane protein insertase | 5842 | 5725 | 5902 | 6007 | 6328 | 6366 | 6404 | 6031 | 5969 | 5965 | 5895 | 5903 |
| K03425 sec-independent protein translocase protein TatE | 81 | 104 | 97 | 102 | 80 | 81 | 66 | 60 | 131 | 125 | 121 | 141 |
| K09490 heat shock 70kDa protein 5 | 0 | 1 | 0 | 0 | 0 | 0 | 0 | 0 | 0 | 1 | 0 | 0 |
| K12257 SecD/SecF fusion protein | 3134 | 3331 | 3515 | 3330 | 7399 | 7358 | 7463 | 7290 | 6275 | 5974 | 6345 | 5940 |
| K13280 signal peptidase I [EC:3.4.21.89] | 95 | 107 | 127 | 100 | 72 | 60 | 60 | 76 | 163 | 172 | 215 | 196 |
| K13301 secretion monitor | 334 | 322 | 301 | 370 | 229 | 235 | 241 | 221 | 459 | 528 | 479 | 498 |
| K13431 signal recognition particle receptor subunit alpha | 0 | 0 | 0 | 0 | 0 | 0 | 0 | 0 | 2 | 0 | 0 | 0 |
| ko04141 Protein processing in endoplasmic reticulum | 8412 | 8499 | 8701 | 8658 | 12448 | 12451 | 12587 | 12521 | 10096 | 10188 | 10311 | 10445 |
| K04079 molecular chaperone HtpG | 8026 | 8029 | 8183 | 8223 | 11366 | 11368 | 11453 | 11419 | 9422 | 9406 | 9524 | 9661 |
| K08852 serine/threonine-protein kinase/endoribonuclease IRE1 [EC:2.7.11.1 3.1.26.-] | 0 | 0 | 0 | 0 | 0 | 0 | 0 | 0 | 0 | 0 | 1 | 0 |
| K09487 heat shock protein 90kDa beta | 1 | 0 | 0 | 0 | 0 | 0 | 0 | 0 | 0 | 0 | 0 | 0 |
| K09523 DnaJ homolog subfamily C member 3 | 21 | 18 | 12 | 25 | 61 | 50 | 56 | 49 | 42 | 49 | 37 | 36 |
| K13525 transitional endoplasmic reticulum ATPase | 3 | 2 | 1 | 1 | 1 | 5 | 4 | 0 | 4 | 16 | 19 | 27 |
| K13993 HSP20 family protein | 341 | 426 | 477 | 387 | 1017 | 1023 | 1068 | 1052 | 603 | 665 | 686 | 678 |
| K13996 protein disulfide-isomerase [EC:5.3.4.1] | 0 | 0 | 0 | 0 | 0 | 0 | 0 | 0 | 0 | 0 | 0 | 1 |
| K14002 DnaJ-related protein SCJ1 | 0 | 0 | 0 | 0 | 0 | 0 | 0 | 0 | 0 | 0 | 1 | 0 |
| K14005 protein transport protein SEC31 | 0 | 0 | 0 | 0 | 0 | 0 | 0 | 0 | 2 | 0 | 0 | 0 |
| K14006 protein transport protein SEC23 | 0 | 0 | 0 | 0 | 0 | 0 | 0 | 0 | 0 | 0 | 3 | 0 |
| K14007 protein transport protein SEC24 | 0 | 2 | 0 | 0 | 0 | 0 | 0 | 0 | 0 | 0 | 0 | 0 |
| K14026 SEL1 protein | 0 | 0 | 0 | 0 | 0 | 0 | 0 | 0 | 0 | 1 | 0 | 0 |
| K16196 eukaryotic translation initiation factor 2-alpha kinase 4 [EC:2.7.11.1] | 0 | 0 | 0 | 0 | 0 | 0 | 0 | 0 | 0 | 1 | 0 | 0 |
| ko04130 SNARE interactions in vesicular transport | 0 | 0 | 0 | 0 | 0 | 0 | 0 | 0 | 0 | 0 | 0 | 1 |
| K08517 vesicle transport protein SEC22 | 0 | 0 | 0 | 0 | 0 | 0 | 0 | 0 | 0 | 0 | 0 | 1 |
| ko04120 Ubiquitin mediated proteolysis | 2 | 4 | 6 | 2 | 0 | 0 | 0 | 0 | 0 | 3 | 5 | 1 |
| K03178 ubiquitin-activating enzyme E1 [EC:6.2.1.45] | 0 | 0 | 0 | 0 | 0 | 0 | 0 | 0 | 0 | 0 | 1 | 0 |
| K03349 anaphase-promoting complex subunit 2 | 0 | 1 | 0 | 0 | 0 | 0 | 0 | 0 | 0 | 0 | 0 | 0 |
| K03355 anaphase-promoting complex subunit 8 | 0 | 0 | 1 | 0 | 0 | 0 | 0 | 0 | 0 | 0 | 0 | 0 |
| K03364 cell division cycle 20-like protein 1, cofactor of APC complex | 0 | 0 | 0 | 0 | 0 | 0 | 0 | 0 | 0 | 2 | 0 | 0 |
| K05633 NEDD4-like E3 ubiquitin-protein ligase WWP1 [EC:2.3.2.26] | 0 | 0 | 2 | 0 | 0 | 0 | 0 | 0 | 0 | 0 | 0 | 0 |
| K10595 E3 ubiquitin-protein ligase HERC2 [EC:2.3.2.26] | 1 | 0 | 2 | 0 | 0 | 0 | 0 | 0 | 0 | 0 | 3 | 0 |
| K10614 E3 ubiquitin-protein ligase HERC3 [EC:2.3.2.26] | 0 | 1 | 1 | 1 | 0 | 0 | 0 | 0 | 0 | 1 | 0 | 1 |
| K10615 E3 ubiquitin-protein ligase HERC4 [EC:2.3.2.26] | 1 | 2 | 0 | 1 | 0 | 0 | 0 | 0 | 0 | 0 | 0 | 0 |
| K10685 ubiquitin-like 1-activating enzyme E1 B [EC:6.2.1.45] | 0 | 0 | 0 | 0 | 0 | 0 | 0 | 0 | 0 | 0 | 1 | 0 |
| ko04122 Sulfur relay system | 30434 | 29942 | 29174 | 29910 | 24195 | 24700 | 24441 | 24495 | 25246 | 24898 | 24555 | 25199 |
| K00566 tRNA-uridine 2-sulfurtransferase [EC:2.8.1.13] | 4897 | 5091 | 5183 | 5058 | 8673 | 8651 | 8798 | 8422 | 7448 | 7019 | 7170 | 7094 |
| K03154 sulfur carrier protein | 598 | 599 | 514 | 556 | 586 | 613 | 750 | 622 | 512 | 505 | 540 | 528 |
| K03636 sulfur-carrier protein | 598 | 593 | 581 | 606 | 322 | 319 | 345 | 372 | 380 | 412 | 370 | 387 |
| K04085 tRNA 2-thiouridine synthesizing protein A [EC:2.8.1.-] | 569 | 582 | 607 | 550 | 336 | 308 | 308 | 339 | 388 | 398 | 378 | 385 |
| K07235 tRNA 2-thiouridine synthesizing protein D [EC:2.8.1.-] | 350 | 364 | 353 | 346 | 197 | 230 | 235 | 182 | 399 | 384 | 411 | 415 |
| K07236 tRNA 2-thiouridine synthesizing protein C | 763 | 722 | 685 | 705 | 424 | 432 | 431 | 435 | 432 | 434 | 411 | 531 |
| K07237 tRNA 2-thiouridine synthesizing protein B | 615 | 633 | 577 | 560 | 361 | 403 | 351 | 374 | 381 | 352 | 344 | 356 |
| K11179 tRNA 2-thiouridine synthesizing protein E [EC:2.8.1.-] | 645 | 631 | 565 | 639 | 401 | 372 | 399 | 439 | 356 | 314 | 347 | 383 |
| K11996 adenylyltransferase and sulfurtransferase [EC:2.7.7.80 2.8.1.11] | 3 | 9 | 5 | 7 | 1 | 1 | 0 | 0 | 7 | 1 | 4 | 1 |
| K21028 molybdopterin synthase sulfurtransferase [EC:2.8.1.11] | 795 | 811 | 622 | 671 | 418 | 490 | 439 | 381 | 1082 | 1073 | 934 | 1009 |
| K21029 molybdopterin-synthase adenylyltransferase [EC:2.7.7.80] | 1608 | 1680 | 1595 | 1721 | 1939 | 2114 | 2095 | 2081 | 1714 | 1598 | 1764 | 1618 |
| K21140 [CysO sulfur-carrier protein]-S-L-cysteine hydrolase [EC:3.13.1.6] | 5 | 12 | 22 | 1 | 11 | 0 | 0 | 0 | 0 | 0 | 10 | 0 |
| K21147 sulfur-carrier protein adenylyltransferase/sulfurtransferase [EC:2.7.7.80 2.7.7.- 2.8.1.11 2.8.1.-] | 5 | 9 | 5 | 7 | 8 | 11 | 4 | 11 | 9 | 10 | 5 | 14 |
| ko03050 Proteasome | 4 | 7 | 4 | 5 | 4 | 5 | 0 | 7 | 22 | 23 | 23 | 31 |
| K03039 26S proteasome regulatory subunit N9 | 0 | 1 | 0 | 0 | 0 | 0 | 0 | 0 | 0 | 0 | 0 | 0 |
| K03063 26S proteasome regulatory subunit T3 | 0 | 0 | 0 | 0 | 0 | 0 | 0 | 0 | 1 | 0 | 1 | 0 |
| K03420 proteasome regulatory subunit | 0 | 2 | 2 | 0 | 0 | 0 | 0 | 0 | 2 | 3 | 8 | 2 |
| K03432 proteasome alpha subunit [EC:3.4.25.1] | 0 | 0 | 0 | 1 | 2 | 1 | 0 | 0 | 6 | 9 | 1 | 6 |
| K03433 proteasome beta subunit [EC:3.4.25.1] | 1 | 0 | 0 | 0 | 0 | 0 | 0 | 0 | 6 | 1 | 7 | 3 |
| K13527 proteasome-associated ATPase | 3 | 4 | 2 | 4 | 2 | 4 | 0 | 7 | 7 | 10 | 6 | 20 |
| ko03018 RNA degradation | 82190 | 82392 | 84199 | 83065 | 95078 | 96446 | 95903 | 94316 | 88501 | 88448 | 88797 | 88041 |
| K00962 polyribonucleotide nucleotidyltransferase [EC:2.7.7.8] | 7679 | 7904 | 8044 | 8058 | 8614 | 8710 | 8825 | 8374 | 8209 | 8316 | 7938 | 8093 |
| K00970 poly(A) polymerase [EC:2.7.7.19] | 3405 | 3310 | 3282 | 3277 | 2105 | 2040 | 2088 | 2169 | 2083 | 2185 | 1965 | 2123 |
| K03628 transcription termination factor Rho | 3808 | 3769 | 4105 | 4006 | 4689 | 4575 | 5157 | 5110 | 4393 | 4267 | 4525 | 4306 |
| K03654 ATP-dependent DNA helicase RecQ [EC:3.6.4.12] | 9850 | 9595 | 10348 | 9987 | 18898 | 18974 | 19017 | 18434 | 15542 | 14692 | 15454 | 14979 |
| K03666 host factor-I protein | 541 | 586 | 606 | 538 | 415 | 376 | 374 | 360 | 325 | 429 | 373 | 382 |
| K03679 exosome complex component RRP4 | 0 | 2 | 0 | 0 | 0 | 0 | 0 | 0 | 2 | 2 | 3 | 11 |
| K03732 ATP-dependent RNA helicase RhlB [EC:3.6.4.13] | 3058 | 2982 | 2950 | 2932 | 1795 | 1837 | 1830 | 1735 | 1790 | 1813 | 1701 | 1762 |
| K04043 molecular chaperone DnaK | 6782 | 6847 | 7192 | 6984 | 7162 | 7302 | 7013 | 6897 | 6986 | 7299 | 7201 | 6902 |
| K04077 chaperonin GroEL | 4258 | 4169 | 4478 | 4105 | 5113 | 5245 | 4692 | 4639 | 5013 | 5069 | 5107 | 4892 |
| K05592 ATP-dependent RNA helicase DeaD [EC:3.6.4.13] | 6496 | 6599 | 6481 | 6510 | 6860 | 7077 | 7188 | 7071 | 7441 | 7488 | 7184 | 7423 |
| K07573 exosome complex component CSL4 | 0 | 0 | 0 | 1 | 0 | 0 | 0 | 0 | 5 | 8 | 9 | 3 |
| K08300 ribonuclease E [EC:3.1.26.12] | 2774 | 2841 | 2769 | 2880 | 1795 | 1858 | 1938 | 1914 | 2096 | 2241 | 2152 | 2136 |
| K08311 putative (di)nucleoside polyphosphate hydrolase [EC:3.6.1.-] | 1125 | 1054 | 1078 | 1139 | 668 | 725 | 649 | 658 | 698 | 743 | 733 | 740 |
| K11600 exosome complex component RRP41 | 0 | 0 | 0 | 0 | 0 | 1 | 3 | 0 | 4 | 4 | 4 | 1 |
| K11927 ATP-dependent RNA helicase RhlE [EC:3.6.4.13] | 3911 | 4039 | 3857 | 4050 | 4521 | 4517 | 4339 | 4461 | 3678 | 3646 | 3848 | 3741 |
| K12573 ribonuclease R [EC:3.1.13.1] | 8142 | 8077 | 8224 | 8081 | 8966 | 8956 | 9063 | 8952 | 7979 | 8218 | 8282 | 8047 |
| K12574 ribonuclease J [EC:3.1.-.-] | 1906 | 1968 | 2014 | 1927 | 3270 | 3316 | 3342 | 3348 | 3590 | 3746 | 3737 | 3665 |
| K12589 exosome complex component RRP42 | 0 | 0 | 0 | 0 | 0 | 5 | 0 | 2 | 4 | 5 | 5 | 1 |
| K12601 superkiller protein 8 | 0 | 0 | 0 | 0 | 1 | 0 | 0 | 0 | 0 | 0 | 0 | 0 |
| K12614 ATP-dependent RNA helicase DDX6/DHH1 [EC:3.6.4.13] | 0 | 0 | 2 | 0 | 0 | 0 | 0 | 0 | 0 | 0 | 0 | 0 |
| K18682 ribonucrease Y [EC:3.1.-.-] | 1753 | 1806 | 1919 | 1803 | 3248 | 3262 | 3137 | 3079 | 2807 | 2799 | 2937 | 2784 |
| Replication and repair | 254371 | 252794 | 253000 | 254176 | 258522 | 260314 | 260498 | 260440 | 256807 | 257921 | 259237 | 258094 |
| ko03030 DNA replication | 78377 | 78897 | 80254 | 79572 | 91059 | 92029 | 92411 | 91046 | 85196 | 84381 | 85484 | 84684 |
| K01972 DNA ligase (NAD+) [EC:6.5.1.2] | 7294 | 7394 | 7403 | 7548 | 7446 | 7596 | 7788 | 7460 | 7698 | 8173 | 8061 | 8046 |
| K02314 replicative DNA helicase [EC:3.6.4.12] | 12109 | 12288 | 12651 | 12770 | 18488 | 18486 | 18875 | 18597 | 13804 | 13601 | 13721 | 13307 |
| K02316 DNA primase [EC:2.7.7.101] | 6773 | 6889 | 6719 | 6761 | 7903 | 7819 | 8041 | 8151 | 7274 | 6722 | 7058 | 7247 |
| K02319 DNA polymerase, archaea type [EC:2.7.7.7] | 0 | 2 | 2 | 0 | 2 | 0 | 2 | 6 | 14 | 30 | 12 | 18 |
| K02322 DNA polymerase II large subunit [EC:2.7.7.7] | 5 | 2 | 0 | 3 | 2 | 0 | 2 | 7 | 9 | 34 | 18 | 34 |
| K02323 DNA polymerase II small subunit [EC:2.7.7.7] | 3 | 0 | 0 | 1 | 0 | 0 | 0 | 0 | 3 | 8 | 8 | 17 |
| K02335 DNA polymerase I [EC:2.7.7.7] | 10609 | 10694 | 10868 | 10636 | 11635 | 11745 | 11911 | 11697 | 10668 | 11017 | 11010 | 10671 |

| | | | | | | | | | | | | |
|---|---|---|---|---|---|---|---|---|---|---|---|---|
| K02337 DNA polymerase III subunit alpha [EC:2.7.7.7] | 12304 | 12500 | 12857 | 12459 | 14834 | 14955 | 14807 | 14744 | 13683 | 13284 | 13590 | 13450 |
| K02338 DNA polymerase III subunit beta [EC:2.7.7.7] | 3544 | 3711 | 3902 | 3862 | 3852 | 3894 | 3772 | 3616 | 4045 | 4105 | 4010 | 4060 |
| K02339 DNA polymerase III subunit chi [EC:2.7.7.7] | 1024 | 964 | 913 | 1029 | 606 | 630 | 545 | 670 | 717 | 679 | 574 | 583 |
| K02340 DNA polymerase III subunit delta [EC:2.7.7.7] | 3246 | 3131 | 3203 | 3116 | 3335 | 3431 | 3507 | 3327 | 3331 | 3048 | 3351 | 3205 |
| K02341 DNA polymerase III subunit delta' [EC:2.7.7.7] | 3618 | 3562 | 3561 | 3527 | 3787 | 3761 | 3873 | 3861 | 3515 | 3566 | 3591 | 3736 |
| K02342 DNA polymerase III subunit epsilon [EC:2.7.7.7] | 3123 | 3318 | 3288 | 3223 | 4529 | 4563 | 4564 | 4333 | 3484 | 3250 | 3467 | 3476 |
| K02343 DNA polymerase III subunit gamma/tau [EC:2.7.7.7] | 5536 | 5208 | 5411 | 5293 | 5960 | 6163 | 5981 | 5726 | 5693 | 5579 | 5643 | 5520 |
| K02344 DNA polymerase III subunit psi [EC:2.7.7.7] | 289 | 255 | 234 | 275 | 161 | 226 | 180 | 208 | 297 | 340 | 346 | 374 |
| K02345 DNA polymerase III subunit theta [EC:2.7.7.7] | 121 | 128 | 142 | 134 | 92 | 95 | 84 | 89 | 199 | 195 | 211 | 202 |
| K02683 DNA primase small subunit [EC:2.7.7.102] | 0 | 0 | 0 | 0 | 5 | 0 | 0 | 0 | 0 | 2 | 0 | 4 |
| K03111 single-strand DNA-binding protein | 2962 | 2949 | 3065 | 3127 | 3031 | 3075 | 3098 | 3041 | 3808 | 3804 | 3864 | 3883 |
| K03469 ribonuclease HI [EC:3.1.26.4] | 1788 | 1825 | 1880 | 1767 | 2550 | 2523 | 2523 | 2689 | 2240 | 2290 | 2234 | 2204 |
| K03470 ribonuclease HII [EC:3.1.26.4] | 1745 | 1797 | 1871 | 1735 | 2183 | 2349 | 2144 | 2191 | 2205 | 2053 | 2232 | 2062 |
| K03471 ribonuclease HIII [EC:3.1.26.4] | 75 | 76 | 78 | 74 | 13 | 27 | 19 | 12 | 112 | 105 | 83 | 103 |
| K03763 DNA polymerase III subunit alpha, Gram-positive type [EC:2.7.7.7] | 2192 | 2181 | 2196 | 2214 | 635 | 679 | 686 | 610 | 2336 | 2415 | 2351 | 2395 |
| K04799 flap endonuclease-1 [EC:3.-.-.-] | 3 | 5 | 0 | 3 | 0 | 0 | 2 | 0 | 9 | 12 | 0 | 11 |
| K04800 replication factor C large subunit | 2 | 0 | 2 | 0 | 5 | 5 | 0 | 2 | 9 | 12 | 9 | 11 |
| K04801 replication factor C small subunit | 3 | 0 | 2 | 1 | 0 | 0 | 2 | 2 | 7 | 9 | 4 | 12 |
| K04802 proliferating cell nuclear antigen | 1 | 0 | 0 | 1 | 1 | 0 | 0 | 1 | 2 | 4 | 3 | 4 |
| K07466 replication factor A1 | 0 | 2 | 0 | 0 | 2 | 0 | 5 | 0 | 11 | 10 | 8 | 14 |
| K10726 replicative DNA helicase Mcm [EC:3.6.4.-] | 3 | 5 | 2 | 3 | 2 | 5 | 0 | 3 | 5 | 8 | 7 | 11 |
| K10747 DNA ligase 1 [EC:6.5.1.1 6.5.1.6 6.5.1.7] | 2 | 9 | 0 | 3 | 0 | 2 | 2 | 0 | 14 | 16 | 10 | 11 |
| K10755 replication factor C subunit 2/4 | 0 | 0 | 0 | 0 | 0 | 0 | 0 | 0 | 0 | 0 | 2 | 0 |
| K18882 DNA primase large subunit | 2 | 0 | 2 | 3 | 0 | 0 | 0 | 3 | 4 | 6 | 3 | 12 |
| ko03410 Base excision repair | 44773 | 44916 | 44980 | 45088 | 43972 | 44468 | 44484 | 43947 | 44065 | 45365 | 45474 | 44824 |
| K01142 exodeoxyribonuclease III [EC:3.1.11.2] | 5333 | 5424 | 5434 | 5283 | 4344 | 4223 | 4228 | 4264 | 3684 | 3715 | 3665 | 3741 |
| K01151 deoxyribonuclease IV [EC:3.1.21.2] | 1465 | 1420 | 1481 | 1487 | 2137 | 2135 | 2176 | 2067 | 2180 | 2289 | 2399 | 2209 |
| K01246 DNA-3-methyladenine glycosylase I [EC:3.2.2.20] | 1589 | 1646 | 1593 | 1659 | 888 | 984 | 914 | 925 | 1197 | 1280 | 1193 | 1320 |
| K01247 DNA-3-methyladenine glycosylase II [EC:3.2.2.21] | 2246 | 2214 | 2037 | 2143 | 1385 | 1312 | 1266 | 1389 | 1281 | 1157 | 1230 | 1286 |
| K03575 A/G-specific adenine glycosylase [EC:3.2.2.31] | 2237 | 2158 | 2127 | 2192 | 2876 | 2750 | 2886 | 2942 | 2958 | 2982 | 2997 | 2932 |
| K03648 uracil-DNA glycosylase [EC:3.2.2.27] | 1481 | 1589 | 1653 | 1620 | 2033 | 1995 | 2067 | 2011 | 2024 | 2182 | 2269 | 2096 |
| K03649 double-stranded uracil-DNA glycosylase [EC:3.2.2.28] | 217 | 194 | 160 | 203 | 69 | 68 | 63 | 56 | 413 | 305 | 380 | 407 |
| K03652 DNA-3-methyladenine glycosylase [EC:3.2.2.21] | 1241 | 1110 | 1012 | 1146 | 624 | 708 | 600 | 628 | 372 | 402 | 381 | 410 |
| K03660 N-glycosylase/DNA lyase [EC:3.2.2.- 4.2.99.18] | 284 | 237 | 283 | 257 | 126 | 114 | 96 | 95 | 298 | 296 | 286 | 291 |
| K05522 endonuclease VIII [EC:3.2.2.- 4.2.99.18] | 439 | 478 | 464 | 488 | 243 | 333 | 289 | 269 | 638 | 669 | 650 | 633 |
| K07462 single-stranded-DNA-specific exonuclease [EC:3.1.-.-] | 5683 | 5786 | 5842 | 5684 | 6305 | 6696 | 6356 | 6466 | 6125 | 6051 | 6218 | 6025 |
| K10563 formamidopyrimidine-DNA glycosylase [EC:3.2.2.23 4.2.99.18] | 2623 | 2534 | 2466 | 2671 | 1660 | 1604 | 1544 | 1577 | 2024 | 2139 | 2013 | 2128 |
| K10771 AP endonuclease 1 [EC:4.2.99.18] | 0 | 0 | 0 | 0 | 0 | 0 | 0 | 2 | 0 | 0 | 0 | 0 |
| K10773 endonuclease III [EC:4.2.99.18] | 1858 | 1881 | 1994 | 1913 | 2106 | 2124 | 2200 | 2017 | 2231 | 2375 | 2409 | 2324 |
| K13529 AraC family transcriptional regulator, regulatory protein of adaptative response / DNA-3-methyladenine glycosylase II [EC:3.2.2.21] | 65 | 40 | 62 | 47 | 3 | 1 | 3 | 3 | 71 | 122 | 102 | 101 |
| K21929 uracil-DNA glycosylase [EC:3.2.2.27] | 103 | 103 | 101 | 104 | 91 | 78 | 93 | 78 | 178 | 179 | 198 | 178 |
| ko03420 Nucleotide excision repair | 68927 | 68654 | 69435 | 69165 | 72260 | 72411 | 73537 | 72615 | 75462 | 78424 | 77456 | 77000 |
| K03657 DNA helicase II / ATP-dependent DNA helicase PcrA [EC:3.6.4.12] | 13959 | 13587 | 13834 | 13842 | 12531 | 12381 | 12758 | 12375 | 18755 | 19996 | 19413 | 19434 |
| K03701 excinuclease ABC subunit A | 12837 | 12866 | 13045 | 12916 | 16770 | 16619 | 16982 | 17180 | 15194 | 15087 | 15327 | 15103 |
| K03702 excinuclease ABC subunit B | 6889 | 7033 | 6994 | 7125 | 6930 | 6852 | 7055 | 6939 | 6883 | 7136 | 6805 | 6969 |
| K03703 excinuclease ABC subunit C | 6235 | 6121 | 6226 | 6094 | 6834 | 7122 | 7032 | 6976 | 6308 | 6712 | 6659 | 6527 |
| K03723 transcription-repair coupling factor (superfamily II helicase) [EC:3.6.4.-] | 11097 | 10946 | 11065 | 10996 | 10106 | 10088 | 9994 | 9974 | 9925 | 10260 | 10148 | 10209 |
| K10841 DNA excision repair protein ERCC-6 | 1 | 2 | 0 | 4 | 3 | 6 | 6 | 10 | 1 | 4 | 7 | 5 |
| ko03430 Mismatch repair | 96434 | 95961 | 97299 | 96882 | 100847 | 101951 | 102336 | 100654 | 105670 | 106202 | 106571 | 106294 |
| K01141 exodeoxyribonuclease I [EC:3.1.11.1] | 3094 | 3153 | 3079 | 3148 | 1940 | 1876 | 1907 | 1952 | 2024 | 1959 | 1898 | 1999 |
| K03555 DNA mismatch repair protein MutS | 8181 | 8083 | 7996 | 8009 | 10025 | 10112 | 10301 | 10184 | 8969 | 8929 | 9114 | 9086 |
| K03572 DNA mismatch repair protein MutL | 5814 | 5784 | 5754 | 5890 | 6807 | 6647 | 6918 | 6824 | 6349 | 6272 | 6196 | 6187 |
| K03573 DNA mismatch repair protein MutH | 424 | 459 | 420 | 436 | 266 | 341 | 326 | 331 | 587 | 544 | 492 | 618 |
| K03601 exodeoxyribonuclease VII large subunit [EC:3.1.11.6] | 5514 | 5199 | 5382 | 5300 | 4797 | 4911 | 4421 | 4515 | 4545 | 4356 | 4419 | 4458 |
| K03602 exodeoxyribonuclease VII small subunit [EC:3.1.11.6] | 975 | 963 | 972 | 928 | 856 | 892 | 852 | 887 | 761 | 789 | 770 | 826 |
| K06223 DNA adenine methylase [EC:2.1.1.72] | 3937 | 3953 | 4012 | 4001 | 3805 | 3804 | 3802 | 3866 | 3212 | 3265 | 3275 | 3256 |
| K07456 DNA mismatch repair protein MutS2 | 3276 | 3307 | 3504 | 3443 | 4979 | 4958 | 5551 | 5323 | 4987 | 5104 | 5149 | 4925 |
| K08737 DNA mismatch repair protein MSH6 | 0 | 0 | 0 | 0 | 0 | 0 | 0 | 0 | 0 | 1 | 0 | 0 |
| K10857 exodeoxyribonuclease X [EC:3.1.11.-] | 321 | 375 | 329 | 390 | 265 | 263 | 252 | 245 | 523 | 468 | 545 | 521 |
| ko03440 Homologous recombination | 115605 | 114607 | 114030 | 114803 | 111260 | 113115 | 111375 | 112172 | 111679 | 110741 | 112283 | 111588 |
| K01159 crossover junction endodeoxyribonuclease RuvC [EC:3.1.22.4] | 1084 | 1191 | 1093 | 1122 | 1634 | 1607 | 1576 | 1556 | 1468 | 1486 | 1439 | 1367 |
| K02317 DNA replication protein DnaT | 350 | 344 | 336 | 358 | 201 | 216 | 252 | 217 | 443 | 460 | 455 | 467 |
| K02686 primosomal replication protein N | 216 | 241 | 200 | 216 | 173 | 162 | 166 | 181 | 347 | 354 | 292 | 341 |
| K03550 holliday junction DNA helicase RuvA [EC:3.6.4.12] | 1507 | 1510 | 1466 | 1461 | 1832 | 1809 | 1759 | 1854 | 1762 | 1757 | 1758 | 1745 |
| K03551 holliday junction DNA helicase RuvB [EC:3.6.4.12] | 2946 | 3039 | 3132 | 3059 | 3829 | 3977 | 3902 | 4018 | 3727 | 3691 | 3896 | 3601 |
| K03553 recombination protein RecA | 3020 | 2886 | 3154 | 3134 | 3405 | 3368 | 3476 | 3560 | 3545 | 3530 | 3761 | 3492 |
| K03581 exodeoxyribonuclease V alpha subunit [EC:3.1.11.5] | 6270 | 6197 | 5845 | 6211 | 3229 | 3318 | 3096 | 3235 | 4359 | 4356 | 4215 | 4450 |
| K03582 exodeoxyribonuclease V beta subunit [EC:3.1.11.5] | 7894 | 7640 | 7007 | 7779 | 4271 | 4573 | 4291 | 4800 | 4797 | 4655 | 4728 | 4680 |
| K03583 exodeoxyribonuclease V gamma subunit [EC:3.1.11.5] | 7570 | 7308 | 6888 | 6984 | 4327 | 4274 | 4037 | 4392 | 4547 | 4392 | 4452 | 4560 |
| K03584 DNA repair protein RecO (recombination protein O) | 1658 | 1664 | 1793 | 1750 | 2446 | 2467 | 2494 | 2419 | 2361 | 2387 | 2417 | 2328 |
| K03629 DNA replication and repair protein RecF | 3793 | 3791 | 3811 | 3940 | 3981 | 4027 | 4090 | 3937 | 3821 | 3946 | 4000 | 4084 |
| K03655 ATP-dependent DNA helicase RecG [EC:3.6.4.12] | 8326 | 8125 | 8243 | 8217 | 8738 | 8822 | 8758 | 8759 | 9776 | 9910 | 9952 | 10112 |
| K04066 primosomal protein N' (replication factor Y) (superfamily II helicase) [EC:3.6.4.-] | 8251 | 8202 | 8099 | 8320 | 9268 | 9339 | 9160 | 9485 | 8474 | 8238 | 8457 | 8377 |
| K04067 primosomal replication protein N'' | 6143 | 5830 | 5430 | 5469 | 3088 | 3090 | 2817 | 3095 | 2208 | 2030 | 2066 | 2156 |
| K06187 recombination protein RecR | 2326 | 2250 | 2051 | 2203 | 2072 | 2152 | 2132 | 2276 | 2132 | 2206 | 2161 | 2234 |
| K10866 DNA repair protein RAD50 [EC:3.6.-.-] | 0 | 0 | 0 | 0 | 2 | 0 | 0 | 0 | 0 | 0 | 0 | 0 |
| K10877 DNA repair and recombination protein RAD54B [EC:3.6.4.-] | 0 | 0 | 0 | 0 | 0 | 1 | 0 | 0 | 0 | 0 | 0 | 0 |
| K10901 bloom syndrome protein [EC:3.6.4.12] | 0 | 0 | 0 | 1 | 0 | 0 | 0 | 0 | 0 | 0 | 0 | 0 |
| ko03450 Non-homologous end-joining | 5222 | 5037 | 4877 | 4964 | 3160 | 3082 | 3141 | 3237 | 3143 | 3099 | 3084 | 3201 |
| K01971 bifunctional non-homologous end joining protein LigD [EC:6.5.1.1] | 3714 | 3624 | 3514 | 3500 | 2674 | 2536 | 2649 | 2671 | 2200 | 2139 | 2135 | 2200 |
| K10979 DNA end-binding protein Ku | 1505 | 1408 | 1363 | 1461 | 484 | 546 | 490 | 566 | 934 | 948 | 949 | 990 |
| ko03460 Fanconi anemia pathway | 0 | 4 | 0 | 1 | 5 | 4 | 5 | 0 | 21 | 31 | 18 | 29 |
| K10896 fanconi anemia group M protein | 0 | 2 | 0 | 0 | 3 | 4 | 0 | 0 | 10 | 21 | 10 | 15 |
| Environmental Information Processing | 1423762 | 1411500 | 1378838 | 1410317 | 953242 | 962025 | 932646 | 956282 | 1095507 | 1116892 | 1087436 | 1117623 |
| Membrane transport | 804952 | 799226 | 778794 | 799027 | 505515 | 511270 | 495820 | 508714 | 653310 | 672004 | 649760 | 670760 |
| ko02010 ABC transporters | 547645 | 543469 | 532152 | 545891 | 332376 | 337586 | 327016 | 334562 | 461008 | 476958 | 462089 | 476914 |
| K01995 branched-chain amino acid transport system ATP-binding protein | 4231 | 4260 | 4059 | 4370 | 2411 | 2442 | 2382 | 2302 | 3121 | 3284 | 3050 | 3227 |
| K01996 branched-chain amino acid transport system ATP-binding protein | 3653 | 3632 | 3751 | 3822 | 2071 | 2168 | 2049 | 2130 | 2987 | 3009 | 2961 | 3022 |
| K01997 branched-chain amino acid transport system permease protein | 5127 | 4999 | 5054 | 5121 | 2665 | 2934 | 2760 | 2881 | 3825 | 3988 | 3665 | 3793 |
| K01998 branched-chain amino acid transport system permease protein | 8864 | 8720 | 8806 | 8876 | 4940 | 4946 | 4720 | 4962 | 5061 | 5479 | 5384 | 5335 |
| K01999 branched-chain amino acid transport system substrate-binding protein | 8994 | 9109 | 8708 | 9021 | 5632 | 5660 | 5267 | 5536 | 6171 | 6501 | 6015 | 6441 |
| K02000 glycine betaine/proline transport system ATP-binding protein [EC:7.6.2.9] | 4639 | 4718 | 4750 | 4732 | 4895 | 4809 | 4933 | 4927 | 3930 | 3915 | 3968 | 3858 |
| K02001 glycine betaine/proline transport system permease protein | 4094 | 4130 | 3946 | 4040 | 3765 | 3730 | 3559 | 3709 | 3061 | 3121 | 3061 | 2990 |
| K02002 glycine betaine/proline transport system substrate-binding protein | 7561 | 7843 | 7293 | 7444 | 5841 | 5722 | 5405 | 5987 | 4243 | 4281 | 4178 | 4184 |
| K02006 cobalt/nickel transport system ATP-binding protein | 802 | 826 | 815 | 843 | 468 | 444 | 414 | 435 | 1051 | 1137 | 1104 | 1130 |
| K02007 cobalt/nickel transport system permease protein | 882 | 824 | 787 | 911 | 390 | 462 | 401 | 389 | 976 | 1053 | 917 | 1070 |
| K02008 cobalt/nickel transport system permease protein | 701 | 636 | 593 | 645 | 316 | 359 | 304 | 304 | 811 | 840 | 706 | 787 |
| K02009 cobalt/nickel transport protein | 360 | 380 | 370 | 375 | 251 | 254 | 229 | 209 | 469 | 467 | 417 | 508 |
| K02010 iron(III) transport system ATP-binding protein [EC:7.2.2.7] | 3959 | 3796 | 3797 | 3971 | 2415 | 2529 | 2219 | 2396 | 3027 | 3084 | 2875 | 3146 |
| K02011 iron(III) transport system permease protein | 7035 | 6903 | 6708 | 7117 | 4153 | 4286 | 4217 | 4327 | 4604 | 4858 | 4768 | 4831 |
| K02012 iron(III) transport system substrate-binding protein | 6176 | 6052 | 5748 | 6226 | 3509 | 3690 | 3512 | 3693 | 3641 | 3779 | 3558 | 3864 |
| K02017 molybdate transport system ATP-binding protein [EC:7.3.2.5] | 2407 | 2456 | 2176 | 2432 | 1414 | 1436 | 1433 | 1414 | 1683 | 1786 | 1695 | 1869 |
| K02018 molybdate transport system permease protein | 1537 | 1587 | 1431 | 1439 | 975 | 886 | 903 | 894 | 1085 | 1080 | 1071 | 1134 |
| K02020 molybdate transport system substrate-binding protein | 2644 | 2442 | 2401 | 2486 | 1364 | 1500 | 1355 | 1530 | 2351 | 2549 | 2406 | 2505 |
| K02036 phosphate transport system ATP-binding protein [EC:7.3.2.1] | 2994 | 3062 | 3102 | 3145 | 2911 | 2966 | 3018 | 2925 | 2859 | 2895 | 2973 | 3007 |
| K02037 phosphate transport system permease protein | 5890 | 5888 | 5899 | 6043 | 5253 | 5409 | 5207 | 5181 | 4692 | 4428 | 4520 | 4454 |
| K02038 phosphate transport system permease protein | 3703 | 3676 | 3791 | 3728 | 3561 | 3609 | 3507 | 3366 | 3364 | 3228 | 3282 | 3321 |
| K02040 phosphate transport system substrate-binding protein | 4868 | 4915 | 4815 | 4837 | 5744 | 5738 | 5693 | 5738 | 4821 | 4756 | 4964 | 4935 |
| K02041 phosphonate transport system ATP-binding protein [EC:7.3.2.2] | 3010 | 2934 | 2815 | 2947 | 1936 | 1922 | 1922 | 2005 | 1683 | 1622 | 1644 | 1670 |
| K02042 phosphonate transport system permease protein | 4162 | 4339 | 3910 | 4053 | 2562 | 2582 | 2608 | 2669 | 2881 | 2864 | 2547 | 2876 |
| K02044 phosphonate transport system substrate-binding protein | 3908 | 3879 | 3761 | 3746 | 2563 | 2546 | 2465 | 2502 | 2132 | 2176 | 1973 | 2202 |
| K02062 thiamine transport system ATP-binding protein | 417 | 456 | 417 | 425 | 311 | 331 | 338 | 273 | 640 | 591 | 627 | 638 |
| K02063 thiamine transport system permease protein | 1025 | 1031 | 1061 | 1113 | 691 | 805 | 694 | 717 | 1416 | 1503 | 1389 | 1544 |
| K02064 thiamine transport system substrate-binding protein | 594 | 607 | 530 | 599 | 368 | 381 | 379 | 371 | 798 | 887 | 786 | 880 |
| K02065 phospholipid/cholesterol/gamma-HCH transport system ATP-binding protein | 3638 | 3659 | 3835 | 3683 | 3431 | 3671 | 3358 | 3531 | 2807 | 2832 | 2817 | 2776 |
| K02066 phospholipid/cholesterol/gamma-HCH transport system permease protein | 4989 | 4914 | 5061 | 5085 | 4252 | 4343 | 4129 | 4256 | 3101 | 3046 | 3050 | 2978 |
| K02067 phospholipid/cholesterol/gamma-HCH transport system substrate-binding protein | 3831 | 3838 | 3776 | 3784 | 3765 | 3884 | 3626 | 3634 | 2808 | 2777 | 2824 | 2757 |
| K02071 D-methionine transport system ATP-binding protein | 6454 | 6312 | 6158 | 6340 | 3960 | 4117 | 3921 | 3972 | 4593 | 4751 | 4576 | 4898 |
| K02072 D-methionine transport system permease protein | 3618 | 3502 | 3307 | 3548 | 2286 | 2320 | 2157 | 2279 | 2441 | 2613 | 2656 | 2777 |
| K02073 D-methionine transport system substrate-binding protein | 6114 | 6299 | 5860 | 6090 | 3799 | 3763 | 3688 | 3716 | 5251 | 5218 | 5050 | 5145 |
| K02193 heme exporter protein A [EC:7.6.2.5] | 293 | 287 | 253 | 274 | 140 | 137 | 130 | 147 | 536 | 565 | 524 | 614 |

| | | | | | | | | | | | | |
|---|---|---|---|---|---|---|---|---|---|---|---|---|
| K02194 heme exporter protein B | 1200 | 1198 | 1051 | 1109 | 553 | 596 | 612 | 635 | 867 | 826 | 794 | 849 |
| K02195 heme exporter protein C | 1473 | 1485 | 1375 | 1377 | 725 | 707 | 724 | 749 | 1010 | 978 | 971 | 987 |
| K02196 heme exporter protein D | 343 | 346 | 287 | 301 | 193 | 177 | 141 | 166 | 268 | 224 | 204 | 289 |
| K02424 L-cystine transport system substrate-binding protein | 1933 | 2054 | 1989 | 2026 | 1320 | 1408 | 1394 | 1345 | 1119 | 1020 | 1200 | 1224 |
| K02471 vitamin B12/bleomycin/antimicrobial peptide transport system ATP-binding/permease protein | 154 | 157 | 138 | 157 | 190 | 180 | 208 | 189 | 259 | 282 | 278 | 267 |
| K03523 biotin transport system substrate-specific component | 554 | 570 | 487 | 537 | 181 | 213 | 182 | 157 | 412 | 533 | 445 | 498 |
| K05643 ATP-binding cassette, subfamily A (ABC1), member 3 | 0 | 0 | 1 | 0 | 0 | 0 | 0 | 0 | 1 | 0 | 0 | 1 |
| K05655 ATP-binding cassette, subfamily B (MDR/TAP), member 8 | 1 | 0 | 0 | 1 | 0 | 0 | 0 | 0 | 0 | 3 | 1 | 0 |
| K05656 ATP-binding cassette, subfamily B (MDR/TAP), member 9 | 1 | 0 | 0 | 1 | 0 | 1 | 0 | 0 | 0 | 0 | 0 | 1 |
| K05657 ATP-binding cassette, subfamily B (MDR/TAP), member 10 | 4 | 2 | 3 | 3 | 0 | 1 | 0 | 0 | 2 | 1 | 6 | 5 |
| K05658 ATP-binding cassette, subfamily B (MDR/TAP), member 1 [EC:7.6.2.2] | 11 | 6 | 5 | 8 | 2 | 3 | 6 | 3 | 16 | 11 | 12 | 12 |
| K05659 ATP-binding cassette, subfamily B (MDR/TAP), member 4 [EC:7.6.2.2] | 1 | 0 | 1 | 0 | 0 | 0 | 0 | 0 | 2 | 2 | 3 | 3 |
| K05661 ATP-binding cassette, subfamily B (MDR/TAP), member 6 | 0 | 0 | 0 | 0 | 1 | 0 | 0 | 0 | 0 | 0 | 1 | 0 |
| K05662 ATP-binding cassette, subfamily B (MDR/TAP), member 7 | 1 | 0 | 0 | 0 | 0 | 0 | 0 | 0 | 0 | 0 | 0 | 0 |
| K05663 ATP-binding cassette, subfamily B, mitochondrial transporter ATM | 0 | 0 | 0 | 0 | 0 | 1 | 0 | 0 | 0 | 0 | 0 | 0 |
| K05666 ATP-binding cassette, subfamily C (CFTR/MRP), member 2 | 0 | 0 | 0 | 0 | 0 | 1 | 0 | 0 | 1 | 1 | 0 | 0 |
| K05673 ATP-binding cassette, subfamily C (CFTR/MRP), member 4 | 0 | 1 | 1 | 1 | 0 | 0 | 0 | 0 | 0 | 1 | 0 | 0 |
| K05685 macrolide transport system ATP-binding/permease protein [EC:3.6.3.-] | 3942 | 3828 | 3702 | 3735 | 2174 | 2211 | 2058 | 2263 | 2446 | 2393 | 2267 | 2274 |
| K05772 tungstate transport system substrate-binding protein | 20 | 19 | 12 | 19 | 17 | 15 | 19 | 16 | 122 | 113 | 93 | 113 |
| K05773 tungstate transport system permease protein | 9 | 12 | 13 | 15 | 8 | 7 | 18 | 19 | 56 | 85 | 78 | 108 |
| K05776 molybdate transport system ATP-binding protein | 1641 | 1674 | 1684 | 1589 | 3015 | 2877 | 3120 | 2958 | 2921 | 2963 | 3065 | 2958 |
| K05813 sn-glycerol 3-phosphate transport system substrate-binding protein | 1650 | 1637 | 1597 | 1651 | 759 | 856 | 794 | 872 | 2238 | 2249 | 2177 | 2258 |
| K05814 sn-glycerol 3-phosphate transport system permease protein | 1486 | 1525 | 1472 | 1455 | 679 | 687 | 686 | 661 | 1939 | 2126 | 1960 | 2094 |
| K05815 sn-glycerol 3-phosphate transport system permease protein | 1430 | 1473 | 1454 | 1432 | 564 | 568 | 631 | 616 | 1862 | 2080 | 1934 | 2062 |
| K05816 sn-glycerol 3-phosphate transport system ATP-binding protein [EC:7.6.2.10] | 1417 | 1455 | 1379 | 1482 | 751 | 813 | 831 | 763 | 1953 | 2111 | 1966 | 2191 |
| K05845 osmoprotectant transport system substrate-binding protein | 1816 | 1829 | 1774 | 1837 | 1056 | 1183 | 1054 | 1004 | 1469 | 1618 | 1480 | 1594 |
| K05846 osmoprotectant transport system permease protein | 2582 | 2539 | 2373 | 2460 | 1624 | 1615 | 1612 | 1734 | 2051 | 2084 | 1966 | 2113 |
| K05847 osmoprotectant transport system ATP-binding protein [EC:7.6.2.9] | 3223 | 3345 | 3177 | 3330 | 1840 | 1857 | 1767 | 1827 | 1939 | 2129 | 1941 | 1995 |
| K06073 vitamin B12 transport system permease protein | 516 | 507 | 527 | 498 | 385 | 371 | 340 | 310 | 728 | 759 | 711 | 728 |
| K06074 vitamin B12 transport system ATP-binding protein [EC:7.6.2.8] | 420 | 399 | 409 | 464 | 312 | 293 | 296 | 260 | 625 | 573 | 558 | 587 |
| K06159 multidrug/microcin transport system ATP-binding/permease protein | 940 | 895 | 894 | 944 | 601 | 727 | 627 | 639 | 1228 | 1354 | 1355 | 1229 |
| K06160 putative pyoverdin transport system ATP-binding/permease protein | 2 | 2 | 2 | 3 | 1 | 1 | 0 | 1 | 0 | 1 | 1 | 1 |
| K06726 D-ribose pyranase [EC:5.4.99.62] | 343 | 288 | 336 | 358 | 208 | 202 | 204 | 183 | 448 | 434 | 452 | 407 |
| K06857 tungstate transport system ATP-binding protein [EC:7.3.2.6] | 8 | 22 | 19 | 25 | 7 | 19 | 23 | 12 | 80 | 63 | 88 | 77 |
| K06858 vitamin B12 transport system substrate-binding protein | 1776 | 1675 | 1598 | 1714 | 942 | 1006 | 967 | 1012 | 983 | 1113 | 981 | 1106 |
| K06861 lipopolysaccharide export system ATP-binding protein [EC:3.6.3.-] | 1919 | 2093 | 1960 | 1967 | 2553 | 2694 | 2615 | 2526 | 2186 | 2174 | 2200 | 2212 |
| K07091 lipopolysaccharide export system permease protein | 3525 | 3823 | 3800 | 3887 | 5367 | 5518 | 5515 | 5296 | 4388 | 4170 | 4460 | 4250 |
| K07122 phospholipid transport system transporter-binding protein | 651 | 716 | 699 | 742 | 448 | 478 | 415 | 376 | 432 | 400 | 449 | 453 |
| K07323 phospholipid transport system substrate-binding protein | 1092 | 1033 | 1004 | 1109 | 739 | 737 | 660 | 691 | 616 | 777 | 672 | 726 |
| K08712 ATP-binding cassette, subfamily G (WHITE), member 2, SNQ2 | 0 | 0 | 0 | 0 | 0 | 2 | 0 | 0 | 0 | 0 | 0 | 0 |
| K09688 capsular polysaccharide transport system permease protein | 162 | 171 | 164 | 187 | 267 | 282 | 247 | 251 | 45 | 50 | 44 | 34 |
| K09689 capsular polysaccharide transport system ATP-binding protein [EC:7.6.2.12] | 163 | 148 | 132 | 112 | 199 | 233 | 203 | 174 | 14 | 9 | 7 | 20 |
| K09690 lipopolysaccharide transport system permease protein | 3240 | 3339 | 3241 | 3092 | 1923 | 1867 | 1836 | 2024 | 1569 | 1523 | 1497 | 1495 |
| K09691 lipopolysaccharide transport system ATP-binding protein | 3037 | 2905 | 2996 | 3065 | 2109 | 2092 | 1949 | 2071 | 1287 | 1291 | 1352 | 1422 |
| K09692 teichoic acid transport system permease protein | 442 | 426 | 451 | 451 | 105 | 125 | 113 | 101 | 313 | 385 | 324 | 385 |
| K09693 teichoic acid transport system ATP-binding protein [EC:7.5.2.4] | 706 | 689 | 668 | 748 | 127 | 150 | 143 | 103 | 546 | 635 | 681 | 740 |
| K09694 lipooligosaccharide transport system permease protein | 0 | 2 | 2 | 0 | 0 | 0 | 0 | 0 | 0 | 0 | 3 | 0 |
| K09695 lipooligosaccharide transport system ATP-binding protein | 3 | 2 | 6 | 3 | 2 | 0 | 0 | 1 | 4 | 7 | 4 | 4 |
| K09696 sodium transport system permease protein | 3 | 12 | 10 | 4 | 7 | 7 | 3 | 4 | 34 | 24 | 21 | 21 |
| K09697 sodium transport system ATP-binding protein [EC:7.2.2.4] | 4 | 10 | 6 | 7 | 2 | 2 | 0 | 4 | 9 | 3 | 24 | 19 |
| K09808 lipoprotein-releasing system permease protein | 4499 | 4629 | 4754 | 4653 | 7028 | 6978 | 7290 | 7387 | 6099 | 6030 | 6108 | 6050 |
| K09810 lipoprotein-releasing system ATP-binding protein [EC:3.6.3.-] | 788 | 877 | 887 | 875 | 1824 | 1829 | 1885 | 1898 | 1662 | 1537 | 1590 | 1589 |
| K09811 cell division transport system permease protein | 3783 | 3733 | 3688 | 3814 | 3684 | 3912 | 3776 | 3795 | 3557 | 3810 | 3908 | 3809 |
| K09812 cell division transport system ATP-binding protein | 2764 | 2846 | 2992 | 2918 | 2654 | 2587 | 2731 | 2653 | 2805 | 2827 | 2885 | 2874 |
| K09815 zinc transport system substrate-binding protein | 4900 | 4844 | 4722 | 4900 | 4342 | 4503 | 4293 | 4560 | 4145 | 3968 | 3963 | 4032 |
| K09816 zinc transport system permease protein | 3819 | 3711 | 3730 | 3775 | 3561 | 3486 | 3358 | 3363 | 3193 | 3144 | 3284 | 3131 |
| K09817 zinc transport system ATP-binding protein [EC:7.2.2.-] | 3825 | 3835 | 3776 | 3825 | 3777 | 3667 | 3650 | 3735 | 3444 | 3195 | 3346 | 3425 |
| K09969 general L-amino acid transport system substrate-binding protein | 1983 | 2075 | 1854 | 1891 | 1450 | 1430 | 1335 | 1372 | 717 | 745 | 696 | 627 |
| K09970 general L-amino acid transport system permease protein | 317 | 288 | 301 | 342 | 445 | 468 | 436 | 439 | 157 | 155 | 170 | 160 |
| K09971 general L-amino acid transport system permease protein | 267 | 273 | 319 | 296 | 456 | 500 | 501 | 477 | 142 | 156 | 132 | 138 |
| K09972 general L-amino acid transport system ATP-binding protein [EC:7.4.2.1] | 173 | 136 | 131 | 150 | 234 | 279 | 202 | 261 | 74 | 52 | 43 | 65 |
| K09996 arginine transport system substrate-binding protein | 702 | 688 | 645 | 707 | 385 | 375 | 413 | 348 | 1173 | 1238 | 1108 | 1197 |
| K09997 arginine transport system substrate-binding protein | 269 | 250 | 275 | 262 | 215 | 276 | 264 | 237 | 268 | 287 | 277 | 235 |
| K09998 arginine transport system permease protein | 393 | 357 | 376 | 334 | 238 | 238 | 213 | 233 | 483 | 569 | 452 | 463 |
| K09999 arginine transport system permease protein | 399 | 389 | 375 | 416 | 281 | 332 | 272 | 294 | 572 | 633 | 586 | 652 |
| K10000 arginine transport system ATP-binding protein [EC:7.4.2.1] | 476 | 418 | 422 | 525 | 276 | 340 | 330 | 341 | 655 | 674 | 638 | 648 |
| K10001 glutamate/aspartate transport system substrate-binding protein | 2101 | 2176 | 2111 | 2150 | 1252 | 1249 | 1245 | 1282 | 1412 | 1395 | 1385 | 1390 |
| K10002 glutamate/aspartate transport system permease protein | 1423 | 1356 | 1303 | 1195 | 818 | 910 | 810 | 814 | 855 | 809 | 813 | 904 |
| K10003 glutamate/aspartate transport system permease protein | 1278 | 1269 | 1297 | 1284 | 811 | 909 | 806 | 821 | 927 | 915 | 888 | 950 |
| K10004 glutamate/aspartate transport system ATP-binding protein [EC:7.4.2.1] | 1324 | 1218 | 1269 | 1350 | 833 | 853 | 809 | 832 | 889 | 945 | 854 | 901 |
| K10005 glutamate transport system substrate-binding protein | 2 | 2 | 5 | 2 | 0 | 2 | 0 | 3 | 4 | 5 | 2 | 11 |
| K10006 glutamate transport system permease protein | 0 | 0 | 2 | 5 | 2 | 3 | 1 | 2 | 4 | 1 | 0 | 4 |
| K10007 glutamate transport system permease protein | 2 | 5 | 9 | 1 | 4 | 6 | 3 | 3 | 7 | 13 | 4 | 20 |
| K10008 glutamate transport system ATP-binding protein [EC:7.4.2.1] | 19 | 20 | 21 | 27 | 5 | 5 | 3 | 7 | 21 | 17 | 13 | 15 |
| K10009 L-cystine transport system permease protein | 1688 | 1671 | 1593 | 1615 | 1186 | 1186 | 1045 | 1154 | 860 | 824 | 796 | 907 |
| K10010 L-cystine transport system ATP-binding protein [EC:7.4.2.1] | 532 | 544 | 523 | 584 | 573 | 605 | 518 | 555 | 570 | 549 | 510 | 533 |
| K10013 lysine/arginine/ornithine transport system substrate-binding protein | 805 | 722 | 734 | 769 | 528 | 644 | 566 | 569 | 978 | 968 | 1002 | 983 |
| K10014 histidine transport system substrate-binding protein | 1528 | 1520 | 1557 | 1549 | 846 | 978 | 899 | 1025 | 1002 | 995 | 941 | 1021 |
| K10015 histidine transport system permease protein | 1239 | 1275 | 1132 | 1183 | 727 | 679 | 665 | 661 | 692 | 822 | 763 | 767 |
| K10016 histidine transport system permease protein | 982 | 1051 | 961 | 1068 | 597 | 616 | 573 | 607 | 767 | 731 | 728 | 843 |
| K10017 histidine transport system ATP-binding protein [EC:7.4.2.1] | 1450 | 1443 | 1320 | 1375 | 900 | 794 | 761 | 842 | 856 | 878 | 856 | 864 |
| K10018 octopine/nopaline transport system substrate-binding protein | 5 | 1 | 4 | 0 | 0 | 0 | 5 | 2 | 6 | 3 | 3 | 9 |
| K10019 octopine/nopaline transport system permease protein | 1 | 0 | 0 | 0 | 0 | 1 | 1 | 0 | 4 | 0 | 5 | 1 |
| K10020 octopine/nopaline transport system permease protein | 2 | 1 | 0 | 0 | 1 | 5 | 2 | 0 | 3 | 1 | 3 | 3 |
| K10021 octopine/nopaline transport system ATP-binding protein [EC:7.4.2.1] | 4 | 4 | 4 | 1 | 0 | 5 | 0 | 1 | 3 | 3 | 1 | 5 |
| K10022 arginine/ornithine transport system substrate-binding protein | 1984 | 1818 | 1846 | 1841 | 994 | 985 | 1047 | 990 | 646 | 586 | 574 | 575 |
| K10023 arginine/ornithine transport system permease protein | 1067 | 1011 | 974 | 1049 | 581 | 598 | 526 | 543 | 322 | 300 | 349 | 280 |
| K10024 arginine/ornithine transport system permease protein | 1109 | 1133 | 1058 | 1018 | 531 | 622 | 577 | 613 | 416 | 393 | 345 | 297 |
| K10025 arginine/ornithine transport system ATP-binding protein [EC:7.4.2.1] | 1030 | 967 | 1065 | 965 | 601 | 562 | 522 | 586 | 330 | 288 | 319 | 320 |
| K10036 glutamine transport system substrate-binding protein | 662 | 663 | 592 | 619 | 548 | 550 | 497 | 488 | 848 | 918 | 855 | 920 |
| K10037 glutamine transport system permease protein | 610 | 637 | 606 | 581 | 440 | 474 | 426 | 407 | 795 | 893 | 811 | 862 |
| K10038 glutamine transport system ATP-binding protein [EC:7.4.2.1] | 527 | 470 | 495 | 473 | 343 | 362 | 293 | 339 | 762 | 687 | 750 | 714 |
| K10039 aspartate/glutamate/glutamine transport system substrate-binding protein | 56 | 65 | 75 | 68 | 19 | 23 | 22 | 39 | 54 | 79 | 60 | 68 |
| K10040 aspartate/glutamate/glutamine transport system permease protein | 109 | 116 | 102 | 108 | 40 | 42 | 40 | 33 | 97 | 116 | 95 | 106 |
| K10041 aspartate/glutamate/glutamine transport system ATP-binding protein [EC:7.4.2.1] | 59 | 73 | 94 | 91 | 14 | 42 | 26 | 33 | 73 | 78 | 66 | 93 |
| K10107 capsular polysaccharide transport system permease protein | 245 | 262 | 263 | 253 | 429 | 421 | 419 | 365 | 93 | 98 | 70 | 77 |
| K10108 maltose/maltodextrin transport system substrate-binding protein | 740 | 770 | 737 | 767 | 558 | 590 | 553 | 576 | 1047 | 1143 | 1009 | 1084 |
| K10109 maltose/maltodextrin transport system permease protein | 942 | 1024 | 994 | 967 | 708 | 750 | 691 | 694 | 1383 | 1417 | 1357 | 1361 |
| K10110 maltose/maltodextrin transport system permease protein | 575 | 591 | 583 | 579 | 388 | 422 | 401 | 420 | 717 | 861 | 848 | 846 |
| K10111 multiple sugar transport system ATP-binding protein [EC:3.6.3.-] | 2239 | 2152 | 2095 | 2158 | 1339 | 1412 | 1414 | 1350 | 2162 | 2200 | 2009 | 2254 |
| K10112 multiple sugar transport system ATP-binding protein | 3863 | 3896 | 3839 | 3936 | 2048 | 2019 | 2003 | 2103 | 2015 | 2306 | 2122 | 2109 |
| K10117 raffinose/stachyose/melibiose transport system substrate-binding protein | 8532 | 8247 | 8648 | 8405 | 1482 | 1482 | 1500 | 1408 | 8794 | 9580 | 9305 | 9575 |
| K10118 raffinose/stachyose/melibiose transport system permease protein | 7026 | 6708 | 6919 | 6980 | 1310 | 1429 | 1341 | 1283 | 7569 | 7993 | 7766 | 7928 |
| K10119 raffinose/stachyose/melibiose transport system permease protein | 6078 | 5831 | 6355 | 6047 | 1162 | 1222 | 1215 | 1083 | 6315 | 7366 | 6612 | 7179 |
| K10188 lactose/L-arabinose transport system substrate-binding protein | 402 | 388 | 429 | 419 | 111 | 105 | 125 | 93 | 756 | 773 | 784 | 813 |
| K10189 lactose/L-arabinose transport system permease protein | 294 | 280 | 308 | 276 | 73 | 74 | 59 | 57 | 636 | 673 | 653 | 653 |
| K10190 lactose/L-arabinose transport system permease protein | 317 | 273 | 280 | 302 | 68 | 83 | 72 | 71 | 597 | 567 | 581 | 643 |
| K10191 lactose/L-arabinose transport system ATP-binding protein | 1 | 2 | 0 | 2 | 0 | 0 | 0 | 0 | 0 | 1 | 1 | 1 |
| K10192 oligogalacturonide transport system substrate-binding protein | 1682 | 1680 | 1706 | 1773 | 366 | 419 | 355 | 334 | 1526 | 1763 | 1694 | 1741 |
| K10193 oligogalacturonide transport system permease protein | 2280 | 2198 | 2197 | 2347 | 385 | 408 | 383 | 384 | 1745 | 1866 | 1826 | 1953 |
| K10194 oligogalacturonide transport system permease protein | 2571 | 2351 | 2307 | 2451 | 374 | 420 | 426 | 415 | 1678 | 2057 | 2008 | 2035 |
| K10195 oligogalacturonide transport system ATP-binding protein | 459 | 477 | 423 | 476 | 108 | 134 | 130 | 123 | 878 | 809 | 823 | 847 |
| K10200 N-acetylglucosamine transport system substrate-binding protein | 137 | 125 | 150 | 136 | 21 | 15 | 16 | 19 | 83 | 91 | 85 | 107 |
| K10201 N-acetylglucosamine transport system permease protein | 204 | 210 | 252 | 226 | 13 | 11 | 22 | 8 | 135 | 152 | 156 | 160 |
| K10202 N-acetylglucosamine transport system permease protein | 454 | 470 | 446 | 439 | 103 | 116 | 129 | 99 | 329 | 361 | 316 | 410 |
| K10227 sorbitol/mannitol transport system substrate-binding protein | 1676 | 1661 | 1470 | 1489 | 772 | 792 | 734 | 878 | 543 | 475 | 420 | 464 |
| K10228 sorbitol/mannitol transport system permease protein | 1324 | 1211 | 1061 | 1132 | 556 | 612 | 569 | 620 | 396 | 383 | 302 | 322 |
| K10229 sorbitol/mannitol transport system permease protein | 1038 | 953 | 930 | 907 | 550 | 525 | 508 | 573 | 328 | 281 | 283 | 317 |
| K10232 alpha-glucoside transport system substrate-binding protein | 0 | 0 | 0 | 1 | 0 | 2 | 0 | 0 | 0 | 0 | 0 | 0 |
| K10233 alpha-glucoside transport system permease protein | 1 | 2 | 0 | 2 | 0 | 1 | 1 | 0 | 3 | 1 | 2 | 2 |
| K10234 alpha-glucoside transport system permease protein | 1 | 0 | 0 | 0 | 4 | 3 | 2 | 1 | 2 | 1 | 4 | 3 |

| | | | | | | | | | | | | |
|---|---|---|---|---|---|---|---|---|---|---|---|---|
| K10235 alpha-glucoside transport system ATP-binding protein | 0 | 1 | 0 | 2 | 1 | 1 | 0 | 0 | 7 | 5 | 3 | 7 |
| K10236 trehalose/maltose transport system substrate-binding protein | 4 | 4 | 2 | 1 | 2 | 0 | 0 | 0 | 3 | 4 | 6 | 4 |
| K10237 trehalose/maltose transport system permease protein | 59 | 61 | 70 | 55 | 11 | 18 | 17 | 7 | 59 | 56 | 53 | 55 |
| K10238 trehalose/maltose transport system permease protein | 7 | 7 | 10 | 9 | 0 | 0 | 0 | 0 | 11 | 2 | 8 | 10 |
| K10240 cellobiose transport system substrate-binding protein | 3 | 5 | 2 | 1 | 0 | 1 | 1 | 0 | 4 | 3 | 8 | 3 |
| K10241 cellobiose transport system permease protein | 464 | 424 | 432 | 450 | 160 | 211 | 191 | 178 | 537 | 592 | 470 | 567 |
| K10242 cellobiose transport system permease protein | 182 | 174 | 199 | 208 | 69 | 66 | 69 | 61 | 239 | 259 | 257 | 283 |
| K10439 ribose transport system substrate-binding protein | 4577 | 4555 | 4434 | 4547 | 2595 | 2701 | 2612 | 2625 | 4447 | 4753 | 4538 | 4753 |
| K10440 ribose transport system permease protein | 4661 | 4416 | 4450 | 4525 | 2291 | 2528 | 2277 | 2299 | 4390 | 4478 | 4297 | 4530 |
| K10441 ribose transport system ATP-binding protein [EC:7.5.2.7] | 6976 | 6946 | 6751 | 6969 | 3754 | 3967 | 3804 | 3849 | 5483 | 5969 | 5577 | 5740 |
| K10537 L-arabinose transport system substrate-binding protein | 600 | 558 | 572 | 596 | 396 | 413 | 403 | 449 | 830 | 898 | 775 | 866 |
| K10538 L-arabinose transport system permease protein | 447 | 460 | 471 | 488 | 299 | 363 | 359 | 372 | 689 | 748 | 727 | 759 |
| K10539 L-arabinose transport system ATP-binding protein [EC:7.5.2.12] | 854 | 843 | 867 | 891 | 551 | 589 | 602 | 622 | 1226 | 1171 | 1274 | 1245 |
| K10540 methyl-galactoside transport system substrate-binding protein | 1930 | 1915 | 1939 | 1943 | 735 | 831 | 683 | 737 | 2077 | 2340 | 2113 | 2231 |
| K10541 methyl-galactoside transport system permease protein | 1704 | 1646 | 1822 | 1711 | 656 | 746 | 727 | 673 | 1809 | 1999 | 1943 | 1987 |
| K10542 methyl-galactoside transport system ATP-binding protein [EC:7.5.2.11] | 2100 | 2132 | 2028 | 2140 | 929 | 1011 | 931 | 938 | 2422 | 2479 | 2372 | 2455 |
| K10543 D-xylose transport system substrate-binding protein | 884 | 844 | 806 | 845 | 467 | 511 | 520 | 482 | 1041 | 1102 | 1021 | 1007 |
| K10544 D-xylose transport system permease protein | 673 | 719 | 699 | 637 | 493 | 464 | 524 | 560 | 643 | 703 | 640 | 672 |
| K10545 D-xylose transport system ATP-binding protein [EC:7.5.2.10] | 1106 | 1188 | 1121 | 1163 | 689 | 759 | 751 | 800 | 1365 | 1528 | 1350 | 1372 |
| K10546 putative multiple sugar transport system substrate-binding protein | 58 | 52 | 62 | 68 | 107 | 77 | 73 | 96 | 81 | 90 | 70 | 80 |
| K10547 putative multiple sugar transport system permease protein | 62 | 50 | 52 | 57 | 65 | 63 | 65 | 87 | 61 | 100 | 104 | 87 |
| K10548 putative multiple sugar transport system ATP-binding protein [EC:7.5.2.-] | 149 | 120 | 113 | 144 | 90 | 110 | 81 | 95 | 117 | 149 | 146 | 148 |
| K10549 D-allose transport system substrate-binding protein | 5 | 7 | 13 | 6 | 3 | 5 | 8 | 6 | 61 | 72 | 83 | 66 |
| K10550 D-allose transport system permease protein | 11 | 7 | 11 | 10 | 6 | 3 | 2 | 7 | 85 | 91 | 79 | 60 |
| K10551 D-allose transport system ATP-binding protein [EC:7.5.2.8] | 29 | 26 | 34 | 23 | 4 | 6 | 16 | 10 | 109 | 99 | 116 | 122 |
| K10552 fructose transport system substrate-binding protein | 0 | 2 | 0 | 0 | 0 | 4 | 0 | 0 | 0 | 6 | 2 | 1 |
| K10553 fructose transport system permease protein | 0 | 0 | 0 | 0 | 3 | 0 | 2 | 0 | 1 | 1 | 0 | 4 |
| K10554 fructose transport system ATP-binding protein | 0 | 0 | 2 | 0 | 0 | 0 | 0 | 0 | 0 | 0 | 2 | 3 |
| K10555 AI-2 transport system substrate-binding protein | 651 | 663 | 638 | 637 | 457 | 511 | 448 | 469 | 823 | 868 | 765 | 857 |
| K10556 AI-2 transport system permease protein | 666 | 612 | 597 | 652 | 418 | 465 | 456 | 459 | 887 | 818 | 809 | 863 |
| K10557 AI-2 transport system permease protein | 663 | 577 | 538 | 637 | 500 | 453 | 429 | 498 | 742 | 702 | 772 | 786 |
| K10558 AI-2 transport system ATP-binding protein | 1018 | 928 | 850 | 970 | 659 | 726 | 725 | 687 | 1258 | 1263 | 1259 | 1267 |
| K10559 rhamnose transport system substrate-binding protein | 732 | 690 | 663 | 764 | 436 | 544 | 492 | 458 | 820 | 902 | 823 | 893 |
| K10560 rhamnose transport system permease protein | 634 | 638 | 548 | 651 | 464 | 529 | 430 | 452 | 826 | 893 | 820 | 849 |
| K10561 rhamnose transport system permease protein | 626 | 607 | 658 | 640 | 402 | 416 | 444 | 435 | 830 | 815 | 749 | 780 |
| K10562 rhamnose transport system ATP-binding protein [EC:7.5.2.-] | 971 | 1032 | 957 | 1092 | 705 | 736 | 714 | 659 | 1216 | 1376 | 1203 | 1293 |
| K10823 oligopeptide transport system ATP-binding protein | 2717 | 2678 | 2653 | 2620 | 878 | 897 | 800 | 793 | 2781 | 3113 | 2881 | 3056 |
| K10824 nickel transport system ATP-binding protein [EC:7.2.2.11] | 536 | 571 | 554 | 629 | 478 | 462 | 463 | 434 | 735 | 760 | 841 | 784 |
| K10829 ferric hydroxamate transport system ATP-binding protein [EC:7.2.2.16] | 654 | 713 | 612 | 690 | 479 | 467 | 477 | 487 | 719 | 724 | 832 | 858 |
| K10830 manganese/zinc transport system ATP-binding protein [EC:7.2.2.5] | 35 | 36 | 33 | 46 | 2 | 4 | 9 | 5 | 43 | 49 | 29 | 44 |
| K11004 ATP-binding cassette, subfamily B, bacterial HlyB/CyaB | 16 | 6 | 9 | 11 | 5 | 5 | 13 | 3 | 126 | 125 | 123 | 129 |
| K11050 multidrug/hemolysin transport system ATP-binding protein | 8 | 6 | 13 | 9 | 0 | 2 | 3 | 3 | 13 | 11 | 8 | 10 |
| K11051 multidrug/hemolysin transport system permease protein | 3 | 3 | 5 | 3 | 2 | 2 | 3 | 0 | 4 | 11 | 12 | 8 |
| K11069 spermidine/putrescine transport system substrate-binding protein | 4047 | 4125 | 4217 | 4157 | 4356 | 4057 | 4167 | 4281 | 3783 | 3881 | 3800 | 3789 |
| K11070 spermidine/putrescine transport system permease protein | 3718 | 3764 | 3752 | 3871 | 3335 | 3248 | 3220 | 3292 | 2712 | 2761 | 2833 | 2914 |
| K11071 spermidine/putrescine transport system permease protein | 4117 | 4067 | 3997 | 3992 | 3241 | 3289 | 3182 | 3386 | 3059 | 3085 | 3023 | 2967 |
| K11072 spermidine/putrescine transport system ATP-binding protein [EC:7.6.2.11] | 5793 | 5782 | 5714 | 5778 | 5372 | 5378 | 5359 | 5353 | 4753 | 4609 | 4770 | 4703 |
| K11073 putrescine transport system substrate-binding protein | 9814 | 9998 | 9252 | 9772 | 5568 | 5291 | 5207 | 5640 | 3824 | 3884 | 3541 | 3705 |
| K11074 putrescine transport system permease protein | 1880 | 1766 | 1725 | 1721 | 1083 | 1057 | 1113 | 1087 | 1159 | 1134 | 1156 | 1065 |
| K11075 putrescine transport system permease protein | 2021 | 2159 | 2083 | 2159 | 1297 | 1319 | 1213 | 1299 | 1148 | 1136 | 1161 | 1119 |
| K11076 putrescine transport system ATP-binding protein | 2267 | 2190 | 2339 | 2242 | 1350 | 1431 | 1263 | 1395 | 1416 | 1444 | 1489 | 1453 |
| K11077 mannopine transport system substrate-binding protein | 2 | 0 | 0 | 0 | 0 | 0 | 0 | 2 | 0 | 4 | 0 | 0 |
| K11078 mannopine transport system permease protein | 1 | 2 | 2 | 0 | 0 | 0 | 0 | 0 | 1 | 3 | 3 | 1 |
| K11079 mannopine transport system permease protein | 0 | 0 | 0 | 0 | 0 | 0 | 0 | 0 | 0 | 1 | 2 | 1 |
| K11080 mannopine transport system ATP-binding protein | 0 | 0 | 1 | 1 | 7 | 0 | 0 | 2 | 2 | 1 | 3 | 0 |
| K11081 2-aminoethylphosphonate transport system substrate-binding protein | 434 | 413 | 469 | 470 | 127 | 113 | 121 | 131 | 835 | 946 | 807 | 871 |
| K11082 2-aminoethylphosphonate transport system permease protein | 253 | 283 | 266 | 298 | 71 | 68 | 70 | 65 | 479 | 489 | 446 | 539 |
| K11083 2-aminoethylphosphonate transport system permease protein | 309 | 333 | 292 | 348 | 84 | 101 | 96 | 69 | 594 | 660 | 609 | 653 |
| K11084 2-aminoethylphosphonate transport system ATP-binding protein | 492 | 491 | 427 | 540 | 151 | 161 | 124 | 134 | 964 | 1048 | 885 | 998 |
| K11085 ATP-binding cassette, subfamily B, bacterial MsbA [EC:3.6.3.-] | 2478 | 2726 | 2768 | 2788 | 5640 | 5360 | 5728 | 5493 | 4930 | 4875 | 4906 | 4912 |
| K11604 manganese/iron transport system substrate-binding protein | 371 | 375 | 350 | 412 | 98 | 100 | 110 | 102 | 786 | 844 | 757 | 753 |
| K11605 manganese/iron transport system permease protein | 316 | 328 | 332 | 333 | 84 | 73 | 93 | 91 | 626 | 718 | 579 | 635 |
| K11606 manganese/iron transport system permease protein | 267 | 286 | 233 | 224 | 75 | 92 | 103 | 89 | 522 | 566 | 564 | 590 |
| K11607 manganese/iron transport system ATP-binding protein | 335 | 319 | 310 | 361 | 70 | 86 | 90 | 91 | 555 | 662 | 572 | 648 |
| K11631 bacitracin transport system ATP-binding protein | 0 | 3 | 3 | 3 | 3 | 5 | 1 | 3 | 2 | 2 | 5 | 3 |
| K11632 bacitracin transport system permease protein | 2 | 3 | 0 | 1 | 0 | 0 | 0 | 0 | 0 | 0 | 0 | 0 |
| K11705 iron/zinc/manganese/copper transport system permease protein | 0 | 0 | 0 | 0 | 0 | 0 | 0 | 0 | 1 | 0 | 0 | 1 |
| K11707 manganese/zinc/iron transport system substrate-binding protein | 4 | 10 | 7 | 13 | 2 | 0 | 2 | 0 | 0 | 1 | 0 | 3 |
| K11708 manganese/zinc/iron transport system permease protein | 2 | 16 | 5 | 19 | 0 | 3 | 0 | 2 | 0 | 0 | 4 | 1 |
| K11709 manganese/zinc/iron transport system permease protein | 8 | 16 | 10 | 23 | 0 | 0 | 3 | 1 | 0 | 2 | 2 | 0 |
| K11710 manganese/zinc/iron transport system ATP- binding protein [EC:7.2.2.5] | 7 | 16 | 5 | 15 | 3 | 4 | 1 | 0 | 0 | 2 | 0 | 3 |
| K11720 lipopolysaccharide export system permease protein | 2712 | 2636 | 2690 | 2789 | 3417 | 3561 | 3445 | 3184 | 2652 | 2747 | 2803 | 2631 |
| K11953 bicarbonate transport system ATP-binding protein [EC:3.6.3.-] | 0 | 0 | 0 | 0 | 0 | 0 | 0 | 0 | 2 | 0 | 1 | 0 | 0 |
| K11955 neutral amino acid transport system permease protein | 0 | 0 | 0 | 1 | 0 | 0 | 0 | 0 | 0 | 0 | 0 | 0 |
| K11958 neutral amino acid transport system ATP-binding protein | 0 | 0 | 1 | 0 | 0 | 1 | 0 | 0 | 0 | 0 | 0 | 1 |
| K11959 urea transport system substrate-binding protein | 2731 | 2773 | 2642 | 2774 | 1360 | 1392 | 1337 | 1460 | 1960 | 1887 | 1851 | 1874 |
| K11960 urea transport system permease protein | 3481 | 3212 | 3082 | 3362 | 1720 | 1594 | 1548 | 1634 | 2183 | 2208 | 2201 | 2193 |
| K11961 urea transport system permease protein | 2023 | 1832 | 1773 | 1819 | 875 | 853 | 797 | 841 | 1416 | 1467 | 1395 | 1514 |
| K11962 urea transport system ATP-binding protein | 1572 | 1565 | 1598 | 1718 | 727 | 680 | 702 | 801 | 1303 | 1163 | 1172 | 1266 |
| K11963 urea transport system ATP-binding protein | 1490 | 1445 | 1381 | 1400 | 637 | 653 | 665 | 721 | 1045 | 1148 | 1113 | 1108 |
| K12368 dipeptide transport system substrate-binding protein | 9050 | 9007 | 8930 | 8954 | 5453 | 5432 | 5421 | 5754 | 3938 | 3966 | 3753 | 3792 |
| K12369 dipeptide transport system permease protein | 2231 | 2235 | 2167 | 2115 | 1169 | 1278 | 1279 | 1286 | 1341 | 1464 | 1351 | 1476 |
| K12370 dipeptide transport system permease protein | 1737 | 1757 | 1639 | 1753 | 1128 | 1109 | 1018 | 996 | 1143 | 1128 | 1156 | 1126 |
| K12371 dipeptide transport system ATP-binding protein | 2117 | 1909 | 1903 | 2064 | 1220 | 1222 | 1236 | 1167 | 1456 | 1318 | 1302 | 1337 |
| K12372 dipeptide transport system ATP-binding protein | 2113 | 2201 | 2082 | 2043 | 1233 | 1304 | 1228 | 1226 | 1328 | 1323 | 1247 | 1265 |
| K12536 ATP-binding cassette, subfamily C, bacterial exporter for protease/lipase | 4110 | 4027 | 3615 | 3782 | 2115 | 1930 | 1898 | 2258 | 1268 | 1166 | 1041 | 1160 |
| K12541 ATP-binding cassette, subfamily C, bacterial LapB | 4305 | 3982 | 3648 | 3962 | 2987 | 2939 | 2738 | 3068 | 1141 | 1017 | 989 | 1073 |
| K13409 ATP-binding cassette, subfamily B, bacterial RaxB | 4471 | 4504 | 4283 | 4386 | 2178 | 2241 | 2051 | 2258 | 2795 | 3111 | 2867 | 2966 |
| K13889 glutathione transport system substrate-binding protein | 1335 | 1288 | 1371 | 1349 | 1235 | 1362 | 1307 | 1313 | 1942 | 1908 | 1894 | 1999 |
| K13890 glutathione transport system permease protein | 541 | 571 | 527 | 599 | 425 | 443 | 408 | 434 | 677 | 733 | 691 | 722 |
| K13891 glutathione transport system permease protein | 579 | 604 | 569 | 556 | 440 | 455 | 384 | 431 | 780 | 801 | 738 | 743 |
| K13892 glutathione transport system ATP-binding protein | 1145 | 1195 | 1065 | 1166 | 843 | 857 | 808 | 770 | 1574 | 1616 | 1630 | 1657 |
| K13893 microcin C transport system substrate-binding protein | 7254 | 7335 | 7122 | 7214 | 4225 | 4138 | 3876 | 4182 | 3407 | 3270 | 3234 | 3416 |
| K13894 microcin C transport system permease protein | 3381 | 3511 | 3283 | 3487 | 1782 | 1881 | 1863 | 1985 | 1751 | 1765 | 1718 | 1709 |
| K13895 microcin C transport system permease protein | 3293 | 3074 | 2934 | 3035 | 1763 | 1678 | 1673 | 1779 | 1640 | 1486 | 1554 | 1525 |
| K13896 microcin C transport system ATP-binding protein | 3696 | 3704 | 3213 | 3397 | 1982 | 1910 | 1851 | 2140 | 2067 | 2101 | 2020 | 2131 |
| K14698 ATP-binding cassette, subfamily B, bacterial IrtA [EC:3.6.3.-] | 1 | 0 | 1 | 0 | 0 | 0 | 0 | 1 | 0 | 0 | 0 | 0 |
| K14699 ATP-binding cassette, subfamily B, bacterial IrtB [EC:3.6.3.-] | 17 | 18 | 19 | 22 | 8 | 10 | 1 | 0 | 1 | 4 | 8 | 9 |
| K15497 molybdate/tungstate transport system ATP-binding protein [EC:7.3.2.5 7.3.2.6] | 0 | 0 | 0 | 0 | 0 | 0 | 0 | 0 | 1 | 0 | 1 | 0 |
| K15580 oligopeptide transport system substrate-binding protein | 5868 | 5926 | 5927 | 6258 | 2553 | 2656 | 2523 | 2388 | 5978 | 6768 | 6461 | 6479 |
| K15581 oligopeptide transport system permease protein | 1627 | 1578 | 1615 | 1588 | 505 | 579 | 572 | 525 | 1586 | 1834 | 1932 | 1868 |
| K15582 oligopeptide transport system permease protein | 1770 | 1658 | 1718 | 1826 | 521 | 638 | 461 | 510 | 1844 | 2110 | 1855 | 1972 |
| K15583 oligopeptide transport system ATP-binding protein | 2354 | 2281 | 2304 | 2407 | 805 | 800 | 785 | 674 | 2312 | 2657 | 2299 | 2453 |
| K15584 nickel transport system substrate-binding protein | 1236 | 1212 | 1136 | 1129 | 836 | 861 | 883 | 799 | 1535 | 1660 | 1515 | 1630 |
| K15585 nickel transport system permease protein | 734 | 709 | 691 | 689 | 597 | 526 | 509 | 510 | 933 | 932 | 944 | 928 |
| K15586 nickel transport system permease protein | 643 | 642 | 630 | 672 | 472 | 477 | 474 | 502 | 832 | 938 | 841 | 903 |
| K15587 nickel transport system ATP-binding protein [EC:7.2.2.11] | 540 | 486 | 446 | 540 | 377 | 443 | 406 | 454 | 614 | 705 | 606 | 692 |
| K15598 putative hydroxymethylpyrimidine transport system substrate-binding protein | 336 | 365 | 284 | 267 | 101 | 110 | 60 | 69 | 706 | 749 | 725 | 708 |
| K15599 putative hydroxymethylpyrimidine transport system permease protein | 230 | 281 | 231 | 229 | 72 | 59 | 68 | 79 | 507 | 581 | 529 | 490 |
| K15600 putative hydroxymethylpyrimidine transport system ATP-binding protein | 290 | 221 | 247 | 296 | 66 | 87 | 64 | 61 | 576 | 545 | 552 | 552 |
| K15770 arabinogalactan oligomer / maltooligosaccharide transport system substrate-binding protein | 1019 | 999 | 1004 | 1027 | 461 | 469 | 496 | 485 | 1235 | 1264 | 1246 | 1317 |
| K15771 arabinogalactan oligomer / maltooligosaccharide transport system permease protein | 1257 | 1202 | 1364 | 1273 | 459 | 528 | 490 | 551 | 1380 | 1434 | 1459 | 1408 |
| K15772 arabinogalactan oligomer / maltooligosaccharide transport system permease protein | 873 | 839 | 875 | 884 | 354 | 399 | 401 | 348 | 1019 | 1051 | 925 | 1044 |
| K16012 ATP-binding cassette, subfamily C, bacterial CydC | 1016 | 1134 | 1010 | 1099 | 666 | 702 | 730 | 676 | 1429 | 1366 | 1358 | 1531 |
| K16013 ATP-binding cassette, subfamily C, bacterial CydD | 1120 | 1174 | 1096 | 1166 | 756 | 812 | 752 | 722 | 1388 | 1588 | 1456 | 1580 |
| K16014 ATP-binding cassette, subfamily C, bacterial CydCD | 41 | 48 | 36 | 53 | 8 | 6 | 8 | 9 | 11 | 23 | 6 | 9 |
| K16199 dipeptide transport system substrate-binding protein | 17 | 8 | 7 | 7 | 0 | 4 | 2 | 2 | 6 | 5 | 7 | 7 |
| K16200 dipeptide transport system permease protein | 1 | 1 | 2 | 1 | 0 | 1 | 0 | 0 | 2 | 2 | 2 | 0 |
| K16201 dipeptide transport system permease protein | 16 | 5 | 16 | 9 | 4 | 2 | 5 | 2 | 13 | 9 | 11 | 5 |
| K16202 dipeptide transport system ATP-binding protein | 19 | 16 | 16 | 13 | 3 | 4 | 5 | 2 | 25 | 24 | 24 | 17 |
| K16299 ATP-binding cassette, subfamily C, bacterial EexD | 17 | 28 | 26 | 9 | 1 | 5 | 3 | 2 | 9 | 14 | 10 | 17 |

| | | | | | | | | | | | | |
|---|---|---|---|---|---|---|---|---|---|---|---|---|
| K16784 biotin transport system ATP-binding protein [EC:3.6.3.-] | 0 | 2 | 0 | 0 | 4 | 3 | 2 | 2 | 6 | 5 | 7 | 0 |
| K16785 energy-coupling factor transport system permease protein | 1679 | 1505 | 1694 | 1719 | 523 | 516 | 523 | 488 | 2110 | 2185 | 2033 | 2177 |
| K16786 energy-coupling factor transport system ATP-binding protein [EC:3.6.3.-] | 793 | 701 | 766 | 724 | 213 | 195 | 197 | 169 | 618 | 800 | 753 | 757 |
| K16787 energy-coupling factor transport system ATP-binding protein [EC:3.6.3.-] | 2391 | 2213 | 2272 | 2302 | 784 | 805 | 811 | 776 | 2586 | 2886 | 2788 | 2956 |
| K16905 fluoroquinolone transport system permease protein | 561 | 498 | 575 | 567 | 63 | 103 | 82 | 67 | 408 | 463 | 454 | 495 |
| K16906 fluoroquinolone transport system permease protein | 406 | 365 | 385 | 418 | 79 | 80 | 64 | 69 | 339 | 347 | 354 | 420 |
| K16907 fluoroquinolone transport system ATP-binding protein [EC:3.6.3.-] | 547 | 594 | 614 | 555 | 101 | 118 | 102 | 86 | 611 | 651 | 712 | 620 |
| K16915 nickel transport protein | 14 | 13 | 19 | 16 | 14 | 14 | 15 | 19 | 13 | 9 | 27 | 19 |
| K16916 putative peptide transport system permease protein | 0 | 0 | 0 | 2 | 0 | 0 | 0 | 2 | 0 | 0 | 0 | 0 |
| K16918 acetoin utilization transport system permease protein | 0 | 0 | 0 | 0 | 0 | 0 | 0 | 2 | 0 | 0 | 0 | 0 |
| K16920 acetoin utilization transport system ATP-binding protein | 0 | 0 | 0 | 0 | 0 | 0 | 0 | 0 | 0 | 0 | 1 | 0 |
| K16956 L-cystine transport system substrate-binding protein | 0 | 2 | 1 | 2 | 0 | 0 | 0 | 0 | 0 | 2 | 0 | 0 |
| K16957 L-cystine transport system substrate-binding protein | 56 | 57 | 55 | 58 | 1 | 4 | 6 | 0 | 27 | 22 | 29 | 31 |
| K16958 L-cystine transport system permease protein | 91 | 65 | 65 | 88 | 4 | 4 | 4 | 3 | 46 | 43 | 48 | 47 |
| K16959 L-cystine transport system permease protein | 23 | 30 | 38 | 35 | 8 | 3 | 7 | 6 | 30 | 22 | 28 | 31 |
| K16960 L-cystine transport system ATP-binding protein [EC:7.4.2.1] | 200 | 191 | 158 | 225 | 15 | 17 | 29 | 20 | 110 | 125 | 119 | 141 |
| K16961 putative S-methylcysteine transport system substrate-binding protein | 126 | 124 | 127 | 110 | 2 | 7 | 3 | 10 | 82 | 76 | 80 | 84 |
| K16962 putative S-methylcysteine transport system permease protein | 158 | 156 | 176 | 144 | 6 | 5 | 11 | 4 | 56 | 97 | 100 | 99 |
| K16963 putative S-methylcysteine transport system ATP-binding protein | 227 | 188 | 188 | 193 | 8 | 11 | 9 | 4 | 94 | 121 | 114 | 137 |
| K17062 arginine/lysine/histidine/glutamine transport system substrate-binding and permease protein | 1 | 2 | 0 | 2 | 1 | 1 | 2 | 0 | 1 | 4 | 2 | 3 |
| K17063 arginine/lysine/histidine/glutamine transport system ATP-binding protein [EC:7.4.2.1] | 0 | 0 | 0 | 0 | 0 | 0 | 0 | 1 | 0 | 0 | 0 | 0 |
| K17073 putative lysine transport system substrate-binding protein | 293 | 314 | 322 | 336 | 83 | 120 | 91 | 72 | 207 | 257 | 219 | 256 |
| K17074 putative lysine transport system permease protein | 567 | 586 | 592 | 564 | 127 | 160 | 140 | 123 | 430 | 564 | 461 | 500 |
| K17076 putative lysine transport system ATP-binding protein [EC:3.6.3.-] | 508 | 486 | 499 | 521 | 159 | 118 | 134 | 118 | 402 | 459 | 385 | 415 |
| K17077 arginine/lysine/histidine transport system permease protein | 478 | 510 | 500 | 532 | 138 | 138 | 126 | 125 | 415 | 489 | 451 | 424 |
| K17202 erythritol transport system substrate-binding protein | 495 | 519 | 555 | 580 | 384 | 354 | 352 | 323 | 700 | 734 | 746 | 728 |
| K17203 erythritol transport system permease protein | 667 | 623 | 598 | 624 | 424 | 421 | 437 | 429 | 878 | 892 | 838 | 823 |
| K17204 erythritol transport system ATP-binding protein | 923 | 980 | 878 | 945 | 604 | 665 | 658 | 582 | 1219 | 1259 | 1134 | 1268 |
| K17205 putative xylitol transport system substrate-binding protein | 3 | 2 | 1 | 1 | 0 | 4 | 0 | 0 | 7 | 6 | 3 | 5 |
| K17206 putative xylitol transport system permease protein | 32 | 40 | 33 | 36 | 3 | 3 | 3 | 1 | 47 | 40 | 20 | 31 |
| K17207 putative xylitol transport system ATP-binding protein | 1 | 1 | 1 | 1 | 2 | 1 | 0 | 2 | 0 | 10 | 5 | 4 |
| K17209 inositol transport system permease protein | 11 | 9 | 19 | 15 | 2 | 4 | 3 | 1 | 7 | 5 | 5 | 7 |
| K17210 inositol transport system ATP-binding protein | 3 | 1 | 1 | 2 | 2 | 2 | 1 | 0 | 0 | 0 | 2 | 1 |
| K17213 inositol transport system substrate-binding protein | 718 | 749 | 698 | 784 | 186 | 163 | 217 | 188 | 1420 | 1440 | 1367 | 1550 |
| K17214 inositol transport system permease protein | 393 | 428 | 392 | 399 | 105 | 124 | 113 | 110 | 845 | 869 | 839 | 872 |
| K17215 inositol transport system ATP-binding protein | 673 | 766 | 624 | 687 | 207 | 217 | 191 | 192 | 1215 | 1213 | 1197 | 1291 |
| K17234 arabinosaccharide transport system substrate-binding protein | 346 | 321 | 325 | 302 | 103 | 131 | 123 | 110 | 408 | 439 | 440 | 479 |
| K17235 arabinosaccharide transport system permease protein | 366 | 294 | 322 | 303 | 100 | 83 | 108 | 100 | 420 | 403 | 374 | 421 |
| K17236 arabinosaccharide transport system permease protein | 253 | 264 | 248 | 264 | 102 | 112 | 119 | 116 | 326 | 343 | 322 | 361 |
| K17237 inositol-phosphate transport system substrate-binding protein | 3 | 0 | 0 | 3 | 4 | 6 | 0 | 2 | 2 | 1 | 5 | 2 |
| K17238 inositol-phosphate transport system permease protein | 5 | 4 | 2 | 3 | 13 | 27 | 10 | 9 | 3 | 5 | 3 | 3 |
| K17239 inositol-phosphate transport system permease protein | 2 | 2 | 0 | 3 | 14 | 8 | 1 | 13 | 1 | 5 | 3 | 1 |
| K17240 inositol-phosphate transport system ATP-binding protein | 32 | 41 | 36 | 43 | 36 | 53 | 40 | 37 | 22 | 44 | 24 | 23 |
| K17241 alpha-1,4-digalacturonate transport system substrate-binding protein | 181 | 154 | 179 | 172 | 61 | 51 | 27 | 32 | 109 | 142 | 123 | 121 |
| K17242 alpha-1,4-digalacturonate transport system permease protein | 251 | 235 | 197 | 226 | 55 | 59 | 45 | 45 | 159 | 159 | 126 | 152 |
| K17243 alpha-1,4-digalacturonate transport system permease protein | 374 | 364 | 377 | 376 | 67 | 59 | 72 | 89 | 233 | 248 | 203 | 257 |
| K17244 putative chitobiose transport system substrate-binding protein | 1 | 2 | 1 | 1 | 0 | 0 | 0 | 0 | 0 | 0 | 0 | 0 |
| K17245 putative chitobiose transport system permease protein | 64 | 65 | 58 | 71 | 17 | 26 | 18 | 17 | 68 | 63 | 65 | 90 |
| K17246 putative chitobiose transport system permease protein | 4 | 5 | 9 | 7 | 2 | 5 | 1 | 0 | 9 | 12 | 10 | 10 |
| K17311 trehalose transport system substrate-binding protein | 1 | 1 | 5 | 3 | 0 | 0 | 0 | 1 | 0 | 0 | 3 | 0 |
| K17312 trehalose transport system permease protein | 3 | 2 | 4 | 3 | 1 | 1 | 1 | 0 | 3 | 2 | 5 | 8 |
| K17313 trehalose transport system permease protein | 1 | 2 | 5 | 1 | 1 | 0 | 1 | 2 | 0 | 0 | 0 | 1 |
| K17315 glucose/mannose transport system substrate-binding protein | 2006 | 1850 | 1807 | 1836 | 1049 | 1011 | 1025 | 1049 | 633 | 543 | 526 | 589 |
| K17316 glucose/mannose transport system permease protein | 1333 | 1391 | 1424 | 1384 | 780 | 719 | 702 | 826 | 452 | 384 | 410 | 404 |
| K17317 glucose/mannose transport system permease protein | 1106 | 1132 | 1131 | 1085 | 611 | 603 | 607 | 655 | 369 | 356 | 349 | 317 |
| K17318 putative aldouronate transport system substrate-binding protein | 9680 | 9526 | 9810 | 9727 | 2005 | 2019 | 2054 | 1975 | 13413 | 13975 | 13772 | 14136 |
| K17319 putative aldouronate transport system permease protein | 11884 | 11815 | 12003 | 11855 | 2376 | 2394 | 2334 | 2297 | 16531 | 17696 | 17315 | 17904 |
| K17320 putative aldouronate transport system permease protein | 11366 | 11243 | 11559 | 11595 | 2314 | 2269 | 2113 | 2060 | 15701 | 16901 | 16601 | 16970 |
| K17321 glycerol transport system substrate-binding protein | 0 | 1 | 0 | 1 | 3 | 0 | 0 | 3 | 7 | 2 | 13 | 3 |
| K17322 glycerol transport system permease protein | 3 | 0 | 1 | 4 | 0 | 0 | 0 | 0 | 4 | 1 | 3 | 1 |
| K17323 glycerol transport system permease protein | 5 | 6 | 6 | 9 | 0 | 1 | 2 | 3 | 4 | 5 | 6 | 10 |
| K17324 glycerol transport system ATP-binding protein | 0 | 1 | 0 | 0 | 0 | 2 | 0 | 0 | 0 | 2 | 3 | 5 |
| K17325 glycerol transport system ATP-binding protein | 3 | 2 | 2 | 3 | 0 | 0 | 2 | 2 | 2 | 1 | 4 | 1 |
| K17326 xylobiose transport system substrate-binding protein | 6 | 6 | 5 | 7 | 7 | 0 | 1 | 2 | 8 | 12 | 13 | 15 |
| K17327 xylobiose transport system permease protein | 0 | 2 | 2 | 2 | 3 | 0 | 0 | 0 | 0 | 4 | 2 | 1 |
| K17328 xylobiose transport system permease protein | 0 | 2 | 3 | 1 | 3 | 2 | 2 | 2 | 0 | 4 | 2 | 0 |
| K17329 N,N'-diacetylchitobiose transport system substrate-binding protein | 16 | 19 | 25 | 17 | 6 | 3 | 6 | 6 | 7 | 14 | 16 | 13 |
| K17330 N,N'-diacetylchitobiose transport system permease protein | 9 | 8 | 9 | 7 | 3 | 3 | 1 | 1 | 10 | 3 | 14 | 10 |
| K17331 N,N'-diacetylchitobiose transport system permease protein | 43 | 36 | 41 | 35 | 4 | 6 | 3 | 9 | 44 | 29 | 35 | 33 |
| K18104 ATP-binding cassette, subfamily B, bacterial AbcA/BmrA [EC:7.6.2.2] | 25 | 12 | 26 | 17 | 7 | 6 | 14 | 13 | 22 | 39 | 28 | 23 |
| K18216 ATP-binding cassette, subfamily B, tetracycline resistant protein | 3 | 3 | 3 | 9 | 0 | 0 | 0 | 0 | 1 | 0 | 2 | 0 |
| K18217 ATP-binding cassette, subfamily B, tetracycline resistant protein | 6 | 0 | 0 | 7 | 0 | 3 | 0 | 3 | 0 | 4 | 1 | 0 |
| K18230 macrolide transport system ATP-binding/permease protein | 0 | 1 | 0 | 1 | 0 | 0 | 1 | 4 | 0 | 1 | 0 | 0 |
| K18231 macrolide transport system ATP-binding/permease protein | 98 | 118 | 87 | 128 | 17 | 37 | 41 | 29 | 73 | 88 | 87 | 106 |
| K18887 ATP-binding cassette, subfamily B, multidrug efflux pump | 618 | 563 | 567 | 561 | 123 | 154 | 120 | 116 | 443 | 468 | 484 | 517 |
| K18888 ATP-binding cassette, subfamily B, multidrug efflux pump | 429 | 420 | 478 | 467 | 147 | 142 | 84 | 93 | 404 | 483 | 378 | 376 |
| K18889 ATP-binding cassette, subfamily B, multidrug efflux pump | 1119 | 1145 | 1148 | 1243 | 758 | 720 | 851 | 786 | 1519 | 1610 | 1552 | 1583 |
| K18890 ATP-binding cassette, subfamily B, multidrug efflux pump | 1264 | 1171 | 1209 | 1243 | 733 | 827 | 855 | 748 | 1743 | 1660 | 1653 | 1763 |
| K18891 ATP-binding cassette, subfamily B, multidrug efflux pump | 79 | 118 | 107 | 88 | 14 | 26 | 28 | 30 | 86 | 110 | 71 | 97 |
| K18892 ATP-binding cassette, subfamily B, multidrug efflux pump | 104 | 64 | 125 | 104 | 47 | 43 | 44 | 31 | 99 | 151 | 114 | 130 |
| K18893 ATP-binding cassette, subfamily B, multidrug efflux pump | 2737 | 2631 | 2668 | 2588 | 1465 | 1468 | 1340 | 1464 | 922 | 827 | 795 | 816 |
| K18895 ATP-binding cassette, subfamily B, salmochelin/enterobactin exporter | 12 | 9 | 5 | 7 | 8 | 2 | 8 | 5 | 221 | 175 | 204 | 213 |
| K19083 bacitracin transport system ATP-binding protein | 0 | 0 | 1 | 0 | 0 | 0 | 0 | 0 | 0 | 1 | 1 | 0 |
| K19226 cationic peptide transport system substrate-binding protein | 850 | 838 | 841 | 928 | 616 | 668 | 619 | 570 | 1259 | 1238 | 1223 | 1323 |
| K19227 cationic peptide transport system permease protein | 462 | 470 | 466 | 546 | 305 | 341 | 343 | 319 | 683 | 736 | 638 | 732 |
| K19228 cationic peptide transport system permease protein | 508 | 443 | 499 | 490 | 276 | 319 | 371 | 308 | 663 | 650 | 665 | 661 |
| K19229 cationic peptide transport system ATP-binding protein | 521 | 521 | 498 | 524 | 350 | 396 | 379 | 389 | 782 | 818 | 786 | 787 |
| K19230 cationic peptide transport system ATP-binding protein | 450 | 431 | 467 | 461 | 273 | 340 | 318 | 297 | 674 | 618 | 640 | 681 |
| K19309 bacitracin transport system ATP-binding protein | 467 | 420 | 429 | 433 | 41 | 49 | 49 | 24 | 414 | 420 | 394 | 434 |
| K19310 bacitracin transport system permease protein | 395 | 401 | 410 | 449 | 40 | 31 | 34 | 27 | 400 | 433 | 407 | 433 |
| K19340 Cu-processing system ATP-binding protein | 1510 | 1468 | 1289 | 1390 | 740 | 679 | 701 | 819 | 458 | 352 | 401 | 415 |
| K19341 Cu-processing system permease protein | 1193 | 1085 | 1012 | 1109 | 486 | 596 | 602 | 747 | 335 | 285 | 333 | 321 |
| K19349 pleuromutilin/lincosamide/streptogramin A transport system ATP-binding/permease protein | 6 | 12 | 7 | 11 | 0 | 0 | 0 | 0 | 21 | 20 | 20 | 18 |
| K19350 lincosamide and streptogramin A transport system ATP-binding/permease protein | 205 | 246 | 217 | 241 | 83 | 98 | 85 | 79 | 343 | 386 | 338 | 386 |
| K19971 manganese/zinc transport system substrate-binding protein | 53 | 71 | 51 | 81 | 3 | 10 | 3 | 3 | 41 | 57 | 38 | 54 |
| K19972 manganese/zinc transport system permease protein | 28 | 27 | 53 | 45 | 9 | 10 | 8 | 3 | 39 | 50 | 45 | 43 |
| K19973 manganese transport system ATP-binding protein [EC:7.2.2.5] | 37 | 30 | 48 | 51 | 19 | 9 | 15 | 12 | 30 | 62 | 44 | 47 |
| K19975 manganese transport system substrate-binding protein | 45 | 45 | 63 | 64 | 9 | 10 | 8 | 4 | 50 | 36 | 36 | 65 |
| K19976 manganese transport system permease protein | 44 | 56 | 35 | 35 | 17 | 21 | 31 | 20 | 49 | 66 | 40 | 45 |
| K20344 ATP-binding cassette, subfamily C, bacteriocin exporter | 328 | 342 | 385 | 340 | 418 | 459 | 468 | 418 | 169 | 239 | 190 | 248 |
| K20386 ATP-binding cassette, subfamily B, bacterial CylB | 131 | 185 | 170 | 151 | 3 | 6 | 12 | 13 | 11 | 15 | 12 | 11 |
| K20459 lantibiotic transport system ATP-binding protein | 166 | 156 | 173 | 163 | 66 | 39 | 41 | 43 | 283 | 276 | 267 | 291 |
| K20460 lantibiotic transport system permease protein | 68 | 70 | 83 | 61 | 39 | 31 | 40 | 29 | 101 | 130 | 112 | 163 |
| K20461 lantibiotic transport system permease protein | 117 | 114 | 136 | 118 | 38 | 44 | 46 | 44 | 199 | 210 | 208 | 206 |
| K20490 lantibiotic transport system ATP-binding protein | 72 | 59 | 74 | 53 | 46 | 51 | 43 | 38 | 106 | 85 | 73 | 88 |
| K20491 lantibiotic transport system permease protein | 75 | 51 | 60 | 64 | 38 | 42 | 46 | 30 | 90 | 106 | 90 | 95 |
| K20492 lantibiotic transport system permease protein | 63 | 59 | 45 | 45 | 25 | 20 | 35 | 36 | 61 | 92 | 102 | 123 |
| K23055 putative lysine/arginine/ornithine/histidine/octopine transport system substrate-binding protein | 79 | 65 | 72 | 62 | 51 | 57 | 29 | 51 | 21 | 22 | 29 | 28 |
| K23056 putative lysine/arginine/ornithine/histidine/octopine transport system permease protein | 1278 | 1172 | 1144 | 1292 | 697 | 652 | 591 | 643 | 374 | 368 | 377 | 387 |
| K23057 putative lysine/arginine/ornithine/histidine/octopine transport system permease protein | 1063 | 1130 | 1077 | 1094 | 618 | 601 | 577 | 577 | 365 | 350 | 327 | 348 |
| K23058 putative lysine/arginine/ornithine/histidine/octopine transport system ATP-binding protein [EC:7.4.2.1] | 1068 | 1164 | 1077 | 1020 | 647 | 612 | 547 | 615 | 383 | 332 | 335 | 311 |
| K23059 arginine/lysine/histidine transporter system substrate-binding protein | 317 | 293 | 356 | 374 | 111 | 89 | 118 | 101 | 267 | 284 | 298 | 299 |
| K23060 arginine/lysine/histidine transport system ATP-binding protein [EC:7.4.2.1] | 366 | 410 | 372 | 345 | 116 | 124 | 148 | 125 | 296 | 380 | 316 | 343 |
| K23061 hydroxyproline transporter system substrate-binding protein | 1154 | 1139 | 1101 | 1055 | 635 | 552 | 589 | 596 | 379 | 341 | 281 | 348 |
| K23062 hydroxyproline transport system permease protein | 953 | 831 | 796 | 783 | 471 | 470 | 443 | 532 | 276 | 227 | 238 | 238 |
| K23063 hydroxyproline transport system permease protein | 840 | 885 | 837 | 815 | 477 | 405 | 408 | 491 | 312 | 236 | 248 | 246 |
| K23064 hydroxyproline transport system ATP-binding protein [EC:7.4.2.1] | 678 | 657 | 583 | 627 | 358 | 366 | 313 | 337 | 209 | 214 | 199 | 179 |
| K23181 ferric citrate transport system substrate-binding protein | 3 | 9 | 0 | 1 | 4 | 3 | 6 | 3 | 51 | 56 | 47 | 50 |
| K23182 ferric citrate transport system permease protein | 5 | 6 | 9 | 3 | 10 | 6 | 0 | 2 | 81 | 87 | 74 | 70 |
| K23183 ferric citrate transport system permease protein | 0 | 1 | 0 | 1 | 1 | 1 | 0 | 2 | 47 | 66 | 38 | 50 |
| K23184 ferric citrate transport system ATP-binding protein [EC:7.2.2.18] | 32 | 24 | 30 | 38 | 18 | 12 | 18 | 19 | 66 | 58 | 56 | 38 |

| | | | | | | | | | | | | |
|---|---|---|---|---|---|---|---|---|---|---|---|---|
| K23185 ferric enterobactin transport system substrate-binding protein | 541 | 605 | 436 | 574 | 394 | 406 | 395 | 414 | 744 | 825 | 785 | 784 |
| K23186 ferric enterobactin transport system permease protein | 588 | 502 | 472 | 572 | 396 | 454 | 427 | 358 | 728 | 718 | 719 | 783 |
| K23187 ferric enterobactin transport system permease protein | 628 | 537 | 481 | 550 | 413 | 371 | 390 | 374 | 772 | 776 | 679 | 848 |
| K23188 ferric enterobactin transport system ATP-binding protein [EC:7.2.2.17] | 587 | 636 | 561 | 595 | 393 | 414 | 381 | 411 | 738 | 737 | 702 | 751 |
| K23227 ferric hydroxamate transport system substrate-binding protein | 575 | 533 | 530 | 597 | 361 | 420 | 353 | 378 | 888 | 857 | 832 | 811 |
| K23228 ferric hydroxamate transport system permease protein | 1162 | 1148 | 1012 | 1230 | 960 | 938 | 819 | 859 | 1493 | 1528 | 1485 | 1597 |
| ko02060 Phosphotransferase system (PTS) | 60928 | 59813 | 58541 | 61350 | 37321 | 38082 | 37156 | 36742 | 66414 | 70443 | 66735 | 69633 |
| K02757 PTS system, beta-glucoside-specific IIC component | 3583 | 3454 | 3436 | 3590 | 2480 | 2627 | 2532 | 2287 | 2783 | 3113 | 2751 | 2912 |
| K02784 phosphocarrier protein HPr | 143 | 157 | 193 | 150 | 107 | 98 | 126 | 124 | 240 | 242 | 240 | 217 |
| K02806 PTS system, nitrogen regulatory IIA component [EC:2.7.1.-] | 1131 | 1124 | 1011 | 1130 | 665 | 675 | 639 | 744 | 696 | 734 | 659 | 679 |
| K08483 phosphotransferase system, enzyme I, PtsI [EC:2.7.3.9] | 1980 | 1872 | 2001 | 2019 | 1128 | 1192 | 1203 | 1097 | 2345 | 2657 | 2402 | 2470 |
| K08484 phosphotransferase system, enzyme I, PtsP [EC:2.7.3.9] | 5022 | 4883 | 4614 | 4885 | 2967 | 2906 | 2937 | 3070 | 3073 | 3027 | 2846 | 3100 |
| K08485 phosphocarrier protein NPr | 161 | 161 | 176 | 178 | 109 | 128 | 123 | 94 | 222 | 277 | 249 | 230 |
| K11183 phosphocarrier protein FPr | 1193 | 1090 | 1054 | 1075 | 671 | 712 | 709 | 739 | 1113 | 1201 | 1106 | 1175 |
| K11200 PTS system, 2-O-A-mannosyl-D-glycerate-specific IIC component | 4 | 7 | 3 | 5 | 19 | 16 | 18 | 26 | 6 | 10 | 17 | 8 |
| K19506 PTS system, fructoselysine/glucoselysine-specific IIA component [EC:2.7.1.-] | 308 | 316 | 283 | 328 | 68 | 79 | 57 | 63 | 442 | 539 | 483 | 471 |
| K19507 PTS system, fructoselysine/glucoselysine-specific IIB component [EC:2.7.1.-] | 327 | 330 | 359 | 348 | 87 | 90 | 127 | 76 | 522 | 529 | 474 | 521 |
| K19508 PTS system, fructoselysine/glucoselysine-specific IIC component | 619 | 601 | 573 | 674 | 127 | 156 | 148 | 155 | 885 | 935 | 814 | 907 |
| K19509 PTS system, fructoselysine/glucoselysine-specific IID component | 902 | 921 | 896 | 888 | 176 | 172 | 171 | 171 | 931 | 1050 | 943 | 1009 |
| K20112 PTS system, galactose-specific IIA component [EC:2.7.1.204] | 29 | 30 | 26 | 27 | 0 | 7 | 1 | 3 | 21 | 16 | 33 | 15 |
| K20113 PTS system, galactose-specific IIB component [EC:2.7.1.204] | 5 | 22 | 19 | 11 | 3 | 5 | 0 | 2 | 11 | 12 | 14 | 12 |
| K20114 PTS system, galactose-specific IIC component | 66 | 47 | 76 | 84 | 16 | 16 | 24 | 16 | 61 | 88 | 71 | 80 |
| ko03070 Bacterial secretion system | 196395 | 195950 | 188110 | 191797 | 135823 | 135607 | 131661 | 137413 | 126014 | 124728 | 121059 | 124342 |
| K02452 general secretion pathway protein C | 1997 | 1941 | 1703 | 1893 | 957 | 952 | 866 | 960 | 1197 | 1203 | 1073 | 1257 |
| K02453 general secretion pathway protein D | 10056 | 9959 | 9079 | 9366 | 4893 | 4857 | 4591 | 5256 | 4562 | 4546 | 4268 | 4511 |
| K02454 general secretion pathway protein E [EC:7.4.2.8] | 7020 | 7015 | 6450 | 6689 | 3444 | 3402 | 3400 | 3687 | 3381 | 3247 | 3139 | 3221 |
| K02455 general secretion pathway protein F | 4872 | 4697 | 4366 | 4396 | 2400 | 2183 | 2113 | 2643 | 2550 | 2268 | 2285 | 2451 |
| K02456 general secretion pathway protein G | 2645 | 2612 | 2544 | 2654 | 1311 | 1203 | 1153 | 1246 | 1191 | 1216 | 1106 | 1158 |
| K02457 general secretion pathway protein H | 1422 | 1483 | 1352 | 1343 | 740 | 623 | 647 | 751 | 795 | 858 | 741 | 748 |
| K02458 general secretion pathway protein I | 1848 | 1651 | 1582 | 1714 | 987 | 888 | 855 | 903 | 800 | 803 | 769 | 795 |
| K02459 general secretion pathway protein J | 2988 | 2992 | 2671 | 2811 | 1520 | 1469 | 1328 | 1553 | 1411 | 1260 | 1171 | 1328 |
| K02460 general secretion pathway protein K | 3033 | 2986 | 2819 | 2924 | 1488 | 1466 | 1390 | 1522 | 1714 | 1605 | 1542 | 1768 |
| K02461 general secretion pathway protein L | 4283 | 4211 | 3641 | 3977 | 1954 | 1956 | 1940 | 2150 | 2394 | 2094 | 2013 | 2227 |
| K02462 general secretion pathway protein M | 1575 | 1474 | 1347 | 1527 | 794 | 740 | 697 | 769 | 778 | 759 | 708 | 869 |
| K02464 general secretion pathway protein O [EC:3.4.23.43 2.1.1.-] | 318 | 383 | 282 | 295 | 81 | 92 | 75 | 82 | 650 | 692 | 673 | 688 |
| K02465 general secretion pathway protein S | 144 | 150 | 171 | 185 | 37 | 41 | 40 | 32 | 303 | 316 | 283 | 318 |
| K03194 type IV secretion system protein VirB1 | 261 | 250 | 226 | 241 | 60 | 78 | 63 | 73 | 546 | 532 | 468 | 498 |
| K03195 type IV secretion system protein VirB10 | 368 | 396 | 363 | 414 | 91 | 108 | 81 | 109 | 826 | 751 | 732 | 786 |
| K03196 type IV secretion system protein VirB11 [EC:7.4.2.8] | 338 | 363 | 340 | 373 | 127 | 118 | 93 | 107 | 718 | 845 | 784 | 813 |
| K03197 type IV secretion system protein VirB2 | 111 | 109 | 97 | 138 | 30 | 36 | 26 | 11 | 218 | 203 | 213 | 224 |
| K03198 type IV secretion system protein VirB3 | 0 | 0 | 0 | 0 | 0 | 0 | 0 | 0 | 3 | 3 | 0 | 2 |
| K03199 type IV secretion system protein VirB4 [EC:7.4.2.8] | 941 | 982 | 1029 | 1041 | 273 | 289 | 274 | 263 | 2076 | 2407 | 2219 | 2147 |
| K03200 type IV secretion system protein VirB5 | 244 | 241 | 277 | 258 | 58 | 77 | 67 | 75 | 547 | 553 | 575 | 574 |
| K03201 type IV secretion system protein VirB6 | 358 | 371 | 401 | 419 | 108 | 114 | 95 | 106 | 782 | 882 | 811 | 792 |
| K03202 type IV secretion system protein VirB7 | 0 | 1 | 1 | 0 | 0 | 0 | 0 | 2 | 0 | 0 | 0 | 0 |
| K03203 type IV secretion system protein VirB8 | 233 | 233 | 248 | 231 | 46 | 76 | 66 | 57 | 461 | 474 | 408 | 463 |
| K03204 type IV secretion system protein VirB9 | 366 | 377 | 379 | 330 | 98 | 109 | 108 | 105 | 726 | 832 | 670 | 788 |
| K03205 type IV secretion system protein VirD4 [EC:7.4.2.8] | 9143 | 9101 | 9102 | 8845 | 4544 | 4597 | 4258 | 4384 | 4998 | 5468 | 5298 | 5521 |
| K03219 type III secretion protein C | 2641 | 2708 | 2581 | 2628 | 1509 | 1411 | 1296 | 1353 | 754 | 791 | 762 | 777 |
| K03221 type III secretion protein F | 326 | 347 | 338 | 362 | 180 | 159 | 201 | 217 | 94 | 119 | 88 | 94 |
| K03222 type III secretion protein J | 1036 | 1027 | 933 | 1034 | 557 | 511 | 517 | 550 | 357 | 317 | 330 | 349 |
| K03223 type III secretion protein L | 43 | 40 | 30 | 41 | 18 | 16 | 10 | 17 | 11 | 7 | 13 | 9 |
| K03224 ATP synthase in type III secretion protein N [EC:7.4.2.8] | 1288 | 1315 | 1270 | 1361 | 719 | 708 | 660 | 752 | 457 | 373 | 401 | 387 |
| K03225 type III secretion protein Q | 1406 | 1287 | 1253 | 1217 | 670 | 684 | 698 | 750 | 394 | 404 | 373 | 397 |
| K03226 type III secretion protein R | 935 | 920 | 787 | 859 | 505 | 496 | 489 | 477 | 290 | 273 | 245 | 300 |
| K03227 type III secretion protein S | 321 | 326 | 297 | 320 | 213 | 142 | 143 | 128 | 66 | 93 | 77 | 88 |
| K03228 type III secretion protein T | 873 | 727 | 741 | 740 | 393 | 420 | 527 | 622 | 251 | 199 | 202 | 227 |
| K03229 type III secretion protein U | 1372 | 1358 | 1271 | 1283 | 662 | 743 | 696 | 765 | 475 | 399 | 396 | 439 |
| K03230 type III secretion protein V | 299 | 290 | 267 | 250 | 156 | 195 | 187 | 175 | 106 | 95 | 95 | 89 |
| K04056 type III secretion protein O | 235 | 248 | 256 | 250 | 159 | 151 | 196 | 205 | 73 | 77 | 60 | 83 |
| K04057 type III secretion protein P | 316 | 308 | 277 | 312 | 151 | 189 | 131 | 161 | 107 | 95 | 87 | 105 |
| K04058 type III secretion protein W | 87 | 72 | 88 | 80 | 44 | 49 | 71 | 87 | 30 | 31 | 21 | 16 |
| K04059 type III secretion protein X | 589 | 495 | 551 | 504 | 241 | 291 | 291 | 286 | 136 | 144 | 144 | 189 |
| K11003 hemolysin D | 10 | 3 | 5 | 7 | 7 | 2 | 5 | 0 | 67 | 67 | 74 | 89 |
| K11891 type VI secretion system protein ImpL | 19635 | 19375 | 18382 | 19132 | 11317 | 11006 | 10706 | 11572 | 9536 | 9422 | 8474 | 9226 |
| K11892 type VI secretion system protein ImpK | 4963 | 4873 | 4359 | 4684 | 2868 | 2840 | 2705 | 2922 | 2267 | 2182 | 2170 | 2241 |
| K11903 type VI secretion system secreted protein Hcp | 2669 | 2663 | 2558 | 2535 | 1564 | 1617 | 1492 | 1616 | 1418 | 1468 | 1323 | 1419 |
| K11904 type VI secretion system secreted protein VgrG | 28524 | 28444 | 27959 | 28775 | 16304 | 16426 | 14901 | 15661 | 11992 | 11726 | 11627 | 11929 |
| K11906 type VI secretion system protein VasD | 2051 | 1977 | 1832 | 1845 | 1161 | 1057 | 1108 | 1107 | 1175 | 1162 | 1092 | 1136 |
| K11907 type VI secretion system protein VasG | 13354 | 13568 | 12620 | 12864 | 8095 | 8383 | 7688 | 8190 | 5367 | 5446 | 5303 | 5438 |
| K11912 serine/threonine-protein kinase PpkA [EC:2.7.11.1] | 3564 | 3692 | 3156 | 3223 | 1850 | 1750 | 1621 | 1957 | 1225 | 982 | 994 | 1128 |
| K11913 type VI secretion system protein | 1861 | 1894 | 1566 | 1652 | 963 | 907 | 976 | 1114 | 684 | 549 | 567 | 584 |
| K11915 serine/threonine protein phosphatase Stp1 [EC:3.1.3.16] | 828 | 852 | 773 | 748 | 397 | 420 | 389 | 540 | 292 | 276 | 273 | 242 |
| K12340 outer membrane protein | 6053 | 6403 | 6047 | 6105 | 7792 | 7869 | 8112 | 8336 | 6229 | 6003 | 6091 | 6137 |
| Signal transduction | 635989 | 629723 | 616992 | 628286 | 464972 | 468313 | 454285 | 465392 | 457522 | 459896 | 452900 | 462323 |
| ko02020 Two-component system | 573175 | 567064 | 554878 | 565370 | 405596 | 407925 | 395160 | 406201 | 400438 | 402988 | 396030 | 404778 |
| K00373 nitrate reductase molybdenum cofactor assembly chaperone NarJ/NarW | 1203 | 1197 | 1214 | 1246 | 660 | 628 | 604 | 620 | 1336 | 1298 | 1270 | 1340 |
| K00575 chemotaxis protein methyltransferase CheR [EC:2.1.1.80] | 1007 | 1058 | 968 | 984 | 586 | 627 | 611 | 628 | 727 | 828 | 770 | 729 |
| K00990 [protein-PII] uridylyltransferase [EC:2.7.7.59] | 5846 | 5799 | 5887 | 5891 | 3794 | 3826 | 3585 | 3552 | 3709 | 3822 | 3730 | 3876 |
| K01104 protein-tyrosine phosphatase [EC:3.1.3.48] | 1739 | 1781 | 1749 | 1683 | 1825 | 1849 | 1788 | 1744 | 1863 | 1914 | 1870 | 1823 |
| K01467 beta-lactamase class C [EC:3.5.2.6] | 4 | 12 | 4 | 1 | 9 | 13 | 18 | 16 | 59 | 78 | 72 | 61 |
| K01545 potassium-transporting ATPase KdpF subunit | 255 | 269 | 257 | 214 | 154 | 148 | 143 | 161 | 83 | 84 | 58 | 91 |
| K01546 potassium-transporting ATPase potassium-binding subunit | 4681 | 4743 | 4778 | 4638 | 5447 | 5568 | 5556 | 5511 | 5831 | 5942 | 6142 | 5851 |
| K01547 potassium-transporting ATPase ATP-binding subunit [EC:7.2.2.6] | 6384 | 6442 | 6277 | 6173 | 6839 | 6599 | 6799 | 6774 | 6674 | 6701 | 6769 | 6628 |
| K01548 potassium-transporting ATPase KdpC subunit | 1639 | 1681 | 1463 | 1622 | 1817 | 1718 | 1662 | 1760 | 1750 | 1630 | 1806 | 1682 |
| K01643 citrate lyase subunit alpha / citrate CoA-transferase [EC:2.8.3.10] | 2215 | 2057 | 2018 | 2201 | 1587 | 1522 | 1457 | 1590 | 3000 | 3135 | 2889 | 3147 |
| K01644 citrate lyase subunit beta / citryl-CoA lyase [EC:4.1.3.34] | 1279 | 1204 | 1137 | 1232 | 829 | 838 | 913 | 788 | 1679 | 1853 | 1660 | 1728 |
| K01646 citrate lyase subunit gamma (acyl carrier protein) | 393 | 428 | 424 | 388 | 262 | 310 | 277 | 319 | 580 | 576 | 553 | 564 |
| K01910 [citrate (pro-3S)-lyase] ligase [EC:6.2.1.22] | 1325 | 1377 | 1404 | 1461 | 953 | 1064 | 1005 | 954 | 2005 | 2097 | 1897 | 1992 |
| K01991 polysaccharide biosynthesis/export protein | 1639 | 1737 | 1927 | 1739 | 3526 | 3493 | 3633 | 3528 | 2772 | 2741 | 2642 | 2675 |
| K02106 short-chain fatty acids transporter | 1166 | 1190 | 1162 | 1158 | 665 | 611 | 595 | 673 | 538 | 506 | 519 | 507 |
| K02253 competence protein ComX | 0 | 0 | 0 | 2 | 0 | 0 | 0 | 0 | 0 | 0 | 0 | 0 |
| K02313 chromosomal replication initiator protein | 5654 | 5570 | 5696 | 5718 | 5497 | 5710 | 5770 | 5762 | 5510 | 5350 | 5596 | 5472 |
| K02398 negative regulator of flagellin synthesis FlgM | 348 | 379 | 371 | 385 | 278 | 285 | 377 | 359 | 146 | 176 | 140 | 178 |
| K02402 flagellar transcriptional activator FlhC | 119 | 90 | 89 | 124 | 138 | 179 | 206 | 180 | 38 | 27 | 34 | 27 |
| K02403 flagellar transcriptional activator FlhD | 137 | 124 | 112 | 99 | 180 | 188 | 156 | 174 | 27 | 25 | 12 | 26 |
| K02405 RNA polymerase sigma factor for flagellar operon FliA | 1624 | 1606 | 1622 | 1620 | 901 | 935 | 860 | 944 | 761 | 865 | 774 | 775 |
| K02406 flagellin | 1133 | 1080 | 1101 | 1137 | 1042 | 1085 | 1172 | 1120 | 509 | 615 | 505 | 519 |
| K02487 type IV pili sensor histidine kinase and response regulator | 1425 | 1259 | 1392 | 1360 | 1160 | 1186 | 1595 | 1706 | 477 | 433 | 411 | 453 |
| K02488 two-component system, cell cycle response regulator [EC:2.7.7.65] | 42 | 40 | 37 | 39 | 8 | 4 | 7 | 9 | 82 | 105 | 79 | 74 |
| K02490 two-component system, response regulator, stage 0 sporulation protein F | 6 | 2 | 15 | 8 | 0 | 2 | 0 | 0 | 4 | 2 | 7 | 4 |
| K02491 two-component system, sporulation sensor kinase A [EC:2.7.13.3] | 0 | 0 | 0 | 0 | 0 | 0 | 0 | 0 | 0 | 0 | 0 | 1 |
| K02556 chemotaxis protein MotA | 2296 | 2160 | 2140 | 2224 | 1195 | 1216 | 1153 | 1194 | 1178 | 1188 | 1087 | 1092 |
| K02584 Nif-specific regulatory protein | 373 | 365 | 307 | 366 | 372 | 452 | 437 | 457 | 54 | 50 | 51 | 80 |
| K02650 type IV pilus assembly protein PilA | 932 | 891 | 868 | 905 | 517 | 470 | 518 | 461 | 325 | 299 | 348 | 302 |
| K02657 twitching motility two-component system response regulator PilG | 745 | 722 | 740 | 703 | 446 | 395 | 378 | 449 | 266 | 221 | 200 | 240 |
| K02658 twitching motility two-component system response regulator PilH | 614 | 617 | 568 | 623 | 321 | 362 | 336 | 334 | 198 | 154 | 169 | 205 |
| K02659 twitching motility protein PilI | 721 | 707 | 689 | 700 | 379 | 446 | 384 | 400 | 257 | 212 | 234 | 196 |
| K02660 twitching motility protein PilJ | 3060 | 3093 | 2998 | 3096 | 1656 | 1807 | 1474 | 1645 | 1002 | 1002 | 976 | 954 |
| K02661 type IV pilus assembly protein PilK | 1620 | 1526 | 1447 | 1448 | 834 | 773 | 805 | 926 | 523 | 436 | 446 | 507 |
| K02667 two-component system, NtrC family, response regulator PilR | 941 | 996 | 981 | 1020 | 564 | 554 | 538 | 629 | 340 | 323 | 318 | 316 |
| K02668 two-component system, NtrC family, sensor histidine kinase PilS [EC:2.7.13.3] | 2767 | 2633 | 2772 | 2693 | 1542 | 1491 | 1446 | 1438 | 924 | 813 | 797 | 829 |
| K03092 RNA polymerase sigma-54 factor | 3637 | 3592 | 3689 | 3611 | 5026 | 5041 | 4837 | 4827 | 4527 | 4342 | 4434 | 4297 |
| K03406 methyl-accepting chemotaxis protein | 47131 | 45884 | 44271 | 45269 | 28990 | 29267 | 25343 | 26712 | 15030 | 14579 | 14353 | 14700 |
| K03407 two-component system, chemotaxis family, sensor kinase CheA [EC:2.7.13.3] | 6775 | 6826 | 6232 | 6213 | 3662 | 3508 | 3228 | 3569 | 2918 | 2742 | 2763 | 2873 |
| K03408 purine-binding chemotaxis protein CheW | 2815 | 2847 | 2804 | 2757 | 1585 | 1549 | 1458 | 1721 | 1115 | 1048 | 1061 | 1007 |
| K03412 two-component system, chemotaxis family, protein-glutamate methylesterase/glutaminase [EC:3.1.1.61 3.5.1.44] | 2468 | 2366 | 2322 | 2355 | 1454 | 1424 | 1550 | 1464 | 970 | 899 | 886 | 913 |
| K03413 two-component system, chemotaxis family, chemotaxis protein CheY | 1552 | 1554 | 1537 | 1537 | 860 | 859 | 805 | 849 | 744 | 738 | 707 | 858 |
| K03415 two-component system, chemotaxis family, chemotaxis protein CheV | 1321 | 1335 | 1327 | 1396 | 863 | 926 | 937 | 859 | 335 | 426 | 378 | 383 |
| K03563 carbon storage regulator | 466 | 545 | 514 | 534 | 239 | 256 | 232 | 216 | 421 | 405 | 389 | 428 |
| K03620 Ni/Fe-hydrogenase 1 B-type cytochrome subunit | 42 | 48 | 65 | 58 | 63 | 66 | 66 | 50 | 105 | 112 | 114 | 131 |

| | | | | | | | | | | | | |
|---|---|---|---|---|---|---|---|---|---|---|---|---|
| K03739 membrane protein involved in D-alanine export | 364 | 304 | 369 | 354 | 408 | 410 | 347 | 319 | 184 | 229 | 187 | 203 |
| K03740 D-alanine transfer protein | 345 | 389 | 353 | 405 | 382 | 404 | 407 | 345 | 141 | 219 | 234 | 224 |
| K03776 aerotaxis receptor | 2704 | 2651 | 2717 | 2803 | 2201 | 2371 | 2140 | 2310 | 794 | 716 | 657 | 739 |
| K04751 nitrogen regulatory protein P-II 1 | 654 | 688 | 678 | 665 | 976 | 906 | 933 | 907 | 980 | 1073 | 1034 | 1059 |
| K04771 serine protease Do [EC:3.4.21.107] | 4021 | 3942 | 3976 | 4213 | 2224 | 2042 | 2155 | 2320 | 2934 | 3065 | 2982 | 3131 |
| K05338 holin-like protein | 37 | 40 | 29 | 20 | 3 | 2 | 0 | 3 | 18 | 25 | 29 | 23 |
| K05339 holin-like protein LrgB | 32 | 33 | 55 | 37 | 0 | 9 | 5 | 5 | 30 | 57 | 25 | 58 |
| K05874 methyl-accepting chemotaxis protein I, serine sensor receptor | 1054 | 1089 | 915 | 993 | 1439 | 1589 | 1398 | 1397 | 79 | 92 | 92 | 85 |
| K05875 methyl-accepting chemotaxis protein II, aspartate sensor receptor | 1214 | 1234 | 1185 | 1240 | 1825 | 1826 | 1731 | 1536 | 88 | 72 | 77 | 74 |
| K05876 methyl-accepting chemotaxis protein III, ribose and galactose sensor receptor | 286 | 312 | 294 | 336 | 419 | 454 | 434 | 447 | 11 | 15 | 10 | 12 |
| K05877 methyl-accepting chemotaxis protein IV, peptide sensor receptor | 263 | 301 | 373 | 308 | 427 | 471 | 371 | 332 | 11 | 12 | 18 | 17 |
| K05964 holo-ACP synthase [EC:2.7.7.61] | 506 | 484 | 479 | 513 | 375 | 439 | 426 | 440 | 537 | 571 | 557 | 571 |
| K05966 triphosphoribosyl-dephospho-CoA synthase [EC:2.4.2.52] | 1206 | 1138 | 1056 | 1147 | 753 | 785 | 756 | 706 | 1436 | 1644 | 1596 | 1640 |
| K06080 RcsF protein | 236 | 246 | 209 | 219 | 163 | 208 | 132 | 160 | 299 | 369 | 308 | 364 |
| K06596 chemosensory pili system protein ChpA (sensor histidine kinase/response regulator) | 10787 | 10711 | 9807 | 10448 | 6017 | 5761 | 5568 | 6414 | 3468 | 3436 | 3240 | 3338 |
| K06597 chemosensory pili system protein ChpB (putative protein-glutamate methylesterase) | 1682 | 1589 | 1510 | 1602 | 838 | 887 | 864 | 956 | 596 | 519 | 447 | 488 |
| K06598 chemosensory pili system protein ChpC | 789 | 724 | 709 | 679 | 402 | 381 | 379 | 452 | 247 | 296 | 248 | 224 |
| K07165 transmembrane sensor | 11201 | 11026 | 10136 | 10621 | 5792 | 5595 | 5582 | 6171 | 3509 | 3182 | 3131 | 3203 |
| K07636 two-component system, OmpR family, phosphate regulon sensor histidine kinase PhoR [EC:2.7.13.3] | 5436 | 5371 | 5230 | 5269 | 6181 | 6321 | 6253 | 6123 | 5350 | 5438 | 5254 | 5301 |
| K07637 two-component system, OmpR family, sensor histidine kinase PhoQ [EC:2.7.13.3] | 2514 | 2553 | 2418 | 2480 | 1419 | 1538 | 1360 | 1547 | 1580 | 1721 | 1630 | 1704 |
| K07638 two-component system, OmpR family, osmolarity sensor histidine kinase EnvZ [EC:2.7.13.3] | 2667 | 2564 | 2445 | 2536 | 1717 | 1698 | 1529 | 1637 | 1828 | 1792 | 1846 | 1845 |
| K07639 two-component system, OmpR family, sensor histidine kinase RstB [EC:2.7.13.3] | 3005 | 2905 | 2744 | 3013 | 1633 | 1711 | 1578 | 1754 | 1631 | 1778 | 1606 | 1757 |
| K07640 two-component system, OmpR family, sensor histidine kinase CpxA [EC:2.7.13.3] | 902 | 901 | 888 | 945 | 663 | 674 | 693 | 671 | 1254 | 1220 | 1253 | 1305 |
| K07641 two-component system, OmpR family, sensor histidine kinase CreC [EC:2.7.13.3] | 1888 | 1911 | 1951 | 1895 | 1237 | 1190 | 1213 | 1188 | 1460 | 1498 | 1346 | 1578 |
| K07642 two-component system, OmpR family, sensor histidine kinase BaeS [EC:2.7.13.3] | 835 | 758 | 756 | 792 | 533 | 573 | 572 | 575 | 1122 | 1184 | 1100 | 1188 |
| K07643 two-component system, OmpR family, sensor histidine kinase BasS [EC:2.7.13.3] | 605 | 604 | 605 | 703 | 436 | 417 | 459 | 427 | 838 | 953 | 881 | 896 |
| K07644 two-component system, OmpR family, heavy metal sensor histidine kinase CusS [EC:2.7.13.3] | 12082 | 11941 | 11475 | 11882 | 7896 | 8077 | 7446 | 7698 | 4518 | 4263 | 4286 | 4445 |
| K07645 two-component system, OmpR family, sensor histidine kinase QseC [EC:2.7.13.3] | 2904 | 2923 | 2697 | 2902 | 1497 | 1433 | 1386 | 1504 | 1753 | 1837 | 1796 | 1958 |
| K07646 two-component system, OmpR family, sensor histidine kinase KdpD [EC:2.7.13.3] | 7688 | 7460 | 7132 | 7380 | 5527 | 5448 | 5583 | 5753 | 7249 | 7325 | 7202 | 7356 |
| K07647 two-component system, OmpR family, sensor histidine kinase TorS [EC:2.7.13.3] | 7 | 11 | 10 | 14 | 20 | 19 | 29 | 25 | 130 | 194 | 179 | 166 |
| K07648 two-component system, OmpR family, aerobic respiration control sensor histidine kinase ArcB [EC:2.7.13.3] | 1322 | 1290 | 1266 | 1369 | 971 | 1050 | 999 | 1015 | 1749 | 1674 | 1689 | 1710 |
| K07649 two-component system, OmpR family, sensor histidine kinase TctE [EC:2.7.13.3] | 2258 | 2167 | 2102 | 2130 | 1163 | 1132 | 1177 | 1208 | 704 | 724 | 609 | 756 |
| K07650 two-component system, OmpR family, sensor histidine kinase CssS [EC:2.7.13.3] | 0 | 0 | 2 | 1 | 2 | 2 | 0 | 0 | 4 | 3 | 7 | 7 |
| K07651 two-component system, OmpR family, sensor histidine kinase ResE [EC:2.7.13.3] | 6 | 9 | 12 | 11 | 2 | 3 | 3 | 1 | 12 | 20 | 22 | 25 |
| K07652 two-component system, OmpR family, sensor histidine kinase VicK [EC:2.7.13.3] | 185 | 167 | 173 | 175 | 140 | 136 | 140 | 159 | 242 | 259 | 228 | 243 |
| K07653 two-component system, OmpR family, sensor histidine kinase MprB [EC:2.7.13.3] | 0 | 3 | 4 | 2 | 0 | 0 | 2 | 0 | 1 | 6 | 2 | 2 |
| K07654 two-component system, OmpR family, sensor histidine kinase MtrB [EC:2.7.13.3] | 8 | 4 | 12 | 8 | 3 | 3 | 1 | 3 | 17 | 17 | 14 | 24 |
| K07656 two-component system, OmpR family, sensor histidine kinase TrcS [EC:2.7.13.3] | 0 | 0 | 0 | 0 | 0 | 0 | 0 | 0 | 1 | 0 | 0 | 0 |
| K07657 two-component system, OmpR family, phosphate regulon response regulator PhoB | 1050 | 1044 | 1057 | 1050 | 749 | 682 | 685 | 679 | 881 | 839 | 759 | 866 |
| K07658 two-component system, OmpR family, alkaline phosphatase synthesis response regulator PhoP | 525 | 429 | 484 | 514 | 107 | 192 | 157 | 150 | 514 | 673 | 648 | 616 |
| K07659 two-component system, OmpR family, phosphate regulon response regulator OmpR | 1247 | 1320 | 1190 | 1257 | 863 | 874 | 850 | 773 | 1042 | 1025 | 1054 | 1046 |
| K07660 two-component system, OmpR family, response regulator PhoP | 1474 | 1545 | 1396 | 1494 | 796 | 825 | 696 | 771 | 1257 | 1375 | 1361 | 1422 |
| K07661 two-component system, OmpR family, response regulator RstA | 1445 | 1473 | 1382 | 1481 | 858 | 836 | 803 | 831 | 881 | 869 | 811 | 845 |
| K07662 two-component system, OmpR family, response regulator CpxR | 449 | 411 | 444 | 421 | 316 | 340 | 369 | 330 | 665 | 678 | 653 | 617 |
| K07663 two-component system, OmpR family, catabolic regulation response regulator CreB | 1541 | 1494 | 1464 | 1485 | 916 | 910 | 918 | 849 | 930 | 864 | 891 | 913 |
| K07664 two-component system, OmpR family, response regulator BaeR | 407 | 415 | 365 | 404 | 262 | 255 | 241 | 258 | 584 | 601 | 561 | 552 |
| K07665 two-component system, OmpR family, copper resistance phosphate regulon response regulator CusR | 5456 | 5225 | 5081 | 5339 | 3454 | 3499 | 3212 | 3447 | 1971 | 1897 | 1874 | 1842 |
| K07666 two-component system, OmpR family, response regulator QseB | 1077 | 1127 | 1131 | 1095 | 667 | 696 | 725 | 683 | 913 | 911 | 931 | 918 |
| K07667 two-component system, OmpR family, KDP operon response regulator KdpE | 2347 | 2055 | 2213 | 2222 | 899 | 922 | 879 | 930 | 2092 | 2061 | 2058 | 2121 |
| K07668 two-component system, OmpR family, response regulator VicR | 101 | 129 | 136 | 124 | 60 | 74 | 76 | 78 | 140 | 148 | 148 | 157 |
| K07669 two-component system, OmpR family, response regulator MprA | 13 | 8 | 10 | 13 | 9 | 21 | 7 | 16 | 12 | 12 | 19 | 11 |
| K07670 two-component system, OmpR family, response regulator MtrA | 9 | 9 | 11 | 2 | 5 | 5 | 1 | 7 | 12 | 10 | 10 | 17 |
| K07671 two-component system, OmpR family, response regulator PrrA | 0 | 0 | 0 | 2 | 0 | 1 | 2 | 0 | 1 | 0 | 2 | 3 |
| K07672 two-component system, OmpR family, response regulator TrcR | 0 | 0 | 0 | 3 | 0 | 0 | 0 | 0 | 3 | 3 | 4 | 1 |
| K07673 two-component system, NarL family, nitrate/nitrite sensor histidine kinase NarX [EC:2.7.13.3] | 2305 | 2392 | 2075 | 2283 | 1526 | 1579 | 1404 | 1505 | 1972 | 1898 | 1819 | 1876 |
| K07674 two-component system, NarL family, nitrate/nitrite sensor histidine kinase NarQ [EC:2.7.13.3] | 344 | 346 | 371 | 385 | 557 | 592 | 548 | 473 | 125 | 104 | 95 | 109 |
| K07675 two-component system, NarL family, sensor histidine kinase UhpB [EC:2.7.13.3] | 592 | 621 | 554 | 625 | 209 | 172 | 174 | 163 | 1355 | 1285 | 1279 | 1314 |
| K07676 two-component system, NarL family, sensor histidine kinase RcsD [EC:2.7.13.3] | 1423 | 1589 | 1433 | 1521 | 1018 | 1012 | 1031 | 1050 | 1930 | 2055 | 1973 | 2023 |
| K07677 two-component system, NarL family, capsular synthesis sensor histidine kinase RcsC [EC:2.7.13.3] | 1648 | 1697 | 1558 | 1743 | 1083 | 1242 | 1206 | 1052 | 2299 | 2465 | 2226 | 2482 |
| K07678 two-component system, NarL family, sensor histidine kinase BarA [EC:2.7.13.3] | 5252 | 5278 | 4976 | 5261 | 3053 | 3115 | 3206 | 3375 | 3477 | 3402 | 3428 | 3438 |
| K07679 two-component system, NarL family, sensor histidine kinase EvgS [EC:2.7.13.3] | 10903 | 10843 | 10873 | 10764 | 5697 | 6016 | 5516 | 5433 | 5573 | 5689 | 5671 | 5932 |
| K07680 two-component system, NarL family, sensor histidine kinase ComP [EC:2.7.13.3] | 2 | 9 | 2 | 1 | 0 | 0 | 0 | 0 | 0 | 0 | 0 | 0 |
| K07681 two-component system, NarL family, vancomycin resistance sensor histidine kinase VraS [EC:2.7.13.3] | 0 | 0 | 0 | 0 | 0 | 0 | 0 | 1 | 0 | 0 | 0 | 0 |
| K07683 two-component system, NarL family, sensor histidine kinase NreB [EC:2.7.13.3] | 0 | 0 | 0 | 0 | 0 | 1 | 2 | 1 | 0 | 3 | 3 | 0 |
| K07684 two-component system, NarL family, nitrate/nitrite response regulator NarL | 1199 | 1132 | 1206 | 1197 | 736 | 774 | 790 | 916 | 828 | 809 | 764 | 753 |
| K07685 two-component system, NarL family, nitrate/nitrite response regulator NarP | 2 | 3 | 5 | 1 | 10 | 9 | 6 | 6 | 42 | 72 | 57 | 36 |
| K07686 two-component system, NarL family, uhpT operon response regulator UhpA | 867 | 871 | 764 | 872 | 438 | 370 | 411 | 409 | 568 | 653 | 645 | 631 |
| K07687 two-component system, NarL family, captular synthesis response regulator RcsB | 2525 | 2576 | 2617 | 2549 | 1445 | 1567 | 1465 | 1414 | 1278 | 1246 | 1185 | 1147 |
| K07688 two-component system, NarL family, response regulator, fimbrial Z protein, FimZ | 124 | 133 | 128 | 145 | 195 | 196 | 224 | 204 | 0 | 1 | 0 | 1 |
| K07689 two-component system, NarL family, invasion response regulator UvrY | 465 | 476 | 443 | 500 | 317 | 374 | 297 | 333 | 522 | 621 | 534 | 651 |
| K07690 two-component system, NarL family, response regulator EvgA | 2519 | 2453 | 2279 | 2346 | 1342 | 1424 | 1295 | 1313 | 1175 | 1105 | 1067 | 1108 |
| K07691 two-component system, NarL family, competent response regulator ComA | 1 | 0 | 0 | 3 | 0 | 0 | 0 | 0 | 0 | 0 | 0 | 0 |
| K07692 two-component system, NarL family, response regulator DegU | 1 | 0 | 0 | 0 | 0 | 0 | 0 | 0 | 0 | 0 | 0 | 0 |
| K07693 two-component system, NarL family, response regulator DesR | 6 | 2 | 9 | 4 | 0 | 2 | 2 | 2 | 11 | 7 | 8 | 4 |
| K07694 two-component system, NarL family, vancomycin resistance associated response regulator VraR | 2 | 2 | 0 | 5 | 2 | 3 | 1 | 3 | 0 | 0 | 0 | 0 |
| K07696 two-component system, NarL family, response regulator NreC | 3 | 6 | 2 | 6 | 3 | 5 | 3 | 7 | 10 | 4 | 2 | 6 |
| K07698 two-component system, sporulation sensor kinase C [EC:2.7.13.3] | 0 | 1 | 0 | 1 | 0 | 0 | 0 | 0 | 5 | 0 | 1 | 1 |
| K07699 two-component system, response regulator, stage 0 sporulation protein A | 490 | 538 | 539 | 552 | 184 | 194 | 187 | 211 | 444 | 541 | 430 | 458 |
| K07700 two-component system, CitB family, cit operon sensor histidine kinase CitA [EC:2.7.13.3] | 1632 | 1609 | 1461 | 1707 | 854 | 936 | 922 | 829 | 2755 | 3000 | 2601 | 2803 |
| K07701 two-component system, CitB family, sensor histidine kinase DcuS [EC:2.7.13.3] | 907 | 960 | 859 | 913 | 628 | 690 | 628 | 630 | 1296 | 1417 | 1205 | 1347 |
| K07702 two-component system, CitB family, response regulator CitB | 706 | 747 | 752 | 689 | 341 | 417 | 411 | 403 | 1129 | 1232 | 1214 | 1217 |
| K07703 two-component system, CitB family, response regulator DcuR | 392 | 364 | 384 | 393 | 232 | 273 | 268 | 312 | 518 | 571 | 520 | 587 |
| K07704 two-component system, LytTR family, sensor histidine kinase LytS [EC:2.7.13.3] | 279 | 287 | 287 | 276 | 86 | 127 | 127 | 95 | 334 | 409 | 359 | 379 |
| K07705 two-component system, LytTR family, response regulator LytT | 96 | 129 | 113 | 104 | 40 | 48 | 47 | 43 | 94 | 115 | 87 | 99 |
| K07706 two-component system, LytTR family, sensor histidine kinase AgrC [EC:2.7.13.3] | 383 | 381 | 359 | 419 | 92 | 98 | 64 | 79 | 556 | 549 | 563 | 536 |
| K07707 two-component system, LytTR family, response regulator AgrA | 145 | 159 | 176 | 162 | 61 | 57 | 70 | 44 | 200 | 243 | 223 | 218 |
| K07708 two-component system, NtrC family, nitrogen regulation sensor histidine kinase GlnL [EC:2.7.13.3] | 2067 | 1994 | 1938 | 2199 | 1347 | 1341 | 1230 | 1308 | 1372 | 1419 | 1375 | 1527 |
| K07709 two-component system, NtrC family, sensor histidine kinase HydH [EC:2.7.13.3] | 859 | 903 | 848 | 904 | 594 | 723 | 623 | 639 | 1271 | 1179 | 1151 | 1246 |
| K07710 two-component system, NtrC family, sensor histidine kinase AtoS [EC:2.7.13.3] | 5 | 5 | 12 | 4 | 19 | 27 | 45 | 16 | 107 | 119 | 102 | 116 |
| K07711 two-component system, NtrC family, sensor histidine kinase GlrK [EC:2.7.13.3] | 846 | 793 | 754 | 867 | 507 | 600 | 485 | 496 | 1214 | 1223 | 1115 | 1248 |
| K07712 two-component system, NtrC family, nitrogen regulation response regulator GlnG | 3562 | 3564 | 3384 | 3535 | 2330 | 2121 | 2127 | 2194 | 2080 | 2148 | 2170 | 2196 |
| K07713 two-component system, NtrC family, response regulator HydG | 3724 | 3810 | 4020 | 3871 | 9125 | 9085 | 9121 | 9160 | 7183 | 6957 | 7422 | 6929 |
| K07714 two-component system, NtrC family, response regulator AtoC | 37 | 24 | 47 | 32 | 42 | 28 | 56 | 32 | 103 | 103 | 98 | 92 |
| K07715 two-component system, NtrC family, response regulator GlrR | 798 | 753 | 745 | 771 | 538 | 629 | 619 | 570 | 1135 | 1146 | 1099 | 1201 |
| K07716 two-component system, cell cycle sensor histidine kinase PleC [EC:2.7.13.3] | 4 | 2 | 3 | 3 | 3 | 6 | 6 | 6 | 6 | 9 | 4 | 7 |
| K07717 two-component system, sensor histidine kinase YcbA [EC:2.7.13.3] | 15 | 12 | 12 | 25 | 5 | 8 | 13 | 7 | 10 | 13 | 3 | 7 |
| K07718 two-component system, sensor histidine kinase YesM [EC:2.7.13.3] | 14983 | 14467 | 15217 | 15053 | 2519 | 2695 | 2646 | 2313 | 16237 | 17870 | 17006 | 17327 |
| K07719 two-component system, response regulator YcbB | 48 | 34 | 34 | 36 | 14 | 13 | 12 | 11 | 32 | 27 | 28 | 14 |
| K07720 two-component system, response regulator YesN | 14237 | 13420 | 14020 | 13981 | 2689 | 2729 | 2700 | 2582 | 14228 | 15383 | 15143 | 15436 |
| K07768 two-component system, OmpR family, sensor histidine kinase SenX3 [EC:2.7.13.3] | 4 | 5 | 5 | 1 | 7 | 6 | 5 | 10 | 11 | 8 | 10 | 15 |
| K07770 two-component system, OmpR family, response regulator CssR | 1 | 1 | 0 | 0 | 3 | 3 | 0 | 0 | 2 | 5 | 2 | 2 |
| K07771 two-component system, OmpR family, response regulator BasR | 351 | 372 | 375 | 411 | 268 | 309 | 289 | 242 | 578 | 510 | 565 | 565 |
| K07772 two-component system, OmpR family, torCAD operon response regulator TorR | 1 | 1 | 2 | 1 | 8 | 7 | 8 | 1 | 28 | 25 | 33 | 33 |
| K07773 two-component system, OmpR family, aerobic respiration control protein ArcA | 340 | 323 | 368 | 332 | 250 | 252 | 248 | 242 | 387 | 454 | 386 | 444 |
| K07774 two-component system, OmpR family, response regulator TctD | 1185 | 1033 | 1016 | 1020 | 589 | 520 | 554 | 613 | 323 | 297 | 325 | 330 |
| K07775 two-component system, OmpR family, response regulator ResD | 11 | 8 | 10 | 7 | 8 | 5 | 2 | 9 | 13 | 26 | 19 | 17 |
| K07776 two-component system, OmpR family, response regulator RegX3 | 6 | 9 | 16 | 8 | 14 | 12 | 8 | 16 | 18 | 31 | 25 | 18 |
| K07777 two-component system, NarL family, sensor histidine kinase DegS [EC:2.7.13.3] | 0 | 0 | 2 | 1 | 0 | 0 | 0 | 0 | 0 | 8 | 7 | 3 |
| K07778 two-component system, NarL family, sensor histidine kinase DesK [EC:2.7.13.3] | 0 | 2 | 0 | 3 | 0 | 2 | 0 | 0 | 0 | 0 | 0 | 1 |
| K07781 LuxR family transcriptional regulator, capsular biosynthesis positive transcription factor | 307 | 303 | 306 | 299 | 247 | 241 | 225 | 214 | 347 | 356 | 378 | 428 |
| K07782 LuxR family transcriptional regulator, quorum-sensing system regulator SdiA | 424 | 382 | 382 | 438 | 284 | 300 | 323 | 273 | 582 | 640 | 640 | 596 |
| K07783 MFS transporter, OPA family, sugar phosphate sensor protein UhpC | 2754 | 3130 | 2828 | 2978 | 3624 | 3411 | 3594 | 3499 | 4205 | 4173 | 4142 | 4237 |
| K07784 MFS transporter, OPA family, hexose phosphate transport protein UhpT | 598 | 650 | 562 | 675 | 150 | 239 | 206 | 188 | 1277 | 1309 | 1252 | 1292 |
| K07786 MFS transporter, DHA2 family, multidrug resistance protein | 3 | 2 | 7 | 7 | 15 | 11 | 17 | 25 | 75 | 87 | 84 | 87 |
| K07787 Cu(I)/Ag(I) efflux system membrane protein CusA/SilA | 8227 | 7974 | 8049 | 7961 | 6558 | 6830 | 6459 | 6582 | 4323 | 4324 | 4314 | 4326 |
| K07788 multidrug efflux pump | 5708 | 5639 | 5393 | 5541 | 3189 | 3289 | 3174 | 3379 | 3614 | 3888 | 3578 | 3835 |
| K07789 multidrug efflux pump | 6155 | 5998 | 5731 | 5962 | 3426 | 3508 | 3372 | 3553 | 3977 | 3756 | 3690 | 3863 |
| K07790 putative membrane protein PagO | 0 | 10 | 14 | 0 | 7 | 0 | 0 | 0 | 0 | 0 | 8 | 0 |
| K07792 anaerobic C4-dicarboxylate transporter DcuB | 1017 | 974 | 987 | 1079 | 694 | 812 | 757 | 799 | 1486 | 1534 | 1493 | 1580 |
| K07793 putative tricarboxylic transport membrane protein | 2971 | 2776 | 2893 | 2822 | 1644 | 1611 | 1339 | 1560 | 1160 | 1117 | 1097 | 1237 |
| K07794 putative tricarboxylic transport membrane protein | 593 | 511 | 638 | 583 | 310 | 307 | 295 | 308 | 210 | 184 | 173 | 192 |

| | | | | | | | | | | | | |
|---|---|---|---|---|---|---|---|---|---|---|---|---|
| K07795 putative tricarboxylic transport membrane protein | 1498 | 1504 | 1424 | 1436 | 824 | 762 | 802 | 808 | 436 | 415 | 431 | 463 |
| K07796 outer membrane protein, Cu(I)/Ag(I) efflux system | 1435 | 1442 | 1375 | 1519 | 1455 | 1506 | 1411 | 1363 | 1331 | 1448 | 1352 | 1512 |
| K07797 multidrug resistance protein K | 7 | 1 | 0 | 7 | 10 | 14 | 10 | 10 | 68 | 57 | 69 | 58 |
| K07798 membrane fusion protein, Cu(I)/Ag(I) efflux system | 3645 | 3550 | 3730 | 3697 | 2795 | 2840 | 2895 | 2727 | 1638 | 1839 | 1722 | 1895 |
| K07799 membrane fusion protein, multidrug efflux system | 2191 | 2156 | 1998 | 2157 | 1314 | 1365 | 1339 | 1338 | 1409 | 1333 | 1279 | 1409 |
| K07800 AgrD protein | 7 | 6 | 12 | 9 | 4 | 3 | 3 | 2 | 4 | 9 | 10 | 5 |
| K07803 zinc resistance-associated protein | 263 | 314 | 301 | 257 | 185 | 225 | 202 | 181 | 434 | 404 | 374 | 421 |
| K07810 Cu(I)/Ag(I) efflux system periplasmic protein CusF | 301 | 326 | 317 | 339 | 336 | 332 | 283 | 295 | 300 | 365 | 314 | 334 |
| K07813 accessory gene regulator B | 200 | 213 | 218 | 209 | 98 | 90 | 97 | 76 | 166 | 192 | 177 | 214 |
| K08082 two-component system, LytTR family, sensor histidine kinase AlgZ [EC:2.7.13.3] | 1786 | 1765 | 1642 | 1757 | 962 | 919 | 925 | 1035 | 582 | 559 | 574 | 565 |
| K08083 two-component system, LytTR family, response regulator AlgR | 1333 | 1430 | 1347 | 1260 | 699 | 665 | 668 | 793 | 400 | 425 | 389 | 417 |
| K08348 formate dehydrogenase-N, alpha subunit [EC:1.17.5.3] | 2136 | 2146 | 2030 | 2134 | 1839 | 1922 | 1813 | 1865 | 2255 | 2343 | 2294 | 2364 |
| K08349 formate dehydrogenase-N, beta subunit | 674 | 709 | 667 | 712 | 719 | 685 | 575 | 649 | 746 | 747 | 807 | 788 |
| K08350 formate dehydrogenase-N, gamma subunit | 436 | 435 | 451 | 473 | 363 | 415 | 423 | 420 | 447 | 494 | 472 | 534 |
| K08372 putative serine protease PepD [EC:3.4.21.-] | 12 | 7 | 13 | 13 | 5 | 13 | 5 | 16 | 16 | 24 | 26 | 29 |
| K08475 two-component system, NtrC family, phosphoglycerate transport system sensor histidine kinase PgtB [EC:2.7.13.3] | 698 | 667 | 692 | 736 | 168 | 170 | 177 | 155 | 1364 | 1409 | 1362 | 1498 |
| K08476 two-component system, NtrC family, phosphoglycerate transport system response regulator PgtA | 479 | 447 | 433 | 437 | 107 | 129 | 129 | 133 | 975 | 900 | 912 | 1006 |
| K08478 phosphoglycerate transport regulatory protein PgtC | 381 | 466 | 431 | 430 | 132 | 108 | 146 | 112 | 883 | 874 | 992 | 951 |
| K08641 zinc D-Ala-D-Ala dipeptidase [EC:3.4.13.22] | 1905 | 1835 | 1716 | 1711 | 2501 | 2550 | 2426 | 2551 | 1714 | 1596 | 1754 | 1630 |
| K08738 cytochrome c | 536 | 445 | 470 | 493 | 285 | 267 | 263 | 266 | 206 | 158 | 146 | 142 |
| K09475 outer membrane pore protein C | 629 | 577 | 622 | 634 | 429 | 474 | 458 | 385 | 856 | 938 | 853 | 866 |
| K09476 outer membrane pore protein F | 794 | 832 | 785 | 825 | 668 | 798 | 790 | 655 | 871 | 921 | 848 | 861 |
| K09477 citrate:succinate antiporter | 16 | 21 | 18 | 18 | 5 | 16 | 10 | 23 | 64 | 73 | 69 | 82 |
| K10125 two-component system, NtrC family, C4-dicarboxylate transport sensor histidine kinase DctB [EC:2.7.13.3] | 2971 | 2775 | 2769 | 2925 | 1530 | 1610 | 1598 | 1635 | 1117 | 1038 | 973 | 1122 |
| K10126 two-component system, NtrC family, C4-dicarboxylate transport response regulator DctD | 4408 | 4410 | 4166 | 4315 | 2438 | 2340 | 2291 | 2415 | 1509 | 1399 | 1404 | 1417 |
| K10697 two-component system, OmpR family, response regulator RpaA | 3 | 2 | 1 | 1 | 0 | 0 | 0 | 0 | 2 | 2 | 2 | 4 |
| K10715 two-component system, sensor histidine kinase RpfC [EC:2.7.13.3] | 2 | 0 | 1 | 1 | 0 | 1 | 1 | 0 | 1 | 0 | 0 | 1 |
| K10851 nitrogen regulatory protein A | 0 | 0 | 0 | 0 | 0 | 1 | 0 | 2 | 0 | 0 | 0 | 0 |
| K10909 two-component system, autoinducer 2 sensor kinase/phosphatase LuxQ [EC:2.7.13.3 3.1.3.-] | 0 | 0 | 0 | 0 | 3 | 2 | 3 | 2 | 1 | 0 | 3 | 1 |
| K10914 CRP/FNR family transcriptional regulator, cyclic AMP receptor protein | 2190 | 2285 | 2142 | 2133 | 1190 | 1085 | 1039 | 1169 | 1181 | 1318 | 1193 | 1221 |
| K10941 sigma-54 dependent transcriptional regulator, flagellar regulatory protein | 1956 | 1955 | 1942 | 1910 | 1072 | 1122 | 987 | 1125 | 619 | 613 | 600 | 573 |
| K10942 two-component system, sensor histidine kinase FlrB [EC:2.7.13.3] | 1791 | 1805 | 1776 | 1728 | 924 | 900 | 741 | 849 | 518 | 512 | 532 | 533 |
| K10943 two-component system, response regulator FlrC | 2189 | 2121 | 2109 | 2065 | 1103 | 1097 | 1031 | 1139 | 757 | 636 | 676 | 663 |
| K11103 aerobic C4-dicarboxylate transport protein | 3967 | 4015 | 4054 | 3946 | 2380 | 2399 | 2250 | 2587 | 2224 | 2270 | 2113 | 2229 |
| K11230 mitogen-activated protein kinase kinase kinase [EC:2.7.11.25] | 0 | 0 | 0 | 0 | 0 | 0 | 0 | 0 | 0 | 0 | 0 | 1 |
| K11231 osomolarity two-component system, sensor histidine kinase SLN1 [EC:2.7.13.3] | 1 | 0 | 0 | 0 | 1 | 0 | 0 | 0 | 0 | 0 | 0 | 0 |
| K11329 two-component system, OmpR family, response regulator RpaB | 1 | 2 | 0 | 0 | 0 | 2 | 1 | 0 | 4 | 2 | 0 | 1 |
| K11354 two-component system, chemotaxis family, sensor kinase Cph1 [EC:2.7.13.3] | 0 | 0 | 0 | 1 | 0 | 0 | 0 | 0 | 3 | 0 | 0 | 0 |
| K11356 two-component system, sensor histidine kinase and response regulator [EC:2.7.13.3] | 1 | 1 | 2 | 0 | 1 | 0 | 1 | 1 | 1 | 1 | 1 | 1 |
| K11357 two-component system, cell cycle sensor histidine kinase DivJ [EC:2.7.13.3] | 2 | 3 | 3 | 3 | 2 | 2 | 4 | 1 | 7 | 3 | 5 | 3 |
| K11382 MFS transporter, OPA family, phosphoglycerate transporter protein | 417 | 459 | 491 | 443 | 84 | 142 | 115 | 108 | 1030 | 1040 | 957 | 974 |
| K11383 two-component system, NtrC family, sensor histidine kinase KinB [EC:2.7.13.3] | 2018 | 1866 | 1847 | 1951 | 1121 | 1099 | 1035 | 1035 | 601 | 570 | 631 | 649 |
| K11384 two-component system, NtrC family, response regulator AlgB | 2262 | 2308 | 2330 | 2359 | 1280 | 1221 | 1265 | 1418 | 696 | 749 | 656 | 731 |
| K11443 two-component system, cell cycle response regulator DivK | 0 | 2 | 0 | 0 | 0 | 0 | 0 | 0 | 0 | 0 | 0 | 0 |
| K11444 two-component system, chemotaxis family, response regulator WspR [EC:2.7.7.65] | 121 | 119 | 118 | 109 | 72 | 81 | 194 | 235 | 38 | 44 | 44 | 42 |
| K11520 two-component system, OmpR family, manganese sensing sensor histidine kinase [EC:2.7.13.3] | 0 | 1 | 0 | 1 | 0 | 0 | 1 | 0 | 0 | 0 | 0 | 1 |
| K11521 two-component system, OmpR family, manganese sensing response regulator | 0 | 1 | 2 | 3 | 1 | 1 | 1 | 0 | 4 | 0 | 3 | 1 |
| K11525 methyl-accepting chemotaxis protein PixJ | 0 | 1 | 1 | 0 | 0 | 0 | 0 | 0 | 0 | 0 | 0 | 0 |
| K11614 two-component system, CitB family, sensor histidine kinase MalK [EC:2.7.13.3] | 63 | 58 | 75 | 58 | 22 | 20 | 27 | 22 | 111 | 151 | 132 | 125 |
| K11615 two-component system, CitB family, response regulator MalR | 54 | 50 | 47 | 62 | 14 | 20 | 10 | 21 | 75 | 86 | 102 | 59 |
| K11616 malate:Na+ symporter | 68 | 55 | 69 | 68 | 63 | 95 | 85 | 86 | 100 | 97 | 85 | 102 |
| K11617 two-component system, NarL family, sensor histidine kinase LiaS [EC:2.7.13.3] | 93 | 66 | 86 | 73 | 24 | 16 | 13 | 24 | 66 | 96 | 80 | 100 |
| K11618 two-component system, NarL family, response regulator LiaR | 44 | 37 | 43 | 39 | 8 | 8 | 9 | 17 | 30 | 55 | 41 | 60 |
| K11624 two-component system, NarL family, response regulator YdfI | 6 | 5 | 3 | 3 | 3 | 0 | 0 | 0 | 6 | 10 | 9 | 13 |
| K11626 putative sodium/glutamine symporter | 1 | 1 | 2 | 1 | 4 | 5 | 2 | 3 | 3 | 3 | 1 | 2 |
| K11629 two-component system, OmpR family, bacitracin resistance sensor histidine kinase BceS [EC:2.7.13.3] | 2 | 2 | 0 | 1 | 0 | 0 | 0 | 0 | 0 | 0 | 0 | 0 |
| K11630 two-component system, OmpR family, bacitracin resistance response regulator BceR | 3 | 0 | 4 | 1 | 2 | 3 | 0 | 2 | 2 | 3 | 1 | 1 |
| K11635 putative ABC transport system ATP-binding protein | 42 | 57 | 41 | 33 | 3 | 21 | 9 | 3 | 35 | 50 | 36 | 49 |
| K11636 putative ABC transport system permease protein | 91 | 94 | 79 | 110 | 8 | 11 | 11 | 25 | 104 | 99 | 64 | 83 |
| K11637 two-component system, CitB family, sensor histidine kinase CitS [EC:2.7.13.3] | 0 | 2 | 0 | 3 | 0 | 1 | 1 | 0 | 1 | 0 | 1 | 2 |
| K11688 C4-dicarboxylate-binding protein DctP | 5009 | 5004 | 4905 | 4852 | 2796 | 2910 | 2662 | 2851 | 1473 | 1631 | 1505 | 1573 |
| K11689 C4-dicarboxylate transporter, DctQ subunit | 2012 | 1980 | 1926 | 1987 | 1166 | 1106 | 988 | 1096 | 695 | 685 | 733 | 713 |
| K11690 C4-dicarboxylate transporter, DctM subunit | 3764 | 3766 | 3629 | 3689 | 2067 | 1970 | 1887 | 2018 | 1633 | 1759 | 1662 | 1698 |
| K11691 two-component system, CitB family, sensor histidine kinase DctS [EC:2.7.13.3] | 0 | 2 | 2 | 3 | 0 | 0 | 0 | 1 | 0 | 2 | 0 | 1 |
| K11692 two-component system, CitB family, response regulator DctR | 8 | 4 | 10 | 5 | 40 | 33 | 37 | 38 | 13 | 17 | 5 | 12 |
| K11711 two-component system, LuxR family, sensor histidine kinase DctS [EC:2.7.13.3] | 4 | 6 | 2 | 6 | 10 | 5 | 8 | 14 | 10 | 4 | 6 | 10 |
| K11712 two-component system, LuxR family, response regulator DctR | 4 | 3 | 7 | 3 | 6 | 5 | 11 | 2 | 5 | 8 | 7 | 6 |
| K12531 ATP-binding cassette, subfamily B, bacterial RtxE | 6 | 9 | 10 | 7 | 4 | 4 | 7 | 5 | 5 | 3 | 1 | 1 |
| K13040 two-component system, LuxR family, sensor histidine kinase TtrS [EC:2.7.13.3] | 190 | 202 | 202 | 195 | 316 | 374 | 358 | 369 | 353 | 320 | 304 | 278 |
| K13041 two-component system, LuxR family, response regulator TtrR | 12 | 17 | 8 | 12 | 16 | 18 | 21 | 25 | 30 | 20 | 17 | 17 |
| K13486 chemotaxis protein methyltransferase WspC | 1918 | 1782 | 1700 | 1699 | 1024 | 957 | 1083 | 1068 | 588 | 608 | 558 | 563 |
| K13487 methyl-accepting chemotaxis protein WspA | 2047 | 2076 | 2052 | 2078 | 1121 | 1108 | 879 | 948 | 673 | 615 | 609 | 665 |
| K13488 chemotaxis-related protein WspB | 774 | 731 | 755 | 784 | 384 | 377 | 337 | 417 | 264 | 212 | 200 | 230 |
| K13489 chemotaxis-related protein WspD | 152 | 135 | 129 | 152 | 80 | 81 | 64 | 101 | 52 | 44 | 45 | 44 |
| K13490 two-component system, chemotaxis family, sensor histidine kinase and response regulator WspE | 3201 | 3067 | 2965 | 3064 | 1621 | 1576 | 1554 | 1858 | 886 | 848 | 934 | 911 |
| K13491 two-component system, chemotaxis family, response regulator WspF [EC:3.1.1.61] | 1658 | 1646 | 1582 | 1636 | 929 | 860 | 850 | 955 | 553 | 481 | 448 | 514 |
| K13532 two-component system, sporulation sensor kinase D [EC:2.7.13.3] | 0 | 0 | 0 | 0 | 0 | 1 | 0 | 0 | 0 | 0 | 0 | 0 |
| K13533 two-component system, sporulation sensor kinase E [EC:2.7.13.3] | 1 | 0 | 0 | 3 | 0 | 0 | 0 | 0 | 0 | 0 | 0 | 0 |
| K13584 two-component system, cell cycle response regulator CtrA | 0 | 0 | 3 | 1 | 0 | 4 | 0 | 1 | 4 | 2 | 0 | 1 |
| K13587 two-component system, cell cycle sensor histidine kinase and response regulator CckA [EC:2.7.13.3] | 2 | 4 | 0 | 1 | 0 | 0 | 0 | 2 | 6 | 8 | 3 | 3 |
| K13588 histidine phosphotransferase ChpT | 0 | 0 | 0 | 1 | 0 | 0 | 0 | 0 | 2 | 0 | 1 | 1 |
| K13589 two-component system, cell cycle response regulator CpdR | 0 | 0 | 0 | 1 | 0 | 0 | 1 | 0 | 0 | 0 | 0 | 1 |
| K13598 two-component system, NtrC family, nitrogen regulation sensor histidine kinase NtrY [EC:2.7.13.3] | 13 | 12 | 5 | 7 | 5 | 2 | 3 | 4 | 7 | 4 | 3 | 7 |
| K13599 two-component system, NtrC family, nitrogen regulation response regulator NtrX | 25 | 12 | 17 | 10 | 8 | 9 | 13 | 11 | 5 | 10 | 8 | 10 |
| K13815 two-component system, response regulator RpfG | 0 | 0 | 0 | 1 | 0 | 0 | 0 | 0 | 0 | 0 | 0 | 0 |
| K13816 DSF synthase | 175 | 160 | 180 | 169 | 243 | 236 | 221 | 220 | 0 | 2 | 0 | 0 |
| K13924 two-component system, chemotaxis family, CheB/CheR fusion protein [EC:2.1.1.80 3.1.1.61] | 112 | 89 | 97 | 76 | 14 | 6 | 13 | 9 | 84 | 69 | 61 | 63 |
| K13927 holo-ACP synthase / triphosphoribosyl-dephospho-CoA synthase [EC:2.7.7.61 2.4.2.52] | 126 | 112 | 130 | 112 | 27 | 25 | 13 | 7 | 144 | 153 | 136 | 152 |
| K14205 phosphatidylglycerol lysyltransferase [EC:2.3.2.3] | 4315 | 4186 | 4250 | 4077 | 2349 | 2292 | 2040 | 2243 | 1571 | 1581 | 1494 | 1675 |
| K14978 two-component system, LuxR family, secretion system sensor histidine kinase SsrA | 1 | 0 | 0 | 1 | 0 | 0 | 0 | 2 | 0 | 0 | 0 | 0 |
| K14979 two-component system, LuxR family, secretion system response regulator SsrB | 1 | 0 | 0 | 0 | 0 | 1 | 0 | 0 | 0 | 0 | 0 | 0 |
| K14980 two-component system, OmpR family, sensor histidine kinase ChvG [EC:2.7.13.3] | 3 | 5 | 3 | 3 | 1 | 0 | 0 | 3 | 6 | 3 | 1 | 8 |
| K14981 two-component system, OmpR family, response regulator ChvI | 3 | 1 | 1 | 1 | 0 | 0 | 1 | 2 | 1 | 0 | 2 | 0 |
| K14982 two-component system, OmpR family, sensor histidine kinase CiaH [EC:2.7.13.3] | 8 | 14 | 10 | 9 | 18 | 25 | 13 | 16 | 37 | 20 | 24 | 27 |
| K14983 two-component system, OmpR family, response regulator CiaR | 8 | 5 | 7 | 13 | 8 | 4 | 17 | 13 | 16 | 30 | 13 | 36 |
| K14986 two-component system, LuxR family, sensor kinase FixL [EC:2.7.13.3] | 3 | 1 | 1 | 1 | 1 | 0 | 0 | 0 | 3 | 1 | 0 | 3 |
| K14987 two-component system, LuxR family, response regulator FixJ | 13 | 14 | 11 | 13 | 27 | 16 | 17 | 23 | 30 | 23 | 17 | 19 |
| K14988 two-component system, NarL family, secretion system sensor histidine kinase SalK | 18 | 29 | 12 | 17 | 37 | 24 | 20 | 14 | 40 | 36 | 44 | 36 |
| K14989 two-component system, NarL family, secretion system response regulator SalR | 21 | 9 | 11 | 10 | 5 | 8 | 17 | 9 | 26 | 19 | 12 | 46 |
| K15011 two-component system, sensor histidine kinase RegB [EC:2.7.13.3] | 816 | 794 | 777 | 872 | 456 | 412 | 406 | 471 | 262 | 285 | 259 | 297 |
| K15012 two-component system, response regulator RegA | 805 | 777 | 754 | 782 | 435 | 406 | 355 | 398 | 245 | 226 | 209 | 210 |
| K15739 D-alanine---(R)-lactate ligase [EC:6.1.2.1] | 0 | 0 | 0 | 0 | 0 | 0 | 2 | 0 | 0 | 0 | 0 | 1 |
| K15861 CRP/FNR family transcriptional regulator, nitrogen fixation regulation protein | 0 | 2 | 1 | 0 | 0 | 0 | 0 | 0 | 0 | 0 | 0 | 0 |
| K16692 tyrosine-protein kinase Etk/Wzc [EC:2.7.10.-] | 572 | 579 | 646 | 669 | 755 | 816 | 679 | 723 | 537 | 611 | 591 | 613 |
| K17060 two-component system, sensor histidine kinase AauS [EC:2.7.13.3] | 430 | 403 | 392 | 431 | 228 | 235 | 162 | 177 | 146 | 114 | 115 | 103 |
| K17061 two-component system, response regulator AauR | 2081 | 2060 | 1921 | 1933 | 1056 | 950 | 1074 | 1117 | 538 | 548 | 655 | 601 |
| K18072 two-component system, OmpR family, sensor kinase ParS [EC:2.7.13.3] | 23 | 26 | 41 | 35 | 24 | 17 | 45 | 66 | 10 | 9 | 13 | 8 |
| K18073 two-component system, OmpR family, response regulator ParR | 381 | 392 | 437 | 393 | 209 | 261 | 207 | 229 | 114 | 127 | 117 | 115 |
| K18093 imipenem/basic amino acid-specific outer membrane pore [EC:3.4.21.-] | 1774 | 1832 | 1813 | 1803 | 1058 | 1049 | 967 | 1047 | 538 | 589 | 539 | 614 |
| K18094 membrane fusion protein, multidrug efflux system | 1211 | 1192 | 1128 | 1114 | 581 | 571 | 554 | 627 | 409 | 337 | 314 | 359 |
| K18095 multidrug efflux pump | 1789 | 1829 | 1692 | 1765 | 983 | 913 | 922 | 1005 | 537 | 457 | 536 | 574 |
| K18321 membrane fusion protein, multidrug efflux system | 0 | 1 | 0 | 1 | 0 | 1 | 0 | 2 | 0 | 1 | 1 | 0 |
| K18322 multidrug efflux pump | 0 | 0 | 0 | 0 | 0 | 0 | 0 | 0 | 1 | 1 | 0 | 0 |
| K18324 multidrug efflux pump | 1838 | 1756 | 1643 | 1870 | 1256 | 1286 | 1225 | 1171 | 2475 | 2588 | 2549 | 2696 |
| K18326 MFS transporter, DHA2 family, multidrug resistance protein | 792 | 755 | 728 | 831 | 505 | 551 | 538 | 550 | 1088 | 1164 | 1055 | 1140 |
| K18344 two-component system, OmpR family, response regulator VanR | 24 | 23 | 26 | 25 | 6 | 12 | 13 | 16 | 9 | 16 | 18 | 13 |
| K18345 two-component system, OmpR family, sensor histidine kinase VanS [EC:2.7.13.3] | 23 | 18 | 11 | 17 | 16 | 15 | 13 | 10 | 6 | 15 | 12 | 8 |
| K18346 vancomycin resistance protein VanW | 1331 | 1389 | 1321 | 1307 | 834 | 756 | 711 | 786 | 469 | 338 | 370 | 435 |
| K18347 D-specific alpha-keto acid dehydrogenase [EC:1.1.1.-] | 0 | 1 | 0 | 1 | 0 | 1 | 3 | 2 | 4 | 0 | 2 | 1 |
| K18348 serine/alanine racemase [EC:5.1.1.18 5.1.1.1] | 24 | 18 | 18 | 13 | 8 | 17 | 13 | 22 | 18 | 12 | 10 | 23 |
| K18349 two-component system, OmpR family, response regulator VanR | 347 | 413 | 406 | 433 | 137 | 166 | 146 | 153 | 388 | 471 | 403 | 429 |

| | | | | | | | | | | | | |
|---|---|---|---|---|---|---|---|---|---|---|---|---|
| K18350 two-component system, OmpR family, sensor histidine kinase VanS [EC:2.7.13.3] | 214 | 207 | 192 | 207 | 95 | 102 | 76 | 76 | 319 | 339 | 331 | 351 |
| K18351 two-component system, OmpR family, sensor histidine kinase VanS [EC:2.7.13.3] | 0 | 0 | 1 | 0 | 0 | 0 | 0 | 0 | 1 | 2 | 2 | 1 |
| K18856 D-alanine---D-serine ligase [EC:6.3.2.35] | 13 | 12 | 19 | 21 | 25 | 12 | 14 | 15 | 9 | 11 | 15 | 12 |
| K18940 two-component system, OmpR family, sensor histidine kinase ArlS [EC:2.7.13.3] | 0 | 1 | 0 | 0 | 1 | 0 | 1 | 0 | 2 | 2 | 2 | 2 |
| K18941 two-component system, OmpR family, response regulator ArlR | 2 | 0 | 2 | 0 | 0 | 0 | 0 | 0 | 0 | 0 | 0 | 0 |
| K18986 two-component system, OmpR family, sensor kinase Ihk [EC:2.7.13.3] | 65 | 69 | 62 | 55 | 58 | 42 | 48 | 51 | 92 | 114 | 87 | 119 |
| K18987 two-component system, OmpR family, response regulator Irr | 0 | 0 | 1 | 0 | 1 | 2 | 0 | 0 | 1 | 2 | 0 | 0 |
| K19078 two-component system, OmpR family, response regulator protein GraR | 0 | 0 | 0 | 0 | 0 | 0 | 0 | 0 | 0 | 0 | 1 | 1 |
| K19082 two-component system, OmpR family, response regulator protein BraR/BceR | 2 | 0 | 0 | 0 | 0 | 0 | 0 | 0 | 2 | 1 | 0 | 1 |
| K19609 two-component system, OmpR family, sensor histidine kinase PfeS [EC:2.7.13.3] | 4136 | 3792 | 3783 | 3753 | 2095 | 2084 | 2050 | 2403 | 1353 | 1106 | 1172 | 1205 |
| K19610 two-component system, OmpR family, response regulator PfeR | 2198 | 2249 | 2038 | 2127 | 1194 | 1092 | 1100 | 1155 | 746 | 675 | 574 | 675 |
| K19611 ferric enterobactin receptor | 7723 | 7692 | 7242 | 7732 | 3954 | 3974 | 3781 | 3943 | 6619 | 6802 | 6502 | 6609 |
| K19641 two-component system, NtrC family, response regulator HupR/HoxA | 1 | 0 | 0 | 0 | 0 | 0 | 0 | 0 | 2 | 1 | 1 | 2 |
| K19661 two-component system, NtrC family, sensor histidine kinase HupT/HoxJ [EC:2.7.13.3] | 3 | 0 | 0 | 1 | 0 | 0 | 0 | 0 | 0 | 0 | 0 | 0 |
| K19691 osomolarity two-component system, sensor histidine kinase NIK1 [EC:2.7.13.3] | 1 | 0 | 0 | 0 | 1 | 0 | 0 | 1 | 0 | 0 | 0 | 0 |
| K20263 two-component system, NarL family, sensor histidine kinase FusK [EC:2.7.13.3] | 360 | 337 | 296 | 332 | 434 | 508 | 500 | 537 | 0 | 0 | 4 | 0 |
| K20264 two-component system, NarL family, response regulator FusR | 134 | 150 | 163 | 123 | 167 | 232 | 208 | 186 | 1 | 0 | 0 | 1 |
| K20340 two-component system, LytTR family, response regulator AgrA2 | 0 | 0 | 0 | 0 | 0 | 0 | 0 | 0 | 1 | 1 | 0 | 1 |
| K20483 lantibiotic biosynthesis protein | 0 | 2 | 2 | 0 | 0 | 4 | 5 | 0 | 0 | 2 | 3 | 3 |
| K20484 lantibiotic biosynthesis protein | 0 | 2 | 0 | 1 | 2 | 0 | 0 | 0 | 2 | 1 | 0 | 0 |
| K20485 ATP-binding cassette, subfamily B, bacterial NisT | 239 | 231 | 259 | 231 | 30 | 39 | 27 | 37 | 508 | 468 | 494 | 527 |
| K20486 lantibiotic leader peptide-processing serine protease [EC:3.4.21.-] | 93 | 124 | 94 | 120 | 10 | 9 | 5 | 12 | 13 | 3 | 0 | 11 |
| K20487 two-component system, OmpR family, lantibiotic biosynthesis sensor histidine kinase NisK/SpaK [EC:2.7.13.3] | 243 | 271 | 234 | 277 | 85 | 98 | 101 | 88 | 506 | 568 | 526 | 584 |
| K20488 two-component system, OmpR family, lantibiotic biosynthesis response regulator NisR/SpaR | 193 | 213 | 200 | 176 | 62 | 77 | 72 | 63 | 331 | 404 | 342 | 320 |
| K20489 lantibiotic immunity protein | 19 | 16 | 10 | 11 | 0 | 0 | 0 | 0 | 0 | 0 | 0 | 3 |
| K20973 two-component system, sensor histidine kinase SagS [EC:2.7.13.3] | 3348 | 3247 | 3174 | 3282 | 1693 | 1726 | 1752 | 1793 | 1090 | 957 | 981 | 984 |
| K20974 two-component system, sensor histidine kinase [EC:2.7.13.3] | 3187 | 2912 | 2857 | 2969 | 1519 | 1512 | 1391 | 1589 | 1175 | 1037 | 965 | 1127 |
| K20975 two-component system, sensor histidine kinase [EC:2.7.13.3] | 3200 | 3285 | 2805 | 2985 | 1644 | 1505 | 1445 | 1694 | 1068 | 838 | 886 | 885 |
| K20976 histidine phosphotransfer protein HptB | 566 | 586 | 522 | 558 | 267 | 308 | 304 | 300 | 203 | 138 | 158 | 154 |
| K20977 two-component system, HptB-dependent secretion and biofilm response regulator | 2332 | 2218 | 2287 | 2321 | 1349 | 1339 | 1234 | 1281 | 758 | 729 | 735 | 713 |
| K20978 HptB-dependent secretion and biofilm anti anti-sigma factor | 464 | 456 | 431 | 389 | 261 | 249 | 225 | 257 | 159 | 142 | 141 | 135 |
| K23236 two-component-system connector protein SafA | 1 | 0 | 0 | 1 | 4 | 2 | 2 | 1 | 13 | 13 | 9 | 17 |
| K23514 RNA polymerase sigma-19 factor, ECF subfamily | 8732 | 8639 | 8155 | 8303 | 6516 | 6306 | 6009 | 6677 | 4279 | 3827 | 4088 | 4100 |
| ko04014 Ras signaling pathway | 0 | 0 | 0 | 0 | 1 | 2 | 0 | 0 | 0 | 0 | 2 | 2 |
| K02677 classical protein kinase C alpha type [EC:2.7.11.13] | 0 | 0 | 0 | 0 | 0 | 1 | 0 | 0 | 0 | 0 | 0 | 0 |
| K03099 son of sevenless | 0 | 0 | 0 | 0 | 0 | 0 | 0 | 0 | 0 | 0 | 1 | 0 |
| K04393 cell division control protein 42 | 0 | 0 | 0 | 0 | 1 | 0 | 0 | 0 | 0 | 0 | 0 | 0 |
| K07889 Ras-related protein Rab-5C | 0 | 0 | 0 | 0 | 0 | 0 | 0 | 0 | 0 | 0 | 0 | 1 |
| K08052 neurofibromin 1 | 0 | 0 | 0 | 0 | 0 | 1 | 0 | 0 | 0 | 0 | 0 | 1 |
| ko04015 Rap1 signaling pathway | 0 | 0 | 0 | 0 | 1 | 1 | 0 | 0 | 0 | 0 | 0 | 0 |
| ko04010 MAPK signaling pathway | 0 | 0 | 1 | 1 | 1 | 2 | 0 | 0 | 0 | 1 | 1 | 2 |
| K04733 interleukin-1 receptor-associated kinase 4 [EC:2.7.11.1] | 0 | 0 | 0 | 1 | 0 | 0 | 0 | 0 | 0 | 1 | 0 | 0 |
| ko04013 MAPK signaling pathway - fly | 3045 | 3028 | 2907 | 2997 | 2796 | 2839 | 2806 | 2673 | 2580 | 2571 | 2580 | 2589 |
| K04564 superoxide dismutase, Fe-Mn family [EC:1.15.1.1] | 3045 | 3028 | 2907 | 2997 | 2796 | 2839 | 2806 | 2673 | 2580 | 2571 | 2579 | 2589 |
| ko04016 MAPK signaling pathway - plant | 17254 | 16503 | 16581 | 17093 | 10411 | 10170 | 9933 | 10375 | 10603 | 10833 | 10503 | 10764 |
| K13413 mitogen-activated protein kinase kinase 4/5 [EC:2.7.12.2] | 0 | 0 | 0 | 0 | 0 | 0 | 0 | 0 | 0 | 0 | 0 | 1 |
| K17686 P-type Cu+ transporter [EC:7.2.2.8] | 8786 | 8204 | 8288 | 8572 | 4794 | 4828 | 4534 | 4873 | 5094 | 5467 | 5198 | 5354 |
| K20716 mitogen-activated protein kinase kinase kinase 17/18 | 0 | 0 | 0 | 0 | 0 | 0 | 0 | 1 | 0 | 0 | 0 | 0 |
| ko04011 MAPK signaling pathway - yeast | 7195 | 7020 | 7036 | 7212 | 4025 | 3826 | 3777 | 3844 | 4174 | 4142 | 4025 | 4047 |
| K04563 cyclin-dependent kinase [EC:2.7.11.22] | 0 | 0 | 0 | 0 | 0 | 0 | 0 | 0 | 0 | 0 | 0 | 1 |
| K06276 3-phosphoinositide dependent protein kinase-1 [EC:2.7.11.1] | 0 | 0 | 0 | 0 | 0 | 0 | 0 | 0 | 1 | 0 | 0 | 0 |
| K06665 general transcriptional corepressor CYC8 | 0 | 0 | 0 | 0 | 0 | 0 | 0 | 0 | 0 | 0 | 1 | 0 |
| K11229 mitogen-activated protein kinase kinase kinase [EC:2.7.11.25] | 0 | 0 | 0 | 0 | 0 | 0 | 0 | 0 | 0 | 0 | 0 | 1 |
| K19806 tyrosine-protein phosphatase 2/3 [EC:3.1.3.48] | 0 | 0 | 0 | 0 | 0 | 0 | 0 | 0 | 0 | 0 | 1 | 0 |
| K19844 GTPase-activating protein BEM2 | 0 | 0 | 0 | 0 | 0 | 0 | 0 | 0 | 0 | 0 | 1 | 0 |
| K23469 G2/mitotic-specific cyclin 2 | 0 | 0 | 2 | 0 | 0 | 0 | 0 | 0 | 0 | 0 | 0 | 0 |
| ko04012 ErbB signaling pathway | 0 | 1 | 0 | 0 | 0 | 1 | 0 | 0 | 0 | 0 | 1 | 0 |
| K07203 serine/threonine-protein kinase mTOR [EC:2.7.11.1] | 0 | 1 | 0 | 0 | 0 | 0 | 0 | 0 | 0 | 0 | 0 | 0 |
| ko04310 Wnt signaling pathway | 0 | 0 | 0 | 0 | 0 | 2 | 0 | 0 | 0 | 0 | 0 | 0 |
| K04496 C-terminal binding protein | 0 | 0 | 0 | 0 | 0 | 1 | 0 | 0 | 0 | 0 | 0 | 0 |
| ko04330 Notch signaling pathway | 0 | 0 | 0 | 0 | 0 | 1 | 0 | 0 | 0 | 0 | 0 | 0 |
| ko04390 Hippo signaling pathway | 1 | 0 | 1 | 0 | 0 | 2 | 0 | 1 | 3 | 0 | 0 | 4 |
| K03982 plasminogen activator inhibitor 1 | 1 | 0 | 1 | 0 | 0 | 0 | 0 | 1 | 3 | 0 | 0 | 3 |
| ko04370 VEGF signaling pathway | 0 | 0 | 0 | 0 | 1 | 1 | 0 | 0 | 0 | 0 | 0 | 0 |
| ko04371 Apelin signaling pathway | 3 | 1 | 1 | 0 | 0 | 0 | 0 | 1 | 3 | 4 | 2 | 6 |
| K07200 5'-AMP-activated protein kinase, regulatory gamma subunit | 0 | 0 | 0 | 0 | 0 | 0 | 0 | 0 | 0 | 0 | 2 | 0 |
| ko04630 Jak-STAT signaling pathway | 0 | 1 | 0 | 0 | 0 | 0 | 0 | 0 | 0 | 0 | 1 | 0 |
| ko04064 NF-kappa B signaling pathway | 0 | 0 | 0 | 1 | 0 | 0 | 0 | 0 | 0 | 1 | 0 | 0 |
| ko04066 HIF-1 signaling pathway | 23815 | 24012 | 23460 | 23637 | 23723 | 24761 | 24150 | 23941 | 23363 | 23128 | 23168 | 23560 |
| ko04068 FoxO signaling pathway | 10256 | 10052 | 9947 | 10208 | 6787 | 6620 | 6625 | 6548 | 6752 | 6676 | 6569 | 6623 |
| ko04020 Calcium signaling pathway | 0 | 0 | 2 | 0 | 0 | 1 | 0 | 0 | 2 | 0 | 1 | 1 |
| K00871 phosphorylase kinase gamma subunit [EC:2.7.11.19] | 0 | 0 | 0 | 0 | 0 | 0 | 0 | 0 | 2 | 0 | 0 | 1 |
| K05853 P-type Ca2+ transporter type 2A [EC:7.2.2.10] | 0 | 0 | 2 | 0 | 0 | 0 | 0 | 0 | 0 | 0 | 1 | 0 |
| ko04070 Phosphatidylinositol signaling system | 7403 | 7692 | 7646 | 7601 | 9057 | 9096 | 8810 | 8743 | 9092 | 9007 | 9065 | 9039 |
| ko04072 Phospholipase D signaling pathway | 927 | 803 | 860 | 901 | 642 | 640 | 545 | 542 | 584 | 584 | 522 | 600 |
| ko04071 Sphingolipid signaling pathway | 1013 | 1084 | 1084 | 1083 | 591 | 585 | 534 | 594 | 350 | 336 | 343 | 310 |
| ko04024 cAMP signaling pathway | 0 | 1 | 3 | 1 | 0 | 2 | 0 | 0 | 0 | 1 | 5 | 1 |
| ko04022 cGMP-PKG signaling pathway | 0 | 0 | 2 | 0 | 0 | 2 | 0 | 0 | 0 | 0 | 1 | 1 |
| ko04151 PI3K-Akt signaling pathway | 8065 | 8053 | 8213 | 8244 | 11411 | 11392 | 11495 | 11451 | 9468 | 9430 | 9559 | 9697 |
| K02091 cyclin-dependent kinase 6 [EC:2.7.11.22] | 0 | 1 | 0 | 0 | 0 | 0 | 0 | 0 | 0 | 0 | 0 | 0 |
| K08266 target of rapamycin complex subunit LST8 | 0 | 0 | 0 | 0 | 0 | 0 | 0 | 1 | 0 | 0 | 0 | 0 |
| ko04152 AMPK signaling pathway | 4959 | 5107 | 5036 | 5074 | 4847 | 5184 | 5014 | 4838 | 5442 | 5465 | 5453 | 5433 |
| K03234 elongation factor 2 | 1 | 3 | 3 | 0 | 3 | 4 | 1 | 2 | 5 | 16 | 14 | 11 |
| ko04150 mTOR signaling pathway | 2 | 1 | 0 | 1 | 0 | 2 | 0 | 1 | 1 | 4 | 1 | 3 |
| ko04075 Plant hormone signal transduction | 0 | 0 | 0 | 0 | 0 | 0 | 0 | 0 | 0 | 0 | 0 | 1 |
| Signaling molecules and interaction | 87 | 86 | 94 | 103 | 126 | 97 | 144 | 114 | 110 | 129 | 87 | 106 |
| ko04080 Neuroactive ligand-receptor interaction | 87 | 86 | 94 | 103 | 126 | 97 | 144 | 114 | 110 | 129 | 87 | 106 |
| K05770 translocator protein | 87 | 86 | 94 | 103 | 126 | 97 | 144 | 114 | 110 | 129 | 87 | 106 |
| Cellular Processes | 852554 | 846686 | 827319 | 841913 | 686255 | 687756 | 673236 | 689112 | 616046 | 615205 | 604625 | 613288 |
| Transport and catabolism | 57136 | 59346 | 61196 | 59445 | 113373 | 112532 | 115402 | 112855 | 94031 | 91192 | 93876 | 91212 |
| ko04144 Endocytosis | 19 | 18 | 28 | 22 | 77 | 67 | 0 | 0 | 47 | 57 | 72 | 47 |
| K10365 capping protein (actin filament) muscle Z-line, beta | 0 | 0 | 0 | 0 | 0 | 0 | 0 | 0 | 0 | 0 | 2 | 0 |
| K12191 charged multivesicular body protein 2A | 0 | 0 | 0 | 0 | 1 | 0 | 0 | 0 | 0 | 0 | 0 | 0 |
| K12481 rabenosyn-5 | 0 | 0 | 0 | 0 | 0 | 0 | 0 | 0 | 0 | 0 | 0 | 1 |
| K17917 sorting nexin-1/2 | 0 | 0 | 0 | 0 | 0 | 0 | 0 | 0 | 0 | 0 | 0 | 1 |
| K17918 sorting nexin-3/12 | 0 | 0 | 1 | 0 | 0 | 0 | 0 | 0 | 0 | 0 | 0 | 0 |
| ko04145 Phagosome | 0 | 0 | 0 | 1 | 0 | 1 | 0 | 0 | 0 | 0 | 0 | 2 |
| ko04142 Lysosome | 26364 | 28255 | 30655 | 28609 | 79743 | 79515 | 81638 | 79073 | 65050 | 62835 | 65419 | 62526 |
| K01279 tripeptidyl-peptidase I [EC:3.4.14.9] | 0 | 1 | 1 | 0 | 0 | 1 | 0 | 0 | 0 | 0 | 1 | 0 |
| K01373 cathepsin F [EC:3.4.22.41] | 3 | 1 | 0 | 1 | 1 | 0 | 1 | 1 | 2 | 3 | 0 | 2 |
| K12381 arylsulfatase G [EC:3.1.6.-] | 0 | 1 | 0 | 1 | 0 | 0 | 0 | 0 | 1 | 8 | 3 | 1 |
| K12399 AP-3 complex subunit sigma | 0 | 0 | 1 | 0 | 0 | 0 | 0 | 0 | 0 | 0 | 0 | 0 |
| K13444 formylglycine-generating enzyme [EC:1.8.3.7] | 14 | 14 | 7 | 9 | 5 | 5 | 2 | 2 | 11 | 12 | 19 | 13 |
| ko04146 Peroxisome | 30480 | 30814 | 30222 | 30538 | 33386 | 32764 | 33610 | 33591 | 28608 | 27944 | 28035 | 28219 |
| K00624 carnitine O-acetyltransferase [EC:2.3.1.7] | 0 | 1 | 0 | 0 | 0 | 2 | 0 | 0 | 0 | 0 | 0 | 0 |
| K04565 superoxide dismutase, Cu-Zn family [EC:1.15.1.1] | 347 | 403 | 322 | 357 | 229 | 235 | 244 | 260 | 524 | 550 | 489 | 563 |
| K13343 peroxin-14 | 0 | 0 | 1 | 0 | 0 | 0 | 0 | 0 | 0 | 0 | 0 | 0 |
| ko04140 Autophagy - animal | 0 | 1 | 0 | 1 | 0 | 1 | 2 | 1 | 5 | 2 | 1 | 4 |
| ko04138 Autophagy - yeast | 272 | 259 | 289 | 275 | 166 | 183 | 154 | 191 | 325 | 356 | 349 | 418 |
| K06655 negative regulator of the PHO system [EC:2.7.11.22] | 0 | 0 | 1 | 0 | 0 | 0 | 0 | 0 | 0 | 0 | 0 | 0 |
| K06902 MFS transporter, UMF1 family | 272 | 258 | 286 | 274 | 165 | 182 | 152 | 188 | 321 | 354 | 349 | 414 |
| K12761 carbon catabolite-derepressing protein kinase [EC:2.7.11.1] | 0 | 0 | 2 | 0 | 0 | 0 | 0 | 0 | 0 | 0 | 0 | 0 |
| K15296 alpha-soluble NSF attachment protein | 0 | 0 | 0 | 0 | 0 | 0 | 0 | 2 | 0 | 0 | 0 | 0 |
| K20179 vacuolar protein sorting-associated protein 11 | 0 | 0 | 0 | 0 | 1 | 0 | 0 | 0 | 0 | 0 | 0 | 0 |
| ko04136 Autophagy - other | 0 | 1 | 0 | 0 | 0 | 0 | 0 | 1 | 0 | 0 | 0 | 0 |
| ko04139 Mitophagy - yeast | 20 | 19 | 26 | 22 | 76 | 69 | 0 | 0 | 47 | 57 | 69 | 44 |
| K08955 ATP-dependent metalloprotease [EC:3.4.24.-] | 0 | 0 | 0 | 0 | 0 | 1 | 0 | 0 | 0 | 0 | 0 | 0 |
| K17774 mitochondrial distribution and morphology protein 10 | 0 | 0 | 2 | 0 | 0 | 0 | 0 | 0 | 0 | 0 | 0 | 0 |
| Cell growth and death | 118730 | 119166 | 120118 | 120010 | 142997 | 142060 | 143408 | 142378 | 122481 | 120894 | 121855 | 120789 |
| ko04110 Cell cycle | 1 | 3 | 1 | 1 | 1 | 0 | 0 | 1 | 2 | 6 | 3 | 4 |
| K02365 separase [EC:3.4.22.49] | 0 | 1 | 0 | 0 | 0 | 0 | 0 | 0 | 0 | 0 | 0 | 0 |

| | | | | | | | | | | | | |
|---|---|---|---|---|---|---|---|---|---|---|---|---|
| ko04111 Cell cycle - yeast | 0 | 2 | 4 | 0 | 0 | 0 | 0 | 0 | 0 | 2 | 1 | 2 |
| K06647 transcription factor MBP1 | 0 | 0 | 0 | 0 | 0 | 0 | 0 | 0 | 0 | 0 | 0 | 1 |
| ko04112 Cell cycle - Caulobacter | 59720 | 60086 | 60909 | 61161 | 70546 | 70690 | 71067 | 69849 | 61788 | 60754 | 61176 | 60755 |
| K01338 ATP-dependent Lon protease [EC:3.4.21.53] | 7617 | 7601 | 7965 | 8096 | 8182 | 8413 | 8263 | 8171 | 8224 | 7962 | 8251 | 8173 |
| K01358 ATP-dependent Clp protease, protease subunit [EC:3.4.21.92] | 3123 | 3062 | 3133 | 3180 | 3074 | 3048 | 3016 | 3028 | 2738 | 2718 | 2749 | 2793 |
| K02651 pilus assembly protein Flp/PilA | 322 | 325 | 346 | 361 | 213 | 188 | 179 | 204 | 125 | 107 | 110 | 98 |
| K03531 cell division protein FtsZ | 3266 | 3160 | 3281 | 3324 | 4560 | 4673 | 4678 | 4325 | 4201 | 4482 | 4094 | 4068 |
| K03544 ATP-dependent Clp protease ATP-binding subunit ClpX | 3450 | 3474 | 3607 | 3551 | 3940 | 3963 | 4181 | 3954 | 4058 | 4384 | 4286 | 4243 |
| K03588 cell division protein FtsW | 4155 | 4338 | 4488 | 4244 | 4947 | 4868 | 4635 | 4880 | 4828 | 4564 | 4411 | 4501 |
| K03589 cell division protein FtsQ | 2766 | 2927 | 2756 | 2849 | 3055 | 2966 | 2992 | 2812 | 2724 | 2693 | 2764 | 2683 |
| K03590 cell division protein FtsA | 2525 | 2803 | 2862 | 2670 | 4311 | 4407 | 4183 | 4209 | 3755 | 3651 | 3748 | 3749 |
| K06985 aspartyl protease family protein | 2432 | 2397 | 2238 | 2323 | 2718 | 2787 | 2804 | 2725 | 1973 | 1729 | 2027 | 1981 |
| K11749 regulator of sigma E protease [EC:3.4.24.-] | 4442 | 4239 | 4221 | 4337 | 4908 | 4637 | 4943 | 4645 | 4369 | 4344 | 4144 | 4345 |
| K13581 modification methylase [EC:2.1.1.72] | 1 | 2 | 2 | 0 | 1 | 0 | 0 | 0 | 1 | 1 | 4 | 2 |
| K13582 localization factor PodJL | 2 | 2 | 2 | 0 | 0 | 2 | 0 | 1 | 6 | 4 | 5 | 7 |
| K13583 GcrA cell cycle regulator | 0 | 0 | 0 | 0 | 2 | 4 | 1 | 1 | 0 | 2 | 1 | 0 |
| K13590 diguanylate cyclase [EC:2.7.7.65] | 4013 | 3997 | 3896 | 3898 | 2315 | 2157 | 2082 | 2171 | 1320 | 1252 | 1212 | 1287 |
| K13592 regulator of CtrA degradation | 2 | 0 | 2 | 0 | 0 | 0 | 2 | 0 | 0 | 0 | 8 | 1 |
| ko04113 Meiosis - yeast | 1713 | 1628 | 1473 | 1628 | 874 | 804 | 827 | 846 | 577 | 476 | 507 | 510 |
| K10878 meiotic recombination protein SPO11 | 0 | 3 | 0 | 0 | 0 | 0 | 0 | 0 | 0 | 0 | 0 | 0 |
| K12762 G protein-coupled receptor GPR1 | 0 | 2 | 0 | 0 | 0 | 2 | 0 | 0 | 0 | 0 | 0 | 0 |
| ko04114 Oocyte meiosis | 0 | 2 | 1 | 0 | 0 | 2 | 0 | 0 | 0 | 0 | 0 | 1 |
| ko04210 Apoptosis | 4779 | 4705 | 4695 | 4691 | 7191 | 7152 | 7076 | 7293 | 5276 | 4819 | 5016 | 4828 |
| K01173 endonuclease G, mitochondrial | 4240 | 4259 | 4225 | 4196 | 6905 | 6884 | 6810 | 7026 | 5063 | 4657 | 4869 | 4680 |
| ko04214 Apoptosis - fly | 4512 | 4329 | 4375 | 4352 | 3495 | 3526 | 3510 | 3422 | 3243 | 3293 | 3203 | 3222 |
| K03386 peroxiredoxin (alkyl hydroperoxide reductase subunit C) [EC:1.11.1.15] | 3889 | 3798 | 3811 | 3756 | 3084 | 3162 | 3103 | 3042 | 2927 | 3006 | 2970 | 2974 |
| ko04215 Apoptosis - multiple species | 536 | 445 | 470 | 493 | 285 | 267 | 263 | 266 | 206 | 158 | 146 | 142 |
| ko04216 Ferroptosis | 10372 | 11098 | 11007 | 10856 | 18986 | 18536 | 19293 | 19142 | 15352 | 14871 | 14985 | 14862 |
| ko04217 Necroptosis | 38168 | 37764 | 38125 | 37814 | 42187 | 41619 | 41898 | 42090 | 36446 | 36833 | 37110 | 36746 |
| K03767 peptidyl-prolyl cis-trans isomerase A (cyclophilin A) [EC:5.2.1.8] | 1305 | 1186 | 1310 | 1288 | 815 | 735 | 717 | 827 | 884 | 877 | 828 | 854 |
| ko04115 p53 signaling pathway | 537 | 446 | 471 | 493 | 285 | 267 | 263 | 267 | 209 | 158 | 146 | 145 |
| ko04218 Cellular senescence | 1 | 2 | 1 | 0 | 2 | 2 | 0 | 1 | 3 | 0 | 0 | 4 |
| Cellular community - eukaryotes | 2 | 0 | 0 | 1 | 2 | 3 | 0 | 1 | 3 | 4 | 6 | 5 |
| ko04510 Focal adhesion | 0 | 0 | 0 | 0 | 1 | 3 | 0 | 0 | 1 | 0 | 1 | 1 |
| ko04520 Adherens junction | 0 | 0 | 0 | 0 | 1 | 0 | 0 | 0 | 0 | 0 | 0 | 0 |
| ko04530 Tight junction | 1 | 0 | 0 | 1 | 2 | 0 | 0 | 1 | 2 | 4 | 5 | 4 |
| ko04540 Gap junction | 0 | 0 | 0 | 0 | 0 | 1 | 0 | 0 | 0 | 0 | 1 | 0 |
| ko04550 Signaling pathways regulating pluripotency of stem cells | 1 | 0 | 0 | 0 | 0 | 0 | 0 | 0 | 0 | 0 | 0 | 0 |
| K14439 SWI/SNF-related matrix-associated actin-dependent regulator of chromatin subfamily A containing DEAD/H box 1 [EC:3.6.4.12] | 1 | 0 | 0 | 0 | 0 | 0 | 0 | 0 | 0 | 0 | 0 | 0 |
| Cellular community - prokaryotes | 561723 | 557523 | 538207 | 551468 | 386790 | 387392 | 376995 | 392329 | 374917 | 376662 | 365223 | 375727 |
| ko02024 Quorum sensing | 269533 | 268066 | 263584 | 268201 | 195748 | 196687 | 193120 | 197908 | 202660 | 205771 | 200171 | 204364 |
| K01318 glutamyl endopeptidase [EC:3.4.21.19] | 0 | 1 | 2 | 0 | 1 | 2 | 0 | 1 | 0 | 1 | 3 | 1 |
| K01399 pseudolysin [EC:3.4.24.26] | 2326 | 2348 | 2320 | 2310 | 1363 | 1294 | 1249 | 1447 | 724 | 714 | 694 | 774 |
| K02031 peptide/nickel transport system ATP-binding protein | 4119 | 4068 | 3953 | 4183 | 1432 | 1678 | 1495 | 1523 | 4947 | 5162 | 4987 | 5172 |
| K02032 peptide/nickel transport system ATP-binding protein | 2972 | 2980 | 2797 | 3048 | 1268 | 1290 | 1220 | 1225 | 4479 | 4652 | 4384 | 4538 |
| K02033 peptide/nickel transport system permease protein | 5025 | 4949 | 4791 | 4998 | 2095 | 2167 | 2083 | 1870 | 6403 | 6572 | 6573 | 6542 |
| K02034 peptide/nickel transport system permease protein | 5217 | 4944 | 5007 | 5244 | 2193 | 2308 | 2107 | 2118 | 6039 | 6250 | 5817 | 6129 |
| K02035 peptide/nickel transport system substrate-binding protein | 10637 | 10571 | 10359 | 10567 | 5462 | 5515 | 5272 | 5446 | 10203 | 10800 | 10154 | 10829 |
| K02052 putative spermidine/putrescine transport system ATP-binding protein | 4397 | 4422 | 4392 | 4387 | 2545 | 2640 | 2494 | 2609 | 2122 | 2153 | 2017 | 2180 |
| K02053 putative spermidine/putrescine transport system permease protein | 1936 | 1972 | 1781 | 2016 | 1152 | 1176 | 1342 | 1333 | 1146 | 1162 | 1175 | 1178 |
| K02054 putative spermidine/putrescine transport system permease protein | 2868 | 2916 | 2878 | 2854 | 1587 | 1707 | 1625 | 1627 | 1424 | 1545 | 1505 | 1541 |
| K02055 putative spermidine/putrescine transport system substrate-binding protein | 8661 | 8595 | 8348 | 8431 | 4986 | 4958 | 4743 | 5005 | 3415 | 3502 | 3450 | 3587 |
| K02250 competence protein ComK | 0 | 3 | 3 | 0 | 0 | 0 | 0 | 0 | 0 | 0 | 2 | 1 |
| K02251 competence protein ComQ | 0 | 0 | 0 | 2 | 0 | 0 | 0 | 3 | 0 | 0 | 0 | 0 |
| K07344 type IV secretion system protein TrbL | 4077 | 3982 | 4026 | 3967 | 2284 | 2189 | 2170 | 2469 | 1272 | 1194 | 1232 | 1206 |
| K08321 3-hydroxy-5-phosphonooxypentane-2,4-dione thiolase [EC:2.3.1.245] | 571 | 535 | 471 | 549 | 402 | 446 | 434 | 458 | 653 | 774 | 712 | 737 |
| K08605 coccolysin [EC:3.4.24.30] | 84 | 75 | 44 | 68 | 9 | 12 | 9 | 7 | 47 | 79 | 76 | 72 |
| K08642 LasA protease [EC:3.4.24.-] | 1740 | 1674 | 1595 | 1614 | 923 | 855 | 776 | 879 | 551 | 524 | 481 | 475 |
| K09823 Fur family transcriptional regulator, zinc uptake regulator | 582 | 587 | 535 | 524 | 402 | 418 | 345 | 346 | 596 | 605 | 636 | 624 |
| K09936 bacterial/archaeal transporter family-2 protein | 2202 | 2199 | 2024 | 2025 | 1037 | 976 | 977 | 1001 | 1693 | 1779 | 1621 | 1608 |
| K11031 thiol-activated cytolysin | 279 | 292 | 312 | 329 | 490 | 405 | 470 | 514 | 30 | 61 | 45 | 47 |
| K11035 hemolysin BL binding component | 0 | 0 | 0 | 0 | 0 | 0 | 0 | 2 | 0 | 0 | 0 | 0 |
| K11063 toxin A/B | 0 | 0 | 0 | 0 | 0 | 0 | 0 | 0 | 0 | 0 | 0 | 1 |
| K11216 autoinducer-2 kinase [EC:2.7.1.189] | 1013 | 968 | 960 | 1053 | 803 | 764 | 828 | 872 | 1197 | 1334 | 1296 | 1361 |
| K11530 (4S)-4-hydroxy-5-phosphonooxypentane-2,3-dione isomerase [EC:5.3.1.32] | 655 | 632 | 611 | 653 | 413 | 395 | 366 | 428 | 355 | 369 | 340 | 364 |
| K11531 lsr operon transcriptional repressor | 647 | 590 | 667 | 621 | 414 | 405 | 416 | 411 | 877 | 765 | 821 | 938 |
| K12990 rhamnosyltransferase [EC:2.4.1.-] | 472 | 458 | 405 | 465 | 215 | 201 | 188 | 244 | 126 | 129 | 87 | 130 |
| K13075 N-acyl homoserine lactone hydrolase [EC:3.1.1.81] | 288 | 292 | 258 | 245 | 97 | 99 | 108 | 95 | 683 | 640 | 581 | 584 |
| K14051 c-di-GMP phosphodiesterase Gmr [EC:3.1.4.52] | 393 | 378 | 372 | 368 | 494 | 618 | 558 | 527 | 101 | 93 | 109 | 95 |
| K14645 serine protease [EC:3.4.21.-] | 5 | 1 | 2 | 1 | 5 | 4 | 9 | 7 | 0 | 2 | 0 | 1 |
| K18099 LuxR family transcriptional regulator, quorum-sensing system regulator RhlR | 1069 | 1022 | 990 | 1045 | 570 | 575 | 612 | 639 | 348 | 339 | 314 | 377 |
| K18100 rhamnosyltransferase subunit A [EC:2.4.1.-] | 1252 | 1155 | 1317 | 1211 | 696 | 728 | 762 | 740 | 406 | 386 | 418 | 376 |
| K18101 rhamnosyltransferase subunit B [EC:2.4.1.-] | 2051 | 2050 | 1964 | 2090 | 1147 | 1151 | 1038 | 1109 | 617 | 608 | 590 | 592 |
| K18139 outer membrane protein, multidrug efflux system | 3509 | 3553 | 3566 | 3778 | 4510 | 4584 | 4565 | 4586 | 4067 | 3807 | 3808 | 3905 |
| K18304 LuxR family transcriptional regulator, quorum-sensing system regulator LasR | 3 | 2 | 2 | 2 | 0 | 3 | 0 | 2 | 0 | 0 | 0 | 0 |
| K18306 membrane fusion protein, multidrug efflux system | 1429 | 1472 | 1246 | 1319 | 772 | 681 | 806 | 905 | 439 | 408 | 441 | 398 |
| K18307 multidrug efflux pump | 2755 | 2747 | 2533 | 2568 | 1499 | 1438 | 1382 | 1667 | 962 | 896 | 741 | 880 |
| K19735 LysR family transcriptional regulator, quorum-sensing system regulator MvfR | 1552 | 1457 | 1456 | 1452 | 853 | 898 | 788 | 859 | 498 | 461 | 428 | 458 |
| K20252 LuxR family transcriptional regulator, quorum-sensing system regulator CinR | 0 | 0 | 0 | 0 | 0 | 0 | 0 | 0 | 0 | 0 | 0 | 1 |
| K20258 PA-I galactophilic lectin | 373 | 406 | 400 | 379 | 226 | 200 | 163 | 214 | 118 | 111 | 120 | 125 |
| K20259 PA-IIL fucose-binding lectin | 500 | 508 | 483 | 554 | 257 | 269 | 267 | 227 | 153 | 194 | 151 | 159 |
| K20265 glutamate:GABA antiporter | 3106 | 3243 | 3093 | 3117 | 4571 | 4411 | 4271 | 4472 | 3520 | 3383 | 3469 | 3350 |
| K20266 type IV secretion system protein TrbJ | 3280 | 3200 | 3279 | 3197 | 1777 | 1788 | 1352 | 1480 | 959 | 948 | 964 | 979 |
| K20271 DeoR family transcriptional regulator, repressor of opine catabolism and conjugal transfer | 0 | 0 | 0 | 0 | 0 | 0 | 0 | 0 | 2 | 4 | 2 | 2 |
| K20276 large repetitive protein | 8703 | 8291 | 7610 | 7945 | 6911 | 7360 | 7690 | 8251 | 1583 | 1572 | 1443 | 1571 |
| K20277 mannose-binding lectin | 564 | 558 | 523 | 563 | 273 | 297 | 287 | 315 | 181 | 143 | 170 | 169 |
| K20345 membrane fusion protein, peptide pheromone/bacteriocin exporter | 129 | 147 | 162 | 132 | 249 | 238 | 254 | 201 | 81 | 128 | 119 | 107 |
| K20373 HTH-type transcriptional regulator, SHP2-responsive activator | 3 | 0 | 2 | 3 | 0 | 0 | 2 | 1 | 0 | 2 | 0 | 0 |
| K20375 HTH-type transcriptional regulator, pheromone-responsive regulator | 0 | 0 | 0 | 0 | 5 | 7 | 3 | 8 | 14 | 14 | 17 | 20 |
| K20377 HTH-type transcriptional regulator, major conjugation operon repressor | 13 | 12 | 6 | 5 | 0 | 0 | 0 | 0 | 0 | 0 | 0 | 0 |
| K20379 sex pheromone cAD1 | 48 | 67 | 58 | 74 | 7 | 13 | 5 | 12 | 51 | 69 | 24 | 58 |
| K20380 HTH-type transcriptional regulator, pheromone-responsive repressor | 69 | 58 | 91 | 76 | 8 | 7 | 5 | 7 | 4 | 0 | 3 | 5 |
| K20381 regulatory protein | 33 | 32 | 32 | 28 | 3 | 4 | 8 | 5 | 2 | 18 | 4 | 4 |
| K20382 aggregation substance | 280 | 308 | 308 | 310 | 20 | 17 | 14 | 23 | 25 | 13 | 17 | 15 |
| K20383 CylL-L protein | 19 | 16 | 30 | 20 | 5 | 0 | 0 | 3 | 0 | 1 | 1 | 0 |
| K20384 CylL-S protein | 24 | 13 | 11 | 19 | 2 | 1 | 2 | 0 | 0 | 2 | 2 | 1 |
| K20385 CylM protein | 210 | 235 | 217 | 247 | 14 | 16 | 9 | 15 | 5 | 16 | 10 | 5 |
| K20387 regulatory protein CylR1 | 9 | 13 | 12 | 10 | 0 | 1 | 0 | 0 | 1 | 1 | 0 | 0 |
| K20388 HTH-type transcriptional regulator, cytolysin regulator | 8 | 9 | 14 | 11 | 0 | 1 | 0 | 0 | 0 | 2 | 3 | 0 |
| K20480 HTH-type transcriptional regulator, quorum sensing regulator NprR | 2 | 0 | 2 | 0 | 0 | 0 | 0 | 3 | 1 | 1 | 2 | 3 |
| K20527 type IV secretion system protein TrbB [EC:7.4.2.8] | 3292 | 3231 | 3157 | 3261 | 1770 | 1713 | 1661 | 1715 | 1064 | 928 | 936 | 944 |
| K20528 type IV secretion system protein TrbC | 1079 | 1060 | 1037 | 1016 | 599 | 615 | 506 | 607 | 321 | 298 | 302 | 334 |
| K20529 type IV secretion system protein TrbD | 750 | 779 | 775 | 730 | 375 | 417 | 398 | 393 | 219 | 212 | 211 | 208 |
| K20530 type IV secretion system protein TrbE [EC:7.4.2.8] | 8042 | 7997 | 7602 | 8030 | 4280 | 4199 | 4173 | 4688 | 2493 | 2407 | 2373 | 2514 |
| K20531 type IV secretion system protein TrbF | 2102 | 2075 | 2012 | 2123 | 1075 | 1043 | 946 | 1198 | 669 | 703 | 613 | 590 |
| K20532 type IV secretion system protein TrbG | 4341 | 4269 | 4134 | 4170 | 2294 | 2134 | 2097 | 2355 | 1294 | 1222 | 1258 | 1195 |
| K20533 type IV secretion system protein TrbI | 5616 | 5365 | 5233 | 5456 | 3040 | 2685 | 2669 | 2981 | 1663 | 1546 | 1653 | 1600 |
| K20539 IclR family transcriptional regulator, blcABC operon repressor | 0 | 0 | 0 | 0 | 2 | 0 | 2 | 0 | 0 | 2 | 2 | 1 |
| ko05111 Biofilm formation - Vibrio cholerae | 86461 | 86502 | 82069 | 84310 | 63006 | 62478 | 61230 | 64076 | 61266 | 61081 | 59257 | 61054 |
| K02463 general secretion pathway protein N | 289 | 350 | 283 | 306 | 65 | 79 | 88 | 88 | 524 | 542 | 532 | 639 |
| K03087 RNA polymerase nonessential primary-like sigma factor | 2240 | 2349 | 2215 | 2383 | 1378 | 1348 | 1313 | 1304 | 1385 | 1440 | 1417 | 1440 |
| K03557 Fis family transcriptional regulator, factor for inversion stimulation protein | 663 | 668 | 625 | 573 | 358 | 397 | 378 | 432 | 374 | 411 | 383 | 351 |
| K03606 putative colanic acid biosysnthesis UDP-glucose lipid carrier transferase | 2299 | 2322 | 2544 | 2432 | 7525 | 7374 | 7668 | 7551 | 5499 | 5040 | 5271 | 5065 |
| K05946 N-acetylglucosaminyldiphosphoundecaprenol N-acetyl-beta-D-mannosaminyltransferase [EC:2.4.1.187] | 599 | 646 | 645 | 638 | 458 | 497 | 522 | 477 | 701 | 819 | 754 | 739 |
| K08604 vibriolysin [EC:3.4.24.25] | 0 | 0 | 0 | 0 | 0 | 0 | 0 | 0 | 1 | 0 | 0 | 0 |
| K10924 MSHA pilin protein MshA | 1 | 2 | 0 | 1 | 0 | 0 | 0 | 0 | 1 | 1 | 0 | 0 |
| K10939 accessory colonization factor AcfD | 8 | 14 | 14 | 20 | 26 | 47 | 35 | 27 | 185 | 231 | 233 | 231 |
| K12276 MSHA biogenesis protein MshE | 1 | 2 | 2 | 1 | 0 | 0 | 1 | 1 | 4 | 0 | 3 | 1 |
| K16554 polysaccharide biosynthesis transport protein | 0 | 0 | 0 | 0 | 3 | 2 | 8 | 2 | 4 | 0 | 3 | 1 |
| K20920 polysaccharide biosynthesis protein VpsM | 0 | 0 | 0 | 0 | 2 | 2 | 0 | 2 | 4 | 0 | 2 | 0 |
| K20922 polysaccharide biosynthesis protein VpsI | 1 | 1 | 0 | 1 | 0 | 0 | 0 | 0 | 1 | 0 | 0 | 0 |

| | | | | | | | | | | | | |
|---|---|---|---|---|---|---|---|---|---|---|---|---|
| K20955 diguanylate cyclase [EC:2.7.7.65] | 0 | 0 | 0 | 0 | 0 | 0 | 0 | 1 | 0 | 0 | 0 | 0 |
| K20956 diguanylate cyclase [EC:2.7.7.65] | 0 | 0 | 1 | 0 | 1 | 0 | 0 | 0 | 0 | 0 | 0 | 1 |
| K20957 diguanylate cyclase [EC:2.7.7.65] | 2 | 2 | 1 | 0 | 0 | 0 | 0 | 0 | 0 | 0 | 1 | 0 |
| K20959 diguanylate cyclase [EC:2.7.7.65] | 0 | 1 | 0 | 2 | 2 | 2 | 0 | 1 | 0 | 0 | 0 | 2 |
| K20962 c-di-GMP phosphodiesterase [EC:3.1.4.52] | 1 | 0 | 1 | 0 | 0 | 0 | 0 | 0 | 0 | 0 | 0 | 0 |
| K20965 c-di-GMP phosphodiesterase [EC:3.1.4.52] | 1 | 0 | 1 | 0 | 0 | 0 | 0 | 0 | 1 | 0 | 2 | 1 |
| K20966 c-di-GMP phosphodiesterase [EC:3.1.4.52] | 9 | 10 | 5 | 7 | 2 | 0 | 0 | 1 | 2 | 4 | 4 | 5 |
| K20988 polysaccharide biosynthesis/export protein VpsN | 0 | 0 | 3 | 0 | 0 | 0 | 0 | 0 | 0 | 0 | 2 | 0 |
| ko02025 Biofilm formation - Pseudomonas aeruginosa | 197505 | 194776 | 185121 | 190346 | 112339 | 111490 | 106507 | 115329 | 80408 | 77735 | 74610 | 78166 |
| K11890 type VI secretion system protein ImpM | 889 | 780 | 674 | 807 | 510 | 538 | 533 | 596 | 243 | 192 | 191 | 187 |
| K11893 type VI secretion system protein ImpJ | 6081 | 5798 | 5460 | 5615 | 3329 | 3372 | 3140 | 3184 | 3333 | 3324 | 3054 | 3176 |
| K11895 type VI secretion system protein ImpH | 5736 | 5720 | 5227 | 5540 | 3380 | 3399 | 3045 | 3343 | 2659 | 2725 | 2684 | 2760 |
| K11900 type VI secretion system protein ImpC | 6497 | 6612 | 6290 | 6482 | 3859 | 4000 | 3646 | 4078 | 2743 | 2594 | 2615 | 2669 |
| K11901 type VI secretion system protein ImpB | 2701 | 2737 | 2551 | 2664 | 1553 | 1578 | 1581 | 1646 | 1116 | 1108 | 1039 | 1186 |
| K11902 type VI secretion system protein ImpA | 3531 | 3402 | 3026 | 3247 | 2030 | 1925 | 1714 | 1852 | 988 | 897 | 830 | 973 |
| K12992 rhamnosyltransferase [EC:2.4.1.-] | 1687 | 1603 | 1482 | 1568 | 787 | 695 | 681 | 687 | 682 | 618 | 590 | 678 |
| K20968 AraC family transcriptional regulator, exoenzyme S synthesis regulatory protein ExsA | 154 | 175 | 190 | 209 | 109 | 111 | 88 | 102 | 49 | 49 | 54 | 50 |
| K20971 two-component system, sensor histidine kinase LadS | 42 | 55 | 38 | 44 | 19 | 28 | 171 | 223 | 16 | 27 | 16 | 17 |
| K20972 two-component system, sensor histidine kinase RetS | 432 | 454 | 404 | 413 | 256 | 226 | 196 | 228 | 168 | 111 | 132 | 149 |
| K20987 polysaccharide biosynthesis/export protein PslD | 108 | 127 | 94 | 103 | 51 | 44 | 46 | 55 | 40 | 27 | 24 | 30 |
| K20997 polysaccharide biosynthesis protein PslA | 1967 | 1897 | 1876 | 1921 | 1082 | 1020 | 982 | 976 | 616 | 555 | 483 | 571 |
| K20998 polysaccharide biosynthesis protein PslE | 6 | 5 | 5 | 7 | 1 | 3 | 3 | 1 | 2 | 2 | 1 | 2 |
| K20999 polysaccharide biosynthesis protein PslF | 1296 | 1364 | 1273 | 1311 | 732 | 668 | 640 | 700 | 456 | 418 | 410 | 421 |
| K21000 polysaccharide biosynthesis protein PslG | 388 | 345 | 288 | 311 | 182 | 186 | 187 | 207 | 116 | 101 | 90 | 88 |
| K21001 polysaccharide biosynthesis protein PslH | 1653 | 1503 | 1378 | 1484 | 734 | 830 | 738 | 826 | 463 | 468 | 409 | 402 |
| K21002 polysaccharide biosynthesis protein PslI | 1437 | 1257 | 1138 | 1263 | 641 | 682 | 558 | 668 | 400 | 385 | 308 | 410 |
| K21003 polysaccharide biosynthesis protein PslJ | 962 | 1052 | 917 | 919 | 489 | 467 | 459 | 510 | 320 | 317 | 243 | 287 |
| K21004 polysaccharide biosynthesis protein PslK | 1704 | 1704 | 1462 | 1638 | 845 | 837 | 743 | 896 | 451 | 448 | 426 | 467 |
| K21005 polysaccharide biosynthesis protein PslL | 374 | 373 | 350 | 350 | 180 | 188 | 167 | 227 | 109 | 91 | 107 | 120 |
| K21006 polysaccharide biosynthesis protein PelA | 3802 | 3722 | 3393 | 3447 | 1959 | 1783 | 1808 | 2021 | 1088 | 1087 | 1116 | 1167 |
| K21007 polysaccharide biosynthesis protein PelB | 3546 | 3424 | 3145 | 3192 | 1730 | 1712 | 1785 | 1823 | 1091 | 945 | 1039 | 976 |
| K21008 polysaccharide biosynthesis protein PelC | 124 | 100 | 91 | 91 | 84 | 58 | 49 | 57 | 39 | 30 | 33 | 34 |
| K21009 polysaccharide biosynthesis protein PelD | 1295 | 1296 | 1201 | 1214 | 676 | 637 | 637 | 775 | 441 | 388 | 373 | 379 |
| K21010 polysaccharide biosynthesis protein PelE | 1086 | 1032 | 1114 | 1066 | 610 | 588 | 563 | 580 | 359 | 373 | 299 | 319 |
| K21011 polysaccharide biosynthesis protein PelF | 2162 | 2091 | 2192 | 2145 | 1161 | 1252 | 1098 | 1154 | 819 | 725 | 715 | 776 |
| K21012 polysaccharide biosynthesis protein PelG | 1961 | 1948 | 2048 | 1961 | 1092 | 1002 | 1044 | 1089 | 649 | 655 | 703 | 667 |
| K21019 diguanylate cyclase [EC:2.7.7.65] | 2142 | 2191 | 2015 | 2029 | 1192 | 1257 | 1121 | 1118 | 714 | 662 | 610 | 657 |
| K21020 diguanylate cyclase [EC:2.7.7.65] | 1607 | 1550 | 1538 | 1575 | 915 | 857 | 787 | 885 | 536 | 497 | 437 | 488 |
| K21021 diguanylate cyclase [EC:2.7.7.65] | 2573 | 2370 | 2307 | 2348 | 1175 | 1145 | 1024 | 1195 | 1552 | 1451 | 1381 | 1551 |
| K21022 diguanylate cyclase [EC:2.7.7.65] | 1871 | 1730 | 1770 | 1663 | 891 | 883 | 899 | 983 | 520 | 596 | 535 | 548 |
| K21023 diguanylate cyclase [EC:2.7.7.65] | 1197 | 1199 | 1131 | 1293 | 1304 | 1303 | 1333 | 1325 | 171 | 186 | 173 | 179 |
| K21024 c-di-GMP phosphodiesterase [EC:3.1.4.52] | 3285 | 3215 | 3094 | 3172 | 1757 | 1800 | 1678 | 1935 | 1033 | 1026 | 1018 | 1060 |
| K21025 multidomain signaling protein FimX | 1367 | 1329 | 1297 | 1347 | 780 | 838 | 688 | 804 | 446 | 451 | 419 | 439 |
| K23127 cyclic-di-GMP-binding protein | 2914 | 2762 | 2722 | 2817 | 1521 | 1517 | 1524 | 1572 | 861 | 928 | 889 | 987 |
| ko02026 Biofilm formation - Escherichia coli | 70504 | 70566 | 68389 | 70531 | 55029 | 56103 | 54450 | 54952 | 66313 | 68170 | 66065 | 68148 |
| K02425 regulator of sigma S factor FliZ | 110 | 107 | 103 | 76 | 164 | 187 | 156 | 145 | 21 | 25 | 33 | 20 |
| K03566 LysR family transcriptional regulator, glycine cleavage system transcriptional activator | 5336 | 5334 | 4976 | 5280 | 3537 | 3561 | 3236 | 3655 | 4335 | 4519 | 4184 | 4455 |
| K03567 glycine cleavage system transcriptional repressor | 1285 | 1360 | 1280 | 1306 | 778 | 846 | 750 | 774 | 1040 | 1076 | 952 | 1115 |
| K04333 LuxR family transcriptional regulator, csgAB operon transcriptional regulatory protein | 401 | 395 | 361 | 403 | 243 | 263 | 252 | 298 | 482 | 524 | 494 | 563 |
| K04334 major curlin subunit | 92 | 97 | 103 | 118 | 122 | 142 | 94 | 69 | 21 | 22 | 32 | 18 |
| K04335 minor curlin subunit | 33 | 33 | 24 | 32 | 51 | 59 | 55 | 48 | 12 | 7 | 11 | 9 |
| K04336 curli production protein | 68 | 67 | 62 | 64 | 108 | 96 | 87 | 122 | 30 | 11 | 29 | 28 |
| K04761 LysR family transcriptional regulator, hydrogen peroxide-inducible genes activator | 3045 | 2983 | 2944 | 3090 | 3729 | 3509 | 3607 | 3738 | 2635 | 2622 | 2625 | 2597 |
| K06204 DnaK suppressor protein | 1816 | 1785 | 1720 | 1691 | 1047 | 1083 | 1068 | 1114 | 1004 | 918 | 918 | 957 |
| K11931 poly-beta-1,6-N-acetyl-D-glucosamine N-deacetylase [EC:3.5.1.-] | 922 | 877 | 860 | 863 | 213 | 241 | 251 | 249 | 1742 | 1737 | 1759 | 1930 |
| K11935 biofilm PGA synthesis protein PgaA | 972 | 1019 | 984 | 945 | 280 | 254 | 303 | 267 | 2140 | 2294 | 2206 | 2247 |
| K11936 poly-beta-1,6-N-acetyl-D-glucosamine synthase [EC:2.4.1.-] | 544 | 533 | 519 | 597 | 181 | 139 | 143 | 147 | 1154 | 1191 | 1123 | 1139 |
| K11937 biofilm PGA synthesis protein PgaD | 145 | 181 | 181 | 180 | 50 | 43 | 55 | 67 | 383 | 408 | 407 | 376 |
| K12687 antigen 43 | 15 | 7 | 5 | 9 | 29 | 33 | 13 | 22 | 114 | 118 | 113 | 115 |
| K18968 diguanylate cyclase [EC:2.7.7.65] | 242 | 207 | 181 | 206 | 353 | 366 | 380 | 329 | 85 | 61 | 77 | 74 |
| K21084 diguanylate cyclase [EC:2.7.7.65] | 773 | 691 | 710 | 683 | 948 | 1094 | 1075 | 966 | 207 | 196 | 282 | 238 |
| K21085 diguanylate cyclase [EC:2.7.7.65] | 887 | 933 | 902 | 845 | 583 | 707 | 616 | 615 | 1160 | 1287 | 1265 | 1284 |
| K21086 c-di-GMP phosphodiesterase [EC:3.1.4.52] | 508 | 444 | 399 | 433 | 341 | 284 | 337 | 307 | 539 | 588 | 554 | 587 |
| K21087 flagellar brake protein | 168 | 129 | 154 | 155 | 244 | 226 | 231 | 198 | 50 | 35 | 38 | 44 |
| K21088 diguanylate cyclase [EC:2.7.7.65] | 602 | 571 | 625 | 672 | 458 | 498 | 442 | 440 | 876 | 847 | 863 | 942 |
| K21089 MerR family transcriptional regulator, activator of the csg genes | 166 | 143 | 146 | 152 | 246 | 222 | 211 | 236 | 28 | 44 | 44 | 39 |
| K21090 c-di-GMP phosphodiesterase [EC:3.1.4.52] | 881 | 837 | 787 | 915 | 663 | 631 | 663 | 622 | 1098 | 1194 | 1177 | 1282 |
| Cell motility | 142652 | 139723 | 136489 | 139293 | 88504 | 90244 | 83421 | 86997 | 62180 | 63149 | 60881 | 62317 |
| ko02030 Bacterial chemotaxis | 98814 | 96934 | 94197 | 95789 | 62468 | 63504 | 58017 | 60325 | 42698 | 42722 | 41364 | 42374 |
| K02410 flagellar motor switch protein FliG | 2057 | 1899 | 1896 | 1947 | 1123 | 1112 | 1064 | 1080 | 1137 | 1219 | 1158 | 1215 |
| K02416 flagellar motor switch protein FliM | 2209 | 2191 | 2104 | 2218 | 1200 | 1299 | 1232 | 1181 | 1128 | 1095 | 1091 | 1117 |
| K02417 flagellar motor switch protein FliN/FliY | 1652 | 1638 | 1632 | 1633 | 683 | 778 | 672 | 734 | 844 | 1017 | 976 | 969 |
| K02557 chemotaxis protein MotB | 3165 | 3098 | 3216 | 3065 | 3340 | 3365 | 3283 | 3221 | 2722 | 2564 | 2653 | 2498 |
| K03409 chemotaxis protein CheX | 2 | 2 | 2 | 0 | 0 | 0 | 0 | 0 | 0 | 0 | 0 | 0 |
| K03410 chemotaxis protein CheC | 346 | 308 | 340 | 356 | 233 | 276 | 268 | 235 | 407 | 392 | 438 | 403 |
| K03411 chemotaxis protein CheD [EC:3.5.1.44] | 1395 | 1213 | 1131 | 1220 | 601 | 629 | 597 | 693 | 659 | 617 | 554 | 586 |
| K03414 chemotaxis protein CheZ | 693 | 632 | 654 | 648 | 427 | 398 | 460 | 462 | 208 | 192 | 161 | 181 |
| ko02040 Flagellar assembly | 55198 | 53757 | 53256 | 54569 | 33501 | 34441 | 32808 | 34082 | 26444 | 27453 | 26412 | 26790 |
| K02386 flagella basal body P-ring formation protein FlgA | 1260 | 1313 | 1262 | 1214 | 721 | 773 | 763 | 858 | 324 | 396 | 333 | 334 |
| K02387 flagellar basal-body rod protein FlgB | 603 | 642 | 678 | 651 | 360 | 327 | 382 | 359 | 298 | 306 | 329 | 348 |
| K02388 flagellar basal-body rod protein FlgC | 985 | 918 | 987 | 986 | 529 | 525 | 496 | 515 | 537 | 563 | 495 | 496 |
| K02389 flagellar basal-body rod modification protein FlgD | 1327 | 1329 | 1377 | 1483 | 775 | 774 | 798 | 792 | 489 | 536 | 464 | 555 |
| K02390 flagellar hook protein FlgE | 1404 | 1372 | 1330 | 1360 | 885 | 884 | 799 | 842 | 770 | 853 | 714 | 795 |
| K02391 flagellar basal-body rod protein FlgF | 1290 | 1304 | 1331 | 1290 | 940 | 881 | 818 | 905 | 359 | 381 | 388 | 404 |
| K02392 flagellar basal-body rod protein FlgG | 1047 | 949 | 1008 | 1009 | 516 | 589 | 514 | 471 | 722 | 899 | 854 | 795 |
| K02393 flagellar L-ring protein precursor FlgH | 1193 | 1160 | 1230 | 1190 | 816 | 759 | 739 | 798 | 409 | 360 | 374 | 388 |
| K02394 flagellar P-ring protein precursor FlgI | 289 | 326 | 316 | 340 | 377 | 424 | 341 | 391 | 106 | 93 | 102 | 87 |
| K02396 flagellar hook-associated protein 1 FlgK | 2307 | 2223 | 2276 | 2288 | 1377 | 1476 | 1344 | 1319 | 1079 | 1208 | 1201 | 1210 |
| K02397 flagellar hook-associated protein 3 FlgL | 2783 | 2621 | 2716 | 2821 | 1400 | 1502 | 1471 | 1585 | 1263 | 1318 | 1193 | 1233 |
| K02399 flagella synthesis protein FlgN | 833 | 726 | 726 | 793 | 489 | 495 | 455 | 474 | 243 | 245 | 231 | 231 |
| K02400 flagellar biosynthesis protein FlhA | 4942 | 4672 | 4684 | 4872 | 2496 | 2614 | 2287 | 2630 | 2674 | 2652 | 2653 | 2788 |
| K02401 flagellar biosynthetic protein FlhB | 1416 | 1383 | 1498 | 1360 | 813 | 892 | 809 | 840 | 803 | 856 | 815 | 832 |
| K02407 flagellar hook-associated protein 2 | 3135 | 3117 | 3015 | 3159 | 1702 | 1771 | 1691 | 1725 | 1307 | 1386 | 1303 | 1353 |
| K02408 flagellar hook-basal body complex protein FliE | 703 | 665 | 713 | 627 | 361 | 381 | 368 | 416 | 250 | 241 | 278 | 290 |
| K02409 flagellar M-ring protein FliF | 3024 | 3158 | 2875 | 3033 | 1790 | 1962 | 1586 | 1752 | 1258 | 1249 | 1128 | 1173 |
| K02411 flagellar assembly protein FliH | 1443 | 1364 | 1227 | 1334 | 813 | 760 | 745 | 851 | 497 | 520 | 491 | 533 |
| K02412 flagellum-specific ATP synthase [EC:7.4.2.8] | 2859 | 2827 | 2609 | 2693 | 1664 | 1647 | 1646 | 1642 | 1368 | 1367 | 1313 | 1294 |
| K02413 flagellar FliJ protein | 549 | 561 | 543 | 564 | 344 | 346 | 368 | 375 | 201 | 180 | 208 | 214 |
| K02414 flagellar hook-length control protein FliK | 1874 | 1908 | 1737 | 1940 | 1185 | 1214 | 1196 | 1357 | 580 | 542 | 567 | 585 |
| K02418 flagellar protein FliO/FliZ | 478 | 477 | 442 | 484 | 342 | 334 | 326 | 311 | 160 | 199 | 176 | 184 |
| K02419 flagellar biosynthetic protein FliP | 944 | 922 | 900 | 941 | 529 | 507 | 526 | 502 | 599 | 668 | 690 | 659 |
| K02420 flagellar biosynthetic protein FliQ | 513 | 469 | 455 | 530 | 262 | 277 | 291 | 282 | 245 | 280 | 250 | 238 |
| K02421 flagellar biosynthetic protein FliR | 851 | 832 | 814 | 817 | 467 | 542 | 513 | 450 | 468 | 498 | 564 | 500 |
| K02422 flagellar protein FliS | 923 | 867 | 868 | 851 | 535 | 510 | 503 | 522 | 409 | 443 | 449 | 410 |
| K02423 flagellar protein FliT | 84 | 65 | 61 | 93 | 119 | 116 | 121 | 121 | 20 | 10 | 22 | 23 |
| K13820 flagellar biosynthetic protein FliR/FlhB | 1 | 0 | 0 | 0 | 0 | 0 | 0 | 0 | 4 | 9 | 4 | 9 |
| K21218 sodium-type flagellar protein MotY | 1398 | 1322 | 1295 | 1394 | 814 | 717 | 737 | 810 | 512 | 404 | 393 | 413 |
| ko04810 Regulation of actin cytoskeleton | 19 | 18 | 24 | 22 | 76 | 69 | 0 | 0 | 47 | 57 | 70 | 44 |
| Organismal Systems | 178529 | 179361 | 178429 | 178268 | 184535 | 185254 | 184739 | 185634 | 165612 | 164450 | 164714 | 163088 |
| Immune system | 13041 | 13177 | 13501 | 13391 | 17039 | 16991 | 17169 | 17009 | 14716 | 14777 | 14722 | 14770 |
| ko04610 Complement and coagulation cascades | 1 | 0 | 1 | 0 | 0 | 0 | 0 | 1 | 3 | 0 | 0 | 3 |
| ko04611 Platelet activation | 0 | 0 | 0 | 0 | 0 | 2 | 0 | 0 | 0 | 0 | 0 | 1 |
| ko04620 Toll-like receptor signaling pathway | 0 | 0 | 0 | 1 | 0 | 0 | 0 | 0 | 0 | 1 | 0 | 0 |
| ko04624 Toll and Imd signaling pathway | 0 | 0 | 0 | 1 | 0 | 0 | 0 | 2 | 0 | 1 | 0 | 0 |
| K10380 ankyrin | 0 | 0 | 0 | 0 | 0 | 0 | 0 | 2 | 0 | 0 | 0 | 0 |
| ko04621 NOD-like receptor signaling pathway | 11995 | 12071 | 12429 | 12323 | 16270 | 16187 | 16457 | 16340 | 13252 | 13406 | 13353 | 13286 |
| K03671 thioredoxin 1 | 2520 | 2662 | 2851 | 2688 | 3470 | 3304 | 3363 | 3367 | 3312 | 3363 | 3317 | 3101 |
| K04612 calcium-sensing receptor | 0 | 0 | 0 | 0 | 0 | 0 | 0 | 0 | 1 | 0 | 0 | 0 |
| ko04622 RIG-I-like receptor signaling pathway | 1025 | 1087 | 1046 | 1046 | 693 | 734 | 712 | 666 | 1413 | 1314 | 1298 | 1436 |
| K11594 ATP-dependent RNA helicase DDX3X [EC:3.6.4.13] | 0 | 0 | 0 | 0 | 0 | 0 | 0 | 0 | 0 | 1 | 0 | 0 |
| ko04650 Natural killer cell mediated cytotoxicity | 0 | 0 | 0 | 0 | 0 | 1 | 0 | 0 | 0 | 0 | 1 | 0 |

| | | | | | | | | | | | | |
|---|---|---|---|---|---|---|---|---|---|---|---|---|
| ko04612 Antigen processing and presentation | 8026 | 8029 | 8184 | 8223 | 11366 | 11368 | 11453 | 11419 | 9422 | 9406 | 9524 | 9662 |
| ko04660 T cell receptor signaling pathway | 0 | 0 | 0 | 0 | 1 | 0 | 0 | 0 | 1 | 0 | 1 | 0 |
| ko04659 Th17 cell differentiation | 8026 | 8030 | 8183 | 8223 | 11366 | 11368 | 11453 | 11419 | 9422 | 9406 | 9524 | 9661 |
| ko04657 IL-17 signaling pathway | 8027 | 8029 | 8183 | 8223 | 11366 | 11368 | 11453 | 11419 | 9422 | 9406 | 9524 | 9661 |
| ko04662 B cell receptor signaling pathway | 0 | 0 | 0 | 0 | 0 | 0 | 0 | 0 | 0 | 0 | 1 | 0 |
| ko04664 Fc epsilon RI signaling pathway | 0 | 0 | 0 | 0 | 0 | 1 | 0 | 0 | 1 | 0 | 1 | 0 |
| ko04666 Fc gamma R-mediated phagocytosis | 19 | 18 | 24 | 22 | 76 | 68 | 0 | 0 | 47 | 57 | 70 | 43 |
| ko04670 Leukocyte transendothelial migration | 0 | 0 | 0 | 0 | 1 | 1 | 0 | 0 | 0 | 0 | 0 | 0 |
| ko04062 Chemokine signaling pathway | 0 | 0 | 0 | 0 | 1 | 0 | 0 | 0 | 0 | 0 | 1 | 0 |
| Endocrine system | 85228 | 86103 | 84534 | 84701 | 89535 | 90344 | 90104 | 91208 | 80633 | 79365 | 80025 | 79369 |
| ko04911 Insulin secretion | 0 | 0 | 0 | 0 | 0 | 1 | 0 | 0 | 0 | 0 | 0 | 0 |
| ko04910 Insulin signaling pathway | 10000 | 10423 | 10406 | 10170 | 14543 | 14946 | 14744 | 14399 | 14021 | 13968 | 14479 | 13865 |
| K07192 flotillin | 895 | 968 | 1003 | 957 | 2053 | 2140 | 2205 | 2095 | 1778 | 1672 | 1812 | 1623 |
| ko04922 Glucagon signaling pathway | 24570 | 24749 | 24363 | 24073 | 24600 | 25086 | 24727 | 24663 | 26096 | 26245 | 26454 | 25811 |
| ko04923 Regulation of lipolysis in adipocytes | 1 | 0 | 0 | 1 | 0 | 1 | 0 | 1 | 0 | 0 | 0 | 1 |
| ko04920 Adipocytokine signaling pathway | 8806 | 9429 | 9308 | 9114 | 17271 | 16719 | 17607 | 17426 | 13721 | 13107 | 13394 | 13271 |
| ko03320 PPAR signaling pathway | 36028 | 36345 | 35679 | 35800 | 36219 | 35948 | 36046 | 37037 | 30356 | 29701 | 29415 | 29738 |
| K08770 ubiquitin C | 1 | 0 | 0 | 0 | 1 | 0 | 0 | 0 | 0 | 0 | 0 | 0 |
| ko04912 GnRH signaling pathway | 0 | 0 | 0 | 0 | 1 | 1 | 0 | 0 | 0 | 0 | 2 | 0 |
| ko04915 Estrogen signaling pathway | 8027 | 8029 | 8184 | 8223 | 11366 | 11368 | 11453 | 11419 | 9422 | 9406 | 9525 | 9662 |
| ko04914 Progesterone-mediated oocyte maturation | 8026 | 8030 | 8184 | 8223 | 11366 | 11368 | 11453 | 11419 | 9422 | 9408 | 9524 | 9661 |
| ko04917 Prolactin signaling pathway | 1321 | 1413 | 1373 | 1488 | 1030 | 1083 | 983 | 977 | 1529 | 1441 | 1495 | 1511 |
| ko04921 Oxytocin signaling pathway | 1 | 3 | 3 | 0 | 3 | 7 | 1 | 2 | 5 | 16 | 16 | 12 |
| ko04926 Relaxin signaling pathway | 0 | 0 | 0 | 0 | 0 | 1 | 0 | 0 | 0 | 0 | 1 | 0 |
| ko04918 Thyroid hormone synthesis | 6142 | 6104 | 5909 | 5851 | 4383 | 4551 | 4584 | 4660 | 4342 | 4239 | 4219 | 4188 |
| ko04919 Thyroid hormone signaling pathway | 6513 | 6573 | 6100 | 6453 | 3388 | 3541 | 3352 | 3634 | 2144 | 1954 | 1809 | 2070 |
| ko04928 Parathyroid hormone synthesis, secretion and action | 0 | 0 | 0 | 0 | 0 | 2 | 0 | 1 | 5 | 1 | 3 | 1 |
| K03626 nascent polypeptide-associated complex subunit alpha | 0 | 0 | 0 | 0 | 0 | 1 | 0 | 1 | 4 | 1 | 2 | 1 |
| ko04916 Melanogenesis | 0 | 0 | 0 | 0 | 0 | 1 | 0 | 0 | 0 | 0 | 0 | 0 |
| ko04614 Renin-angiotensin system | 1101 | 1213 | 1233 | 1176 | 3977 | 4022 | 4125 | 4042 | 3029 | 2895 | 3260 | 2845 |
| K01322 prolyl oligopeptidase [EC:3.4.21.26] | 1101 | 1208 | 1233 | 1176 | 3977 | 4022 | 4125 | 4042 | 3028 | 2895 | 3257 | 2845 |
| K01392 thimet oligopeptidase [EC:3.4.24.15] | 0 | 0 | 0 | 0 | 0 | 0 | 0 | 0 | 1 | 0 | 0 | 0 |
| K08780 carboxypeptidase A3 [EC:3.4.17.1] | 0 | 5 | 0 | 0 | 0 | 0 | 0 | 0 | 0 | 0 | 3 | 0 |
| ko04925 Aldosterone synthesis and secretion | 0 | 0 | 0 | 0 | 0 | 1 | 0 | 0 | 0 | 0 | 0 | 0 |
| ko04927 Cortisol synthesis and secretion | 0 | 0 | 1 | 1 | 0 | 0 | 0 | 0 | 6 | 4 | 10 | 12 |
| K14349 neutral cholesterol ester hydrolase 1 [EC:3.1.1.-] | 0 | 0 | 1 | 1 | 0 | 0 | 0 | 0 | 6 | 4 | 10 | 12 |
| Circulatory system | 3668 | 3691 | 3846 | 3809 | 2174 | 2301 | 2108 | 2052 | 1263 | 1256 | 1169 | 1147 |
| ko04260 Cardiac muscle contraction | 3668 | 3691 | 3846 | 3809 | 2174 | 2298 | 2108 | 2052 | 1263 | 1256 | 1169 | 1146 |
| ko04261 Adrenergic signaling in cardiomyocytes | 0 | 0 | 2 | 0 | 0 | 3 | 0 | 0 | 0 | 0 | 1 | 1 |
| ko04270 Vascular smooth muscle contraction | 0 | 0 | 0 | 0 | 0 | 3 | 0 | 0 | 0 | 0 | 0 | 1 |
| Digestive system | 7127 | 7634 | 7782 | 7664 | 15544 | 15802 | 15992 | 15651 | 12931 | 12646 | 12883 | 12540 |
| ko04970 Salivary secretion | 0 | 0 | 0 | 0 | 0 | 1 | 0 | 0 | 0 | 0 | 0 | 0 |
| ko04971 Gastric acid secretion | 0 | 0 | 0 | 0 | 0 | 1 | 0 | 0 | 0 | 0 | 0 | 0 |
| ko04972 Pancreatic secretion | 0 | 6 | 2 | 2 | 0 | 2 | 0 | 0 | 2 | 1 | 7 | 8 |
| ko04976 Bile secretion | 12 | 7 | 8 | 10 | 2 | 4 | 6 | 3 | 27 | 29 | 28 | 36 |
| ko04973 Carbohydrate digestion and absorption | 3056 | 3194 | 3262 | 3327 | 4622 | 4866 | 4740 | 4750 | 5104 | 5028 | 5146 | 5167 |
| ko04974 Protein digestion and absorption | 3112 | 3443 | 3550 | 3343 | 10262 | 10368 | 10610 | 10263 | 7405 | 7158 | 7322 | 6934 |
| K01278 dipeptidyl-peptidase 4 [EC:3.4.14.5] | 3112 | 3437 | 3550 | 3343 | 10262 | 10368 | 10609 | 10261 | 7405 | 7158 | 7319 | 6931 |
| K14208 Xaa-Pro aminopeptidase 2 [EC:3.4.11.9] | 0 | 0 | 0 | 0 | 0 | 0 | 0 | 1 | 0 | 0 | 0 | 1 |
| K14210 solute carrier family 3 (neutral and basic amino acid transporter), member 1 | 0 | 0 | 0 | 0 | 0 | 0 | 1 | 1 | 0 | 0 | 0 | 2 |
| K16628 collagen, type VII, alpha | 0 | 1 | 0 | 0 | 0 | 0 | 0 | 0 | 0 | 0 | 0 | 0 |
| ko04975 Fat digestion and absorption | 0 | 1 | 0 | 2 | 1 | 1 | 2 | 0 | 2 | 2 | 3 | 8 |
| ko04979 Cholesterol metabolism | 87 | 86 | 95 | 104 | 126 | 97 | 144 | 114 | 116 | 133 | 97 | 118 |
| ko04978 Mineral absorption | 860 | 903 | 866 | 879 | 531 | 465 | 490 | 521 | 283 | 300 | 296 | 289 |
| K07213 copper chaperone | 860 | 903 | 866 | 879 | 531 | 465 | 490 | 521 | 283 | 300 | 296 | 289 |
| Excretory system | 2043 | 1923 | 2018 | 2096 | 2893 | 3054 | 3289 | 2914 | 3694 | 3538 | 3853 | 3655 |
| ko04962 Vasopressin-regulated water reabsorption | 0 | 0 | 0 | 0 | 0 | 1 | 0 | 0 | 0 | 0 | 0 | 1 |
| K12462 Rho GDP-dissociation inhibitor | 0 | 0 | 0 | 0 | 0 | 1 | 0 | 0 | 0 | 0 | 0 | 0 |
| ko04960 Aldosterone-regulated sodium reabsorption | 0 | 0 | 0 | 0 | 0 | 1 | 0 | 0 | 1 | 0 | 0 | 0 |
| ko04961 Endocrine and other factor-regulated calcium reabsorption | 0 | 0 | 0 | 0 | 0 | 1 | 0 | 0 | 0 | 0 | 0 | 0 |
| ko04964 Proximal tubule bicarbonate reclamation | 2043 | 1923 | 2018 | 2095 | 2893 | 3051 | 3289 | 2914 | 3693 | 3538 | 3853 | 3654 |
| ko04966 Collecting duct acid secretion | 0 | 0 | 0 | 1 | 0 | 1 | 0 | 0 | 0 | 0 | 0 | 0 |
| Nervous system | 27075 | 26686 | 26746 | 26510 | 28824 | 28324 | 28853 | 29209 | 22373 | 22202 | 22656 | 22066 |
| ko04724 Glutamatergic synapse | 23491 | 22949 | 23048 | 22848 | 24057 | 23576 | 24216 | 24286 | 18930 | 18842 | 19237 | 18793 |
| ko04727 GABAergic synapse | 24531 | 24020 | 24134 | 23917 | 27376 | 26908 | 27543 | 27662 | 21506 | 21326 | 21862 | 21312 |
| ko04725 Cholinergic synapse | 0 | 0 | 0 | 0 | 0 | 1 | 0 | 0 | 0 | 0 | 0 | 0 |
| ko04728 Dopaminergic synapse | 2541 | 2665 | 2610 | 2588 | 1445 | 1412 | 1307 | 1544 | 864 | 874 | 791 | 750 |
| ko04726 Serotonergic synapse | 2541 | 2665 | 2610 | 2588 | 1445 | 1410 | 1307 | 1544 | 864 | 874 | 791 | 749 |
| ko04720 Long-term potentiation | 0 | 0 | 0 | 0 | 0 | 3 | 0 | 0 | 0 | 0 | 0 | 1 |
| ko04730 Long-term depression | 0 | 0 | 0 | 0 | 0 | 1 | 0 | 0 | 0 | 0 | 0 | 0 |
| ko04723 Retrograde endocannabinoid signaling | 1 | 0 | 2 | 1 | 2 | 4 | 3 | 1 | 2 | 1 | 1 | 4 |
| ko04721 Synaptic vesicle cycle | 2 | 1 | 0 | 3 | 0 | 1 | 0 | 2 | 0 | 0 | 0 | 0 |
| K05038 solute carrier family 6 (neurotransmitter transporter, glycine) member 5/9 | 2 | 1 | 0 | 2 | 0 | 0 | 0 | 0 | 0 | 0 | 0 | 0 |
| ko04722 Neurotrophin signaling pathway | 0 | 0 | 0 | 1 | 1 | 1 | 0 | 0 | 1 | 1 | 1 | 0 |
| Sensory system | 0 | 0 | 0 | 0 | 0 | 3 | 0 | 0 | 0 | 0 | 0 | 1 |
| ko04745 Phototransduction - fly | 0 | 0 | 0 | 0 | 0 | 1 | 0 | 0 | 0 | 0 | 0 | 0 |
| ko04750 Inflammatory mediator regulation of TRP channels | 0 | 0 | 0 | 0 | 0 | 3 | 0 | 0 | 0 | 0 | 0 | 1 |
| Development and regeneration | 908 | 786 | 838 | 879 | 568 | 573 | 545 | 542 | 537 | 528 | 452 | 557 |
| ko04320 Dorso-ventral axis formation | 0 | 0 | 0 | 0 | 0 | 0 | 0 | 0 | 0 | 0 | 1 | 0 |
| ko04360 Axon guidance | 0 | 0 | 0 | 0 | 1 | 1 | 0 | 0 | 0 | 0 | 0 | 0 |
| ko04361 Axon regeneration | 908 | 786 | 836 | 879 | 567 | 572 | 545 | 542 | 537 | 527 | 451 | 557 |
| ko04380 Osteoclast differentiation | 0 | 0 | 2 | 0 | 0 | 0 | 0 | 0 | 0 | 1 | 0 | 0 |
| Aging | 33006 | 32777 | 33300 | 33166 | 31413 | 31363 | 30776 | 30146 | 30974 | 31053 | 30962 | 30479 |
| ko04211 Longevity regulating pathway | 10220 | 10031 | 9917 | 10187 | 6744 | 6597 | 6583 | 6517 | 6706 | 6656 | 6534 | 6590 |
| ko04212 Longevity regulating pathway - worm | 24384 | 24109 | 24720 | 24458 | 22093 | 22192 | 21304 | 21081 | 21443 | 21743 | 21589 | 21177 |
| ko04213 Longevity regulating pathway - multiple species | 18842 | 18699 | 18497 | 18895 | 16064 | 15768 | 16055 | 15582 | 16237 | 15966 | 15905 | 15892 |
| K03695 ATP-dependent Clp protease ATP-binding subunit ClpB | 6561 | 6640 | 6785 | 6721 | 8216 | 8134 | 8397 | 7957 | 8426 | 8282 | 8373 | 8226 |
| Environmental adaptation | 48101 | 48891 | 48185 | 48451 | 50342 | 50325 | 50690 | 51443 | 43711 | 43716 | 42953 | 43764 |
| ko04710 Circadian rhythm | 0 | 0 | 0 | 0 | 0 | 0 | 0 | 0 | 0 | 0 | 2 | 0 |
| ko04713 Circadian entrainment | 0 | 0 | 0 | 0 | 0 | 1 | 0 | 0 | 0 | 0 | 0 | 0 |
| ko04714 Thermogenesis | 17786 | 18246 | 18318 | 18105 | 22356 | 21963 | 22394 | 22413 | 17264 | 16738 | 16719 | 16712 |
| K11450 lysine-specific histone demethylase 1A [EC:1.-.-.-] | 0 | 0 | 0 | 0 | 0 | 0 | 0 | 0 | 0 | 0 | 1 | 0 |
| K18160 NADH dehydrogenase [ubiquinone] 1 alpha subcomplex assembly factor 2 | 0 | 0 | 0 | 0 | 0 | 0 | 0 | 0 | 0 | 0 | 0 | 1 |
| ko04626 Plant-pathogen interaction | 30315 | 30645 | 29867 | 30346 | 27986 | 28361 | 28296 | 29030 | 26447 | 26978 | 26234 | 27052 |
| K13408 membrane fusion protein | 716 | 677 | 699 | 736 | 296 | 308 | 270 | 242 | 1111 | 1166 | 1097 | 1167 |
| Human Diseases | 433129 | 432509 | 428098 | 432004 | 385339 | 389210 | 379746 | 382930 | 388756 | 390610 | 386542 | 389318 |
| Cancer: overview | 58098 | 57711 | 56996 | 57707 | 57992 | 58696 | 58091 | 58929 | 54959 | 53497 | 54153 | 53819 |
| ko05200 Pathways in cancer | 24479 | 24180 | 23868 | 24491 | 21025 | 20992 | 20510 | 21131 | 18509 | 18069 | 18110 | 18426 |
| ko05202 Transcriptional misregulation in cancer | 1 | 1 | 1 | 1 | 0 | 0 | 0 | 1 | 0 | 2 | 0 | 1 |
| K00069 15-hydroxyprostaglandin dehydrogenase (NAD) [EC:1.1.1.141] | 1 | 0 | 0 | 1 | 0 | 0 | 0 | 1 | 0 | 0 | 0 | 0 |
| ko05206 MicroRNAs in cancer | 4450 | 4428 | 4579 | 4488 | 8528 | 8938 | 9136 | 8783 | 8220 | 7815 | 8123 | 8010 |
| ko05205 Proteoglycans in cancer | 2269 | 2348 | 2292 | 2492 | 3098 | 3100 | 3018 | 3176 | 2862 | 2799 | 2860 | 2743 |
| ko05204 Chemical carcinogenesis | 10340 | 10167 | 9884 | 10361 | 6013 | 5970 | 5699 | 5945 | 6581 | 6308 | 6186 | 6437 |
| ko05203 Viral carcinogenesis | 7147 | 7132 | 7138 | 6837 | 6773 | 6674 | 6742 | 6952 | 7213 | 6987 | 7129 | 6912 |
| K11407 histone deacetylase 6 [EC:3.5.1.98] | 0 | 0 | 0 | 0 | 0 | 0 | 0 | 0 | 2 | 0 | 0 | 0 |
| K15979 staphylococcal nuclease domain-containing protein 1 | 0 | 0 | 2 | 0 | 0 | 0 | 0 | 0 | 0 | 0 | 0 | 0 |
| ko05230 Central carbon metabolism in cancer | 25119 | 24949 | 24698 | 24529 | 25968 | 26445 | 26612 | 26688 | 25725 | 25043 | 25647 | 24969 |
| ko05231 Choline metabolism in cancer | 929 | 804 | 860 | 901 | 645 | 647 | 550 | 544 | 593 | 593 | 526 | 611 |
| ko05235 PD-L1 expression and PD-1 checkpoint pathway in cancer | 0 | 1 | 0 | 0 | 0 | 0 | 0 | 0 | 0 | 0 | 0 | 0 |
| Cancer: specific types | 25303 | 25079 | 24709 | 25390 | 21636 | 21648 | 21092 | 21700 | 19748 | 19317 | 19222 | 19753 |
| ko05210 Colorectal cancer | 536 | 446 | 470 | 493 | 285 | 267 | 263 | 266 | 206 | 159 | 147 | 142 |
| ko05212 Pancreatic cancer | 0 | 2 | 0 | 0 | 1 | 0 | 0 | 0 | 0 | 0 | 0 | 0 |
| ko05225 Hepatocellular carcinoma | 10749 | 10616 | 10336 | 10896 | 6255 | 6142 | 5882 | 6398 | 6401 | 6103 | 6082 | 6189 |
| ko05226 Gastric cancer | 11 | 7 | 5 | 8 | 2 | 3 | 6 | 3 | 16 | 11 | 13 | 12 |
| ko05214 Glioma | 0 | 2 | 0 | 0 | 0 | 1 | 0 | 0 | 0 | 0 | 1 | 0 |
| ko05221 Acute myeloid leukemia | 0 | 1 | 0 | 0 | 0 | 0 | 0 | 0 | 0 | 0 | 1 | 0 |
| ko05220 Chronic myeloid leukemia | 0 | 1 | 0 | 0 | 0 | 1 | 0 | 0 | 0 | 0 | 1 | 0 |
| ko05218 Melanoma | 0 | 1 | 0 | 0 | 0 | 0 | 0 | 0 | 0 | 0 | 0 | 0 |
| ko05211 Renal cell carcinoma | 5167 | 5090 | 4879 | 4879 | 3119 | 3214 | 2912 | 3048 | 2480 | 2401 | 2359 | 2434 |
| ko05219 Bladder cancer | 813 | 893 | 836 | 891 | 609 | 653 | 576 | 566 | 1222 | 1237 | 1100 | 1315 |
| ko05215 Prostate cancer | 8027 | 8030 | 8183 | 8223 | 11366 | 11368 | 11453 | 11419 | 9423 | 9406 | 9525 | 9661 |

| | | | | | | | | | | | | |
|---|---|---|---|---|---|---|---|---|---|---|---|---|
| ko05213 Endometrial cancer | 0 | 0 | 0 | 0 | 0 | 0 | 0 | 0 | 1 | 0 | 1 | 0 |
| ko05224 Breast cancer | 0 | 2 | 0 | 0 | 0 | 0 | 0 | 0 | 0 | 0 | 1 | 0 |
| ko05222 Small cell lung cancer | 536 | 446 | 470 | 493 | 285 | 267 | 263 | 266 | 206 | 158 | 146 | 142 |
| ko05223 Non-small cell lung cancer | 0 | 1 | 0 | 0 | 0 | 1 | 0 | 0 | 1 | 0 | 1 | 0 |
| Immune disease | 2829 | 2939 | 2985 | 3050 | 2582 | 2557 | 2683 | 2528 | 3418 | 3756 | 3780 | 3630 |
| ko05323 Rheumatoid arthritis | 0 | 0 | 2 | 1 | 0 | 1 | 0 | 0 | 0 | 1 | 0 | 0 |
| ko05340 Primary immunodeficiency | 2829 | 2939 | 2983 | 3049 | 2582 | 2556 | 2683 | 2528 | 3418 | 3755 | 3780 | 3630 |
| Neurodegenerative disease | 21284 | 21341 | 21199 | 21477 | 15092 | 15136 | 14898 | 14720 | 13950 | 14007 | 13605 | 13904 |
| ko05010 Alzheimer disease | 10716 | 10903 | 10958 | 10933 | 8115 | 8302 | 8068 | 7943 | 6715 | 6795 | 6579 | 6748 |
| ko05012 Parkinson disease | 4209 | 4144 | 4327 | 4309 | 2465 | 2573 | 2378 | 2319 | 1484 | 1422 | 1334 | 1298 |
| ko05014 Amyotrophic lateral sclerosis (ALS) | 8059 | 7850 | 7802 | 8040 | 4462 | 4260 | 4284 | 4370 | 4856 | 4793 | 4588 | 4706 |
| K04575 amyotrophic lateral sclerosis 2 protein | 1 | 0 | 0 | 0 | 0 | 0 | 0 | 0 | 0 | 0 | 0 | 0 |
| ko05016 Huntington disease | 7601 | 7579 | 7554 | 7663 | 5490 | 5645 | 5425 | 5252 | 4589 | 4546 | 4403 | 4453 |
| ko05020 Prion diseases | 347 | 404 | 322 | 357 | 229 | 235 | 244 | 260 | 524 | 551 | 489 | 563 |
| Substance dependence | 2541 | 2665 | 2610 | 2588 | 1445 | 1412 | 1307 | 1544 | 866 | 874 | 792 | 750 |
| ko05030 Cocaine addiction | 2541 | 2665 | 2610 | 2588 | 1445 | 1409 | 1307 | 1544 | 864 | 874 | 791 | 749 |
| ko05031 Amphetamine addiction | 2541 | 2665 | 2610 | 2588 | 1445 | 1412 | 1307 | 1544 | 864 | 874 | 791 | 750 |
| ko05032 Morphine addiction | 0 | 0 | 0 | 0 | 0 | 1 | 0 | 0 | 0 | 0 | 0 | 0 |
| ko05034 Alcoholism | 2541 | 2665 | 2610 | 2588 | 1445 | 1411 | 1307 | 1544 | 866 | 874 | 792 | 750 |
| Cardiovascular disease | 25987 | 26132 | 26042 | 26290 | 26068 | 25658 | 25621 | 26356 | 23567 | 23257 | 23178 | 23344 |
| ko05418 Fluid shear stress and atherosclerosis | 25451 | 25687 | 25570 | 25797 | 25783 | 25391 | 25358 | 26090 | 23361 | 23099 | 23029 | 23202 |
| ko05410 Hypertrophic cardiomyopathy (HCM) | 0 | 0 | 2 | 0 | 0 | 0 | 0 | 0 | 0 | 0 | 3 | 0 |
| ko05412 Arrhythmogenic right ventricular cardiomyopathy (ARVC) | 0 | 0 | 2 | 0 | 0 | 0 | 0 | 0 | 0 | 0 | 1 | 0 |
| ko05414 Dilated cardiomyopathy (DCM) | 0 | 0 | 2 | 0 | 0 | 0 | 0 | 0 | 0 | 0 | 1 | 0 |
| ko05416 Viral myocarditis | 536 | 445 | 470 | 493 | 285 | 267 | 263 | 266 | 206 | 158 | 146 | 142 |
| Endocrine and metabolic disease | 36409 | 36823 | 36940 | 36118 | 39470 | 39773 | 38709 | 38709 | 35804 | 35918 | 36416 | 35662 |
| ko04930 Type II diabetes mellitus | 7811 | 7857 | 7847 | 7535 | 9322 | 9295 | 9412 | 9660 | 9167 | 8776 | 9166 | 8828 |
| ko04940 Type I diabetes mellitus | 5297 | 5240 | 5561 | 5171 | 8431 | 8576 | 8015 | 8013 | 7589 | 7551 | 7731 | 7408 |
| ko04932 Non-alcoholic fatty liver disease (NAFLD) | 4204 | 4136 | 4314 | 4303 | 2460 | 2567 | 2373 | 2318 | 1474 | 1415 | 1317 | 1295 |
| ko04931 Insulin resistance | 13929 | 14500 | 14337 | 14229 | 16139 | 16120 | 15997 | 15669 | 15085 | 15771 | 15836 | 15682 |
| ko04933 AGE-RAGE signaling pathway in diabetic complications | 1 | 0 | 1 | 0 | 1 | 1 | 0 | 1 | 3 | 0 | 0 | 3 |
| ko04934 Cushing syndrome | 5167 | 5091 | 4880 | 4880 | 3118 | 3214 | 2912 | 3048 | 2486 | 2405 | 2368 | 2446 |
| Infectious disease: bacterial | 119846 | 119132 | 117207 | 118561 | 97186 | 98763 | 93431 | 94477 | 97836 | 98567 | 97925 | 99311 |
| ko05110 Vibrio cholerae infection | 2165 | 2130 | 2117 | 2034 | 1324 | 1294 | 1182 | 1336 | 631 | 667 | 615 | 599 |
| K10954 zona occludens toxin | 2165 | 2130 | 2117 | 2033 | 1324 | 1292 | 1182 | 1336 | 630 | 667 | 615 | 599 |
| ko05120 Epithelial cell signaling in Helicobacter pylori infection | 11809 | 11476 | 11592 | 11615 | 11994 | 12219 | 12309 | 11842 | 14163 | 14596 | 14320 | 14570 |
| K03191 acid-activated urea channel | 0 | 0 | 2 | 0 | 0 | 0 | 0 | 1 | 1 | 0 | 2 | 1 |
| K08303 putative protease [EC:3.4.-.-] | 7471 | 7308 | 7728 | 7446 | 9980 | 10396 | 10485 | 9874 | 10527 | 10825 | 10809 | 10761 |
| ko05130 Pathogenic Escherichia coli infection | 0 | 0 | 0 | 0 | 1 | 1 | 0 | 0 | 0 | 0 | 0 | 0 |
| ko05132 Salmonella infection | 9131 | 9041 | 9116 | 9238 | 10716 | 11184 | 11068 | 10743 | 10668 | 10515 | 10636 | 10673 |
| K05916 nitric oxide dioxygenase [EC:1.14.12.17] | 2505 | 2462 | 2425 | 2496 | 1545 | 1580 | 1450 | 1563 | 1474 | 1608 | 1521 | 1602 |
| K12264 anaerobic nitric oxide reductase flavorubredoxin | 996 | 911 | 928 | 945 | 594 | 705 | 639 | 569 | 1269 | 1263 | 1201 | 1319 |
| K12265 nitric oxide reductase FlRd-NAD(+) reductase [EC:1.18.1.-] | 697 | 748 | 688 | 748 | 500 | 529 | 516 | 466 | 995 | 926 | 884 | 963 |
| K12266 anaerobic nitric oxide reductase transcription regulator | 1898 | 1817 | 1883 | 1921 | 1113 | 1245 | 1096 | 1227 | 1595 | 1581 | 1539 | 1613 |
| K13771 Rrf2 family transcriptional regulator, nitric oxide-sensitive transcriptional repressor | 311 | 298 | 248 | 276 | 208 | 230 | 226 | 247 | 414 | 385 | 383 | 441 |
| ko05131 Shigellosis | 0 | 0 | 0 | 0 | 1 | 1 | 0 | 0 | 7 | 6 | 3 | 16 |
| K11011 enterotoxin Set1A | 0 | 0 | 0 | 0 | 0 | 1 | 0 | 0 | 6 | 6 | 3 | 15 |
| K11012 enterotoxin Set1B | 0 | 0 | 0 | 0 | 0 | 0 | 0 | 0 | 1 | 0 | 0 | 1 |
| ko05135 Yersinia infection | 1736 | 1863 | 1731 | 1773 | 1145 | 1101 | 899 | 1005 | 1011 | 1007 | 965 | 969 |
| K13735 adhesin/invasin | 20 | 26 | 22 | 25 | 112 | 133 | 156 | 129 | 418 | 442 | 407 | 435 |
| K23475 translocator | 1062 | 1181 | 1060 | 1081 | 629 | 560 | 436 | 543 | 361 | 301 | 317 | 307 |
| K23476 translocator | 635 | 638 | 625 | 644 | 328 | 341 | 307 | 333 | 185 | 206 | 172 | 184 |
| ko05133 Pertussis | 61860 | 61731 | 59032 | 60500 | 37204 | 37618 | 34012 | 35350 | 40093 | 39986 | 39353 | 41042 |
| K07326 hemolysin activation/secretion protein | 4645 | 4473 | 4114 | 4386 | 2414 | 2311 | 2314 | 2582 | 1429 | 1361 | 1346 | 1379 |
| K07345 major type 1 subunit fimbrin (pilin) | 3088 | 3162 | 3141 | 3209 | 1823 | 1884 | 1741 | 1802 | 3340 | 3214 | 3111 | 3378 |
| K07347 outer membrane usher protein | 21757 | 21819 | 20940 | 21618 | 11494 | 11469 | 11072 | 11356 | 17377 | 17890 | 17470 | 18304 |
| K07389 cytolysin-activating lysine-acyltransferase [EC:2.3.1.-] | 12 | 16 | 23 | 22 | 3 | 2 | 1 | 3 | 49 | 67 | 56 | 85 |
| K15125 filamentous hemagglutinin | 12530 | 12223 | 11243 | 11692 | 6340 | 6318 | 3629 | 4209 | 4250 | 3865 | 3819 | 3969 |
| ko05134 Legionellosis | 16792 | 16528 | 16951 | 17054 | 19206 | 19665 | 19000 | 19207 | 17522 | 17716 | 17866 | 17391 |
| K03596 GTP-binding protein LepA | 5434 | 5432 | 5318 | 5588 | 5745 | 5835 | 5734 | 6005 | 5719 | 5834 | 5837 | 5767 |
| K07874 Ras-related protein Rab-1A | 0 | 0 | 1 | 0 | 0 | 0 | 0 | 0 | 0 | 0 | 0 | 0 |
| ko05150 Staphylococcus aureus infection | 5693 | 5531 | 5629 | 5534 | 3697 | 3689 | 3404 | 3453 | 2345 | 2470 | 2361 | 2544 |
| K11041 exfoliative toxin A/B | 123 | 129 | 131 | 134 | 33 | 39 | 30 | 24 | 128 | 148 | 122 | 157 |
| ko05152 Tuberculosis | 16444 | 16376 | 16955 | 16420 | 18304 | 18552 | 17661 | 17540 | 17026 | 17283 | 17418 | 16872 |
| ko05100 Bacterial invasion of epithelial cells | 163 | 176 | 155 | 156 | 151 | 171 | 179 | 155 | 516 | 607 | 553 | 623 |
| K13730 internalin A | 143 | 150 | 133 | 131 | 38 | 38 | 23 | 26 | 98 | 165 | 146 | 188 |
| Infectious disease: viral | 7800 | 7696 | 7715 | 7465 | 7242 | 7093 | 7209 | 7385 | 7570 | 7323 | 7407 | 7205 |
| ko05166 Human T-cell leukemia virus 1 infection | 87 | 92 | 95 | 103 | 128 | 97 | 144 | 115 | 113 | 133 | 92 | 109 |
| K08874 transformation/transcription domain-associated protein | 0 | 0 | 0 | 0 | 0 | 0 | 0 | 0 | 0 | 0 | 2 | 0 |
| K11304 histone acetyltransferase HTATIP [EC:2.3.1.48] | 0 | 2 | 0 | 0 | 0 | 0 | 0 | 0 | 0 | 0 | 0 | 0 |
| ko05170 Human immunodeficiency virus 1 infection | 536 | 446 | 470 | 494 | 285 | 268 | 263 | 266 | 206 | 159 | 146 | 142 |
| ko05162 Measles | 536 | 446 | 471 | 495 | 285 | 268 | 265 | 266 | 210 | 161 | 146 | 147 |
| ko05164 Influenza A | 557 | 464 | 482 | 520 | 346 | 319 | 321 | 315 | 252 | 209 | 183 | 182 |
| ko05161 Hepatitis B | 537 | 445 | 470 | 495 | 286 | 268 | 263 | 267 | 208 | 164 | 150 | 146 |
| ko05160 Hepatitis C | 536 | 446 | 470 | 494 | 285 | 268 | 265 | 266 | 210 | 160 | 147 | 146 |
| ko05168 Herpes simplex virus 1 infection | 544 | 455 | 475 | 499 | 286 | 270 | 267 | 268 | 212 | 165 | 151 | 152 |
| K09228 KRAB domain-containing zinc finger protein | 0 | 0 | 0 | 0 | 0 | 0 | 0 | 0 | 0 | 0 | 1 | 0 |
| ko05163 Human cytomegalovirus infection | 536 | 447 | 470 | 493 | 285 | 268 | 263 | 266 | 206 | 158 | 147 | 142 |
| ko05167 Kaposi sarcoma-associated herpesvirus infection | 536 | 447 | 470 | 493 | 285 | 267 | 263 | 266 | 206 | 158 | 146 | 142 |
| ko05169 Epstein-Barr virus infection | 536 | 447 | 470 | 494 | 285 | 267 | 263 | 266 | 207 | 159 | 147 | 142 |
| ko05165 Human papillomavirus infection | 7147 | 7133 | 7132 | 6837 | 6768 | 6675 | 6742 | 6952 | 7203 | 6975 | 7126 | 6906 |
| K12236 transcriptional repressor NF-X1 | 0 | 0 | 0 | 0 | 1 | 0 | 0 | 0 | 0 | 0 | 0 | 0 |
| Infectious disease: parasitic | 5030 | 4709 | 4864 | 4918 | 2754 | 2895 | 2836 | 2946 | 3399 | 3389 | 3225 | 3430 |
| ko05146 Amoebiasis | 340 | 367 | 352 | 346 | 212 | 226 | 250 | 212 | 661 | 651 | 628 | 681 |
| K13963 serpin B | 10 | 16 | 9 | 9 | 5 | 3 | 0 | 3 | 21 | 25 | 25 | 28 |
| ko05145 Toxoplasmosis | 536 | 445 | 471 | 494 | 285 | 267 | 263 | 266 | 207 | 159 | 146 | 143 |
| ko05140 Leishmaniasis | 0 | 0 | 0 | 1 | 0 | 0 | 0 | 0 | 0 | 1 | 0 | 0 |
| ko05142 Chagas disease (American trypanosomiasis) | 3862 | 3649 | 3788 | 3823 | 2109 | 2269 | 2185 | 2340 | 2426 | 2493 | 2357 | 2502 |
| K01354 oligopeptidase B [EC:3.4.21.83] | 3861 | 3649 | 3787 | 3822 | 2109 | 2269 | 2185 | 2339 | 2423 | 2492 | 2357 | 2499 |
| ko05143 African trypanosomiasis | 4153 | 3897 | 4040 | 4078 | 2257 | 2403 | 2323 | 2467 | 2528 | 2579 | 2451 | 2603 |
| K14475 inhibitor of cysteine peptidase | 292 | 248 | 253 | 256 | 148 | 133 | 138 | 128 | 104 | 87 | 94 | 104 |
| Drug resistance: antimicrobial | 184012 | 183965 | 182503 | 184152 | 164647 | 166555 | 163321 | 164537 | 171064 | 173124 | 169995 | 171510 |
| ko01501 beta-Lactam resistance | 102035 | 102455 | 102559 | 102669 | 94665 | 95957 | 94437 | 94402 | 97066 | 98109 | 96493 | 96791 |
| K02171 BlaI family transcriptional regulator, penicillinase repressor | 97 | 106 | 78 | 86 | 18 | 30 | 35 | 22 | 109 | 98 | 93 | 101 |
| K02172 bla regulator protein blaR1 | 591 | 533 | 503 | 572 | 135 | 133 | 131 | 124 | 538 | 651 | 579 | 630 |
| K03585 membrane fusion protein, multidrug efflux system | 8515 | 8742 | 9237 | 8856 | 13268 | 13321 | 12784 | 12883 | 10656 | 10421 | 10443 | 10230 |
| K08218 MFS transporter, PAT family, beta-lactamase induction signal transducer AmpG | 5180 | 5314 | 5286 | 5070 | 7425 | 7460 | 7782 | 7326 | 5981 | 5833 | 5979 | 5694 |
| K17837 metallo-beta-lactamase class B [EC:3.5.2.6] | 291 | 311 | 328 | 278 | 748 | 675 | 720 | 706 | 470 | 414 | 425 | 420 |
| K17838 beta-lactamase class D [EC:3.5.2.6] | 0 | 0 | 0 | 0 | 1 | 0 | 0 | 1 | 0 | 0 | 0 | 0 |
| K17850 LysR family transcriptional regulator, regulator of gene expression of beta-lactamase | 1319 | 1327 | 1274 | 1225 | 808 | 690 | 701 | 754 | 420 | 403 | 406 | 380 |
| K18129 TetR/AcrR family transcriptional regulator, mexXY operon repressor | 847 | 798 | 781 | 835 | 450 | 413 | 417 | 433 | 256 | 255 | 216 | 250 |
| K18130 TetR/AcrR family transcriptional regulator, transcriptional repressor NalC | 1008 | 1018 | 970 | 979 | 566 | 526 | 469 | 577 | 295 | 272 | 281 | 318 |
| K18131 MarR family transcriptional regulator, repressor of the mexAB-oprM multidrug resistance operon | 596 | 569 | 578 | 600 | 318 | 330 | 324 | 322 | 208 | 193 | 186 | 172 |
| K18135 TetR/AcrR family transcriptional regulator, repressor of the mexAB-oprM multidrug resistance operon | 561 | 533 | 594 | 528 | 269 | 276 | 247 | 310 | 176 | 165 | 132 | 155 |
| K18138 multidrug efflux pump | 16452 | 16175 | 16335 | 16226 | 15802 | 15639 | 15705 | 15659 | 14086 | 13988 | 13943 | 13763 |
| K18143 two-component system, OmpR family, sensor histidine kinase AdeS [EC:2.7.13.3] | 520 | 506 | 584 | 551 | 383 | 431 | 363 | 385 | 692 | 841 | 793 | 827 |
| K18144 two-component system, OmpR family, response regulator AdeR | 382 | 388 | 364 | 379 | 258 | 266 | 268 | 260 | 521 | 592 | 484 | 550 |
| K18145 membrane fusion protein, multidrug efflux system | 0 | 0 | 0 | 0 | 0 | 0 | 0 | 1 | 0 | 0 | 0 | 1 |
| K18146 multidrug efflux pump | 1558 | 1599 | 1564 | 1695 | 1131 | 1248 | 1149 | 1089 | 2260 | 2288 | 2179 | 2404 |
| K18148 release factor H-coupled RctB family protein | 1908 | 1823 | 1604 | 1842 | 1036 | 949 | 936 | 1024 | 707 | 696 | 600 | 653 |
| K18150 antirepressor for MexR | 217 | 231 | 233 | 219 | 126 | 117 | 116 | 141 | 67 | 64 | 68 | 72 |
| K18699 beta-lactamase class A SHV [EC:3.5.2.6] | 24 | 31 | 16 | 22 | 8 | 7 | 6 | 12 | 47 | 46 | 41 | 41 |
| K18767 beta-lactamase class A CTX-M [EC:3.5.2.6] | 0 | 0 | 0 | 0 | 0 | 0 | 0 | 0 | 2 | 0 | 0 | 0 |
| K18796 beta-lactamase class A LEN [EC:3.5.2.6] | 2 | 0 | 2 | 0 | 0 | 1 | 4 | 4 | 1 | 0 | 0 | 0 |
| K18973 beta-lactamase class D OXA-50 [EC:3.5.2.6] | 1345 | 1441 | 1369 | 1386 | 777 | 784 | 795 | 861 | 488 | 443 | 417 | 412 |
| K19096 beta-lactamase class C CMY-2 [EC:3.5.2.6] | 0 | 0 | 0 | 0 | 2 | 0 | 2 | 0 | 1 | 2 | 0 | 1 |
| K19215 beta-lactamase class C ACT/MIR [EC:3.5.2.6] | 0 | 0 | 0 | 0 | 0 | 0 | 0 | 0 | 1 | 0 | 0 | 0 |
| K19216 metallo-beta-lactamase class B IND [EC:3.5.2.6] | 0 | 0 | 0 | 1 | 2 | 0 | 0 | 2 | 1 | 0 | 3 | 0 |
| K20320 beta-lactamase class C PDC [EC:3.5.2.6] | 1933 | 1961 | 1829 | 1890 | 1072 | 1061 | 941 | 1141 | 663 | 612 | 555 | 576 |
| K21277 beta-lactamase class D OXA-60 [EC:3.5.2.6] | 1 | 0 | 0 | 0 | 0 | 0 | 0 | 0 | 0 | 0 | 0 | 0 |
| ko01502 Vancomycin resistance | 24717 | 24886 | 24649 | 24660 | 28824 | 29198 | 29360 | 28813 | 27169 | 27398 | 27118 | 26786 |

| | | | | | | | | | | | | |
|---|---|---|---|---|---|---|---|---|---|---|---|---|
| ko01503 Cationic antimicrobial peptide (CAMP) resistance | 88684 | 88362 | 87392 | 88438 | 78253 | 78507 | 76377 | 78495 | 77924 | 78165 | 76991 | 78186 |
| K01406 serralysin [EC:3.4.24.40] | 1684 | 1682 | 1566 | 1582 | 973 | 863 | 896 | 888 | 576 | 534 | 516 | 536 |
| K01448 N-acetylmuramoyl-L-alanine amidase [EC:3.5.1.28] | 8921 | 8949 | 8994 | 8960 | 9021 | 9132 | 8962 | 9053 | 8809 | 8877 | 8693 | 8769 |
| K03673 thiol:disulfide interchange protein DsbA | 1492 | 1669 | 1702 | 1634 | 1083 | 1021 | 1004 | 1129 | 1373 | 1488 | 1472 | 1500 |
| K06079 copper homeostasis protein (lipoprotein) | 492 | 465 | 477 | 445 | 337 | 397 | 331 | 424 | 519 | 638 | 558 | 569 |
| K12962 undecaprenyl phosphate-alpha-L-ara4N flippase subunit ArnE | 792 | 702 | 674 | 779 | 443 | 404 | 418 | 448 | 462 | 424 | 408 | 459 |
| K12963 undecaprenyl phosphate-alpha-L-ara4N flippase subunit ArnF | 466 | 423 | 344 | 453 | 297 | 276 | 262 | 299 | 374 | 387 | 397 | 404 |
| K13632 AraC family transcriptional regulator, mar-sox-rob regulon activator | 194 | 175 | 199 | 177 | 163 | 146 | 125 | 155 | 289 | 301 | 264 | 253 |
| K19236 L,D-transpeptidase YcfS | 522 | 519 | 506 | 488 | 311 | 389 | 396 | 373 | 811 | 723 | 734 | 739 |
| K19238 signal transduction protein PmrD | 173 | 113 | 168 | 149 | 101 | 101 | 110 | 94 | 213 | 217 | 225 | 176 |
| Drug resistance: antineoplastic | 39097 | 38159 | 37889 | 38521 | 32773 | 32581 | 32259 | 32942 | 31499 | 30544 | 30126 | 30545 |
| ko01521 EGFR tyrosine kinase inhibitor resistance | 2 | 1 | 0 | 0 | 0 | 2 | 0 | 0 | 0 | 4 | 1 | 4 |
| ko01524 Platinum drug resistance | 17388 | 16450 | 16525 | 17194 | 9668 | 9634 | 9148 | 9739 | 10322 | 10424 | 10019 | 10432 |
| ko01523 Antifolate resistance | 21707 | 21708 | 21364 | 21327 | 23105 | 22946 | 23111 | 23203 | 21178 | 20117 | 20106 | 20109 |
| ko01522 Endocrine resistance | 0 | 1 | 0 | 0 | 0 | 0 | 0 | 0 | 0 | 0 | 1 | 0 |
| Brite Hierarchies | 5418349 | 5398942 | 5339635 | 5401726 | 4741005 | 4780733 | 4715107 | 4749119 | 4925928 | 4956758 | 4890240 | 4953017 |
| Protein families: metabolism | 1096552 | 1092366 | 1082616 | 1095412 | 995638 | 1004854 | 990434 | 993473 | 995825 | 1000562 | 989552 | 999807 |
| ko01001 Protein kinases | 152919 | 150861 | 146140 | 150476 | 94896 | 96148 | 93043 | 95912 | 109312 | 110809 | 107760 | 111659 |
| K00936 two-component system, sensor histidine kinase PdtaS [EC:2.7.13.3] | 132 | 137 | 125 | 126 | 37 | 31 | 35 | 26 | 168 | 201 | 183 | 176 |
| K02476 two-component system, CitB family, sensor kinase [EC:2.7.13.3] | 87 | 77 | 71 | 74 | 12 | 20 | 11 | 14 | 93 | 101 | 81 | 85 |
| K02478 two-component system, LytTR family, sensor kinase [EC:2.7.13.3] | 2011 | 1853 | 1857 | 1941 | 1283 | 1324 | 1340 | 1293 | 2767 | 2768 | 2596 | 2731 |
| K02480 two-component system, NarL family, sensor kinase [EC:2.7.13.3] | 3 | 0 | 3 | 3 | 0 | 3 | 0 | 1 | 1 | 11 | 2 | 2 |
| K02482 two-component system, NtrC family, sensor kinase [EC:2.7.13.3] | 1771 | 1853 | 1886 | 1852 | 994 | 1065 | 983 | 1013 | 621 | 611 | 650 | 649 |
| K02484 two-component system, OmpR family, sensor kinase [EC:2.7.13.3] | 1004 | 1054 | 1149 | 1077 | 3603 | 3500 | 3623 | 3438 | 2652 | 2681 | 2773 | 2671 |
| K02486 two-component system, sensor kinase [EC:2.7.13.3] | 3 | 2 | 0 | 1 | 0 | 0 | 0 | 0 | 0 | 0 | 1 | 1 |
| K04757 serine/threonine-protein kinase RsbW [EC:2.7.11.1] | 136 | 106 | 136 | 149 | 39 | 43 | 57 | 36 | 242 | 251 | 261 | 270 |
| K07154 serine/threonine-protein kinase HipA [EC:2.7.11.1] | 1927 | 2100 | 2118 | 2082 | 5065 | 5138 | 5352 | 5097 | 4650 | 4574 | 4782 | 4675 |
| K08857 NIMA (never in mitosis gene a)-related kinase 1/4/5 [EC:2.7.11.1] | 0 | 0 | 0 | 0 | 0 | 1 | 0 | 0 | 0 | 0 | 0 | 0 |
| K08869 aarF domain-containing kinase | 0 | 0 | 0 | 0 | 1 | 0 | 0 | 0 | 0 | 0 | 0 | 0 |
| K08884 serine/threonine protein kinase, bacterial [EC:2.7.11.1] | 49 | 38 | 46 | 39 | 14 | 9 | 7 | 10 | 123 | 105 | 114 | 149 |
| K11527 two-component system, sensor histidine kinase and response regulator [EC:2.7.13.3] | 688 | 736 | 834 | 687 | 2331 | 2388 | 2448 | 2186 | 1672 | 1561 | 1782 | 1513 |
| K12132 eukaryotic-like serine/threonine-protein kinase [EC:2.7.11.1] | 1372 | 1307 | 1454 | 1366 | 732 | 856 | 773 | 668 | 1051 | 1149 | 1051 | 1170 |
| K15904 bifunctional N6-L-threonylcarbamoyladenine synthase / protein kinase Bud32 [EC:2.3.1.234 2.7.11.1] | 0 | 0 | 5 | 1 | 2 | 4 | 5 | 2 | 3 | 3 | 8 | 10 |
| K16309 serine/threonine-protein kinase greatwall [EC:2.7.11.1] | 0 | 0 | 2 | 0 | 0 | 0 | 0 | 0 | 4 | 2 | 2 | 1 |
| K17752 serine/threonine-protein kinase RsbT [EC:2.7.11.1] | 1 | 0 | 0 | 0 | 0 | 0 | 0 | 0 | 0 | 0 | 0 | 0 |
| K19852 serine/threonine protein kinase KIN1/2 [EC:2.7.11.1] | 0 | 0 | 0 | 0 | 0 | 1 | 0 | 0 | 0 | 0 | 0 | 0 |
| ko01009 Protein phosphatases and associated proteins | 15991 | 16129 | 16357 | 16336 | 22340 | 22960 | 22565 | 22176 | 20202 | 20375 | 20321 | 20436 |
| K15732 RNA polymerase II subunit A C-terminal domain phosphatase [EC:3.1.3.16] | 0 | 0 | 0 | 0 | 0 | 0 | 0 | 0 | 0 | 0 | 1 | 0 |
| K17623 pseudouridine 5'-phosphatase [EC:3.1.3.96] | 0 | 0 | 2 | 0 | 0 | 1 | 0 | 0 | 2 | 3 | 3 | 1 |
| K20074 PPM family protein phosphatase [EC:3.1.3.16] | 643 | 549 | 621 | 629 | 366 | 376 | 383 | 302 | 419 | 523 | 432 | 464 |
| K21814 serine/threonine-protein phosphatase CPPED1 [EC:3.1.3.16] | 3 | 1 | 1 | 0 | 1 | 0 | 0 | 2 | 0 | 0 | 3 | 0 |
| ko01002 Peptidases | 281229 | 280455 | 280046 | 282141 | 280291 | 281848 | 280342 | 278403 | 278351 | 278636 | 277235 | 277455 |
| K00683 glutaminyl-peptide cyclotransferase [EC:2.3.2.5] | 0 | 1 | 0 | 0 | 0 | 0 | 0 | 0 | 0 | 0 | 0 | 0 |
| K01258 tripeptide aminopeptidase [EC:3.4.11.4] | 2499 | 2497 | 2450 | 2486 | 3425 | 3429 | 3393 | 3298 | 3409 | 3442 | 3514 | 3493 |
| K01261 glutamyl aminopeptidase [EC:3.4.11.7] | 70 | 76 | 74 | 70 | 26 | 25 | 16 | 20 | 73 | 108 | 87 | 61 |
| K01262 Xaa-Pro aminopeptidase [EC:3.4.11.9] | 6820 | 7053 | 7095 | 7041 | 10081 | 10301 | 10219 | 10033 | 8131 | 8448 | 8328 | 8038 |
| K01265 methionyl aminopeptidase [EC:3.4.11.18] | 4600 | 4762 | 4781 | 5020 | 5587 | 5587 | 5539 | 5544 | 5322 | 5500 | 5452 | 5382 |
| K01266 D-aminopeptidase [EC:3.4.11.19] | 2205 | 2118 | 2051 | 2071 | 941 | 852 | 815 | 988 | 984 | 928 | 953 | 1052 |
| K01267 aspartyl aminopeptidase [EC:3.4.11.21] | 2173 | 2142 | 2113 | 2248 | 999 | 919 | 928 | 955 | 1107 | 1066 | 1151 | 1042 |
| K01268 aminopeptidase I [EC:3.4.11.22] | 0 | 0 | 2 | 0 | 0 | 0 | 0 | 0 | 0 | 0 | 0 | 0 |
| K01271 Xaa-Pro dipeptidase [EC:3.4.13.9] | 1815 | 1787 | 1660 | 1755 | 974 | 984 | 1030 | 909 | 1597 | 1885 | 1646 | 1640 |
| K01273 membrane dipeptidase [EC:3.4.13.19] | 2581 | 2349 | 2467 | 2544 | 1111 | 1233 | 1118 | 1135 | 1729 | 1838 | 1638 | 1805 |
| K01277 dipeptidyl-peptidase III [EC:3.4.14.4] | 1280 | 1431 | 1481 | 1341 | 4377 | 4244 | 4422 | 4242 | 3024 | 2818 | 3128 | 2955 |
| K01281 X-Pro dipeptidyl-peptidase [EC:3.4.14.11] | 4 | 2 | 2 | 5 | 3 | 12 | 3 | 3 | 4 | 2 | 5 | 3 |
| K01284 peptidyl-dipeptidase Dcp [EC:3.4.15.5] | 3812 | 4056 | 4237 | 4133 | 10238 | 9931 | 10208 | 10020 | 8067 | 7774 | 7922 | 7596 |
| K01293 Gly-Xaa carboxypeptidase [EC:3.4.17.4] | 0 | 0 | 0 | 0 | 2 | 0 | 0 | 1 | 2 | 2 | 2 | 2 |
| K01295 glutamate carboxypeptidase [EC:3.4.17.11] | 2183 | 2282 | 2121 | 2304 | 1588 | 1468 | 1385 | 1435 | 978 | 952 | 870 | 960 |
| K01297 muramoyltetrapeptide carboxypeptidase [EC:3.4.17.13] | 2724 | 2711 | 2622 | 2706 | 3036 | 3001 | 2891 | 2989 | 2721 | 2585 | 2753 | 2657 |
| K01299 carboxypeptidase Taq [EC:3.4.17.19] | 1355 | 1364 | 1301 | 1289 | 580 | 646 | 640 | 610 | 1509 | 1579 | 1616 | 1653 |
| K01303 acylaminoacyl-peptidase [EC:3.4.19.1] | 3 | 2 | 0 | 1 | 0 | 0 | 0 | 0 | 3 | 3 | 3 | 3 |
| K01304 pyroglutamyl-peptidase [EC:3.4.19.3] | 325 | 303 | 308 | 312 | 109 | 103 | 133 | 122 | 611 | 637 | 554 | 630 |
| K01305 beta-aspartyl-dipeptidase (metallo-type) [EC:3.4.19.-] | 205 | 201 | 220 | 171 | 194 | 211 | 231 | 214 | 332 | 376 | 374 | 358 |
| K01308 g-D-glutamyl-meso-diaminopimelate peptidase [EC:3.4.19.11] | 192 | 158 | 178 | 175 | 26 | 27 | 23 | 22 | 146 | 191 | 171 | 177 |
| K01337 lysyl endopeptidase [EC:3.4.21.50] | 2300 | 2223 | 1934 | 2176 | 1256 | 1150 | 1180 | 1302 | 761 | 693 | 643 | 676 |
| K01342 subtilisin [EC:3.4.21.62] | 0 | 0 | 0 | 1 | 2 | 1 | 1 | 2 | 3 | 2 | 0 | 0 |
| K01355 omptin [EC:3.4.23.49] | 7 | 5 | 3 | 2 | 3 | 0 | 12 | 0 | 99 | 80 | 103 | 99 |
| K01356 repressor LexA [EC:3.4.21.88] | 2122 | 2065 | 2086 | 2199 | 937 | 968 | 882 | 933 | 4131 | 4493 | 4364 | 4183 |
| K01361 lactocepin [EC:3.4.21.96] | 1 | 0 | 2 | 1 | 46 | 37 | 31 | 29 | 2 | 0 | 1 | 3 |
| K01372 bleomycin hydrolase [EC:3.4.22.40] | 1630 | 1700 | 1912 | 1721 | 5149 | 5117 | 5161 | 4992 | 3613 | 3709 | 3770 | 3509 |
| K01387 microbial collagenase [EC:3.4.24.3] | 2 | 2 | 4 | 0 | 0 | 2 | 0 | 0 | 0 | 0 | 0 | 1 |
| K01400 bacillolysin [EC:3.4.24.28] | 0 | 2 | 2 | 0 | 3 | 2 | 0 | 0 | 2 | 0 | 2 | 0 |
| K01407 protease III [EC:3.4.24.55] | 1799 | 1776 | 1758 | 1826 | 1263 | 1314 | 1298 | 1308 | 2398 | 2473 | 2408 | 2555 |
| K01414 oligopeptidase A [EC:3.4.24.70] | 3793 | 3749 | 3609 | 3777 | 2339 | 2380 | 2282 | 2412 | 2827 | 2680 | 2780 | 2850 |
| K01419 ATP-dependent HslUV protease, peptidase subunit HslV [EC:3.4.25.2] | 1209 | 1214 | 1198 | 1261 | 710 | 740 | 668 | 664 | 796 | 832 | 799 | 770 |
| K01423 beta-barrel assembly-enhancing protease [EC:3.4.-.-] | 1117 | 1147 | 1082 | 1200 | 845 | 875 | 829 | 873 | 1072 | 1077 | 1102 | 1148 |
| K01436 amidohydrolase [EC:3.5.1.-] | 67 | 57 | 57 | 55 | 31 | 34 | 30 | 27 | 36 | 56 | 49 | 58 |
| K02278 prepilin peptidase CpaA [EC:3.4.23.43] | 794 | 789 | 749 | 819 | 422 | 352 | 354 | 429 | 315 | 315 | 280 | 323 |
| K02506 leader peptidase HopD [EC:3.4.23.43] | 344 | 329 | 294 | 368 | 198 | 217 | 274 | 193 | 490 | 458 | 465 | 472 |
| K02654 leader peptidase (prepilin peptidase) / N-methyltransferase [EC:3.4.23.43 2.1.1.-] | 1657 | 1600 | 1687 | 1618 | 928 | 870 | 802 | 772 | 782 | 793 | 770 | 830 |
| K03152 protein deglycase [EC:3.5.1.124] | 1006 | 1040 | 1095 | 1081 | 1443 | 1372 | 1442 | 1468 | 1758 | 1645 | 1604 | 1688 |
| K03503 DNA polymerase V [EC:3.4.21.-] | 777 | 776 | 717 | 754 | 839 | 828 | 873 | 803 | 443 | 457 | 473 | 453 |
| K03568 TldD protein | 2812 | 2709 | 2586 | 2765 | 1562 | 1627 | 1647 | 1739 | 1859 | 1962 | 1776 | 1968 |
| K03592 PmbA protein | 2257 | 2097 | 2137 | 2134 | 1273 | 1341 | 1461 | 1273 | 1801 | 1655 | 1593 | 1751 |
| K03605 hydrogenase maturation protease [EC:3.4.23.-] | 160 | 146 | 132 | 161 | 216 | 266 | 241 | 204 | 123 | 105 | 101 | 107 |
| K03797 carboxyl-terminal processing protease [EC:3.4.21.102] | 10362 | 10318 | 10581 | 10739 | 15231 | 15344 | 15241 | 14813 | 12139 | 11830 | 12439 | 11888 |
| K03798 cell division protease FtsH [EC:3.4.24.-] | 7788 | 7625 | 8090 | 7725 | 7320 | 7558 | 7310 | 6910 | 8155 | 8566 | 8405 | 8347 |
| K03799 heat shock protein HtpX [EC:3.4.24.-] | 1932 | 1906 | 1866 | 1996 | 2826 | 2657 | 2691 | 2523 | 2365 | 2311 | 2436 | 2342 |
| K04076 ATP-dependent Lon protease [EC:3.4.21.53] | 6 | 1 | 3 | 3 | 2 | 3 | 1 | 0 | 10 | 16 | 19 | 14 |
| K04691 serine protease DegS [EC:3.4.21.-] | 2563 | 2511 | 2407 | 2530 | 1527 | 1551 | 1497 | 1434 | 1497 | 1601 | 1489 | 1543 |
| K04770 Lon-like ATP-dependent protease [EC:3.4.21.-] | 960 | 994 | 886 | 1025 | 696 | 752 | 709 | 675 | 1367 | 1431 | 1364 | 1427 |
| K04772 serine protease DegQ [EC:3.4.21.-] | 898 | 908 | 896 | 996 | 557 | 748 | 584 | 656 | 1320 | 1255 | 1277 | 1285 |
| K04773 protease IV [EC:3.4.21.-] | 2343 | 2493 | 2529 | 2537 | 4719 | 4893 | 4974 | 4673 | 4443 | 4195 | 4303 | 4345 |
| K04774 serine protease SohB [EC:3.4.21.-] | 1739 | 1727 | 1810 | 1757 | 988 | 1051 | 1047 | 1104 | 1009 | 1067 | 962 | 1068 |
| K04775 protease YdgD [EC:3.4.21.-] | 426 | 452 | 523 | 548 | 278 | 367 | 321 | 320 | 655 | 687 | 656 | 634 |
| K05520 protease I [EC:3.5.1.124] | 3325 | 3317 | 3206 | 3126 | 1795 | 1778 | 1762 | 1856 | 1793 | 1858 | 1688 | 1838 |
| K05995 dipeptidase E [EC:3.4.13.21] | 245 | 282 | 261 | 261 | 127 | 120 | 103 | 117 | 655 | 688 | 620 | 668 |
| K05998 pseudomonalisin [EC:3.4.21.100] | 0 | 0 | 0 | 0 | 0 | 1 | 0 | 0 | 0 | 0 | 0 | 0 |
| K06012 spore protease [EC:3.4.24.78] | 522 | 512 | 516 | 473 | 178 | 132 | 145 | 140 | 361 | 377 | 360 | 366 |
| K06336 spore coat-associated protein N | 14 | 12 | 11 | 16 | 5 | 5 | 5 | 10 | 21 | 15 | 27 | 18 |
| K06383 stage II sporulation protein GA (sporulation sigma-E factor processing peptidase) [EC:3.4.23.-] | 255 | 256 | 264 | 247 | 93 | 77 | 93 | 62 | 221 | 221 | 217 | 227 |
| K06399 stage IV sporulation protein B [EC:3.4.21.116] | 460 | 501 | 521 | 502 | 144 | 134 | 133 | 131 | 434 | 462 | 466 | 418 |
| K06402 stage IV sporulation protein FB [EC:3.4.24.-] | 1 | 2 | 0 | 2 | 2 | 1 | 0 | 1 | 3 | 6 | 7 | 7 |
| K06972 presequence protease [EC:3.4.24.-] | 1432 | 1431 | 1487 | 1523 | 322 | 349 | 336 | 356 | 1362 | 1408 | 1413 | 1327 |
| K06974 archaemetzincin [EC:3.4.-.-] | 7 | 5 | 12 | 9 | 34 | 30 | 34 | 33 | 20 | 29 | 19 | 32 |
| K07010 putative glutamine amidotransferase | 1625 | 1647 | 1621 | 1727 | 771 | 777 | 850 | 897 | 719 | 718 | 752 | 745 |
| K07177 Lon-like protease | 58 | 56 | 69 | 77 | 15 | 16 | 19 | 21 | 68 | 59 | 61 | 75 |
| K07261 penicillin-insensitive murein DD-endopeptidase [EC:3.4.24.-] | 525 | 535 | 449 | 560 | 320 | 324 | 374 | 323 | 725 | 644 | 637 | 754 |
| K07262 serine-type D-Ala-D-Ala endopeptidase (penicillin-binding protein 7) [EC:3.4.21.-] | 1929 | 1981 | 1959 | 1940 | 1148 | 1242 | 1197 | 1245 | 1207 | 1224 | 1228 | 1228 |
| K07263 zinc protease [EC:3.4.24.-] | 6389 | 6457 | 6391 | 6431 | 8190 | 8264 | 8277 | 8382 | 6039 | 5494 | 5684 | 5420 |
| K07284 sortase A [EC:3.4.22.70] | 239 | 220 | 234 | 270 | 76 | 142 | 88 | 101 | 356 | 440 | 352 | 402 |
| K07386 putative endopeptidase [EC:3.4.24.-] | 1875 | 2064 | 2107 | 1953 | 5104 | 4971 | 5077 | 5037 | 4074 | 3876 | 3854 | 3675 |
| K07387 metalloprotease [EC:3.4.24.-] | 1875 | 1812 | 1737 | 1753 | 2475 | 2545 | 2418 | 2552 | 2068 | 2023 | 1947 | 1970 |
| K07991 archaeal preflagellin peptidase FlaK [EC:3.4.23.52] | 0 | 0 | 0 | 0 | 0 | 2 | 2 | 0 | 0 | 3 | 3 | 4 |
| K08258 staphopain A [EC:3.4.22.48] | 87 | 60 | 77 | 57 | 3 | 4 | 1 | 7 | 66 | 100 | 59 | 82 |
| K08315 hydrogenase 3 maturation protease [EC:3.4.23.51] | 312 | 307 | 266 | 295 | 182 | 195 | 196 | 223 | 399 | 401 | 420 | 447 |
| K08326 aminopeptidase [EC:3.4.11.-] | 9 | 5 | 6 | 8 | 6 | 12 | 15 | 9 | 27 | 39 | 39 | 48 |
| K08566 plasminogen activator [EC:3.4.23.48] | 0 | 0 | 0 | 0 | 0 | 0 | 2 | 0 | 0 | 2 | 0 | 4 |
| K08600 sortase B [EC:3.4.22.71] | 625 | 623 | 645 | 666 | 196 | 188 | 234 | 174 | 663 | 712 | 681 | 750 |
| K08602 oligoendopeptidase F [EC:3.4.24.-] | 601 | 636 | 703 | 641 | 219 | 265 | 247 | 215 | 608 | 652 | 620 | 607 |
| K08640 zinc D-Ala-D-Ala carboxypeptidase [EC:3.4.17.14] | 23 | 20 | 12 | 34 | 31 | 28 | 37 | 18 | 61 | 39 | 72 | 71 |
| K08651 thermitase [EC:3.4.21.66] | 3 | 2 | 0 | 2 | 2 | 5 | 3 | 5 | 0 | 0 | 0 | 2 |

| | | | | | | | | | | | | |
|---|---|---|---|---|---|---|---|---|---|---|---|---|
| K08659 dipeptidase [EC:3.4.-.-] | 78 | 86 | 97 | 70 | 111 | 149 | 146 | 112 | 98 | 105 | 115 | 101 |
| K08675 ATP-dependent Lon protease [EC:3.4.21.53] | 1 | 1 | 2 | 0 | 0 | 0 | 0 | 3 | 0 | 1 | 0 | 1 |
| K08676 tricorn protease [EC:3.4.21.-] | 432 | 429 | 472 | 380 | 594 | 614 | 591 | 607 | 39 | 82 | 76 | 98 |
| K08956 AFG3 family protein [EC:3.4.24.-] | 0 | 0 | 1 | 1 | 0 | 0 | 1 | 0 | 1 | 0 | 1 | 0 |
| K09612 alkaline phosphatase isozyme conversion protein [EC:3.4.11.-] | 654 | 690 | 565 | 665 | 465 | 463 | 486 | 424 | 838 | 893 | 842 | 877 |
| K09933 MtfA peptidase | 1509 | 1478 | 1471 | 1503 | 858 | 940 | 830 | 934 | 1024 | 1037 | 1000 | 960 |
| K11873 ubiquitin carboxyl-terminal hydrolase 10 [EC:3.4.19.12] | 0 | 0 | 0 | 0 | 0 | 0 | 0 | 0 | 1 | 0 | 0 | 0 |
| K12940 aminobenzoyl-glutamate utilization protein A | 943 | 959 | 980 | 936 | 529 | 600 | 494 | 577 | 1501 | 1484 | 1403 | 1592 |
| K12941 aminobenzoyl-glutamate utilization protein B | 920 | 955 | 853 | 936 | 260 | 320 | 257 | 301 | 1455 | 1445 | 1389 | 1514 |
| K13049 carboxypeptidase PM20D1 [EC:3.4.17.-] | 391 | 380 | 420 | 385 | 123 | 135 | 105 | 133 | 755 | 730 | 702 | 782 |
| K13274 cell wall-associated protease [EC:3.4.21.-] | 0 | 0 | 1 | 0 | 0 | 0 | 0 | 0 | 1 | 0 | 2 | 0 |
| K13275 major intracellular serine protease [EC:3.4.21.-] | 1 | 0 | 1 | 0 | 0 | 2 | 1 | 0 | 0 | 0 | 0 | 0 |
| K13277 minor extracellular protease Epr [EC:3.4.21.-] | 1 | 1 | 0 | 1 | 0 | 0 | 0 | 1 | 3 | 1 | 2 | 3 |
| K13282 cyanophycinase [EC:3.4.15.6] | 2 | 0 | 0 | 4 | 0 | 0 | 0 | 0 | 2 | 3 | 4 | 9 |
| K13617 protein phosphatase methylesterase 1 [EC:3.1.1.89] | 0 | 1 | 0 | 0 | 0 | 0 | 0 | 0 | 0 | 0 | 0 | 0 |
| K13694 murein DD-endopeptidase / murein LD-carboxypeptidase [EC:3.4.-.- 3.4.17.13] | 626 | 682 | 669 | 680 | 1330 | 1339 | 1365 | 1374 | 1176 | 1239 | 1166 | 1212 |
| K13695 probable lipoprotein NlpC | 306 | 264 | 250 | 325 | 169 | 226 | 215 | 220 | 454 | 422 | 435 | 474 |
| K14358 secernin | 33 | 26 | 30 | 27 | 18 | 29 | 22 | 5 | 30 | 24 | 49 | 31 |
| K14664 IAA-amino acid hydrolase [EC:3.5.1.-] | 50 | 49 | 37 | 37 | 15 | 18 | 14 | 10 | 29 | 35 | 45 | 40 |
| K14665 amidohydrolase [EC:3.5.1.-] | 99 | 105 | 107 | 96 | 66 | 76 | 79 | 100 | 150 | 130 | 118 | 160 |
| K16203 D-amino peptidase [EC:3.4.11.-] | 307 | 278 | 286 | 296 | 38 | 42 | 52 | 64 | 279 | 295 | 254 | 285 |
| K16291 L,D-transpeptidase ErfK/SrfK | 1765 | 1865 | 1835 | 1934 | 1045 | 1039 | 1088 | 1173 | 1196 | 1150 | 1097 | 1200 |
| K16922 putative peptide zinc metalloprotease protein | 464 | 402 | 417 | 446 | 713 | 704 | 673 | 661 | 44 | 63 | 60 | 66 |
| K17733 peptidoglycan LD-endopeptidase CwlK [EC:3.4.-.-] | 334 | 319 | 389 | 343 | 32 | 18 | 28 | 36 | 53 | 54 | 44 | 48 |
| K17734 serine protease AprX [EC:3.4.21.-] | 892 | 797 | 761 | 819 | 103 | 107 | 109 | 103 | 529 | 629 | 516 | 618 |
| K18015 deubiquitinase [EC:3.4.22.-] | 0 | 0 | 0 | 0 | 8 | 6 | 7 | 4 | 0 | 1 | 3 | 1 |
| K18530 putative aminopeptidase FrvX [EC:3.4.11.-] | 0 | 14 | 7 | 0 | 6 | 5 | 0 | 0 | 0 | 0 | 13 | 0 |
| K18545 streptogrisin B [EC:3.4.21.81] | 1 | 0 | 0 | 0 | 0 | 0 | 0 | 0 | 0 | 0 | 0 | 0 |
| K18574 Xaa-Xaa-Pro tripeptidyl-peptidase [EC:3.4.14.12] | 0 | 0 | 0 | 0 | 0 | 0 | 0 | 0 | 0 | 0 | 0 | 1 |
| K18988 serine-type D-Ala-D-Ala carboxypeptidase/endopeptidase [EC:3.4.16.4 3.4.21.-] | 667 | 674 | 636 | 719 | 467 | 538 | 521 | 514 | 873 | 937 | 834 | 1066 |
| K19223 peptidoglycan DL-endopeptidase LytF [EC:3.4.-.-] | 0 | 0 | 0 | 0 | 0 | 0 | 0 | 0 | 0 | 1 | 0 | 0 |
| K19224 peptidoglycan DL-endopeptidase LytE [EC:3.4.-.-] | 2 | 1 | 0 | 3 | 0 | 0 | 0 | 0 | 0 | 0 | 0 | 0 |
| K19225 rhomboid protease GluP [EC:3.4.21.105] | 353 | 338 | 399 | 368 | 76 | 61 | 147 | 107 | 287 | 300 | 320 | 298 |
| K19234 L,D-transpeptidase YnhG | 508 | 548 | 466 | 562 | 351 | 389 | 342 | 351 | 715 | 748 | 827 | 736 |
| K19235 L,D-transpeptidase YbiS | 580 | 550 | 585 | 605 | 365 | 413 | 410 | 378 | 804 | 916 | 820 | 888 |
| K19303 murein DD-endopeptidase [EC:3.4.-.-] | 369 | 352 | 381 | 376 | 247 | 283 | 252 | 256 | 505 | 494 | 484 | 516 |
| K19304 murein DD-endopeptidase [EC:3.4.24.-] | 764 | 809 | 732 | 753 | 484 | 502 | 565 | 436 | 1041 | 1053 | 1015 | 1025 |
| K19689 aminopeptidase [EC:3.4.11.-] | 220 | 221 | 231 | 227 | 118 | 120 | 109 | 115 | 369 | 397 | 361 | 350 |
| K19701 aminopeptidase YwaD [EC:3.4.11.6 3.4.11.10] | 4 | 3 | 2 | 1 | 0 | 0 | 2 | 0 | 4 | 9 | 2 | 4 |
| K20609 putative aminopeptidase [EC:3.4.11.-] | 7 | 3 | 3 | 1 | 30 | 20 | 18 | 10 | 13 | 17 | 14 | 12 |
| K20742 gamma-D-glutamyl-L-lysine dipeptidyl-peptidase [EC:3.4.14.13] | 8 | 3 | 1 | 4 | 4 | 1 | 1 | 0 | 0 | 1 | 5 | 1 |
| K21429 immunomodulating metalloprotease | 4119 | 4025 | 4030 | 4143 | 2436 | 2246 | 2231 | 2494 | 1320 | 1279 | 1297 | 1231 |
| K21470 L,D-transpeptidase YcbB | 1717 | 1737 | 1747 | 1725 | 3271 | 3384 | 3089 | 3146 | 3391 | 3151 | 3376 | 3247 |
| K21471 peptidoglycan DL-endopeptidase CwlO [EC:3.4.-.-] | 867 | 841 | 961 | 899 | 204 | 206 | 199 | 146 | 826 | 922 | 839 | 940 |
| K21472 peptidoglycan LD-endopeptidase LytH [EC:3.4.-.-] | 1764 | 1799 | 1830 | 1889 | 715 | 731 | 708 | 652 | 994 | 1137 | 1006 | 970 |
| K21473 peptidoglycan DL-endopeptidase RipA [EC:3.4.-.-] | 45 | 37 | 51 | 36 | 7 | 5 | 12 | 6 | 19 | 16 | 18 | 20 |
| K21604 jasmonoyl-L-amino acid hydrolase [EC:3.5.1.127] | 35 | 21 | 20 | 19 | 7 | 4 | 2 | 7 | 10 | 10 | 20 | 14 |
| K21613 uncharacterized hydrolase [EC:3.-.-.-] | 651 | 676 | 666 | 657 | 403 | 457 | 462 | 392 | 873 | 863 | 914 | 941 |
| K22684 metacaspase-1 [EC:3.4.22.-] | 5 | 5 | 3 | 5 | 21 | 17 | 16 | 10 | 13 | 16 | 10 | 7 |
| ko01003 Glycosyltransferases | 81413 | 80648 | 79395 | 80901 | 66846 | 67819 | 66539 | 67375 | 64413 | 65616 | 63893 | 65524 |
| K00712 poly(glycerol-phosphate) alpha-glucosyltransferase [EC:2.4.1.52] | 1 | 4 | 3 | 2 | 14 | 17 | 5 | 12 | 4 | 5 | 7 | 4 |
| K00752 hyaluronan synthase [EC:2.4.1.212] | 3 | 5 | 2 | 5 | 0 | 0 | 0 | 0 | 8 | 12 | 8 | 17 |
| K02852 UDP-N-acetyl-D-mannosaminouronate:lipid I N-acetyl-D-mannosaminouronosyltransferase [EC:2.4.1.180] | 560 | 511 | 479 | 515 | 365 | 373 | 357 | 363 | 635 | 726 | 589 | 642 |
| K02853 enterobacterial common antigen polymerase [EC:2.4.1.-] | 809 | 899 | 795 | 846 | 605 | 632 | 657 | 614 | 1127 | 1164 | 1181 | 1109 |
| K03669 membrane glycosyltransferase [EC:2.4.1.-] | 5774 | 5722 | 5474 | 5489 | 3213 | 3386 | 3125 | 3361 | 3286 | 3310 | 3247 | 3280 |
| K05841 sterol 3beta-glucosyltransferase [EC:2.4.1.173] | 0 | 0 | 0 | 1 | 0 | 0 | 0 | 1 | 0 | 0 | 0 | 0 |
| K07272 rhamnosyltransferase [EC:2.4.1.-] | 0 | 1 | 11 | 7 | 2 | 5 | 4 | 2 | 4 | 15 | 8 | 7 |
| K12246 poly-alpha-2,8 sialosyl sialyltransferase [EC:2.4.-.-] | 2 | 2 | 0 | 3 | 1 | 2 | 5 | 2 | 42 | 46 | 47 | 44 |
| K12582 dTDP-N-acetylfucosamine:lipid II N-acetylfucosaminyltransferase [EC:2.4.1.325] | 708 | 674 | 674 | 725 | 513 | 549 | 543 | 522 | 952 | 1073 | 1006 | 938 |
| K12984 (heptosyl)LPS beta-1,4-glucosyltransferase [EC:2.4.1.-] | 457 | 513 | 487 | 553 | 316 | 328 | 323 | 307 | 576 | 681 | 585 | 713 |
| K12987 alpha-1,6-rhamnosyltransferase [EC:2.4.1.-] | 1395 | 1444 | 1276 | 1335 | 791 | 721 | 757 | 811 | 387 | 395 | 405 | 424 |
| K12988 alpha-1,3-rhamnosyltransferase [EC:2.4.1.-] | 1512 | 1450 | 1517 | 1459 | 845 | 886 | 809 | 891 | 526 | 454 | 438 | 470 |
| K12989 mannosyltransferase [EC:2.4.1.-] | 0 | 0 | 0 | 1 | 0 | 0 | 0 | 0 | 1 | 3 | 0 | 0 |
| K12993 alpha-1,2-mannosyltransferase/rhamnosyltransferase [EC:2.4.1.-] | 2502 | 2467 | 2362 | 2314 | 1352 | 1344 | 1294 | 1437 | 815 | 745 | 719 | 759 |
| K12994 mannosyl-N-acetyl-alpha-D-glucosaminyl-diphospho-ditrans,octacis-undecaprenol 3-alpha-mannosyltransferase / alpha-1,3-rhamnosyltransferase [EC | 1773 | 1621 | 1670 | 1624 | 918 | 947 | 883 | 1001 | 528 | 487 | 546 | 512 |
| K12995 N-acetyl-alpha-D-glucosaminyl-diphospho-ditrans,octacis-undecaprenol 3-alpha-mannosyltransferase / rhamnosyltransferase [EC:2.4.1.348 2.4.1.-] | 1877 | 1871 | 1885 | 1945 | 1994 | 1942 | 1898 | 1889 | 1242 | 1149 | 1209 | 1134 |
| K12996 rhamnosyltransferase [EC:2.4.1.-] | 8 | 5 | 8 | 11 | 51 | 59 | 49 | 63 | 30 | 21 | 41 | 26 |
| K12997 rhamnosyltransferase [EC:2.4.1.-] | 25 | 33 | 31 | 44 | 8 | 10 | 12 | 10 | 3 | 7 | 1 | 6 |
| K12999 glucosyltransferase [EC:2.4.1.-] | 2 | 0 | 4 | 0 | 1 | 2 | 0 | 0 | 0 | 1 | 1 | 3 |
| K13002 glycosyltransferase [EC:2.4.1.-] | 101 | 100 | 110 | 89 | 100 | 122 | 156 | 136 | 19 | 11 | 12 | 26 |
| K13004 galacturonosyltransferase [EC:2.4.1.-] | 67 | 69 | 69 | 70 | 30 | 24 | 15 | 16 | 64 | 84 | 100 | 95 |
| K13007 Fuc2NAc and GlcNAc transferase [EC:2.4.1.-] | 1379 | 1391 | 1452 | 1529 | 765 | 789 | 868 | 858 | 467 | 453 | 411 | 414 |
| K13678 1,2-diacylglycerol-3-alpha-glucose alpha-1,2-galactosyltransferase [EC:2.4.1.-] | 24 | 55 | 40 | 29 | 7 | 17 | 9 | 5 | 96 | 95 | 86 | 90 |
| K13683 putative colanic acid biosynthesis glycosyltransferase [EC:2.4.-.-] | 132 | 133 | 147 | 175 | 189 | 201 | 181 | 198 | 56 | 35 | 45 | 52 |
| K13684 putative colanic acid biosynthesis glycosyltransferase [EC:2.4.-.-] | 268 | 254 | 259 | 296 | 320 | 354 | 352 | 336 | 70 | 76 | 61 | 74 |
| K13688 cyclic beta-1,2-glucan synthetase [EC:2.4.1.-] | 19 | 17 | 10 | 11 | 3 | 9 | 0 | 3 | 20 | 19 | 13 | 17 |
| K16555 succinoglycan biosynthesis protein ExoO [EC:2.4.-.-] | 0 | 0 | 0 | 0 | 1 | 0 | 1 | 0 | 1 | 0 | 0 | 0 |
| K16698 teichuronic acid biosynthesis glycosyltransferase TuaG [EC:2.4.-.-] | 93 | 97 | 103 | 96 | 37 | 30 | 28 | 25 | 129 | 114 | 90 | 101 |
| K16702 amylovoran biosynthesis glycosyltransferase AmsE [EC:2.4.-.-] | 1 | 1 | 2 | 0 | 0 | 0 | 0 | 0 | 3 | 9 | 10 | 5 |
| K16703 colanic acid/amylovoran biosynthesis glycosyltransferase [EC:2.4.-.-] | 251 | 243 | 225 | 259 | 311 | 370 | 350 | 342 | 45 | 53 | 87 | 53 |
| K17248 N-acetylgalactosamine-N,N'-diacetylbacillosaminyl-diphospho-undecaprenol 4-alpha-N-acetylgalactosaminyltransferase [EC:2.4.1.291] | 0 | 0 | 0 | 0 | 0 | 0 | 0 | 1 | 0 | 0 | 0 | 0 |
| K17249 GalNAc-alpha-(1->4)-GalNAc-alpha-(1->3)-diNAcBac-PP-undecaprenol alpha-1,4-N-acetyl-D-galactosaminyltransferase [EC:2.4.1.292] | 1 | 0 | 0 | 0 | 0 | 0 | 0 | 0 | 0 | 0 | 0 | 0 |
| K19354 heptose III glucuronosyltransferase [EC:2.4.1.-] | 8 | 9 | 0 | 5 | 14 | 27 | 12 | 10 | 60 | 50 | 56 | 79 |
| K19423 glycosyltransferase EpsE [EC:2.4.-.-] | 4 | 6 | 12 | 9 | 4 | 2 | 5 | 1 | 4 | 2 | 2 | 2 |
| K19424 glycosyltransferase EpsF [EC:2.4.-.-] | 53 | 46 | 31 | 45 | 19 | 22 | 16 | 21 | 37 | 35 | 25 | 31 |
| K19425 glycosyltransferase EpsH [EC:2.4.-.-] | 20 | 19 | 19 | 28 | 27 | 26 | 23 | 23 | 11 | 10 | 13 | 13 |
| K20151 alpha(1,3/1,4) fucosyltransferase [EC:2.4.1.65 2.4.1.152] | 53 | 58 | 67 | 71 | 153 | 123 | 146 | 128 | 133 | 114 | 116 | 165 |
| K20926 cellobiuronic acid synthase [EC:2.4.1.-] | 7 | 6 | 3 | 7 | 3 | 5 | 3 | 1 | 6 | 8 | 3 | 12 |
| K21306 dolichyl-phosphooligosaccharide-protein glycotransferase [EC:2.4.99.21] | 0 | 8 | 0 | 1 | 3 | 0 | 0 | 3 | 19 | 20 | 4 | 31 |
| K21363 UDP-Glc:alpha-D-GlcNAc-glucosaminyl-diphosphoundecaprenol beta-1,3-glucosyltransferase [EC:2.4.1.305] | 0 | 0 | 0 | 0 | 0 | 0 | 0 | 0 | 0 | 0 | 1 | 0 |
| K21365 UDP-GalNAc:alpha-D-GalNAc-diphosphoundecaprenol alpha-1,3-N-acetylgalactosaminyltransferase [EC:2.4.1.306] | 0 | 2 | 0 | 1 | 0 | 2 | 0 | 0 | 0 | 0 | 3 | 1 |
| K21367 GDP-Fuc:beta-D-Gal-1,3-alpha-D-GalNAc-1,3-alpha-GalNAc-diphosphoundecaprenol alpha-1,2-fucosyltransferase [EC:2.4.1.308] | 1 | 1 | 3 | 0 | 1 | 0 | 0 | 1 | 0 | 0 | 1 | 0 |
| K22708 poly(ribitol-phosphate) beta-N-acetylglucosaminyltransferase [EC:2.4.1.355] | 0 | 0 | 0 | 0 | 0 | 0 | 0 | 1 | 0 | 0 | 0 | 0 |
| ko01005 Lipopolysaccharide biosynthesis proteins | 106121 | 105764 | 104351 | 106555 | 92358 | 93522 | 88724 | 90743 | 84571 | 85176 | 84139 | 85816 |
| K00996 undecaprenyl-phosphate galactose phosphotransferase [EC:2.7.8.6] | 176 | 186 | 169 | 186 | 22 | 20 | 19 | 15 | 185 | 199 | 178 | 208 |
| K05789 chain length determinant protein (polysaccharide antigen chain regulator) | 1372 | 1425 | 1609 | 1488 | 846 | 894 | 792 | 821 | 466 | 501 | 505 | 517 |
| K08280 lipopolysaccharide O-acetyltransferase [EC:2.3.1.-] | 0 | 0 | 0 | 0 | 2 | 2 | 0 | 0 | 2 | 0 | 0 | 0 |
| K13006 UDP-perosamine 4-acetyltransferase [EC:2.3.1.-] | 0 | 0 | 0 | 1 | 4 | 1 | 0 | 0 | 1 | 2 | 1 | 3 |
| K13012 O-antigen biosynthesis protein WbqP | 5 | 4 | 7 | 10 | 9 | 18 | 8 | 14 | 79 | 62 | 51 | 56 |
| K13013 O-antigen biosynthesis protein WbqV | 0 | 0 | 0 | 0 | 0 | 0 | 0 | 0 | 1 | 0 | 0 | 0 |
| K19714 3-deoxy-alpha-D-manno-octulosonate 8-oxidase [EC:1.1.3.48] | 32 | 19 | 15 | 22 | 1 | 6 | 9 | 3 | 14 | 18 | 29 | 20 |
| K19715 8-amino-3,8-dideoxy-alpha-D-manno-octulosonate transaminase [EC:2.6.1.109] | 0 | 0 | 0 | 0 | 0 | 0 | 0 | 1 | 0 | 1 | 0 | 3 |
| K20444 O-antigen biosynthesis protein [EC:2.4.1.-] | 3451 | 3254 | 3218 | 3467 | 1715 | 1947 | 1683 | 1721 | 2063 | 2399 | 2174 | 2365 |
| K20534 polyisoprenyl-phosphate glycosyltransferase [EC:2.4.-.-] | 2786 | 2690 | 2512 | 2767 | 1356 | 1267 | 1227 | 1395 | 1040 | 1060 | 1003 | 1086 |
| ko01011 Peptidoglycan biosynthesis and degradation proteins | 148799 | 148842 | 148108 | 148413 | 144175 | 147682 | 146254 | 145208 | 151487 | 152531 | 150435 | 151970 |
| K03806 N-acetyl-anhydromuramoyl-L-alanine amidase [EC:3.5.1.28] | 1257 | 1321 | 1334 | 1370 | 793 | 854 | 743 | 744 | 834 | 832 | 819 | 804 |
| K03980 putative peptidoglycan lipid II flippase | 1049 | 946 | 983 | 978 | 785 | 744 | 899 | 831 | 1328 | 1457 | 1435 | 1417 |
| K06078 murein lipoprotein | 52 | 46 | 54 | 45 | 35 | 31 | 36 | 48 | 68 | 65 | 61 | 70 |
| K08304 membrane-bound lytic murein transglycosylase A [EC:4.2.2.-] | 2578 | 2546 | 2307 | 2447 | 1643 | 1585 | 1555 | 1665 | 1517 | 1503 | 1298 | 1509 |
| K08305 membrane-bound lytic murein transglycosylase B [EC:4.2.2.-] | 6399 | 6316 | 5935 | 6045 | 3613 | 3608 | 3456 | 3555 | 2774 | 2766 | 2568 | 2806 |
| K08306 membrane-bound lytic murein transglycosylase C [EC:4.2.2.-] | 660 | 636 | 677 | 744 | 462 | 539 | 457 | 468 | 1027 | 1055 | 1023 | 995 |
| K08307 membrane-bound lytic murein transglycosylase D [EC:4.2.2.-] | 3954 | 4082 | 3947 | 4012 | 4535 | 4958 | 4568 | 4863 | 4119 | 4108 | 4071 | 4031 |
| K08308 membrane-bound lytic murein transglycosylase E [EC:4.2.2.-] | 341 | 342 | 309 | 334 | 225 | 247 | 250 | 210 | 434 | 458 | 528 | 464 |
| K08309 soluble lytic murein transglycosylase [EC:4.2.2.-] | 4161 | 4061 | 4057 | 4074 | 2548 | 2539 | 2463 | 2631 | 2696 | 2668 | 2511 | 2775 |
| K11066 N-acetylmuramoyl-L-alanine amidase [EC:3.5.1.28] | 1389 | 1384 | 1358 | 1305 | 823 | 827 | 862 | 808 | 896 | 896 | 906 | 1006 |
| K18691 membrane-bound lytic murein transglycosylase F [EC:4.2.2.-] | 3857 | 3880 | 3906 | 3752 | 4371 | 4449 | 4389 | 4619 | 3985 | 3831 | 3899 | 3760 |
| K21467 penicillin-binding protein 3 | 618 | 579 | 636 | 643 | 195 | 184 | 225 | 230 | 716 | 713 | 707 | 734 |
| K21468 penicillin-binding protein 4B | 2 | 1 | 1 | 2 | 0 | 0 | 0 | 0 | 0 | 0 | 0 | 0 |
| ko01004 Lipid biosynthesis proteins | 107647 | 107776 | 105211 | 106853 | 93393 | 92039 | 91907 | 94053 | 91152 | 90904 | 89886 | 91401 |
| K00666 fatty-acyl-CoA synthase [EC:6.2.1.-] | 13534 | 13471 | 13154 | 13175 | 10580 | 10545 | 10079 | 11129 | 6811 | 6501 | 6701 | 6805 |
| K12431 phthioceranic/hydroxyphthioceranic acid synthase [EC:2.3.1.287] | 0 | 0 | 0 | 0 | 0 | 1 | 0 | 0 | 0 | 0 | 0 | 0 |

| | | | | | | | | | | | | |
|---|---|---|---|---|---|---|---|---|---|---|---|---|
| K12436 polyketide synthase 12 | 1 | 1 | 0 | 0 | 0 | 0 | 2 | 1 | 2 | 2 | 0 | 1 |
| K12441 phthiocerol/phenolphthiocerol synthesis type-I polyketide synthase B | 0 | 0 | 0 | 0 | 0 | 0 | 0 | 0 | 0 | 0 | 0 | 1 |
| K12443 phthiocerol/phenolphthiocerol synthesis type-I polyketide synthase D | 0 | 0 | 0 | 0 | 0 | 2 | 0 | 0 | 1 | 0 | 0 | 1 |
| K12500 thioesterase III [EC:3.1.2.-] | 226 | 213 | 210 | 212 | 173 | 197 | 150 | 175 | 314 | 306 | 303 | 300 |
| K12507 acyl-CoA synthetase [EC:6.2.1.-] | 1121 | 1025 | 1003 | 1199 | 746 | 864 | 756 | 735 | 607 | 677 | 616 | 660 |
| K13611 polyketide synthase PksJ | 230 | 202 | 236 | 214 | 43 | 38 | 64 | 37 | 4 | 4 | 3 | 9 |
| K13614 polyketide synthase PksN | 20 | 16 | 23 | 17 | 18 | 9 | 20 | 10 | 7 | 3 | 9 | 15 |
| ko01008 Polyketide biosynthesis proteins | 32760 | 31711 | 29474 | 30693 | 15551 | 15491 | 14954 | 15681 | 27362 | 27072 | 26152 | 27715 |
| K05374 yersiniabactin synthetase, thioesterase component | 316 | 272 | 276 | 262 | 107 | 92 | 91 | 73 | 571 | 625 | 621 | 644 |
| K12242 pyochelin biosynthetic protein PchC | 1242 | 1262 | 1098 | 1186 | 614 | 572 | 659 | 700 | 436 | 322 | 383 | 394 |
| K15395 hybrid polyketide synthase / nonribosomal peptide synthetase FtdB | 0 | 0 | 0 | 0 | 0 | 0 | 0 | 0 | 0 | 0 | 1 | 0 |
| K15644 coronafacic acid polyketide synthase Cfa6 | 0 | 0 | 1 | 0 | 0 | 0 | 0 | 0 | 1 | 0 | 0 | 0 |
| K16167 alkylresorcinol/alkylpyrone synthase | 3 | 3 | 2 | 0 | 0 | 0 | 0 | 0 | 0 | 0 | 0 | 0 |
| ko01006 Prenyltransferases | 17179 | 16907 | 17291 | 17177 | 19862 | 19971 | 20297 | 19622 | 18072 | 18543 | 18348 | 18473 |
| ko01007 Amino acid related enzymes | 214733 | 214798 | 216352 | 217549 | 215793 | 218266 | 216152 | 215217 | 213973 | 214937 | 213659 | 214340 |
| K01894 glutamyl-Q tRNA(Asp) synthetase [EC:6.1.1.-] | 845 | 870 | 743 | 838 | 574 | 587 | 566 | 603 | 880 | 875 | 880 | 887 |
| K02805 dTDP-4-amino-4,6-dideoxygalactose transaminase [EC:2.6.1.59] | 810 | 811 | 730 | 867 | 581 | 637 | 644 | 628 | 1307 | 1313 | 1244 | 1311 |
| K03918 L-lysine 6-transaminase [EC:2.6.1.36] | 0 | 3 | 0 | 0 | 0 | 0 | 0 | 0 | 2 | 0 | 1 | 4 |
| K09469 2-aminoethylphosphonate-pyruvate transaminase | 1 | 1 | 0 | 2 | 1 | 2 | 1 | 0 | 2 | 5 | 3 | 7 |
| K10907 aminotransferase [EC:2.6.1.-] | 857 | 901 | 966 | 948 | 387 | 395 | 421 | 346 | 962 | 958 | 984 | 1002 |
| K14261 alanine-synthesizing transaminase [EC:2.6.1.-] | 3003 | 2884 | 2815 | 2903 | 1682 | 1769 | 1641 | 1842 | 1819 | 1690 | 1727 | 1872 |
| K14287 methionine transaminase [EC:2.6.1.88] | 2671 | 2695 | 2605 | 2714 | 1498 | 1677 | 1599 | 1631 | 1569 | 1556 | 1663 | 1710 |
| ko00199 Cytochrome P450 | 0 | 1 | 0 | 0 | 12 | 15 | 5 | 5 | 23 | 25 | 29 | 24 |
| K15629 fatty-acid peroxygenase [EC:1.11.2.4] | 0 | 1 | 0 | 0 | 12 | 15 | 5 | 5 | 23 | 23 | 29 | 24 |
| K21119 cytochrome P450 family 130 | 0 | 0 | 0 | 0 | 0 | 0 | 0 | 0 | 0 | 2 | 0 | 0 |
| ko00194 Photosynthesis proteins | 26142 | 26229 | 27048 | 26875 | 24434 | 24776 | 24578 | 24154 | 24174 | 24255 | 24515 | 24112 |
| K02116 ATP synthase protein I | 1064 | 987 | 1040 | 1044 | 541 | 583 | 555 | 517 | 615 | 593 | 625 | 641 |
| Protein families: genetic information processing | 1747091 | 1747644 | 1751205 | 1755209 | 1724736 | 1740347 | 1733333 | 1727195 | 1747550 | 1755511 | 1742577 | 1751148 |
| ko03000 Transcription factors | 267307 | 264211 | 259048 | 263978 | 192182 | 194183 | 190278 | 194023 | 243829 | 248530 | 239279 | 248388 |
| K00375 GntR family transcriptional regulator / MocR family aminotransferase | 9309 | 9057 | 8594 | 8770 | 5533 | 6034 | 5760 | 5940 | 6407 | 6969 | 6832 | 6897 |
| K01420 CRP/FNR family transcriptional regulator, anaerobic regulatory protein | 915 | 958 | 938 | 957 | 594 | 719 | 649 | 622 | 1045 | 1013 | 1019 | 1034 |
| K01926 redox-sensing transcriptional repressor | 1048 | 1051 | 1164 | 1089 | 1553 | 1544 | 1630 | 1537 | 1544 | 1535 | 1526 | 1594 |
| K01994 LuxR family transcriptional regulator, transcriptional regulator of spore coat protein | 0 | 0 | 0 | 0 | 0 | 0 | 0 | 0 | 0 | 2 | 0 | 3 |
| K02019 molybdate transport system regulatory protein | 1759 | 1695 | 1606 | 1642 | 1053 | 1053 | 1001 | 1031 | 1222 | 1209 | 1084 | 1227 |
| K02043 GntR family transcriptional regulator, phosphonate transport system regulatory protein | 1247 | 1213 | 1124 | 1115 | 571 | 562 | 586 | 582 | 888 | 910 | 878 | 962 |
| K02076 Fur family transcriptional regulator, zinc uptake regulator | 21 | 30 | 38 | 29 | 5 | 5 | 8 | 6 | 22 | 25 | 24 | 29 |
| K02081 DeoR family transcriptional regulator, aga operon transcriptional repressor | 441 | 377 | 391 | 366 | 526 | 607 | 559 | 517 | 129 | 139 | 129 | 164 |
| K02099 AraC family transcriptional regulator, arabinose operon regulatory protein | 972 | 952 | 836 | 1021 | 555 | 537 | 571 | 593 | 1444 | 1513 | 1410 | 1574 |
| K02103 GntR family transcriptional regulator, arabinose operon transcriptional repressor | 444 | 456 | 435 | 469 | 118 | 85 | 115 | 92 | 416 | 470 | 387 | 413 |
| K02167 TetR/AcrR family transcriptional regulator, transcriptional repressor of bet genes | 452 | 396 | 369 | 385 | 177 | 190 | 196 | 178 | 548 | 542 | 538 | 545 |
| K02430 DeoR family transcriptional regulator, L-fucose operon activator | 446 | 498 | 463 | 488 | 274 | 299 | 318 | 292 | 542 | 581 | 511 | 611 |
| K02436 DeoR family transcriptional regulator, galactitol utilization operon repressor | 2 | 6 | 0 | 3 | 3 | 9 | 9 | 16 | 34 | 37 | 39 | 50 |
| K02443 glycerol uptake operon antiterminator | 214 | 198 | 211 | 265 | 113 | 133 | 109 | 99 | 572 | 584 | 559 | 565 |
| K02444 DeoR family transcriptional regulator, glycerol-3-phosphate regulon repressor | 2405 | 2299 | 2339 | 2391 | 1580 | 1724 | 1588 | 1556 | 1733 | 1634 | 1528 | 1741 |
| K02468 DeoR family transcriptional regulator, glucitol operon repressor | 446 | 452 | 399 | 481 | 347 | 270 | 327 | 288 | 564 | 638 | 614 | 668 |
| K02508 AraC family transcriptional regulator, 4-hydroxyphenylacetate 3-monooxygenase operon regulatory protein | 609 | 614 | 614 | 629 | 267 | 250 | 280 | 262 | 842 | 795 | 820 | 865 |
| K02521 LysR family transcriptional regulator, positive regulator for ilvC | 628 | 644 | 563 | 699 | 414 | 501 | 434 | 434 | 838 | 881 | 861 | 880 |
| K02525 LacI family transcriptional regulator, kdg operon repressor | 1294 | 1214 | 1176 | 1171 | 751 | 780 | 798 | 798 | 906 | 901 | 922 | 955 |
| K02529 LacI family transcriptional regulator | 15200 | 15226 | 15889 | 15612 | 18828 | 18535 | 19422 | 18441 | 25315 | 25807 | 25165 | 25397 |
| K02530 DeoR family transcriptional regulator, lactose phosphotransferase system repressor | 102 | 93 | 76 | 84 | 39 | 33 | 32 | 23 | 115 | 132 | 119 | 117 |
| K02531 transcriptional antiterminator | 1 | 4 | 8 | 0 | 6 | 0 | 0 | 0 | 0 | 0 | 7 | 0 |
| K02538 activator of the mannose operon, transcriptional antiterminator | 485 | 537 | 509 | 560 | 560 | 634 | 589 | 522 | 288 | 363 | 296 | 302 |
| K02562 mannitol operon repressor | 357 | 391 | 287 | 388 | 240 | 316 | 272 | 280 | 535 | 479 | 468 | 568 |
| K02565 N-acetylglucosamine repressor | 755 | 714 | 672 | 704 | 462 | 526 | 483 | 493 | 1003 | 1034 | 925 | 1058 |
| K02616 phenylacetic acid degradation operon negative regulatory protein | 336 | 336 | 297 | 358 | 89 | 91 | 77 | 102 | 649 | 733 | 553 | 662 |
| K02623 LysR family transcriptional regulator, pca operon transcriptional activator | 2094 | 1934 | 1843 | 1834 | 1044 | 1014 | 940 | 1142 | 754 | 631 | 590 | 623 |
| K02624 IclR family transcriptional regulator, pca regulon regulatory protein | 1333 | 1389 | 1333 | 1337 | 505 | 530 | 460 | 460 | 1800 | 1921 | 1929 | 1984 |
| K02647 carbohydrate diacid regulator | 3542 | 3281 | 3274 | 3415 | 1611 | 1548 | 1451 | 1536 | 2814 | 2985 | 2706 | 2889 |
| K02688 transcriptional regulator, propionate catabolism operon regulatory protein | 2419 | 2375 | 2375 | 2296 | 1329 | 1282 | 1200 | 1335 | 789 | 809 | 801 | 779 |
| K02854 AraC family transcriptional regulator, L-rhamnose operon transcriptional activator RhaR | 541 | 562 | 526 | 550 | 379 | 419 | 397 | 441 | 842 | 877 | 744 | 858 |
| K02855 AraC family transcriptional regulator, L-rhamnose operon regulatory protein RhaS | 643 | 627 | 558 | 637 | 381 | 442 | 461 | 378 | 711 | 676 | 650 | 770 |
| K03402 transcriptional regulator of arginine metabolism | 938 | 1002 | 1025 | 1003 | 1326 | 1408 | 1364 | 1216 | 1418 | 1484 | 1462 | 1469 |
| K03435 LacI family transcriptional regulator, fructose operon transcriptional repressor | 2642 | 2519 | 2343 | 2462 | 1260 | 1224 | 1308 | 1438 | 2130 | 1987 | 2055 | 2111 |
| K03436 DeoR family transcriptional regulator, fructose operon transcriptional repressor | 390 | 361 | 358 | 398 | 59 | 70 | 66 | 63 | 687 | 786 | 739 | 799 |
| K03477 DeoR family transcriptional regulator, ulaG and ulaABCDEF operon transcriptional repressor | 430 | 507 | 420 | 475 | 333 | 419 | 300 | 349 | 613 | 632 | 624 | 622 |
| K03480 transcriptional antiterminator | 14 | 15 | 15 | 21 | 4 | 26 | 18 | 21 | 39 | 41 | 26 | 37 |
| K03481 RpiR family transcriptional regulator, glv operon transcriptional regulator | 48 | 35 | 53 | 59 | 35 | 46 | 18 | 34 | 68 | 90 | 63 | 85 |
| K03482 GntR family transcriptional regulator, glv operon transcriptional regulator | 913 | 822 | 834 | 874 | 445 | 428 | 389 | 430 | 1025 | 1186 | 1084 | 1201 |
| K03483 mannitol operon transcriptional antiterminator | 170 | 141 | 124 | 177 | 24 | 33 | 33 | 45 | 163 | 172 | 121 | 177 |
| K03484 LacI family transcriptional regulator, sucrose operon repressor | 1877 | 1921 | 1864 | 1948 | 596 | 621 | 631 | 614 | 1817 | 2005 | 1937 | 1965 |
| K03485 LacI family transcriptional regulator, trehalose operon repressor | 602 | 663 | 621 | 622 | 455 | 425 | 473 | 446 | 839 | 911 | 808 | 950 |
| K03486 GntR family transcriptional regulator, trehalose operon transcriptional repressor | 509 | 446 | 463 | 471 | 89 | 83 | 76 | 74 | 434 | 428 | 456 | 435 |
| K03487 LacI family transcriptional regulator, asc operon repressor | 833 | 959 | 876 | 872 | 780 | 795 | 779 | 763 | 895 | 901 | 836 | 911 |
| K03488 beta-glucoside operon transcriptional antiterminator | 1341 | 1343 | 1304 | 1307 | 783 | 849 | 640 | 728 | 1215 | 1409 | 1253 | 1326 |
| K03489 GntR family transcriptional regulator, transcriptional regulator of bglA | 4 | 0 | 0 | 3 | 7 | 0 | 4 | 1 | 4 | 4 | 2 | 3 |
| K03490 AraC family transcriptional regulator, dual regulator of chb operon | 522 | 523 | 517 | 550 | 304 | 324 | 326 | 331 | 688 | 672 | 662 | 743 |
| K03491 lichenan operon transcriptional antiterminator | 706 | 659 | 752 | 671 | 108 | 111 | 99 | 94 | 518 | 729 | 623 | 540 |
| K03492 GntR family transcriptional regulator | 429 | 494 | 458 | 446 | 355 | 360 | 326 | 302 | 575 | 601 | 611 | 617 |
| K03493 transcriptional antiterminator | 2 | 0 | 0 | 0 | 0 | 0 | 0 | 0 | 0 | 0 | 0 | 0 |
| K03497 ParB family transcriptional regulator, chromosome partitioning protein | 15665 | 15894 | 15757 | 15732 | 16344 | 16071 | 16291 | 16292 | 17335 | 17504 | 17321 | 17147 |
| K03556 LuxR family transcriptional regulator, maltose regulon positive regulatory protein | 7906 | 7558 | 7507 | 7853 | 4618 | 4863 | 4585 | 4834 | 6551 | 6701 | 6608 | 6660 |
| K03576 LysR family transcriptional regulator, regulator for metE and metH | 1590 | 1569 | 1516 | 1598 | 967 | 933 | 849 | 928 | 1705 | 1765 | 1515 | 1662 |
| K03577 TetR/AcrR family transcriptional regulator, acrAB operon repressor | 254 | 254 | 227 | 287 | 213 | 235 | 220 | 247 | 292 | 308 | 273 | 283 |
| K03603 GntR family transcriptional regulator, negative regulator for fad regulon and positive regulator of fabA | 383 | 451 | 394 | 405 | 275 | 340 | 268 | 273 | 580 | 621 | 579 | 587 |
| K03604 LacI family transcriptional regulator, purine nucleotide synthesis repressor | 647 | 622 | 590 | 579 | 382 | 450 | 410 | 477 | 884 | 843 | 874 | 895 |
| K03705 heat-inducible transcriptional repressor | 665 | 576 | 645 | 689 | 234 | 210 | 235 | 234 | 563 | 665 | 599 | 610 |
| K03706 transcriptional pleiotropic repressor | 375 | 380 | 400 | 433 | 94 | 91 | 104 | 74 | 415 | 454 | 398 | 396 |
| K03708 transcriptional regulator of stress and heat shock response | 59 | 61 | 66 | 64 | 31 | 45 | 32 | 39 | 48 | 67 | 80 | 69 |
| K03709 DtxR family transcriptional regulator, Mn-dependent transcriptional regulator | 31 | 44 | 37 | 48 | 16 | 5 | 9 | 7 | 41 | 62 | 50 | 44 |
| K03710 GntR family transcriptional regulator | 2062 | 1893 | 2166 | 2014 | 2147 | 2146 | 2146 | 2076 | 2555 | 2516 | 2466 | 2615 |
| K03711 Fur family transcriptional regulator, ferric uptake regulator | 1984 | 2028 | 2114 | 2121 | 2719 | 2748 | 2758 | 2728 | 2502 | 2496 | 2515 | 2459 |
| K03712 MarR family transcriptional regulator, multiple antibiotic resistance protein MarR | 317 | 283 | 329 | 361 | 313 | 307 | 276 | 312 | 287 | 370 | 368 | 367 |
| K03713 MerR family transcriptional regulator, glutamine synthetase repressor | 16 | 14 | 26 | 23 | 4 | 4 | 5 | 10 | 17 | 39 | 26 | 19 |
| K03717 LysR family transcriptional regulator, transcriptional activator of nhaA | 515 | 464 | 452 | 483 | 424 | 395 | 368 | 352 | 588 | 612 | 583 | 641 |
| K03718 Lrp/AsnC family transcriptional regulator, regulator for asnA, asnC and gidA | 912 | 804 | 882 | 912 | 2303 | 2351 | 2342 | 2291 | 1691 | 1681 | 1785 | 1643 |
| K03719 Lrp/AsnC family transcriptional regulator, leucine-responsive regulatory protein | 4752 | 4699 | 4659 | 4629 | 3543 | 3579 | 3433 | 3728 | 2945 | 2961 | 2735 | 2940 |
| K03720 TrpR family transcriptional regulator, trp operon repressor | 280 | 228 | 221 | 289 | 160 | 193 | 143 | 136 | 353 | 385 | 349 | 348 |
| K03721 transcriptional regulator of aroF, aroG, tyrA and aromatic amino acid transport | 3259 | 3225 | 3068 | 3243 | 1801 | 1903 | 1878 | 1867 | 2029 | 1954 | 1871 | 1916 |
| K03755 AraC family transcriptional regulator, transcriptional activator of adiA | 2 | 4 | 4 | 1 | 8 | 16 | 17 | 13 | 73 | 86 | 39 | 48 |
| K03764 MetJ family transcriptional regulator, methionine regulon repressor | 289 | 220 | 269 | 229 | 143 | 179 | 178 | 140 | 389 | 412 | 340 | 415 |
| K03765 transcriptional activator of cad operon | 524 | 524 | 532 | 578 | 220 | 192 | 189 | 187 | 1145 | 1240 | 1092 | 1273 |
| K03892 ArsR family transcriptional regulator, arsenate/arsenite/antimonite-responsive transcriptional repressor | 768 | 801 | 725 | 782 | 640 | 574 | 602 | 628 | 395 | 487 | 452 | 423 |
| K03973 phage shock protein C | 186 | 201 | 192 | 214 | 123 | 133 | 107 | 137 | 262 | 301 | 251 | 287 |
| K03974 psp operon transcriptional activator | 506 | 562 | 455 | 549 | 331 | 407 | 365 | 337 | 726 | 702 | 791 | 778 |
| K04033 AraC family transcriptional regulator, ethanolamine operon transcriptional activator | 462 | 405 | 393 | 414 | 138 | 131 | 105 | 144 | 888 | 819 | 844 | 947 |
| K04769 AbrB family transcriptional regulator, stage V sporulation protein T | 413 | 381 | 389 | 353 | 96 | 130 | 97 | 92 | 297 | 356 | 293 | 326 |
| K05311 central glycolytic genes regulator | 107 | 114 | 93 | 114 | 40 | 38 | 41 | 35 | 140 | 190 | 180 | 172 |
| K05346 deoxyribonucleoside regulator | 95 | 67 | 65 | 78 | 24 | 43 | 58 | 36 | 131 | 135 | 127 | 188 |
| K05372 AraC family transcriptional regulator | 881 | 801 | 852 | 907 | 364 | 318 | 379 | 326 | 990 | 1071 | 918 | 1021 |
| K05499 LacI family transcriptional regulator, repressor for deo operon, udp, cdd, tsx, nupC, and nupG | 897 | 901 | 885 | 924 | 500 | 600 | 547 | 501 | 1311 | 1331 | 1177 | 1285 |
| K05501 TetR/AcrR family transcriptional regulator | 448 | 418 | 442 | 449 | 323 | 362 | 335 | 322 | 610 | 689 | 600 | 652 |
| K05527 BolA family transcriptional regulator, general stress-responsive regulator | 693 | 604 | 658 | 655 | 370 | 416 | 390 | 431 | 438 | 446 | 444 | 449 |
| K05596 LysR family transcriptional regulator, chromosome initiation inhibitor | 2185 | 2005 | 2022 | 2015 | 1295 | 1169 | 1227 | 1277 | 1244 | 1254 | 1151 | 1296 |
| K05785 transcriptional antiterminator RfaH | 325 | 303 | 315 | 280 | 221 | 262 | 215 | 214 | 441 | 481 | 432 | 477 |
| K05798 LysR family transcriptional regulator, transcriptional activator for leuABCD operon | 631 | 586 | 559 | 580 | 444 | 450 | 426 | 378 | 818 | 828 | 821 | 865 |
| K05799 GntR family transcriptional regulator, transcriptional repressor for pyruvate dehydrogenase complex | 1913 | 1776 | 1829 | 1864 | 1126 | 1087 | 990 | 1157 | 1770 | 1741 | 1701 | 1701 |
| K05800 Lrp/AsnC family transcriptional regulator | 907 | 955 | 900 | 848 | 525 | 492 | 525 | 544 | 589 | 681 | 600 | 534 |
| K05804 AraC family transcriptional regulator, mar-sox-rob regulon activator | 811 | 796 | 848 | 787 | 568 | 591 | 543 | 559 | 1148 | 1174 | 1103 | 1115 |
| K05817 LysR family transcriptional regulator, hca operon transcriptional activator | 1 | 1 | 1 | 0 | 5 | 10 | 6 | 3 | 1 | 5 | 2 | 5 |
| K05818 IclR family transcriptional regulator, mhp operon transcriptional activator | 366 | 314 | 301 | 303 | 79 | 80 | 113 | 94 | 635 | 593 | 576 | 635 |
| K05836 GntR family transcriptional regulator, histidine utilization repressor | 469 | 470 | 410 | 486 | 294 | 350 | 312 | 339 | 656 | 687 | 602 | 716 |

| | | | | | | | | | | | | |
|---|---|---|---|---|---|---|---|---|---|---|---|---|
| K05880 transcriptional activator for dhaKLM operon | 1099 | 1168 | 1021 | 1091 | 748 | 740 | 710 | 702 | 1570 | 1670 | 1537 | 1688 |
| K06075 MarR family transcriptional regulator, transcriptional regulator for hemolysin | 1202 | 1224 | 1185 | 1150 | 749 | 881 | 836 | 901 | 837 | 883 | 834 | 862 |
| K06140 regulator of nucleoside diphosphate kinase | 869 | 869 | 888 | 859 | 505 | 513 | 527 | 532 | 581 | 578 | 579 | 564 |
| K06145 LacI family transcriptional regulator, gluconate utilization system Gnt-I transcriptional repressor | 946 | 846 | 819 | 983 | 781 | 782 | 804 | 701 | 940 | 993 | 943 | 988 |
| K06146 LacI family transcriptional regulator, gluconate utilization system Gnt-II transcriptional activator | 194 | 216 | 229 | 202 | 289 | 314 | 288 | 348 | 53 | 72 | 69 | 47 |
| K06283 putative DeoR family transcriptional regulator, stage III sporulation protein D | 171 | 159 | 152 | 154 | 33 | 38 | 33 | 42 | 99 | 143 | 110 | 128 |
| K06284 AbrB family transcriptional regulator, transcriptional pleiotropic regulator of transition state genes | 36 | 27 | 45 | 30 | 21 | 27 | 34 | 23 | 21 | 27 | 23 | 24 |
| K06314 prespore-specific regulator | 0 | 0 | 0 | 2 | 0 | 0 | 0 | 0 | 0 | 0 | 0 | 0 |
| K06599 AraC family transcriptional regulator, chemosensory pili system protein ChpD | 1354 | 1370 | 1297 | 1287 | 691 | 695 | 708 | 754 | 480 | 392 | 402 | 380 |
| K06608 DeoR family transcriptional regulator, myo-inositol catabolism operon repressor | 54 | 72 | 62 | 61 | 11 | 4 | 2 | 4 | 129 | 95 | 94 | 113 |
| K06714 arginine utilization regulatory protein | 2205 | 2290 | 2324 | 2166 | 1238 | 1197 | 1210 | 1227 | 841 | 856 | 779 | 927 |
| K07110 XRE family transcriptional regulator, fatty acid utilization regulator | 0 | 0 | 2 | 2 | 0 | 2 | 0 | 0 | 1 | 3 | 2 | 1 |
| K07471 probable transcriptional regulator | 7 | 2 | 1 | 3 | 8 | 2 | 6 | 5 | 27 | 32 | 26 | 29 |
| K07506 AraC family transcriptional regulator | 2283 | 2435 | 2223 | 2323 | 2497 | 2490 | 2469 | 2449 | 2270 | 2133 | 2111 | 2142 |
| K07592 LysR family transcriptional regulator, tdc operon transcriptional activator | 337 | 326 | 344 | 355 | 123 | 107 | 78 | 99 | 784 | 697 | 761 | 788 |
| K07721 ArsR family transcriptional regulator | 0 | 0 | 0 | 0 | 0 | 2 | 1 | 0 | 4 | 13 | 0 | 2 |
| K07722 CopG family transcriptional regulator, nickel-responsive regulator | 313 | 316 | 291 | 332 | 241 | 256 | 234 | 248 | 402 | 435 | 420 | 432 |
| K07724 Ner family transcriptional regulator | 230 | 215 | 224 | 198 | 135 | 174 | 151 | 122 | 288 | 323 | 309 | 313 |
| K07726 putative transcriptional regulator | 375 | 392 | 386 | 348 | 204 | 196 | 236 | 231 | 555 | 534 | 540 | 600 |
| K07727 putative transcriptional regulator | 1650 | 1610 | 1691 | 1666 | 859 | 864 | 785 | 816 | 817 | 786 | 812 | 796 |
| K07728 putative transcriptional regulator | 0 | 2 | 0 | 1 | 0 | 1 | 2 | 0 | 4 | 3 | 3 | 6 |
| K07729 putative transcriptional regulator | 680 | 661 | 606 | 603 | 329 | 299 | 270 | 293 | 565 | 607 | 583 | 618 |
| K07730 putative transcriptional regulator | 0 | 0 | 0 | 1 | 0 | 0 | 1 | 1 | 1 | 11 | 3 | 7 |
| K07731 putative transcriptional regulator | 0 | 0 | 0 | 0 | 0 | 0 | 0 | 0 | 0 | 0 | 0 | 1 |
| K07733 prophage regulatory protein | 778 | 800 | 838 | 800 | 484 | 523 | 498 | 504 | 938 | 928 | 865 | 784 |
| K07734 transcriptional regulator | 1001 | 972 | 997 | 940 | 548 | 533 | 527 | 640 | 300 | 325 | 291 | 321 |
| K07735 putative transcriptional regulator | 1710 | 1643 | 1611 | 1638 | 2014 | 2002 | 2035 | 1907 | 1651 | 1693 | 1745 | 1657 |
| K07736 CarD family transcriptional regulator | 197 | 222 | 196 | 190 | 109 | 93 | 129 | 112 | 314 | 341 | 303 | 332 |
| K07738 transcriptional repressor NrdR | 1386 | 1371 | 1278 | 1391 | 705 | 655 | 657 | 704 | 852 | 959 | 873 | 954 |
| K07978 GntR family transcriptional regulator | 0 | 0 | 0 | 0 | 0 | 0 | 0 | 0 | 1 | 0 | 0 | 0 |
| K07979 GntR family transcriptional regulator | 615 | 547 | 562 | 584 | 162 | 186 | 188 | 160 | 663 | 721 | 657 | 742 |
| K08277 transcriptional activator CaiF | 98 | 88 | 80 | 92 | 134 | 147 | 136 | 145 | 41 | 48 | 23 | 27 |
| K08365 MerR family transcriptional regulator, mercuric resistance operon regulatory protein | 940 | 860 | 973 | 945 | 657 | 697 | 741 | 642 | 254 | 225 | 218 | 267 |
| K09017 TetR/AcrR family transcriptional regulator | 2680 | 2595 | 2579 | 2547 | 1491 | 1509 | 1511 | 1731 | 1198 | 1234 | 1095 | 1211 |
| K09681 LysR family transcriptional regulator, transcription activator of glutamate synthase operon | 8 | 13 | 6 | 17 | 2 | 0 | 2 | 0 | 23 | 29 | 30 | 28 |
| K09682 MarR family transcriptional regulator, protease production regulatory protein HPr | 0 | 0 | 0 | 1 | 0 | 0 | 0 | 0 | 0 | 0 | 0 | 0 |
| K09683 hut operon positive regulatory protein | 36 | 29 | 30 | 38 | 0 | 0 | 0 | 0 | 60 | 45 | 63 | 40 |
| K09684 PucR family transcriptional regulator, purine catabolism regulatory protein | 2309 | 2297 | 2247 | 2329 | 1075 | 1071 | 1027 | 1109 | 978 | 1023 | 872 | 892 |
| K09685 purine operon repressor | 102 | 103 | 98 | 125 | 30 | 38 | 32 | 23 | 140 | 128 | 132 | 142 |
| K09720 HTH-type transcriptional regulator, global nitrogen regulator NrpRI | 0 | 5 | 1 | 5 | 2 | 0 | 0 | 2 | 9 | 19 | 7 | 6 |
| K09825 Fur family transcriptional regulator, peroxide stress response regulator | 526 | 520 | 595 | 540 | 942 | 1091 | 994 | 953 | 910 | 867 | 1021 | 900 |
| K09826 Fur family transcriptional regulator, iron response regulator | 0 | 0 | 0 | 0 | 0 | 4 | 0 | 2 | 0 | 1 | 0 | 0 |
| K10711 GntR family transcriptional regulator, frlABCD operon transcriptional regulator | 216 | 181 | 165 | 185 | 271 | 261 | 241 | 204 | 10 | 25 | 25 | 21 |
| K10778 AraC family transcriptional regulator, regulatory protein of adaptative response / methylated-DNA-[protein]-cysteine methyltransferase [EC:2.1.1.63] | 2735 | 2835 | 2665 | 2753 | 3023 | 3153 | 3160 | 2983 | 3282 | 3094 | 3324 | 3313 |
| K10947 PadR family transcriptional regulator, regulatory protein PadR | 595 | 625 | 646 | 667 | 677 | 695 | 788 | 685 | 927 | 855 | 892 | 966 |
| K10972 LysR family transcriptional regulator, transcriptional activator of the allD operon | 3 | 1 | 1 | 1 | 15 | 13 | 14 | 12 | 39 | 51 | 40 | 38 |
| K10973 IclR family transcriptional regulator, negative regulator of allantoin and glyoxylate utilization operons | 2 | 0 | 1 | 1 | 5 | 5 | 12 | 8 | 43 | 60 | 44 | 58 |
| K11474 GntR family transcriptional regulator, glc operon transcriptional activator | 1331 | 1272 | 1239 | 1355 | 697 | 723 | 700 | 695 | 522 | 462 | 433 | 468 |
| K11475 GntR family transcriptional regulator, vanillate catabolism transcriptional regulator | 1216 | 1202 | 1188 | 1210 | 675 | 698 | 644 | 732 | 403 | 392 | 322 | 425 |
| K11534 DeoR family transcriptional regulator, deoxyribose operon repressor | 858 | 793 | 824 | 843 | 597 | 649 | 543 | 687 | 688 | 742 | 690 | 802 |
| K11908 sigma-54 dependent transcriptional regulator | 2 | 0 | 0 | 1 | 1 | 2 | 1 | 0 | 1 | 1 | 0 | 1 |
| K11914 sigma-54 dependent transcriptional regulator | 2204 | 2156 | 1992 | 2087 | 1183 | 1171 | 1138 | 1145 | 636 | 672 | 587 | 641 |
| K11917 sigma-54 dependent transcriptional regulator | 1387 | 1339 | 1202 | 1289 | 712 | 771 | 659 | 824 | 408 | 411 | 350 | 362 |
| K11920 AraC family transcriptional regulator | 0 | 2 | 0 | 0 | 10 | 9 | 8 | 9 | 0 | 3 | 1 | 1 |
| K11921 LysR family transcriptional regulator, cyn operon transcriptional activator | 1490 | 1403 | 1367 | 1450 | 671 | 672 | 626 | 731 | 1051 | 1017 | 1000 | 1073 |
| K11922 GntR family transcriptional regulator, mannosyl-D-glycerate transport/metabolism system repressor | 3 | 13 | 12 | 9 | 14 | 32 | 24 | 29 | 11 | 9 | 16 | 9 |
| K11924 DtxR family transcriptional regulator, manganese transport regulator | 267 | 238 | 269 | 291 | 177 | 218 | 187 | 169 | 356 | 413 | 333 | 408 |
| K11925 SgrR family transcriptional regulator | 977 | 1097 | 997 | 1046 | 722 | 777 | 704 | 763 | 1284 | 1509 | 1336 | 1456 |
| K11926 sigma factor-binding protein Crl | 201 | 203 | 231 | 216 | 134 | 172 | 135 | 163 | 311 | 343 | 328 | 321 |
| K12113 LacI family transcriptional regulator, ebg operon repressor | 2 | 5 | 5 | 4 | 16 | 16 | 14 | 7 | 57 | 57 | 78 | 55 |
| K12146 hydrogenase-4 transcriptional activator | 362 | 356 | 389 | 385 | 594 | 663 | 554 | 597 | 4 | 2 | 3 | 2 |
| K12243 AraC family transcriptional regulator, transcriptional activator of the genes for pyochelin and ferripyochelin receptors | 1694 | 1614 | 1363 | 1358 | 812 | 723 | 705 | 851 | 489 | 431 | 406 | 439 |
| K13530 AraC family transcriptional regulator, regulatory protein of adaptative response / methylphosphotriester-DNA alkyltransferase methyltransferase [EC | 31 | 39 | 37 | 27 | 5 | 5 | 4 | 3 | 25 | 50 | 34 | 39 |
| K13631 AraC family transcriptional regulator, mar-sox-rob regulon activator | 425 | 415 | 406 | 405 | 285 | 307 | 295 | 299 | 601 | 707 | 634 | 543 |
| K13633 AraC family transcriptional regulator, transcriptional activator FtrA | 552 | 509 | 518 | 496 | 372 | 329 | 308 | 356 | 712 | 767 | 715 | 745 |
| K13634 LysR family transcriptional regulator, cys regulon transcriptional activator | 1743 | 1650 | 1649 | 1647 | 1023 | 975 | 1027 | 959 | 1086 | 1240 | 1135 | 1152 |
| K13635 LysR family transcriptional regulator, cys regulon transcriptional activator | 542 | 604 | 563 | 538 | 370 | 424 | 406 | 378 | 771 | 850 | 799 | 796 |
| K13636 LysR family transcriptional regulator, D-serine deaminase activator | 951 | 1001 | 915 | 931 | 483 | 628 | 516 | 562 | 1407 | 1467 | 1399 | 1549 |
| K13637 GntR family transcriptional regulator, uxu operon transcriptional repressor | 411 | 387 | 385 | 457 | 294 | 347 | 322 | 326 | 568 | 579 | 586 | 533 |
| K13638 MerR family transcriptional regulator, Zn(II)-responsive regulator of zntA | 273 | 243 | 283 | 294 | 188 | 202 | 210 | 193 | 379 | 372 | 400 | 385 |
| K13639 MerR family transcriptional regulator, redox-sensitive transcriptional activator SoxR | 430 | 442 | 467 | 464 | 321 | 311 | 301 | 301 | 441 | 422 | 431 | 436 |
| K13640 MerR family transcriptional regulator, heat shock protein HspR | 8 | 4 | 6 | 2 | 2 | 1 | 3 | 5 | 9 | 7 | 8 | 9 |
| K13641 IclR family transcriptional regulator, acetate operon repressor | 549 | 564 | 536 | 596 | 324 | 401 | 437 | 373 | 729 | 780 | 796 | 760 |
| K13642 CRP/FNR family transcriptional regulator, transcriptional activator FtrB | 0 | 0 | 0 | 0 | 0 | 0 | 0 | 0 | 2 | 2 | 0 | 0 |
| K13643 Rrf2 family transcriptional regulator, iron-sulfur cluster assembly transcription factor | 1029 | 1074 | 1121 | 1060 | 628 | 673 | 639 | 662 | 684 | 671 | 680 | 689 |
| K13652 AraC family transcriptional regulator | 1440 | 1448 | 1509 | 1406 | 2899 | 2870 | 2959 | 3080 | 2712 | 2637 | 2795 | 2586 |
| K13653 AraC family transcriptional regulator | 2752 | 2527 | 2769 | 2701 | 624 | 631 | 636 | 503 | 3002 | 3142 | 2975 | 3238 |
| K13654 GntR family transcriptional regulator, colanic acid and biofilm gene transcriptional regulator | 108 | 124 | 118 | 98 | 183 | 159 | 183 | 166 | 9 | 11 | 8 | 11 |
| K13655 HTH-type transcriptional regulator / antitoxin MqsA | 1 | 0 | 0 | 3 | 0 | 0 | 0 | 2 | 0 | 0 | 0 | 0 |
| K13770 TetR/AcrR family transcriptional regulator, fatty acid metabolism regulator protein | 0 | 0 | 0 | 0 | 2 | 1 | 3 | 1 | 0 | 1 | 0 | 0 |
| K13772 Rrf2 family transcriptional regulator, iron-responsive regulator | 0 | 0 | 0 | 1 | 0 | 0 | 0 | 0 | 0 | 2 | 0 | 2 |
| K13794 LysR family transcriptional regulator, regulatory protein for tcuABC | 1 | 4 | 0 | 1 | 2 | 5 | 5 | 2 | 2 | 0 | 1 | 2 |
| K13928 LysR family transcriptional regulator, malonate utilization transcriptional regulator | 2058 | 2132 | 1855 | 1914 | 1156 | 1164 | 1100 | 1190 | 1110 | 1072 | 1034 | 1115 |
| K14056 HTH-type transcriptional regulator, repressor for puuD | 158 | 184 | 197 | 192 | 46 | 49 | 46 | 52 | 422 | 443 | 401 | 360 |
| K14057 LysR family transcriptional regulator, regulator of abg operon | 285 | 255 | 275 | 304 | 89 | 69 | 62 | 87 | 626 | 601 | 524 | 567 |
| K14063 AraC family transcriptional regulator, positive regulator of tynA and feaB | 364 | 409 | 364 | 415 | 111 | 101 | 93 | 109 | 755 | 753 | 628 | 794 |
| K14348 GntR family transcriptional regulator, L-lactate dehydrogenase operon regulator | 514 | 523 | 473 | 525 | 383 | 393 | 414 | 386 | 707 | 710 | 775 | 750 |
| K14414 transcriptional regulatory protein RtcR | 3081 | 2907 | 2952 | 2869 | 2046 | 2024 | 2123 | 2023 | 922 | 899 | 870 | 908 |
| K15545 transcriptional regulator of PTS gene | 624 | 713 | 682 | 710 | 466 | 511 | 452 | 453 | 944 | 937 | 928 | 966 |
| K15546 XRE family transcriptional regulator, aerobic/anaerobic benzoate catabolism transcriptional regulator | 0 | 0 | 0 | 0 | 0 | 0 | 0 | 0 | 1 | 3 | 1 | 0 |
| K15735 GntR family transcriptional regulator, carbon starvation induced regulator | 291 | 246 | 270 | 292 | 82 | 83 | 82 | 87 | 631 | 556 | 540 | 551 |
| K15773 HTH-type transcriptional regulator / antitoxin HipB | 104 | 93 | 86 | 110 | 24 | 32 | 29 | 25 | 230 | 208 | 220 | 242 |
| K15782 Lrp/AsnC family transcriptional regulator, regulator of ectoine-degradation genes | 0 | 0 | 0 | 1 | 0 | 0 | 0 | 0 | 0 | 3 | 0 | 0 |
| K15835 RpiR family transcriptional regulator, murPQ operon repressor | 0 | 1 | 0 | 0 | 12 | 4 | 12 | 10 | 4 | 4 | 7 | 6 |
| K15836 formate hydrogenlyase transcriptional activator | 799 | 820 | 848 | 831 | 249 | 255 | 253 | 244 | 1875 | 1788 | 1830 | 1844 |
| K15973 MarR family transcriptional regulator, 2-MHQ and catechol-resistance regulon repressor | 15 | 26 | 22 | 21 | 3 | 4 | 1 | 7 | 17 | 27 | 15 | 21 |
| K15974 MarR family transcriptional regulator, negative regulator of the multidrug operon emrRAB | 303 | 287 | 284 | 359 | 245 | 224 | 259 | 232 | 357 | 479 | 399 | 491 |
| K16135 LysR family transcriptional regulator, transcriptional activator for dmlA | 340 | 302 | 325 | 347 | 86 | 93 | 102 | 93 | 691 | 658 | 669 | 794 |
| K16136 LacI family transcriptional regulator, maltose regulon regulatory protein | 510 | 526 | 475 | 531 | 350 | 448 | 395 | 348 | 648 | 678 | 622 | 651 |
| K16137 TetR/AcrR family transcriptional regulator, transcriptional repressor for nem operon | 2816 | 2808 | 2603 | 2828 | 1539 | 1488 | 1469 | 1676 | 1666 | 1636 | 1537 | 1655 |
| K16138 TetR/AcrR family transcriptional regulator, repressor for uid operon | 93 | 95 | 97 | 89 | 47 | 33 | 34 | 47 | 291 | 277 | 264 | 300 |
| K16326 CRP/FNR family transcriptional regulator, putaive post-exponential-phase nitrogen-starvation regulator | 82 | 108 | 102 | 81 | 154 | 151 | 160 | 152 | 55 | 36 | 46 | 35 |
| K16516 LysR family transcriptional regulator, regulator for genes of the gallate degradation pathway | 539 | 546 | 459 | 542 | 160 | 143 | 165 | 163 | 1019 | 1031 | 900 | 1061 |
| K17472 Rrf2 family transcriptional regulator, cysteine metabolism repressor | 0 | 0 | 0 | 0 | 0 | 0 | 0 | 0 | 3 | 2 | 3 | 5 |
| K17473 sigma-54 dependent transcriptional regulator, dga operon transcriptional activator | 146 | 150 | 157 | 185 | 19 | 24 | 17 | 22 | 177 | 201 | 180 | 167 |
| K17640 LacI family transcriptional regulator, xylobiose transport system transcriptional regulator | 0 | 0 | 2 | 0 | 0 | 0 | 0 | 0 | 0 | 1 | 0 | 0 |
| K17736 AraC family transcriptional regulator, carnitine catabolism transcriptional activator | 178 | 162 | 181 | 136 | 91 | 69 | 68 | 107 | 64 | 45 | 39 | 52 |
| K17737 LysR family transcriptional regulator, carnitine catabolism transcriptional activator | 1583 | 1494 | 1344 | 1464 | 876 | 885 | 780 | 877 | 960 | 921 | 931 | 891 |
| K18140 TetR/AcrR family transcriptional regulator, acrEF/envCD operon repressor | 248 | 265 | 264 | 277 | 84 | 97 | 93 | 65 | 496 | 534 | 496 | 579 |
| K18294 TetR/AcrR family transcriptional regulator, mexCD-oprJ operon repressor | 375 | 361 | 357 | 379 | 222 | 151 | 212 | 205 | 113 | 123 | 93 | 100 |
| K18297 LysR family transcriptional regulator, mexEF-oprN operon transcriptional activator | 1431 | 1406 | 1343 | 1457 | 770 | 666 | 584 | 661 | 451 | 429 | 367 | 426 |
| K18301 TetR/AcrR family transcriptional regulator, mexJK operon transcriptional repressor | 791 | 755 | 713 | 741 | 303 | 327 | 342 | 355 | 639 | 650 | 581 | 685 |
| K18325 AraC family of transcriptional regulator, multidrug resistance transcriptional activator | 415 | 364 | 421 | 395 | 278 | 318 | 288 | 273 | 545 | 526 | 537 | 501 |
| K18476 TetR/AcrR family transcriptional regulator, tetracycline repressor protein | 0 | 2 | 3 | 0 | 1 | 0 | 0 | 0 | 0 | 0 | 1 | 0 |
| K18531 putative frv operon regulatory protein | 0 | 1 | 0 | 0 | 14 | 12 | 12 | 15 | 0 | 6 | 0 | 7 |
| K18830 HTH-type transcriptional regulator / antitoxin PezA | 3 | 3 | 1 | 1 | 3 | 4 | 3 | 4 | 3 | 10 | 13 | 11 |
| K18831 HTH-type transcriptional regulator / antitoxin HigA | 1513 | 1698 | 1699 | 1754 | 1589 | 1501 | 1509 | 1481 | 1919 | 1935 | 1765 | 1914 |
| K18900 LysR family transcriptional regulator, regulator for bpeEF and oprC | 188 | 239 | 233 | 263 | 46 | 65 | 56 | 49 | 347 | 419 | 357 | 428 |
| K18907 GntR family transcriptional regulator, regulator for abcA and norABC | 12 | 11 | 13 | 11 | 10 | 15 | 13 | 14 | 28 | 27 | 24 | 36 |
| K18909 MarR family transcriptional regulator, repressor for mepA | 0 | 0 | 1 | 0 | 0 | 0 | 0 | 2 | 5 | 1 | 3 | 0 |
| K18918 RHH-type transcriptional regulator, rel operon repressor / antitoxin RelB | 105 | 93 | 93 | 106 | 125 | 119 | 110 | 97 | 4 | 0 | 4 | 7 |

| | | | | | | | | | | | | |
|---|---|---|---|---|---|---|---|---|---|---|---|---|
| K18954 AraC family transcriptional regulator, transcriptional activator of pobA | 1636 | 1678 | 1646 | 1698 | 1068 | 995 | 958 | 1004 | 1139 | 1024 | 1027 | 1111 |
| K18955 WhiB family transcriptional regulator, redox-sensing transcriptional regulator | 0 | 0 | 5 | 5 | 0 | 2 | 1 | 0 | 8 | 4 | 3 | 5 |
| K18991 AraC family transcriptional regulator, activator of mtrCDE | 2147 | 2092 | 2058 | 2063 | 1157 | 1247 | 1101 | 1235 | 1226 | 1255 | 1218 | 1254 |
| K19057 MerR family transcriptional regulator, mercuric resistance operon regulatory protein | 735 | 698 | 748 | 757 | 518 | 535 | 545 | 497 | 202 | 156 | 162 | 170 |
| K19242 LysR family transcriptional regulator, repressor for citA | 36 | 32 | 33 | 35 | 5 | 4 | 2 | 3 | 66 | 57 | 76 | 70 |
| K19333 IclR family transcriptional regulator, KDG regulon repressor | 604 | 566 | 528 | 578 | 405 | 318 | 276 | 321 | 728 | 710 | 693 | 743 |
| K19335 TetR/AcrR family transcriptional regulator, repressor for divergent bdcA | 171 | 199 | 211 | 215 | 57 | 49 | 40 | 52 | 349 | 389 | 394 | 439 |
| K19337 RpiR family transcriptional regulator, carbohydrate utilization regulator | 2487 | 2482 | 2446 | 2409 | 1462 | 1504 | 1405 | 1444 | 1361 | 1258 | 1208 | 1338 |
| K19338 LysR family transcriptional regulator, nitrogen assimilation regulatory protein | 598 | 612 | 559 | 589 | 397 | 428 | 397 | 355 | 786 | 839 | 752 | 865 |
| K19339 NosR/NirI family transcriptional regulator, nitrous oxide reductase regulator | 17 | 11 | 20 | 24 | 11 | 23 | 6 | 7 | 10 | 14 | 16 | 17 |
| K19449 XRE family transcriptional regulator, master regulator for biofilm formation | 3 | 0 | 6 | 3 | 0 | 0 | 2 | 0 | 11 | 12 | 15 | 13 |
| K19505 sigma-54 dependent transcriptional regulator, gfr operon transcriptional activator | 1226 | 1318 | 1189 | 1235 | 348 | 389 | 363 | 329 | 2257 | 2398 | 2388 | 2390 |
| K19575 MerR family transcriptional regulator, activator of bmr gene | 4 | 3 | 11 | 8 | 0 | 0 | 1 | 1 | 3 | 4 | 3 | 1 |
| K19587 Rrf2 family transcriptional regulator, repressor of oqxAB | 309 | 301 | 262 | 321 | 200 | 222 | 181 | 181 | 418 | 374 | 373 | 423 |
| K19591 MerR family transcriptional regulator, copper efflux regulator | 883 | 924 | 823 | 891 | 564 | 507 | 515 | 551 | 558 | 589 | 553 | 522 |
| K19592 MerR family transcriptional regulator, gold-responsive activator of gol and ges genes | 0 | 0 | 0 | 0 | 0 | 0 | 3 | 2 | 0 | 2 | 3 | 1 |
| K19736 TetR/AcrR family transcriptional regulator, regulator of autoinduction and epiphytic fitness | 0 | 0 | 0 | 0 | 0 | 1 | 0 | 0 | 0 | 0 | 1 | 0 |
| K19775 GntR family transcriptional regulator, hexuronate regulon transcriptional repressor | 389 | 421 | 397 | 395 | 343 | 376 | 345 | 325 | 509 | 514 | 541 | 629 |
| K19776 GntR family transcriptional regulator, galactonate operon transcriptional repressor | 459 | 399 | 391 | 417 | 335 | 351 | 332 | 280 | 648 | 680 | 597 | 637 |
| K21405 sigma-54 dependent transcriptional regulator, acetoin dehydrogenase operon transcriptional activator AcoR | 5752 | 5607 | 5087 | 5380 | 2951 | 2989 | 2833 | 3226 | 3039 | 3090 | 2870 | 2998 |
| K21555 CRP/FNR family transcriptional regulator, carbon monoxide oxidation system transcription regulator | 3 | 2 | 5 | 7 | 3 | 4 | 4 | 2 | 4 | 24 | 7 | 6 |
| K21556 CRP/FNR family transcriptional regulator, polysaccharide utilization system transcription regulator | 400 | 442 | 459 | 449 | 1511 | 1418 | 1504 | 1403 | 1017 | 1073 | 1108 | 987 |
| K21557 HTH-type transcriptional regulator, polysaccharide utilization system transcription regulator | 963 | 1144 | 1196 | 1033 | 3593 | 3511 | 3602 | 3562 | 2633 | 2385 | 2519 | 2414 |
| K21562 CRP/FNR family transcriptional regulator, anaerobic regulatory protein | 4 | 2 | 8 | 11 | 20 | 17 | 10 | 17 | 25 | 37 | 31 | 30 |
| K21563 CRP/FNR family transcriptional regulator, dissimilatory nitrate respiration regulator | 463 | 437 | 409 | 390 | 245 | 227 | 210 | 254 | 143 | 108 | 102 | 120 |
| K21564 CRP/FNR family transcriptional regulator, nitrogen oxide reductase regulator | 2 | 2 | 2 | 0 | 0 | 0 | 0 | 0 | 0 | 5 | 0 | 2 |
| K21600 CsoR family transcriptional regulator, copper-sensing transcriptional repressor | 163 | 189 | 152 | 167 | 95 | 93 | 84 | 109 | 219 | 270 | 240 | 254 |
| K21602 IclR family transcriptional regulator, carbohydrate utilization repressor | 216 | 196 | 232 | 205 | 302 | 317 | 291 | 310 | 129 | 114 | 120 | 133 |
| K21645 LysR family transcriptional regulator, hypochlorite-specific transcription factor HypT | 569 | 574 | 557 | 560 | 240 | 268 | 252 | 260 | 761 | 834 | 717 | 817 |
| K21686 XRE family transcriptional regulator, fatty acid utilization regulator | 0 | 0 | 0 | 0 | 2 | 2 | 0 | 0 | 4 | 7 | 2 | 4 |
| K21697 LuxR family transcriptional regulator, quorum-sensing system regulator VjbR | 2 | 0 | 0 | 3 | 0 | 0 | 0 | 0 | 1 | 4 | 3 | 0 |
| K21698 LysR family transcriptional regulator, transcriptional activator for aaeXAB operon | 654 | 569 | 597 | 550 | 405 | 443 | 465 | 430 | 779 | 767 | 789 | 831 |
| K21699 LysR family transcriptional regulator, transcriptional activator for bauABCD operon | 1574 | 1559 | 1439 | 1481 | 865 | 879 | 793 | 902 | 497 | 495 | 436 | 447 |
| K21701 AraC family transcriptional regulator, transcriptional activator for feuABC-ybbA operon | 36 | 35 | 42 | 35 | 8 | 5 | 8 | 10 | 39 | 61 | 33 | 47 |
| K21703 LysR family transcriptional regulator, low CO2-responsive transcriptional regulator | 1 | 1 | 2 | 1 | 0 | 0 | 0 | 0 | 1 | 0 | 2 | 2 |
| K21711 LysR family transcriptional regulator, putative pyruvate carboxylase regulator | 1531 | 1623 | 1536 | 1638 | 926 | 874 | 869 | 982 | 551 | 468 | 536 | 550 |
| K21742 LysR family transcriptional regulator, transcriptional activator for ttdABT operon | 1 | 5 | 5 | 3 | 14 | 12 | 29 | 5 | 51 | 62 | 47 | 58 |
| K21743 MerR family transcriptional regulator, multidrug-efflux activator | 2 | 2 | 2 | 1 | 0 | 0 | 0 | 0 | 4 | 3 | 2 | 1 |
| K21744 MerR family transcriptional regulator, thiopeptide resistance regulator | 8 | 5 | 3 | 2 | 1 | 0 | 1 | 1 | 7 | 3 | 9 | 5 |
| K21745 MerR family transcriptional regulator, aldehyde-responsive regulator | 2 | 1 | 6 | 2 | 0 | 0 | 0 | 0 | 3 | 2 | 8 | 6 |
| K21746 AraC family transcriptional regulator, reactive chlorine species (RCS)-specific activator of rcl operon | 104 | 94 | 126 | 105 | 41 | 29 | 55 | 40 | 125 | 163 | 143 | 184 |
| K21755 LysR family transcriptional regulator, salicylic acid-responsive activator of bsdBCD | 171 | 185 | 186 | 191 | 289 | 299 | 284 | 298 | 5 | 6 | 16 | 10 |
| K21757 LysR family transcriptional regulator, benzoate and cis,cis-muconate-responsive activator of ben and cat genes | 1013 | 1029 | 897 | 1018 | 549 | 441 | 522 | 602 | 346 | 335 | 277 | 321 |
| K21825 AraC family transcriptional regulator, L-arginine-responsive activator | 1454 | 1494 | 1451 | 1452 | 851 | 778 | 675 | 859 | 486 | 456 | 427 | 410 |
| K21826 AraC family transcriptional regulator, glycine betaine-responsive activator | 1634 | 1675 | 1498 | 1590 | 872 | 917 | 769 | 943 | 550 | 462 | 492 | 512 |
| K21828 CRP/FNR family transcriptional regulator, arginine deiminase pathway regulator | 39 | 45 | 37 | 49 | 21 | 15 | 8 | 17 | 43 | 49 | 49 | 49 |
| K21829 D-arginine utilization repressor | 877 | 929 | 956 | 927 | 529 | 461 | 447 | 536 | 314 | 294 | 272 | 282 |
| K21885 ArsR family transcriptional regulator, cadmium/lead-responsive transcriptional repressor | 0 | 0 | 0 | 0 | 0 | 0 | 0 | 0 | 1 | 1 | 1 | 0 |
| K21900 LysR family transcriptional regulator, transcriptional activator of the cysJI operon | 31 | 23 | 42 | 31 | 2 | 4 | 2 | 9 | 47 | 60 | 35 | 54 |
| K21901 LuxR family transcriptional regulator, dicarboxylate transport regulator | 4 | 5 | 2 | 5 | 4 | 5 | 2 | 4 | 31 | 51 | 39 | 32 |
| K21903 ArsR family transcriptional regulator, lead/cadmium/zinc/bismuth-responsive transcriptional repressor | 197 | 185 | 187 | 187 | 100 | 103 | 119 | 123 | 220 | 210 | 177 | 187 |
| K21905 AraC family transcriptional regulator, glutamate-dependent acid resistance regulator | 3 | 1 | 4 | 1 | 10 | 15 | 5 | 4 | 47 | 41 | 31 | 30 |
| K21906 AraC family transcriptional regulator, glutamate-dependent acid resistance regulator | 0 | 0 | 2 | 1 | 11 | 9 | 5 | 10 | 64 | 68 | 42 | 57 |
| K21907 LuxR family transcriptional regulator, glutamate-dependent acid resistance regulator | 1 | 2 | 2 | 0 | 9 | 1 | 11 | 5 | 23 | 28 | 40 | 34 |
| K21959 LysR family transcriptional regulator, cell division regulator | 1 | 2 | 2 | 2 | 0 | 0 | 1 | 0 | 0 | 0 | 0 | 0 |
| K21960 LysR family transcriptional regulator, regulator of the ytmI operon | 0 | 2 | 3 | 3 | 0 | 0 | 1 | 0 | 0 | 2 | 4 | 0 |
| K21961 TetR/AcrR family transcriptional regulator, ethionamide resistance regulator | 107 | 117 | 101 | 102 | 0 | 4 | 0 | 0 | 59 | 62 | 77 | 74 |
| K21963 LuxR family transcriptional regulator, Mat/Ecp fimbriae transcriptional regulator | 106 | 114 | 104 | 104 | 37 | 32 | 35 | 51 | 243 | 224 | 210 | 266 |
| K21972 MerR family transcriptional regulator, repressor of blue light- and temperature-responsive genes | 436 | 369 | 387 | 442 | 348 | 342 | 328 | 282 | 572 | 583 | 559 | 567 |
| K22041 TetR/AcrR family transcriptional regulator, copper-responsive repressor | 338 | 370 | 358 | 351 | 228 | 257 | 291 | 240 | 521 | 626 | 596 | 621 |
| K22043 ArsR family transcriptional regulator, zinc-responsive transcriptional repressor | 21 | 18 | 8 | 17 | 10 | 9 | 8 | 8 | 21 | 25 | 34 | 27 |
| K22103 DeoR family transcriptional regulator, carbon catabolite repression regulator | 49 | 40 | 44 | 45 | 3 | 15 | 15 | 7 | 47 | 40 | 50 | 36 |
| K22104 GntR family transcriptional regulator, sialic acid-inducible nan operon repressor | 225 | 178 | 185 | 209 | 261 | 261 | 250 | 267 | 67 | 64 | 75 | 74 |
| K22105 TetR/AcrR family transcriptional regulator, fatty acid biosynthesis regulator | 440 | 451 | 406 | 447 | 309 | 371 | 337 | 313 | 596 | 592 | 591 | 581 |
| K22106 TetR/AcrR family transcriptional regulator, repressor of fatR-cypB operon | 11 | 23 | 12 | 7 | 8 | 21 | 10 | 4 | 7 | 29 | 26 | 34 |
| K22108 TetR/AcrR family transcriptional regulator, cholesterol catabolism regulator | 7 | 2 | 2 | 4 | 2 | 7 | 3 | 3 | 17 | 17 | 18 | 13 |
| K22293 GntR family transcriptional regulator, rspAB operon transcriptional repressor | 491 | 540 | 528 | 516 | 277 | 276 | 273 | 245 | 594 | 617 | 627 | 577 |
| K22296 MarR family transcriptional regulator, lower aerobic nicotinate degradation pathway regulator | 639 | 660 | 636 | 621 | 310 | 295 | 286 | 317 | 176 | 158 | 202 | 175 |
| K22299 HTH-type transcriptional regulator, competence development regulator | 1 | 1 | 0 | 1 | 0 | 0 | 0 | 0 | 1 | 1 | 0 | 4 |
| K22300 HTH-type transcriptional regulator, cell division transcriptional repressor | 73 | 51 | 38 | 87 | 93 | 110 | 103 | 108 | 2 | 4 | 1 | 0 |
| K22302 transcriptional repressor of cell division inhibition gene dicB | 54 | 61 | 59 | 58 | 73 | 76 | 87 | 73 | 6 | 13 | 10 | 12 |
| K22486 transcriptional regulator HilA, main transcriptional regulator of SPI1 | 0 | 0 | 0 | 0 | 14 | 7 | 24 | 16 | 7 | 2 | 5 | 4 |
| K22489 MarR family transcriptional regulator, temperature-dependent positive regulator of motility | 264 | 266 | 242 | 243 | 146 | 195 | 174 | 184 | 261 | 356 | 309 | 325 |
| K22491 MerR family transcriptional regulator, light-induced transcriptional regulator | 1562 | 1494 | 1423 | 1382 | 828 | 789 | 719 | 868 | 451 | 430 | 423 | 420 |
| K22650 luxR family transcriptional regulator, regulator of transport and utilization of aryl beta-glucosides | 376 | 417 | 387 | 427 | 309 | 289 | 331 | 288 | 539 | 556 | 538 | 615 |
| K23015 GntR family transcriptional regulator, transcriptional activator for L-galactonate catabolism | 226 | 253 | 202 | 216 | 277 | 304 | 301 | 273 | 47 | 49 | 64 | 57 |
| K23237 AraC family transcriptional regulator, melibiose operon regulatory protein | 383 | 424 | 318 | 418 | 115 | 129 | 111 | 99 | 845 | 923 | 815 | 913 |
| K23238 RpiR family transcriptional regulator, repressor of rpiB and als operon | 6 | 2 | 6 | 8 | 6 | 3 | 3 | 3 | 50 | 70 | 58 | 68 |
| K23239 FrmR/RcnR family transcriptional regulator, repressor of frmRAB operon | 188 | 157 | 157 | 177 | 89 | 131 | 94 | 114 | 211 | 221 | 186 | 200 |
| K23240 FrmR/RcnR family transcriptional regulator, repressor of rcnA expression | 152 | 188 | 193 | 182 | 143 | 105 | 129 | 96 | 264 | 236 | 278 | 254 |
| K23253 PucR family transcriptional regulator, proline-responsive transcriptional activator | 0 | 1 | 0 | 0 | 0 | 0 | 0 | 0 | 4 | 1 | 0 | 3 |
| K23356 HTH-type transcriptional regulator, sugar sensing transcriptional regulator | 100 | 85 | 99 | 96 | 14 | 14 | 5 | 1 | 134 | 134 | 177 | 149 |
| ko03021 Transcription machinery | 91360 | 91530 | 92565 | 91532 | 99939 | 100209 | 100092 | 99903 | 93164 | 93260 | 93398 | 91380 |
| K02600 transcription termination/antitermination protein NusA | 3852 | 3876 | 4055 | 3982 | 4996 | 5027 | 5018 | 4994 | 4688 | 4834 | 4430 | 4619 |
| K02601 transcription termination/antitermination protein NusG | 1699 | 1740 | 1750 | 1844 | 2017 | 1937 | 2063 | 1895 | 1895 | 2097 | 2048 | 1954 |
| K03057 transcription factor S | 0 | 0 | 0 | 0 | 2 | 0 | 0 | 1 | 3 | 1 | 3 | 4 |
| K03086 RNA polymerase primary sigma factor | 4796 | 4817 | 4923 | 4757 | 4180 | 4189 | 3963 | 3918 | 5070 | 5076 | 5251 | 5143 |
| K03088 RNA polymerase sigma-70 factor, ECF subfamily | 10471 | 10767 | 11039 | 10236 | 19908 | 19856 | 20345 | 20347 | 14507 | 13939 | 14524 | 13727 |
| K03089 RNA polymerase sigma-32 factor | 2116 | 1990 | 2008 | 1957 | 1295 | 1174 | 1166 | 1295 | 1234 | 1216 | 1223 | 1243 |
| K03090 RNA polymerase sigma-B factor | 9 | 11 | 19 | 17 | 10 | 18 | 23 | 9 | 22 | 25 | 13 | 26 |
| K03091 RNA polymerase sporulation-specific sigma factor | 1368 | 1381 | 1413 | 1465 | 440 | 513 | 549 | 463 | 1341 | 1642 | 1383 | 1469 |
| K03093 RNA polymerase sigma factor | 15 | 21 | 16 | 20 | 6 | 2 | 3 | 2 | 15 | 16 | 13 | 14 |
| K03580 ATP-dependent helicase HepA [EC:3.6.4.-] | 5931 | 5578 | 5652 | 5672 | 3367 | 3260 | 3305 | 3401 | 3801 | 3939 | 3582 | 3725 |
| K03597 sigma-E factor negative regulatory protein RseA | 1367 | 1357 | 1251 | 1286 | 825 | 849 | 753 | 801 | 862 | 885 | 745 | 818 |
| K03598 sigma-E factor negative regulatory protein RseB | 1815 | 1772 | 1773 | 1862 | 1129 | 1112 | 1076 | 1139 | 1118 | 1136 | 1112 | 1136 |
| K03599 stringent starvation protein A | 627 | 650 | 684 | 619 | 433 | 480 | 485 | 490 | 623 | 644 | 605 | 619 |
| K03600 stringent starvation protein B | 909 | 870 | 854 | 929 | 597 | 570 | 540 | 480 | 584 | 631 | 612 | 636 |
| K03624 transcription elongation factor GreA | 1361 | 1416 | 1469 | 1489 | 1704 | 1734 | 1819 | 1771 | 1903 | 2010 | 1963 | 1927 |
| K03625 transcription antitermination protein NusB | 1827 | 1955 | 1773 | 1809 | 2594 | 2555 | 2623 | 2622 | 2472 | 2152 | 2232 | 2204 |
| K03803 sigma-E factor negative regulatory protein RseC | 1193 | 1252 | 1206 | 1194 | 1374 | 1356 | 1318 | 1361 | 1118 | 1300 | 1176 | 1128 |
| K04749 anti-sigma B factor antagonist | 28 | 14 | 23 | 19 | 7 | 6 | 7 | 6 | 7 | 15 | 10 | 17 |
| K04760 transcription elongation factor GreB | 1196 | 1112 | 1079 | 1057 | 655 | 678 | 655 | 604 | 739 | 715 | 687 | 803 |
| K07315 phosphoserine phosphatase RsbU/P [EC:3.1.3.3] | 314 | 301 | 294 | 310 | 104 | 74 | 73 | 93 | 253 | 245 | 212 | 273 |
| K11367 chromodomain-helicase-DNA-binding protein 1 [EC:3.6.4.12] | 0 | 0 | 0 | 2 | 0 | 0 | 0 | 0 | 0 | 0 | 0 | 0 |
| K14845 RAT1-interacting protein | 0 | 0 | 0 | 0 | 1 | 0 | 0 | 0 | 0 | 0 | 0 | 0 |
| K19000 Rho-binding antiterminator | 691 | 706 | 688 | 728 | 438 | 415 | 413 | 427 | 402 | 383 | 399 | 387 |
| ko03019 Messenger RNA biogenesis | 85737 | 86214 | 86968 | 86768 | 87635 | 89767 | 88744 | 87845 | 86439 | 87295 | 86668 | 86568 |
| K02553 regulator of ribonuclease activity A | 1094 | 1047 | 1058 | 1070 | 610 | 699 | 597 | 656 | 649 | 682 | 650 | 746 |
| K03698 3'-5' exoribonuclease [EC:3.1.-.-] | 699 | 771 | 717 | 749 | 197 | 218 | 203 | 201 | 705 | 783 | 718 | 778 |
| K06958 RNase adapter protein RapZ | 2133 | 2021 | 1891 | 2018 | 1151 | 1189 | 1254 | 1178 | 1477 | 1667 | 1552 | 1633 |
| K08301 ribonuclease G [EC:3.1.26.-] | 4036 | 4064 | 4091 | 4058 | 4447 | 4957 | 4809 | 4779 | 4419 | 4149 | 4240 | 4181 |
| K09893 regulator of ribonuclease activity B | 269 | 254 | 236 | 306 | 165 | 213 | 190 | 161 | 413 | 422 | 368 | 356 |
| K10357 myosin V | 0 | 0 | 0 | 0 | 0 | 0 | 1 | 0 | 0 | 2 | 0 | 0 |
| K11493 regulator of chromosome condensation | 1 | 0 | 0 | 0 | 0 | 0 | 0 | 0 | 0 | 0 | 0 | 0 |
| K18692 ATP-dependent RNA helicase CshB [EC:3.6.4.13] | 63 | 58 | 70 | 60 | 20 | 32 | 14 | 17 | 80 | 103 | 77 | 63 |
| K18765 RNase E specificity factor CsrD | 1201 | 1178 | 1107 | 1260 | 779 | 908 | 878 | 865 | 1598 | 1676 | 1591 | 1669 |
| ko03041 Spliceosome | 19 | 17 | 19 | 12 | 0 | 2 | 2 | 0 | 30 | 31 | 32 | 43 |
| K12736 peptidylprolyl isomerase domain and WD repeat-containing protein 1 [EC:5.2.1.8] | 0 | 0 | 0 | 0 | 0 | 0 | 0 | 0 | 0 | 0 | 2 | 0 |
| K13177 ATP-dependent RNA helicase DDX1 [EC:3.6.4.13] | 0 | 0 | 0 | 0 | 0 | 0 | 0 | 0 | 1 | 0 | 0 | 0 |
| K13179 ATP-dependent RNA helicase DDX18/HAS1 [EC:3.6.4.13] | 1 | 0 | 0 | 0 | 0 | 0 | 0 | 0 | 0 | 0 | 0 | 0 |
| K17043 ATP-dependent RNA helicase DDX43 [EC:3.6.4.13] | 0 | 0 | 0 | 0 | 0 | 0 | 0 | 0 | 0 | 1 | 0 | 0 |

| | | | | | | | | | | | | |
|---|---|---|---|---|---|---|---|---|---|---|---|---|
| ko03011 Ribosome | 78384 | 79588 | 82698 | 81135 | 88612 | 88928 | 90151 | 86988 | 84998 | 85396 | 85577 | 85767 |
| K17437 large subunit ribosomal protein L15 | 0 | 2 | 0 | 0 | 0 | 0 | 0 | 0 | 0 | 0 | 0 | 0 |
| ko03009 Ribosome biogenesis | 156988 | 157286 | 156854 | 157650 | 151246 | 152400 | 152627 | 152332 | 156586 | 157365 | 155923 | 156824 |
| K00561 23S rRNA (adenine-N6)-dimethyltransferase [EC:2.1.1.184] | 387 | 525 | 544 | 531 | 1488 | 1520 | 1652 | 1535 | 1109 | 1014 | 1086 | 1057 |
| K00563 23S rRNA (guanine745-N1)-methyltransferase [EC:2.1.1.187] | 1715 | 1771 | 1581 | 1725 | 1103 | 1081 | 962 | 1076 | 1131 | 1173 | 1109 | 1027 |
| K00564 16S rRNA (guanine1207-N2)-methyltransferase [EC:2.1.1.172] | 2307 | 2357 | 2243 | 2350 | 1472 | 1287 | 1285 | 1411 | 1444 | 1447 | 1446 | 1554 |
| K00783 23S rRNA (pseudouridine1915-N3)-methyltransferase [EC:2.1.1.177] | 1706 | 1733 | 1758 | 1731 | 1843 | 1790 | 1747 | 1937 | 1801 | 1795 | 1750 | 1732 |
| K02427 23S rRNA (uridine2552-2'-O)-methyltransferase [EC:2.1.1.166] | 1198 | 1119 | 1212 | 1206 | 791 | 820 | 765 | 800 | 883 | 905 | 838 | 904 |
| K02528 16S rRNA (adenine1518-N6/adenine1519-N6)-dimethyltransferase [EC:2.1.1.182] | 2840 | 2744 | 2843 | 2791 | 2884 | 2837 | 2986 | 2976 | 2859 | 2919 | 2871 | 2888 |
| K02687 ribosomal protein L11 methyltransferase [EC:2.1.1.-] | 3044 | 3116 | 3033 | 2853 | 2949 | 3191 | 3011 | 2864 | 2822 | 3033 | 2822 | 2906 |
| K02834 ribosome-binding factor A | 1415 | 1372 | 1395 | 1364 | 1406 | 1428 | 1422 | 1405 | 1434 | 1519 | 1452 | 1430 |
| K02860 16S rRNA processing protein RimM | 1884 | 1769 | 1911 | 1727 | 1912 | 1957 | 2016 | 1936 | 1810 | 1803 | 1780 | 1895 |
| K02972 stationary-phase-induced ribosome-associated protein | 83 | 100 | 72 | 99 | 54 | 60 | 75 | 64 | 114 | 115 | 141 | 136 |
| K03212 23S rRNA (uracil747-C5)-methyltransferase [EC:2.1.1.189] | 667 | 660 | 627 | 634 | 388 | 396 | 420 | 410 | 898 | 914 | 864 | 861 |
| K03214 RNA methyltransferase, TrmH family [EC:2.1.1.-] | 603 | 602 | 632 | 619 | 444 | 438 | 490 | 467 | 852 | 921 | 819 | 912 |
| K03215 23S rRNA (uracil1939-C5)-methyltransferase [EC:2.1.1.190] | 3738 | 3895 | 3815 | 3824 | 2919 | 2987 | 2903 | 2860 | 3392 | 3412 | 3256 | 3518 |
| K03218 23S rRNA (guanosine2251-2'-O)-methyltransferase [EC:2.1.1.185] | 2222 | 2177 | 2222 | 2212 | 2608 | 2628 | 2459 | 2465 | 2464 | 2569 | 2542 | 2530 |
| K03438 16S rRNA (cytosine1402-N4)-methyltransferase [EC:2.1.1.199] | 3676 | 3332 | 3688 | 3503 | 3563 | 3629 | 3523 | 3542 | 3202 | 3291 | 3425 | 3279 |
| K03500 16S rRNA (cytosine967-C5)-methyltransferase [EC:2.1.1.176] | 3831 | 3714 | 3490 | 3820 | 2201 | 2162 | 2065 | 2225 | 2557 | 2671 | 2748 | 2654 |
| K03501 16S rRNA (guanine527-N7)-methyltransferase [EC:2.1.1.170] | 2268 | 2318 | 2287 | 2254 | 2086 | 2248 | 2229 | 2172 | 1902 | 1924 | 2016 | 1886 |
| K03595 GTPase | 2743 | 2781 | 2980 | 2758 | 3136 | 3332 | 3297 | 3167 | 3167 | 3242 | 3250 | 3196 |
| K03665 GTPase | 4234 | 4324 | 4318 | 4224 | 4706 | 4776 | 4748 | 4865 | 4318 | 4320 | 4511 | 4365 |
| K03789 [ribosomal protein S18]-alanine N-acetyltransferase [EC:2.3.1.266] | 1433 | 1342 | 1252 | 1394 | 701 | 706 | 750 | 798 | 933 | 1000 | 977 | 925 |
| K03790 [ribosomal protein S5]-alanine N-acetyltransferase [EC:2.3.1.267] | 1739 | 1768 | 1748 | 1725 | 919 | 863 | 894 | 971 | 1371 | 1368 | 1392 | 1449 |
| K03812 ribosome modulation factor | 317 | 334 | 336 | 327 | 198 | 210 | 196 | 201 | 197 | 191 | 210 | 219 |
| K03817 ribosomal-protein-serine acetyltransferase [EC:2.3.1.-] | 891 | 884 | 826 | 816 | 445 | 484 | 465 | 486 | 617 | 578 | 563 | 603 |
| K03977 GTPase | 3157 | 3311 | 3353 | 3288 | 4117 | 4545 | 4325 | 4196 | 4472 | 4614 | 4647 | 4528 |
| K03979 GTPase [EC:3.6.5.-] | 3851 | 3708 | 3807 | 3814 | 4099 | 3948 | 4206 | 4146 | 3938 | 4054 | 3870 | 3836 |
| K04795 fibrillarin-like pre-rRNA processing protein | 0 | 0 | 0 | 0 | 0 | 0 | 0 | 2 | 0 | 2 | 2 | 4 |
| K05590 ATP-dependent RNA helicase SrmB [EC:3.6.4.13] | 697 | 702 | 673 | 744 | 555 | 507 | 453 | 526 | 1047 | 1027 | 952 | 1046 |
| K05591 ATP-independent RNA helicase DbpA [EC:3.6.4.13] | 1614 | 1530 | 1575 | 1627 | 1080 | 1110 | 1032 | 1125 | 1311 | 1362 | 1322 | 1465 |
| K05808 putative sigma-54 modulation protein | 1176 | 1125 | 1165 | 1177 | 1177 | 1091 | 1125 | 1208 | 1105 | 1153 | 1164 | 1192 |
| K05809 ribosome-associated inhibitor A | 241 | 213 | 201 | 194 | 155 | 196 | 167 | 144 | 315 | 337 | 333 | 358 |
| K05844 ribosomal protein S6--L-glutamate ligase [EC:6.3.2.-] | 1068 | 1145 | 1117 | 1151 | 782 | 760 | 788 | 736 | 967 | 943 | 954 | 895 |
| K06174 ATP-binding cassette, sub-family E, member 1 | 3 | 0 | 0 | 1 | 0 | 0 | 2 | 2 | 4 | 16 | 20 | 7 |
| K06177 tRNA pseudouridine32 synthase / 23S rRNA pseudouridine746 synthase [EC:5.4.99.28 5.4.99.29] | 2909 | 2844 | 2782 | 2807 | 3718 | 3804 | 3797 | 3649 | 3400 | 3258 | 3295 | 3219 |
| K06178 23S rRNA pseudouridine2605 synthase [EC:5.4.99.22] | 776 | 825 | 836 | 837 | 672 | 727 | 914 | 986 | 1026 | 1058 | 966 | 1054 |
| K06179 23S rRNA pseudouridine955/2504/2580 synthase [EC:5.4.99.24] | 2298 | 2421 | 2435 | 2420 | 1432 | 1486 | 1360 | 1382 | 1738 | 1676 | 1669 | 1669 |
| K06180 23S rRNA pseudouridine1911/1915/1917 synthase [EC:5.4.99.23] | 4835 | 4864 | 5154 | 5023 | 7249 | 7418 | 7297 | 7164 | 6469 | 6289 | 6697 | 6448 |
| K06181 23S rRNA pseudouridine2457 synthase [EC:5.4.99.20] | 1468 | 1385 | 1289 | 1334 | 812 | 761 | 657 | 754 | 881 | 847 | 828 | 882 |
| K06182 23S rRNA pseudouridine2604 synthase [EC:5.4.99.21] | 1838 | 1787 | 1745 | 1838 | 939 | 997 | 924 | 1067 | 1365 | 1263 | 1300 | 1261 |
| K06183 16S rRNA pseudouridine516 synthase [EC:5.4.99.19] | 1702 | 1784 | 1681 | 1669 | 1029 | 986 | 1031 | 1011 | 1247 | 1161 | 1208 | 1283 |
| K06442 23S rRNA (cytidine1920-2'-O)/16S rRNA (cytidine1409-2'-O)-methyltransferase [EC:2.1.1.226 2.1.1.227] | 506 | 462 | 464 | 530 | 172 | 176 | 156 | 171 | 521 | 568 | 491 | 537 |
| K06941 23S rRNA (adenine2503-C2)-methyltransferase [EC:2.1.1.192] | 3673 | 3546 | 3857 | 3715 | 3873 | 3928 | 3996 | 4010 | 4139 | 4110 | 4153 | 4117 |
| K06942 ribosome-binding ATPase | 3863 | 4025 | 4076 | 4048 | 4638 | 4857 | 4673 | 4580 | 4149 | 4293 | 4458 | 4260 |
| K06948 30S ribosome assembly GTPase | 73 | 69 | 65 | 63 | 14 | 19 | 33 | 27 | 70 | 97 | 64 | 77 |
| K06961 ribosomal RNA assembly protein | 0 | 0 | 0 | 1 | 2 | 0 | 0 | 0 | 2 | 3 | 3 | 10 |
| K06968 23S rRNA (cytidine2498-2'-O)-methyltransferase [EC:2.1.1.186] | 1380 | 1378 | 1330 | 1370 | 782 | 839 | 804 | 843 | 1156 | 1178 | 1200 | 1160 |
| K06969 23S rRNA (cytosine1962-C5)-methyltransferase [EC:2.1.1.191] | 5367 | 5358 | 5434 | 5248 | 4752 | 4722 | 5108 | 4930 | 4377 | 4525 | 4490 | 4397 |
| K06970 23S rRNA (adenine1618-N6)-methyltransferase [EC:2.1.1.181] | 1050 | 1094 | 1058 | 1061 | 2195 | 2005 | 2242 | 2102 | 2139 | 2097 | 2165 | 2200 |
| K07042 probable rRNA maturation factor | 1168 | 1132 | 1099 | 1115 | 689 | 670 | 647 | 694 | 897 | 866 | 814 | 960 |
| K07056 16S rRNA (cytidine1402-2'-O)-methyltransferase [EC:2.1.1.198] | 2360 | 2439 | 2542 | 2491 | 3846 | 3812 | 3961 | 3827 | 3485 | 3378 | 3588 | 3534 |
| K07115 23S rRNA (adenine2030-N6)-methyltransferase [EC:2.1.1.266] | 2488 | 2585 | 2502 | 2484 | 3029 | 3112 | 3006 | 3140 | 2672 | 2410 | 2452 | 2559 |
| K07320 ribosomal protein L3 glutamine methyltransferase [EC:2.1.1.298] | 950 | 934 | 854 | 1025 | 637 | 610 | 614 | 582 | 944 | 1003 | 989 | 965 |
| K07566 L-threonylcarbamoyladenylate synthase [EC:2.7.7.87] | 2186 | 2058 | 2045 | 2078 | 2294 | 2129 | 2321 | 2396 | 2386 | 2436 | 2375 | 2350 |
| K07569 RNA-binding protein | 2 | 2 | 0 | 1 | 0 | 0 | 0 | 0 | 1 | 5 | 0 | 1 |
| K07574 RNA-binding protein | 909 | 947 | 924 | 944 | 594 | 547 | 522 | 484 | 636 | 617 | 568 | 612 |
| K08316 16S rRNA (guanine966-N2)-methyltransferase [EC:2.1.1.171] | 1305 | 1275 | 1174 | 1206 | 781 | 726 | 734 | 779 | 855 | 865 | 821 | 898 |
| K09136 ribosomal protein S12 methylthiotransferase accessory factor | 1723 | 1553 | 1572 | 1660 | 1002 | 1086 | 1117 | 946 | 1650 | 1751 | 1604 | 1749 |
| K09140 pre-rRNA-processing protein TSR3 | 0 | 0 | 0 | 0 | 0 | 0 | 0 | 0 | 0 | 3 | 0 | 6 |
| K09710 ribosome-associated protein | 972 | 986 | 1121 | 1041 | 1256 | 1132 | 1270 | 1204 | 1169 | 1183 | 1182 | 1151 |
| K09748 ribosome maturation factor RimP | 1563 | 1659 | 1778 | 1699 | 1892 | 1865 | 1858 | 1850 | 1638 | 1709 | 1759 | 1612 |
| K09761 16S rRNA (uracil1498-N3)-methyltransferase [EC:2.1.1.193] | 2612 | 2711 | 2533 | 2473 | 2778 | 2717 | 2697 | 2762 | 2569 | 2778 | 2569 | 2638 |
| K09889 ribosome-associated protein | 1107 | 1103 | 1070 | 1132 | 662 | 656 | 681 | 701 | 807 | 719 | 684 | 763 |
| K11145 ribonuclease III family protein [EC:3.1.26.-] | 192 | 218 | 210 | 209 | 71 | 48 | 71 | 53 | 215 | 214 | 188 | 238 |
| K11391 23S rRNA (guanine1835-N2)-methyltransferase [EC:2.1.1.174] | 1882 | 1977 | 1885 | 1922 | 1237 | 1187 | 1182 | 1239 | 1438 | 1342 | 1361 | 1447 |
| K11392 16S rRNA (cytosine1407-C5)-methyltransferase [EC:2.1.1.178] | 771 | 771 | 762 | 835 | 515 | 538 | 575 | 556 | 989 | 1089 | 1088 | 1174 |
| K12297 23S rRNA (guanine2445-N2)-methyltransferase / 23S rRNA (guanine2069-N7)-methyltransferase [EC:2.1.1.173 2.1.1.264] | 3654 | 3731 | 3517 | 3678 | 2190 | 2156 | 2110 | 2054 | 2569 | 2571 | 2568 | 2485 |
| K14441 ribosomal protein S12 methylthiotransferase [EC:2.8.4.4] | 3914 | 3764 | 3882 | 3909 | 4323 | 4251 | 4270 | 4288 | 3926 | 3895 | 3793 | 3982 |
| K14540 ribosome biogenesis GTPase A | 439 | 432 | 442 | 416 | 160 | 155 | 180 | 157 | 496 | 476 | 435 | 490 |
| K14761 ribosome-associated protein | 264 | 280 | 292 | 269 | 160 | 156 | 172 | 156 | 357 | 334 | 368 | 392 |
| K14762 ribosome-associated protein | 191 | 199 | 187 | 186 | 180 | 165 | 152 | 149 | 263 | 243 | 272 | 244 |
| K14768 U3 small nucleolar RNA-associated protein 7 | 0 | 0 | 0 | 0 | 0 | 0 | 0 | 0 | 0 | 1 | 0 | 0 |
| K14779 ATP-dependent RNA helicase DDX52/ROK1 [EC:3.6.4.13] | 0 | 4 | 0 | 0 | 0 | 0 | 0 | 0 | 0 | 0 | 0 | 0 |
| K14782 protein AATF/BFR2 | 0 | 0 | 0 | 1 | 0 | 0 | 0 | 0 | 0 | 0 | 0 | 0 |
| K14797 essential nuclear protein 1 | 0 | 0 | 0 | 0 | 0 | 0 | 0 | 0 | 0 | 0 | 0 | 1 |
| K14810 ATP-dependent RNA helicase DDX56/DBP9 [EC:3.6.4.13] | 0 | 0 | 0 | 0 | 0 | 0 | 0 | 0 | 0 | 0 | 1 | 0 |
| K14811 ATP-dependent RNA helicase DBP3 [EC:3.6.4.13] | 0 | 0 | 2 | 0 | 0 | 0 | 0 | 0 | 0 | 0 | 0 | 0 |
| K14831 protein MAK16 | 0 | 0 | 0 | 0 | 0 | 0 | 0 | 0 | 0 | 2 | 0 | 0 |
| K14832 ribosome biogenesis protein MAK21 | 0 | 0 | 0 | 0 | 0 | 0 | 0 | 0 | 0 | 0 | 2 | 0 |
| K14846 ribosome production factor 1 | 0 | 2 | 0 | 0 | 0 | 0 | 0 | 0 | 0 | 0 | 0 | 0 |
| K14859 ribosome biogenesis protein SSF1/2 | 0 | 0 | 0 | 0 | 0 | 0 | 0 | 0 | 0 | 0 | 0 | 1 |
| K15508 21S rRNA (uridine2791-2'-O)-methyltransferase [EC:2.1.1.168] | 0 | 0 | 0 | 0 | 0 | 0 | 0 | 0 | 0 | 2 | 0 | 0 |
| K15632 23S rRNA (adenine-C8)-methyltransferase [EC:2.1.1.224] | 6 | 5 | 6 | 4 | 1 | 1 | 1 | 0 | 24 | 24 | 12 | 17 |
| K15984 16S rRNA (guanine1516-N2)-methyltransferase [EC:2.1.1.242] | 1978 | 1901 | 1731 | 1854 | 1159 | 1043 | 1029 | 1072 | 1251 | 1240 | 1198 | 1181 |
| K18850 50S ribosomal protein L16 3-hydroxylase [EC:1.14.11.47] | 2001 | 2026 | 2005 | 2115 | 1361 | 1375 | 1365 | 1313 | 2059 | 2221 | 1922 | 2059 |
| K20224 importin-9 | 0 | 2 | 0 | 0 | 0 | 0 | 0 | 0 | 0 | 0 | 0 | 0 |
| K21514 23S rRNA (uridine2479-2'-O)-methyltransferase [EC:2.1.1.208] | 3 | 0 | 2 | 3 | 0 | 0 | 0 | 0 | 3 | 2 | 0 | 1 |
| ko03016 Transfer RNA biogenesis | 302875 | 303504 | 304679 | 304679 | 300276 | 304600 | 302449 | 299449 | 299014 | 301908 | 298864 | 300575 |
| K00554 tRNA (guanine37-N1)-methyltransferase [EC:2.1.1.228] | 2309 | 2356 | 2488 | 2380 | 2298 | 2431 | 2449 | 2318 | 2469 | 2459 | 2568 | 2435 |
| K00555 tRNA (guanine26-N2/guanine27-N2)-dimethyltransferase [EC:2.1.1.215 2.1.1.216] | 0 | 0 | 3 | 1 | 0 | 0 | 0 | 2 | 1 | 10 | 6 | 12 |
| K00556 tRNA (guanosine-2'-O-)-methyltransferase [EC:2.1.1.34] | 501 | 449 | 479 | 489 | 280 | 389 | 320 | 331 | 622 | 713 | 666 | 732 |
| K00557 tRNA (uracil-5-)-methyltransferase [EC:2.1.1.35] | 2379 | 2429 | 2258 | 2376 | 1396 | 1418 | 1365 | 1391 | 1479 | 1487 | 1401 | 1410 |
| K00773 queuine tRNA-ribosyltransferase [EC:2.4.2.29] | 3471 | 3449 | 3732 | 3538 | 3572 | 3588 | 3801 | 3557 | 3601 | 3978 | 3763 | 3837 |
| K00989 ribonuclease PH [EC:2.7.7.56] | 1901 | 1809 | 1898 | 1739 | 1220 | 1198 | 1143 | 1142 | 1229 | 1153 | 1142 | 1249 |
| K01147 exoribonuclease II [EC:3.1.13.1] | 1254 | 1236 | 1101 | 1171 | 947 | 1022 | 943 | 947 | 1731 | 1792 | 1685 | 1739 |
| K01166 ribonuclease T2 [EC:4.6.1.19] | 1067 | 1102 | 1176 | 997 | 638 | 593 | 582 | 571 | 354 | 350 | 328 | 354 |
| K01169 ribonuclease I (enterobacter ribonuclease) [EC:4.6.1.21] | 471 | 446 | 490 | 435 | 333 | 335 | 331 | 284 | 604 | 637 | 624 | 689 |
| K01170 tRNA-intron endonuclease, archaea type [EC:4.6.1.16] | 1 | 0 | 0 | 0 | 0 | 2 | 0 | 2 | 0 | 0 | 3 | 1 |
| K01409 N6-L-threonylcarbamoyladenine synthase [EC:2.3.1.234] | 3919 | 3728 | 3748 | 3765 | 3464 | 3511 | 3381 | 3491 | 3469 | 3477 | 3510 | 3602 |
| K01855 tRNA pseudouridine38/39 synthase [EC:5.4.99.45] | 0 | 0 | 0 | 0 | 1 | 0 | 0 | 0 | 0 | 0 | 0 | 0 |
| K01975 RNA 2',3'-cyclic 3'-phosphodiesterase [EC:3.1.4.58] | 1238 | 1253 | 1214 | 1248 | 683 | 746 | 618 | 735 | 827 | 746 | 735 | 727 |
| K02533 tRNA/rRNA methyltransferase [EC:2.1.1.-] | 412 | 523 | 544 | 475 | 358 | 418 | 394 | 338 | 744 | 682 | 717 | 750 |
| K03177 tRNA pseudouridine55 synthase [EC:5.4.99.25] | 3185 | 3226 | 3247 | 3218 | 3285 | 3292 | 3270 | 3183 | 3147 | 3086 | 3099 | 2940 |
| K03216 tRNA (cytidine/uridine-2'-O-)-methyltransferase [EC:2.1.1.207] | 1502 | 1471 | 1452 | 1472 | 794 | 819 | 819 | 739 | 1133 | 1074 | 1011 | 1135 |
| K03437 RNA methyltransferase, TrmH family | 895 | 875 | 993 | 828 | 1444 | 1530 | 1492 | 1379 | 1365 | 1390 | 1497 | 1401 |
| K03439 tRNA (guanine-N7-)-methyltransferase [EC:2.1.1.33] | 2645 | 2568 | 2538 | 2650 | 2927 | 3203 | 3055 | 3025 | 2691 | 2661 | 2640 | 2658 |
| K03495 tRNA uridine 5-carboxymethylaminomethyl modification enzyme | 4585 | 4452 | 4727 | 4489 | 6370 | 6530 | 6640 | 6370 | 6461 | 6698 | 6582 | 6403 |
| K03536 ribonuclease P protein component [EC:3.1.26.5] | 1390 | 1400 | 1413 | 1404 | 1515 | 1574 | 1557 | 1644 | 1507 | 1587 | 1556 | 1538 |
| K03650 tRNA modification GTPase [EC:3.6.-.-] | 5245 | 5145 | 5334 | 5165 | 5227 | 5432 | 5572 | 5267 | 5242 | 5197 | 5115 | 5092 |
| K03683 ribonuclease T [EC:3.1.13.-] | 1206 | 1177 | 1123 | 1232 | 658 | 680 | 608 | 601 | 821 | 783 | 775 | 743 |
| K03684 ribonuclease D [EC:3.1.13.5] | 2461 | 2455 | 2464 | 2476 | 1428 | 1437 | 1363 | 1400 | 1420 | 1478 | 1459 | 1494 |
| K03976 Cys-tRNA(Pro)/Cys-tRNA(Cys) deacylase [EC:3.1.1.-] | 1484 | 1554 | 1501 | 1628 | 1610 | 1637 | 1604 | 1655 | 1513 | 1592 | 1545 | 1508 |
| K04075 tRNA(Ile)-lysidine synthase [EC:6.3.4.19] | 4295 | 4540 | 4403 | 4425 | 4550 | 4591 | 4358 | 4527 | 4201 | 4345 | 4268 | 4313 |
| K04094 methylenetetrahydrofolate--tRNA-(uracil-5-)-methyltransferase [EC:2.1.1.74] | 171 | 138 | 129 | 128 | 100 | 79 | 109 | 118 | 142 | 180 | 194 | 180 |
| K05539 tRNA-dihydrouridine synthase A [EC:1.-.-.-] | 2108 | 2041 | 2088 | 2054 | 1246 | 1393 | 1225 | 1335 | 1432 | 1492 | 1309 | 1420 |
| K05540 tRNA-dihydrouridine synthase B [EC:1.-.-.-] | 2158 | 2087 | 2076 | 2209 | 1304 | 1279 | 1299 | 1325 | 1369 | 1382 | 1391 | 1369 |
| K05541 tRNA-dihydrouridine synthase C [EC:1.-.-.-] | 1338 | 1233 | 1209 | 1311 | 737 | 821 | 704 | 776 | 1028 | 996 | 964 | 1127 |
| K06020 energy-dependent translational throttle protein EttA | 3895 | 3784 | 3420 | 3593 | 2238 | 2243 | 2264 | 2346 | 2187 | 2318 | 2202 | 2414 |
| K06168 tRNA-2-methylthio-N6-dimethylallyladenosine synthase [EC:2.8.4.3] | 4371 | 4415 | 4476 | 4488 | 5198 | 5262 | 5421 | 5208 | 5044 | 5183 | 5037 | 5027 |
| K06169 tRNA-(ms[2]io[6]A)-hydroxylase [EC:1.-.-.-] | 1316 | 1370 | 1276 | 1277 | 778 | 750 | 804 | 782 | 935 | 935 | 872 | 979 |

| | | | | | | | | | | | | |
|---|---|---|---|---|---|---|---|---|---|---|---|---|
| K06173 tRNA pseudouridine38-40 synthase [EC:5.4.99.12] | 2952 | 2849 | 2992 | 2847 | 2860 | 2760 | 2775 | 2878 | 2869 | 3071 | 3129 | 3044 |
| K06175 tRNA pseudouridine65 synthase [EC:5.4.99.26] | 447 | 487 | 442 | 504 | 326 | 358 | 365 | 359 | 632 | 709 | 684 | 691 |
| K06176 tRNA pseudouridine13 synthase [EC:5.4.99.27] | 2191 | 2075 | 1950 | 2030 | 1262 | 1215 | 1247 | 1257 | 1285 | 1333 | 1310 | 1371 |
| K06917 tRNA 2-selenouridine synthase [EC:2.9.1.-] | 966 | 1044 | 864 | 987 | 441 | 453 | 453 | 390 | 1082 | 1272 | 1228 | 1173 |
| K06925 tRNA threonylcarbamoyladenosine biosynthesis protein TsaE | 1487 | 1554 | 1632 | 1545 | 1645 | 1713 | 1693 | 1641 | 1565 | 1583 | 1611 | 1686 |
| K06932 tRNA(Ile2)-agmatinylcytidine synthase [EC:6.3.4.22] | 0 | 0 | 1 | 0 | 0 | 3 | 0 | 0 | 4 | 3 | 4 | 10 |
| K06957 tRNA(Met) cytidine acetyltransferase [EC:2.3.1.193] | 1210 | 1216 | 1130 | 1247 | 780 | 881 | 846 | 852 | 1600 | 1572 | 1515 | 1665 |
| K06967 tRNA (adenine22-N1)-methyltransferase [EC:2.1.1.217] | 319 | 306 | 308 | 294 | 74 | 84 | 103 | 99 | 331 | 370 | 326 | 345 |
| K06980 tRNA-modifying protein YgfZ | 2214 | 2098 | 1908 | 2032 | 1305 | 1267 | 1147 | 1339 | 1371 | 1365 | 1241 | 1467 |
| K07050 misacylated tRNA(Ala) deacylase [EC:3.1.1.-] | 0 | 0 | 0 | 0 | 0 | 0 | 2 | 1 | 0 | 0 | 0 | 1 |
| K07055 tRNA wybutosine-synthesizing protein 2 [EC:2.5.1.114] | 0 | 0 | 0 | 1 | 0 | 2 | 3 | 0 | 4 | 0 | 2 | 2 |
| K07254 tRNA (cytidine56-2'-O)-methyltransferase [EC:2.1.1.206] | 0 | 0 | 0 | 1 | 0 | 0 | 0 | 1 | 4 | 5 | 0 | 3 |
| K07442 tRNA (adenine57-N1/adenine58-N1)-methyltransferase catalytic subunit [EC:2.1.1.219 2.1.1.220] | 12 | 7 | 11 | 10 | 7 | 7 | 8 | 2 | 13 | 37 | 14 | 32 |
| K07446 tRNA (guanine10-N2)-dimethyltransferase [EC:2.1.1.213] | 0 | 0 | 0 | 1 | 0 | 2 | 0 | 0 | 0 | 5 | 2 | 1 |
| K07558 tRNA nucleotidyltransferase (CCA-adding enzyme) [EC:2.7.7.72] | 0 | 0 | 0 | 1 | 0 | 0 | 0 | 4 | 2 | 8 | 3 | 12 |
| K07559 putative RNA 2'-phosphotransferase [EC:2.7.1.-] | 1318 | 1255 | 1185 | 1190 | 770 | 710 | 686 | 773 | 477 | 493 | 435 | 415 |
| K07560 D-aminoacyl-tRNA deacylase [EC:3.1.1.96] | 1624 | 1610 | 1722 | 1659 | 1842 | 1928 | 1805 | 1900 | 1831 | 1754 | 1727 | 1835 |
| K07568 S-adenosylmethionine:tRNA ribosyltransferase-isomerase [EC:2.4.99.17] | 4985 | 5151 | 5224 | 5086 | 6857 | 6924 | 6796 | 6793 | 6108 | 6029 | 6146 | 6029 |
| K07583 tRNA pseudouridine synthase 10 [EC:5.4.99.25] | 0 | 3 | 2 | 1 | 0 | 4 | 0 | 2 | 2 | 13 | 3 | 17 |
| K07739 elongator complex protein 3 [EC:2.3.1.48] | 1 | 2 | 3 | 0 | 1 | 2 | 0 | 5 | 2 | 32 | 9 | 14 |
| K09119 KEOPS complex subunit Cgi121 | 1 | 0 | 0 | 0 | 0 | 0 | 0 | 0 | 4 | 4 | 0 | 3 |
| K09765 epoxyqueuosine reductase [EC:1.17.99.6] | 705 | 692 | 706 | 666 | 1261 | 1245 | 1383 | 1305 | 1133 | 1081 | 1126 | 1125 |
| K11991 tRNA(adenine34) deaminase [EC:3.5.4.33] | 878 | 956 | 1041 | 898 | 1274 | 1308 | 1193 | 1280 | 1482 | 1511 | 1445 | 1498 |
| K13628 iron-sulfur cluster assembly protein | 704 | 718 | 822 | 688 | 434 | 475 | 418 | 452 | 519 | 443 | 441 | 484 |
| K14058 tRNA 2-thiocytidine biosynthesis protein TtcA | 1599 | 1596 | 1539 | 1664 | 1029 | 1082 | 994 | 1020 | 1192 | 1257 | 1197 | 1287 |
| K14415 tRNA-splicing ligase RtcB (3'-phosphate/5'-hydroxy nucleic acid ligase) [EC:6.5.1.8] | 4402 | 4455 | 4513 | 4269 | 5924 | 6137 | 6341 | 6268 | 4766 | 4684 | 4784 | 4593 |
| K14742 tRNA threonylcarbamoyladenosine biosynthesis protein TsaB | 2116 | 2176 | 2063 | 2054 | 2301 | 2286 | 2358 | 2328 | 2116 | 2093 | 2042 | 2056 |
| K14864 tRNA (cytidine32/guanosine34-2'-O)-methyltransferase [EC:2.1.1.205] | 0 | 2 | 0 | 0 | 0 | 0 | 0 | 0 | 0 | 0 | 0 | 0 |
| K15256 tRNA (cmo5U34)-methyltransferase [EC:2.1.1.-] | 2422 | 2505 | 2461 | 2510 | 1463 | 1504 | 1551 | 1534 | 1177 | 1153 | 1134 | 1122 |
| K15257 tRNA (mo5U34)-methyltransferase [EC:2.1.1.-] | 2078 | 1976 | 1987 | 1950 | 1073 | 1149 | 1088 | 1170 | 1225 | 1229 | 1274 | 1261 |
| K15396 tRNA (cytidine32/uridine32-2'-O)-methyltransferase [EC:2.1.1.200] | 1616 | 1642 | 1614 | 1610 | 937 | 958 | 905 | 896 | 1019 | 1041 | 976 | 1059 |
| K15429 tRNA (guanine37-N1)-methyltransferase [EC:2.1.1.228] | 0 | 0 | 0 | 0 | 0 | 0 | 0 | 0 | 0 | 5 | 2 | 6 |
| K15437 aminoacyl tRNA synthase complex-interacting multifunctional protein 1 | 0 | 0 | 0 | 0 | 0 | 0 | 0 | 0 | 0 | 0 | 1 | 0 |
| K15441 tRNA-specific adenosine deaminase 2 [EC:3.5.4.-] | 1 | 0 | 0 | 0 | 0 | 0 | 0 | 0 | 0 | 0 | 0 | 0 |
| K15449 tRNA wybutosine-synthesizing protein 1 [EC:4.1.3.44] | 2 | 0 | 2 | 1 | 2 | 0 | 0 | 0 | 5 | 10 | 7 | 3 |
| K15451 tRNA wybutosine-synthesizing protein 4 [EC:2.1.1.290 2.3.1.231] | 2 | 0 | 0 | 0 | 0 | 0 | 0 | 0 | 0 | 0 | 0 | 0 |
| K15460 tRNA1Val (adenine37-N6)-methyltransferase [EC:2.1.1.223] | 805 | 855 | 873 | 895 | 1688 | 1712 | 1779 | 1483 | 1629 | 1481 | 1464 | 1581 |
| K15461 tRNA 5-methylaminomethyl-2-thiouridine biosynthesis bifunctional protein [EC:2.1.1.61 1.5.-.-] | 3428 | 3557 | 3154 | 3451 | 1987 | 1951 | 1793 | 2212 | 2179 | 2056 | 2018 | 2122 |
| K15865 threonylcarbamoyladenosine tRNA methylthiotransferase CDKAL1 [EC:2.8.4.5] | 0 | 0 | 0 | 0 | 0 | 0 | 0 | 0 | 13 | 8 | 3 | 12 |
| K16317 tRNA (pseudouridine54-N1)-methyltransferase [EC:2.1.1.257] | 0 | 0 | 0 | 0 | 0 | 0 | 0 | 0 | 0 | 0 | 0 | 1 |
| K18707 threonylcarbamoyladenosine tRNA methylthiotransferase MtaB [EC:2.8.4.5] | 1447 | 1439 | 1428 | 1422 | 2969 | 2905 | 3015 | 2868 | 2547 | 2476 | 2679 | 2497 |
| K18779 7-cyano-7-deazaguanine tRNA-ribosyltransferase [EC:2.4.2.48] | 2 | 2 | 0 | 2 | 3 | 0 | 5 | 0 | 9 | 12 | 3 | 13 |
| K18828 tRNA(fMet)-specific endonuclease VapC [EC:3.1.-.-] | 591 | 672 | 708 | 659 | 559 | 582 | 625 | 547 | 156 | 152 | 129 | 163 |
| K18979 epoxyqueuosine reductase [EC:1.17.99.6] | 2505 | 2511 | 2408 | 2521 | 1329 | 1379 | 1310 | 1447 | 1552 | 1693 | 1556 | 1643 |
| K19055 Ala-tRNA(Pro) deacylase [EC:3.1.1.-] | 1179 | 1217 | 1220 | 1228 | 717 | 676 | 661 | 639 | 848 | 908 | 821 | 966 |
| K21947 tRNA-5-methyluridine54 2-sulfurtransferase [EC:2.8.1.15] | 16 | 16 | 20 | 11 | 8 | 7 | 6 | 3 | 15 | 27 | 8 | 24 |
| K22132 tRNA threonylcarbamoyladenosine dehydratase | 2366 | 2404 | 2285 | 2282 | 2480 | 2449 | 2497 | 2398 | 2386 | 2431 | 2393 | 2330 |
| K22446 tRNA (cytosine49-C5)-methyltransferase [EC:2.1.1.-] | 1 | 2 | 0 | 0 | 0 | 0 | 0 | 0 | 0 | 4 | 4 | 1 | 1 |
| K22900 tRNA (adenine37-N6)-methyltransferase [EC:2.1.1.-] | 1267 | 1126 | 1073 | 1183 | 809 | 726 | 722 | 780 | 924 | 922 | 918 | 921 |
| K22901 tRNA (cytosine40_48-C5)-methyltransferase [EC:2.1.1.-] | 0 | 0 | 0 | 0 | 0 | 0 | 0 | 0 | 0 | 0 | 1 | 0 |
| ko03012 Translation factors | 74099 | 73560 | 74132 | 74198 | 79172 | 80030 | 80308 | 79532 | 75911 | 75938 | 75714 | 75841 |
| K01056 peptidyl-tRNA hydrolase, PTH1 family [EC:3.1.1.29] | 2220 | 2180 | 2234 | 2166 | 2362 | 2450 | 2222 | 2353 | 2148 | 2124 | 2197 | 2125 |
| K02355 elongation factor G | 13885 | 14034 | 14571 | 14373 | 16119 | 16274 | 16431 | 15978 | 15295 | 15373 | 15119 | 14999 |
| K02356 elongation factor P | 2301 | 2163 | 2274 | 2298 | 2447 | 2455 | 2424 | 2482 | 2651 | 2804 | 2664 | 2685 |
| K02357 elongation factor Ts | 2096 | 2094 | 2153 | 2163 | 3033 | 3178 | 3099 | 2950 | 3111 | 3183 | 3199 | 3084 |
| K02358 elongation factor Tu | 7248 | 7245 | 7465 | 7340 | 6619 | 7047 | 6673 | 6791 | 6268 | 6267 | 6172 | 6165 |
| K02493 release factor glutamine methyltransferase [EC:2.1.1.297] | 2942 | 2843 | 2800 | 2863 | 2979 | 3066 | 3051 | 3084 | 2841 | 2665 | 2896 | 2810 |
| K02518 translation initiation factor IF-1 | 698 | 680 | 684 | 679 | 767 | 838 | 869 | 800 | 851 | 838 | 812 | 805 |
| K02519 translation initiation factor IF-2 | 7933 | 7841 | 8057 | 7814 | 9232 | 9372 | 10195 | 10116 | 8034 | 8185 | 8140 | 8210 |
| K02520 translation initiation factor IF-3 | 1846 | 1811 | 1875 | 1803 | 1995 | 1863 | 2155 | 2079 | 1886 | 1849 | 1968 | 1942 |
| K02835 peptide chain release factor 1 | 3934 | 3744 | 3922 | 4025 | 4182 | 4000 | 4196 | 3990 | 4103 | 4015 | 3868 | 3936 |
| K02836 peptide chain release factor 2 | 3520 | 3553 | 3531 | 3568 | 3587 | 3577 | 3639 | 3611 | 3541 | 3525 | 3545 | 3490 |
| K02837 peptide chain release factor 3 | 4478 | 4339 | 4490 | 4542 | 5279 | 5364 | 5340 | 5320 | 5193 | 5041 | 5117 | 5186 |
| K02838 ribosome recycling factor | 1991 | 1899 | 2039 | 1971 | 2089 | 2159 | 2101 | 2094 | 2013 | 2032 | 2016 | 1978 |
| K02839 peptide chain release factor | 1409 | 1582 | 1480 | 1465 | 1783 | 1789 | 1758 | 1611 | 1276 | 1213 | 1293 | 1278 |
| K03232 elongation factor 1-beta | 0 | 0 | 0 | 0 | 0 | 0 | 0 | 0 | 0 | 0 | 0 | 1 |
| K03235 elongation factor 3 | 0 | 0 | 1 | 0 | 0 | 0 | 0 | 0 | 0 | 1 | 0 | 0 |
| K03263 translation initiation factor 5A | 0 | 0 | 0 | 0 | 0 | 3 | 0 | 2 | 0 | 0 | 3 | 0 |
| K03833 selenocysteine-specific elongation factor | 4451 | 4348 | 3745 | 4191 | 2615 | 2412 | 2452 | 2563 | 2609 | 2592 | 2504 | 2768 |
| K04568 elongation factor P--(R)-beta-lysine ligase [EC:6.3.1.-] | 693 | 652 | 621 | 664 | 468 | 463 | 506 | 444 | 885 | 972 | 888 | 900 |
| K04794 peptidyl-tRNA hydrolase, PTH2 family [EC:3.1.1.29] | 2 | 1 | 0 | 1 | 0 | 5 | 0 | 3 | 0 | 7 | 3 | 3 |
| K06158 ATP-binding cassette, subfamily F, member 3 | 9144 | 9213 | 8978 | 9047 | 10940 | 11018 | 10571 | 10559 | 9939 | 9971 | 10093 | 10068 |
| K06927 diphthine-ammonia ligase [EC:6.3.1.14] | 2 | 3 | 0 | 0 | 0 | 4 | 2 | 0 | 1 | 2 | 3 | 7 |
| K07561 2-(3-amino-3-carboxypropyl)histidine synthase [EC:2.5.1.108] | 2 | 0 | 0 | 0 | 0 | 2 | 0 | 0 | 4 | 3 | 3 | 9 |
| K09890 alternative ribosome-rescue factor | 339 | 374 | 355 | 366 | 191 | 197 | 188 | 187 | 307 | 277 | 283 | 308 |
| K09906 elongation factor P hydroxylase [EC:1.14.-.-] | 345 | 318 | 333 | 316 | 220 | 213 | 217 | 216 | 426 | 518 | 486 | 490 |
| K15029 translation initiation factor 3 subunit L | 0 | 0 | 0 | 0 | 0 | 0 | 0 | 0 | 0 | 0 | 0 | 1 |
| K15034 ribosome-associated protein | 868 | 840 | 836 | 866 | 536 | 483 | 439 | 484 | 506 | 482 | 473 | 473 |
| K19810 L-lysine 2,3-aminomutase [EC:5.4.3.-] | 667 | 653 | 614 | 653 | 434 | 534 | 462 | 449 | 969 | 940 | 903 | 995 |
| ko03110 Chaperones and folding catalysts | 140562 | 142302 | 144193 | 142461 | 171690 | 171961 | 172907 | 172002 | 156854 | 155575 | 157192 | 155640 |
| K02199 cytochrome c biogenesis protein CcmG, thiol:disulfide interchange protein DsbE | 775 | 734 | 716 | 743 | 370 | 356 | 351 | 409 | 583 | 533 | 556 | 562 |
| K03611 disulfide bond formation protein DsbB | 2288 | 2317 | 2178 | 2264 | 1297 | 1316 | 1217 | 1419 | 1114 | 1200 | 1066 | 1070 |
| K03667 ATP-dependent HslUV protease ATP-binding subunit HslU | 3231 | 3335 | 3160 | 3331 | 1963 | 2013 | 1682 | 1877 | 2028 | 2121 | 1976 | 2085 |
| K03672 thioredoxin 2 [EC:1.8.1.8] | 271 | 243 | 248 | 244 | 170 | 188 | 178 | 181 | 354 | 402 | 352 | 412 |
| K03674 glutaredoxin 1 | 132 | 163 | 162 | 154 | 100 | 94 | 102 | 105 | 192 | 210 | 193 | 215 |
| K03675 glutaredoxin 2 | 77 | 99 | 105 | 132 | 179 | 176 | 214 | 200 | 36 | 31 | 28 | 33 |
| K03676 glutaredoxin 3 | 644 | 665 | 631 | 638 | 396 | 462 | 390 | 395 | 400 | 409 | 417 | 428 |
| K03686 molecular chaperone DnaJ | 3291 | 3442 | 3241 | 3286 | 3702 | 3566 | 3480 | 3493 | 3163 | 3138 | 3235 | 3430 |
| K03687 molecular chaperone GrpE | 1148 | 1061 | 1093 | 1097 | 1375 | 1433 | 1686 | 1548 | 1564 | 1559 | 1566 | 1466 |
| K03694 ATP-dependent Clp protease ATP-binding subunit ClpA | 2891 | 2798 | 2889 | 2884 | 1959 | 2008 | 1995 | 1932 | 2427 | 2450 | 2275 | 2439 |
| K03696 ATP-dependent Clp protease ATP-binding subunit ClpC | 7391 | 7401 | 7383 | 7143 | 7670 | 7260 | 7681 | 7673 | 6183 | 6162 | 6199 | 6048 |
| K03697 ATP-dependent Clp protease ATP-binding subunit ClpE | 105 | 128 | 122 | 122 | 42 | 40 | 37 | 34 | 128 | 142 | 149 | 157 |
| K03768 peptidyl-prolyl cis-trans isomerase B (cyclophilin B) [EC:5.2.1.8] | 1812 | 1814 | 1727 | 1862 | 913 | 981 | 957 | 1050 | 1434 | 1525 | 1478 | 1496 |
| K03769 peptidyl-prolyl cis-trans isomerase C [EC:5.2.1.8] | 2848 | 2795 | 2684 | 2660 | 1829 | 1882 | 1821 | 1887 | 1363 | 1339 | 1305 | 1406 |
| K03770 peptidyl-prolyl cis-trans isomerase D [EC:5.2.1.8] | 3607 | 3514 | 3713 | 3647 | 6022 | 6216 | 6396 | 6170 | 5246 | 5137 | 5271 | 5307 |
| K03771 peptidyl-prolyl cis-trans isomerase SurA [EC:5.2.1.8] | 2606 | 2845 | 2940 | 2787 | 6809 | 6968 | 7175 | 6967 | 5754 | 5251 | 5598 | 5336 |
| K03772 FKBP-type peptidyl-prolyl cis-trans isomerase FkpA [EC:5.2.1.8] | 570 | 563 | 568 | 561 | 429 | 449 | 446 | 467 | 772 | 666 | 741 | 750 |
| K03773 FKBP-type peptidyl-prolyl cis-trans isomerase FklB [EC:5.2.1.8] | 4047 | 4211 | 4211 | 4211 | 4598 | 4473 | 4611 | 4597 | 3453 | 3389 | 3466 | 3387 |
| K03774 FKBP-type peptidyl-prolyl cis-trans isomerase SlpA [EC:5.2.1.8] | 1015 | 997 | 1002 | 958 | 659 | 680 | 583 | 579 | 665 | 605 | 626 | 626 |
| K03775 FKBP-type peptidyl-prolyl cis-trans isomerase SlyD [EC:5.2.1.8] | 1341 | 1398 | 1393 | 1362 | 1788 | 1812 | 1794 | 1755 | 1505 | 1578 | 1499 | 1576 |
| K03805 thiol:disulfide interchange protein DsbG | 1360 | 1373 | 1257 | 1336 | 771 | 798 | 768 | 917 | 910 | 840 | 866 | 889 |
| K03981 thiol:disulfide interchange protein DsbC [EC:5.3.4.1] | 1206 | 1139 | 1100 | 1155 | 823 | 893 | 800 | 897 | 897 | 894 | 806 | 864 |
| K04044 molecular chaperone HscA | 2858 | 2831 | 2744 | 2776 | 1949 | 1810 | 1913 | 1930 | 2169 | 2209 | 2164 | 2199 |
| K04045 molecular chaperone HscC | 958 | 926 | 943 | 958 | 564 | 685 | 687 | 681 | 623 | 636 | 659 | 714 |
| K04046 hypothetical chaperone protein | 2847 | 2912 | 2814 | 2860 | 1786 | 1710 | 1713 | 1813 | 1877 | 1904 | 1855 | 2044 |
| K04078 chaperonin GroES | 1021 | 1006 | 1131 | 1045 | 1123 | 1054 | 1122 | 1138 | 1109 | 1134 | 1057 | 1110 |
| K04080 molecular chaperone IbpA | 1158 | 1147 | 1054 | 1226 | 714 | 697 | 725 | 715 | 1010 | 1004 | 932 | 1025 |
| K04081 molecular chaperone IbpB | 273 | 298 | 252 | 294 | 246 | 208 | 233 | 194 | 411 | 372 | 373 | 450 |
| K04082 molecular chaperone HscB | 947 | 885 | 892 | 875 | 545 | 516 | 580 | 558 | 633 | 662 | 597 | 633 |
| K04083 molecular chaperone Hsp33 | 2333 | 2375 | 2267 | 2387 | 1362 | 1363 | 1397 | 1357 | 1732 | 1706 | 1658 | 1770 |
| K04084 thiol:disulfide interchange protein DsbD [EC:1.8.1.8] | 8690 | 8762 | 8465 | 8512 | 10895 | 10633 | 10815 | 11208 | 8294 | 7788 | 8407 | 8030 |
| K04086 ATP-dependent Clp protease ATP-binding subunit ClpL | 11 | 22 | 16 | 19 | 38 | 11 | 21 | 16 | 34 | 55 | 27 | 30 |
| K04651 hydrogenase nickel incorporation protein HypA/HybF | 379 | 389 | 383 | 383 | 268 | 325 | 278 | 260 | 521 | 532 | 512 | 480 |
| K04652 hydrogenase nickel incorporation protein HypB | 723 | 692 | 661 | 726 | 434 | 481 | 444 | 464 | 1073 | 1112 | 1002 | 1084 |
| K04762 ribosome-associated heat shock protein Hsp15 | 1310 | 1230 | 1350 | 1189 | 1441 | 1513 | 1475 | 1480 | 1186 | 1142 | 1166 | 1132 |
| K04797 prefoldin alpha subunit | 0 | 0 | 0 | 0 | 2 | 0 | 2 | 3 | 0 | 0 | 1 | 2 |
| K04798 prefoldin beta subunit | 0 | 0 | 0 | 0 | 0 | 1 | 0 | 0 | 0 | 0 | 0 | 2 |
| K05516 curved DNA-binding protein | 910 | 966 | 1015 | 889 | 1932 | 1999 | 1974 | 1944 | 2081 | 1960 | 2004 | 1986 |
| K05801 DnaJ like chaperone protein | 1693 | 1554 | 1547 | 1615 | 1126 | 1068 | 931 | 1094 | 1067 | 1045 | 1042 | 1108 |
| K05838 putative thioredoxin | 2040 | 1861 | 1836 | 1929 | 1050 | 1158 | 1026 | 1220 | 1234 | 1101 | 1165 | 1162 |
| K06006 periplasmic protein CpxP/Spy | 551 | 547 | 480 | 578 | 375 | 400 | 443 | 378 | 710 | 671 | 722 | 765 |

| | | | | | | | | | | | | |
|---|---|---|---|---|---|---|---|---|---|---|---|---|
| K07346 fimbrial chaperone protein | 3012 | 3090 | 2860 | 3003 | 1673 | 1702 | 1671 | 1796 | 1535 | 1499 | 1517 | 1563 |
| K07390 monothiol glutaredoxin | 676 | 663 | 678 | 691 | 387 | 394 | 370 | 454 | 395 | 411 | 375 | 448 |
| K07533 foldase protein PrsA [EC:5.2.1.8] | 418 | 429 | 374 | 437 | 128 | 117 | 100 | 102 | 404 | 471 | 430 | 439 |
| K09488 TNF receptor-associated protein 1 | 3 | 0 | 2 | 3 | 1 | 2 | 0 | 3 | 3 | 4 | 2 | 1 |
| K09510 DnaJ homolog subfamily B member 4 | 0 | 0 | 0 | 0 | 0 | 1 | 0 | 0 | 0 | 0 | 0 | 0 |
| K09569 FK506-binding protein 2 [EC:5.2.1.8] | 4 | 2 | 2 | 3 | 6 | 3 | 8 | 4 | 4 | 4 | 14 | 3 |
| K09575 FK506-binding protein 9/10 [EC:5.2.1.8] | 4 | 4 | 4 | 2 | 5 | 7 | 2 | 3 | 5 | 7 | 9 | 5 |
| K09796 periplasmic copper chaperone A | 784 | 760 | 733 | 742 | 419 | 394 | 346 | 478 | 256 | 258 | 220 | 264 |
| K19777 acid stress chaperone HdeA | 0 | 0 | 0 | 0 | 3 | 2 | 2 | 0 | 12 | 13 | 11 | 11 |
| K19778 acid stress chaperone HdeB | 238 | 302 | 256 | 246 | 76 | 79 | 84 | 73 | 509 | 573 | 545 | 563 |
| K21804 protein N-lysine methyltransferase METTL21A [EC:2.1.1.-] | 0 | 0 | 0 | 0 | 1 | 0 | 0 | 0 | 0 | 0 | 0 | 0 |
| K22447 archaeal chaperonin | 5 | 2 | 6 | 3 | 5 | 1 | 5 | 2 | 5 | 25 | 8 | 23 |
| K23349 putative dimethyl sulfoxide reductase chaperone | 339 | 352 | 319 | 345 | 239 | 259 | 283 | 257 | 523 | 496 | 512 | 539 |
| ko04131 Membrane trafficking | 33126 | 33650 | 33901 | 33584 | 38836 | 38843 | 39242 | 39087 | 39286 | 38833 | 39803 | 39386 |
| K08332 vacuolar protein 8 | 0 | 0 | 2 | 0 | 0 | 0 | 0 | 0 | 0 | 0 | 0 | 0 |
| K08518 syntaxin-binding protein 5 | 0 | 0 | 0 | 0 | 0 | 0 | 0 | 0 | 0 | 4 | 0 | 0 |
| K10402 kinesin family member 20 | 0 | 0 | 0 | 0 | 0 | 0 | 0 | 1 | 0 | 0 | 0 | 0 |
| K17922 sorting nexin-8 | 0 | 2 | 0 | 0 | 0 | 0 | 0 | 0 | 0 | 0 | 0 | 0 |
| K19950 protein transport protein SEC9 | 0 | 0 | 1 | 0 | 0 | 0 | 0 | 0 | 0 | 0 | 0 | 0 |
| K19998 sec1 family domain-containing protein 1 | 0 | 0 | 0 | 0 | 0 | 0 | 0 | 0 | 0 | 0 | 2 | 0 |
| K20280 trafficking protein particle complex subunit 5 | 0 | 0 | 0 | 0 | 0 | 0 | 0 | 0 | 0 | 0 | 0 | 3 |
| K20352 p24 family protein delta-1 | 0 | 0 | 0 | 0 | 0 | 0 | 0 | 0 | 0 | 0 | 2 | 0 |
| K20456 oxysterol-binding protein 1 | 0 | 0 | 1 | 0 | 0 | 0 | 0 | 0 | 0 | 0 | 0 | 0 |
| K21440 ankyrin repeat domain-containing protein 50 | 0 | 0 | 0 | 0 | 1 | 2 | 0 | 0 | 0 | 2 | 0 | 1 |
| K23481 vacuolar protein sorting-associated protein 1 | 0 | 0 | 1 | 0 | 0 | 0 | 0 | 0 | 0 | 0 | 0 | 0 |
| ko04121 Ubiquitin system | 7 | 16 | 14 | 9 | 4 | 2 | 2 | 1 | 11 | 5 | 10 | 3 |
| K10273 F-box and leucine-rich repeat protein 7 | 0 | 0 | 3 | 0 | 0 | 0 | 0 | 0 | 0 | 0 | 0 | 0 |
| K11997 tripartite motif-containing protein 2/3 | 0 | 0 | 0 | 0 | 0 | 0 | 0 | 0 | 0 | 0 | 1 | 0 |
| K12025 tripartite motif-containing protein 52 | 0 | 0 | 0 | 0 | 0 | 0 | 0 | 0 | 0 | 1 | 0 | 1 |
| K13570 prokaryotic ubiquitin-like protein Pup | 0 | 1 | 0 | 0 | 1 | 0 | 2 | 0 | 3 | 0 | 0 | 0 |
| K15071 recyclin-1 | 0 | 1 | 0 | 0 | 0 | 0 | 0 | 0 | 0 | 0 | 0 | 0 |
| K22372 E3 ISG15--protein ligase [EC:2.3.2.-] | 1 | 0 | 0 | 0 | 0 | 0 | 0 | 1 | 0 | 0 | 0 | 0 |
| K22382 WD repeat-containing protein 26 | 0 | 1 | 0 | 0 | 0 | 0 | 0 | 0 | 0 | 0 | 0 | 0 |
| K23333 E3 ubiquitin-protein transferase RMND5 [EC:2.3.2.27] | 0 | 0 | 0 | 0 | 0 | 1 | 0 | 0 | 0 | 0 | 0 | 0 |
| ko03051 Proteasome | 8035 | 8045 | 8198 | 8234 | 11373 | 11387 | 11460 | 11438 | 9455 | 9451 | 9563 | 9728 |
| K13571 proteasome accessory factor A [EC:6.3.1.19] | 2 | 5 | 6 | 4 | 0 | 5 | 6 | 3 | 5 | 17 | 12 | 20 |
| K13573 proteasome accessory factor C | 0 | 1 | 0 | 0 | 0 | 0 | 0 | 0 | 0 | 1 | 0 | 0 |
| K20814 Pup amidohydrolase [EC:3.5.1.119] | 3 | 3 | 4 | 2 | 3 | 9 | 1 | 9 | 6 | 4 | 4 | 15 |
| ko03032 DNA replication proteins | 151073 | 152131 | 154561 | 153379 | 177539 | 179508 | 179646 | 177943 | 166333 | 165020 | 164970 | 164458 |
| K02315 DNA replication protein DnaC | 2707 | 2698 | 2756 | 2779 | 1833 | 1810 | 1708 | 1755 | 1843 | 1918 | 1784 | 1820 |
| K02469 DNA gyrase subunit A [EC:5.6.2.2] | 9652 | 9879 | 9806 | 9889 | 9055 | 9280 | 9264 | 8947 | 9623 | 9891 | 9375 | 9607 |
| K02470 DNA gyrase subunit B [EC:5.6.2.2] | 8651 | 8590 | 8771 | 8696 | 8339 | 8597 | 8787 | 8460 | 8779 | 8915 | 8840 | 8797 |
| K02621 topoisomerase IV subunit A [EC:5.6.2.2] | 6129 | 6165 | 6441 | 6389 | 9185 | 9111 | 9095 | 9237 | 7947 | 7459 | 7545 | 7576 |
| K02622 topoisomerase IV subunit B [EC:5.6.2.2] | 5161 | 5311 | 5380 | 5383 | 6896 | 6807 | 6965 | 6869 | 5736 | 5672 | 5541 | 5578 |
| K03163 DNA topoisomerase I [EC:5.6.2.1] | 0 | 0 | 0 | 0 | 0 | 0 | 0 | 0 | 0 | 0 | 0 | 1 |
| K03166 DNA topoisomerase VI subunit A [EC:5.6.2.2] | 0 | 0 | 1 | 0 | 2 | 4 | 2 | 0 | 8 | 10 | 10 | 12 |
| K03167 DNA topoisomerase VI subunit B [EC:5.6.2.2] | 1 | 2 | 4 | 0 | 0 | 0 | 0 | 5 | 5 | 8 | 8 | 12 |
| K03168 DNA topoisomerase I [EC:5.6.2.1] | 10958 | 10964 | 10899 | 10779 | 10407 | 10381 | 10044 | 10254 | 8920 | 8753 | 8391 | 8541 |
| K03169 DNA topoisomerase III [EC:5.6.2.1] | 10565 | 10897 | 11225 | 10763 | 20071 | 20242 | 20277 | 20056 | 18116 | 17946 | 17799 | 17620 |
| K03346 replication initiation and membrane attachment protein | 72 | 69 | 69 | 80 | 21 | 22 | 22 | 15 | 90 | 86 | 82 | 66 |
| K03530 DNA-binding protein HU-beta | 1924 | 1902 | 1999 | 1869 | 2945 | 2860 | 2951 | 2919 | 2340 | 2348 | 2316 | 2349 |
| K03645 negative modulator of initiation of replication | 320 | 347 | 344 | 421 | 280 | 256 | 215 | 257 | 567 | 579 | 581 | 540 |
| K04764 integration host factor subunit alpha | 570 | 644 | 628 | 608 | 338 | 365 | 343 | 366 | 383 | 389 | 369 | 380 |
| K05787 DNA-binding protein HU-alpha | 648 | 629 | 600 | 639 | 384 | 392 | 353 | 396 | 395 | 378 | 363 | 362 |
| K05788 integration host factor subunit beta | 708 | 646 | 669 | 657 | 451 | 449 | 399 | 392 | 378 | 437 | 400 | 424 |
| K09723 DNA replication factor GINS | 0 | 0 | 0 | 0 | 2 | 0 | 2 | 0 | 2 | 4 | 4 | 7 |
| K10725 archaeal cell division control protein 6 | 0 | 3 | 2 | 0 | 2 | 2 | 0 | 3 | 8 | 8 | 13 | 20 |
| K10730 ATP-dependent DNA helicase Q4 [EC:3.6.4.12] | 0 | 0 | 0 | 0 | 0 | 0 | 0 | 0 | 0 | 1 | 0 | 0 |
| K10748 DNA replication terminus site-binding protein | 502 | 498 | 474 | 503 | 321 | 375 | 372 | 352 | 625 | 712 | 691 | 721 |
| K10763 DnaA-homolog protein | 1473 | 1357 | 1325 | 1521 | 816 | 920 | 807 | 923 | 888 | 961 | 964 | 990 |
| K10900 werner syndrome ATP-dependent helicase [EC:3.6.4.12] | 0 | 0 | 0 | 0 | 0 | 0 | 0 | 0 | 0 | 1 | 0 | 0 |
| K11144 primosomal protein DnaI | 48 | 51 | 56 | 58 | 5 | 12 | 8 | 20 | 70 | 77 | 47 | 61 |
| K18996 replication initiation protein RepC | 0 | 0 | 0 | 0 | 0 | 0 | 0 | 1 | 11 | 2 | 3 | 4 |
| K19783 DNA polymerase alpha-associated DNA helicase A [EC:3.6.4.12] | 0 | 0 | 0 | 1 | 0 | 0 | 0 | 0 | 0 | 0 | 0 | 0 |
| ko03036 Chromosome and associated proteins | 178583 | 179508 | 180946 | 180709 | 189001 | 190387 | 188782 | 189035 | 187482 | 187266 | 185156 | 185336 |
| K02499 tetrapyrrole methylase family protein / MazG family protein | 198 | 204 | 203 | 193 | 83 | 105 | 109 | 109 | 224 | 241 | 253 | 259 |
| K03466 DNA segregation ATPase FtsK/SpoIIIE, S-DNA-T family | 7822 | 7851 | 8020 | 8036 | 8695 | 8577 | 8809 | 8863 | 8163 | 8314 | 8328 | 8095 |
| K03496 chromosome partitioning protein | 10687 | 10580 | 10690 | 11012 | 10427 | 10397 | 10027 | 10084 | 10784 | 10712 | 10585 | 10757 |
| K03528 cell division protein ZipA | 1434 | 1300 | 1413 | 1334 | 867 | 818 | 897 | 928 | 826 | 821 | 744 | 866 |
| K03529 chromosome segregation protein | 5563 | 5471 | 5316 | 5630 | 2683 | 2762 | 2556 | 2873 | 2641 | 2860 | 2749 | 2900 |
| K03569 rod shape-determining protein MreB and related proteins | 3986 | 3959 | 3848 | 3778 | 3953 | 4016 | 4010 | 4027 | 3957 | 4028 | 4044 | 4099 |
| K03570 rod shape-determining protein MreC | 1859 | 1920 | 1876 | 1977 | 2738 | 2820 | 2834 | 2649 | 2788 | 2695 | 2794 | 2720 |
| K03571 rod shape-determining protein MreD | 1215 | 1230 | 1163 | 1186 | 689 | 653 | 642 | 700 | 836 | 954 | 852 | 853 |
| K03586 cell division protein FtsL | 626 | 637 | 614 | 642 | 444 | 441 | 383 | 364 | 461 | 435 | 432 | 446 |
| K03591 cell division protein FtsN | 422 | 389 | 385 | 427 | 242 | 334 | 341 | 328 | 555 | 535 | 512 | 541 |
| K03593 ATP-binding protein involved in chromosome partitioning | 1548 | 1698 | 1765 | 1759 | 3216 | 3212 | 3222 | 3220 | 2954 | 2846 | 2830 | 2717 |
| K03608 cell division topological specificity factor | 638 | 683 | 660 | 709 | 347 | 338 | 306 | 333 | 470 | 478 | 472 | 474 |
| K03609 septum site-determining protein MinD | 2157 | 2167 | 2114 | 2092 | 1138 | 1037 | 1118 | 1101 | 1607 | 1654 | 1599 | 1588 |
| K03610 septum site-determining protein MinC | 1811 | 1723 | 1808 | 1886 | 1014 | 910 | 934 | 930 | 1209 | 1229 | 1156 | 1194 |
| K03632 chromosome partition protein MukB | 2380 | 2490 | 2360 | 2512 | 1641 | 1864 | 1810 | 1696 | 3661 | 3591 | 3356 | 3642 |
| K03633 chromosome partition protein MukF | 716 | 758 | 721 | 699 | 571 | 495 | 512 | 523 | 920 | 1040 | 1009 | 1000 |
| K03733 integrase/recombinase XerC | 2830 | 2719 | 2859 | 2801 | 3299 | 3292 | 3348 | 3424 | 3124 | 2899 | 2919 | 2916 |
| K03746 DNA-binding protein H-NS | 311 | 350 | 343 | 343 | 278 | 323 | 289 | 255 | 418 | 433 | 434 | 485 |
| K03804 chromosome partition protein MukE | 398 | 362 | 372 | 393 | 264 | 274 | 284 | 273 | 539 | 560 | 563 | 576 |
| K03978 GTP-binding protein | 1740 | 1747 | 1829 | 1743 | 2027 | 2042 | 2009 | 1885 | 1869 | 1917 | 2015 | 2011 |
| K04047 starvation-inducible DNA-binding protein | 1332 | 1243 | 1314 | 1233 | 1646 | 1664 | 1662 | 1716 | 1400 | 1359 | 1383 | 1304 |
| K04074 cell division initiation protein | 54 | 57 | 58 | 74 | 30 | 38 | 41 | 34 | 56 | 100 | 78 | 63 |
| K04095 cell filamentation protein | 558 | 556 | 473 | 542 | 329 | 395 | 342 | 307 | 678 | 689 | 606 | 773 |
| K04763 integrase/recombinase XerD | 2633 | 2665 | 2764 | 2729 | 3232 | 3281 | 3336 | 3182 | 3359 | 3467 | 3275 | 3226 |
| K05589 cell division protein FtsB | 565 | 550 | 545 | 542 | 357 | 354 | 354 | 401 | 356 | 369 | 341 | 351 |
| K05837 rod shape determining protein RodA | 3830 | 4058 | 4098 | 4093 | 4749 | 4924 | 4912 | 4739 | 4470 | 4271 | 4329 | 4254 |
| K05896 segregation and condensation protein A | 1765 | 1740 | 1570 | 1756 | 824 | 844 | 811 | 782 | 768 | 874 | 796 | 841 |
| K06024 segregation and condensation protein B | 1625 | 1579 | 1523 | 1568 | 862 | 847 | 909 | 993 | 746 | 728 | 681 | 680 |
| K06286 septation ring formation regulator | 106 | 114 | 104 | 124 | 42 | 31 | 32 | 23 | 98 | 159 | 102 | 136 |
| K06899 nucleoid-associated protein | 3001 | 3015 | 2958 | 2856 | 2158 | 2235 | 2085 | 2182 | 1552 | 1710 | 1701 | 1715 |
| K06916 cell division protein ZapE | 5184 | 5146 | 4922 | 5163 | 2751 | 2688 | 2647 | 2840 | 3225 | 3138 | 3113 | 3099 |
| K09772 cell division inhibitor SepF | 254 | 234 | 224 | 251 | 83 | 108 | 77 | 50 | 215 | 255 | 248 | 238 |
| K09888 cell division protein ZapA | 1161 | 1130 | 1153 | 1071 | 1082 | 1038 | 1005 | 1141 | 1042 | 1125 | 1000 | 1023 |
| K09892 cell division protein ZapB | 683 | 651 | 669 | 659 | 401 | 398 | 382 | 383 | 373 | 415 | 384 | 380 |
| K11267 sister chromatid cohesion protein PDS5 | 0 | 0 | 0 | 0 | 1 | 0 | 0 | 0 | 0 | 0 | 0 | 0 |
| K11274 chromosome transmission fidelity protein 4 | 0 | 0 | 0 | 0 | 0 | 2 | 0 | 0 | 0 | 0 | 0 | 0 |
| K11396 chromatin modification-related protein YNG2 | 0 | 2 | 0 | 0 | 0 | 0 | 0 | 0 | 0 | 0 | 0 | 0 |
| K11494 RCC1 and BTB domain-containing protein | 4 | 3 | 2 | 2 | 0 | 0 | 0 | 0 | 1 | 0 | 1 | 1 |
| K11673 actin-related protein 8 | 0 | 0 | 0 | 0 | 0 | 0 | 0 | 0 | 0 | 0 | 1 | 0 |
| K11685 DNA-binding protein StpA | 137 | 142 | 138 | 158 | 59 | 62 | 42 | 48 | 342 | 334 | 353 | 372 |
| K11940 heat shock protein HspQ | 213 | 164 | 184 | 199 | 118 | 143 | 140 | 135 | 271 | 276 | 281 | 293 |
| K12961 DnaA initiator-associating protein | 327 | 407 | 377 | 389 | 286 | 269 | 258 | 243 | 574 | 617 | 444 | 555 |
| K13052 cell division protein DivIC | 51 | 31 | 30 | 43 | 2 | 21 | 3 | 9 | 48 | 35 | 34 | 36 |
| K13053 cell division inhibitor SulA | 986 | 994 | 1042 | 1014 | 536 | 609 | 628 | 577 | 632 | 546 | 539 | 598 |
| K14440 SWI/SNF-related matrix-associated actin-dependent regulator of chromatin subfamily A-like protein 1 [EC:3.6.4.12] | 3 | 0 | 2 | 1 | 2 | 2 | 4 | 5 | 0 | 1 | 2 | 0 |
| K14967 COMPASS component BRE2 | 0 | 0 | 0 | 0 | 1 | 0 | 0 | 0 | 0 | 0 | 0 | 0 |
| K16570 gamma-tubulin complex component 3 | 0 | 0 | 0 | 0 | 0 | 2 | 0 | 1 | 0 | 0 | 0 | 1 |
| K16732 protein regulator of cytokinesis 1 | 0 | 1 | 0 | 0 | 0 | 0 | 0 | 0 | 0 | 0 | 0 | 0 |
| K16803 cytoskeleton-associated protein 5 | 0 | 0 | 0 | 0 | 0 | 0 | 0 | 0 | 0 | 0 | 2 | 0 |
| K18657 cell division protein ZapC | 275 | 245 | 294 | 309 | 216 | 249 | 218 | 201 | 384 | 379 | 353 | 418 |
| K18778 cell division protein ZapD | 488 | 500 | 512 | 500 | 376 | 397 | 387 | 360 | 695 | 692 | 622 | 705 |
| K18997 chaperone modulatory protein CbpM | 131 | 131 | 139 | 126 | 35 | 43 | 35 | 48 | 281 | 346 | 266 | 331 |
| K19678 intraflagellar transport protein 80 | 0 | 0 | 0 | 0 | 0 | 0 | 1 | 0 | 0 | 1 | 0 | 0 |
| K19788 obg-like ATPase 1 | 0 | 0 | 0 | 0 | 2 | 0 | 0 | 0 | 0 | 0 | 0 | 1 |
| K20092 chromodomain-helicase-DNA-binding protein 1-like [EC:3.6.4.12] | 5 | 6 | 5 | 3 | 23 | 27 | 22 | 21 | 14 | 17 | 19 | 17 |
| K23398 activating signal cointegrator 1 | 11 | 9 | 17 | 5 | 21 | 22 | 20 | 19 | 18 | 22 | 8 | 20 |

| | | | | | | | | | | | | |
|---|---|---|---|---|---|---|---|---|---|---|---|---|
| ko03400 DNA repair and recombination proteins | 411948 | 410576 | 410376 | 412556 | 402842 | 407222 | 405997 | 403014 | 407858 | 410156 | 410379 | 408771 |
| K00567 methylated-DNA-[protein]-cysteine S-methyltransferase [EC:2.1.1.63] | 1684 | 1680 | 1778 | 1632 | 1960 | 1943 | 1952 | 1883 | 1808 | 1833 | 1819 | 1922 |
| K01160 crossover junction endodeoxyribonuclease RusA [EC:3.1.22.4] | 104 | 100 | 114 | 121 | 143 | 192 | 169 | 161 | 68 | 77 | 81 | 61 |
| K01554 8-oxo-dGDP phosphatase [EC:3.6.1.58] | 0 | 0 | 0 | 0 | 0 | 0 | 0 | 0 | 1 | 1 | 2 | 1 |
| K01669 deoxyribodipyrimidine photo-lyase [EC:4.1.99.3] | 3191 | 3186 | 3009 | 3114 | 1839 | 1875 | 1794 | 2010 | 1818 | 1818 | 1839 | 2016 |
| K02336 DNA polymerase II [EC:2.7.7.7] | 3949 | 3857 | 3791 | 3818 | 2259 | 2170 | 2250 | 2253 | 2646 | 2819 | 2737 | 2800 |
| K02346 DNA polymerase IV [EC:2.7.7.7] | 3076 | 3189 | 3143 | 3199 | 3500 | 3641 | 3594 | 3485 | 4058 | 4154 | 4266 | 4237 |
| K02347 DNA polymerase (family X) | 3 | 0 | 2 | 1 | 0 | 0 | 0 | 0 | 0 | 0 | 0 | 0 |
| K03502 DNA polymerase V | 2395 | 2456 | 2395 | 2510 | 1939 | 2308 | 2196 | 2052 | 2687 | 2746 | 2620 | 2658 |
| K03546 DNA repair protein SbcC/Rad50 | 7414 | 7508 | 7485 | 7672 | 9226 | 9480 | 9783 | 9107 | 8859 | 8461 | 8697 | 8576 |
| K03547 DNA repair protein SbcD/Mre11 | 4770 | 4754 | 4857 | 4903 | 4971 | 4938 | 5068 | 4854 | 4891 | 5084 | 5054 | 4930 |
| K03552 holliday junction resolvase Hjr [EC:3.1.22.4] | 0 | 3 | 3 | 1 | 2 | 7 | 12 | 6 | 3 | 3 | 0 | 0 |
| K03554 recombination associated protein RdgC | 1589 | 1600 | 1554 | 1630 | 977 | 1008 | 934 | 997 | 1097 | 1119 | 1108 | 1044 |
| K03565 regulatory protein | 1907 | 1844 | 1895 | 1923 | 2082 | 2144 | 2134 | 1936 | 1905 | 1726 | 1908 | 1809 |
| K03574 8-oxo-dGTP diphosphatase [EC:3.6.1.55] | 3398 | 3384 | 3271 | 3459 | 1725 | 1766 | 1818 | 1754 | 1884 | 1891 | 1799 | 1900 |
| K03631 DNA repair protein RecN (Recombination protein N) | 5665 | 6045 | 5666 | 5678 | 6048 | 5758 | 5833 | 5811 | 5582 | 5245 | 5239 | 5293 |
| K03656 ATP-dependent DNA helicase Rep [EC:3.6.4.12] | 4877 | 4823 | 4625 | 4869 | 2931 | 3008 | 3013 | 2892 | 3073 | 3046 | 2926 | 3036 |
| K03658 DNA helicase IV [EC:3.6.4.12] | 1268 | 1208 | 1245 | 1205 | 781 | 895 | 904 | 839 | 1518 | 1585 | 1447 | 1473 |
| K03700 recombination protein U | 278 | 261 | 240 | 266 | 82 | 72 | 94 | 67 | 217 | 292 | 202 | 265 |
| K03722 ATP-dependent DNA helicase DinG [EC:3.6.4.12] | 6344 | 6227 | 6006 | 6172 | 3595 | 3628 | 3661 | 3753 | 4745 | 4902 | 4800 | 4769 |
| K03724 ATP-dependent helicase Lhr and Lhr-like helicase [EC:3.6.4.-] | 11378 | 11248 | 10753 | 11170 | 6527 | 6567 | 6273 | 6589 | 4624 | 4305 | 4271 | 4528 |
| K03919 DNA oxidative demethylase [EC:1.14.11.33] | 360 | 378 | 370 | 378 | 251 | 327 | 315 | 282 | 523 | 506 | 556 | 546 |
| K04483 DNA repair protein RadA | 0 | 2 | 0 | 0 | 0 | 0 | 2 | 0 | 2 | 12 | 3 | 5 |
| K04484 DNA repair protein RadB | 1 | 0 | 2 | 1 | 1 | 0 | 0 | 0 | 4 | 0 | 3 | 5 |
| K04485 DNA repair protein RadA/Sms | 4215 | 4142 | 4280 | 4225 | 4866 | 4942 | 4895 | 4985 | 4669 | 4670 | 4839 | 4679 |
| K05982 deoxyribonuclease V [EC:3.1.21.7] | 429 | 427 | 470 | 471 | 346 | 302 | 310 | 318 | 544 | 505 | 547 | 586 |
| K05984 excinuclease Cho [EC:3.1.25.-] | 434 | 463 | 432 | 481 | 293 | 372 | 314 | 283 | 623 | 606 | 602 | 684 |
| K07455 recombination protein RecT | 311 | 371 | 410 | 416 | 188 | 222 | 214 | 189 | 172 | 177 | 141 | 180 |
| K07458 DNA mismatch endonuclease, patch repair protein [EC:3.1.-.-] | 603 | 668 | 632 | 659 | 1445 | 1404 | 1500 | 1500 | 1348 | 1290 | 1293 | 1430 |
| K10906 exodeoxyribonuclease VIII [EC:3.1.11.-] | 78 | 75 | 59 | 96 | 0 | 4 | 3 | 1 | 5 | 3 | 1 | 2 |
| K12149 DNA-damage-inducible protein I | 143 | 160 | 134 | 156 | 107 | 89 | 114 | 119 | 251 | 246 | 262 | 263 |
| K13531 methylated-DNA-[protein]-cysteine S-methyltransferase [EC:2.1.1.63] | 25 | 15 | 10 | 18 | 16 | 16 | 6 | 4 | 16 | 16 | 17 | 15 |
| K14160 protein ImuA | 0 | 0 | 0 | 0 | 2 | 0 | 0 | 2 | 2 | 2 | 3 | 0 |
| K14161 protein ImuB | 2137 | 2024 | 2029 | 2181 | 1190 | 1066 | 1057 | 1241 | 700 | 647 | 658 | 624 |
| K14162 error-prone DNA polymerase [EC:2.7.7.7] | 4188 | 4165 | 4180 | 4136 | 2446 | 2297 | 2223 | 2362 | 1445 | 1469 | 1233 | 1340 |
| K14623 DNA-damage-inducible protein D | 781 | 872 | 797 | 828 | 1713 | 1611 | 1765 | 1641 | 1923 | 1788 | 1764 | 1874 |
| K14635 ATP-dependent DNA helicase MPH1 [EC:3.6.4.12] | 0 | 0 | 0 | 0 | 0 | 0 | 0 | 1 | 0 | 0 | 0 | 0 |
| K15083 DNA repair protein RAD16 | 0 | 0 | 0 | 0 | 0 | 0 | 0 | 0 | 0 | 1 | 0 | 0 |
| K15342 CRISP-associated protein Cas1 | 1315 | 1290 | 1343 | 1315 | 871 | 839 | 829 | 833 | 1332 | 1443 | 1448 | 1433 |
| K15505 DNA repair protein RAD5 [EC:3.6.4.-] | 0 | 0 | 1 | 0 | 0 | 0 | 0 | 0 | 0 | 0 | 0 | 0 |
| K16898 ATP-dependent helicase/nuclease subunit A [EC:3.1.-.- 3.6.4.12] | 1673 | 1588 | 1653 | 1628 | 528 | 494 | 488 | 500 | 1500 | 1730 | 1529 | 1664 |
| K16899 ATP-dependent helicase/nuclease subunit B [EC:3.1.-.- 3.6.4.12] | 1490 | 1466 | 1439 | 1459 | 511 | 519 | 484 | 459 | 1288 | 1617 | 1398 | 1557 |
| K17816 8-oxo-dGTP diphosphatase / 2-hydroxy-dATP diphosphatase [EC:3.6.1.55 3.6.1.56] | 2 | 0 | 0 | 0 | 0 | 2 | 0 | 0 | 0 | 0 | 0 | 0 |
| K19789 DNA repair protein RadD | 1025 | 1036 | 895 | 1019 | 670 | 772 | 725 | 633 | 1307 | 1431 | 1413 | 1422 |
| K22585 DNA repair protein NreA | 0 | 0 | 2 | 1 | 0 | 1 | 0 | 0 | 8 | 7 | 8 | 12 |
| K22803 structural maintenance of chromosomes protein 5 | 0 | 0 | 0 | 0 | 0 | 0 | 0 | 0 | 0 | 0 | 1 | 0 |
| K23258 8-oxo-dGTP phosphatase [EC:3.6.1.-] | 6 | 5 | 7 | 1 | 7 | 2 | 8 | 7 | 8 | 10 | 12 | 9 |
| K23387 golgi to ER traffic protein 4 | 0 | 1 | 0 | 0 | 0 | 0 | 0 | 0 | 0 | 0 | 0 | 0 |
| ko03029 Mitochondrial biogenesis | 136544 | 136136 | 139445 | 138257 | 146275 | 148240 | 147226 | 146676 | 136714 | 137422 | 136852 | 136371 |
| K07152 protein SCO1/2 | 1060 | 1040 | 963 | 1046 | 608 | 552 | 571 | 653 | 332 | 327 | 334 | 319 |
| K07277 outer membrane protein insertion porin family | 7870 | 7983 | 8335 | 8113 | 9370 | 9474 | 9494 | 9579 | 9689 | 9646 | 9526 | 9529 |
| K10908 DNA-directed RNA polymerase, mitochondrial [EC:2.7.7.6] | 0 | 0 | 0 | 0 | 0 | 0 | 0 | 1 | 0 | 0 | 0 | 0 |
| K14998 surfeit locus 1 family protein | 3 | 2 | 1 | 0 | 1 | 2 | 0 | 0 | 1 | 4 | 3 | 4 |
| K15980 acyl-CoA dehydrogenase family member 9 [EC:1.3.99.-] | 1 | 0 | 1 | 1 | 0 | 0 | 0 | 0 | 1 | 0 | 0 | 0 |
| K17662 cytochrome b pre-mRNA-processing protein 3 | 0 | 0 | 1 | 0 | 3 | 0 | 1 | 0 | 3 | 2 | 1 | 0 |
| K17675 ATP-dependent RNA helicase SUPV3L1/SUV3 [EC:3.6.4.13] | 113 | 93 | 97 | 106 | 81 | 89 | 79 | 68 | 71 | 97 | 80 | 79 |
| K17677 ATP-dependent helicase IRC3 [EC:3.6.4.-] | 0 | 0 | 2 | 0 | 0 | 0 | 0 | 0 | 0 | 0 | 0 | 0 |
| K17679 ATP-dependent RNA helicase MSS116, mitochondrial [EC:3.6.4.13] | 0 | 0 | 1 | 0 | 0 | 0 | 0 | 0 | 0 | 0 | 0 | 0 |
| K17680 twinkle protein [EC:3.6.4.12] | 390 | 474 | 454 | 389 | 551 | 604 | 508 | 568 | 55 | 62 | 94 | 63 |
| K17768 mitochondrial import receptor subunit TOM70 | 0 | 2 | 0 | 0 | 0 | 0 | 0 | 0 | 0 | 0 | 0 | 0 |
| K17807 mitochondrial translocator assembly and maintenance protein 41 | 0 | 0 | 1 | 0 | 0 | 0 | 0 | 0 | 0 | 0 | 0 | 0 |
| K17970 mitochondrial division protein 1 | 0 | 0 | 0 | 0 | 0 | 0 | 0 | 0 | 0 | 2 | 0 | 0 |
| K21594 translation factor GUF1, mitochondrial [EC:3.6.5.-] | 1 | 0 | 3 | 0 | 3 | 2 | 1 | 6 | 0 | 3 | 0 | 0 |
| K22140 dynamin-like GTPase MGM1, mitochondrial | 0 | 0 | 0 | 0 | 0 | 1 | 0 | 0 | 0 | 0 | 2 | 0 |
| K23164 reticulon-4-interacting protein 1, mitochondrial | 0 | 0 | 0 | 0 | 0 | 1 | 0 | 0 | 0 | 0 | 0 | 0 |
| Protein families: signaling and cellular processes | 3127866 | 3111743 | 3058229 | 3107297 | 2539537 | 2559854 | 2507899 | 2546416 | 2706008 | 2727809 | 2678755 | 2727025 |
| ko02000 Transporters | 1995062 | 1986872 | 1951522 | 1985628 | 1667605 | 1682584 | 1655177 | 1673328 | 1836688 | 1854918 | 1822917 | 1858054 |
| K00427 L-lactate permease | 737 | 674 | 752 | 728 | 201 | 229 | 209 | 219 | 1513 | 1507 | 1530 | 1450 |
| K01551 arsenite/tail-anchored protein-transporting ATPase [EC:7.3.2.7 7.3.-.-] | 1647 | 1525 | 1526 | 1657 | 2639 | 2848 | 2865 | 2629 | 534 | 457 | 558 | 528 |
| K01604 methylmalonyl-CoA decarboxylase subunit alpha [EC:7.2.4.3] | 1567 | 1633 | 1821 | 1719 | 3293 | 3430 | 3455 | 3250 | 3173 | 3189 | 3215 | 3053 |
| K01989 putative ABC transport system substrate-binding protein | 2450 | 2435 | 2356 | 2419 | 1299 | 1313 | 1229 | 1270 | 1383 | 1400 | 1364 | 1430 |
| K01990 ABC-2 type transport system ATP-binding protein | 15758 | 15670 | 15717 | 16004 | 15108 | 15149 | 15054 | 14821 | 19362 | 19930 | 19765 | 19895 |
| K01992 ABC-2 type transport system permease protein | 20466 | 20595 | 20855 | 20092 | 28545 | 28480 | 28914 | 28787 | 27918 | 27762 | 27673 | 27841 |
| K02003 putative ABC transport system ATP-binding protein | 12512 | 12289 | 12210 | 12554 | 11986 | 11650 | 11653 | 11749 | 13459 | 13674 | 13478 | 14122 |
| K02004 putative ABC transport system permease protein | 22775 | 22823 | 22800 | 22817 | 23143 | 23266 | 23652 | 23184 | 25120 | 25598 | 25341 | 25783 |
| K02013 iron complex transport system ATP-binding protein [EC:7.2.2.-] | 9086 | 8997 | 8694 | 8831 | 8791 | 9058 | 8858 | 8963 | 9168 | 9179 | 9113 | 9242 |
| K02014 iron complex outermembrane recepter protein | 74685 | 75008 | 71861 | 73145 | 55899 | 54729 | 53641 | 57046 | 42642 | 40910 | 40986 | 41452 |
| K02015 iron complex transport system permease protein | 15581 | 15376 | 14840 | 15255 | 13547 | 13429 | 13594 | 13981 | 13661 | 13463 | 13634 | 14060 |
| K02016 iron complex transport system substrate-binding protein | 10307 | 10105 | 9747 | 10109 | 9270 | 9194 | 8895 | 9323 | 9705 | 9900 | 10011 | 9985 |
| K02021 putative ABC transport system ATP-binding protein | 0 | 0 | 0 | 0 | 0 | 0 | 0 | 1 | 0 | 0 | 0 | 0 |
| K02024 maltoporin | 2821 | 2761 | 2790 | 2888 | 1824 | 1984 | 1809 | 1807 | 3894 | 4069 | 3954 | 4337 |
| K02025 multiple sugar transport system permease protein | 15750 | 14967 | 15375 | 15583 | 3515 | 3592 | 3316 | 3153 | 14524 | 15690 | 14914 | 15610 |
| K02026 multiple sugar transport system permease protein | 14983 | 14174 | 14635 | 14438 | 3272 | 3438 | 3271 | 3016 | 13812 | 15708 | 14519 | 15180 |
| K02027 multiple sugar transport system substrate-binding protein | 15180 | 15003 | 15202 | 15533 | 3487 | 3748 | 3633 | 3343 | 14730 | 16296 | 15827 | 15828 |
| K02028 polar amino acid transport system ATP-binding protein [EC:7.4.2.1] | 4117 | 3916 | 3601 | 4002 | 2076 | 2095 | 2159 | 1978 | 4158 | 4258 | 4157 | 4270 |
| K02029 polar amino acid transport system permease protein | 7663 | 7491 | 7173 | 7462 | 4810 | 4782 | 4689 | 4730 | 5456 | 5759 | 5260 | 5718 |
| K02030 polar amino acid transport system substrate-binding protein | 16306 | 16048 | 15482 | 16113 | 9011 | 9297 | 8854 | 9580 | 9705 | 9683 | 9481 | 9826 |
| K02049 NitT/TauT family transport system ATP-binding protein | 5457 | 5256 | 5017 | 5182 | 2847 | 3150 | 2750 | 2977 | 3519 | 3559 | 3351 | 3740 |
| K02050 NitT/TauT family transport system permease protein | 5152 | 5028 | 4570 | 4919 | 2990 | 3106 | 2963 | 3000 | 3271 | 3301 | 3097 | 3453 |
| K02051 NitT/TauT family transport system substrate-binding protein | 6936 | 6693 | 6507 | 6882 | 5262 | 5221 | 5086 | 5341 | 6081 | 6169 | 5933 | 6209 |
| K02056 simple sugar transport system ATP-binding protein [EC:7.5.2.-] | 8099 | 7744 | 7591 | 7783 | 4083 | 4169 | 4000 | 4108 | 6604 | 6868 | 6519 | 6897 |
| K02057 simple sugar transport system permease protein | 9377 | 8980 | 8559 | 9025 | 4311 | 4480 | 4352 | 4369 | 6831 | 7336 | 6895 | 7344 |
| K02058 simple sugar transport system substrate-binding protein | 3564 | 3368 | 3234 | 3404 | 2143 | 2125 | 1991 | 2156 | 2765 | 2740 | 2582 | 2697 |
| K02068 putative ABC transport system ATP-binding protein | 699 | 673 | 715 | 701 | 1391 | 1238 | 1296 | 1243 | 1431 | 1430 | 1452 | 1511 |
| K02069 putative ABC transport system permease protein | 805 | 820 | 843 | 889 | 1650 | 1626 | 1614 | 1681 | 1781 | 1867 | 1749 | 1688 |
| K02074 zinc/manganese transport system ATP-binding protein | 528 | 513 | 453 | 543 | 300 | 307 | 326 | 322 | 667 | 842 | 721 | 743 |
| K02075 zinc/manganese transport system permease protein | 497 | 487 | 498 | 512 | 311 | 334 | 297 | 341 | 650 | 616 | 630 | 669 |
| K02077 zinc/manganese transport system substrate-binding protein | 508 | 519 | 525 | 549 | 382 | 375 | 359 | 368 | 609 | 624 | 660 | 682 |
| K02100 MFS transporter, SP family, arabinose:H+ symporter | 774 | 766 | 711 | 848 | 534 | 556 | 565 | 508 | 1242 | 1235 | 1169 | 1190 |
| K02168 choline/glycine/proline betaine transport protein | 6289 | 5989 | 6187 | 6310 | 3389 | 3299 | 3194 | 3425 | 3303 | 3496 | 3280 | 3383 |
| K02198 cytochrome c-type biogenesis protein CcmF | 3415 | 3319 | 3064 | 3154 | 1693 | 1799 | 1799 | 1980 | 2540 | 2477 | 2511 | 2569 |
| K02205 arginine/ornithine permease | 2115 | 2003 | 1885 | 2062 | 1103 | 1098 | 1027 | 1210 | 753 | 662 | 558 | 559 |
| K02429 MFS transporter, FHS family, L-fucose permease | 3322 | 3598 | 3642 | 3406 | 8546 | 8920 | 8712 | 8532 | 6475 | 6569 | 6630 | 6628 |
| K02440 glycerol uptake facilitator protein | 1451 | 1445 | 1401 | 1525 | 756 | 877 | 762 | 780 | 1612 | 1701 | 1705 | 1704 |
| K02445 MFS transporter, OPA family, glycerol-3-phosphate transporter | 3479 | 3529 | 3410 | 3697 | 1864 | 1876 | 1869 | 1937 | 2129 | 2075 | 1933 | 2240 |
| K02511 MFS transporter, ACS family, 4-hydroxyphenylacetate permease | 519 | 587 | 569 | 554 | 163 | 178 | 151 | 141 | 1037 | 1171 | 1021 | 1068 |
| K02526 2-keto-3-deoxygluconate permease | 1009 | 1019 | 1072 | 1107 | 1190 | 1197 | 1110 | 1054 | 875 | 915 | 826 | 859 |
| K02532 MFS transporter, OHS family, lactose permease | 3396 | 3314 | 3081 | 3273 | 1889 | 2050 | 1948 | 1842 | 4822 | 5035 | 4644 | 4988 |
| K02550 glycolate permease | 7 | 9 | 10 | 11 | 19 | 16 | 19 | 21 | 68 | 86 | 77 | 82 |
| K02598 nitrite transporter | 195 | 198 | 255 | 234 | 262 | 290 | 295 | 283 | 159 | 160 | 125 | 144 |
| K02625 MFS transporter, MHS family, dicarboxylic acid transporter PcaT | 1967 | 1839 | 1849 | 1861 | 1031 | 1060 | 1006 | 1131 | 657 | 610 | 558 | 592 |
| K02824 uracil permease | 3912 | 4241 | 4036 | 4220 | 4112 | 4080 | 3809 | 3998 | 4230 | 4086 | 4011 | 3993 |
| K02856 L-rhamnose-H+ transport protein | 1029 | 1107 | 1097 | 1055 | 2250 | 2243 | 2406 | 2276 | 2463 | 2356 | 2394 | 2293 |
| K03282 large conductance mechanosensitive channel | 1276 | 1359 | 1399 | 1412 | 1342 | 1243 | 1547 | 1501 | 1300 | 1271 | 1271 | 1318 |
| K03284 magnesium transporter | 4464 | 4535 | 4778 | 4542 | 5996 | 6222 | 6328 | 6111 | 5704 | 5578 | 5543 | 5563 |
| K03286 OmpA-OmpF porin, OOP family | 2269 | 2375 | 2168 | 2248 | 1424 | 1456 | 1416 | 1379 | 1483 | 1376 | 1453 | 1404 |
| K03288 MFS transporter, MHS family, citrate/tricarballylate:H+ symporter | 768 | 759 | 664 | 774 | 208 | 211 | 258 | 217 | 1543 | 1462 | 1396 | 1574 |
| K03289 MFS transporter, NHS family, nucleoside permease | 693 | 790 | 748 | 840 | 538 | 589 | 536 | 522 | 1117 | 1183 | 1131 | 1152 |
| K03290 MFS transporter, SHS family, sialic acid transporter | 332 | 326 | 291 | 330 | 494 | 504 | 476 | 481 | 119 | 99 | 98 | 111 |
| K03291 MFS transporter, SET family, sugar efflux transporter | 1111 | 1162 | 1003 | 1151 | 673 | 607 | 623 | 608 | 1754 | 1864 | 1807 | 1925 |
| K03297 small multidrug resistance pump | 588 | 581 | 486 | 542 | 374 | 371 | 331 | 376 | 187 | 174 | 177 | 207 |

| | | | | | | | | | | | | |
|---|---|---|---|---|---|---|---|---|---|---|---|---|
| K03298 drug/metabolite transporter, DME family | 1 | 2 | 2 | 3 | 3 | 1 | 2 | 0 | 3 | 2 | 3 | 4 |
| K03299 gluconate:H+ symporter, GntP family | 5605 | 5622 | 5726 | 5728 | 4772 | 4715 | 4596 | 4503 | 4696 | 4761 | 4722 | 4740 |
| K03303 lactate permease | 3725 | 3487 | 3765 | 3611 | 4623 | 4677 | 4534 | 4498 | 3263 | 3066 | 3295 | 3077 |
| K03304 tellurite resistance protein | 500 | 510 | 533 | 550 | 348 | 346 | 381 | 338 | 855 | 734 | 733 | 737 |
| K03312 glutamate:Na+ symporter, ESS family | 2624 | 2500 | 2466 | 2458 | 1298 | 1495 | 1243 | 1414 | 1744 | 1808 | 1587 | 1820 |
| K03313 Na+:H+ antiporter, NhaA family | 1835 | 1825 | 1790 | 1764 | 3329 | 3245 | 3426 | 3241 | 2793 | 2685 | 2725 | 2706 |
| K03314 Na+:H+ antiporter, NhaB family | 1214 | 1199 | 1188 | 1134 | 500 | 480 | 443 | 419 | 1496 | 1456 | 1372 | 1491 |
| K03315 Na+:H+ antiporter, NhaC family | 2338 | 2446 | 2629 | 2346 | 3157 | 3165 | 3298 | 3195 | 3753 | 3865 | 3740 | 3579 |
| K03318 tyrosine permease | 0 | 2 | 1 | 2 | 0 | 2 | 0 | 0 | 0 | 0 | 1 | 0 |
| K03320 ammonium transporter, Amt family | 5470 | 5257 | 5260 | 5303 | 3946 | 4077 | 4340 | 4357 | 4263 | 4364 | 4142 | 4165 |
| K03321 sulfate permease, SulP family | 8560 | 8442 | 8326 | 8405 | 8420 | 8404 | 8154 | 8151 | 7011 | 6808 | 6710 | 6796 |
| K03322 manganese transport protein | 5824 | 5632 | 5244 | 5491 | 5080 | 5096 | 5105 | 5301 | 4239 | 4111 | 3976 | 4098 |
| K03324 phosphate:Na+ symporter | 10364 | 10191 | 9896 | 10012 | 8454 | 8660 | 7977 | 8418 | 7764 | 7604 | 7419 | 7624 |
| K03325 arsenite transporter | 409 | 473 | 505 | 451 | 1508 | 1367 | 1427 | 1385 | 1207 | 1178 | 1188 | 1129 |
| K03326 C4-dicarboxylate transporter, DcuC family | 1668 | 1709 | 1655 | 1711 | 1528 | 1685 | 1545 | 1617 | 2607 | 2640 | 2617 | 2735 |
| K03327 multidrug resistance protein, MATE family | 9981 | 9976 | 9482 | 9723 | 10092 | 9956 | 10059 | 10504 | 8810 | 8311 | 8595 | 8584 |
| K03329 amino acid exporter | 2 | 0 | 0 | 1 | 7 | 8 | 3 | 12 | 39 | 37 | 41 | 31 |
| K03442 small conductance mechanosensitive channel | 2944 | 2959 | 2712 | 2837 | 3050 | 3270 | 3217 | 3236 | 2817 | 2883 | 2812 | 2819 |
| K03444 MFS transporter, SP family, sugar porter, other | 0 | 2 | 5 | 0 | 0 | 2 | 2 | 0 | 2 | 0 | 0 | 0 |
| K03445 MFS transporter, DHA1 family, purine ribonucleoside efflux pump | 797 | 824 | 734 | 786 | 559 | 617 | 570 | 516 | 1272 | 1245 | 1148 | 1264 |
| K03446 MFS transporter, DHA2 family, multidrug resistance protein | 9206 | 8877 | 8779 | 8837 | 5603 | 5810 | 5574 | 5735 | 4422 | 4657 | 4267 | 4615 |
| K03449 MFS transporter, CP family, cyanate transporter | 5985 | 6012 | 5416 | 5870 | 3325 | 3363 | 3250 | 3636 | 3339 | 3247 | 3187 | 3375 |
| K03459 formate transporter | 0 | 0 | 0 | 0 | 11 | 7 | 8 | 7 | 5 | 4 | 8 | 3 |
| K03468 p-hydroxybenzoic acid efflux pump subunit AaeB | 1189 | 1229 | 1197 | 1251 | 923 | 997 | 850 | 872 | 1650 | 1737 | 1655 | 1666 |
| K03498 trk system potassium uptake protein | 6540 | 6150 | 6474 | 6269 | 5396 | 5499 | 5585 | 5508 | 6476 | 6752 | 6624 | 6798 |
| K03499 trk system potassium uptake protein | 6363 | 6470 | 6526 | 6353 | 6863 | 6957 | 6645 | 6717 | 7071 | 7199 | 6705 | 7179 |
| K03535 MFS transporter, ACS family, glucarate transporter | 1619 | 1656 | 1678 | 1786 | 1413 | 1614 | 1457 | 1437 | 2039 | 1958 | 1896 | 2057 |
| K03543 membrane fusion protein, multidrug efflux system | 7714 | 7735 | 7525 | 7461 | 6920 | 6632 | 6362 | 6499 | 5176 | 5201 | 4948 | 5145 |
| K03548 putative permease | 2158 | 2040 | 2087 | 2085 | 1276 | 1260 | 1227 | 1277 | 1273 | 1292 | 1279 | 1291 |
| K03549 KUP system potassium uptake protein | 4586 | 4672 | 4569 | 4721 | 3150 | 3239 | 3066 | 3190 | 3050 | 3007 | 2901 | 3025 |
| K03559 biopolymer transport protein ExbD | 2244 | 2081 | 2169 | 2197 | 3395 | 3416 | 3444 | 3359 | 2556 | 2630 | 2631 | 2471 |
| K03560 biopolymer transport protein TolR | 1607 | 1454 | 1406 | 1486 | 844 | 839 | 803 | 889 | 738 | 686 | 657 | 725 |
| K03561 biopolymer transport protein ExbB | 6350 | 6278 | 6177 | 6103 | 9149 | 8671 | 9081 | 8991 | 6613 | 6411 | 6445 | 6502 |
| K03562 biopolymer transport protein TolQ | 1405 | 1417 | 1511 | 1494 | 932 | 940 | 853 | 895 | 893 | 959 | 937 | 915 |
| K03640 peptidoglycan-associated lipoprotein | 984 | 1048 | 1071 | 1114 | 689 | 603 | 669 | 661 | 719 | 752 | 740 | 699 |
| K03641 TolB protein | 2490 | 2392 | 2443 | 2496 | 1436 | 1609 | 1506 | 1480 | 1707 | 1703 | 1614 | 1722 |
| K03643 LPS-assembly lipoprotein | 1026 | 1053 | 957 | 1058 | 656 | 662 | 532 | 633 | 714 | 723 | 733 | 764 |
| K03646 colicin import membrane protein | 1151 | 1163 | 1235 | 1255 | 669 | 696 | 661 | 648 | 1004 | 1074 | 1001 | 992 |
| K03756 putrescine:ornithine antiporter | 183 | 167 | 175 | 148 | 246 | 264 | 231 | 265 | 310 | 330 | 315 | 345 |
| K03757 cadaverine:lysine antiporter | 724 | 696 | 654 | 695 | 396 | 380 | 357 | 353 | 1388 | 1519 | 1472 | 1418 |
| K03758 arginine:ornithine antiporter / lysine permease | 3851 | 3720 | 3725 | 3926 | 2251 | 2275 | 2097 | 2397 | 2066 | 2062 | 1916 | 2060 |
| K03759 arginine:agmatine antiporter | 41 | 54 | 43 | 41 | 29 | 34 | 24 | 16 | 166 | 187 | 193 | 181 |
| K03761 MFS transporter, MHS family, alpha-ketoglutarate permease | 2882 | 3020 | 2856 | 2870 | 1727 | 1758 | 1736 | 1748 | 1683 | 1785 | 1751 | 1797 |
| K03762 MFS transporter, MHS family, proline/betaine transporter | 1432 | 1378 | 1372 | 1480 | 1380 | 1417 | 1344 | 1273 | 1318 | 1444 | 1196 | 1353 |
| K03811 nicotinamide mononucleotide transporter | 1727 | 1670 | 1593 | 1689 | 1730 | 1744 | 2044 | 2038 | 1677 | 1723 | 1669 | 1723 |
| K03832 periplasmic protein TonB | 5242 | 5066 | 5309 | 5333 | 8523 | 8533 | 8739 | 8627 | 6280 | 6128 | 6475 | 6249 |
| K03834 tyrosine-specific transport protein | 671 | 660 | 586 | 702 | 468 | 543 | 483 | 438 | 938 | 1060 | 986 | 1014 |
| K03835 tryptophan-specific transport protein | 3550 | 3747 | 3515 | 3612 | 2640 | 2562 | 2478 | 2591 | 1885 | 1904 | 1754 | 1912 |
| K03836 low affinity tryptophan permease | 2 | 6 | 5 | 8 | 17 | 25 | 13 | 13 | 98 | 105 | 102 | 90 |
| K03837 serine transporter | 1399 | 1389 | 1342 | 1364 | 1083 | 1141 | 992 | 1048 | 1845 | 2085 | 1828 | 1884 |
| K03838 threonine transporter | 385 | 417 | 384 | 378 | 121 | 125 | 130 | 111 | 875 | 885 | 883 | 877 |
| K03893 arsenical pump membrane protein | 2613 | 2575 | 2335 | 2528 | 1977 | 2109 | 2184 | 2086 | 1312 | 1388 | 1373 | 1355 |
| K04744 LPS-assembly protein | 6170 | 6260 | 6331 | 6229 | 3878 | 3815 | 3648 | 3765 | 3632 | 3815 | 3658 | 3821 |
| K04758 ferrous iron transport protein A | 991 | 1040 | 991 | 951 | 457 | 508 | 485 | 495 | 801 | 922 | 842 | 920 |
| K04759 ferrous iron transport protein B | 10206 | 10046 | 10215 | 10310 | 9331 | 9607 | 9350 | 9480 | 10819 | 10835 | 10638 | 10977 |
| K05020 glycine betaine transporter | 153 | 115 | 118 | 114 | 105 | 121 | 119 | 133 | 227 | 259 | 265 | 265 |
| K05041 solute carrier family 6 (neurotransmitter transporter, creatine) member 8 | 3 | 2 | 0 | 0 | 0 | 0 | 0 | 0 | 0 | 0 | 0 | 0 |
| K05245 L-carnitine/gamma-butyrobetaine antiporter | 389 | 319 | 354 | 308 | 530 | 588 | 507 | 538 | 108 | 121 | 133 | 130 |
| K05340 glucose uptake protein | 757 | 796 | 807 | 864 | 2381 | 2347 | 2435 | 2335 | 1661 | 1621 | 1716 | 1618 |
| K05373 MFS transporter, putative signal transducer | 485 | 478 | 485 | 512 | 123 | 158 | 162 | 147 | 1052 | 1205 | 1132 | 1154 |
| K05517 nucleoside-specific channel-forming protein | 2010 | 2036 | 1878 | 1806 | 1184 | 1119 | 1063 | 1280 | 1248 | 1271 | 1165 | 1244 |
| K05548 MFS transporter, AAHS family, benzoate transport protein | 2175 | 2258 | 2068 | 2029 | 1153 | 1221 | 1182 | 1266 | 749 | 659 | 622 | 676 |
| K05559 multicomponent K+:H+ antiporter subunit A | 3964 | 3873 | 3827 | 3762 | 2130 | 2136 | 1861 | 2126 | 1275 | 1204 | 1092 | 1171 |
| K05560 multicomponent K+:H+ antiporter subunit C | 529 | 491 | 473 | 459 | 295 | 331 | 267 | 280 | 159 | 155 | 136 | 152 |
| K05561 multicomponent K+:H+ antiporter subunit D | 2215 | 2234 | 2132 | 2167 | 1164 | 1166 | 1210 | 1241 | 707 | 623 | 683 | 711 |
| K05562 multicomponent K+:H+ antiporter subunit E | 653 | 649 | 657 | 753 | 364 | 367 | 299 | 265 | 229 | 215 | 197 | 200 |
| K05563 multicomponent K+:H+ antiporter subunit F | 367 | 361 | 393 | 369 | 205 | 188 | 207 | 184 | 126 | 103 | 108 | 124 |
| K05564 multicomponent K+:H+ antiporter subunit G | 490 | 542 | 457 | 536 | 300 | 219 | 276 | 278 | 177 | 134 | 180 | 156 |
| K05565 multicomponent Na+:H+ antiporter subunit A | 1 | 2 | 2 | 1 | 0 | 0 | 0 | 0 | 0 | 0 | 0 | 0 |
| K05566 multicomponent Na+:H+ antiporter subunit B | 119 | 121 | 108 | 113 | 50 | 43 | 56 | 40 | 148 | 138 | 151 | 124 |
| K05567 multicomponent Na+:H+ antiporter subunit C | 74 | 58 | 65 | 64 | 31 | 23 | 12 | 22 | 99 | 104 | 79 | 87 |
| K05568 multicomponent Na+:H+ antiporter subunit D | 592 | 495 | 509 | 530 | 230 | 240 | 191 | 195 | 597 | 684 | 635 | 623 |
| K05569 multicomponent Na+:H+ antiporter subunit E | 82 | 89 | 72 | 74 | 27 | 40 | 22 | 22 | 117 | 121 | 117 | 106 |
| K05570 multicomponent Na+:H+ antiporter subunit F | 21 | 19 | 9 | 17 | 7 | 11 | 14 | 19 | 26 | 24 | 19 | 17 |
| K05571 multicomponent Na+:H+ antiporter subunit G | 29 | 19 | 22 | 23 | 16 | 10 | 35 | 11 | 38 | 30 | 23 | 26 |
| K05595 multiple antibiotic resistance protein | 2176 | 2216 | 2201 | 2133 | 3491 | 3475 | 3544 | 3680 | 2647 | 2602 | 2768 | 2655 |
| K05777 putative thiamine transport system substrate-binding protein | 450 | 451 | 431 | 454 | 108 | 103 | 99 | 134 | 941 | 868 | 864 | 971 |
| K05778 putative thiamine transport system permease protein | 460 | 473 | 431 | 509 | 95 | 102 | 121 | 146 | 997 | 927 | 918 | 929 |
| K05779 putative thiamine transport system ATP-binding protein | 233 | 226 | 194 | 270 | 57 | 55 | 65 | 74 | 499 | 499 | 513 | 581 |
| K05782 benzoate membrane transport protein | 2391 | 2293 | 2342 | 2302 | 1343 | 1337 | 1270 | 1283 | 1405 | 1384 | 1340 | 1466 |
| K05786 chloramphenicol-sensitive protein RarD | 2529 | 2480 | 2332 | 2411 | 1582 | 1562 | 1541 | 1496 | 1380 | 1367 | 1365 | 1341 |
| K05802 potassium-dependent mechanosensitive channel | 7130 | 7157 | 6815 | 7210 | 4297 | 4188 | 4067 | 4150 | 4362 | 4353 | 4331 | 4320 |
| K05807 outer membrane protein assembly factor BamD | 2483 | 2383 | 2464 | 2572 | 2697 | 2900 | 2799 | 2791 | 2383 | 2225 | 2295 | 2322 |
| K05819 MFS transporter, AAHS family, 3-hydroxyphenylpropionic acid transporter | 423 | 463 | 368 | 427 | 104 | 115 | 107 | 112 | 832 | 805 | 825 | 811 |
| K05820 MFS transporter, PPP family, 3-phenylpropionic acid transporter | 3792 | 3807 | 3481 | 3587 | 2134 | 2038 | 1975 | 2213 | 2035 | 1943 | 1807 | 2210 |
| K05832 putative ABC transport system permease protein | 2073 | 2138 | 2061 | 2103 | 1077 | 1077 | 1084 | 1111 | 1065 | 1157 | 1156 | 1130 |
| K05833 putative ABC transport system ATP-binding protein | 1715 | 1895 | 1835 | 1729 | 897 | 891 | 877 | 931 | 988 | 1219 | 1040 | 1030 |
| K05834 homoserine/homoserine lactone efflux protein | 1323 | 1336 | 1248 | 1327 | 835 | 816 | 759 | 868 | 787 | 784 | 793 | 821 |
| K05835 threonine efflux protein | 367 | 302 | 313 | 317 | 263 | 311 | 262 | 266 | 381 | 433 | 374 | 388 |
| K06076 long-chain fatty acid transport protein | 5265 | 5057 | 5127 | 5149 | 5055 | 5389 | 5338 | 5314 | 4313 | 4297 | 4295 | 4267 |
| K06141 MFS transporter, TsgA protein | 452 | 470 | 498 | 463 | 138 | 186 | 157 | 163 | 1062 | 1034 | 982 | 1028 |
| K06144 universal stress protein B | 204 | 205 | 174 | 185 | 173 | 127 | 137 | 144 | 279 | 308 | 287 | 280 |
| K06147 ATP-binding cassette, subfamily B, bacterial | 22047 | 21960 | 22237 | 22378 | 8890 | 9173 | 8919 | 8750 | 20780 | 22532 | 21125 | 22209 |
| K06148 ATP-binding cassette, subfamily C, bacterial | 2179 | 2109 | 1971 | 1957 | 1109 | 1051 | 1085 | 1215 | 726 | 694 | 631 | 734 |
| K06155 Gnt-I system high-affinity gluconate transporter | 769 | 846 | 784 | 900 | 514 | 674 | 592 | 580 | 1052 | 1145 | 1024 | 1103 |
| K06156 Gnt-I system low-affinity gluconate transporter | 971 | 998 | 988 | 1027 | 688 | 645 | 606 | 657 | 1271 | 1266 | 1196 | 1203 |
| K06157 Gnt-II system L-idonate transporter | 286 | 308 | 283 | 296 | 441 | 477 | 406 | 385 | 66 | 85 | 87 | 74 |
| K06186 outer membrane protein assembly factor BamE | 1024 | 951 | 928 | 986 | 597 | 572 | 537 | 577 | 530 | 497 | 522 | 544 |
| K06188 aquaporin Z | 1922 | 1985 | 1892 | 1945 | 2176 | 2047 | 2312 | 2160 | 1848 | 1771 | 1921 | 1871 |
| K06189 magnesium and cobalt transporter | 1574 | 1670 | 1604 | 1632 | 1055 | 1062 | 986 | 1008 | 1180 | 1208 | 1142 | 1286 |
| K06196 cytochrome c-type biogenesis protein | 606 | 634 | 590 | 653 | 95 | 114 | 95 | 97 | 529 | 541 | 563 | 590 |
| K06199 fluoride exporter | 1325 | 1276 | 1200 | 1287 | 1434 | 1457 | 1453 | 1418 | 1045 | 1025 | 1093 | 1103 |
| K06212 formate transporter | 742 | 646 | 600 | 681 | 607 | 723 | 704 | 688 | 741 | 802 | 697 | 712 |
| K06213 magnesium transporter | 3583 | 3604 | 3747 | 3881 | 4179 | 4139 | 4066 | 4025 | 4859 | 5068 | 4903 | 5137 |
| K06216 putative ribose uptake protein | 53 | 54 | 58 | 62 | 8 | 14 | 3 | 3 | 39 | 48 | 40 | 41 |
| K06296 spore germination protein KB | 0 | 0 | 0 | 0 | 0 | 2 | 0 | 0 | 4 | 0 | 0 | 3 |
| K06311 spore germination protein | 0 | 2 | 0 | 0 | 0 | 0 | 0 | 0 | 0 | 0 | 0 | 0 |
| K06409 stage V sporulation protein B | 1486 | 1357 | 1446 | 1434 | 344 | 440 | 394 | 372 | 1237 | 1491 | 1341 | 1361 |
| K06518 holin-like protein | 889 | 902 | 853 | 887 | 1175 | 1176 | 1018 | 1138 | 1276 | 1178 | 1263 | 1269 |
| K06600 chemosensory pili system protein ChpE | 961 | 924 | 865 | 920 | 483 | 485 | 521 | 535 | 279 | 235 | 243 | 317 |
| K06609 MFS transporter, SP family, major inositol transporter | 0 | 0 | 0 | 0 | 0 | 0 | 0 | 0 | 0 | 0 | 1 | 0 |
| K06610 MFS transporter, SP family, inositol transporter | 0 | 0 | 0 | 1 | 8 | 4 | 3 | 0 | 4 | 2 | 4 | 0 |
| K06895 L-lysine exporter family protein LysE/ArgO | 648 | 553 | 562 | 613 | 358 | 360 | 354 | 354 | 602 | 664 | 690 | 635 |
| K06901 putative MFS transporter, AGZA family, xanthine/uracil permease | 8447 | 8269 | 8272 | 8372 | 4564 | 4755 | 4457 | 4680 | 6169 | 6558 | 6238 | 6662 |
| K07027 glycosyltransferase 2 family protein | 2826 | 2906 | 2784 | 2757 | 3259 | 3314 | 3280 | 3353 | 3120 | 2927 | 2981 | 2958 |
| K07084 putative amino acid transporter | 1745 | 1692 | 1648 | 1745 | 944 | 958 | 843 | 888 | 609 | 512 | 510 | 554 |
| K07114 Ca-activated chloride channel homolog | 6032 | 5957 | 6004 | 5771 | 8887 | 8877 | 8976 | 8950 | 6670 | 6443 | 6626 | 6404 |
| K07156 copper resistance protein C | 366 | 353 | 356 | 330 | 332 | 362 | 295 | 339 | 318 | 318 | 283 | 354 |
| K07221 phosphate-selective porin OprO and OprP | 3407 | 3515 | 3584 | 3566 | 2265 | 2136 | 2099 | 2261 | 1072 | 1092 | 1034 | 1033 |
| K07224 iron uptake system component EfeO | 664 | 688 | 576 | 669 | 374 | 416 | 497 | 400 | 883 | 843 | 861 | 872 |
| K07230 periplasmic iron binding protein | 121 | 134 | 102 | 122 | 190 | 229 | 159 | 214 | 5 | 10 | 8 | 7 |

| | | | | | | | | | | | | |
|---|---|---|---|---|---|---|---|---|---|---|---|---|
| K07238 zinc transporter, ZIP family | 2672 | 2497 | 2512 | 2581 | 1457 | 1481 | 1406 | 1416 | 1696 | 1668 | 1648 | 1782 |
| K07240 chromate transporter | 4549 | 4438 | 4569 | 4682 | 6586 | 6715 | 6742 | 6610 | 4796 | 4602 | 4623 | 4666 |
| K07241 nickel/cobalt transporter (NiCoT) family protein | 396 | 426 | 354 | 482 | 107 | 114 | 114 | 104 | 806 | 792 | 792 | 827 |
| K07243 high-affinity iron transporter | 4349 | 4305 | 4301 | 4236 | 2865 | 2995 | 2734 | 2947 | 1833 | 1732 | 1686 | 1842 |
| K07244 mgtE-like transporter | 0 | 3 | 0 | 0 | 3 | 0 | 2 | 3 | 6 | 3 | 1 | 12 |
| K07245 copper resistance protein D | 808 | 785 | 772 | 836 | 707 | 851 | 852 | 758 | 685 | 662 | 734 | 736 |
| K07267 porin | 5767 | 5637 | 5273 | 5421 | 2752 | 2954 | 2669 | 2936 | 2663 | 2380 | 2478 | 2559 |
| K07274 MipA family protein | 379 | 403 | 409 | 461 | 310 | 312 | 327 | 349 | 632 | 647 | 643 | 622 |
| K07278 translocation and assembly module TamA | 3693 | 3687 | 3613 | 3750 | 2206 | 2290 | 2105 | 2196 | 2327 | 2400 | 2346 | 2417 |
| K07279 autotransporter family porin | 8 | 9 | 4 | 15 | 42 | 32 | 26 | 38 | 182 | 173 | 200 | 205 |
| K07287 outer membrane protein assembly factor BamC | 1507 | 1515 | 1549 | 1611 | 964 | 1045 | 996 | 1039 | 1101 | 1230 | 1169 | 1173 |
| K07300 Ca2+:H+ antiporter | 547 | 549 | 518 | 584 | 411 | 342 | 384 | 387 | 754 | 805 | 764 | 780 |
| K07301 cation:H+ antiporter | 1288 | 1339 | 1463 | 1313 | 2777 | 2802 | 2966 | 2833 | 2666 | 2661 | 2800 | 2536 |
| K07311 Tat-targeted selenate reductase subunit YnfG | 3 | 1 | 6 | 1 | 4 | 11 | 0 | 5 | 29 | 22 | 21 | 23 |
| K07312 Tat-targeted selenate reductase subunit YnfH | 452 | 487 | 461 | 486 | 313 | 373 | 377 | 330 | 662 | 617 | 590 | 671 |
| K07490 ferrous iron transport protein C | 337 | 371 | 368 | 374 | 215 | 189 | 283 | 294 | 262 | 269 | 244 | 245 |
| K07507 putative Mg2+ transporter-C (MgtC) family protein | 4279 | 4250 | 4062 | 4178 | 3228 | 3304 | 3154 | 3275 | 2850 | 2748 | 2811 | 2671 |
| K07552 MFS transporter, DHA1 family, multidrug resistance protein | 4107 | 4112 | 3907 | 4048 | 2400 | 2347 | 2044 | 2122 | 2941 | 3107 | 2914 | 2976 |
| K07791 anaerobic C4-dicarboxylate transporter DcuA | 1670 | 1719 | 1808 | 1732 | 3848 | 3807 | 3894 | 3723 | 3296 | 3329 | 3405 | 3388 |
| K07862 serine/threonine transporter | 1389 | 1466 | 1322 | 1382 | 945 | 1031 | 933 | 958 | 1512 | 1526 | 1429 | 1530 |
| K08137 MFS transporter, SP family, galactose:H+ symporter | 816 | 819 | 849 | 888 | 578 | 668 | 575 | 592 | 1198 | 1214 | 1147 | 1232 |
| K08138 MFS transporter, SP family, xylose:H+ symportor | 766 | 889 | 1001 | 881 | 2773 | 2805 | 2782 | 2921 | 1965 | 1919 | 1840 | 1696 |
| K08151 MFS transporter, DHA1 family, tetracycline resistance protein | 1810 | 1775 | 1744 | 1658 | 1063 | 1012 | 1009 | 1050 | 648 | 545 | 620 | 573 |
| K08153 MFS transporter, DHA1 family, multidrug resistance protein | 444 | 410 | 402 | 387 | 91 | 89 | 65 | 63 | 256 | 304 | 229 | 266 |
| K08154 MFS transporter, DHA1 family, 2-module integral membrane pump EmrD | 805 | 812 | 785 | 777 | 553 | 607 | 602 | 521 | 1095 | 1081 | 1046 | 1096 |
| K08156 MFS transporter, DHA1 family, arabinose polymer utilization protein | 2020 | 2117 | 2115 | 2082 | 5723 | 5901 | 5862 | 5657 | 4823 | 4399 | 4809 | 4498 |
| K08159 MFS transporter, DHA1 family, L-arabinose/isopropyl-beta-D-thiogalactopyranoside export protein | 2370 | 2392 | 2283 | 2355 | 1492 | 1494 | 1296 | 1450 | 1560 | 1493 | 1397 | 1538 |
| K08160 MFS transporter, DHA1 family, multidrug/chloramphenicol efflux transport protein | 622 | 621 | 594 | 718 | 480 | 499 | 495 | 477 | 932 | 926 | 976 | 919 |
| K08161 MFS transporter, DHA1 family, multidrug resistance protein | 1496 | 1515 | 1426 | 1555 | 1056 | 1081 | 1085 | 1050 | 1979 | 2209 | 2010 | 2150 |
| K08162 MFS transporter, DHA1 family, multidrug resistance protein | 710 | 730 | 675 | 713 | 497 | 494 | 499 | 473 | 990 | 979 | 1005 | 1069 |
| K08163 MFS transporter, DHA1 family, multidrug resistance protein | 435 | 512 | 470 | 473 | 140 | 168 | 136 | 162 | 994 | 1105 | 1029 | 1075 |
| K08164 MFS transporter, DHA1 family, putative efflux transporter | 1 | 0 | 0 | 0 | 0 | 0 | 0 | 0 | 0 | 0 | 0 | 0 |
| K08167 MFS transporter, DHA2 family, multidrug resistance protein | 3046 | 2960 | 2684 | 2761 | 1607 | 1615 | 1478 | 1523 | 1715 | 1812 | 1695 | 1680 |
| K08168 MFS transporter, DHA2 family, metal-tetracycline-proton antiporter | 2 | 2 | 4 | 2 | 30 | 21 | 24 | 24 | 31 | 39 | 25 | 18 |
| K08169 MFS transporter, DHA2 family, multidrug resistance protein | 2454 | 2420 | 2516 | 2664 | 3939 | 3998 | 4139 | 3952 | 4453 | 4349 | 4165 | 4232 |
| K08170 MFS transporter, DHA2 family, multidrug resistance protein | 524 | 482 | 476 | 485 | 126 | 124 | 119 | 116 | 959 | 1017 | 1034 | 977 |
| K08172 MFS transporter, MHS family, shikimate and dehydroshikimate transport protein | 682 | 727 | 641 | 691 | 496 | 551 | 474 | 515 | 1064 | 1095 | 1129 | 1157 |
| K08173 MFS transporter, MHS family, metabolite:H+ symporter | 281 | 225 | 245 | 260 | 349 | 406 | 361 | 339 | 58 | 73 | 78 | 70 |
| K08177 MFS transporter, OFA family, oxalate/formate antiporter | 837 | 818 | 808 | 837 | 644 | 684 | 556 | 555 | 1122 | 1207 | 1074 | 1205 |
| K08178 MFS transporter, SHS family, lactate transporter | 420 | 393 | 410 | 411 | 100 | 135 | 115 | 105 | 882 | 843 | 855 | 900 |
| K08191 MFS transporter, ACS family, hexuronate transporter | 4812 | 4750 | 4790 | 4884 | 5614 | 5871 | 5823 | 5646 | 5393 | 5181 | 5238 | 5342 |
| K08194 MFS transporter, ACS family, D-galactonate transporter | 4318 | 4192 | 3928 | 4193 | 2663 | 2682 | 2609 | 2775 | 3572 | 3683 | 3550 | 3564 |
| K08195 MFS transporter, AAHS family, 4-hydroxybenzoate transporter | 5676 | 5946 | 5437 | 5874 | 3160 | 3015 | 3076 | 3209 | 3449 | 3327 | 3229 | 3577 |
| K08196 MFS transporter, AAHS family, cis,cis-muconate transporter | 1895 | 1847 | 1806 | 1889 | 1084 | 1057 | 971 | 1074 | 682 | 555 | 535 | 610 |
| K08217 MFS transporter, DHA3 family, macrolide efflux protein | 544 | 618 | 580 | 635 | 383 | 414 | 411 | 416 | 607 | 638 | 619 | 669 |
| K08219 MFS transporter, UMF2 family, putative MFS family transporter protein | 646 | 617 | 628 | 673 | 433 | 459 | 417 | 431 | 862 | 960 | 878 | 882 |
| K08221 MFS transporter, ACDE family, multidrug resistance protein | 0 | 0 | 0 | 0 | 14 | 6 | 3 | 8 | 14 | 16 | 13 | 16 |
| K08222 MFS transporter, YQGE family, putative transporter | 835 | 948 | 959 | 875 | 2049 | 1975 | 2264 | 2199 | 1833 | 1780 | 1897 | 1733 |
| K08223 MFS transporter, FSR family, fosmidomycin resistance protein | 1553 | 1424 | 1517 | 1532 | 2943 | 2971 | 3050 | 2794 | 2851 | 2852 | 3026 | 2785 |
| K08224 MFS transporter, YNFM family, putative membrane transport protein | 2471 | 2357 | 2302 | 2268 | 1321 | 1349 | 1332 | 1426 | 1562 | 1522 | 1542 | 1520 |
| K08225 MFS transporter, ENTS family, enterobactin (siderophore) exporter | 736 | 682 | 659 | 691 | 490 | 578 | 489 | 497 | 852 | 897 | 874 | 959 |
| K08227 MFS transporter, LPLT family, lysophospholipid transporter | 783 | 727 | 670 | 736 | 467 | 522 | 469 | 444 | 1039 | 1058 | 899 | 1042 |
| K08344 suppressor for copper-sensitivity B | 719 | 703 | 660 | 725 | 200 | 209 | 205 | 200 | 1387 | 1391 | 1399 | 1476 |
| K08363 mercuric ion transport protein | 707 | 685 | 727 | 671 | 543 | 539 | 518 | 531 | 176 | 158 | 179 | 171 |
| K08364 periplasmic mercuric ion binding protein | 658 | 709 | 699 | 685 | 1024 | 1006 | 941 | 931 | 594 | 542 | 591 | 484 |
| K08368 MFS transporter, putative metabolite transport protein | 2728 | 2606 | 2384 | 2582 | 1693 | 1739 | 1652 | 1684 | 876 | 841 | 756 | 890 |
| K08369 MFS transporter, putative metabolite:H+ symporter | 2502 | 2455 | 2304 | 2381 | 1454 | 1571 | 1501 | 1554 | 3001 | 3285 | 3250 | 3410 |
| K08715 inward rectifier potassium channel | 1 | 0 | 0 | 0 | 0 | 0 | 0 | 0 | 0 | 0 | 0 | 0 |
| K08717 urea transporter | 960 | 938 | 892 | 915 | 545 | 516 | 703 | 775 | 325 | 268 | 273 | 288 |
| K08721 outer membrane protein, multidrug efflux system | 2188 | 2349 | 2128 | 2155 | 1251 | 1200 | 1137 | 1227 | 709 | 657 | 680 | 745 |
| K08970 nickel/cobalt transporter (NicO) family protein | 1990 | 2003 | 1784 | 2054 | 1725 | 1627 | 1665 | 1653 | 2097 | 2082 | 2084 | 2054 |
| K08978 bacterial/archaeal transporter family protein | 1151 | 1029 | 1120 | 1103 | 613 | 684 | 606 | 613 | 560 | 563 | 580 | 578 |
| K08994 ion channel-forming bestrophin family protein | 501 | 474 | 452 | 536 | 443 | 390 | 352 | 339 | 733 | 651 | 700 | 694 |
| K09013 Fe-S cluster assembly ATP-binding protein | 1350 | 1340 | 1424 | 1388 | 2165 | 2184 | 2178 | 2234 | 2334 | 2228 | 2247 | 2268 |
| K09016 putative pyrimidine permease RutG | 521 | 524 | 474 | 484 | 151 | 157 | 141 | 140 | 1082 | 1159 | 1007 | 1098 |
| K09771 small multidrug resistance family-3 protein | 736 | 649 | 695 | 657 | 425 | 380 | 363 | 398 | 399 | 464 | 439 | 501 |
| K09774 lipopolysaccharide export system protein LptA | 1153 | 1114 | 1195 | 1200 | 732 | 799 | 736 | 729 | 734 | 752 | 753 | 741 |
| K09800 translocation and assembly module TamB | 7231 | 7086 | 6797 | 7026 | 4087 | 4143 | 3906 | 4241 | 5067 | 5007 | 4828 | 5004 |
| K09820 manganese/iron transport system ATP-binding protein | 2 | 2 | 2 | 3 | 2 | 2 | 0 | 4 | 3 | 4 | 2 | 1 |
| K10120 fructooligosaccharide transport system substrate-binding protein | 434 | 417 | 431 | 461 | 25 | 18 | 25 | 10 | 423 | 398 | 408 | 418 |
| K10121 fructooligosaccharide transport system permease protein | 35 | 33 | 23 | 42 | 4 | 6 | 8 | 3 | 14 | 18 | 16 | 17 |
| K10122 fructooligosaccharide transport system permease protein | 20 | 19 | 21 | 17 | 4 | 5 | 1 | 5 | 15 | 9 | 16 | 9 |
| K10124 carbohydrate-specific outer membrane porin | 1033 | 1038 | 1012 | 1033 | 796 | 877 | 770 | 800 | 1536 | 1629 | 1494 | 1580 |
| K10716 voltage-gated potassium channel | 2068 | 2113 | 2008 | 2033 | 2630 | 2547 | 2537 | 2537 | 2502 | 2393 | 2517 | 2513 |
| K10820 galactofuranose transport system ATP-binding protein [EC:7.5.2.9] | 1190 | 1202 | 1120 | 1138 | 867 | 838 | 828 | 808 | 1484 | 1583 | 1334 | 1567 |
| K10974 cytosine permease | 2611 | 2544 | 2304 | 2460 | 1504 | 1425 | 1407 | 1384 | 1740 | 1765 | 1615 | 1718 |
| K10975 allantoin permease | 96 | 102 | 71 | 82 | 20 | 30 | 28 | 23 | 108 | 136 | 124 | 112 |
| K11005 hemolysin A | 10 | 1 | 12 | 9 | 3 | 5 | 2 | 2 | 123 | 128 | 138 | 146 |
| K11102 proton glutamate symport protein | 3256 | 3243 | 3174 | 3259 | 1928 | 1965 | 1833 | 1944 | 1870 | 1913 | 1796 | 1852 |
| K11104 melibiose permease | 675 | 667 | 642 | 729 | 209 | 227 | 196 | 195 | 1299 | 1279 | 1275 | 1357 |
| K11105 cell volume regulation protein A | 4807 | 4894 | 4807 | 4941 | 4982 | 5047 | 4974 | 5104 | 4248 | 4185 | 4234 | 4255 |
| K11106 L-tartrate/succinate antiporter | 798 | 782 | 741 | 754 | 566 | 600 | 508 | 486 | 1107 | 1199 | 1079 | 1092 |
| K11189 phosphocarrier protein | 4139 | 4082 | 3843 | 3940 | 2073 | 2265 | 2245 | 2483 | 1867 | 1826 | 1730 | 1855 |
| K11201 PTS system, fructose-specific IIA-like component [EC:2.7.1.-] | 0 | 0 | 0 | 0 | 8 | 10 | 1 | 5 | 0 | 2 | 6 | 2 |
| K11202 PTS system, fructose-specific IIB-like component [EC:2.7.1.-] | 104 | 98 | 130 | 129 | 35 | 40 | 50 | 37 | 115 | 153 | 160 | 154 |
| K11203 PTS system, fructose-specific IIC-like component | 437 | 427 | 418 | 414 | 120 | 146 | 100 | 100 | 372 | 398 | 326 | 372 |
| K11249 cysteine/O-acetylserine efflux protein | 316 | 371 | 348 | 362 | 267 | 311 | 255 | 244 | 451 | 502 | 498 | 485 |
| K11250 leucine efflux protein | 1332 | 1308 | 1304 | 1362 | 924 | 928 | 861 | 931 | 695 | 726 | 717 | 689 |
| K11535 nucleoside transport protein | 637 | 758 | 613 | 790 | 498 | 510 | 519 | 471 | 923 | 1043 | 967 | 1113 |
| K11537 MFS transporter, NHS family, xanthosine permease | 734 | 697 | 889 | 794 | 2586 | 2546 | 2624 | 2477 | 1910 | 1885 | 1846 | 1833 |
| K11719 lipopolysaccharide export system protein LptC | 1199 | 1257 | 1220 | 1217 | 684 | 782 | 751 | 741 | 773 | 832 | 743 | 797 |
| K11732 phenylalanine-specific permease | 822 | 820 | 685 | 911 | 607 | 703 | 631 | 576 | 1100 | 1085 | 1069 | 1265 |
| K11733 lysine-specific permease | 3317 | 3324 | 3147 | 3271 | 1952 | 2005 | 1976 | 1934 | 1810 | 1895 | 1841 | 1898 |
| K11734 aromatic amino acid transport protein AroP | 4928 | 4803 | 4785 | 4885 | 2879 | 2979 | 2781 | 2953 | 3294 | 3275 | 3292 | 3163 |
| K11735 GABA permease | 2451 | 2330 | 2318 | 2325 | 1182 | 1130 | 1112 | 1188 | 1768 | 1658 | 1569 | 1686 |
| K11736 proline-specific permease ProY | 726 | 752 | 738 | 778 | 543 | 581 | 556 | 564 | 1024 | 1015 | 1024 | 1054 |
| K11737 D-serine/D-alanine/glycine transporter | 812 | 843 | 839 | 833 | 662 | 626 | 582 | 598 | 1216 | 1215 | 1181 | 1258 |
| K11738 L-asparagine permease | 764 | 763 | 821 | 732 | 544 | 552 | 565 | 527 | 933 | 1122 | 1054 | 1044 |
| K11741 quaternary ammonium compound-resistance protein SugE | 2736 | 2873 | 2904 | 2805 | 2151 | 2151 | 1962 | 2153 | 1583 | 1496 | 1510 | 1516 |
| K11742 spermidine export protein MdtI | 583 | 564 | 551 | 497 | 321 | 362 | 318 | 346 | 351 | 341 | 340 | 341 |
| K11743 spermidine export protein MdtJ | 681 | 703 | 674 | 663 | 370 | 428 | 371 | 405 | 403 | 367 | 377 | 409 |
| K11745 glutathione-regulated potassium-efflux system ancillary protein KefC | 1066 | 1113 | 1000 | 1157 | 785 | 870 | 810 | 790 | 1430 | 1461 | 1365 | 1476 |
| K11746 glutathione-regulated potassium-efflux system ancillary protein KefF | 376 | 338 | 347 | 366 | 224 | 246 | 227 | 237 | 474 | 457 | 471 | 494 |
| K11747 glutathione-regulated potassium-efflux system protein KefB | 1131 | 1108 | 1079 | 1168 | 828 | 835 | 866 | 875 | 1501 | 1564 | 1485 | 1578 |
| K11748 glutathione-regulated potassium-efflux system ancillary protein KefG | 396 | 393 | 345 | 372 | 270 | 274 | 263 | 273 | 531 | 516 | 611 | 517 |
| K11815 multidrug resistance protein EbrB | 1 | 0 | 0 | 0 | 0 | 0 | 0 | 0 | 0 | 0 | 0 | 0 |
| K11928 sodium/proline symporter | 3526 | 3470 | 3322 | 3456 | 2168 | 2094 | 2041 | 2086 | 2142 | 2037 | 2141 | 2240 |
| K11929 outer membrane pore protein E | 669 | 672 | 611 | 706 | 480 | 559 | 503 | 453 | 934 | 858 | 767 | 872 |
| K11934 outer membrane protein X | 429 | 441 | 366 | 432 | 401 | 416 | 387 | 396 | 334 | 391 | 378 | 434 |
| K11939 inner membrane transporter RhtA | 1759 | 1708 | 1571 | 1692 | 979 | 970 | 912 | 971 | 1051 | 1009 | 1013 | 1112 |
| K12041 solute carrier family 9 (sodium/hydrogen exchanger), member 6/7 | 0 | 0 | 2 | 0 | 0 | 0 | 0 | 0 | 0 | 0 | 0 | 0 |
| K12299 MFS transporter, ACS family, probable galactarate transporter | 802 | 754 | 692 | 874 | 560 | 584 | 610 | 517 | 1076 | 1146 | 1064 | 1121 |
| K12516 putative surface-exposed virulence protein | 8 | 9 | 8 | 3 | 11 | 6 | 13 | 7 | 119 | 135 | 130 | 104 |
| K12537 membrane fusion protein, protease secretion system | 3507 | 3194 | 3022 | 3187 | 1725 | 1696 | 1522 | 1898 | 1048 | 915 | 926 | 958 |
| K12538 outer membrane protein, protease secretion system | 3813 | 3736 | 3479 | 3506 | 2096 | 1862 | 1751 | 2108 | 1215 | 1067 | 1041 | 1146 |
| K12542 membrane fusion protein, adhesin transport system | 1898 | 1985 | 1790 | 1904 | 1516 | 1568 | 1388 | 1430 | 543 | 526 | 473 | 513 |
| K12543 outer membrane protein, adhesin transport system | 2372 | 2280 | 2195 | 2200 | 1685 | 1638 | 1471 | 1687 | 565 | 632 | 553 | 649 |
| K12545 heme acquisition protein HasA | 323 | 301 | 275 | 278 | 178 | 155 | 131 | 155 | 115 | 76 | 83 | 94 |
| K12678 autotransporter family porin | 110 | 95 | 119 | 101 | 62 | 56 | 58 | 62 | 217 | 236 | 214 | 242 |
| K12684 serine protease autotransporter [EC:3.4.21.-] | 12 | 7 | 19 | 19 | 26 | 39 | 14 | 25 | 251 | 314 | 302 | 304 |
| K12685 subtilase-type serine protease [EC:3.4.21.-] | 3560 | 3382 | 3283 | 3310 | 1978 | 1900 | 1729 | 2046 | 1219 | 1184 | 1140 | 1208 |

| | | | | | | | | | | | | |
|---|---|---|---|---|---|---|---|---|---|---|---|---|
| K12686 outer membrane lipase/esterase | 3023 | 3036 | 2692 | 2824 | 1583 | 1613 | 1628 | 1830 | 960 | 854 | 853 | 918 |
| K13283 ferrous-iron efflux pump FieF | 571 | 533 | 556 | 591 | 362 | 379 | 365 | 389 | 748 | 716 | 731 | 771 |
| K13629 D-serine transporter | 849 | 836 | 843 | 911 | 613 | 673 | 618 | 571 | 1189 | 1184 | 1246 | 1257 |
| K13888 membrane fusion protein, macrolide-specific efflux system | 1819 | 1798 | 1699 | 1714 | 1124 | 1061 | 1021 | 1118 | 976 | 1055 | 922 | 960 |
| K13929 malonate decarboxylase alpha subunit [EC:2.3.1.187] | 4267 | 4201 | 3664 | 3937 | 2221 | 2263 | 2049 | 2294 | 2142 | 2155 | 2003 | 2192 |
| K13931 malonate decarboxylase delta subunit | 710 | 752 | 639 | 624 | 395 | 382 | 353 | 369 | 389 | 373 | 371 | 359 |
| K13936 malonate transporter | 481 | 525 | 456 | 540 | 389 | 415 | 397 | 405 | 665 | 655 | 594 | 648 |
| K14052 putrescine importer | 2357 | 2203 | 2308 | 2165 | 1119 | 1108 | 941 | 1029 | 1503 | 1436 | 1446 | 1586 |
| K14053 outer membrane protein G | 3 | 2 | 6 | 8 | 8 | 12 | 4 | 10 | 48 | 54 | 44 | 33 |
| K14062 outer membrane protein N | 367 | 332 | 357 | 370 | 90 | 111 | 76 | 79 | 725 | 772 | 728 | 779 |
| K14347 solute carrier family 10 (sodium/bile acid cotransporter), member 7 | 2024 | 1896 | 1808 | 1979 | 1126 | 1148 | 1127 | 1186 | 1285 | 1177 | 1238 | 1183 |
| K14392 sodium/pantothenate symporter | 1100 | 1092 | 1007 | 1050 | 753 | 824 | 770 | 630 | 1480 | 1581 | 1473 | 1478 |
| K14393 cation/acetate symporter | 5003 | 4954 | 5039 | 5245 | 2927 | 2925 | 2803 | 3006 | 2501 | 2619 | 2594 | 2736 |
| K14430 phosphate transporter | 0 | 0 | 0 | 0 | 0 | 0 | 0 | 0 | 1 | 0 | 0 | 0 |
| K14445 solute carrier family 13 (sodium-dependent dicarboxylate transporter), member 2/3/5 | 1047 | 1003 | 1145 | 1080 | 3168 | 3097 | 3173 | 3183 | 2600 | 2458 | 2565 | 2654 |
| K14717 solute carrier family 39 (zinc transporter), member 11 | 13 | 7 | 11 | 6 | 8 | 14 | 9 | 12 | 12 | 8 | 7 | 9 |
| K15108 solute carrier family 25 (mitochondrial thiamine pyrophosphate transporter), member 19 | 0 | 0 | 0 | 0 | 0 | 0 | 0 | 0 | 1 | 0 | 0 | 0 |
| K15110 solute carrier family 25 (mitochondrial 2-oxodicarboxylate transporter), member 21 | 0 | 0 | 0 | 0 | 1 | 0 | 0 | 0 | 0 | 0 | 0 | 0 |
| K15268 O-acetylserine/cysteine efflux transporter | 1970 | 1899 | 1784 | 1844 | 1094 | 1159 | 1026 | 1139 | 1144 | 1139 | 1145 | 1101 |
| K15269 probable blue pigment (indigoidine) exporter | 338 | 371 | 350 | 373 | 77 | 92 | 91 | 65 | 639 | 748 | 750 | 750 |
| K15270 S-adenosylmethionine uptake transporter | 3 | 4 | 3 | 3 | 3 | 4 | 8 | 3 | 11 | 6 | 3 | 8 |
| K15272 solute carrier family 35 (UDP-sugar transporter), member A1/2/3 | 0 | 0 | 1 | 0 | 0 | 0 | 0 | 0 | 0 | 0 | 0 | 0 |
| K15280 solute carrier family 35, member C2 | 1 | 0 | 0 | 0 | 0 | 0 | 0 | 0 | 0 | 0 | 0 | 0 |
| K15547 multidrug resistance protein MdtO | 1209 | 1181 | 1105 | 1197 | 803 | 791 | 725 | 748 | 1715 | 1738 | 1620 | 1700 |
| K15548 p-hydroxybenzoic acid efflux pump subunit AaeA | 588 | 583 | 604 | 638 | 420 | 472 | 424 | 408 | 799 | 858 | 736 | 893 |
| K15549 membrane fusion protein, multidrug efflux system | 603 | 579 | 582 | 574 | 438 | 392 | 416 | 359 | 829 | 853 | 891 | 970 |
| K15550 outer membrane protein, multidrug efflux system | 845 | 824 | 754 | 723 | 483 | 506 | 463 | 470 | 1068 | 1080 | 1119 | 1095 |
| K15721 pesticin/yersiniabactin receptor | 675 | 770 | 727 | 803 | 208 | 210 | 203 | 192 | 1625 | 1637 | 1605 | 1605 |
| K15725 outer membrane protein, cobalt-zinc-cadmium efflux system | 3821 | 3508 | 3564 | 3627 | 2026 | 2041 | 1891 | 1983 | 1160 | 1073 | 964 | 1092 |
| K15726 cobalt-zinc-cadmium resistance protein CzcA | 10540 | 10333 | 10489 | 10435 | 12447 | 12757 | 12708 | 12826 | 8708 | 8118 | 8519 | 7979 |
| K15727 membrane fusion protein, cobalt-zinc-cadmium efflux system | 3308 | 3243 | 3487 | 3257 | 3506 | 3594 | 3318 | 3525 | 2360 | 2243 | 2316 | 2102 |
| K15738 ABC transport system ATP-binding/permease protein | 4465 | 4334 | 4355 | 4388 | 5181 | 5133 | 5240 | 5111 | 5337 | 5521 | 5472 | 5435 |
| K15987 K(+)-stimulated pyrophosphate-energized sodium pump | 2949 | 3148 | 3194 | 3059 | 5609 | 5644 | 5686 | 5608 | 4930 | 4979 | 5001 | 4975 |
| K16052 MscS family membrane protein | 41 | 39 | 44 | 41 | 30 | 23 | 27 | 19 | 133 | 142 | 124 | 155 |
| K16053 miniconductance mechanosensitive channel | 3802 | 3897 | 3754 | 3889 | 4336 | 4531 | 4417 | 4567 | 4739 | 4543 | 4543 | 4538 |
| K16074 zinc transporter | 3579 | 3712 | 3694 | 3537 | 1817 | 1849 | 1837 | 1964 | 2257 | 2114 | 2118 | 2162 |
| K16075 magnesium transporter | 0 | 0 | 0 | 0 | 0 | 0 | 0 | 0 | 0 | 0 | 2 | 0 |
| K16076 outer membrane porin protein LC | 5 | 2 | 2 | 8 | 8 | 13 | 8 | 16 | 86 | 126 | 108 | 100 |
| K16077 sucrose porin | 1426 | 1420 | 1377 | 1528 | 1026 | 984 | 959 | 967 | 1849 | 1936 | 1811 | 1921 |
| K16079 outer membrane immunogenic protein | 0 | 2 | 0 | 5 | 0 | 4 | 2 | 3 | 4 | 6 | 3 | 7 |
| K16081 alginate production protein | 2042 | 2184 | 1876 | 2048 | 1185 | 1066 | 1038 | 1187 | 642 | 636 | 518 | 663 |
| K16087 hemoglobin/transferrin/lactoferrin receptor protein | 11754 | 11828 | 10885 | 11545 | 6456 | 6321 | 6136 | 6781 | 5365 | 5210 | 5219 | 5470 |
| K16088 outer-membrane receptor for ferric coprogen and ferric-rhodotorulic acid | 11032 | 11074 | 10190 | 10380 | 6219 | 6107 | 5617 | 6412 | 3692 | 3173 | 3140 | 3454 |
| K16089 outer membrane receptor for ferrienterochelin and colicins | 9650 | 9626 | 9649 | 9262 | 18389 | 18612 | 18690 | 18428 | 13848 | 13639 | 13875 | 13395 |
| K16090 catecholate siderophore receptor | 7795 | 7608 | 7613 | 7529 | 4198 | 3861 | 3736 | 3973 | 3959 | 3960 | 3790 | 3974 |
| K16091 Fe(3+) dicitrate transport protein | 4296 | 4454 | 4305 | 4162 | 2450 | 2335 | 2314 | 2538 | 1569 | 1572 | 1447 | 1505 |
| K16092 vitamin B12 transporter | 3982 | 3985 | 3489 | 3573 | 2120 | 2109 | 2119 | 2368 | 2572 | 2604 | 2510 | 2561 |
| K16139 glucuronide carrier protein | 12 | 5 | 3 | 6 | 8 | 14 | 8 | 9 | 48 | 28 | 45 | 31 |
| K16140 putative glucuronide porin | 2 | 1 | 3 | 1 | 28 | 16 | 19 | 10 | 70 | 59 | 61 | 61 |
| K16152 heme acquisition protein HasR | 4277 | 4068 | 3673 | 3748 | 2095 | 2051 | 1948 | 2268 | 1231 | 1105 | 1182 | 1218 |
| K16169 xanthine permease | 104 | 88 | 94 | 89 | 23 | 34 | 29 | 16 | 105 | 90 | 94 | 106 |
| K16209 lactose/raffinose/galactose permease | 30 | 31 | 37 | 27 | 32 | 36 | 25 | 27 | 90 | 89 | 77 | 103 |
| K16210 oligogalacturonide transporter | 2025 | 1971 | 1977 | 2177 | 1797 | 1802 | 1558 | 1764 | 2202 | 2400 | 2296 | 2392 |
| K16211 maltose/moltooligosaccharide transporter | 154 | 201 | 199 | 193 | 267 | 276 | 245 | 280 | 20 | 44 | 34 | 33 |
| K16234 histidine transporter | 2961 | 2887 | 2786 | 3074 | 1468 | 1397 | 1407 | 1581 | 1741 | 1851 | 1706 | 1702 |
| K16235 S-methylmethionine transporter | 554 | 523 | 518 | 590 | 146 | 175 | 170 | 133 | 1091 | 1137 | 1082 | 1159 |
| K16238 ethanolamine permease | 2211 | 2222 | 2167 | 2177 | 1272 | 1349 | 1308 | 1319 | 1561 | 1571 | 1406 | 1577 |
| K16248 probable glucitol transport protein GutA | 139 | 129 | 111 | 126 | 189 | 165 | 164 | 155 | 460 | 495 | 454 | 437 |
| K16261 yeast amino acid transporter | 0 | 2 | 0 | 0 | 2 | 0 | 0 | 0 | 0 | 0 | 0 | 0 |
| K16263 amino acid efflux transporter | 781 | 746 | 740 | 817 | 535 | 603 | 491 | 572 | 987 | 1030 | 974 | 976 |
| K16264 cobalt-zinc-cadmium efflux system protein | 2777 | 2676 | 2739 | 2675 | 3202 | 3400 | 3319 | 3370 | 2347 | 2211 | 2272 | 2250 |
| K16300 membrane fusion protein, epimerase transport system | 5 | 2 | 0 | 3 | 0 | 0 | 0 | 0 | 0 | 0 | 0 | 0 |
| K16301 deferrochelatase/peroxidase EfeB [EC:1.11.1.-] | 753 | 689 | 732 | 740 | 509 | 564 | 549 | 529 | 983 | 1098 | 1025 | 1033 |
| K16321 high-affinity gluconate transporter | 2 | 0 | 1 | 1 | 8 | 21 | 8 | 13 | 11 | 16 | 3 | 7 |
| K16322 low-affinity inorganic phosphate transporter | 2740 | 2768 | 2554 | 2665 | 1778 | 1712 | 1721 | 1689 | 2081 | 2077 | 1966 | 2023 |
| K16323 purine nucleoside transport protein | 0 | 0 | 3 | 0 | 0 | 0 | 0 | 0 | 0 | 0 | 0 | 0 |
| K16324 putative pseudouridine transporter | 365 | 343 | 362 | 381 | 530 | 665 | 602 | 675 | 62 | 84 | 79 | 71 |
| K16325 nucleoside permease | 81 | 110 | 72 | 76 | 115 | 126 | 98 | 94 | 62 | 58 | 77 | 68 |
| K16331 sulfate permease | 3 | 0 | 0 | 0 | 0 | 0 | 0 | 0 | 0 | 0 | 0 | 0 |
| K16345 xanthine permease XanP | 7074 | 7194 | 6663 | 6988 | 6412 | 6577 | 6115 | 6552 | 5634 | 5409 | 5464 | 5535 |
| K16346 xanthine permease XanQ | 3 | 9 | 1 | 6 | 9 | 27 | 24 | 19 | 93 | 89 | 80 | 87 |
| K16347 entericidin A | 76 | 84 | 75 | 83 | 68 | 67 | 55 | 48 | 132 | 119 | 131 | 107 |
| K16348 entericidin B | 121 | 95 | 103 | 127 | 71 | 69 | 73 | 62 | 150 | 177 | 157 | 156 |
| K16693 enterobacterial common antigen flippase | 772 | 783 | 755 | 716 | 510 | 580 | 507 | 539 | 1031 | 1175 | 1156 | 1214 |
| K16695 lipopolysaccharide exporter | 261 | 287 | 239 | 274 | 454 | 431 | 432 | 372 | 67 | 54 | 61 | 52 |
| K16788 niacin transporter | 62 | 40 | 34 | 45 | 11 | 11 | 11 | 12 | 77 | 62 | 40 | 49 |
| K16789 thiamine transporter | 297 | 319 | 337 | 349 | 130 | 145 | 118 | 111 | 257 | 311 | 257 | 294 |
| K16923 energy-coupling factor transport system substrate-specific component | 268 | 293 | 325 | 314 | 89 | 77 | 113 | 91 | 253 | 304 | 242 | 273 |
| K16924 energy-coupling factor transport system substrate-specific component | 233 | 288 | 224 | 243 | 46 | 41 | 55 | 43 | 262 | 274 | 245 | 258 |
| K16925 energy-coupling factor transport system substrate-specific component | 45 | 39 | 50 | 44 | 4 | 10 | 8 | 7 | 30 | 72 | 45 | 24 |
| K16926 energy-coupling factor transport system substrate-specific component | 160 | 164 | 178 | 185 | 138 | 135 | 100 | 135 | 524 | 506 | 497 | 528 |
| K16927 energy-coupling factor transport system substrate-specific component | 52 | 53 | 65 | 69 | 50 | 36 | 58 | 60 | 169 | 150 | 156 | 125 |
| K16928 energy-coupling factor transport system substrate-specific component | 0 | 0 | 0 | 0 | 0 | 0 | 0 | 1 | 0 | 1 | 1 | 0 |
| K16929 energy-coupling factor transport system substrate-specific component | 0 | 0 | 0 | 0 | 0 | 1 | 2 | 1 | 1 | 0 | 2 | 1 |
| K17713 outer membrane protein assembly factor BamB | 2467 | 2626 | 2483 | 2550 | 1556 | 1606 | 1471 | 1444 | 1601 | 1665 | 1475 | 1637 |
| K17938 peptide/bleomycin uptake transporter | 703 | 706 | 671 | 704 | 548 | 559 | 525 | 521 | 922 | 1119 | 991 | 1038 |
| K18141 membrane fusion protein, multidrug efflux system | 470 | 504 | 400 | 507 | 163 | 130 | 172 | 136 | 976 | 966 | 938 | 990 |
| K18142 multidrug efflux pump | 1346 | 1276 | 1220 | 1315 | 443 | 455 | 439 | 424 | 2742 | 2595 | 2697 | 2740 |
| K18214 MFS transporter, DHA3 family, tetracycline resistance protein | 17 | 19 | 23 | 11 | 6 | 6 | 5 | 20 | 20 | 14 | 13 | 31 |
| K18218 tetracycline resistance efflux pump | 1 | 2 | 1 | 1 | 4 | 7 | 3 | 3 | 0 | 1 | 2 | 1 |
| K18295 membrane fusion protein, multidrug efflux system | 1844 | 1843 | 1576 | 1675 | 978 | 994 | 798 | 905 | 603 | 550 | 477 | 581 |
| K18296 multidrug efflux pump | 5157 | 5119 | 4830 | 4644 | 2762 | 2580 | 2597 | 2930 | 1517 | 1564 | 1515 | 1626 |
| K18298 membrane fusion protein, multidrug efflux system | 1727 | 1726 | 1573 | 1633 | 914 | 911 | 862 | 892 | 501 | 442 | 436 | 505 |
| K18299 multidrug efflux pump | 3503 | 3461 | 3456 | 3280 | 1730 | 1782 | 1641 | 1913 | 1095 | 989 | 934 | 984 |
| K18300 outer membrane protein, multidrug efflux system | 708 | 678 | 673 | 677 | 347 | 345 | 366 | 339 | 202 | 200 | 175 | 222 |
| K18302 membrane fusion protein, multidrug efflux system | 1510 | 1522 | 1426 | 1530 | 834 | 803 | 722 | 928 | 562 | 503 | 514 | 547 |
| K18303 multidrug efflux pump | 4838 | 4849 | 4860 | 4685 | 2706 | 2615 | 2506 | 2689 | 1626 | 1543 | 1534 | 1619 |
| K18305 transmembrane protein | 796 | 831 | 738 | 792 | 438 | 428 | 369 | 480 | 226 | 240 | 224 | 227 |
| K18308 outer membrane protein, multidrug efflux system | 2500 | 2302 | 1989 | 2175 | 1150 | 1119 | 1062 | 1269 | 758 | 616 | 607 | 677 |
| K18480 cholesterol transport system auxiliary component | 1666 | 1617 | 1651 | 1627 | 864 | 849 | 828 | 968 | 464 | 459 | 480 | 459 |
| K18552 MFS transporter, DHA1 family, chloramphenicol/florfenicol resistance protein | 2 | 3 | 0 | 0 | 0 | 4 | 0 | 0 | 4 | 4 | 1 | 3 |
| K18567 MFS transporter, DHA1 family, purine base/nucleoside efflux pump | 2720 | 2793 | 2529 | 2663 | 1706 | 1555 | 1700 | 1797 | 786 | 781 | 720 | 775 |
| K18814 putative inorganic carbon (hco3(-)) transporter | 73 | 58 | 50 | 61 | 24 | 16 | 25 | 26 | 64 | 104 | 82 | 82 |
| K18898 membrane fusion protein, multidrug efflux system | 3 | 4 | 3 | 3 | 9 | 9 | 25 | 10 | 61 | 97 | 99 | 76 |
| K18899 multidrug efflux pump | 13 | 16 | 5 | 15 | 36 | 49 | 46 | 36 | 194 | 192 | 177 | 197 |
| K18901 membrane fusion protein, multidrug efflux system | 0 | 0 | 0 | 0 | 1 | 0 | 0 | 0 | 2 | 2 | 3 | 7 |
| K18902 multidrug efflux pump | 5 | 2 | 9 | 3 | 4 | 6 | 6 | 6 | 4 | 3 | 2 | 7 |
| K18903 outer membrane protein, multidrug efflux system | 1 | 0 | 0 | 0 | 0 | 0 | 0 | 2 | 0 | 0 | 0 | 0 |
| K18904 outer membrane protein, multidrug efflux system | 0 | 0 | 0 | 0 | 3 | 7 | 3 | 3 | 4 | 2 | 3 | 3 |
| K18924 paired small multidrug resistance pump | 20 | 18 | 19 | 25 | 8 | 5 | 2 | 3 | 18 | 19 | 20 | 23 |
| K18925 paired small multidrug resistance pump | 12 | 12 | 11 | 13 | 0 | 2 | 1 | 0 | 13 | 10 | 8 | 18 |
| K18926 MFS transporter, DHA2 family, lincomycin resistance protein | 6 | 4 | 4 | 9 | 0 | 5 | 4 | 8 | 14 | 10 | 12 | 10 |
| K18934 MFS transporter, DHA2 family, multidrug resistance protein | 0 | 0 | 0 | 1 | 0 | 0 | 0 | 0 | 0 | 0 | 0 | 0 |
| K18967 diguanylate cyclase [EC:2.7.7.65] | 3 | 3 | 3 | 2 | 5 | 3 | 3 | 3 | 3 | 7 | 3 | 4 |
| K18975 small multidrug resistance pump | 0 | 0 | 0 | 0 | 0 | 0 | 0 | 0 | 0 | 0 | 1 | 0 |
| K19058 mercuric ion transport protein | 76 | 68 | 74 | 68 | 93 | 132 | 126 | 105 | 0 | 0 | 2 | 0 |
| K19059 mercuric ion transport protein | 428 | 399 | 460 | 426 | 321 | 320 | 352 | 338 | 75 | 113 | 77 | 117 |
| K19231 fibronectin-binding autotransporter adhesin | 70 | 94 | 88 | 86 | 16 | 5 | 13 | 13 | 146 | 180 | 155 | 177 |
| K19416 modulator of FtsH protease | 1165 | 1066 | 1023 | 1037 | 596 | 650 | 603 | 642 | 692 | 714 | 610 | 686 |
| K19419 transmembrane protein EpsG | 0 | 0 | 0 | 1 | 3 | 0 | 9 | 1 | 0 | 0 | 1 | 1 |
| K19540 fructoselysine transporter | 285 | 251 | 274 | 290 | 409 | 525 | 487 | 449 | 4 | 5 | 8 | 10 |

| | | | | | | | | | | | | |
|---|---|---|---|---|---|---|---|---|---|---|---|---|
| K19564 choline transport protein | 0 | 1 | 0 | 0 | 0 | 0 | 0 | 0 | 0 | 0 | 0 | 0 |
| K19577 MFS transporter, DHA1 family, inner membrane transport protein | 1819 | 1927 | 1716 | 1769 | 1115 | 1031 | 988 | 1089 | 1335 | 1314 | 1284 | 1352 |
| K19578 MFS transporter, DHA1 family, multidrug resistance protein | 1 | 0 | 0 | 0 | 0 | 0 | 0 | 0 | 0 | 0 | 0 | 0 |
| K19585 multidrug efflux pump | 1737 | 1835 | 1736 | 1906 | 1176 | 1195 | 1225 | 1122 | 2360 | 2527 | 2457 | 2600 |
| K19586 membrane fusion protein, multidrug efflux system | 676 | 729 | 657 | 708 | 510 | 495 | 484 | 506 | 917 | 983 | 930 | 1019 |
| K19593 outer membrane protein, multidrug efflux system | 2324 | 2202 | 2142 | 2210 | 1140 | 1156 | 1117 | 1242 | 708 | 686 | 648 | 689 |
| K19594 gold/copper resistance efflux pump | 4810 | 4654 | 4384 | 4469 | 2411 | 2498 | 2313 | 2695 | 1578 | 1404 | 1378 | 1433 |
| K19595 membrane fusion protein, gold/copper resistance efflux system | 1475 | 1431 | 1351 | 1420 | 736 | 783 | 769 | 892 | 465 | 454 | 389 | 465 |
| K20510 malonyl-S-ACP:biotin-protein carboxyltransferase subunit MadC [EC:2.1.3.10] | 154 | 131 | 147 | 148 | 0 | 0 | 0 | 0 | 63 | 47 | 43 | 52 |
| K20511 malonyl-S-ACP:biotin-protein carboxyltransferase subunit MadD [EC:2.1.3.10] | 114 | 149 | 125 | 121 | 0 | 0 | 0 | 0 | 50 | 73 | 45 | 63 |
| K20541 cellulose synthase operon protein B | 3483 | 3388 | 3153 | 3575 | 2859 | 3129 | 2934 | 2973 | 3830 | 3870 | 3600 | 3979 |
| K20543 cellulose synthase operon protein C | 5956 | 5625 | 5580 | 5802 | 4591 | 4988 | 4641 | 4464 | 6641 | 6664 | 6308 | 6730 |
| K20989 urea-proton symporter | 0 | 0 | 0 | 0 | 1 | 0 | 0 | 0 | 0 | 0 | 0 | 0 |
| K21133 multidrug efflux pump | 4551 | 4252 | 4024 | 4131 | 2364 | 2151 | 2238 | 2248 | 1529 | 1381 | 1231 | 1287 |
| K21134 multidrug efflux pump | 5407 | 5214 | 4966 | 5043 | 2961 | 2924 | 2791 | 3106 | 1781 | 1635 | 1702 | 1726 |
| K21135 membrane fusion protein, multidrug efflux system | 1551 | 1572 | 1311 | 1445 | 816 | 777 | 682 | 811 | 458 | 443 | 414 | 401 |
| K21136 membrane fusion protein, multidrug efflux system | 2139 | 2025 | 2078 | 2036 | 1268 | 1158 | 1042 | 1212 | 701 | 628 | 591 | 689 |
| K21137 membrane fusion protein, multidrug efflux system | 1166 | 1275 | 1124 | 1192 | 640 | 615 | 518 | 628 | 404 | 367 | 368 | 363 |
| K21393 TRAP-type transport system large permease protein | 48 | 40 | 42 | 39 | 22 | 33 | 33 | 26 | 90 | 148 | 101 | 131 |
| K21394 TRAP-type transport system small permease protein | 42 | 37 | 45 | 44 | 5 | 8 | 22 | 5 | 71 | 49 | 79 | 75 |
| K21395 TRAP-type transport system periplasmic protein | 246 | 231 | 234 | 243 | 73 | 61 | 66 | 75 | 336 | 331 | 294 | 336 |
| K21449 trimeric autotransporter adhesin | 907 | 972 | 1031 | 956 | 528 | 548 | 587 | 565 | 725 | 732 | 663 | 705 |
| K21572 starch-binding outer membrane protein, SusD/RagB family | 80802 | 88133 | 95078 | 87060 | 284791 | 283905 | 291892 | 281873 | 205554 | 195781 | 207048 | 194994 |
| K21573 TonB-dependent starch-binding outer membrane protein SusC | 2916 | 3156 | 3624 | 3262 | 9615 | 9950 | 9757 | 9546 | 6681 | 6419 | 6768 | 6321 |
| K21864 voltage-dependent calcium channel | 0 | 0 | 0 | 0 | 0 | 0 | 0 | 0 | 0 | 0 | 2 | 0 |
| K21989 calcium permeable stress-gated cation channel | 1 | 2 | 0 | 0 | 0 | 0 | 0 | 0 | 0 | 0 | 0 | 0 |
| K21993 formate transporter | 142 | 132 | 141 | 134 | 55 | 33 | 61 | 43 | 100 | 169 | 97 | 145 |
| K22044 moderate conductance mechanosensitive channel | 3033 | 2840 | 2838 | 2889 | 1720 | 1925 | 1662 | 1825 | 2260 | 2308 | 2200 | 2408 |
| K22051 miniconductance mechanosensitive channel | 2190 | 2053 | 2020 | 2133 | 1534 | 1606 | 1534 | 1483 | 2747 | 2896 | 2845 | 2999 |
| K22110 oligogalacturonate-specific porin family protein | 572 | 591 | 551 | 600 | 543 | 523 | 520 | 549 | 608 | 598 | 626 | 602 |
| K22112 putative amide transporter protein | 724 | 735 | 771 | 671 | 396 | 395 | 349 | 393 | 218 | 222 | 185 | 238 |
| K22736 vacuolar iron transporter family protein | 1234 | 1274 | 1111 | 1197 | 842 | 731 | 772 | 874 | 435 | 333 | 309 | 402 |
| K22737 erythrin-vacuolar iron transport family protein | 0 | 0 | 2 | 0 | 2 | 0 | 0 | 0 | 2 | 2 | 3 | 1 |
| K23016 MFS transporter, ACS family, L-galactonate transporter | 307 | 336 | 311 | 324 | 383 | 480 | 468 | 401 | 0 | 2 | 3 | 2 |
| K23025 H+-translocating diphosphatase [EC:7.1.3.1] | 0 | 0 | 0 | 0 | 0 | 0 | 1 | 0 | 0 | 0 | 1 | 1 |
| K23241 MFS transporter, DHA1 family, multidrug resistance protein | 1213 | 1227 | 1111 | 1224 | 647 | 632 | 646 | 631 | 1739 | 1821 | 1798 | 1996 |
| K23242 manganese efflux pump family protein | 491 | 449 | 446 | 461 | 381 | 470 | 408 | 453 | 532 | 500 | 452 | 524 |
| K23329 ATP-binding cassette, subfamily B, vacuolar membrane transporter HMT1/ACLQ | 1 | 0 | 0 | 1 | 0 | 0 | 0 | 1 | 0 | 0 | 1 | 0 |
| K23360 methylmalonyl-CoA decarboxylase subunit epsilon | 1 | 0 | 2 | 1 | 0 | 0 | 0 | 0 | 0 | 0 | 2 | 0 |
| K23508 galactofuranose transport system substrate-binding protein | 819 | 758 | 685 | 770 | 599 | 611 | 543 | 522 | 849 | 840 | 908 | 975 |
| K23509 galactofuranose transport system permease protein | 1460 | 1424 | 1348 | 1472 | 1070 | 1162 | 1051 | 1059 | 1787 | 1782 | 1862 | 1960 |
| K23511 riboflavin transport system permease protein | 0 | 1 | 1 | 1 | 0 | 0 | 0 | 0 | 0 | 0 | 0 | 1 |
| ko02044 Secretion system | 441359 | 438686 | 422560 | 433004 | 275189 | 275811 | 264933 | 278999 | 294480 | 296581 | 285975 | 295358 |
| K02221 YggT family protein | 1270 | 1255 | 1331 | 1325 | 776 | 822 | 770 | 673 | 895 | 846 | 840 | 831 |
| K02236 leader peptidase (prepilin peptidase) / N-methyltransferase [EC:3.4.23.43 2.1.1.-] | 45 | 35 | 65 | 49 | 14 | 10 | 11 | 17 | 36 | 54 | 69 | 44 |
| K02237 competence protein ComEA | 862 | 857 | 907 | 877 | 470 | 479 | 425 | 548 | 606 | 706 | 740 | 590 |
| K02238 competence protein ComEC | 5582 | 5679 | 5610 | 5771 | 5749 | 5730 | 5808 | 5924 | 5824 | 5868 | 5581 | 5861 |
| K02240 competence protein ComFA | 61 | 58 | 66 | 65 | 14 | 19 | 30 | 16 | 55 | 106 | 73 | 101 |
| K02241 competence protein ComFB | 78 | 72 | 52 | 59 | 6 | 12 | 7 | 3 | 66 | 76 | 62 | 87 |
| K02242 competence protein ComFC | 26 | 40 | 40 | 30 | 7 | 10 | 12 | 8 | 36 | 43 | 36 | 36 |
| K02243 competence protein ComGA | 48 | 67 | 48 | 56 | 20 | 16 | 17 | 28 | 59 | 106 | 56 | 72 |
| K02244 competence protein ComGB | 42 | 53 | 49 | 45 | 14 | 18 | 11 | 12 | 67 | 75 | 75 | 62 |
| K02245 competence protein ComGC | 8 | 13 | 15 | 11 | 5 | 5 | 4 | 3 | 14 | 19 | 13 | 17 |
| K02246 competence protein ComGD | 19 | 30 | 16 | 19 | 7 | 8 | 7 | 6 | 16 | 16 | 32 | 31 |
| K02248 competence protein ComGF | 25 | 19 | 23 | 33 | 2 | 7 | 12 | 5 | 22 | 24 | 22 | 26 |
| K02279 pilus assembly protein CpaB | 1644 | 1726 | 1559 | 1580 | 883 | 899 | 866 | 931 | 508 | 491 | 546 | 523 |
| K02280 pilus assembly protein CpaC | 1963 | 2012 | 1871 | 2014 | 1151 | 1172 | 1191 | 1185 | 703 | 682 | 657 | 694 |
| K02281 pilus assembly protein CpaD | 9 | 12 | 4 | 9 | 17 | 16 | 6 | 23 | 19 | 27 | 29 | 28 |
| K02282 pilus assembly protein CpaE | 1922 | 1979 | 1836 | 1854 | 1066 | 1064 | 1063 | 1120 | 650 | 622 | 659 | 660 |
| K02283 pilus assembly protein CpaF [EC:7.4.2.8] | 2441 | 2351 | 2445 | 2288 | 1303 | 1331 | 1301 | 1374 | 1062 | 1009 | 1058 | 1100 |
| K02450 general secretion pathway protein A | 6 | 9 | 4 | 6 | 5 | 4 | 2 | 5 | 100 | 94 | 87 | 102 |
| K02451 general secretion pathway protein B | 211 | 187 | 154 | 186 | 45 | 37 | 56 | 58 | 404 | 450 | 412 | 404 |
| K02504 protein transport protein HofB | 747 | 643 | 688 | 817 | 515 | 541 | 480 | 490 | 1004 | 975 | 1050 | 944 |
| K02505 protein transport protein HofC | 839 | 758 | 674 | 756 | 534 | 524 | 565 | 559 | 998 | 993 | 1055 | 1121 |
| K02507 protein transport protein HofQ | 744 | 760 | 809 | 834 | 551 | 659 | 568 | 497 | 1068 | 1183 | 1058 | 1170 |
| K02652 type IV pilus assembly protein PilB | 3107 | 3329 | 3181 | 3372 | 1677 | 1692 | 1623 | 1628 | 1576 | 1719 | 1750 | 1548 |
| K02653 type IV pilus assembly protein PilC | 2609 | 2592 | 2709 | 2666 | 1346 | 1407 | 1221 | 1262 | 1426 | 1633 | 1494 | 1578 |
| K02655 type IV pilus assembly protein PilE | 664 | 723 | 689 | 706 | 358 | 343 | 371 | 380 | 225 | 255 | 222 | 211 |
| K02656 type IV pilus assembly protein PilF | 704 | 709 | 688 | 672 | 393 | 406 | 389 | 406 | 232 | 226 | 244 | 200 |
| K02662 type IV pilus assembly protein PilM | 80 | 72 | 96 | 65 | 47 | 48 | 49 | 37 | 84 | 101 | 90 | 93 |
| K02663 type IV pilus assembly protein PilN | 705 | 681 | 673 | 715 | 383 | 389 | 410 | 398 | 246 | 234 | 271 | 257 |
| K02664 type IV pilus assembly protein PilO | 1119 | 1078 | 1086 | 1136 | 565 | 620 | 536 | 600 | 366 | 380 | 399 | 399 |
| K02665 type IV pilus assembly protein PilP | 838 | 821 | 801 | 822 | 503 | 481 | 466 | 444 | 268 | 244 | 273 | 263 |
| K02666 type IV pilus assembly protein PilQ | 3243 | 3343 | 3424 | 3316 | 1798 | 1941 | 1768 | 1912 | 1191 | 1195 | 1087 | 1144 |
| K02669 twitching motility protein PilT | 4499 | 4540 | 4495 | 4507 | 2850 | 2944 | 2850 | 2954 | 2897 | 2960 | 2669 | 2906 |
| K02670 twitching motility protein PilU | 2022 | 2053 | 2007 | 1953 | 1153 | 1164 | 1074 | 1120 | 629 | 661 | 597 | 654 |
| K02671 type IV pilus assembly protein PilV | 804 | 812 | 878 | 844 | 469 | 513 | 482 | 495 | 257 | 305 | 286 | 280 |
| K02672 type IV pilus assembly protein PilW | 1246 | 1279 | 1272 | 1317 | 805 | 804 | 704 | 814 | 427 | 389 | 423 | 398 |
| K02673 type IV pilus assembly protein PilX | 869 | 893 | 962 | 919 | 544 | 588 | 511 | 534 | 274 | 295 | 296 | 262 |
| K02674 type IV pilus assembly protein PilY1 | 5374 | 5554 | 5888 | 5865 | 3216 | 3352 | 3295 | 3347 | 1958 | 1851 | 1810 | 1761 |
| K02676 type IV pilus assembly protein PilZ | 457 | 442 | 394 | 459 | 250 | 262 | 239 | 255 | 142 | 118 | 148 | 146 |
| K02679 prepilin peptidase dependent protein A | 415 | 336 | 373 | 397 | 236 | 222 | 231 | 239 | 528 | 516 | 542 | 576 |
| K02680 prepilin peptidase dependent protein B | 303 | 296 | 277 | 309 | 206 | 205 | 205 | 202 | 443 | 396 | 431 | 460 |
| K02681 prepilin peptidase dependent protein C | 180 | 185 | 157 | 172 | 128 | 125 | 136 | 118 | 278 | 242 | 251 | 271 |
| K02682 prepilin peptidase dependent protein D | 238 | 219 | 214 | 235 | 185 | 192 | 145 | 167 | 277 | 329 | 298 | 304 |
| K03220 type III secretion protein D | 1960 | 2043 | 1946 | 1958 | 1006 | 1043 | 1019 | 1063 | 559 | 554 | 595 | 527 |
| K04049 type III secretion protein B | 480 | 482 | 417 | 417 | 246 | 220 | 236 | 304 | 141 | 125 | 105 | 141 |
| K04050 type III secretion protein E | 269 | 240 | 238 | 213 | 137 | 122 | 127 | 120 | 77 | 67 | 69 | 77 |
| K04051 type III secretion protein G | 530 | 498 | 511 | 548 | 292 | 278 | 239 | 291 | 184 | 163 | 139 | 163 |
| K04052 type III secretion protein H | 634 | 573 | 549 | 568 | 298 | 340 | 314 | 277 | 214 | 153 | 158 | 181 |
| K04053 type III secretion protein I | 480 | 427 | 474 | 454 | 257 | 226 | 253 | 274 | 155 | 156 | 126 | 133 |
| K04054 type III secretion protein K | 833 | 731 | 695 | 725 | 458 | 385 | 379 | 455 | 228 | 203 | 229 | 233 |
| K04060 type III secretion protein Y | 366 | 442 | 382 | 433 | 213 | 218 | 225 | 236 | 97 | 113 | 108 | 103 |
| K04061 flagellar biosynthesis protein | 668 | 709 | 647 | 675 | 307 | 291 | 308 | 347 | 340 | 340 | 304 | 361 |
| K04337 curli production assembly/transport component CsgE | 72 | 91 | 65 | 77 | 99 | 113 | 103 | 112 | 19 | 20 | 13 | 31 |
| K04338 curli production assembly/transport component CsgF | 85 | 65 | 67 | 95 | 124 | 120 | 130 | 114 | 21 | 20 | 21 | 12 |
| K06214 curli production assembly/transport component CsgG | 145 | 143 | 156 | 156 | 256 | 256 | 279 | 229 | 34 | 58 | 48 | 48 |
| K07332 archaeal flagellar protein FlaI | 0 | 0 | 2 | 0 | 0 | 3 | 0 | 0 | 13 | 6 | 4 | 15 |
| K07333 archaeal flagellar protein FlaJ | 0 | 2 | 2 | 0 | 2 | 4 | 0 | 4 | 2 | 13 | 5 | 18 |
| K07348 minor fimbrial subunit | 351 | 376 | 313 | 359 | 229 | 241 | 253 | 208 | 537 | 552 | 521 | 514 |
| K07349 minor fimbrial subunit | 209 | 216 | 205 | 217 | 60 | 60 | 66 | 43 | 451 | 497 | 446 | 452 |
| K07350 minor fimbrial subunit | 594 | 595 | 575 | 622 | 475 | 575 | 463 | 427 | 815 | 808 | 783 | 810 |
| K08084 type IV fimbrial biogenesis protein FimT | 749 | 724 | 766 | 811 | 435 | 453 | 407 | 421 | 211 | 267 | 241 | 244 |
| K08085 type IV fimbrial biogenesis protein FimU | 1037 | 1111 | 1084 | 1065 | 546 | 557 | 547 | 557 | 338 | 326 | 285 | 328 |
| K11894 type VI secretion system protein ImpI | 1788 | 1779 | 1793 | 1822 | 1072 | 979 | 1050 | 1038 | 521 | 517 | 508 | 469 |
| K11896 type VI secretion system protein ImpG | 7259 | 6984 | 6620 | 6908 | 4112 | 4121 | 4073 | 4382 | 4259 | 3990 | 3906 | 4111 |
| K11897 type VI secretion system protein ImpF | 1013 | 990 | 920 | 877 | 612 | 609 | 652 | 696 | 283 | 246 | 257 | 270 |
| K11898 type VI secretion system protein ImpE | 1214 | 1170 | 1168 | 1164 | 820 | 783 | 767 | 810 | 330 | 290 | 296 | 345 |
| K11905 type VI secretion system protein | 884 | 939 | 876 | 899 | 492 | 511 | 549 | 536 | 632 | 568 | 569 | 581 |
| K11909 type VI secretion system protein VasI | 57 | 50 | 53 | 73 | 65 | 83 | 90 | 71 | 4 | 4 | 3 | 1 |
| K11910 type VI secretion system protein VasJ | 3055 | 3169 | 3021 | 2997 | 1900 | 1825 | 1796 | 1820 | 1686 | 1766 | 1787 | 1765 |
| K11911 type VI secretion system protein VasL | 832 | 864 | 771 | 887 | 593 | 589 | 595 | 580 | 1001 | 1023 | 1088 | 1022 |
| K11916 serine/threonine-protein kinase Stk1 [EC:2.7.11.-] | 1429 | 1315 | 1285 | 1352 | 660 | 724 | 653 | 727 | 430 | 418 | 450 | 378 |
| K11918 type VI secretion system protein | 575 | 636 | 604 | 576 | 309 | 301 | 315 | 360 | 195 | 161 | 172 | 193 |
| K11919 type VI secretion system lysozyme-related protein | 618 | 575 | 526 | 529 | 279 | 297 | 288 | 357 | 193 | 160 | 172 | 182 |
| K12055 chromosome partitioning related protein ParA | 1450 | 1464 | 1542 | 1390 | 1276 | 1288 | 1293 | 1221 | 589 | 555 | 576 | 531 |
| K12056 conjugal transfer mating pair stabilization protein TraG | 1460 | 1455 | 1528 | 1536 | 543 | 590 | 616 | 525 | 8713 | 9573 | 8826 | 8891 |
| K12057 conjugal transfer pilus assembly protein TraF | 386 | 429 | 459 | 472 | 141 | 176 | 184 | 149 | 2633 | 2918 | 2545 | 2720 |
| K12058 conjugal transfer mating pair stabilization protein TraN | 603 | 660 | 710 | 685 | 267 | 254 | 281 | 252 | 4068 | 4484 | 3969 | 4072 |
| K12059 conjugal transfer pilus assembly protein TrbC | 174 | 206 | 223 | 226 | 46 | 86 | 90 | 73 | 1236 | 1342 | 1316 | 1423 |
| K12060 conjugal transfer pilus assembly protein TraU | 532 | 533 | 484 | 536 | 192 | 217 | 211 | 188 | 3139 | 3322 | 3187 | 3294 |

| | | | | | | | | | | | | |
|---|---|---|---|---|---|---|---|---|---|---|---|---|
| K12061 conjugal transfer pilus assembly protein TraW | 344 | 329 | 387 | 379 | 139 | 169 | 151 | 110 | 2208 | 2538 | 2168 | 2183 |
| K12062 conjugal transfer pilin signal peptidase TrbI | 230 | 192 | 231 | 229 | 82 | 76 | 81 | 83 | 1263 | 1423 | 1173 | 1319 |
| K12063 conjugal transfer ATP-binding protein TraC | 1448 | 1289 | 1445 | 1470 | 629 | 563 | 541 | 526 | 8524 | 9390 | 8573 | 9016 |
| K12064 conjugal transfer pilus assembly protein TraV | 80 | 89 | 111 | 106 | 50 | 44 | 37 | 30 | 479 | 547 | 529 | 584 |
| K12065 conjugal transfer pilus assembly protein TraB | 658 | 631 | 660 | 682 | 232 | 297 | 252 | 235 | 4093 | 4190 | 3884 | 4169 |
| K12066 conjugal transfer pilus assembly protein TraK | 245 | 267 | 269 | 289 | 98 | 120 | 110 | 114 | 1486 | 1606 | 1461 | 1680 |
| K12067 conjugal transfer pilus assembly protein TraE | 347 | 302 | 294 | 294 | 108 | 144 | 100 | 115 | 1776 | 1856 | 1764 | 1735 |
| K12068 conjugal transfer pilus assembly protein TraL | 187 | 153 | 161 | 169 | 50 | 62 | 72 | 64 | 862 | 985 | 961 | 957 |
| K12069 conjugal transfer pilus assembly protein TraA | 2 | 1 | 2 | 3 | 0 | 0 | 0 | 0 | 0 | 0 | 1 | 0 |
| K12070 conjugal transfer pilus assembly protein TraI | 0 | 2 | 0 | 0 | 0 | 0 | 0 | 0 | 0 | 0 | 2 | 1 |
| K12071 conjugal transfer pilus assembly protein TraD | 31 | 30 | 12 | 21 | 42 | 54 | 54 | 54 | 48 | 37 | 44 | 38 |
| K12072 conjugative transfer pilus assembly protein TraH | 829 | 766 | 803 | 838 | 369 | 393 | 383 | 365 | 4616 | 4805 | 4626 | 4823 |
| K12203 defect in organelle trafficking protein DotB [EC:7.2.4.8] | 0 | 1 | 0 | 1 | 0 | 0 | 0 | 0 | 2 | 0 | 1 | 1 |
| K12204 defect in organelle trafficking protein DotC | 0 | 0 | 0 | 0 | 0 | 0 | 0 | 0 | 1 | 0 | 0 | 1 |
| K12209 intracellular multiplication protein IcmE | 0 | 0 | 0 | 0 | 0 | 1 | 0 | 0 | 0 | 0 | 0 | 0 |
| K12213 intracellular multiplication protein IcmK | 0 | 0 | 0 | 0 | 0 | 2 | 0 | 0 | 0 | 0 | 0 | 0 |
| K12217 intracellular multiplication protein IcmO [EC:7.2.4.8] | 4 | 3 | 9 | 4 | 3 | 1 | 2 | 2 | 17 | 10 | 10 | 20 |
| K12218 intracellular multiplication protein IcmP | 0 | 0 | 0 | 0 | 0 | 0 | 0 | 0 | 0 | 1 | 0 | 0 |
| K12268 accessory secretory protein Asp1 | 0 | 0 | 1 | 0 | 0 | 2 | 0 | 0 | 0 | 0 | 1 | 0 |
| K12269 accessory secretory protein Asp2 | 0 | 1 | 0 | 1 | 3 | 2 | 4 | 1 | 2 | 0 | 1 | 0 |
| K12270 accessory secretory protein Asp3 | 0 | 0 | 0 | 0 | 0 | 0 | 0 | 1 | 0 | 0 | 1 | 0 |
| K12278 MSHA biogenesis protein MshG | 0 | 1 | 0 | 1 | 0 | 0 | 0 | 0 | 0 | 2 | 0 | 1 |
| K12288 pilus assembly protein HofM | 482 | 560 | 490 | 540 | 379 | 387 | 387 | 409 | 700 | 770 | 626 | 692 |
| K12289 pilus assembly protein HofN | 399 | 326 | 339 | 363 | 290 | 297 | 236 | 237 | 465 | 460 | 458 | 485 |
| K12290 pilus assembly protein HofO | 267 | 252 | 213 | 263 | 141 | 200 | 188 | 179 | 318 | 326 | 309 | 385 |
| K12291 pilus assembly protein HofP | 183 | 234 | 186 | 217 | 131 | 146 | 138 | 169 | 332 | 326 | 292 | 291 |
| K12510 tight adherence protein B | 1546 | 1547 | 1517 | 1612 | 747 | 739 | 741 | 763 | 702 | 601 | 717 | 752 |
| K12511 tight adherence protein C | 1387 | 1424 | 1258 | 1294 | 750 | 740 | 741 | 842 | 481 | 401 | 449 | 408 |
| K12517 major pilin subunit PapA | 0 | 1 | 0 | 1 | 2 | 0 | 0 | 3 | 0 | 4 | 1 | 1 |
| K12518 outer membrane usher protein PapC | 7 | 8 | 9 | 11 | 41 | 43 | 40 | 44 | 127 | 120 | 115 | 170 |
| K12519 chaperone protein PapD | 2 | 4 | 6 | 4 | 13 | 10 | 11 | 6 | 39 | 35 | 40 | 44 |
| K12520 minor pilin subunit PapE | 0 | 0 | 0 | 0 | 0 | 0 | 0 | 0 | 1 | 2 | 1 | 0 |
| K12521 minor pilin subunit PapF | 0 | 0 | 1 | 0 | 1 | 0 | 0 | 0 | 1 | 0 | 0 | 0 |
| K12522 adhesin PapG | 0 | 0 | 2 | 1 | 1 | 1 | 1 | 0 | 4 | 3 | 2 | 0 |
| K12523 minor pilin subunit PapK | 2 | 0 | 0 | 0 | 1 | 0 | 3 | 0 | 2 | 0 | 1 | 1 |
| K21964 Mat/Ecp fimbriae major subunit | 307 | 285 | 281 | 296 | 74 | 55 | 81 | 67 | 482 | 459 | 475 | 488 |
| K21965 Mat/Ecp fimbriae periplasmic chaperone | 282 | 290 | 274 | 322 | 79 | 79 | 77 | 76 | 530 | 493 | 552 | 604 |
| K21966 Mat/Ecp fimbriae outer membrane usher protein | 1241 | 1337 | 1164 | 1303 | 288 | 292 | 308 | 273 | 2025 | 2231 | 2137 | 2207 |
| K21967 Mat/Ecp fimbriae adhesin | 818 | 797 | 726 | 816 | 180 | 216 | 184 | 210 | 1462 | 1379 | 1346 | 1484 |
| K21968 Mat/Ecp fimbriae periplasmic chaperone | 356 | 362 | 305 | 389 | 78 | 93 | 98 | 97 | 596 | 584 | 540 | 614 |
| K22487 type III secretion system protein | 0 | 3 | 2 | 0 | 3 | 0 | 0 | 0 | 0 | 0 | 2 | 0 |
| K22488 type III secretion system protein | 0 | 0 | 0 | 0 | 16 | 17 | 1 | 9 | 5 | 0 | 2 | 0 |
| K22504 type III secretion system outer membrane ring protein | 0 | 0 | 0 | 0 | 0 | 0 | 0 | 0 | 0 | 0 | 2 | 0 |
| K22505 type III secretion system inner membrane ring protein | 0 | 0 | 0 | 0 | 3 | 0 | 2 | 4 | 0 | 0 | 0 | 1 |
| K22506 type III secretion system ATPase [EC:7.4.2.8] | 0 | 0 | 1 | 0 | 0 | 0 | 0 | 0 | 0 | 0 | 0 | 0 |
| K22507 type III secretion system export apparatus protein | 1 | 0 | 0 | 0 | 5 | 6 | 0 | 8 | 0 | 0 | 0 | 0 |
| K22508 type III secretion system export apparatus protein | 0 | 0 | 0 | 0 | 0 | 3 | 5 | 4 | 0 | 2 | 1 | 0 |
| K22509 type III secretion system export apparatus protein | 0 | 0 | 0 | 0 | 7 | 5 | 3 | 9 | 0 | 0 | 2 | 1 |
| K22510 type III secretion system export apparatus switch protein | 0 | 0 | 0 | 0 | 2 | 6 | 7 | 3 | 1 | 0 | 0 | 2 |
| K22514 type III secretion system protein | 0 | 0 | 0 | 0 | 1 | 0 | 0 | 0 | 0 | 0 | 1 | 0 |
| ko02042 Bacterial toxins | 23408 | 23457 | 23290 | 23428 | 22753 | 22816 | 23118 | 23209 | 16945 | 16365 | 16465 | 16328 |
| K03699 putative hemolysin | 1844 | 1908 | 1958 | 1829 | 2836 | 2775 | 2919 | 2865 | 2850 | 2870 | 2960 | 2878 |
| K11008 cytotoxic necrotizing factor 1 | 8 | 2 | 5 | 7 | 11 | 3 | 7 | 18 | 142 | 200 | 165 | 172 |
| K11009 murine toxin | 68 | 62 | 60 | 68 | 84 | 115 | 110 | 106 | 0 | 0 | 0 | 0 |
| K11020 exotoxin A [EC:2.4.2.-] | 449 | 411 | 367 | 392 | 219 | 191 | 215 | 234 | 125 | 143 | 117 | 120 |
| K11021 insecticidal toxin complex protein TccC | 0 | 0 | 0 | 1 | 0 | 0 | 0 | 0 | 0 | 0 | 0 | 1 |
| K11045 cAMP factor | 0 | 1 | 0 | 0 | 1 | 0 | 0 | 0 | 0 | 0 | 0 | 0 |
| K11068 hemolysin III | 1605 | 1595 | 1702 | 1604 | 2012 | 2058 | 2089 | 1950 | 2044 | 1928 | 2080 | 1953 |
| K11139 hemolysin E | 3 | 0 | 0 | 3 | 14 | 6 | 3 | 6 | 12 | 15 | 16 | 17 |
| K16636 exoenzyme T | 1645 | 1636 | 1654 | 1585 | 931 | 971 | 883 | 906 | 611 | 563 | 504 | 547 |
| K16637 adenylate cyclase ExoY | 883 | 822 | 758 | 784 | 407 | 406 | 379 | 433 | 219 | 241 | 210 | 228 |
| K16638 exoenzyme U | 2600 | 2537 | 2781 | 2767 | 1489 | 1465 | 1541 | 1463 | 894 | 846 | 823 | 783 |
| ko02022 Two-component system | 238634 | 234562 | 229833 | 235177 | 148390 | 149747 | 146323 | 150491 | 171306 | 173655 | 169638 | 173985 |
| K02475 two-component system, CitB family, response regulator | 28 | 28 | 22 | 31 | 3 | 5 | 7 | 8 | 39 | 32 | 22 | 28 |
| K02477 two-component system, LytTR family, response regulator | 1946 | 1959 | 2124 | 2112 | 3918 | 3979 | 3964 | 3940 | 4079 | 3921 | 4098 | 3833 |
| K02479 two-component system, NarL family, response regulator | 1 | 0 | 0 | 0 | 0 | 0 | 0 | 0 | 0 | 0 | 0 | 0 |
| K02481 two-component system, NtrC family, response regulator | 1696 | 1764 | 1813 | 1731 | 5340 | 5384 | 5591 | 5339 | 3940 | 3617 | 3923 | 3697 |
| K02483 two-component system, OmpR family, response regulator | 417 | 424 | 431 | 435 | 212 | 230 | 235 | 227 | 964 | 1038 | 908 | 931 |
| K02485 two-component system, response regulator | 526 | 519 | 580 | 550 | 424 | 451 | 380 | 410 | 772 | 860 | 767 | 832 |
| K07183 two-component system, response regulator / RNA-binding antiterminator | 567 | 590 | 501 | 586 | 306 | 305 | 294 | 262 | 190 | 167 | 156 | 135 |
| K07814 putative two-component system response regulator | 2584 | 2576 | 2470 | 2532 | 1460 | 1332 | 1260 | 1385 | 1243 | 1283 | 1250 | 1303 |
| K21603 two-component system, OmpR family, flagellar system response regulator FtcR | 0 | 2 | 1 | 0 | 0 | 0 | 0 | 3 | 0 | 0 | 3 | 0 |
| K22010 two-component system, response regulator PdtaR | 375 | 311 | 358 | 338 | 87 | 118 | 88 | 85 | 281 | 334 | 316 | 332 |
| K22067 two-component system, oxyanion-binding sensor | 1789 | 1729 | 1805 | 1741 | 921 | 952 | 885 | 1069 | 575 | 459 | 509 | 470 |
| ko02035 Bacterial motility proteins | 232520 | 230031 | 224589 | 229169 | 138908 | 141101 | 132665 | 138650 | 104936 | 105096 | 102415 | 105206 |
| K02385 flagellar protein FlbD | 115 | 100 | 128 | 101 | 29 | 24 | 31 | 26 | 76 | 84 | 77 | 90 |
| K02395 flagellar protein FlgJ | 2085 | 2045 | 2044 | 2046 | 1262 | 1285 | 1280 | 1276 | 695 | 616 | 636 | 608 |
| K02404 flagellar biosynthesis protein FlhF | 1539 | 1606 | 1463 | 1459 | 811 | 808 | 687 | 809 | 511 | 418 | 442 | 452 |
| K02415 flagellar FliL protein | 995 | 949 | 995 | 905 | 620 | 617 | 769 | 754 | 295 | 333 | 323 | 332 |
| K02675 type IV pilus assembly protein PilY2 | 429 | 515 | 555 | 548 | 311 | 306 | 293 | 286 | 157 | 171 | 165 | 173 |
| K03516 flagellar protein FlhE | 81 | 81 | 74 | 93 | 145 | 105 | 113 | 121 | 20 | 22 | 25 | 17 |
| K04562 flagellar biosynthesis protein FlhG | 1170 | 1152 | 1156 | 1169 | 628 | 668 | 604 | 655 | 386 | 325 | 364 | 406 |
| K06595 heam-based aerotactic trancducer | 2 | 0 | 6 | 3 | 0 | 0 | 0 | 0 | 0 | 1 | 0 | 0 |
| K06601 flagellar protein FlbT | 0 | 0 | 1 | 0 | 0 | 0 | 0 | 0 | 0 | 0 | 2 | 1 |
| K06602 flagellar protein FlaF | 0 | 0 | 0 | 1 | 0 | 0 | 0 | 0 | 0 | 0 | 0 | 0 |
| K06603 flagellar protein FlaG | 524 | 565 | 530 | 558 | 286 | 299 | 316 | 312 | 176 | 154 | 179 | 165 |
| K07351 fimbrial protein | 242 | 236 | 240 | 264 | 103 | 91 | 98 | 50 | 549 | 588 | 532 | 606 |
| K07352 type 1 fimbrial protein | 0 | 4 | 7 | 0 | 2 | 0 | 0 | 0 | 0 | 0 | 3 | 0 |
| K07353 fimbrial chaperone protein | 0 | 0 | 0 | 0 | 9 | 4 | 8 | 7 | 0 | 0 | 3 | 1 |
| K07354 outer membrane usher protein | 0 | 0 | 0 | 0 | 16 | 11 | 21 | 16 | 0 | 12 | 6 | 5 |
| K07355 fimbrial-like protein | 0 | 0 | 0 | 0 | 5 | 8 | 1 | 2 | 0 | 0 | 3 | 2 |
| K07356 fimbrial protein | 0 | 0 | 0 | 0 | 14 | 16 | 8 | 5 | 24 | 15 | 14 | 7 |
| K10564 chemotaxis protein MotC | 1 | 2 | 0 | 1 | 3 | 0 | 0 | 0 | 5 | 0 | 3 | 5 |
| K10565 chemotaxis protein MotD | 0 | 1 | 0 | 0 | 3 | 0 | 4 | 0 | 0 | 6 | 1 | 5 |
| K13626 flagellar assembly factor FliW | 288 | 251 | 253 | 279 | 68 | 75 | 94 | 89 | 271 | 317 | 280 | 288 |
| K18475 lysine-N-methylase [EC:2.1.1.-] | 169 | 167 | 198 | 191 | 76 | 89 | 104 | 88 | 193 | 234 | 219 | 223 |
| ko04812 Cytoskeleton proteins | 42353 | 42608 | 42643 | 42945 | 43096 | 42923 | 42383 | 42182 | 48325 | 49299 | 48277 | 48294 |
| K06990 MEMO1 family protein | 7 | 3 | 4 | 7 | 0 | 2 | 3 | 0 | 28 | 24 | 16 | 27 |
| K10401 kinesin family member 18/19 | 0 | 0 | 0 | 0 | 2 | 0 | 0 | 0 | 0 | 0 | 0 | 1 |
| K17292 tubulin-specific chaperone A | 0 | 0 | 0 | 0 | 0 | 0 | 0 | 0 | 1 | 0 | 0 | 0 |
| K18640 plasmid segregation protein ParM | 1591 | 1612 | 1576 | 1704 | 900 | 971 | 832 | 877 | 4905 | 5481 | 5150 | 5121 |
| K19755 radial spoke head protein 1 | 20 | 23 | 43 | 31 | 100 | 99 | 0 | 0 | 73 | 54 | 88 | 68 |
| ko04147 Exosome | 228110 | 227877 | 228494 | 227107 | 244287 | 245232 | 242597 | 244941 | 227812 | 225518 | 225574 | 224253 |
| K01482 dimethylargininase [EC:3.5.3.18] | 72 | 69 | 75 | 84 | 44 | 33 | 47 | 41 | 39 | 34 | 25 | 39 |
| K02503 histidine triad (HIT) family protein | 2078 | 2105 | 2006 | 2175 | 1888 | 1911 | 1912 | 1875 | 1717 | 1648 | 1665 | 1624 |
| K03098 apolipoprotein D and lipocalin family protein | 1350 | 1343 | 1257 | 1291 | 739 | 686 | 719 | 714 | 880 | 819 | 889 | 861 |
| K03522 electron transfer flavoprotein alpha subunit | 3999 | 4137 | 3983 | 3921 | 4340 | 4137 | 4150 | 4411 | 3167 | 2835 | 3044 | 3117 |
| K17086 transmembrane 9 superfamily member 2/4 | 0 | 0 | 0 | 0 | 0 | 0 | 0 | 0 | 0 | 0 | 0 | 1 |
| K17255 Rab GDP dissociation inhibitor | 0 | 1 | 0 | 0 | 0 | 0 | 0 | 0 | 0 | 0 | 0 | 0 |
| K18222 arylsulfatase E [EC:3.1.6.-] | 0 | 0 | 0 | 0 | 0 | 0 | 0 | 0 | 0 | 0 | 0 | 1 |
| K18258 thiomorpholine-carboxylate dehydrogenase [EC:1.5.1.25] | 0 | 0 | 0 | 1 | 0 | 1 | 0 | 2 | 0 | 0 | 1 | 1 |
| K20011 peroxiredoxin 3 [EC:1.11.1.15] | 1 | 0 | 0 | 0 | 0 | 0 | 0 | 0 | 0 | 0 | 0 | 0 |
| ko02048 Prokaryotic defense system | 80503 | 80809 | 82773 | 82175 | 83336 | 84742 | 84850 | 83760 | 85210 | 87396 | 85610 | 86848 |
| K00571 site-specific DNA-methyltransferase (adenine-specific) [EC:2.1.1.72] | 969 | 927 | 1063 | 970 | 1725 | 1765 | 1852 | 1666 | 2782 | 2965 | 2811 | 2818 |
| K01153 type I restriction enzyme, R subunit [EC:3.1.21.3] | 6070 | 6071 | 6312 | 6229 | 4517 | 4612 | 4472 | 4430 | 3152 | 3438 | 3379 | 3438 |
| K01154 type I restriction enzyme, S subunit [EC:3.1.21.3] | 3298 | 3282 | 3607 | 3402 | 4432 | 4464 | 4710 | 4430 | 2626 | 2662 | 2779 | 2731 |
| K01155 type II restriction enzyme [EC:3.1.21.4] | 132 | 142 | 166 | 143 | 167 | 174 | 183 | 169 | 21 | 29 | 21 | 22 |
| K01156 type III restriction enzyme [EC:3.1.21.5] | 130 | 140 | 160 | 173 | 24 | 37 | 28 | 37 | 292 | 405 | 358 | 334 |
| K03427 type I restriction enzyme M protein [EC:2.1.1.72] | 4622 | 4539 | 4928 | 4761 | 4681 | 4879 | 4800 | 4757 | 3089 | 2895 | 2892 | 3091 |
| K03970 phage shock protein B | 115 | 116 | 105 | 114 | 76 | 79 | 103 | 77 | 193 | 188 | 175 | 168 |
| K05839 haemolysin expression modulating protein | 305 | 289 | 319 | 347 | 168 | 194 | 161 | 163 | 934 | 1020 | 982 | 1052 |

| | | | | | | | | | | | | |
|---|---|---|---|---|---|---|---|---|---|---|---|---|
| K06218 mRNA interferase RelE/StbE | 131 | 143 | 134 | 106 | 78 | 104 | 65 | 90 | 24 | 25 | 33 | 38 |
| K07012 CRISPR-associated endonuclease/helicase Cas3 [EC:3.1.-.- 3.6.4.-] | 3000 | 3144 | 3248 | 3259 | 1391 | 1428 | 1465 | 1285 | 3308 | 3491 | 3414 | 3467 |
| K07016 CRISPR-associated protein Csm1 | 0 | 0 | 1 | 1 | 2 | 0 | 1 | 4 | 6 | 7 | 8 | 5 |
| K07060 endoribonuclease Nob1 [EC:3.1.-.-] | 0 | 0 | 2 | 3 | 0 | 0 | 0 | 0 | 2 | 4 | 2 | 0 |
| K07062 toxin FitB [EC:3.1.-.-] | 13 | 18 | 23 | 15 | 21 | 16 | 16 | 31 | 20 | 36 | 21 | 28 |
| K07171 mRNA interferase MazF [EC:3.1.-.-] | 665 | 570 | 576 | 614 | 303 | 261 | 225 | 254 | 608 | 801 | 666 | 700 |
| K07172 antitoxin MazE | 2 | 1 | 1 | 0 | 8 | 10 | 9 | 6 | 10 | 8 | 19 | 17 |
| K07316 adenine-specific DNA-methyltransferase [EC:2.1.1.72] | 85 | 68 | 86 | 81 | 53 | 60 | 50 | 35 | 218 | 300 | 237 | 218 |
| K07317 adenine-specific DNA-methyltransferase [EC:2.1.1.72] | 5 | 3 | 4 | 6 | 8 | 12 | 2 | 5 | 7 | 3 | 6 | 10 |
| K07318 adenine-specific DNA-methyltransferase [EC:2.1.1.72] | 365 | 392 | 406 | 399 | 69 | 90 | 56 | 64 | 164 | 251 | 243 | 262 |
| K07319 adenine-specific DNA-methyltransferase [EC:2.1.1.72] | 642 | 623 | 586 | 548 | 446 | 502 | 480 | 476 | 2052 | 2373 | 2144 | 2259 |
| K07334 toxin HigB-1 | 498 | 497 | 479 | 488 | 614 | 538 | 561 | 596 | 373 | 421 | 407 | 416 |
| K07339 mRNA interferase HicA [EC:3.1.-.-] | 4 | 7 | 5 | 5 | 8 | 5 | 10 | 4 | 9 | 28 | 20 | 20 |
| K07341 death on curing protein | 192 | 217 | 214 | 221 | 78 | 64 | 77 | 71 | 357 | 375 | 410 | 421 |
| K07448 restriction system protein | 416 | 449 | 465 | 435 | 286 | 325 | 285 | 281 | 198 | 248 | 213 | 219 |
| K07451 5-methylcytosine-specific restriction enzyme A [EC:3.1.21.-] | 592 | 575 | 617 | 648 | 969 | 1022 | 1004 | 915 | 332 | 364 | 342 | 331 |
| K07452 5-methylcytosine-specific restriction enzyme B [EC:3.1.21.-] | 976 | 882 | 931 | 880 | 1191 | 1248 | 1286 | 1183 | 666 | 679 | 671 | 713 |
| K07464 CRISPR-associated exonuclease Cas4 [EC:3.1.12.1] | 237 | 181 | 231 | 237 | 216 | 204 | 227 | 207 | 276 | 297 | 285 | 336 |
| K07473 DNA-damage-inducible protein J | 405 | 428 | 389 | 425 | 200 | 188 | 152 | 181 | 855 | 935 | 932 | 869 |
| K07746 antitoxin ParD1/3/4 | 563 | 526 | 596 | 570 | 524 | 551 | 557 | 608 | 202 | 197 | 216 | 192 |
| K09002 CRISPR-associated protein Csm3 | 1 | 2 | 3 | 1 | 2 | 0 | 2 | 5 | 10 | 3 | 7 | 14 |
| K09127 CRISPR-associated protein Cmr3 | 2 | 0 | 0 | 1 | 0 | 0 | 0 | 0 | 0 | 0 | 0 | 0 |
| K09159 antitoxin CptB | 614 | 572 | 533 | 551 | 336 | 362 | 346 | 355 | 384 | 365 | 326 | 367 |
| K09951 CRISPR-associated protein Cas2 | 239 | 234 | 265 | 259 | 170 | 164 | 174 | 175 | 351 | 378 | 387 | 437 |
| K09952 CRISPR-associated endonuclease Csn1 [EC:3.1.-.-] | 255 | 268 | 262 | 308 | 85 | 113 | 80 | 89 | 211 | 285 | 251 | 215 |
| K12148 biofilm regulator BssS | 220 | 168 | 174 | 199 | 154 | 184 | 163 | 189 | 208 | 180 | 180 | 219 |
| K12151 multiple stress resistance protein BhsA | 161 | 148 | 171 | 172 | 146 | 162 | 146 | 149 | 169 | 173 | 168 | 192 |
| K13650 MqsR-controlled colanic acid and biofilm protein A | 93 | 109 | 85 | 90 | 25 | 31 | 34 | 36 | 215 | 214 | 241 | 211 |
| K13651 motility quorum-sensing regulator / GCU-specific mRNA interferase toxin | 2 | 4 | 2 | 1 | 0 | 0 | 2 | 0 | 0 | 0 | 0 | 1 |
| K16214 UDP-N-acetylglucosamine kinase [EC:2.7.1.176] | 47 | 34 | 49 | 29 | 24 | 32 | 24 | 23 | 30 | 29 | 30 | 28 |
| K18829 antitoxin VapB | 471 | 559 | 551 | 583 | 362 | 446 | 398 | 403 | 159 | 169 | 140 | 144 |
| K18837 cytoskeleton-binding toxin CbtA and related proteins | 8 | 9 | 12 | 7 | 8 | 8 | 3 | 12 | 89 | 81 | 87 | 101 |
| K18838 cytoskeleton bundling-enhancing protein CbeA and related proteins | 4 | 9 | 1 | 2 | 3 | 10 | 5 | 5 | 73 | 65 | 74 | 66 |
| K18839 toxin GhoT | 0 | 1 | 1 | 0 | 0 | 2 | 0 | 2 | 2 | 10 | 13 | 3 |
| K18840 antitoxin GhoS | 0 | 1 | 1 | 0 | 5 | 1 | 4 | 7 | 26 | 7 | 13 | 12 |
| K18841 mRNA interferase ChpB [EC:3.1.-.-] | 67 | 64 | 62 | 62 | 78 | 97 | 81 | 86 | 4 | 0 | 0 | 1 |
| K18842 antitoxin ChpS | 0 | 0 | 0 | 0 | 0 | 0 | 4 | 4 | 2 | 0 | 0 | 2 |
| K18843 antitoxin HicB | 8 | 11 | 9 | 13 | 24 | 22 | 18 | 17 | 34 | 40 | 51 | 44 |
| K18862 small toxic polypeptide LdrA/B/C/D | 5 | 0 | 1 | 1 | 17 | 14 | 17 | 16 | 20 | 23 | 38 | 27 |
| K18919 protein HokC/D | 3 | 2 | 2 | 1 | 4 | 5 | 3 | 3 | 21 | 26 | 16 | 17 |
| K18920 protein HokA | 82 | 100 | 75 | 110 | 24 | 40 | 43 | 24 | 166 | 227 | 170 | 179 |
| K18921 protein HokB | 1 | 0 | 4 | 0 | 3 | 4 | 3 | 7 | 16 | 20 | 17 | 23 |
| K18922 protein HokE | 101 | 84 | 82 | 93 | 35 | 28 | 28 | 23 | 195 | 206 | 185 | 204 |
| K18923 antitoxin StbD | 87 | 95 | 83 | 90 | 137 | 155 | 120 | 109 | 21 | 10 | 12 | 15 |
| K19046 CRISPR system Cascade subunit CasB | 212 | 250 | 213 | 217 | 74 | 79 | 77 | 80 | 524 | 464 | 508 | 485 |
| K19048 toxic protein SymE | 328 | 325 | 320 | 318 | 230 | 254 | 267 | 240 | 331 | 392 | 379 | 364 |
| K19075 CRISPR-associated protein Cst2 | 0 | 4 | 3 | 0 | 4 | 0 | 0 | 0 | 0 | 0 | 5 | 0 |
| K19076 CRISPR-associated protein Cmr2 | 1 | 2 | 0 | 1 | 0 | 0 | 0 | 0 | 0 | 0 | 0 | 0 |
| K19088 CRISPR-associated protein Cst1 | 2 | 0 | 1 | 0 | 0 | 0 | 1 | 0 | 0 | 0 | 0 | 0 |
| K19090 CRISPR-associated protein Cas5t | 1 | 3 | 2 | 0 | 1 | 0 | 0 | 0 | 0 | 0 | 0 | 0 |
| K19091 CRISPR-associated endoribonuclease Cas6 [EC:3.1.-.-] | 4 | 6 | 9 | 11 | 13 | 5 | 8 | 9 | 74 | 72 | 108 | 112 |
| K19092 toxin ParE1/3/4 | 884 | 862 | 808 | 875 | 697 | 810 | 677 | 800 | 295 | 288 | 286 | 292 |
| K19114 CRISPR-associated protein Csh1 | 3 | 7 | 2 | 3 | 7 | 3 | 6 | 7 | 62 | 47 | 50 | 63 |
| K19115 CRISPR-associated protein Csh2 | 4 | 11 | 18 | 7 | 9 | 15 | 13 | 15 | 101 | 122 | 82 | 109 |
| K19116 CRISPR-associated protein Cas5h | 9 | 9 | 8 | 8 | 14 | 5 | 13 | 6 | 64 | 94 | 77 | 66 |
| K19117 CRISPR-associated protein Csd1 | 418 | 477 | 486 | 464 | 494 | 488 | 492 | 504 | 395 | 467 | 404 | 413 |
| K19118 CRISPR-associated protein Csd2 | 626 | 604 | 657 | 607 | 266 | 301 | 298 | 268 | 380 | 482 | 420 | 497 |
| K19119 CRISPR-associated protein Cas5d | 442 | 428 | 396 | 457 | 213 | 182 | 182 | 197 | 318 | 367 | 299 | 320 |
| K19123 CRISPR system Cascade subunit CasA | 542 | 554 | 562 | 581 | 168 | 186 | 221 | 188 | 1130 | 1049 | 1098 | 1288 |
| K19124 CRISPR system Cascade subunit CasC | 390 | 358 | 405 | 389 | 130 | 142 | 150 | 147 | 763 | 836 | 833 | 844 |
| K19125 CRISPR system Cascade subunit CasD | 279 | 300 | 285 | 252 | 121 | 74 | 87 | 75 | 455 | 568 | 479 | 504 |
| K19126 CRISPR system Cascade subunit CasE | 241 | 236 | 233 | 280 | 80 | 84 | 93 | 68 | 494 | 544 | 547 | 580 |
| K19127 CRISPR-associated protein Csy1 | 1283 | 1323 | 1287 | 1318 | 720 | 671 | 708 | 714 | 469 | 496 | 438 | 445 |
| K19128 CRISPR-associated protein Csy2 | 593 | 554 | 576 | 522 | 287 | 330 | 312 | 312 | 195 | 216 | 198 | 217 |
| K19129 CRISPR-associated protein Csy3 | 1103 | 1008 | 1021 | 1060 | 568 | 535 | 568 | 602 | 365 | 346 | 369 | 341 |
| K19130 CRISPR-associated endonuclease Csy4 [EC:3.1.-.-] | 763 | 774 | 734 | 770 | 422 | 424 | 378 | 432 | 243 | 263 | 283 | 224 |
| K19134 CRISPR-associated protein Csx10 | 68 | 54 | 60 | 47 | 2 | 1 | 0 | 3 | 0 | 0 | 0 | 0 |
| K19137 CRISPR-associated protein Csn2 | 38 | 35 | 28 | 39 | 6 | 13 | 10 | 12 | 32 | 46 | 33 | 32 |
| K19138 CRISPR-associated protein Csm2 | 0 | 0 | 1 | 1 | 0 | 0 | 0 | 3 | 0 | 0 | 2 | 1 |
| K19139 CRISPR-associated protein Csm4 | 0 | 2 | 0 | 0 | 0 | 0 | 0 | 0 | 2 | 3 | 3 | 9 |
| K19140 CRISPR-associated protein Csm5 | 2 | 0 | 0 | 1 | 0 | 0 | 2 | 0 | 1 | 0 | 0 | 2 |
| K19142 CRISPR-associated protein Cmr6 | 0 | 0 | 2 | 2 | 0 | 0 | 0 | 0 | 0 | 1 | 0 | 0 |
| K19145 CRISPR-associated protein Csx16 | 1 | 0 | 0 | 0 | 0 | 0 | 0 | 1 | 0 | 0 | 0 | 0 |
| K19147 5-methylcytosine-specific restriction enzyme subunit McrC | 576 | 628 | 600 | 619 | 724 | 762 | 824 | 804 | 264 | 225 | 229 | 287 |
| K19148 small toxic protein TisB | 31 | 25 | 22 | 41 | 12 | 6 | 13 | 9 | 52 | 44 | 64 | 55 |
| K19154 small toxic protein ShoB | 1 | 1 | 1 | 0 | 0 | 0 | 0 | 2 | 4 | 3 | 5 | 5 |
| K19155 toxin YhaV [EC:3.1.-.-] | 0 | 0 | 3 | 0 | 6 | 3 | 2 | 4 | 21 | 19 | 19 | 28 |
| K19156 antitoxin PrlF | 0 | 0 | 2 | 0 | 1 | 4 | 2 | 2 | 11 | 9 | 19 | 20 |
| K19157 mRNA interferase YafQ [EC:3.1.-.-] | 440 | 424 | 436 | 460 | 184 | 244 | 199 | 189 | 864 | 849 | 854 | 886 |
| K19158 toxin YoeB [EC:3.1.-.-] | 53 | 62 | 69 | 70 | 30 | 38 | 42 | 50 | 29 | 30 | 34 | 21 |
| K19159 antitoxin YefM | 2 | 2 | 5 | 7 | 3 | 5 | 3 | 2 | 21 | 21 | 13 | 14 |
| K19162 hha toxicity modulator TomB | 214 | 249 | 218 | 249 | 179 | 164 | 183 | 182 | 339 | 322 | 335 | 307 |
| K19163 toxin CcdB | 191 | 178 | 156 | 192 | 148 | 174 | 133 | 141 | 202 | 188 | 188 | 194 |
| K19164 antitoxin CcdA | 68 | 59 | 54 | 68 | 89 | 95 | 90 | 82 | 16 | 23 | 24 | 15 |
| K19165 antitoxin Phd | 113 | 123 | 128 | 130 | 41 | 36 | 42 | 33 | 250 | 278 | 278 | 295 |
| K19166 mRNA interferase HigB [EC:3.1.-.-] | 376 | 448 | 375 | 419 | 694 | 710 | 722 | 703 | 1137 | 1240 | 1206 | 1132 |
| K19167 protein AbiQ | 2 | 1 | 0 | 3 | 3 | 4 | 12 | 8 | 0 | 0 | 0 | 0 |
| K19168 toxin CptA | 899 | 967 | 915 | 861 | 532 | 509 | 500 | 557 | 524 | 599 | 522 | 544 |
| K19169 DNA sulfur modification protein DndB | 8 | 7 | 5 | 6 | 18 | 28 | 33 | 24 | 26 | 28 | 18 | 20 |
| K19170 DNA sulfur modification protein DndC | 6 | 9 | 6 | 10 | 7 | 1 | 10 | 9 | 32 | 28 | 36 | 26 |
| K19171 DNA sulfur modification protein DndD | 55 | 52 | 69 | 62 | 8 | 3 | 8 | 3 | 154 | 158 | 168 | 173 |
| K19172 DNA sulfur modification protein DndE | 3 | 10 | 10 | 12 | 1 | 0 | 1 | 2 | 8 | 2 | 6 | 8 |
| K19175 DNA phosphorothioation-dependent restriction protein DptH | 3 | 5 | 3 | 6 | 1 | 2 | 2 | 3 | 0 | 0 | 0 | 0 |
| K19334 biofilm protein TabA | 460 | 487 | 457 | 493 | 246 | 267 | 263 | 266 | 579 | 544 | 547 | 539 |
| K19686 ribonuclease VapC [EC:3.1.-.-] | 0 | 0 | 0 | 0 | 0 | 0 | 0 | 0 | 0 | 0 | 2 | 0 | 1 |
| K19687 antitoxin VapB | 0 | 0 | 0 | 0 | 0 | 0 | 0 | 0 | 1 | 0 | 0 | 1 |
| K19688 biofilm regulator BssR | 207 | 231 | 236 | 231 | 166 | 186 | 153 | 163 | 376 | 349 | 333 | 310 |
| K21487 toxin YobL [EC:3.1.-.-] | 0 | 8 | 9 | 2 | 10 | 0 | 0 | 0 | 0 | 0 | 6 | 0 |
| K21488 antitoxin YobK | 98 | 79 | 99 | 98 | 363 | 309 | 381 | 373 | 267 | 227 | 279 | 261 |
| K21493 toxin YxiD [EC:3.1.-.-] | 1 | 0 | 0 | 5 | 0 | 0 | 0 | 5 | 0 | 0 | 0 | 0 |
| K21495 antitoxin FitA | 1 | 0 | 0 | 0 | 0 | 0 | 0 | 0 | 0 | 1 | 1 | 1 |
| K21498 antitoxin HigA-1 | 717 | 762 | 780 | 726 | 776 | 737 | 707 | 771 | 517 | 510 | 503 | 507 |
| ko04030 G protein-coupled receptors | 0 | 2 | 0 | 0 | 0 | 2 | 0 | 0 | 1 | 0 | 0 | 0 |
| ko04031 GTP-binding proteins | 0 | 0 | 1 | 0 | 1 | 0 | 0 | 1 | 0 | 0 | 0 | 1 |
| ko04090 CD molecules | 3123 | 3443 | 3555 | 3351 | 10264 | 10371 | 10615 | 10264 | 7421 | 7170 | 7332 | 6944 |
| ko00536 Glycosaminoglycan binding proteins | 12786 | 12488 | 11519 | 12000 | 6736 | 6628 | 4019 | 4555 | 4275 | 3915 | 3848 | 4035 |
| ko00537 Glycosylphosphatidylinositol (GPI)-anchored proteins | 11270 | 11437 | 11492 | 11569 | 11459 | 11673 | 11658 | 11731 | 11295 | 11507 | 11411 | 11418 |
| ko01504 Antimicrobial resistance genes | 137682 | 137214 | 134550 | 136095 | 112296 | 112557 | 108757 | 112156 | 98321 | 96351 | 95857 | 97439 |
| K00638 chloramphenicol O-acetyltransferase type B [EC:2.3.1.28] | 906 | 991 | 926 | 948 | 535 | 503 | 464 | 543 | 334 | 271 | 271 | 300 |
| K00662 aminoglycoside 3-N-acetyltransferase [EC:2.3.1.81] | 164 | 149 | 137 | 154 | 12 | 16 | 14 | 10 | 162 | 156 | 193 | 164 |
| K00663 aminoglycoside 6'-N-acetyltransferase [EC:2.3.1.82] | 0 | 2 | 3 | 1 | 0 | 0 | 0 | 0 | 4 | 13 | 7 | 5 |
| K00897 kanamycin kinase [EC:2.7.1.95] | 3 | 3 | 5 | 7 | 2 | 2 | 2 | 2 | 0 | 2 | 2 | 0 |
| K00984 streptomycin 3-adenylyltransferase [EC:2.7.7.47] | 38 | 48 | 41 | 39 | 62 | 46 | 56 | 57 | 108 | 83 | 111 | 104 |
| K04343 streptomycin 6-kinase [EC:2.7.1.72] | 1036 | 978 | 994 | 1000 | 564 | 558 | 512 | 485 | 298 | 282 | 325 | 319 |
| K05593 aminoglycoside 6-adenylyltransferase [EC:2.7.7.-] | 1101 | 1093 | 1223 | 1150 | 1838 | 1733 | 1781 | 1774 | 2144 | 1993 | 2083 | 2078 |
| K06880 erythromycin esterase [EC:3.1.1.-] | 445 | 430 | 472 | 493 | 25 | 31 | 23 | 32 | 389 | 445 | 428 | 436 |
| K06979 macrolide phosphotransferase | 0 | 0 | 0 | 0 | 2 | 2 | 0 | 2 | 4 | 0 | 3 | 4 |
| K10673 streptomycin 3-kinase [EC:2.7.1.87] | 0 | 0 | 0 | 0 | 0 | 0 | 0 | 0 | 1 | 2 | 2 | 1 |
| K17910 aminoglycoside 2''-phosphotransferase [EC:2.7.1.190] | 0 | 0 | 0 | 0 | 0 | 0 | 0 | 1 | 0 | 0 | 0 | 0 |
| K18220 ribosomal protection tetracycline resistance protein | 2052 | 1884 | 2078 | 2086 | 2379 | 2257 | 2367 | 2180 | 2889 | 2838 | 2833 | 2991 |
| K18234 virginiamycin A acetyltransferase [EC:2.3.1.-] | 737 | 843 | 840 | 842 | 1399 | 1438 | 1457 | 1398 | 1430 | 1402 | 1523 | 1385 |

| | | | | | | | | | | | | |
|---|---|---|---|---|---|---|---|---|---|---|---|---|
| K18235 virginiamycin B lyase [EC:4.2.99.-] | 1648 | 1626 | 1535 | 1623 | 949 | 925 | 486 | 554 | 488 | 520 | 517 | 545 |
| K18554 chloramphenicol 3-O phosphotransferase [EC:2.7.1.-] | 0 | 0 | 0 | 1 | 3 | 0 | 0 | 0 | 0 | 0 | 0 | 0 |
| K18555 fluoroquinolone resistance protein | 0 | 0 | 0 | 0 | 0 | 0 | 0 | 0 | 5 | 2 | 0 | 4 |
| K18815 aminoglycoside 6'-N-acetyltransferase I [EC:2.3.1.82] | 33 | 42 | 40 | 38 | 11 | 15 | 8 | 7 | 72 | 63 | 58 | 68 |
| K18816 aminoglycoside 6'-N-acetyltransferase I [EC:2.3.1.82] | 4 | 2 | 4 | 6 | 20 | 27 | 19 | 24 | 6 | 7 | 7 | 4 |
| K19271 chloramphenicol O-acetyltransferase type A [EC:2.3.1.28] | 1312 | 1279 | 1413 | 1391 | 1704 | 1679 | 1651 | 1761 | 1813 | 1853 | 1870 | 1871 |
| K19272 aminoglycoside 3'-phosphotransferase I [EC:2.7.1.95] | 0 | 0 | 0 | 0 | 0 | 0 | 1 | 2 | 1 | 6 | 0 | 0 |
| K19273 streptothricin acetyltransferase [EC:2.3.-.-] | 54 | 56 | 50 | 53 | 3 | 7 | 2 | 2 | 0 | 5 | 2 | 3 |
| K19275 aminoglycoside 3-N-acetyltransferase II [EC:2.3.1.81] | 0 | 0 | 1 | 0 | 0 | 0 | 0 | 0 | 0 | 0 | 5 | 0 |
| K19278 aminoglycoside 6'-N-acetyltransferase Ib [EC:2.3.1.82] | 3 | 2 | 3 | 4 | 25 | 19 | 24 | 22 | 0 | 1 | 1 | 0 |
| K19299 aminoglycoside 3'-phosphotransferase III [EC:2.7.1.95] | 93 | 66 | 69 | 103 | 11 | 6 | 9 | 15 | 1 | 2 | 5 | 7 |
| K19300 aminoglycoside 3'-phosphotransferase II [EC:2.7.1.95] | 1371 | 1300 | 1270 | 1315 | 705 | 745 | 672 | 740 | 422 | 348 | 424 | 424 |
| K19545 lincosamide nucleotidyltransferase A/C/D/E | 30 | 44 | 53 | 45 | 120 | 135 | 114 | 109 | 21 | 40 | 24 | 23 |
| K19883 bifunctional aminoglycoside 6'-N-acetyltransferase / aminoglycoside 2''-phosphotransferase [EC:2.3.1.82 2.7.1.190] | 11 | 5 | 9 | 9 | 26 | 30 | 34 | 23 | 0 | 4 | 8 | 4 |
| K21252 fosfomycin resistance protein FosX | 1 | 2 | 0 | 0 | 0 | 0 | 2 | 1 | 4 | 2 | 0 | 0 |
| K21253 glutathione S-transferase fosA [EC:2.5.1.18] | 691 | 611 | 558 | 628 | 347 | 302 | 284 | 348 | 205 | 175 | 167 | 211 |
| K21265 glutathione S-transferase fosA5 [EC:2.5.1.18] | 235 | 258 | 248 | 282 | 181 | 164 | 158 | 171 | 388 | 348 | 317 | 361 |
| Not Included in Pathway or Brite | 1286209 | 1286157 | 1270584 | 1287391 | 1247267 | 1255851 | 1233992 | 1242086 | 1221688 | 1222085 | 1215842 | 1223116 |
| Unclassified: metabolism | 532051 | 531480 | 522219 | 529695 | 514625 | 518736 | 514880 | 515915 | 514840 | 515478 | 511658 | 516219 |
| ko99980 Enzymes with EC numbers | 437737 | 437440 | 431257 | 436711 | 442535 | 445609 | 443797 | 442622 | 435652 | 435982 | 433944 | 435628 |
| K00046 gluconate 5-dehydrogenase [EC:1.1.1.69] | 1400 | 1484 | 1627 | 1316 | 2431 | 2388 | 2568 | 2357 | 2272 | 2269 | 2350 | 2215 |
| K00063 D-arabinose 1-dehydrogenase [EC:1.1.1.117] | 11 | 10 | 6 | 3 | 0 | 0 | 0 | 0 | 1 | 2 | 3 | 2 |
| K00091 dihydroflavonol-4-reductase [EC:1.1.1.219] | 1278 | 1344 | 1248 | 1261 | 680 | 704 | 681 | 688 | 650 | 614 | 587 | 565 |
| K00098 L-idonate 5-dehydrogenase [EC:1.1.1.264] | 222 | 215 | 218 | 245 | 339 | 347 | 365 | 307 | 63 | 78 | 81 | 60 |
| K00119 alcohol dehydrogenase (nicotinoprotein) [EC:1.1.99.36] | 0 | 1 | 0 | 0 | 0 | 0 | 0 | 0 | 0 | 0 | 0 | 1 |
| K00153 S-(hydroxymethyl)mycothiol dehydrogenase [EC:1.1.1.306] | 1 | 0 | 0 | 0 | 0 | 1 | 0 | 0 | 0 | 0 | 0 | 0 |
| K00154 coniferyl-aldehyde dehydrogenase [EC:1.2.1.68] | 2579 | 2568 | 2297 | 2440 | 1396 | 1421 | 1201 | 1385 | 753 | 772 | 759 | 796 |
| K00179 indolepyruvate ferredoxin oxidoreductase, alpha subunit [EC:1.2.7.8] | 1241 | 1392 | 1526 | 1362 | 3002 | 3313 | 3246 | 3171 | 2618 | 2545 | 2492 | 2465 |
| K00180 indolepyruvate ferredoxin oxidoreductase, beta subunit [EC:1.2.7.8] | 407 | 459 | 474 | 475 | 1148 | 1194 | 1137 | 1105 | 1002 | 877 | 909 | 900 |
| K00219 2,4-dienoyl-CoA reductase (NADPH2) [EC:1.3.1.34] | 7287 | 7058 | 6376 | 7036 | 3916 | 3939 | 3732 | 4163 | 3648 | 3562 | 3319 | 3687 |
| K00221 alkylmercury lyase [EC:4.99.1.2] | 0 | 0 | 0 | 0 | 0 | 0 | 0 | 0 | 0 | 0 | 1 | 3 |
| K00298 N5-(carboxyethyl)ornithine synthase [EC:1.5.1.24] | 1 | 2 | 1 | 3 | 19 | 23 | 28 | 15 | 0 | 0 | 0 | 1 |
| K00311 electron-transferring-flavoprotein dehydrogenase [EC:1.5.5.1] | 2197 | 2191 | 2092 | 2242 | 1201 | 1200 | 1126 | 1233 | 696 | 625 | 712 | 678 |
| K00313 electron transfer flavoprotein-quinone oxidoreductase [EC:1.5.5.-] | 614 | 578 | 641 | 627 | 905 | 938 | 1021 | 911 | 369 | 342 | 347 | 313 |
| K00344 NADPH:quinone reductase [EC:1.6.5.5] | 3174 | 3057 | 2843 | 3074 | 1914 | 1986 | 2004 | 2169 | 1662 | 1443 | 1452 | 1557 |
| K00346 Na+-transporting NADH:ubiquinone oxidoreductase subunit A [EC:7.2.1.1] | 2033 | 1991 | 2147 | 2017 | 2131 | 2081 | 2104 | 2109 | 1554 | 1414 | 1458 | 1484 |
| K00347 Na+-transporting NADH:ubiquinone oxidoreductase subunit B [EC:7.2.1.1] | 2884 | 2973 | 3022 | 2953 | 3406 | 3504 | 3470 | 3595 | 3299 | 3214 | 3213 | 3120 |
| K00348 Na+-transporting NADH:ubiquinone oxidoreductase subunit C [EC:7.2.1.1] | 1606 | 1610 | 1737 | 1753 | 1971 | 2027 | 2094 | 2018 | 1811 | 1826 | 1787 | 1764 |
| K00349 Na+-transporting NADH:ubiquinone oxidoreductase subunit D [EC:7.2.1.1] | 1341 | 1369 | 1288 | 1220 | 1709 | 1678 | 1751 | 1678 | 1709 | 1518 | 1532 | 1583 |
| K00350 Na+-transporting NADH:ubiquinone oxidoreductase subunit E [EC:7.2.1.1] | 1432 | 1380 | 1506 | 1402 | 1692 | 1676 | 1604 | 1723 | 1578 | 1518 | 1526 | 1495 |
| K00351 Na+-transporting NADH:ubiquinone oxidoreductase subunit F [EC:7.2.1.1] | 1102 | 1198 | 1197 | 1204 | 2510 | 2644 | 2621 | 2560 | 2428 | 2301 | 2363 | 2376 |
| K00354 NADPH2 dehydrogenase [EC:1.6.99.1] | 15 | 5 | 7 | 7 | 3 | 5 | 3 | 3 | 15 | 8 | 9 | 7 |
| K00428 cytochrome c peroxidase [EC:1.11.1.5] | 1524 | 1414 | 1517 | 1574 | 955 | 865 | 836 | 845 | 595 | 562 | 552 | 585 |
| K00433 non-heme chloroperoxidase [EC:1.11.1.10] | 1314 | 1203 | 1202 | 1274 | 521 | 517 | 487 | 511 | 1216 | 1162 | 1258 | 1182 |
| K00437 [NiFe] hydrogenase large subunit [EC:1.12.2.1] | 14 | 16 | 11 | 15 | 3 | 5 | 6 | 2 | 9 | 32 | 15 | 21 |
| K00520 mercuric reductase [EC:1.16.1.1] | 3453 | 3328 | 3393 | 3341 | 2355 | 2357 | 2394 | 2327 | 962 | 928 | 848 | 875 |
| K00532 ferredoxin hydrogenase [EC:1.12.7.2] | 0 | 2 | 5 | 0 | 0 | 0 | 0 | 0 | 13 | 15 | 10 | 14 |
| K00533 ferredoxin hydrogenase large subunit [EC:1.12.7.2] | 30 | 33 | 33 | 32 | 23 | 31 | 45 | 43 | 40 | 40 | 57 | 50 |
| K00534 ferredoxin hydrogenase small subunit [EC:1.12.7.2] | 2 | 2 | 2 | 1 | 2 | 2 | 1 | 1 | 1 | 0 | 0 | 0 |
| K00537 arsenate reductase (glutaredoxin) [EC:1.20.4.1] | 1549 | 1533 | 1649 | 1645 | 1707 | 1747 | 1747 | 1747 | 1614 | 1466 | 1612 | 1653 |
| K00573 protein-L-isoaspartate(D-aspartate) O-methyltransferase [EC:2.1.1.77] | 1440 | 1403 | 1296 | 1384 | 823 | 873 | 822 | 821 | 827 | 851 | 827 | 826 |
| K00574 cyclopropane-fatty-acyl-phospholipid synthase [EC:2.1.1.79] | 4102 | 3968 | 3878 | 4032 | 2403 | 2470 | 2445 | 2540 | 3089 | 3323 | 2982 | 3308 |
| K00598 trans-aconitate 2-methyltransferase [EC:2.1.1.144] | 1470 | 1536 | 1405 | 1499 | 775 | 763 | 732 | 774 | 958 | 1036 | 968 | 1080 |
| K00612 carbamoyltransferase [EC:2.1.3.-] | 4985 | 4813 | 4695 | 4599 | 2587 | 2488 | 2580 | 2818 | 1604 | 1484 | 1378 | 1500 |
| K00633 galactoside O-acetyltransferase [EC:2.3.1.18] | 179 | 178 | 188 | 173 | 106 | 146 | 237 | 220 | 287 | 283 | 309 | 318 |
| K00661 maltose O-acetyltransferase [EC:2.3.1.79] | 1866 | 1709 | 1868 | 1765 | 2611 | 2835 | 2581 | 2498 | 3133 | 3199 | 3257 | 3178 |
| K00675 N-hydroxyarylamine O-acetyltransferase [EC:2.3.1.118] | 1849 | 1860 | 1832 | 1827 | 1146 | 1063 | 1028 | 1103 | 1285 | 1193 | 1203 | 1176 |
| K00684 leucyl/phenylalanyl-tRNA---protein transferase [EC:2.3.2.6] | 1330 | 1296 | 1304 | 1409 | 867 | 799 | 863 | 780 | 953 | 838 | 850 | 924 |
| K00754 L-malate glycosyltransferase [EC:2.4.1.-] | 3 | 0 | 1 | 2 | 2 | 3 | 0 | 1 | 2 | 0 | 0 | 1 |
| K00786 glycosyltransferase [EC:2.4.-.-] | 34 | 18 | 47 | 31 | 5 | 0 | 9 | 5 | 30 | 32 | 36 | 29 |
| K00809 deoxyhypusine synthase [EC:2.5.1.46] | 1 | 2 | 2 | 3 | 2 | 2 | 1 | 0 | 3 | 10 | 2 | 12 |
| K00862 erythritol kinase (D-erythritol 1-phosphate-forming) [EC:2.7.1.215] | 1 | 1 | 0 | 0 | 0 | 2 | 0 | 0 | 4 | 1 | 0 | 3 |
| K00906 isocitrate dehydrogenase kinase/phosphatase [EC:2.7.11.5 3.1.3.-] | 3721 | 3643 | 3445 | 3612 | 2290 | 2252 | 2120 | 2079 | 2278 | 2493 | 2246 | 2354 |
| K00924 kinase [EC:2.7.1.-] | 1 | 0 | 0 | 0 | 0 | 0 | 0 | 0 | 0 | 0 | 0 | 0 |
| K00982 [glutamine synthetase] adenylyltransferase / [glutamine synthetase]-adenylyl-L-tyrosine phosphorylase [EC:2.7.7.42 2.7.7.89] | 6777 | 6719 | 6589 | 6610 | 3951 | 4214 | 4150 | 4148 | 4419 | 4080 | 3935 | 4426 |
| K00986 RNA-directed DNA polymerase [EC:2.7.7.49] | 5336 | 5187 | 5345 | 5269 | 2996 | 2902 | 2742 | 2906 | 1716 | 1725 | 1833 | 1793 |
| K01002 phosphoglycerol transferase [EC:2.7.8.20] | 1309 | 1476 | 1377 | 1487 | 1057 | 1048 | 1014 | 982 | 2004 | 2203 | 2075 | 2255 |
| K01023 arylsulfate sulfotransferase [EC:2.8.2.22] | 2643 | 2566 | 2575 | 2522 | 1939 | 1994 | 1948 | 1921 | 2644 | 2590 | 2688 | 2558 |
| K01050 cholinesterase [EC:3.1.1.8] | 4 | 2 | 2 | 1 | 0 | 0 | 0 | 0 | 4 | 4 | 3 | 3 |
| K01066 acetyl esterase [EC:3.1.1.-] | 2189 | 2314 | 2216 | 2225 | 1520 | 1457 | 1434 | 1513 | 1507 | 1518 | 1397 | 1528 |
| K01073 acyl-CoA hydrolase [EC:3.1.2.20] | 0 | 0 | 0 | 1 | 0 | 0 | 0 | 0 | 0 | 1 | 0 | 0 |
| K01090 protein phosphatase [EC:3.1.3.16] | 4 | 4 | 2 | 1 | 2 | 0 | 2 | 0 | 0 | 0 | 0 | 0 |
| K01118 FMN-dependent NADH-azoreductase [EC:1.7.1.17] | 2408 | 2436 | 2353 | 2345 | 1337 | 1369 | 1327 | 1450 | 1638 | 1596 | 1487 | 1602 |
| K01133 choline-sulfatase [EC:3.1.6.6] | 3914 | 3944 | 3938 | 3804 | 4225 | 4374 | 4159 | 4385 | 3245 | 3226 | 3194 | 3086 |
| K01138 uncharacterized sulfatase [EC:3.1.6.-] | 31 | 28 | 34 | 48 | 48 | 67 | 66 | 51 | 187 | 241 | 209 | 183 |
| K01143 exodeoxyribonuclease (lambda-induced) [EC:3.1.11.3] | 0 | 0 | 0 | 1 | 5 | 5 | 4 | 8 | 2 | 1 | 3 | 1 |
| K01144 exodeoxyribonuclease V [EC:3.1.11.5] | 232 | 236 | 248 | 277 | 501 | 483 | 529 | 538 | 202 | 238 | 234 | 229 |
| K01150 deoxyribonuclease I [EC:3.1.21.1] | 1512 | 1506 | 1472 | 1503 | 915 | 947 | 822 | 929 | 976 | 992 | 1024 | 1030 |
| K01174 micrococcal nuclease [EC:3.1.31.1] | 76 | 44 | 50 | 70 | 4 | 1 | 9 | 13 | 39 | 46 | 34 | 52 |
| K01175 esterase [EC:3.1.-.-] | 485 | 499 | 456 | 487 | 350 | 358 | 288 | 331 | 572 | 705 | 634 | 715 |
| K01180 endo-1,3(4)-beta-glucanase [EC:3.2.1.6] | 0 | 0 | 0 | 0 | 0 | 0 | 0 | 1 | 2 | 0 | 0 | 0 |
| K01181 endo-1,4-beta-xylanase [EC:3.2.1.8] | 1114 | 1306 | 1424 | 1342 | 4610 | 4266 | 4559 | 4452 | 3280 | 3120 | 3293 | 3066 |
| K01185 lysozyme [EC:3.2.1.17] | 1091 | 1088 | 1207 | 1150 | 2493 | 2542 | 2547 | 2546 | 1575 | 1490 | 1500 | 1635 |
| K01215 glucan 1,6-alpha-glucosidase [EC:3.2.1.70] | 136 | 188 | 167 | 195 | 46 | 32 | 42 | 24 | 208 | 229 | 206 | 239 |
| K01219 beta-agarase [EC:3.2.1.81] | 1 | 4 | 1 | 0 | 0 | 0 | 0 | 0 | 0 | 0 | 1 | 0 |
| K01224 arabinogalactan endo-1,4-beta-galactosidase [EC:3.2.1.89] | 1045 | 1043 | 1061 | 1079 | 986 | 1009 | 1044 | 975 | 916 | 977 | 927 | 1003 |
| K01235 alpha-glucuronidase [EC:3.2.1.139] | 1903 | 2004 | 2062 | 1937 | 5592 | 5467 | 5654 | 5562 | 4759 | 4623 | 4946 | 4528 |
| K01250 pyrimidine-specific ribonucleoside hydrolase [EC:3.2.-.-] | 946 | 853 | 924 | 922 | 505 | 487 | 478 | 455 | 974 | 1056 | 966 | 1101 |
| K01274 D-alanyl-D-alanine dipeptidase [EC:3.4.13.-] | 0 | 0 | 0 | 0 | 0 | 0 | 1 | 0 | 0 | 0 | 2 | 3 |
| K01286 D-alanyl-D-alanine carboxypeptidase [EC:3.4.16.4] | 104 | 72 | 99 | 75 | 8 | 16 | 17 | 10 | 76 | 120 | 119 | 108 |
| K01447 N-acetylmuramoyl-L-alanine amidase [EC:3.5.1.28] | 1357 | 1312 | 1259 | 1281 | 1170 | 1172 | 1032 | 1220 | 735 | 673 | 731 | 714 |
| K01449 N-acetylmuramoyl-L-alanine amidase [EC:3.5.1.28] | 340 | 370 | 338 | 429 | 110 | 129 | 125 | 84 | 275 | 349 | 329 | 365 |
| K01461 N-acyl-D-glutamate deacylase [EC:3.5.1.82] | 0 | 0 | 0 | 0 | 0 | 0 | 0 | 0 | 1 | 0 | 0 | 0 |
| K01462 peptide deformylase [EC:3.5.1.88] | 2994 | 2874 | 3015 | 2838 | 2477 | 2504 | 2611 | 2642 | 2395 | 2516 | 2458 | 2478 |
| K01463 N-acetylglucosamine malate deacetylase 1 [EC:3.5.1.-] | 25 | 18 | 18 | 33 | 1 | 0 | 0 | 0 | 61 | 60 | 57 | 62 |
| K01475 isatin hydrolase [EC:3.5.2.20] | 0 | 0 | 1 | 1 | 1 | 2 | 1 | 0 | 1 | 1 | 0 | 0 |
| K01526 guanosine-diphosphatase [EC:3.6.1.42] | 0 | 0 | 2 | 0 | 0 | 0 | 0 | 0 | 0 | 0 | 0 | 0 |
| K01529 inosine/xanthosine triphosphatase [EC:3.6.1.-] | 294 | 308 | 337 | 367 | 220 | 299 | 223 | 233 | 437 | 457 | 415 | 423 |
| K01530 phospholipid-translocating ATPase [EC:7.6.2.1] | 0 | 2 | 0 | 0 | 0 | 0 | 0 | 0 | 0 | 0 | 0 | 0 |
| K01531 P-type Mg2+ transporter [EC:7.2.2.14] | 5853 | 5767 | 5504 | 5708 | 3056 | 3254 | 3186 | 3088 | 5485 | 6163 | 5833 | 5893 |
| K01533 P-type Cu2+ transporter [EC:7.2.2.9] | 4687 | 4700 | 4836 | 4818 | 6208 | 6204 | 6110 | 6128 | 4162 | 4113 | 4461 | 4127 |
| K01534 Cd2+/Zn2+-exporting ATPase [EC:3.6.3.3 7.2.2.12] | 7640 | 7815 | 7493 | 7801 | 7676 | 7587 | 7833 | 7728 | 7780 | 8037 | 7768 | 7759 |
| K01536 P-type Na+/K+ transporter [EC:7.2.2.3 7.2.2.-] | 1 | 0 | 0 | 0 | 0 | 0 | 0 | 0 | 0 | 0 | 1 | 1 |
| K01537 P-type Ca2+ transporter type 2C [EC:7.2.2.10] | 5918 | 5708 | 6117 | 5789 | 7139 | 7144 | 7414 | 7174 | 7673 | 7943 | 8130 | 8045 |
| K01567 peptidoglycan-N-acetylmuramic acid deacetylase [EC:3.5.1.-] | 338 | 295 | 314 | 330 | 68 | 92 | 72 | 89 | 279 | 303 | 291 | 288 |
| K01616 2-oxoglutarate decarboxylase [EC:4.1.1.71] | 0 | 0 | 0 | 0 | 0 | 0 | 0 | 1 | 0 | 0 | 0 | 0 |
| K01671 sulfofructosephosphate aldolase [EC:4.1.2.57] | 202 | 179 | 214 | 207 | 346 | 345 | 308 | 352 | 45 | 53 | 37 | 58 |
| K01683 D-arabinonate dehydratase [EC:4.2.1.5] | 8 | 4 | 2 | 5 | 1 | 2 | 0 | 1 | 3 | 5 | 3 | 2 |
| K01726 gamma-carbonic anhydrase [EC:4.2.1.-] | 2 | 1 | 4 | 1 | 0 | 0 | 43 | 40 | 0 | 0 | 1 | 0 |
| K01727 hyaluronate lyase [EC:4.2.2.1] | 2388 | 2249 | 2327 | 2332 | 510 | 486 | 426 | 441 | 1451 | 1716 | 1519 | 1573 |
| K01751 diaminopropionate ammonia-lyase [EC:4.3.1.15] | 1205 | 1265 | 1182 | 1222 | 236 | 277 | 319 | 247 | 1537 | 1733 | 1579 | 1721 |
| K01766 cysteine sulfinate desulfinase [EC:4.4.1.-] | 2127 | 2152 | 1959 | 2084 | 1324 | 1267 | 1226 | 1307 | 1465 | 1437 | 1416 | 1570 |
| K01780 histidine racemase [EC:5.1.1.24] | 42 | 33 | 52 | 59 | 9 | 9 | 3 | 7 | 108 | 93 | 98 | 93 |
| K01802 peptidylprolyl isomerase [EC:5.2.1.8] | 720 | 677 | 683 | 649 | 710 | 615 | 546 | 548 | 415 | 417 | 421 | 453 |
| K01811 alpha-D-xyloside xylohydrolase [EC:3.2.1.177] | 8507 | 8600 | 8912 | 8899 | 13729 | 13646 | 13943 | 13554 | 14911 | 14534 | 14856 | 14297 |
| K01932 gamma-polyglutamate synthase [EC:6.3.2.-] | 3 | 2 | 2 | 3 | 0 | 1 | 0 | 1 | 0 | 0 | 5 | 1 |
| K01974 RNA 3'-terminal phosphate cyclase (ATP) [EC:6.5.1.4] | 44 | 40 | 36 | 25 | 29 | 37 | 133 | 151 | 60 | 65 | 61 | 53 |
| K02082 tagatose-6-phosphate ketose/aldose isomerase [EC:5.-.-.-] | 298 | 326 | 325 | 378 | 436 | 418 | 428 | 409 | 97 | 79 | 86 | 122 |
| K02182 carnitine-CoA ligase [EC:6.2.1.48] | 328 | 304 | 345 | 314 | 479 | 515 | 527 | 477 | 102 | 77 | 97 | 94 |
| K02217 ferritin [EC:1.16.3.2] | 951 | 945 | 1036 | 992 | 1462 | 1385 | 1402 | 1386 | 1315 | 1297 | 1262 | 1309 |
| K02334 DNA polymerase bacteriophage-type [EC:2.7.7.7] | 230 | 210 | 235 | 214 | 217 | 217 | 243 | 220 | 522 | 525 | 512 | 542 |

| | | | | | | | | | | | | |
|---|---|---|---|---|---|---|---|---|---|---|---|---|
| K02352 lactonase [EC:3.1.1.-] | 239 | 238 | 271 | 287 | 71 | 54 | 61 | 65 | 561 | 643 | 581 | 607 |
| K02431 L-fucose mutarotase [EC:5.1.3.29] | 340 | 334 | 285 | 339 | 91 | 107 | 119 | 128 | 639 | 668 | 638 | 632 |
| K02467 arabinose 5-phosphate isomerase [EC:5.3.1.13] | 566 | 590 | 548 | 637 | 390 | 398 | 424 | 414 | 852 | 775 | 841 | 856 |
| K02558 UDP-N-acetylmuramate: L-alanyl-gamma-D-glutamyl-meso-diaminopimelate ligase [EC:6.3.2.45] | 3195 | 3127 | 2938 | 3029 | 1924 | 1959 | 1902 | 1978 | 2038 | 2079 | 1810 | 2016 |
| K02846 N-methyl-L-tryptophan oxidase [EC:1.5.3.-] | 573 | 612 | 586 | 605 | 402 | 416 | 393 | 449 | 884 | 907 | 877 | 888 |
| K03207 colanic acid biosynthesis protein WcaH [EC:3.6.1.-] | 110 | 63 | 69 | 74 | 129 | 129 | 127 | 141 | 29 | 36 | 31 | 26 |
| K03387 alkyl hydroperoxide reductase subunit F [EC:1.6.4.-] | 2467 | 2508 | 2509 | 2517 | 4401 | 4369 | 4239 | 4367 | 3952 | 3967 | 3993 | 3895 |
| K03424 TatD DNase family protein [EC:3.1.21.-] | 6516 | 6707 | 6477 | 6518 | 6537 | 6717 | 6481 | 6635 | 6397 | 6483 | 6037 | 6214 |
| K03478 chitin disaccharide deacetylase [EC:3.5.1.105] | 460 | 460 | 435 | 469 | 283 | 324 | 294 | 299 | 590 | 673 | 657 | 669 |
| K03518 aerobic carbon-monoxide dehydrogenase small subunit [EC:1.2.5.3] | 751 | 741 | 709 | 706 | 173 | 204 | 204 | 197 | 516 | 554 | 535 | 512 |
| K03519 aerobic carbon-monoxide dehydrogenase medium subunit [EC:1.2.5.3] | 86 | 80 | 80 | 74 | 14 | 12 | 15 | 20 | 58 | 62 | 71 | 47 |
| K03520 aerobic carbon-monoxide dehydrogenase large subunit [EC:1.2.5.3] | 0 | 1 | 2 | 1 | 0 | 0 | 0 | 1 | 0 | 3 | 2 | 3 |
| K03534 L-rhamnose mutarotase [EC:5.1.3.32] | 827 | 854 | 836 | 928 | 1732 | 1736 | 1785 | 1735 | 1475 | 1547 | 1571 | 1534 |
| K03564 peroxiredoxin Q/BCP [EC:1.11.1.15] | 1489 | 1524 | 1465 | 1572 | 1628 | 1607 | 1641 | 1589 | 1507 | 1512 | 1552 | 1468 |
| K03578 ATP-dependent helicase HrpA [EC:3.6.4.13] | 8193 | 8066 | 7896 | 8087 | 4979 | 4854 | 4632 | 5044 | 5089 | 5048 | 4972 | 5300 |
| K03579 ATP-dependent helicase HrpB [EC:3.6.4.13] | 2635 | 2606 | 2287 | 2537 | 1634 | 1599 | 1786 | 1847 | 2407 | 2238 | 2337 | 2415 |
| K03614 Na+-translocating ferredoxin:NAD+ oxidoreductase subunit D [EC:7.2.1.2] | 1788 | 1881 | 1906 | 1932 | 2678 | 2690 | 2784 | 2637 | 2813 | 2798 | 2721 | 2671 |
| K03615 Na+-translocating ferredoxin:NAD+ oxidoreductase subunit C [EC:7.2.1.2] | 2458 | 2390 | 2561 | 2449 | 3378 | 3511 | 3813 | 3847 | 3713 | 3866 | 3575 | 3671 |
| K03616 Na+-translocating ferredoxin:NAD+ oxidoreductase subunit B [EC:7.2.1.2] | 1459 | 1487 | 1442 | 1434 | 2233 | 2227 | 2099 | 2110 | 2132 | 2182 | 2129 | 2002 |
| K03716 spore photoproduct lyase [EC:4.1.99.14] | 384 | 345 | 417 | 334 | 77 | 69 | 82 | 63 | 409 | 459 | 388 | 458 |
| K03725 archaea-specific helicase [EC:3.6.4.-] | 0 | 2 | 3 | 2 | 1 | 3 | 3 | 2 | 4 | 13 | 2 | 10 |
| K03726 helicase [EC:3.6.4.-] | 847 | 850 | 781 | 872 | 330 | 352 | 337 | 314 | 2230 | 2165 | 2111 | 2204 |
| K03727 ATP-dependent RNA helicase HelY [EC:3.6.4.-] | 4 | 3 | 2 | 5 | 3 | 4 | 3 | 7 | 10 | 23 | 12 | 11 |
| K03734 FAD:protein FMN transferase [EC:2.7.1.180] | 4499 | 4658 | 4617 | 4611 | 5895 | 5853 | 6076 | 5804 | 5568 | 5231 | 5525 | 5470 |
| K03741 arsenate reductase (thioredoxin) [EC:1.20.4.4] | 1149 | 1264 | 1242 | 1212 | 1661 | 1661 | 1592 | 1723 | 1019 | 983 | 1065 | 1011 |
| K03793 pteridine reductase [EC:1.5.1.33] | 0 | 0 | 0 | 0 | 2 | 0 | 1 | 0 | 0 | 6 | 3 | 2 |
| K03802 cyanophycin synthetase [EC:6.3.2.29 6.3.2.30] | 5 | 3 | 3 | 2 | 0 | 0 | 0 | 0 | 15 | 9 | 13 | 22 |
| K03813 molybdenum transport protein [EC:2.4.2.-] | 20 | 26 | 24 | 46 | 67 | 43 | 45 | 59 | 114 | 85 | 94 | 82 |
| K03818 putative colanic acid biosynthesis acetyltransferase WcaF [EC:2.3.1.-] | 244 | 242 | 252 | 252 | 489 | 524 | 530 | 516 | 216 | 239 | 248 | 243 |
| K03819 putative colanic acid biosynthesis acetyltransferase WcaB [EC:2.3.1.-] | 91 | 83 | 84 | 100 | 137 | 169 | 118 | 138 | 19 | 23 | 17 | 23 |
| K03820 apolipoprotein N-acyltransferase [EC:2.3.1.269] | 3562 | 3430 | 3239 | 3323 | 2136 | 2127 | 2022 | 2093 | 2118 | 2050 | 1963 | 2117 |
| K03824 putative acetyltransferase [EC:2.3.1.-] | 1658 | 1765 | 1588 | 1743 | 1061 | 1021 | 1018 | 958 | 844 | 849 | 894 | 904 |
| K03825 putative acetyltransferase [EC:2.3.1.-] | 765 | 758 | 745 | 792 | 500 | 496 | 536 | 496 | 628 | 615 | 516 | 544 |
| K03826 putative acetyltransferase [EC:2.3.1.-] | 542 | 528 | 579 | 557 | 221 | 272 | 252 | 274 | 643 | 657 | 612 | 706 |
| K03827 putative acetyltransferase [EC:2.3.1.-] | 2075 | 2110 | 2133 | 2053 | 3205 | 3421 | 3404 | 3456 | 2816 | 2875 | 2777 | 2809 |
| K03828 putative acetyltransferase [EC:2.3.1.-] | 319 | 353 | 312 | 320 | 203 | 219 | 248 | 234 | 421 | 437 | 361 | 465 |
| K03829 putative acetyltransferase [EC:2.3.1.-] | 198 | 174 | 178 | 175 | 139 | 193 | 161 | 133 | 200 | 227 | 220 | 254 |
| K03830 putative acetyltransferase [EC:2.3.1.-] | 162 | 203 | 203 | 227 | 64 | 90 | 69 | 99 | 448 | 434 | 476 | 434 |
| K03924 MoxR-like ATPase [EC:3.6.3.-] | 8085 | 8174 | 8050 | 7997 | 10280 | 10356 | 10491 | 10375 | 8894 | 8407 | 8921 | 8443 |
| K03928 carboxylesterase [EC:3.1.1.1] | 61 | 73 | 69 | 55 | 8 | 16 | 4 | 11 | 90 | 97 | 81 | 95 |
| K03929 para-nitrobenzyl esterase [EC:3.1.1.-] | 1389 | 1248 | 1218 | 1308 | 723 | 748 | 743 | 767 | 1806 | 1921 | 1763 | 1807 |
| K03930 putative tributyrin esterase [EC:3.1.1.-] | 59 | 60 | 74 | 51 | 18 | 16 | 12 | 16 | 48 | 87 | 52 | 68 |
| K04068 anaerobic ribonucleoside-triphosphate reductase activating protein [EC:1.97.1.4] | 1883 | 1836 | 1744 | 1872 | 2029 | 1969 | 1939 | 1999 | 1817 | 1731 | 1769 | 1794 |
| K04069 pyruvate formate lyase activating enzyme [EC:1.97.1.4] | 3999 | 4058 | 3947 | 3956 | 4240 | 4306 | 4357 | 4217 | 5173 | 5430 | 5107 | 5439 |
| K04070 putative pyruvate formate lyase activating enzyme [EC:1.97.1.4] | 488 | 459 | 417 | 466 | 115 | 167 | 162 | 154 | 442 | 462 | 500 | 501 |
| K04087 membrane protease subunit HflC [EC:3.4.-.-] | 2837 | 2926 | 2818 | 2785 | 1755 | 1698 | 1630 | 1845 | 1619 | 1679 | 1492 | 1761 |
| K04088 membrane protease subunit HflK [EC:3.4.-.-] | 2223 | 2315 | 2242 | 2270 | 1347 | 1503 | 1307 | 1356 | 1477 | 1508 | 1526 | 1487 |
| K04766 acetoin utilization protein AcuA [EC:2.3.1.-] | 2 | 0 | 0 | 0 | 0 | 0 | 0 | 0 | 0 | 0 | 0 | 0 |
| K04844 hypothetical glycosyl hydrolase [EC:3.2.1.-] | 505 | 454 | 476 | 451 | 165 | 200 | 161 | 133 | 340 | 425 | 432 | 447 |
| K04940 opine dehydrogenase [EC:1.5.1.28] | 64 | 76 | 57 | 46 | 132 | 133 | 138 | 114 | 99 | 103 | 98 | 123 |
| K05303 O-methyltransferase [EC:2.1.1.-] | 14 | 16 | 10 | 15 | 10 | 13 | 20 | 11 | 0 | 0 | 1 | 2 |
| K05521 ADP-ribosyl-[dinitrogen reductase] hydrolase [EC:3.2.2.24] | 48 | 55 | 52 | 45 | 62 | 111 | 64 | 89 | 0 | 0 | 0 | 0 |
| K05715 2-phosphoglycerate kinase [EC:2.7.2.-] | 0 | 0 | 2 | 0 | 0 | 0 | 0 | 1 | 2 | 3 | 1 | 9 |
| K05716 cyclic 2,3-diphosphoglycerate synthetase [EC:4.6.1.-] | 0 | 0 | 0 | 0 | 0 | 0 | 0 | 0 | 4 | 14 | 6 | 9 |
| K05810 polyphenol oxidase [EC:1.10.3.-] | 2883 | 2920 | 2931 | 2928 | 2973 | 2759 | 2945 | 2829 | 3200 | 2995 | 3186 | 3137 |
| K05882 aryl-alcohol dehydrogenase (NADP+) [EC:1.1.1.91] | 0 | 0 | 0 | 1 | 0 | 0 | 0 | 1 | 0 | 0 | 0 | 0 |
| K05886 serine 3-dehydrogenase (NADP+) [EC:1.1.1.276] | 0 | 4 | 2 | 1 | 0 | 0 | 2 | 1 | 1 | 0 | 1 | 0 |
| K05888 L-gulono-1,4-lactone dehydrogenase [EC:1.1.2.-] | 2708 | 2757 | 2733 | 2740 | 1838 | 1748 | 1722 | 1763 | 894 | 788 | 829 | 738 |
| K05910 NADH peroxidase [EC:1.11.1.1] | 55 | 65 | 72 | 48 | 18 | 25 | 24 | 28 | 94 | 84 | 91 | 93 |
| K05919 superoxide reductase [EC:1.15.1.2] | 692 | 606 | 611 | 641 | 251 | 274 | 300 | 253 | 447 | 462 | 480 | 522 |
| K05922 quinone-reactive Ni/Fe-hydrogenase large subunit [EC:1.12.5.1] | 0 | 0 | 2 | 0 | 0 | 0 | 0 | 0 | 0 | 0 | 0 | 1 |
| K05970 sialate O-acetylesterase [EC:3.1.1.53] | 7102 | 7807 | 8283 | 7434 | 23314 | 23653 | 24167 | 23593 | 17506 | 16863 | 18058 | 16656 |
| K05985 ribonuclease M5 [EC:3.1.26.8] | 93 | 71 | 93 | 92 | 38 | 49 | 59 | 41 | 59 | 82 | 87 | 93 |
| K05989 alpha-L-rhamnosidase [EC:3.2.1.40] | 8454 | 8889 | 9485 | 8975 | 23443 | 22912 | 23681 | 22780 | 19571 | 18763 | 19771 | 18592 |
| K06015 N-acyl-D-amino-acid deacylase [EC:3.5.1.81] | 1069 | 1055 | 967 | 1045 | 259 | 245 | 278 | 228 | 1460 | 1673 | 1548 | 1674 |
| K06023 HPr kinase/phosphorylase [EC:2.7.11.- 2.7.4.-] | 580 | 577 | 505 | 563 | 199 | 192 | 205 | 180 | 552 | 690 | 550 | 592 |
| K06113 arabinan endo-1,5-alpha-L-arabinosidase [EC:3.2.1.99] | 2654 | 2818 | 2955 | 2839 | 9865 | 9926 | 9938 | 9754 | 7232 | 7007 | 7367 | 6918 |
| K06137 pyrroloquinoline-quinone synthase [EC:1.3.3.11] | 1346 | 1239 | 1188 | 1226 | 601 | 586 | 536 | 598 | 840 | 840 | 824 | 890 |
| K06139 PqqA peptide cyclase [EC:1.21.98.4] | 380 | 368 | 387 | 413 | 111 | 112 | 98 | 115 | 832 | 826 | 754 | 815 |
| K06221 2,5-diketo-D-gluconate reductase A [EC:1.1.1.346] | 674 | 642 | 681 | 677 | 343 | 459 | 427 | 382 | 884 | 987 | 907 | 984 |
| K06222 2,5-diketo-D-gluconate reductase B [EC:1.1.1.346] | 1712 | 1695 | 1471 | 1676 | 835 | 791 | 705 | 892 | 1092 | 1113 | 1113 | 1023 |
| K06324 spore coat protein A, manganese oxidase [EC:1.16.3.3] | 3 | 2 | 5 | 1 | 0 | 0 | 0 | 0 | 0 | 4 | 3 | 6 |
| K06379 stage II sporulation protein AB (anti-sigma F factor) [EC:2.7.11.1] | 217 | 264 | 249 | 204 | 84 | 70 | 73 | 63 | 208 | 224 | 209 | 267 |
| K06382 stage II sporulation protein E [EC:3.1.3.16] | 808 | 797 | 789 | 758 | 229 | 280 | 275 | 236 | 598 | 671 | 648 | 653 |
| K06441 ferredoxin hydrogenase gamma subunit [EC:1.12.7.2] | 31 | 44 | 33 | 29 | 45 | 63 | 65 | 49 | 52 | 50 | 53 | 44 |
| K06607 myo-inositol catabolism protein IolS [EC:1.1.1.-] | 3 | 0 | 2 | 1 | 5 | 2 | 1 | 5 | 10 | 26 | 7 | 9 |
| K06864 pyridinium-3,5-biscarboxylic acid mononucleotide sulfurtransferase [EC:4.4.1.37] | 381 | 357 | 372 | 331 | 89 | 97 | 113 | 108 | 357 | 404 | 431 | 437 |
| K06898 pyridinium-3,5-biscarboxylic acid mononucleotide synthase [EC:2.5.1.143] | 422 | 452 | 372 | 390 | 148 | 142 | 151 | 139 | 388 | 465 | 455 | 420 |
| K06937 7,8-dihydro-6-hydroxymethylpterin dimethyltransferase [EC:2.1.1.-] | 76 | 76 | 59 | 76 | 49 | 38 | 42 | 48 | 155 | 191 | 166 | 198 |
| K06988 8-hydroxy-5-deazaflavin:NADPH oxidoreductase [EC:1.5.1.40] | 3 | 2 | 2 | 5 | 5 | 2 | 11 | 5 | 0 | 8 | 5 | 6 |
| K07037 cyclic-di-AMP phosphodiesterase PgpH [EC:3.1.4.-] | 1483 | 1586 | 1688 | 1598 | 4884 | 4914 | 4959 | 4555 | 3560 | 3388 | 3684 | 3574 |
| K07053 3',5'-nucleoside bisphosphate phosphatase [EC:3.1.3.97] | 2618 | 2540 | 2427 | 2630 | 1148 | 1195 | 1147 | 1211 | 2052 | 2180 | 2044 | 2116 |
| K07067 diadenylate cyclase [EC:2.7.7.85] | 3 | 11 | 3 | 1 | 2 | 0 | 0 | 0 | 2 | 9 | 7 | 7 |
| K07094 putative glycerol-1-phosphate prenyltransferase [EC:2.5.1.-] | 2 | 0 | 0 | 0 | 0 | 0 | 0 | 0 | 0 | 0 | 0 | 0 |
| K07107 acyl-CoA thioester hydrolase [EC:3.1.2.-] | 5006 | 5099 | 4991 | 5009 | 4382 | 4330 | 4180 | 4439 | 3229 | 3235 | 3166 | 3101 |
| K07116 acyl-homoserine-lactone acylase [EC:3.5.1.97] | 6946 | 6846 | 6694 | 6684 | 3790 | 3704 | 3469 | 3880 | 2128 | 2102 | 1932 | 2052 |
| K07132 ATP-dependent target DNA activator [EC:3.6.1.3] | 41 | 44 | 44 | 46 | 86 | 92 | 70 | 71 | 90 | 96 | 86 | 111 |
| K07147 methionine sulfoxide reductase catalytic subunit [EC:1.8.-.-] | 2309 | 2305 | 2279 | 2321 | 1399 | 1487 | 1383 | 1508 | 1380 | 1394 | 1444 | 1535 |
| K07223 porphyrinogen peroxidase [EC:1.11.1.-] | 2214 | 2286 | 2287 | 2207 | 2640 | 2633 | 2674 | 2761 | 2402 | 2182 | 2260 | 2279 |
| K07229 ferric-chelate reductase (NADPH) [EC:1.16.1.9] | 471 | 491 | 421 | 531 | 338 | 393 | 380 | 372 | 537 | 684 | 632 | 679 |
| K07271 lipopolysaccharide cholinephosphotransferase [EC:2.7.8.-] | 1789 | 1805 | 2012 | 1893 | 3883 | 3959 | 3993 | 3810 | 3262 | 3129 | 3477 | 3214 |
| K07302 isoquinoline 1-oxidoreductase subunit alpha [EC:1.3.99.16] | 1157 | 1148 | 1094 | 1097 | 609 | 593 | 569 | 615 | 388 | 299 | 346 | 354 |
| K07303 isoquinoline 1-oxidoreductase subunit beta [EC:1.3.99.16] | 6365 | 6231 | 5595 | 5976 | 3307 | 3043 | 2916 | 3420 | 2050 | 1745 | 1688 | 1832 |
| K07304 peptide-methionine (S)-S-oxide reductase [EC:1.8.4.11] | 1547 | 1401 | 1379 | 1397 | 918 | 850 | 828 | 929 | 1025 | 1005 | 1000 | 969 |
| K07305 peptide-methionine (R)-S-oxide reductase [EC:1.8.4.12] | 828 | 831 | 789 | 764 | 505 | 499 | 512 | 456 | 441 | 495 | 424 | 482 |
| K07313 serine/threonine protein phosphatase 1 [EC:3.1.3.16] | 529 | 509 | 488 | 530 | 444 | 422 | 401 | 417 | 562 | 682 | 615 | 646 |
| K07314 serine/threonine protein phosphatase 2 [EC:3.1.3.16] | 5 | 0 | 2 | 3 | 2 | 6 | 8 | 7 | 41 | 52 | 35 | 45 |
| K07336 PKHD-type hydroxylase [EC:1.14.11.-] | 1480 | 1352 | 1441 | 1433 | 694 | 649 | 727 | 702 | 885 | 887 | 811 | 893 |
| K07393 glutathionyl-hydroquinone reductase [EC:1.8.5.7] | 2078 | 2050 | 2029 | 1989 | 1375 | 1436 | 1384 | 1305 | 1169 | 1227 | 1174 | 1200 |
| K07444 putative N6-adenine-specific DNA methylase [EC:2.1.1.-] | 1328 | 1508 | 1730 | 1459 | 3244 | 3254 | 3212 | 3313 | 2748 | 2702 | 2790 | 2707 |
| K07447 putative holliday junction resolvase [EC:3.1.-.-] | 1589 | 1502 | 1547 | 1521 | 1526 | 1582 | 1562 | 1601 | 1512 | 1428 | 1532 | 1517 |
| K07469 aldehyde oxidoreductase [EC:1.2.99.7] | 137 | 137 | 121 | 151 | 100 | 102 | 120 | 92 | 306 | 315 | 264 | 294 |
| K07503 endonuclease [EC:3.1.-.-] | 2 | 8 | 3 | 5 | 3 | 0 | 2 | 3 | 4 | 10 | 12 | 16 |
| K07557 archaeosine synthase [EC:2.6.1.97] | 0 | 0 | 2 | 1 | 3 | 0 | 0 | 0 | 0 | 13 | 2 | 9 |
| K07588 LAO/AO transport system kinase [EC:2.7.-.-] | 856 | 896 | 866 | 823 | 2535 | 2576 | 2512 | 2484 | 2025 | 1870 | 1868 | 1889 |
| K07749 formyl-CoA transferase [EC:2.8.3.16] | 1174 | 1233 | 1188 | 1137 | 636 | 691 | 687 | 702 | 491 | 478 | 418 | 457 |
| K07755 arsenite methyltransferase [EC:2.1.1.137] | 12 | 23 | 22 | 21 | 8 | 14 | 13 | 16 | 16 | 39 | 25 | 28 |
| K07757 sugar-phosphatase [EC:3.1.3.23] | 410 | 481 | 442 | 446 | 315 | 380 | 351 | 315 | 675 | 727 | 637 | 694 |
| K08282 non-specific serine/threonine protein kinase [EC:2.7.11.1] | 23 | 40 | 27 | 28 | 14 | 16 | 8 | 12 | 31 | 31 | 20 | 48 |
| K08286 protein-serine/threonine kinase [EC:2.7.11.-] | 0 | 2 | 0 | 0 | 2 | 0 | 0 | 0 | 0 | 0 | 0 | 0 |
| K08296 phosphohistidine phosphatase [EC:3.1.3.-] | 991 | 911 | 847 | 946 | 517 | 507 | 500 | 611 | 624 | 550 | 562 | 595 |
| K08297 crotonobetainyl-CoA dehydrogenase [EC:1.3.8.13] | 261 | 276 | 262 | 272 | 424 | 433 | 410 | 411 | 82 | 89 | 60 | 106 |
| K08298 L-carnitine CoA-transferase [EC:2.8.3.21] | 237 | 280 | 269 | 336 | 404 | 418 | 469 | 401 | 159 | 127 | 127 | 154 |
| K08299 crotonobetainyl-CoA hydratase [EC:4.2.1.149] | 193 | 160 | 183 | 180 | 209 | 288 | 275 | 247 | 68 | 72 | 41 | 60 |
| K08313 fructose-6-phosphate aldolase 1 [EC:4.1.2.-] | 3 | 5 | 3 | 6 | 19 | 10 | 16 | 13 | 31 | 43 | 44 | 44 |
| K08314 fructose-6-phosphate aldolase 2 [EC:4.1.2.-] | 460 | 437 | 458 | 428 | 321 | 405 | 310 | 343 | 548 | 597 | 572 | 612 |
| K08317 uncharacterized oxidoreductase [EC:1.1.-.-] | 1006 | 971 | 837 | 933 | 819 | 823 | 750 | 771 | 770 | 835 | 810 | 865 |
| K08319 L-threonate 2-dehydrogenase [EC:1.1.1.411] | 283 | 334 | 335 | 351 | 91 | 97 | 80 | 83 | 740 | 694 | 677 | 711 |
| K08320 (d)CTP diphosphatase [EC:3.6.1.65] | 261 | 230 | 213 | 225 | 176 | 152 | 162 | 142 | 283 | 328 | 244 | 348 |
| K08961 chondroitin-sulfate-ABC endolyase/exolyase [EC:4.2.2.20 4.2.2.21] | 2667 | 3073 | 3195 | 2990 | 11208 | 11056 | 11143 | 10593 | 8386 | 8326 | 8746 | 7861 |
| K09001 anhydro-N-acetylmuramic acid kinase [EC:2.7.1.170] | 2615 | 2434 | 2385 | 2605 | 1548 | 1526 | 1572 | 1526 | 1496 | 1564 | 1512 | 1589 |

| | | | | | | | | | | | | |
|---|---|---|---|---|---|---|---|---|---|---|---|---|
| K09022 2-iminobutanoate/2-iminopropanoate deaminase [EC:3.5.99.10] | 895 | 1002 | 1112 | 1053 | 1081 | 1150 | 1163 | 1061 | 1491 | 1648 | 1644 | 1632 |
| K09121 pyridinium-3,5-bisthiocarboxylic acid mononucleotide nickel chelatase [EC:4.99.1.12] | 481 | 478 | 503 | 481 | 138 | 143 | 165 | 173 | 593 | 605 | 680 | 654 |
| K09773 [pyruvate, water dikinase]-phosphate phosphotransferase / [pyruvate, water dikinase] kinase [EC:2.7.4.28 2.7.11.33] | 1691 | 1659 | 1670 | 1703 | 1027 | 1026 | 1025 | 1059 | 1182 | 1208 | 1131 | 1132 |
| K09809 CDP-glycerol glycerophosphotransferase [EC:2.7.8.12] | 324 | 326 | 315 | 306 | 57 | 68 | 61 | 74 | 403 | 485 | 479 | 526 |
| K10231 kojibiose phosphorylase [EC:2.4.1.230] | 357 | 395 | 400 | 382 | 90 | 81 | 66 | 76 | 229 | 260 | 270 | 251 |
| K10253 DOPA 4,5-dioxygenase [EC:1.14.99.-] | 629 | 646 | 563 | 607 | 344 | 363 | 337 | 322 | 183 | 174 | 180 | 179 |
| K10254 oleate hydratase [EC:4.2.1.53] | 1406 | 1432 | 1438 | 1490 | 284 | 357 | 339 | 264 | 1574 | 1767 | 1599 | 1642 |
| K10530 L-lactate oxidase [EC:1.1.3.2] | 6 | 9 | 5 | 7 | 12 | 16 | 7 | 6 | 92 | 76 | 71 | 97 |
| K10531 L-ornithine N5-monooxygenase [EC:1.14.13.195 1.14.13.196] | 1533 | 1560 | 1452 | 1460 | 848 | 810 | 731 | 833 | 483 | 478 | 412 | 456 |
| K10670 glycine/sarcosine/betaine reductase complex component A [EC:1.21.4.2 1.21.4.3 1.21.4.4] | 50 | 38 | 37 | 36 | 23 | 27 | 19 | 19 | 23 | 50 | 11 | 33 |
| K10671 glycine reductase complex component B subunit alpha and beta [EC:1.21.4.2] | 88 | 96 | 108 | 118 | 42 | 44 | 38 | 34 | 35 | 81 | 48 | 38 |
| K10672 glycine reductase complex component B subunit gamma [EC:1.21.4.2] | 143 | 153 | 146 | 150 | 57 | 36 | 62 | 25 | 73 | 81 | 56 | 74 |
| K10677 inulin fructotransferase (DFA-I-forming) [EC:4.2.2.17] | 0 | 0 | 0 | 0 | 0 | 1 | 0 | 0 | 1 | 0 | 0 | 1 |
| K10708 fructoselysine 6-phosphate deglycase [EC:3.5.-.-] | 292 | 371 | 319 | 353 | 366 | 394 | 379 | 378 | 139 | 132 | 122 | 122 |
| K10709 fructoselysine 3-epimerase [EC:5.1.3.41] | 223 | 196 | 227 | 227 | 319 | 296 | 294 | 282 | 15 | 25 | 23 | 21 |
| K10710 fructoselysine 6-kinase [EC:2.7.1.218] | 255 | 266 | 259 | 299 | 299 | 307 | 292 | 283 | 113 | 86 | 102 | 90 |
| K10819 histidine kinase [EC:2.7.13.3] | 7 | 2 | 5 | 12 | 5 | 7 | 10 | 9 | 2 | 1 | 1 | 0 |
| K11065 thiol peroxidase, atypical 2-Cys peroxiredoxin [EC:1.11.1.15] | 558 | 650 | 699 | 657 | 1192 | 1159 | 1213 | 1180 | 1020 | 1136 | 1144 | 1097 |
| K11161 retinol dehydrogenase 13 [EC:1.1.1.300] | 0 | 0 | 0 | 0 | 0 | 1 | 0 | 0 | 1 | 1 | 0 | 0 |
| K11169 dehydrogenase/reductase SDR family member 13 [EC:1.1.-.-] | 0 | 0 | 0 | 1 | 0 | 0 | 0 | 0 | 0 | 0 | 0 | 0 |
| K11206 deaminated glutathione amidase [EC:3.5.1.128] | 2079 | 2083 | 1942 | 2014 | 1278 | 1159 | 1171 | 1176 | 1291 | 1347 | 1301 | 1204 |
| K11209 GSH-dependent disulfide-bond oxidoreductase [EC:1.8.4.-] | 2309 | 2186 | 2276 | 2329 | 1322 | 1247 | 1215 | 1210 | 1819 | 1932 | 1765 | 1850 |
| K11750 esterase FrsA [EC:3.1.-.-] | 685 | 748 | 678 | 752 | 574 | 593 | 532 | 520 | 989 | 1042 | 968 | 1070 |
| K11933 NADH oxidoreductase Hcr [EC:1.-.-.-] | 574 | 563 | 536 | 576 | 420 | 375 | 421 | 381 | 794 | 806 | 751 | 746 |
| K11938 HMP-PP phosphatase [EC:3.6.1.-] | 548 | 526 | 522 | 608 | 397 | 430 | 452 | 392 | 791 | 864 | 807 | 843 |
| K11941 glucans biosynthesis protein C [EC:2.1.-.-] | 524 | 530 | 557 | 555 | 386 | 444 | 415 | 361 | 845 | 772 | 759 | 828 |
| K11942 isobutyryl-CoA mutase [EC:5.4.99.13] | 8 | 3 | 2 | 4 | 3 | 2 | 5 | 2 | 0 | 0 | 0 | 0 |
| K12136 hydrogenase-4 component A [EC:1.-.-.-] | 141 | 150 | 125 | 141 | 163 | 213 | 209 | 202 | 4 | 13 | 4 | 15 |
| K12137 hydrogenase-4 component B [EC:1.-.-.-] | 526 | 528 | 521 | 586 | 755 | 835 | 714 | 667 | 185 | 233 | 193 | 208 |
| K12138 hydrogenase-4 component C [EC:1.-.-.-] | 211 | 202 | 207 | 191 | 329 | 302 | 361 | 303 | 3 | 1 | 2 | 2 |
| K12139 hydrogenase-4 component D [EC:1.-.-.-] | 325 | 292 | 273 | 309 | 430 | 438 | 501 | 442 | 4 | 3 | 0 | 4 |
| K12140 hydrogenase-4 component E [EC:1.-.-.-] | 139 | 120 | 146 | 165 | 162 | 223 | 182 | 175 | 39 | 37 | 50 | 51 |
| K12141 hydrogenase-4 component F [EC:1.-.-.-] | 360 | 361 | 315 | 362 | 406 | 464 | 499 | 454 | 92 | 90 | 99 | 101 |
| K12142 hydrogenase-4 component G [EC:1.-.-.-] | 380 | 385 | 308 | 369 | 520 | 650 | 511 | 477 | 4 | 7 | 8 | 7 |
| K12144 hydrogenase-4 component I [EC:1.-.-.-] | 175 | 170 | 138 | 145 | 210 | 254 | 231 | 220 | 4 | 1 | 0 | 0 |
| K12145 hydrogenase-4 component J [EC:1.-.-.-] | 90 | 97 | 67 | 87 | 130 | 157 | 115 | 112 | 1 | 0 | 0 | 1 |
| K12152 phosphatase NudJ [EC:3.6.1.-] | 235 | 221 | 250 | 254 | 206 | 198 | 183 | 174 | 365 | 385 | 342 | 459 |
| K12267 peptide methionine sulfoxide reductase msrA/msrB [EC:1.8.4.11 1.8.4.12] | 1258 | 1155 | 1219 | 1256 | 1974 | 1920 | 1988 | 2000 | 1989 | 2016 | 2176 | 2045 |
| K12374 arylsulfatase D/F/H [EC:3.1.6.-] | 0 | 1 | 0 | 0 | 0 | 0 | 0 | 0 | 0 | 1 | 0 | 0 |
| K12375 arylsulfatase I/J [EC:3.1.6.-] | 0 | 0 | 0 | 0 | 0 | 2 | 0 | 0 | 0 | 0 | 0 | 0 |
| K12376 arylsulfatase K [EC:3.1.6.-] | 0 | 0 | 0 | 0 | 1 | 2 | 4 | 2 | 2 | 0 | 1 | 0 |
| K12410 NAD-dependent deacetylase [EC:3.5.1.-] | 3727 | 3641 | 3516 | 3529 | 3251 | 3230 | 3116 | 3200 | 3085 | 3024 | 3058 | 3239 |
| K12508 feruloyl-CoA synthase [EC:6.2.1.34] | 3 | 0 | 0 | 0 | 0 | 0 | 0 | 2 | 0 | 0 | 0 | 0 |
| K12700 non-specific riboncleoside hydrolase [EC:3.2.-.-] | 621 | 690 | 580 | 594 | 396 | 469 | 464 | 424 | 797 | 927 | 829 | 878 |
| K12944 nucleoside triphosphatase [EC:3.6.1.-] | 268 | 276 | 242 | 270 | 203 | 184 | 158 | 171 | 367 | 356 | 363 | 375 |
| K12945 GDP-mannose pyrophosphatase NudK [EC:3.6.1.-] | 149 | 132 | 118 | 106 | 127 | 203 | 183 | 152 | 52 | 59 | 57 | 58 |
| K12950 manganese/zinc-transporting P-type ATPase C [EC:7.2.2.-] | 3 | 0 | 4 | 1 | 3 | 0 | 2 | 0 | 2 | 0 | 3 | 1 |
| K12952 cation-transporting P-type ATPase E [EC:7.2.2.-] | 474 | 470 | 474 | 522 | 191 | 248 | 201 | 203 | 711 | 815 | 734 | 799 |
| K12953 cation-transporting P-type ATPase F [EC:7.2.2.-] | 1 | 1 | 0 | 1 | 0 | 0 | 0 | 0 | 0 | 0 | 0 | 1 |
| K12954 cation-transporting P-type ATPase G [EC:7.2.2.-] | 0 | 0 | 0 | 0 | 0 | 0 | 0 | 0 | 1 | 0 | 2 | 1 |
| K12957 uncharacterized zinc-type alcohol dehydrogenase-like protein [EC:1.-.-.-] | 704 | 720 | 665 | 731 | 453 | 494 | 505 | 407 | 1040 | 939 | 951 | 1015 |
| K13043 N-succinyl-L-ornithine transcarbamylase [EC:2.1.3.11] | 643 | 716 | 765 | 623 | 2214 | 2179 | 2297 | 2300 | 1597 | 1503 | 1646 | 1513 |
| K13059 N-acetylhexosamine 1-kinase [EC:2.7.1.162] | 3 | 2 | 2 | 0 | 0 | 1 | 2 | 0 | 9 | 6 | 3 | 15 |
| K13069 diguanylate cyclase [EC:2.7.7.65] | 583 | 610 | 559 | 540 | 373 | 454 | 425 | 460 | 875 | 868 | 928 | 1014 |
| K13243 c-di-GMP-specific phosphodiesterase [EC:3.1.4.52] | 13 | 10 | 6 | 9 | 21 | 36 | 31 | 36 | 106 | 142 | 121 | 131 |
| K13244 c-di-GMP-specific phosphodiesterase [EC:3.1.4.52] | 1 | 3 | 2 | 0 | 12 | 16 | 17 | 10 | 38 | 57 | 51 | 44 |
| K13252 putrescine carbamoyltransferase [EC:2.1.3.6] | 32 | 23 | 20 | 36 | 13 | 11 | 8 | 15 | 40 | 50 | 60 | 42 |
| K13278 60kDa lysophospholipase [EC:3.1.1.5 3.1.1.47 3.5.1.1] | 0 | 0 | 0 | 0 | 0 | 0 | 0 | 0 | 0 | 1 | 0 | 0 |
| K13281 UV DNA damage endonuclease [EC:3.-.-.-] | 10 | 22 | 7 | 11 | 5 | 5 | 2 | 3 | 39 | 20 | 37 | 31 |
| K13292 phosphatidylglycerol---prolipoprotein diacylglyceryl transferase [EC:2.5.1.145] | 2433 | 2358 | 2257 | 2205 | 1350 | 1376 | 1290 | 1310 | 1667 | 1664 | 1780 | 1825 |
| K13574 uncharacterized oxidoreductase [EC:1.1.1.-] | 845 | 843 | 818 | 982 | 828 | 892 | 858 | 810 | 971 | 932 | 957 | 950 |
| K13670 putative glycosyltransferase [EC:2.4.-.-] | 2 | 0 | 0 | 0 | 0 | 0 | 0 | 0 | 0 | 0 | 0 | 0 |
| K13714 bifunctional autolysin [EC:3.5.1.28 3.2.1.96] | 0 | 0 | 0 | 0 | 0 | 0 | 0 | 0 | 3 | 0 | 0 | 0 |
| K13727 phenolic acid decarboxylase [EC:4.1.1.-] | 332 | 315 | 305 | 329 | 257 | 232 | 252 | 232 | 442 | 449 | 405 | 456 |
| K13930 triphosphoribosyl-dephospho-CoA synthase [EC:2.4.2.52] | 987 | 1034 | 843 | 958 | 652 | 680 | 577 | 608 | 726 | 777 | 743 | 769 |
| K13932 malonate decarboxylase beta subunit [EC:4.1.1.87] | 1760 | 1736 | 1443 | 1601 | 986 | 940 | 967 | 932 | 968 | 1014 | 925 | 968 |
| K13933 malonate decarboxylase gamma subunit [EC:4.1.1.87] | 2006 | 1959 | 1751 | 1856 | 1022 | 1142 | 1156 | 1139 | 1008 | 969 | 881 | 1052 |
| K13934 phosphoribosyl-dephospho-CoA transferase [EC:2.7.7.66] | 1238 | 1289 | 1117 | 1231 | 623 | 647 | 664 | 750 | 724 | 719 | 647 | 761 |
| K13935 malonate decarboxylase epsilon subunit [EC:2.3.1.39] | 1411 | 1301 | 1106 | 1296 | 783 | 752 | 783 | 779 | 895 | 901 | 865 | 939 |
| K13979 uncharacterized zinc-type alcohol dehydrogenase-like protein [EC:1.-.-.-] | 2377 | 2393 | 2160 | 2239 | 1450 | 1561 | 1395 | 1448 | 1446 | 1495 | 1328 | 1401 |
| K14189 uncharacterized oxidoreductase [EC:1.-.-.-] | 134 | 126 | 139 | 143 | 202 | 195 | 205 | 187 | 0 | 2 | 0 | 0 |
| K14351 arylacetamide deacetylase-like 3/4 [EC:3.1.1.-] | 0 | 0 | 0 | 1 | 0 | 0 | 0 | 0 | 0 | 0 | 0 | 0 |
| K14587 protein sgcE [EC:5.1.3.-] | 1 | 2 | 0 | 0 | 5 | 1 | 1 | 4 | 3 | 6 | 3 | 1 |
| K14659 chitooligosaccharide deacetylase [EC:3.5.1.-] | 1 | 0 | 0 | 0 | 0 | 0 | 0 | 0 | 0 | 0 | 0 | 0 |
| K14744 prophage endopeptidase [EC:3.4.-.-] | 470 | 487 | 519 | 492 | 537 | 560 | 497 | 462 | 154 | 130 | 135 | 112 |
| K15509 sulfopropanediol 3-dehydrogenase [EC:1.1.1.308] | 2 | 2 | 2 | 1 | 0 | 0 | 0 | 0 | 8 | 6 | 1 | 5 |
| K15515 sulfoacetaldehyde dehydrogenase [EC:1.2.1.81] | 47 | 33 | 60 | 53 | 6 | 11 | 3 | 6 | 15 | 21 | 12 | 20 |
| K15524 mannosylglycerate hydrolase [EC:3.2.1.170] | 33 | 33 | 23 | 22 | 40 | 38 | 56 | 32 | 22 | 54 | 32 | 39 |
| K15527 cysteate synthase [EC:2.5.1.76] | 3 | 6 | 6 | 1 | 14 | 8 | 3 | 10 | 0 | 2 | 0 | 0 |
| K15531 oligosaccharide reducing-end xylanase [EC:3.2.1.156] | 116 | 120 | 119 | 120 | 47 | 78 | 71 | 89 | 291 | 304 | 289 | 286 |
| K15532 unsaturated rhamnogalacturonyl hydrolase [EC:3.2.1.172] | 4181 | 4169 | 4269 | 4227 | 5527 | 5703 | 5706 | 5562 | 5684 | 5877 | 5818 | 5777 |
| K15533 1,3-beta-galactosyl-N-acetylhexosamine phosphorylase [EC:2.4.1.211] | 1281 | 1343 | 1348 | 1305 | 273 | 262 | 263 | 227 | 1080 | 1238 | 1138 | 1138 |
| K15534 beta-D-galactosyl-(1->4)-L-rhamnose phosphorylase [EC:2.4.1.247] | 175 | 164 | 167 | 156 | 22 | 36 | 27 | 12 | 155 | 134 | 160 | 181 |
| K15538 glycoprotein endo-alpha-1,2-mannosidase [EC:3.2.1.130] | 0 | 0 | 1 | 2 | 3 | 2 | 3 | 1 | 0 | 1 | 1 | 0 |
| K15915 undecaprenyl phosphate N,N'-diacetylbacillosamine 1-phosphate transferase [EC:2.7.8.36] | 10 | 12 | 14 | 11 | 3 | 3 | 9 | 3 | 7 | 7 | 2 | 6 |
| K15922 sulfoquinovosidase [EC:3.2.1.199] | 31 | 23 | 16 | 31 | 4 | 8 | 7 | 9 | 81 | 53 | 61 | 71 |
| K15924 glucuronoarabinoxylan endo-1,4-beta-xylanase [EC:3.2.1.136] | 0 | 0 | 2 | 0 | 0 | 0 | 0 | 3 | 0 | 2 | 0 | 0 |
| K16212 4-O-beta-D-mannosyl-D-glucose phosphorylase [EC:2.4.1.281] | 996 | 1052 | 1141 | 1055 | 2836 | 2799 | 2814 | 2735 | 2303 | 2132 | 2201 | 2223 |
| K16213 cellobiose epimerase [EC:5.1.3.11] | 263 | 288 | 255 | 268 | 202 | 226 | 236 | 205 | 181 | 223 | 220 | 189 |
| K16704 dTDP-4-amino-4,6-dideoxy-D-galactose acyltransferase [EC:2.3.1.210] | 449 | 425 | 412 | 505 | 294 | 351 | 294 | 309 | 559 | 582 | 576 | 587 |
| K16868 tellurite methyltransferase [EC:2.1.1.265] | 407 | 393 | 403 | 368 | 241 | 232 | 226 | 244 | 553 | 592 | 569 | 585 |
| K16869 octanoyl-[GcvH]:protein N-octanoyltransferase [EC:2.3.1.204] | 40 | 47 | 48 | 35 | 8 | 6 | 8 | 4 | 55 | 78 | 47 | 54 |
| K16872 beta-ketodecanoyl-[acyl-carrier-protein] synthase [EC:2.3.1.207] | 1575 | 1648 | 1593 | 1633 | 986 | 866 | 846 | 918 | 541 | 477 | 479 | 445 |
| K17064 D-2-hydroxyacid dehydrogenase (NADP+) [EC:1.1.1.272] | 2 | 2 | 1 | 0 | 3 | 1 | 0 | 0 | 0 | 0 | 0 | 0 |
| K17624 endo-alpha-N-acetylgalactosaminidase [EC:3.2.1.97] | 190 | 246 | 235 | 237 | 45 | 30 | 27 | 31 | 193 | 265 | 224 | 272 |
| K17735 carnitine 3-dehydrogenase [EC:1.1.1.108] | 1264 | 1286 | 1051 | 1161 | 673 | 593 | 609 | 650 | 383 | 366 | 343 | 383 |
| K17758 ADP-dependent NAD(P)H-hydrate dehydratase [EC:4.2.1.136] | 4835 | 4787 | 4670 | 4685 | 5119 | 5273 | 5251 | 5197 | 4768 | 4831 | 4810 | 4949 |
| K17759 NAD(P)H-hydrate epimerase [EC:5.1.99.6] | 3 | 5 | 5 | 12 | 7 | 16 | 14 | 10 | 13 | 8 | 15 | 12 |
| K17760 quinohemoprotein ethanol dehydrogenase [EC:1.1.9.1] | 1 | 0 | 0 | 0 | 0 | 0 | 0 | 0 | 0 | 0 | 0 | 0 |
| K17810 D-aspartate ligase [EC:6.3.1.12] | 38 | 16 | 19 | 26 | 17 | 28 | 12 | 21 | 41 | 26 | 30 | 31 |
| K17869 NADH oxidase (H2O-forming) [EC:1.6.3.4] | 0 | 0 | 2 | 0 | 2 | 0 | 0 | 0 | 0 | 0 | 0 | 0 |
| K17870 NADH oxidase (H2O2-forming) [EC:1.6.3.3] | 3 | 2 | 2 | 5 | 0 | 0 | 3 | 0 | 3 | 13 | 7 | 2 |
| K17883 mycothione reductase [EC:1.8.1.15] | 1 | 0 | 1 | 1 | 1 | 0 | 0 | 0 | 0 | 0 | 0 | 0 |
| K17884 archaetidylinositol phosphate synthase [EC:2.7.8.39] | 0 | 0 | 0 | 0 | 0 | 2 | 0 | 0 | 5 | 8 | 0 | 3 |
| K17948 N-acetylneuraminate epimerase [EC:5.1.3.24] | 0 | 10 | 5 | 7 | 9 | 23 | 8 | 10 | 63 | 92 | 62 | 82 |
| K17992 NADP-reducing hydrogenase subunit HndB [EC:1.12.1.3] | 232 | 181 | 208 | 213 | 55 | 37 | 42 | 56 | 144 | 158 | 133 | 152 |
| K17998 iron-hydrogenase subunit beta [EC:1.12.1.4] | 0 | 0 | 0 | 0 | 0 | 0 | 0 | 1 | 0 | 0 | 0 | 0 |
| K18006 [NiFe] hydrogenase diaphorase moiety small subunit [EC:1.12.1.2] | 0 | 0 | 0 | 0 | 0 | 1 | 0 | 0 | 0 | 0 | 0 | 0 |
| K18008 [NiFe] hydrogenase small subunit [EC:1.12.2.1] | 12 | 9 | 7 | 8 | 3 | 0 | 2 | 7 | 4 | 21 | 4 | 15 |
| K18016 membrane-bound hydrogenase subunit alpha [EC:1.12.7.2] | 2 | 0 | 0 | 0 | 0 | 0 | 0 | 0 | 0 | 0 | 0 | 0 |
| K18197 rhamnogalacturonan endolyase [EC:4.2.2.23] | 409 | 430 | 437 | 418 | 559 | 633 | 655 | 618 | 40 | 87 | 56 | 95 |
| K18198 rhamnogalacturonan exolyase [EC:4.2.2.24] | 2 | 0 | 1 | 0 | 0 | 0 | 0 | 0 | 0 | 0 | 0 | 0 |
| K18200 peptidylamidoglycolate lyase [EC:4.3.2.5] | 0 | 0 | 0 | 1 | 0 | 0 | 0 | 0 | 0 | 0 | 0 | 0 |
| K18205 non-reducing end beta-L-arabinofuranosidase [EC:3.2.1.185] | 16 | 23 | 18 | 13 | 3 | 11 | 3 | 12 | 13 | 38 | 24 | 19 |
| K18206 beta-L-arabinobiosidase [EC:3.2.1.187] | 4 | 5 | 2 | 4 | 9 | 9 | 2 | 3 | 31 | 34 | 27 | 61 |
| K18237 ribose 1,5-bisphosphate isomerase [EC:5.3.1.29] | 96 | 96 | 108 | 97 | 20 | 17 | 26 | 17 | 47 | 82 | 74 | 68 |
| K18244 acyl-CoA dehydrogenase [EC:1.3.99.-] | 20 | 10 | 26 | 10 | 2 | 3 | 3 | 1 | 64 | 54 | 53 | 50 |
| K18313 succinyl-CoA---D-citramalate CoA-transferase [EC:2.8.3.20] | 1 | 0 | 1 | 0 | 0 | 0 | 0 | 0 | 0 | 0 | 0 | 0 |
| K18330 NADP-reducing hydrogenase subunit HndA [EC:1.12.1.3] | 340 | 374 | 464 | 401 | 910 | 859 | 874 | 962 | 802 | 737 | 705 | 741 |
| K18331 NADP-reducing hydrogenase subunit HndC [EC:1.12.1.3] | 1805 | 1795 | 1997 | 1877 | 3768 | 3833 | 3955 | 3649 | 3432 | 3426 | 3621 | 3376 |
| K18332 NADP-reducing hydrogenase subunit HndD [EC:1.12.1.3] | 2259 | 2162 | 2308 | 2232 | 3601 | 3634 | 3721 | 3530 | 3414 | 3548 | 3414 | 3424 |

| | | | | | | | | | | | | |
|---|---|---|---|---|---|---|---|---|---|---|---|---|
| K18367 CoA-dependent NAD(P)H sulfur oxidoreductase [EC:1.8.1.18] | 2 | 3 | 0 | 1 | 2 | 1 | 4 | 3 | 1 | 1 | 2 | 1 |
| K18425 3-hydroxy-D-aspartate aldolase [EC:4.1.3.41] | 0 | 1 | 0 | 1 | 0 | 0 | 0 | 0 | 12 | 1 | 3 | 1 |
| K18427 4-hydroxyphenylacetate decarboxylase large subunit [EC:4.1.1.83] | 75 | 76 | 56 | 58 | 0 | 0 | 0 | 0 | 127 | 141 | 136 | 128 |
| K18428 4-hydroxyphenylacetate decarboxylase small subunit [EC:4.1.1.83] | 2 | 1 | 1 | 3 | 0 | 1 | 0 | 2 | 10 | 7 | 4 | 10 |
| K18446 triphosphatase [EC:3.6.1.25] | 883 | 838 | 868 | 859 | 640 | 624 | 602 | 603 | 1147 | 1168 | 1140 | 1228 |
| K18456 8-oxoguanine deaminase [EC:3.5.4.32] | 2422 | 2261 | 2266 | 2211 | 1384 | 1171 | 1214 | 1393 | 770 | 669 | 713 | 767 |
| K18457 L-proline amide hydrolase [EC:3.5.1.101] | 0 | 0 | 0 | 0 | 0 | 0 | 0 | 1 | 0 | 0 | 0 | 0 |
| K18459 guanidinopropionase [EC:3.5.3.17] | 1435 | 1475 | 1350 | 1441 | 791 | 781 | 794 | 882 | 523 | 424 | 417 | 478 |
| K18478 sulfofructose kinase [EC:2.7.1.184] | 239 | 207 | 197 | 216 | 302 | 351 | 379 | 352 | 87 | 96 | 84 | 93 |
| K18479 sulfoquinovose isomerase [EC:5.3.1.31] | 0 | 0 | 0 | 3 | 0 | 2 | 0 | 1 | 2 | 0 | 0 | 4 |
| K18540 (R)-amidase [EC:3.5.1.100] | 998 | 1001 | 945 | 1018 | 570 | 550 | 527 | 573 | 341 | 310 | 253 | 318 |
| K18561 FAD-dependent fumarate reductase [EC:1.3.8.-] | 0 | 0 | 0 | 0 | 1 | 0 | 0 | 0 | 0 | 0 | 0 | 0 |
| K18581 unsaturated chondroitin disaccharide hydrolase [EC:3.2.1.180] | 4075 | 3950 | 4098 | 4085 | 4389 | 5040 | 4713 | 4714 | 4703 | 4787 | 4786 | 4757 |
| K18650 exo-poly-alpha-galacturonosidase [EC:3.2.1.82] | 3 | 2 | 3 | 2 | 19 | 16 | 13 | 13 | 0 | 0 | 1 | 2 |
| K18672 diadenylate cyclase [EC:2.7.7.85] | 482 | 444 | 477 | 471 | 131 | 148 | 148 | 130 | 470 | 495 | 477 | 512 |
| K18673 beta-glucoside kinase [EC:2.7.1.85] | 373 | 359 | 375 | 372 | 101 | 85 | 98 | 136 | 708 | 822 | 736 | 731 |
| K18678 phytol kinase [EC:2.7.1.182] | 3 | 9 | 6 | 8 | 9 | 16 | 7 | 15 | 24 | 19 | 17 | 25 |
| K18700 fluoroacetyl-CoA thioesterase [EC:3.1.2.29] | 0 | 1 | 0 | 0 | 1 | 0 | 0 | 0 | 3 | 2 | 2 | 1 |
| K18702 CoA:oxalate CoA-transferase [EC:2.8.3.19] | 19 | 12 | 16 | 17 | 36 | 40 | 40 | 25 | 73 | 95 | 102 | 79 |
| K18703 succinate---hydroxymethylglutarate CoA-transferase [EC:2.8.3.13] | 0 | 1 | 1 | 1 | 1 | 2 | 1 | 3 | 0 | 0 | 1 | 1 |
| K18704 CDP-ribitol ribitolphosphotransferase / teichoic acid ribitol-phosphate polymerase [EC:2.7.8.14 2.7.8.47] | 0 | 5 | 2 | 6 | 2 | 5 | 4 | 3 | 1 | 4 | 1 | 10 |
| K18783 nigerose phosphorylase [EC:2.4.1.279] | 15 | 11 | 26 | 17 | 6 | 3 | 7 | 2 | 10 | 23 | 21 | 15 |
| K18785 beta-1,4-mannooligosaccharide/beta-1,4-mannosyl-N-acetylglucosamine phosphorylase [EC:2.4.1.319 2.4.1.320] | 836 | 884 | 941 | 912 | 1959 | 2064 | 2139 | 1945 | 1626 | 1590 | 1598 | 1437 |
| K18786 cellobionic acid phosphorylase [EC:2.4.1.321] | 3 | 2 | 0 | 0 | 2 | 1 | 0 | 1 | 0 | 1 | 3 | 1 |
| K18820 oligosaccharide 4-alpha-D-glucosyltransferase [EC:2.4.1.161] | 0 | 0 | 0 | 1 | 0 | 0 | 0 | 0 | 0 | 0 | 0 | 0 |
| K18855 UDP-N-acetylglucosamine 3-dehydrogenase [EC:1.1.1.374] | 2 | 2 | 6 | 3 | 1 | 2 | 0 | 2 | 6 | 15 | 9 | 9 |
| K18910 D-psicose/D-tagatose/L-ribulose 3-epimerase [EC:5.1.3.30 5.1.3.31] | 328 | 309 | 293 | 329 | 16 | 15 | 14 | 7 | 262 | 284 | 247 | 237 |
| K18915 ferric-chelate reductase [NAD(P)H] [EC:1.16.1.10] | 0 | 0 | 0 | 0 | 0 | 0 | 0 | 0 | 0 | 0 | 0 | 1 |
| K18916 phosphonate dehydrogenase [EC:1.20.1.1] | 365 | 315 | 290 | 326 | 506 | 506 | 471 | 483 | 57 | 55 | 65 | 54 |
| K18931 AMP phosphorylase [EC:2.4.2.57] | 0 | 0 | 0 | 0 | 0 | 0 | 0 | 0 | 0 | 1 | 0 | 1 |
| K19049 chondroitin AC lyase [EC:4.2.2.5] | 0 | 0 | 2 | 0 | 0 | 4 | 1 | 1 | 5 | 2 | 3 | 1 |
| K19050 heparin lyase [EC:4.2.2.7] | 9 | 8 | 13 | 8 | 33 | 28 | 24 | 24 | 27 | 20 | 28 | 30 |
| K19052 heparan-sulfate lyase [EC:4.2.2.8] | 1029 | 1138 | 1205 | 1119 | 4108 | 4055 | 4208 | 3910 | 3002 | 2930 | 3086 | 2943 |
| K19068 UDP-2-acetamido-2,6-beta-L-arabino-hexul-4-ose reductase [EC:1.1.1.367] | 468 | 489 | 495 | 457 | 117 | 117 | 106 | 141 | 271 | 272 | 259 | 251 |
| K19179 glycerophosphoinositol glycerophosphodiesterase [EC:3.1.4.44] | 0 | 0 | 3 | 0 | 1 | 0 | 0 | 1 | 0 | 0 | 0 | 0 |
| K19181 1,5-anhydro-D-fructose reductase (1,5-anhydro-D-mannitol-forming) [EC:1.1.1.292] | 0 | 0 | 0 | 0 | 0 | 0 | 4 | 2 | 0 | 4 | 5 | 0 |
| K19245 sulfonate dioxygenase [EC:1.14.11.-] | 0 | 0 | 2 | 0 | 0 | 0 | 0 | 0 | 0 | 0 | 0 | 0 |
| K19265 L-glyceraldehyde 3-phosphate reductase [EC:1.1.1.-] | 2247 | 2265 | 2198 | 2288 | 3333 | 3411 | 3333 | 3393 | 3030 | 2946 | 3251 | 3143 |
| K19270 sugar-phosphatase [EC:3.1.3.23] | 764 | 736 | 758 | 750 | 543 | 492 | 537 | 511 | 967 | 996 | 985 | 1079 |
| K19356 lytic cellulose monooxygenase (C1-hydroxylating) [EC:1.14.99.54] | 0 | 0 | 0 | 0 | 0 | 0 | 0 | 0 | 0 | 0 | 0 | 1 |
| K19386 NADH kinase [EC:2.7.1.86] | 0 | 0 | 0 | 0 | 0 | 0 | 0 | 0 | 0 | 0 | 0 | 1 |
| K19405 protein arginine kinase [EC:2.7.14.1] | 374 | 413 | 458 | 393 | 93 | 103 | 76 | 63 | 265 | 295 | 275 | 273 |
| K19422 glycosyltransferase EpsD [EC:2.4.-.-] | 8 | 15 | 6 | 11 | 41 | 27 | 31 | 30 | 14 | 20 | 6 | 13 |
| K19426 pyruvyl transferase EpsI [EC:2.-.-.-] | 0 | 0 | 0 | 0 | 0 | 0 | 0 | 1 | 0 | 0 | 0 | 0 |
| K19427 glycosyltransferase EpsJ [EC:2.4.-.-] | 67 | 69 | 58 | 67 | 44 | 34 | 40 | 34 | 86 | 116 | 87 | 85 |
| K19428 sugar transferase EpsL [EC:2.-.-.-] | 28 | 30 | 26 | 37 | 5 | 8 | 10 | 5 | 14 | 21 | 11 | 18 |
| K19429 acetyltransferase EpsM [EC:2.3.1.-] | 93 | 77 | 98 | 68 | 90 | 88 | 70 | 82 | 9 | 11 | 26 | 25 |
| K19430 pyridoxal phosphate-dependent aminotransferase EpsN [EC:2.6.1.-] | 6 | 2 | 3 | 3 | 1 | 1 | 3 | 2 | 3 | 2 | 4 | 1 |
| K19431 pyruvyl transferase EpsO [EC:2.-.-.-] | 17 | 18 | 14 | 11 | 3 | 2 | 3 | 1 | 0 | 0 | 3 | 0 |
| K19597 Au+-exporting ATPase [EC:3.6.1.-] | 0 | 0 | 0 | 0 | 0 | 0 | 0 | 2 | 0 | 0 | 0 | 0 |
| K19711 L-serine 3-dehydrogenase (NAD+) [EC:1.1.1.387] | 1331 | 1218 | 1237 | 1220 | 647 | 638 | 668 | 784 | 447 | 349 | 424 | 411 |
| K19713 thiosulfate dehydrogenase [EC:1.8.2.2] | 4279 | 4310 | 3943 | 4238 | 2287 | 2265 | 2103 | 2299 | 1317 | 1244 | 1268 | 1299 |
| K19802 L-Ala-D/L-Glu epimerase [EC:5.1.1.20] | 1838 | 1821 | 1871 | 1845 | 2858 | 3037 | 2860 | 2800 | 2730 | 2870 | 2637 | 2666 |
| K19814 glutamate 2,3-aminomutase [EC:5.4.3.9] | 0 | 2 | 0 | 1 | 0 | 0 | 0 | 0 | 0 | 1 | 0 | 1 |
| K19817 coenzyme F420H2 oxidase [EC:1.5.3.22] | 0 | 2 | 0 | 3 | 5 | 1 | 3 | 3 | 13 | 19 | 19 | 17 |
| K19954 alcohol dehydrogenase [EC:1.1.1.-] | 25 | 27 | 23 | 28 | 6 | 6 | 4 | 7 | 16 | 11 | 24 | 15 |
| K19955 alcohol dehydrogenase [EC:1.1.1.-] | 1095 | 1015 | 1041 | 1089 | 309 | 343 | 401 | 283 | 1282 | 1228 | 1274 | 1215 |
| K19967 D-threonine aldolase [EC:4.1.2.42] | 0 | 0 | 0 | 0 | 0 | 0 | 0 | 0 | 1 | 1 | 1 | 2 |
| K19997 GlcNAc-P-P-Und epimerase [EC:5.1.3.26] | 119 | 88 | 125 | 108 | 140 | 150 | 132 | 151 | 9 | 14 | 12 | 9 |
| K20025 (R)-2-hydroxyisocaproyl-CoA dehydratase alpha subunit [EC:4.2.1.157] | 30 | 18 | 19 | 18 | 0 | 0 | 0 | 0 | 6 | 7 | 6 | 8 |
| K20026 (R)-2-hydroxyisocaproyl-CoA dehydratase beta subunit [EC:4.2.1.157] | 16 | 17 | 8 | 15 | 8 | 15 | 10 | 16 | 5 | 7 | 4 | 5 |
| K20037 choline trimethylamine-lyase activating enzyme [EC:1.97.1.-] | 310 | 303 | 308 | 310 | 74 | 61 | 66 | 69 | 173 | 214 | 201 | 195 |
| K20038 choline trimethylamine-lyase [EC:4.3.99.4] | 970 | 829 | 896 | 885 | 319 | 331 | 312 | 299 | 600 | 627 | 677 | 633 |
| K20141 2-oxoglutarate carboxylase small subunit [EC:6.4.1.7] | 1 | 0 | 0 | 0 | 0 | 0 | 0 | 0 | 1 | 0 | 0 | 0 |
| K20201 protein arginine phosphatase [EC:3.9.1.2] | 2 | 1 | 3 | 1 | 0 | 0 | 0 | 1 | 0 | 4 | 1 | 1 |
| K20202 sulfide dehydrogenase subunit alpha [EC:1.8.1.19] | 1 | 1 | 1 | 1 | 5 | 2 | 3 | 3 | 3 | 3 | 3 | 2 |
| K20203 sulfide dehydrogenase subunit beta [EC:1.8.1.19] | 0 | 0 | 0 | 0 | 0 | 0 | 0 | 0 | 0 | 0 | 0 | 1 |
| K20215 diphthine synthase [EC:2.1.1.98] | 3 | 0 | 2 | 0 | 0 | 2 | 0 | 3 | 5 | 3 | 3 | 9 |
| K20238 sphingolipid C9-methyltransferase [EC:2.1.1.317] | 1 | 0 | 0 | 0 | 0 | 0 | 0 | 0 | 0 | 0 | 0 | 0 |
| K20415 ACP-SH:acetate ligase [EC:6.2.1.35] | 135 | 150 | 139 | 145 | 0 | 0 | 0 | 2 | 46 | 57 | 48 | 53 |
| K20429 dTDP-4-amino-4,6-dideoxy-D-glucose transaminase [EC:2.6.1.33] | 0 | 0 | 0 | 1 | 0 | 0 | 0 | 0 | 0 | 0 | 0 | 0 |
| K20525 oligo-alginate lyase [EC:4.2.2.26] | 2 | 1 | 2 | 1 | 2 | 1 | 2 | 3 | 1 | 1 | 1 | 1 |
| K20542 endoglucanase [EC:3.2.1.4] | 1658 | 1639 | 1503 | 1619 | 1401 | 1366 | 1349 | 1361 | 1882 | 1905 | 1827 | 1965 |
| K20548 3,6-anhydro-alpha-L-galactose dehydrogenase [EC:1.2.1.92] | 0 | 0 | 0 | 0 | 0 | 0 | 0 | 0 | 0 | 0 | 1 | 0 |
| K20549 3,6-anhydro-L-galactonate cycloisomerase [EC:5.5.1.25] | 0 | 1 | 0 | 0 | 1 | 0 | 0 | 1 | 0 | 0 | 2 | 0 |
| K20757 threo-3-hydroxy-D-aspartate ammonia-lyase [EC:4.3.1.27] | 0 | 0 | 0 | 0 | 0 | 0 | 0 | 0 | 0 | 1 | 0 | 1 |
| K20817 N-acyl-D-aspartate deacylase [EC:3.5.1.83] | 0 | 0 | 1 | 0 | 0 | 0 | 0 | 0 | 0 | 0 | 0 | 0 |
| K20848 fructan beta-(2,1)-fructosidase [EC:3.2.1.153] | 0 | 0 | 0 | 0 | 0 | 0 | 0 | 0 | 1 | 0 | 0 | 1 |
| K20882 (R)-2-hydroxy-4-methylpentanoate CoA-transferase [EC:2.8.3.24] | 15 | 12 | 17 | 29 | 3 | 9 | 12 | 11 | 45 | 36 | 40 | 48 |
| K20883 alpha-1,2-colitosyltransferase [EC:2.4.1.341] | 0 | 4 | 2 | 1 | 4 | 2 | 0 | 3 | 0 | 0 | 0 | 1 |
| K20885 beta-1,2-mannobiose phosphorylase / 1,2-beta-oligomannan phosphorylase [EC:2.4.1.339 2.4.1.340] | 434 | 470 | 475 | 473 | 1860 | 1711 | 1771 | 1767 | 1243 | 1209 | 1280 | 1280 |
| K20902 D-erythrulose 4-kinase [EC:2.7.1.210] | 2 | 0 | 0 | 2 | 3 | 0 | 0 | 2 | 14 | 17 | 16 | 17 |
| K20903 (R)-2-hydroxyglutaryl-CoA dehydratase subunit alpha [EC:4.2.1.167] | 34 | 32 | 40 | 34 | 62 | 76 | 56 | 55 | 19 | 44 | 34 | 26 |
| K20904 (R)-2-hydroxyglutaryl-CoA dehydratase subunit beta [EC:4.2.1.167] | 20 | 26 | 18 | 23 | 58 | 59 | 40 | 33 | 13 | 30 | 24 | 23 |
| K20937 1-butanol dehydrogenase (quinone) [EC:1.1.5.11] | 1 | 0 | 2 | 0 | 0 | 1 | 0 | 0 | 0 | 0 | 0 | 0 |
| K20938 long-chain alkane monooxygenase [EC:1.14.14.28] | 2111 | 2125 | 1864 | 2073 | 981 | 978 | 819 | 918 | 1628 | 1667 | 1558 | 1693 |
| K21054 D-erythritol 1-phosphate dehydrogenase [EC:1.1.1.402] | 0 | 0 | 0 | 0 | 0 | 0 | 2 | 0 | 0 | 1 | 2 | 2 |
| K21055 3-deoxy-D-glycero-D-galacto-nononate 9-phosphatase [EC:3.1.3.103] | 0 | 2 | 0 | 0 | 0 | 0 | 0 | 2 | 0 | 2 | 0 | 0 |
| K21065 beta-1,2-mannosidase [EC:3.2.1.197] | 1849 | 1943 | 2090 | 1816 | 7206 | 6993 | 7155 | 6757 | 5423 | 5253 | 5241 | 4939 |
| K21132 alpha-mannan endo-1,2-alpha-mannanase / glycoprotein endo-alpha-1,2-mannosidase [EC:3.2.1.198 3.2.1.130] | 1 | 2 | 1 | 3 | 3 | 5 | 7 | 3 | 0 | 0 | 3 | 1 |
| K21279 3-deoxy-D-glycero-D-galacto-nononate 9-phosphate synthase [EC:2.5.1.132] | 1 | 0 | 0 | 0 | 2 | 0 | 0 | 0 | 0 | 0 | 0 | 0 |
| K21285 teichoic acid glycerol-phosphate primase [EC:2.7.8.44] | 1 | 2 | 0 | 1 | 0 | 1 | 0 | 0 | 0 | 0 | 0 | 0 |
| K21298 1,2-beta-oligoglucan phosphorylase [EC:2.4.1.333] | 171 | 189 | 181 | 189 | 33 | 16 | 45 | 31 | 143 | 158 | 135 | 128 |
| K21350 sucrose 6(F)-phosphate phosphorylase [EC:2.4.1.329] | 2 | 0 | 0 | 3 | 0 | 0 | 1 | 0 | 5 | 5 | 8 | 17 |
| K21351 sucrose:sucrose fructosyltransferase [EC:2.4.1.99] | 0 | 0 | 0 | 0 | 0 | 0 | 0 | 1 | 0 | 0 | 0 | 0 |
| K21355 1,2-alpha-glucosylglycerol phosphorylase [EC:2.4.1.332] | 9 | 20 | 14 | 13 | 0 | 0 | 0 | 0 | 4 | 9 | 8 | 8 |
| K21399 2-amino-4-ketopentanoate thiolase alpha subunit [EC:2.3.1.263] | 1 | 0 | 2 | 0 | 0 | 0 | 0 | 0 | 0 | 0 | 1 | 0 |
| K21400 2-amino-4-ketopentanoate thiolase beta subunit [EC:2.3.1.263] | 2 | 2 | 0 | 3 | 2 | 2 | 1 | 0 | 5 | 0 | 2 | 4 |
| K21416 acetoin:2,6-dichlorophenolindophenol oxidoreductase subunit alpha [EC:1.1.1.-] | 1743 | 1772 | 1547 | 1733 | 841 | 809 | 868 | 885 | 1267 | 1199 | 1108 | 1233 |
| K21417 acetoin:2,6-dichlorophenolindophenol oxidoreductase subunit beta [EC:1.1.1.-] | 1971 | 1961 | 1848 | 2024 | 1099 | 1025 | 956 | 1098 | 1342 | 1433 | 1286 | 1388 |
| K21420 leucyl-tRNA---protein transferase [EC:2.3.2.29] | 958 | 973 | 953 | 997 | 486 | 489 | 501 | 515 | 309 | 297 | 285 | 260 |
| K21430 aldose sugar dehydrogenase [EC:1.1.5.-] | 2312 | 2431 | 2248 | 2329 | 1314 | 1253 | 1365 | 1341 | 1382 | 1308 | 1327 | 1522 |
| K21481 heme oxygenase (mycobilin-producing) [EC:1.14.99.57] | 0 | 0 | 0 | 1 | 0 | 0 | 0 | 0 | 0 | 0 | 0 | 0 |
| K21511 capsid assembly protease [EC:3.4.21.-] | 0 | 0 | 0 | 0 | 15 | 27 | 9 | 8 | 3 | 7 | 3 | 4 |
| K21512 terminase, large subunit [EC:3.1.21.4] | 0 | 3 | 2 | 1 | 35 | 12 | 19 | 22 | 8 | 9 | 14 | 15 |
| K21527 adenine modification enzyme [EC:2.3.1.-] | 0 | 8 | 3 | 0 | 5 | 0 | 0 | 0 | 0 | 0 | 6 | 0 |
| K21567 ferredoxin/flavodoxin---NADP+ reductase [EC:1.18.1.2 1.19.1.1] | 53 | 83 | 83 | 83 | 18 | 19 | 24 | 20 | 56 | 89 | 77 | 74 |
| K21575 neopullulanase [EC:3.2.1.135] | 1353 | 1464 | 1661 | 1451 | 4922 | 4939 | 4903 | 4610 | 3386 | 3258 | 3585 | 3364 |
| K21576 glycine/sarcosine/betaine reductase complex component C subunit alpha [EC:1.21.4.2 1.21.4.3 1.21.4.4] | 56 | 58 | 76 | 58 | 35 | 26 | 22 | 43 | 64 | 65 | 54 | 61 |
| K21577 glycine/sarcosine/betaine reductase complex component C subunit beta [EC:1.21.4.2 1.21.4.3 1.21.4.4] | 95 | 75 | 84 | 91 | 31 | 41 | 48 | 31 | 58 | 81 | 69 | 73 |
| K21578 betaine reductase complex component B subunit alpha [EC:1.21.4.4] | 38 | 56 | 47 | 41 | 8 | 5 | 8 | 10 | 28 | 19 | 21 | 22 |
| K21579 betaine reductase complex component B subunit beta [EC:1.21.4.4] | 40 | 36 | 37 | 38 | 8 | 21 | 7 | 2 | 21 | 32 | 27 | 19 |
| K21583 sarcosine reductase complex component B subunit alpha [EC:1.21.4.3] | 52 | 22 | 39 | 35 | 8 | 10 | 10 | 10 | 23 | 38 | 25 | 15 |
| K21584 sarcosine reductase complex component B subunit beta [EC:1.21.4.3] | 5 | 14 | 15 | 14 | 1 | 1 | 3 | 3 | 11 | 26 | 13 | 9 |
| K21592 teichoic acid ribitol-phosphate primase [EC:2.7.8.46] | 1 | 2 | 1 | 1 | 0 | 0 | 0 | 0 | 0 | 0 | 0 | 1 |
| K21616 D-altritol 5-dehydrogenase [EC:1.1.1.407] | 0 | 0 | 0 | 1 | 2 | 2 | 0 | 0 | 4 | 4 | 2 | 3 |
| K21702 galactitol 2-dehydrogenase (L-tagatose-forming) [EC:1.1.1.406] | 0 | 2 | 0 | 0 | 0 | 0 | 0 | 0 | 0 | 1 | 0 | 0 |
| K21713 lytic chitin monoxygenase [EC:1.14.99.53] | 53 | 33 | 40 | 35 | 5 | 7 | 11 | 6 | 39 | 59 | 24 | 52 |
| K21750 3-deoxy-D-glycero-D-galacto-nononate cytidylyltransferase [EC:2.7.7.92] | 2 | 1 | 0 | 0 | 2 | 2 | 0 | 0 | 0 | 0 | 0 | 3 |
| K21887 cation-transporting P-type ATPase J [EC:7.2.2.-] | 0 | 1 | 2 | 0 | 0 | 1 | 1 | 1 | 1 | 1 | 1 | 1 |
| K21909 D-erythrulose 1-phosphate 3-epimerase [EC:5.1.3.38] | 16 | 13 | 9 | 23 | 16 | 17 | 20 | 16 | 34 | 39 | 37 | 41 |

| | | | | | | | | | | | | |
|---|---|---|---|---|---|---|---|---|---|---|---|---|
| K21910 L-erythrulose 1-phosphate isomerase [EC:5.3.1.33] | 14 | 23 | 17 | 16 | 1 | 5 | 4 | 2 | 6 | 10 | 11 | 3 |
| K21911 D-erythrulose 4-phosphate isomerase [EC:5.3.1.34] | 1 | 0 | 0 | 0 | 0 | 0 | 0 | 2 | 1 | 1 | 4 | 1 |
| K21935 beta-lysine N6-acetyltransferase [EC:2.3.1.264] | 0 | 3 | 0 | 0 | 0 | 0 | 0 | 0 | 0 | 0 | 1 | 3 |
| K21936 anaerobilin synthase [EC:2.1.1.342] | 124 | 118 | 121 | 143 | 75 | 91 | 85 | 95 | 272 | 232 | 225 | 278 |
| K21948 3-dehydrotetronate 4-kinase [EC:2.7.1.217] | 586 | 476 | 494 | 531 | 131 | 113 | 141 | 125 | 1019 | 999 | 1008 | 1070 |
| K22015 formate dehydrogenase (acceptor) [EC:1.17.99.7] | 2567 | 2482 | 2373 | 2812 | 1900 | 1930 | 1858 | 1796 | 3643 | 3605 | 3464 | 3741 |
| K22024 4-phospho-D-threonate 3-dehydrogenase / 4-phospho-D-erythronate 3-dehydrogenase [EC:1.1.1.408 1.1.1.409] | 2366 | 2228 | 2299 | 2442 | 969 | 1135 | 1081 | 954 | 2452 | 2680 | 2449 | 2569 |
| K22025 D-erythronate 2-dehydrogenase [EC:1.1.1.410] | 0 | 0 | 0 | 0 | 0 | 0 | 0 | 0 | 0 | 3 | 3 | 4 |
| K22027 indole-3-acetate monooxygenase [EC:1.14.13.235] | 2 | 0 | 0 | 3 | 0 | 0 | 0 | 0 | 0 | 0 | 0 | 0 |
| K22078 protein-glucosylgalactosylhydroxylysine glucosidase [EC:3.2.1.107] | 3 | 5 | 10 | 4 | 21 | 16 | 20 | 11 | 10 | 8 | 8 | 4 |
| K22129 D-threonate/D-erythronate kinase [EC:2.7.1.219 2.7.1.220] | 2485 | 2773 | 2557 | 2700 | 1411 | 1377 | 1439 | 1418 | 2868 | 2997 | 2876 | 2961 |
| K22130 3-dehydro-4-phosphotetronate decarboxylase [EC:4.1.1.104] | 87 | 65 | 62 | 76 | 20 | 20 | 12 | 23 | 126 | 141 | 97 | 110 |
| K22131 2-dehydrotetronate isomerase [EC:5.3.1.35] | 1647 | 1570 | 1483 | 1623 | 792 | 790 | 745 | 878 | 927 | 996 | 925 | 988 |
| K22209 D(-)-tartrate dehydratase [EC:4.2.1.81] | 1216 | 1172 | 1118 | 1222 | 661 | 691 | 596 | 666 | 386 | 399 | 372 | 390 |
| K22213 6-methylsalicylate decarboxylase [EC:4.1.1.52] | 3 | 4 | 0 | 1 | 0 | 0 | 0 | 0 | 0 | 0 | 0 | 0 |
| K22247 polysialic-acid O-acetyltransferase [EC:2.3.1.136] | 2 | 1 | 0 | 1 | 0 | 0 | 0 | 2 | 9 | 16 | 13 | 12 |
| K22250 poly(3-hydroxyoctanoate) depolymerase [EC:3.1.1.76] | 1413 | 1314 | 1315 | 1364 | 768 | 727 | 766 | 834 | 428 | 437 | 423 | 428 |
| K22278 peptidoglycan-N-acetylglucosamine deacetylase [EC:3.5.1.104] | 885 | 742 | 835 | 823 | 238 | 200 | 188 | 162 | 607 | 727 | 739 | 692 |
| K22303 alpha-ketoglutarate-dependent sulfate ester dioxygenase [EC:1.14.11.-] | 1557 | 1533 | 1458 | 1545 | 994 | 1019 | 927 | 1031 | 488 | 387 | 418 | 417 |
| K22306 glucosyl-3-phosphoglycerate phosphatase [EC:3.1.3.85] | 3 | 5 | 2 | 0 | 0 | 2 | 1 | 2 | 2 | 4 | 3 | 6 |
| K22310 L-ornithine Nalpha-acyltransferase [EC:2.3.2.30] | 0 | 0 | 0 | 0 | 0 | 0 | 1 | 0 | 0 | 4 | 2 | 0 |
| K22338 formate dehydrogenase (NAD+, ferredoxin) subunit A [EC:1.17.1.11] | 0 | 9 | 10 | 0 | 3 | 0 | 0 | 0 | 0 | 0 | 12 | 0 |
| K22339 formate dehydrogenase (NAD+, ferredoxin) subunit B [EC:1.17.1.11] | 2 | 1 | 2 | 1 | 2 | 1 | 3 | 0 | 0 | 2 | 4 | 2 |
| K22340 formate dehydrogenase (NAD+, ferredoxin) subunit C [EC:1.17.1.11] | 0 | 0 | 1 | 1 | 1 | 4 | 1 | 0 | 4 | 2 | 0 | 3 |
| K22341 formate dehydrogenase (NAD+, ferredoxin) subunit [EC:1.17.1.11] | 3 | 1 | 4 | 3 | 6 | 2 | 6 | 5 | 4 | 8 | 4 | 9 |
| K22342 dimethylamine monooxygenase subunit A [EC:1.14.13.238] | 0 | 0 | 0 | 0 | 1 | 0 | 0 | 0 | 0 | 0 | 0 | 0 |
| K22343 dimethylamine monooxygenase subunit B [EC:1.14.13.238] | 2 | 2 | 0 | 0 | 0 | 0 | 0 | 1 | 0 | 0 | 0 | 0 |
| K22344 dimethylamine monooxygenase subunit C [EC:1.14.13.238] | 3 | 0 | 1 | 0 | 0 | 0 | 0 | 0 | 0 | 0 | 0 | 0 |
| K22348 manganese oxidase [EC:1.16.3.3] | 107 | 134 | 116 | 104 | 14 | 6 | 13 | 12 | 218 | 221 | 206 | 181 |
| K22349 manganese oxidase [EC:1.16.3.3] | 3 | 0 | 3 | 13 | 0 | 0 | 0 | 0 | 0 | 0 | 0 | 0 |
| K22405 NADH oxidase (H2O-forming) [EC:1.6.3.4] | 251 | 254 | 250 | 231 | 30 | 29 | 31 | 28 | 332 | 317 | 327 | 396 |
| K22409 N-acetylmuramoyl-L-alanine amidase [EC:3.5.1.28] | 0 | 0 | 0 | 0 | 0 | 0 | 0 | 0 | 0 | 0 | 0 | 1 |
| K22430 caffeyl-CoA reductase-Etf complex subunit CarC [EC:1.3.1.108] | 41 | 38 | 36 | 38 | 0 | 2 | 2 | 1 | 24 | 16 | 15 | 21 |
| K22431 caffeyl-CoA reductase-Etf complex subunit CarD [EC:1.3.1.108] | 60 | 63 | 36 | 58 | 4 | 1 | 8 | 3 | 34 | 22 | 19 | 28 |
| K22432 caffeyl-CoA reductase-Etf complex subunit CarE [EC:1.3.1.108] | 547 | 504 | 554 | 557 | 115 | 124 | 116 | 91 | 331 | 406 | 299 | 294 |
| K22438 trans-aconitate 3-methyltransferase [EC:2.1.1.145] | 0 | 0 | 0 | 0 | 0 | 0 | 0 | 0 | 0 | 1 | 0 | 1 |
| K22441 diamine N-acetyltransferase [EC:2.3.1.57] | 225 | 182 | 188 | 186 | 61 | 66 | 88 | 65 | 440 | 463 | 440 | 453 |
| K22443 carnitine monooxygenase subunit [EC:1.14.13.239] | 441 | 392 | 430 | 467 | 131 | 141 | 91 | 89 | 757 | 877 | 851 | 952 |
| K22444 carnitine monooxygenase subunit [EC:1.14.13.239] | 342 | 357 | 310 | 368 | 99 | 103 | 90 | 88 | 724 | 717 | 700 | 763 |
| K22452 protein-glutamine gamma-glutamyltransferase [EC:2.3.2.13] | 2555 | 2479 | 2331 | 2436 | 1367 | 1238 | 1219 | 1351 | 772 | 723 | 640 | 727 |
| K22465 5-hydroxybenzimidazole synthase [EC:4.1.99.23] | 33 | 15 | 26 | 31 | 51 | 54 | 50 | 33 | 78 | 63 | 76 | 93 |
| K22551 N-methyl-L-proline demethylase [EC:1.5.3.-] | 0 | 0 | 0 | 0 | 0 | 0 | 1 | 0 | 1 | 0 | 1 | 1 |
| K22579 rifampicin phosphotransferase [EC:2.7.9.6] | 74 | 64 | 72 | 80 | 17 | 12 | 20 | 10 | 36 | 51 | 44 | 52 |
| K22589 threo-3-hydroxy-L-aspartate ammonia-lyase [EC:4.3.1.16] | 1546 | 1492 | 1398 | 1419 | 733 | 774 | 671 | 816 | 466 | 401 | 456 | 493 |
| K22597 glucosylglycerate phosphorylase [EC:2.4.1.352] | 280 | 231 | 243 | 227 | 90 | 54 | 104 | 76 | 224 | 226 | 197 | 218 |
| K22616 ornithine lipid ester-linked acyl 2-hydroxylase [EC:1.14.11.58] | 3 | 0 | 0 | 0 | 2 | 0 | 0 | 3 | 2 | 2 | 3 | 1 |
| K22617 lyso-ornithine lipid O-acyltransferase [EC:2.3.1.270] | 1232 | 1225 | 1100 | 1147 | 626 | 644 | 519 | 707 | 407 | 332 | 340 | 314 |
| K22704 hopanoid C-3 methylase [EC:2.1.1.-] | 4 | 11 | 9 | 9 | 3 | 0 | 3 | 8 | 12 | 9 | 10 | 9 |
| K22721 inositol phosphorylceramide mannosyltransferase catalytic subunit [EC:2.4.1.-] | 1 | 0 | 0 | 0 | 0 | 1 | 0 | 0 | 0 | 0 | 0 | 0 |
| K22757 glutaminyl-peptide cyclotransferase [EC:2.3.2.5] | 6 | 2 | 2 | 5 | 6 | 0 | 3 | 4 | 2 | 0 | 0 | 1 |
| K22837 SAMP-activating enzyme [EC:2.7.7.100] | 1 | 0 | 0 | 1 | 0 | 1 | 1 | 1 | 1 | 1 | 0 | 0 |
| K22874 L-tyrosine isonitrile desaturase/decarboxylase [EC:1.14.20.9 1.14.20.10] | 1210 | 1114 | 991 | 1065 | 569 | 529 | 489 | 630 | 324 | 298 | 351 | 346 |
| K22906 phloretin hydrolase [EC:3.7.1.4] | 7 | 7 | 11 | 7 | 0 | 0 | 3 | 2 | 11 | 6 | 7 | 8 |
| K22927 cyclic-di-AMP phosphodiesterase [EC:3.1.4.59] | 1506 | 1463 | 1428 | 1521 | 424 | 439 | 463 | 380 | 1292 | 1346 | 1280 | 1312 |
| K22952 L-tyrosine isonitrile synthase [EC:4.1.99.24] | 1014 | 954 | 870 | 963 | 472 | 449 | 443 | 550 | 275 | 275 | 248 | 281 |
| K23078 menaquinone C8-methyltransferase [EC:2.1.1.350] | 98 | 86 | 129 | 104 | 168 | 145 | 167 | 139 | 158 | 151 | 139 | 162 |
| K23107 1-deoxyxylulose-5-phosphate synthase [EC:1.1.-.-] | 1120 | 1163 | 1062 | 1238 | 532 | 645 | 647 | 563 | 1736 | 1869 | 1761 | 1775 |
| K23110 (R)-phenyllactyl-CoA dehydratase alpha subunit [EC:4.2.1.-] | 12 | 4 | 3 | 5 | 0 | 0 | 0 | 0 | 1 | 3 | 3 | 3 |
| K23111 (R)-phenyllactyl-CoA dehydratase beta subunit [EC:4.2.1.-] | 3 | 2 | 2 | 1 | 0 | 0 | 0 | 0 | 0 | 0 | 0 | 0 |
| K23122 staphylopine dehydrogenase [EC:1.5.1.52] | 1 | 0 | 0 | 0 | 0 | 0 | 0 | 0 | 0 | 0 | 0 | 0 |
| K23147 aminoimidazole riboside kinase [EC:2.7.1.223] | 14 | 18 | 17 | 15 | 0 | 0 | 0 | 0 | 7 | 14 | 6 | 8 |
| K23148 cytidine diphosphoramidate kinase [EC:2.7.1.224] | 0 | 0 | 0 | 0 | 2 | 0 | 3 | 3 | 5 | 0 | 3 | 4 |
| K23149 5-phospho-D-xylono-1,4-lactonase [EC:3.1.1.104] | 721 | 538 | 637 | 595 | 51 | 55 | 40 | 40 | 413 | 406 | 411 | 404 |
| K23244 D-apiose dehydrogenase [EC:1.1.1.-] | 0 | 0 | 0 | 1 | 0 | 2 | 2 | 1 | 4 | 3 | 1 | 4 |
| K23245 D-apionate oxidoisomerase [EC:1.1.1.-] | 49 | 35 | 37 | 27 | 7 | 7 | 5 | 2 | 78 | 90 | 84 | 71 |
| K23246 D-apionate lactonase [EC:3.1.1.-] | 2 | 0 | 0 | 0 | 0 | 3 | 5 | 2 | 4 | 5 | 4 | 3 |
| K23247 3-oxo-isoapionate kinase [EC:2.7.1.-] | 0 | 2 | 0 | 0 | 2 | 5 | 3 | 0 | 2 | 4 | 8 | 0 |
| K23248 3-oxo-isoapionate decarboxylase [EC:4.1.1.-] | 0 | 0 | 0 | 0 | 0 | 0 | 0 | 0 | 2 | 0 | 1 | 0 |
| K23254 P-type Ag+ transporter [EC:7.2.2.15] | 1406 | 1455 | 1422 | 1368 | 2061 | 2280 | 2102 | 2054 | 136 | 151 | 132 | 139 |
| K23256 NADPH-dependent curcumin reductase [EC:1.3.1.-] | 2926 | 2835 | 2619 | 2773 | 1537 | 1548 | 1491 | 1730 | 1431 | 1375 | 1420 | 1417 |
| K23271 erythritol/L-threitol dehydrogenase [EC:1.1.1.-] | 284 | 317 | 294 | 287 | 65 | 67 | 57 | 56 | 423 | 453 | 443 | 460 |
| K23300 menaquinone reductase, multiheme cytochrome c subunit [EC:1.97.-.-] | 6 | 4 | 4 | 1 | 0 | 2 | 3 | 0 | 2 | 1 | 2 | 0 |
| K23301 menaquinone reductase, iron-sulfur cluster-binding subunit [EC:1.97.-.-] | 3 | 5 | 6 | 6 | 0 | 0 | 0 | 2 | 0 | 2 | 2 | 1 |
| K23463 (Z)-2-((N-methylformamido)methylene)-5-hydroxybutyrolactone dehydrogenase [EC:1.2.1.-] | 0 | 0 | 0 | 0 | 0 | 0 | 0 | 1 | 0 | 1 | 0 |
| K23518 O-acetyl-ADP-ribose deacetylase [EC:3.1.1.106] | 1245 | 1351 | 1195 | 1330 | 787 | 816 | 851 | 777 | 871 | 771 | 839 | 835 |
| ko99981 Carbohydrate metabolism | 11010 | 11083 | 10880 | 11239 | 10813 | 10748 | 10565 | 10942 | 10698 | 10732 | 10570 | 11169 |
| K00782 L-lactate dehydrogenase complex protein LldG | 497 | 610 | 628 | 646 | 1531 | 1552 | 1497 | 1447 | 1043 | 934 | 1018 | 949 |
| K02566 NagD protein | 566 | 628 | 507 | 599 | 359 | 382 | 296 | 316 | 720 | 699 | 708 | 826 |
| K04767 acetoin utilization protein AcuB | 35 | 31 | 37 | 27 | 3 | 4 | 8 | 10 | 64 | 56 | 67 | 52 |
| K06206 sugar fermentation stimulation protein A | 1827 | 1807 | 1709 | 1809 | 894 | 908 | 888 | 922 | 1329 | 1388 | 1331 | 1447 |
| K06605 myo-inositol catabolism protein IolH | 425 | 414 | 357 | 401 | 105 | 83 | 93 | 93 | 710 | 779 | 770 | 823 |
| K11184 catabolite repression HPr-like protein | 71 | 80 | 82 | 82 | 19 | 15 | 13 | 30 | 82 | 72 | 77 | 77 |
| K11930 periplasmic protein TorT | 3 | 5 | 1 | 7 | 14 | 5 | 15 | 8 | 58 | 59 | 44 | 43 |
| K17641 beta-xylosidase | 0 | 0 | 0 | 0 | 0 | 0 | 0 | 0 | 0 | 1 | 0 | 0 |
| K18928 L-lactate dehydrogenase complex protein LldE | 633 | 646 | 727 | 714 | 1774 | 1785 | 1858 | 1781 | 1144 | 1196 | 1229 | 1207 |
| K18929 L-lactate dehydrogenase complex protein LldF | 1218 | 1283 | 1415 | 1334 | 3268 | 3389 | 3335 | 3391 | 2271 | 2088 | 2229 | 2231 |
| K18930 D-lactate dehydrogenase | 4663 | 4520 | 4436 | 4560 | 2455 | 2246 | 2166 | 2554 | 1450 | 1448 | 1307 | 1498 |
| K19504 glucoselysine-6-phosphate deglycase | 534 | 561 | 509 | 528 | 149 | 170 | 164 | 150 | 975 | 996 | 915 | 1101 |
| K19510 fructoselysine-6-phosphate deglycase | 535 | 490 | 468 | 532 | 236 | 189 | 216 | 221 | 805 | 967 | 812 | 861 |
| K22111 9-O-acetyl-N-acetylneuraminic acid deacetylase | 3 | 6 | 1 | 0 | 6 | 17 | 16 | 19 | 46 | 48 | 63 | 54 |
| K22136 bacillithiol synthase | 0 | 2 | 2 | 0 | 0 | 2 | 0 | 0 | 1 | 1 | 0 | 0 |
| K22216 dehydratase ilvD1 | 0 | 0 | 1 | 0 | 0 | 1 | 0 | 0 | 0 | 0 | 0 | 0 |
| ko99982 Energy metabolism | 32087 | 31907 | 30917 | 31470 | 25007 | 25227 | 24480 | 25537 | 26157 | 26545 | 25641 | 26431 |
| K00183 prokaryotic molybdopterin-containing oxidoreductase family, molybdopterin binding subunit | 47 | 54 | 51 | 64 | 45 | 27 | 43 | 30 | 111 | 157 | 151 | 146 |
| K00184 prokaryotic molybdopterin-containing oxidoreductase family, iron-sulfur binding subunit | 54 | 44 | 41 | 24 | 41 | 25 | 22 | 23 | 90 | 118 | 77 | 82 |
| K00185 prokaryotic molybdopterin-containing oxidoreductase family, membrane subunit | 46 | 55 | 44 | 37 | 24 | 23 | 33 | 30 | 81 | 106 | 91 | 86 |
| K02164 nitric oxide reductase NorE protein | 956 | 849 | 809 | 873 | 508 | 497 | 459 | 520 | 343 | 281 | 294 | 275 |
| K02379 FdhD protein | 2163 | 2000 | 2080 | 2116 | 1271 | 1285 | 1243 | 1360 | 1223 | 1312 | 1297 | 1304 |
| K02381 FdrA protein | 49 | 51 | 51 | 41 | 45 | 29 | 37 | 32 | 156 | 180 | 187 | 192 |
| K02448 nitric oxide reductase NorD protein | 2748 | 2754 | 2487 | 2552 | 1662 | 1454 | 1497 | 1697 | 951 | 779 | 765 | 839 |
| K02569 cytochrome c-type protein NapC | 485 | 492 | 540 | 514 | 273 | 299 | 254 | 293 | 191 | 195 | 198 | 197 |
| K02570 periplasmic nitrate reductase NapD | 430 | 487 | 433 | 435 | 232 | 236 | 241 | 259 | 143 | 195 | 143 | 171 |
| K02571 periplasmic nitrate reductase NapE | 245 | 244 | 267 | 238 | 149 | 97 | 144 | 142 | 82 | 60 | 60 | 62 |
| K02572 ferredoxin-type protein NapF | 785 | 699 | 721 | 704 | 409 | 318 | 265 | 319 | 270 | 220 | 204 | 222 |
| K02573 ferredoxin-type protein NapG | 91 | 76 | 55 | 65 | 132 | 103 | 111 | 121 | 180 | 179 | 167 | 163 |
| K02574 ferredoxin-type protein NapH | 65 | 72 | 83 | 78 | 94 | 111 | 103 | 106 | 164 | 177 | 157 | 135 |
| K02585 nitrogen fixation protein NifB | 240 | 258 | 257 | 323 | 397 | 438 | 399 | 386 | 8 | 23 | 10 | 17 |
| K02587 nitrogenase molybdenum-cofactor synthesis protein NifE | 332 | 319 | 317 | 308 | 350 | 444 | 405 | 376 | 34 | 36 | 19 | 24 |
| K02592 nitrogenase molybdenum-iron protein NifN | 242 | 285 | 235 | 264 | 387 | 403 | 396 | 364 | 0 | 1 | 0 | 4 |
| K02593 nitrogen fixation protein NifT | 37 | 37 | 37 | 31 | 58 | 57 | 61 | 65 | 0 | 0 | 2 | 0 |
| K02595 nitrogenase-stabilizing/protective protein | 31 | 48 | 44 | 39 | 72 | 70 | 54 | 61 | 0 | 0 | 0 | 0 |
| K02596 nitrogen fixation protein NifX | 224 | 198 | 199 | 212 | 281 | 294 | 301 | 296 | 0 | 0 | 2 | 2 |
| K02597 nitrogen fixation protein NifZ | 96 | 88 | 65 | 80 | 128 | 96 | 114 | 98 | 1 | 0 | 0 | 0 |
| K03521 electron transfer flavoprotein beta subunit | 2888 | 2882 | 2826 | 2865 | 3078 | 3008 | 3178 | 3203 | 2425 | 2488 | 2183 | 2362 |
| K03612 Na+-translocating ferredoxin:NAD+ oxidoreductase subunit G | 1198 | 1201 | 1230 | 1233 | 1844 | 1753 | 1698 | 1635 | 1737 | 1588 | 1722 | 1750 |
| K03613 Na+-translocating ferredoxin:NAD+ oxidoreductase subunit E | 2284 | 2209 | 2139 | 2262 | 2028 | 2279 | 2037 | 2198 | 2026 | 2053 | 1965 | 2027 |
| K03617 Na+-translocating ferredoxin:NAD+ oxidoreductase subunit A | 1423 | 1509 | 1487 | 1414 | 1588 | 1696 | 1604 | 1723 | 1571 | 1552 | 1627 | 1471 |
| K03839 flavodoxin I | 1026 | 1055 | 1072 | 1014 | 2319 | 2327 | 2357 | 2386 | 1809 | 1753 | 1786 | 1793 |
| K03840 flavodoxin II | 397 | 392 | 368 | 414 | 230 | 269 | 227 | 253 | 519 | 640 | 512 | 594 |
| K03855 ferredoxin like protein | 55 | 76 | 81 | 59 | 96 | 114 | 115 | 91 | 24 | 24 | 20 | 35 |
| K04013 cytochrome c-type protein NrfB | 1 | 2 | 0 | 0 | 5 | 10 | 8 | 9 | 41 | 31 | 26 | 28 |

| | | | | | | | | | | | | |
|---|---|---|---|---|---|---|---|---|---|---|---|---|
| K04014 protein NrfC | 53 | 41 | 40 | 46 | 107 | 101 | 93 | 110 | 117 | 128 | 151 | 155 |
| K04015 protein NrfD | 62 | 83 | 56 | 58 | 101 | 128 | 93 | 125 | 146 | 146 | 193 | 163 |
| K04016 cytochrome c-type biogenesis protein NrfE | 8 | 3 | 10 | 2 | 9 | 17 | 13 | 17 | 99 | 98 | 83 | 95 |
| K04017 formate-dependent nitrite reductase complex subunit NrfF | 0 | 3 | 0 | 1 | 6 | 4 | 4 | 3 | 14 | 16 | 14 | 39 |
| K04018 formate-dependent nitrite reductase complex subunit NrfG | 0 | 0 | 2 | 4 | 8 | 4 | 3 | 5 | 30 | 41 | 47 | 40 |
| K04488 nitrogen fixation protein NifU and related proteins | 1371 | 1245 | 1308 | 1302 | 569 | 719 | 593 | 635 | 850 | 1048 | 992 | 1011 |
| K04747 nitric oxide reductase NorF protein | 0 | 0 | 0 | 0 | 0 | 0 | 0 | 0 | 0 | 2 | 1 | 0 |
| K04748 nitric oxide reductase NorQ protein | 792 | 824 | 817 | 748 | 443 | 353 | 368 | 479 | 285 | 308 | 281 | 273 |
| K04755 ferredoxin, 2Fe-2S | 716 | 790 | 740 | 728 | 478 | 489 | 497 | 446 | 473 | 514 | 478 | 498 |
| K05337 ferredoxin | 25 | 27 | 25 | 29 | 25 | 27 | 17 | 17 | 20 | 37 | 14 | 28 |
| K05524 ferredoxin | 906 | 840 | 823 | 864 | 483 | 513 | 462 | 506 | 286 | 257 | 261 | 277 |
| K05796 electron transport protein HydN | 565 | 573 | 517 | 598 | 364 | 336 | 340 | 346 | 897 | 867 | 860 | 931 |
| K06205 MioC protein | 1006 | 959 | 934 | 967 | 546 | 593 | 611 | 559 | 607 | 639 | 550 | 585 |
| K06862 energy-converting hydrogenase B subunit Q | 0 | 0 | 0 | 0 | 0 | 0 | 0 | 0 | 1 | 1 | 5 | 2 |
| K09165 dodecin | 413 | 389 | 403 | 363 | 208 | 232 | 188 | 251 | 136 | 137 | 116 | 142 |
| K11107 ferredoxin | 160 | 178 | 155 | 179 | 122 | 117 | 95 | 102 | 216 | 236 | 202 | 213 |
| K12143 hydrogenase-4 component H | 126 | 121 | 111 | 122 | 178 | 225 | 173 | 162 | 5 | 7 | 2 | 5 |
| K12262 cytochrome b561 | 1339 | 1296 | 1250 | 1196 | 844 | 868 | 895 | 902 | 470 | 428 | 482 | 499 |
| K13795 citrate/tricarballylate utilization protein | 1 | 4 | 0 | 0 | 2 | 0 | 0 | 0 | 0 | 0 | 3 | 0 |
| K13796 tricarballylate dehydrogenase | 2 | 1 | 2 | 0 | 1 | 0 | 3 | 2 | 0 | 0 | 0 | 1 |
| K13819 NifU-like protein | 188 | 163 | 171 | 155 | 217 | 220 | 235 | 201 | 6 | 13 | 11 | 10 |
| K14086 ech hydrogenase subunit A | 59 | 52 | 73 | 57 | 18 | 16 | 20 | 19 | 66 | 110 | 105 | 86 |
| K14087 ech hydrogenase subunit B | 12 | 17 | 17 | 13 | 3 | 5 | 7 | 6 | 12 | 15 | 8 | 7 |
| K14088 ech hydrogenase subunit C | 24 | 33 | 26 | 24 | 6 | 5 | 3 | 3 | 55 | 46 | 40 | 34 |
| K14089 ech hydrogenase subunit D | 22 | 19 | 26 | 17 | 5 | 4 | 5 | 4 | 34 | 44 | 46 | 31 |
| K14090 ech hydrogenase subunit E | 77 | 103 | 100 | 72 | 25 | 29 | 16 | 19 | 144 | 123 | 161 | 136 |
| K14091 ech hydrogenase subunit F | 7 | 5 | 6 | 7 | 1 | 1 | 5 | 9 | 9 | 18 | 11 | 5 |
| K14092 energy-converting hydrogenase A subunit A | 0 | 0 | 0 | 1 | 0 | 0 | 0 | 1 | 0 | 1 | 1 | 1 |
| K14093 energy-converting hydrogenase A subunit B | 0 | 0 | 0 | 0 | 0 | 0 | 0 | 0 | 1 | 2 | 3 | 8 |
| K14094 energy-converting hydrogenase A subunit C | 0 | 0 | 0 | 1 | 0 | 0 | 0 | 0 | 0 | 0 | 0 | 2 |
| K14095 energy-converting hydrogenase A subunit D | 0 | 0 | 0 | 1 | 0 | 0 | 0 | 0 | 2 | 2 | 1 | 3 |
| K14096 energy-converting hydrogenase A subunit E | 0 | 0 | 0 | 0 | 0 | 0 | 0 | 3 | 0 | 2 | 0 | 3 |
| K14097 energy-converting hydrogenase A subunit F | 0 | 0 | 0 | 0 | 0 | 0 | 0 | 0 | 0 | 3 | 1 | 4 |
| K14098 energy-converting hydrogenase A subunit G | 0 | 2 | 0 | 0 | 0 | 0 | 0 | 0 | 4 | 1 | 2 | 3 |
| K14099 energy-converting hydrogenase A subunit H | 0 | 3 | 0 | 0 | 0 | 0 | 0 | 2 | 10 | 5 | 1 | 4 |
| K14100 energy-converting hydrogenase A subunit I | 0 | 0 | 0 | 0 | 0 | 0 | 0 | 0 | 0 | 1 | 2 | 0 |
| K14101 energy-converting hydrogenase A subunit J | 0 | 2 | 0 | 0 | 2 | 0 | 2 | 0 | 2 | 1 | 3 | 13 |
| K14102 energy-converting hydrogenase A subunit K | 0 | 1 | 0 | 0 | 0 | 0 | 0 | 2 | 0 | 0 | 2 | 1 |
| K14103 energy-converting hydrogenase A subunit L | 0 | 1 | 0 | 0 | 0 | 0 | 2 | 0 | 0 | 0 | 2 | 1 |
| K14104 energy-converting hydrogenase A subunit M | 0 | 1 | 0 | 1 | 0 | 0 | 0 | 1 | 0 | 3 | 3 | 1 |
| K14105 energy-converting hydrogenase A subunit N | 0 | 0 | 0 | 1 | 0 | 0 | 0 | 0 | 4 | 7 | 3 | 6 |
| K14106 energy-converting hydrogenase A subunit O | 3 | 0 | 3 | 1 | 0 | 0 | 0 | 0 | 4 | 5 | 11 | 12 |
| K14107 energy-converting hydrogenase A subunit P | 0 | 0 | 2 | 1 | 1 | 0 | 0 | 0 | 0 | 5 | 5 | 9 |
| K14108 energy-converting hydrogenase A subunit Q | 2 | 0 | 1 | 3 | 2 | 0 | 0 | 0 | 2 | 17 | 8 | 7 |
| K14109 energy-converting hydrogenase A subunit R | 0 | 0 | 2 | 0 | 0 | 2 | 0 | 2 | 4 | 9 | 5 | 10 |
| K14110 energy-converting hydrogenase B subunit A | 0 | 0 | 0 | 0 | 0 | 0 | 0 | 1 | 3 | 5 | 0 | 4 |
| K14111 energy-converting hydrogenase B subunit B | 0 | 2 | 0 | 0 | 0 | 0 | 0 | 0 | 3 | 1 | 0 | 0 |
| K14112 energy-converting hydrogenase B subunit C | 0 | 0 | 0 | 1 | 0 | 1 | 0 | 3 | 2 | 1 | 1 | 0 |
| K14113 energy-converting hydrogenase B subunit D | 0 | 0 | 0 | 0 | 0 | 1 | 0 | 0 | 0 | 0 | 0 | 4 |
| K14114 energy-converting hydrogenase B subunit E | 0 | 0 | 0 | 0 | 0 | 0 | 0 | 2 | 0 | 4 | 3 | 2 |
| K14115 energy-converting hydrogenase B subunit F | 2 | 2 | 2 | 1 | 2 | 1 | 2 | 2 | 2 | 17 | 8 | 7 |
| K14116 energy-converting hydrogenase B subunit G | 0 | 0 | 2 | 0 | 0 | 0 | 0 | 1 | 0 | 0 | 0 | 1 |
| K14117 energy-converting hydrogenase B subunit H | 0 | 0 | 0 | 0 | 0 | 0 | 0 | 0 | 0 | 0 | 0 | 1 |
| K14118 energy-converting hydrogenase B subunit I | 0 | 0 | 0 | 1 | 0 | 0 | 0 | 0 | 4 | 1 | 3 | 4 |
| K14119 energy-converting hydrogenase B subunit J | 0 | 0 | 0 | 1 | 0 | 0 | 0 | 0 | 2 | 3 | 2 | 2 |
| K14120 energy-converting hydrogenase B subunit K | 0 | 2 | 0 | 1 | 0 | 2 | 1 | 2 | 3 | 10 | 8 | 13 |
| K14121 energy-converting hydrogenase B subunit L | 0 | 0 | 0 | 0 | 0 | 0 | 3 | 0 | 0 | 2 | 0 | 2 |
| K14122 energy-converting hydrogenase B subunit M | 0 | 0 | 0 | 0 | 0 | 0 | 1 | 2 | 0 | 0 | 0 | 1 |
| K14123 energy-converting hydrogenase B subunit N | 0 | 0 | 0 | 1 | 2 | 0 | 0 | 2 | 7 | 14 | 5 | 9 |
| K14124 energy-converting hydrogenase B subunit O | 0 | 0 | 2 | 0 | 0 | 4 | 0 | 0 | 5 | 7 | 2 | 3 |
| K14125 energy-converting hydrogenase B subunit P | 0 | 0 | 0 | 0 | 0 | 0 | 0 | 2 | 0 | 0 | 3 | 1 |
| K15536 soluble cytochrome b562 | 240 | 268 | 242 | 240 | 159 | 196 | 140 | 165 | 321 | 328 | 318 | 350 |
| K15790 nitrogen fixation protein NifQ | 105 | 93 | 104 | 118 | 157 | 163 | 132 | 158 | 2 | 0 | 0 | 0 |
| K15827 formate hydrogenlyase subunit 2 | 229 | 270 | 211 | 268 | 69 | 104 | 80 | 71 | 496 | 512 | 501 | 509 |
| K15828 formate hydrogenlyase subunit 3 | 737 | 794 | 738 | 790 | 208 | 233 | 220 | 227 | 1608 | 1565 | 1468 | 1523 |
| K15829 formate hydrogenlyase subunit 4 | 323 | 340 | 342 | 332 | 97 | 115 | 107 | 110 | 732 | 792 | 752 | 776 |
| K15830 formate hydrogenlyase subunit 5 | 695 | 716 | 592 | 726 | 220 | 187 | 199 | 221 | 1385 | 1455 | 1428 | 1481 |
| K15831 formate hydrogenlyase subunit 6 | 223 | 202 | 218 | 217 | 84 | 65 | 54 | 51 | 443 | 460 | 469 | 462 |
| K15832 formate hydrogenlyase subunit 7 | 295 | 319 | 294 | 330 | 99 | 117 | 99 | 81 | 634 | 675 | 589 | 643 |
| K16183 methylamine methyltransferase corrinoid activation protein | 0 | 0 | 0 | 0 | 0 | 0 | 0 | 1 | 2 | 0 | 1 | 1 |
| K16885 quinone-modifying oxidoreductase, subunit QmoA | 7 | 12 | 8 | 3 | 3 | 4 | 6 | 0 | 13 | 13 | 24 | 15 |
| K16886 quinone-modifying oxidoreductase, subunit QmoB | 13 | 16 | 13 | 17 | 3 | 13 | 8 | 3 | 28 | 32 | 15 | 31 |
| K16887 quinone-modifying oxidoreductase, subunit QmoC | 4 | 9 | 12 | 7 | 4 | 4 | 2 | 1 | 12 | 17 | 11 | 19 |
| K19344 cytochrome c55X | 340 | 371 | 313 | 378 | 197 | 197 | 174 | 178 | 124 | 110 | 89 | 120 |
| K19345 protein NirF | 1962 | 1923 | 1797 | 1896 | 1074 | 963 | 993 | 1129 | 649 | 515 | 588 | 633 |
| K19824 rubrerythrin | 0 | 0 | 0 | 0 | 0 | 0 | 2 | 0 | 1 | 4 | 4 | 4 |
| K21739 probable pyridine nucleotide-disulfide oxidoreductase | 315 | 330 | 367 | 296 | 98 | 82 | 98 | 102 | 476 | 508 | 505 | 458 |
| K23077 dissimilatory sulfite reductase related protein | 0 | 5 | 1 | 1 | 2 | 0 | 4 | 0 | 0 | 2 | 3 | 9 |
| K23302 menaquinone reductase, molybdopterin-binding-like subunit | 14 | 6 | 7 | 17 | 6 | 5 | 3 | 4 | 2 | 1 | 5 | 5 |
| K23303 menaquinone reductase, integral membrane subunit | 6 | 12 | 13 | 7 | 3 | 0 | 3 | 3 | 7 | 0 | 1 | 3 |
| ko99983 Lipid metabolism | 5661 | 5491 | 5247 | 5338 | 3110 | 3156 | 2915 | 3363 | 2883 | 2833 | 2881 | 2938 |
| K09456 putative acyl-CoA dehydrogenase | 3808 | 3802 | 3586 | 3613 | 2055 | 2168 | 2037 | 2324 | 2255 | 2283 | 2330 | 2352 |
| K11729 acyl-CoA dehydrogenase family member 10 | 3 | 5 | 2 | 1 | 2 | 0 | 1 | 1 | 1 | 2 | 3 | 2 |
| K11730 acyl-CoA dehydrogenase family member 11 | 2 | 0 | 0 | 0 | 0 | 0 | 0 | 0 | 0 | 0 | 0 | 0 |
| K16229 glycopeptidolipid biosynthesis protein | 1 | 0 | 0 | 1 | 0 | 0 | 0 | 0 | 0 | 0 | 0 | 0 |
| K21830 phosphorylcholine phosphatase | 1847 | 1684 | 1659 | 1723 | 1053 | 988 | 877 | 1038 | 627 | 548 | 548 | 584 |
| ko99984 Nucleotide metabolism | 442 | 456 | 365 | 422 | 216 | 225 | 235 | 219 | 423 | 448 | 390 | 434 |
| K03647 protein involved in ribonucleotide reduction | 239 | 254 | 191 | 201 | 166 | 161 | 183 | 154 | 322 | 278 | 251 | 316 |
| K15051 DNA-entry nuclease | 203 | 202 | 174 | 221 | 50 | 64 | 52 | 65 | 101 | 170 | 139 | 118 |
| ko99985 Amino acid metabolism | 8786 | 8702 | 8725 | 8805 | 5157 | 5479 | 5434 | 5242 | 10754 | 11042 | 10603 | 11020 |
| K04023 ethanolamine transporter | 935 | 968 | 891 | 952 | 337 | 359 | 382 | 326 | 1344 | 1419 | 1261 | 1349 |
| K04024 ethanolamine utilization protein EutJ | 560 | 611 | 650 | 585 | 165 | 206 | 184 | 142 | 886 | 869 | 833 | 869 |
| K04025 ethanolamine utilization protein EutK | 188 | 199 | 198 | 209 | 45 | 68 | 52 | 59 | 348 | 369 | 394 | 418 |
| K04026 ethanolamine utilization protein EutL | 316 | 353 | 341 | 327 | 93 | 88 | 103 | 97 | 619 | 719 | 689 | 714 |
| K04027 ethanolamine utilization protein EutM | 174 | 134 | 149 | 152 | 46 | 38 | 35 | 41 | 255 | 236 | 230 | 328 |
| K04028 ethanolamine utilization protein EutN | 304 | 297 | 288 | 274 | 62 | 73 | 90 | 75 | 403 | 443 | 431 | 434 |
| K04029 ethanolamine utilization protein EutP | 560 | 568 | 488 | 563 | 256 | 290 | 278 | 289 | 821 | 811 | 880 | 863 |
| K04030 ethanolamine utilization protein EutQ | 476 | 410 | 511 | 471 | 108 | 108 | 139 | 106 | 751 | 688 | 755 | 758 |
| K04031 ethanolamine utilization protein EutS | 521 | 505 | 473 | 536 | 215 | 245 | 256 | 192 | 721 | 789 | 708 | 694 |
| K05806 leu operon leader peptide | 15 | 11 | 16 | 14 | 24 | 30 | 29 | 24 | 3 | 2 | 3 | 4 |
| K06203 CysZ protein | 992 | 1003 | 1048 | 1041 | 600 | 644 | 688 | 701 | 771 | 951 | 872 | 866 |
| K06997 PLP dependent protein | 2492 | 2416 | 2403 | 2443 | 2487 | 2564 | 2523 | 2429 | 2526 | 2434 | 2382 | 2419 |
| K08234 glyoxylase I family protein | 351 | 333 | 368 | 376 | 211 | 261 | 194 | 198 | 583 | 550 | 495 | 547 |
| K10795 D-proline reductase (dithiol)-stabilizing protein PrdD | 0 | 0 | 0 | 0 | 0 | 0 | 0 | 0 | 0 | 1 | 1 | 1 |
| K10796 D-proline reductase (dithiol)-stabilizing protein PrdE | 1 | 0 | 2 | 1 | 0 | 0 | 0 | 1 | 1 | 0 | 0 | 2 |
| K11325 L-cysteine/cystine lyase | 20 | 16 | 9 | 17 | 0 | 0 | 0 | 0 | 13 | 16 | 8 | 10 |
| K14054 protein MpaA | 407 | 366 | 419 | 407 | 248 | 253 | 227 | 295 | 568 | 605 | 525 | 565 |
| K18802 glutamine amidotransferase | 0 | 0 | 2 | 0 | 0 | 0 | 0 | 0 | 0 | 0 | 0 | 0 |
| K21700 beta-alanine degradation protein BauB | 474 | 512 | 469 | 437 | 260 | 252 | 254 | 267 | 141 | 140 | 136 | 179 |
| ko99986 Glycan metabolism | 10957 | 10977 | 10358 | 10400 | 6818 | 6824 | 6637 | 7029 | 4686 | 4649 | 4306 | 4691 |
| K03208 colanic acid biosynthesis glycosyl transferase WcaI | 283 | 249 | 249 | 272 | 387 | 377 | 428 | 416 | 72 | 61 | 66 | 74 |
| K07270 glycosyl transferase, family 25 | 0 | 2 | 0 | 0 | 2 | 5 | 0 | 0 | 2 | 5 | 6 | 3 |
| K08992 lipopolysaccharide assembly protein A | 169 | 159 | 195 | 177 | 116 | 112 | 151 | 117 | 226 | 257 | 218 | 203 |
| K13620 putative colanic acid polymerase | 242 | 203 | 212 | 269 | 363 | 363 | 428 | 366 | 64 | 73 | 69 | 75 |
| K16568 exopolysaccharide production protein ExoZ | 0 | 0 | 0 | 3 | 0 | 0 | 0 | 2 | 0 | 1 | 4 | 2 |
| K16707 UDP-galactose-lipid carrier transferase | 0 | 0 | 0 | 0 | 0 | 0 | 0 | 0 | 1 | 1 | 1 | 0 |
| K16710 colanic acid/amylovoran biosynthesis protein | 254 | 275 | 254 | 211 | 352 | 415 | 369 | 384 | 61 | 69 | 55 | 53 |
| K16711 colanic acid biosynthesis protein WcaM | 237 | 293 | 303 | 287 | 407 | 458 | 414 | 406 | 46 | 68 | 63 | 66 |
| K19292 alginate biosynthesis protein AlgK | 2119 | 2251 | 1994 | 1996 | 1179 | 1054 | 1142 | 1199 | 686 | 570 | 620 | 666 |
| K19293 alginate biosynthesis protein AlgX | 2317 | 2281 | 2067 | 2044 | 1252 | 1245 | 1105 | 1257 | 657 | 649 | 566 | 716 |
| K19294 alginate O-acetyltransferase complex protein AlgI | 2905 | 2923 | 2859 | 2846 | 1401 | 1421 | 1227 | 1436 | 1339 | 1369 | 1214 | 1385 |

| | | | | | | | | | | | | |
|---|---|---|---|---|---|---|---|---|---|---|---|---|
| K19295 alginate O-acetyltransferase complex protein AlgJ | 940 | 820 | 815 | 776 | 445 | 450 | 437 | 498 | 265 | 233 | 257 | 258 |
| K19296 alginate O-acetyltransferase complex protein AlgF | 828 | 795 | 720 | 811 | 414 | 407 | 400 | 444 | 249 | 237 | 231 | 227 |
| K19421 polysaccharide biosynthesis protein EpsC | 3 | 5 | 4 | 3 | 0 | 0 | 0 | 0 | 9 | 2 | 11 | 6 |
| K19804 lipopolysaccharide assembly protein B | 660 | 720 | 686 | 705 | 500 | 517 | 536 | 503 | 1009 | 1054 | 923 | 957 |
| K21688 resuscitation-promoting factor RpfB | 0 | 1 | 0 | 0 | 0 | 0 | 0 | 0 | 1 | 0 | 0 | 2 | 0 |
| ko99987 Cofactor metabolism | 9552 | 9579 | 9355 | 9836 | 7410 | 7885 | 7472 | 7714 | 8999 | 8926 | 8830 | 9030 |
| K02173 putative kinase | 8 | 5 | 6 | 9 | 9 | 9 | 18 | 7 | 37 | 40 | 35 | 46 |
| K02498 HemY protein | 1673 | 1694 | 1649 | 1721 | 1018 | 1073 | 956 | 1053 | 1411 | 1322 | 1310 | 1286 |
| K02859 riboflavin biosynthesis RibT protein | 0 | 0 | 0 | 1 | 2 | 5 | 3 | 2 | 4 | 11 | 3 | 4 |
| K03688 ubiquinone biosynthesis protein | 3437 | 3350 | 3343 | 3506 | 2011 | 2166 | 1986 | 2102 | 2506 | 2505 | 2440 | 2672 |
| K03690 ubiquinone biosynthesis protein UbiJ | 1501 | 1498 | 1442 | 1507 | 903 | 909 | 818 | 920 | 839 | 915 | 880 | 968 |
| K03753 molybdopterin-guanine dinucleotide biosynthesis adapter protein | 348 | 364 | 358 | 346 | 263 | 321 | 296 | 273 | 491 | 500 | 488 | 528 |
| K12942 aminobenzoyl-glutamate transport protein | 1792 | 1920 | 1830 | 2002 | 2818 | 3046 | 3053 | 2965 | 3486 | 3403 | 3416 | 3301 |
| K13580 magnesium chelatase subunit ChlD-like protein | 793 | 748 | 727 | 744 | 386 | 356 | 342 | 392 | 225 | 229 | 254 | 223 |
| K18588 coenzyme Q-binding protein COQ10 | 0 | 0 | 0 | 0 | 0 | 0 | 0 | 0 | 0 | 1 | 4 | 2 |
| ko99988 Secondary metabolism | 8587 | 8248 | 7845 | 8055 | 5006 | 4968 | 4726 | 5012 | 5616 | 5568 | 5590 | 5798 |
| K03932 polyhydroxybutyrate depolymerase | 1713 | 1536 | 1419 | 1562 | 883 | 862 | 828 | 862 | 497 | 445 | 491 | 477 |
| K04753 suppressor of ftsI | 933 | 890 | 915 | 957 | 660 | 658 | 582 | 640 | 1174 | 1329 | 1272 | 1352 |
| K06219 S-adenosylmethionine-dependent methyltransferase | 1726 | 1709 | 1682 | 1670 | 1021 | 924 | 920 | 993 | 1251 | 1221 | 1243 | 1274 |
| K14588 blue copper oxidase | 888 | 899 | 861 | 929 | 657 | 704 | 667 | 687 | 1214 | 1270 | 1261 | 1256 |
| K18004 HQNO biosynthesis monooxygenase PqsL | 2094 | 2040 | 1780 | 1816 | 1056 | 1059 | 1014 | 1103 | 702 | 560 | 547 | 590 |
| K19784 chromate reductase, NAD(P)H dehydrogenase (quinone) | 1232 | 1174 | 1188 | 1121 | 729 | 759 | 715 | 727 | 778 | 743 | 776 | 849 |
| K23376 staphyloferrin B biosynthesis citrate synthase | 1 | 0 | 0 | 0 | 0 | 2 | 0 | 0 | 0 | 0 | 0 | 0 |
| ko99999 Others | 7232 | 7597 | 7270 | 7419 | 8553 | 8615 | 8619 | 8235 | 8972 | 8753 | 8903 | 9080 |
| K06999 phospholipase/carboxylesterase | 1532 | 1591 | 1478 | 1500 | 903 | 878 | 830 | 859 | 1010 | 991 | 977 | 1071 |
| K09181 acetyltransferase | 2728 | 2774 | 2786 | 2800 | 5107 | 5347 | 5341 | 4874 | 5500 | 5338 | 5534 | 5512 |
| K15975 glyoxalase family protein | 153 | 165 | 177 | 173 | 32 | 21 | 33 | 30 | 193 | 185 | 172 | 164 |
| K15977 putative oxidoreductase | 1315 | 1458 | 1401 | 1423 | 1623 | 1594 | 1601 | 1549 | 1785 | 1775 | 1745 | 1792 |
| K21831 cholinesterase | 1504 | 1609 | 1428 | 1523 | 888 | 775 | 814 | 923 | 484 | 464 | 475 | 541 |
| Unclassified: genetic information processing | 125734 | 126742 | 126387 | 127572 | 140283 | 141546 | 129488 | 132441 | 130762 | 129906 | 130795 | 129454 |
| ko99973 Transcription | 15183 | 15122 | 15143 | 15212 | 12004 | 12068 | 11761 | 11714 | 13002 | 12850 | 12937 | 13005 |
| K02466 glucitol operon activator protein | 236 | 226 | 205 | 275 | 156 | 200 | 206 | 168 | 357 | 356 | 309 | 341 |
| K02589 nitrogen regulatory protein PII 1 | 89 | 64 | 65 | 85 | 11 | 14 | 16 | 6 | 128 | 145 | 157 | 146 |
| K02590 nitrogen regulatory protein PII 2 | 70 | 65 | 64 | 60 | 12 | 3 | 12 | 11 | 124 | 133 | 118 | 120 |
| K03622 archaea-specific DNA-binding protein | 0 | 0 | 0 | 0 | 0 | 0 | 0 | 0 | 1 | 2 | 2 | 1 |
| K03623 ribonuclease inhibitor | 191 | 147 | 160 | 157 | 127 | 136 | 125 | 115 | 243 | 241 | 245 | 246 |
| K03627 putative transcription factor | 0 | 0 | 0 | 0 | 0 | 0 | 0 | 0 | 3 | 2 | 0 | 4 |
| K03704 cold shock protein | 3198 | 3242 | 3230 | 3209 | 1739 | 1778 | 1720 | 1675 | 2136 | 2203 | 2145 | 2261 |
| K04752 nitrogen regulatory protein P-II 2 | 828 | 845 | 826 | 839 | 528 | 577 | 496 | 553 | 523 | 542 | 480 | 490 |
| K04796 small nuclear ribonucleoprotein | 0 | 0 | 0 | 0 | 0 | 0 | 0 | 0 | 0 | 2 | 5 | 1 |
| K06959 protein Tex | 8174 | 8079 | 8125 | 8135 | 8233 | 8177 | 8038 | 7973 | 7530 | 7357 | 7519 | 7467 |
| K07591 threonine dehydratase operon activator protein | 2 | 2 | 1 | 2 | 2 | 5 | 3 | 7 | 22 | 21 | 23 | 28 |
| K07740 regulator of sigma D | 1251 | 1183 | 1199 | 1257 | 773 | 743 | 723 | 791 | 795 | 742 | 748 | 757 |
| K07741 anti-repressor protein | 780 | 846 | 914 | 817 | 270 | 292 | 289 | 314 | 313 | 294 | 326 | 314 |
| K07744 transcriptional regulator | 0 | 5 | 0 | 1 | 0 | 0 | 3 | 0 | 6 | 0 | 3 | 6 |
| K15833 formate hydrogenlyase regulatory protein HycA | 199 | 213 | 188 | 187 | 80 | 63 | 59 | 56 | 422 | 392 | 452 | 434 |
| K15834 formate hydrogenlyase maturation protein HycH | 143 | 174 | 146 | 162 | 53 | 63 | 45 | 39 | 364 | 371 | 359 | 353 |
| K16509 regulatory protein spx | 22 | 31 | 20 | 26 | 20 | 17 | 26 | 6 | 35 | 47 | 46 | 36 |
| ko99974 Translation | 7135 | 7095 | 7049 | 7456 | 3574 | 3570 | 3155 | 3681 | 4750 | 4677 | 4701 | 4977 |
| K07570 general stress protein 13 | 30 | 30 | 21 | 23 | 4 | 12 | 4 | 4 | 39 | 29 | 24 | 32 |
| K07571 S1 RNA binding domain protein | 43 | 51 | 79 | 56 | 26 | 25 | 42 | 29 | 46 | 58 | 54 | 79 |
| K07572 putative nucleotide binding protein | 0 | 0 | 0 | 0 | 0 | 0 | 0 | 0 | 1 | 1 | 1 | 6 |
| K07575 PUA domain protein | 0 | 0 | 0 | 0 | 1 | 0 | 0 | 0 | 2 | 5 | 2 | 7 |
| K07576 metallo-beta-lactamase family protein | 1451 | 1491 | 1410 | 1539 | 626 | 593 | 498 | 581 | 966 | 955 | 944 | 1033 |
| K07577 putative mRNA 3-end processing factor | 1 | 0 | 2 | 3 | 2 | 1 | 0 | 0 | 4 | 3 | 0 | 4 |
| K07579 putative methylase | 0 | 0 | 1 | 0 | 2 | 2 | 2 | 0 | 0 | 4 | 3 | 4 |
| K07580 Zn-ribbon RNA-binding protein | 0 | 0 | 0 | 1 | 0 | 0 | 0 | 0 | 0 | 0 | 0 | 1 |
| K07581 RNA-binding protein | 0 | 0 | 0 | 0 | 0 | 2 | 2 | 0 | 2 | 0 | 3 | 1 |
| K07585 tRNA methyltransferase | 0 | 0 | 0 | 0 | 0 | 1 | 0 | 2 | 0 | 5 | 2 | 4 |
| K13926 ribosome-dependent ATPase | 5610 | 5523 | 5536 | 5834 | 2913 | 2934 | 2607 | 3065 | 3690 | 3617 | 3668 | 3806 |
| ko99975 Protein processing | 39934 | 39715 | 38004 | 39530 | 28186 | 28122 | 27285 | 28845 | 30048 | 30436 | 29983 | 30432 |
| K02380 FdhE protein | 2172 | 2080 | 1804 | 2003 | 1326 | 1295 | 1121 | 1346 | 1298 | 1401 | 1301 | 1340 |
| K03187 urease accessory protein | 837 | 860 | 692 | 825 | 338 | 380 | 318 | 348 | 713 | 699 | 649 | 694 |
| K03188 urease accessory protein | 704 | 640 | 628 | 695 | 229 | 278 | 225 | 260 | 825 | 846 | 772 | 878 |
| K03189 urease accessory protein | 753 | 712 | 732 | 763 | 332 | 340 | 290 | 326 | 732 | 745 | 699 | 716 |
| K03190 urease accessory protein | 1661 | 1691 | 1567 | 1561 | 777 | 772 | 718 | 885 | 1071 | 1095 | 1133 | 1147 |
| K03192 urease accessory protein | 1014 | 927 | 789 | 829 | 468 | 462 | 464 | 566 | 283 | 248 | 216 | 294 |
| K03545 trigger factor | 2970 | 2978 | 3203 | 3027 | 4323 | 4187 | 4149 | 4091 | 4512 | 4673 | 4584 | 4511 |
| K03664 SsrA-binding protein | 1445 | 1452 | 1504 | 1503 | 1531 | 1572 | 1577 | 1569 | 1572 | 1574 | 1558 | 1503 |
| K03668 heat shock protein HslJ | 875 | 982 | 892 | 927 | 527 | 572 | 523 | 563 | 550 | 579 | 600 | 549 |
| K04063 lipoyl-dependent peroxidase | 1865 | 1814 | 1716 | 1835 | 1299 | 1194 | 1143 | 1380 | 847 | 847 | 806 | 840 |
| K04653 hydrogenase expression/formation protein HypC | 240 | 207 | 163 | 217 | 200 | 198 | 220 | 219 | 208 | 212 | 198 | 215 |
| K04654 hydrogenase expression/formation protein HypD | 892 | 832 | 798 | 831 | 569 | 633 | 633 | 666 | 1122 | 1180 | 1156 | 1184 |
| K04655 hydrogenase expression/formation protein HypE | 784 | 722 | 642 | 805 | 518 | 555 | 538 | 563 | 1014 | 1010 | 1103 | 987 |
| K04656 hydrogenase maturation protein HypF | 1584 | 1610 | 1452 | 1601 | 1084 | 1150 | 1095 | 1064 | 2073 | 2184 | 2055 | 2228 |
| K06191 glutaredoxin-like protein NrdH | 193 | 161 | 172 | 184 | 111 | 114 | 134 | 108 | 203 | 238 | 200 | 215 |
| K06891 ATP-dependent Clp protease adaptor protein ClpS | 603 | 555 | 572 | 559 | 322 | 364 | 291 | 325 | 388 | 411 | 457 | 386 |
| K07388 hydrogenase expression/formation protein | 0 | 3 | 0 | 1 | 0 | 0 | 2 | 0 | 4 | 11 | 8 | 10 |
| K07391 magnesium chelatase family protein | 5000 | 5358 | 4996 | 5242 | 5169 | 5203 | 5196 | 5379 | 4843 | 4585 | 4800 | 4818 |
| K07392 AAA family ATPase | 0 | 3 | 1 | 0 | 1 | 0 | 0 | 0 | 4 | 8 | 3 | 5 |
| K07394 SM-20-related protein | 998 | 922 | 829 | 974 | 509 | 527 | 517 | 546 | 292 | 318 | 280 | 329 |
| K07395 putative proteasome-type protease | 1543 | 1441 | 1519 | 1431 | 868 | 914 | 816 | 836 | 829 | 829 | 859 | 849 |
| K07396 putative protein-disulfide isomerase | 2293 | 2330 | 2132 | 2287 | 1385 | 1337 | 1298 | 1461 | 660 | 599 | 680 | 666 |
| K07397 putative redox protein | 1619 | 1547 | 1626 | 1695 | 991 | 939 | 889 | 868 | 769 | 743 | 659 | 765 |
| K07399 cytochrome c biogenesis protein | 2 | 0 | 1 | 0 | 0 | 0 | 0 | 0 | 0 | 0 | 0 | 0 |
| K07400 Fe/S biogenesis protein NfuA | 1065 | 997 | 1079 | 1079 | 636 | 645 | 617 | 643 | 702 | 724 | 722 | 684 |
| K07401 selenoprotein W-related protein | 614 | 674 | 593 | 615 | 369 | 337 | 315 | 372 | 423 | 434 | 361 | 426 |
| K07402 xanthine dehydrogenase accessory factor | 3681 | 3781 | 3636 | 3683 | 1946 | 1816 | 1901 | 1966 | 1680 | 1816 | 1688 | 1680 |
| K07403 membrane-bound serine protease (ClpP class) | 2182 | 2230 | 1918 | 2020 | 1090 | 1063 | 1109 | 1218 | 687 | 649 | 611 | 686 |
| K11208 GST-like protein | 303 | 319 | 329 | 326 | 239 | 259 | 187 | 275 | 469 | 520 | 442 | 479 |
| K16511 adapter protein MecA 1/2 | 453 | 403 | 473 | 413 | 110 | 97 | 112 | 95 | 287 | 344 | 357 | 358 |
| K17247 methionine sulfoxide reductase heme-binding subunit | 1366 | 1285 | 1346 | 1388 | 885 | 879 | 847 | 877 | 843 | 792 | 897 | 854 |
| K19411 protein arginine kinase activator | 223 | 199 | 200 | 211 | 34 | 40 | 40 | 30 | 145 | 122 | 129 | 136 |
| ko99976 Replication and repair | 62956 | 64271 | 65678 | 64812 | 96117 | 97350 | 86849 | 87782 | 82278 | 81169 | 82483 | 80293 |
| K01146 protein Xni | 334 | 361 | 356 | 381 | 300 | 312 | 326 | 338 | 407 | 493 | 441 | 493 |
| K02086 DNA replication protein | 40 | 49 | 39 | 37 | 4 | 5 | 14 | 12 | 30 | 61 | 44 | 37 |
| K03630 DNA repair protein RadC | 4666 | 4671 | 4599 | 4662 | 4330 | 4426 | 4258 | 4192 | 3804 | 3836 | 3747 | 3607 |
| K04096 DNA processing protein | 2482 | 2663 | 2427 | 2633 | 3419 | 3454 | 3585 | 3557 | 3576 | 3545 | 3314 | 3431 |
| K06909 phage terminase large subunit | 536 | 440 | 546 | 478 | 118 | 114 | 99 | 108 | 272 | 337 | 260 | 304 |
| K06919 putative DNA primase/helicase | 1894 | 1913 | 1949 | 1919 | 1590 | 1725 | 1574 | 1701 | 1041 | 1162 | 1199 | 1186 |
| K06993 ribonuclease H-related protein | 8 | 17 | 15 | 21 | 3 | 7 | 7 | 4 | 6 | 18 | 4 | 16 |
| K07357 type 1 fimbriae regulatory protein FimB | 254 | 246 | 275 | 300 | 84 | 90 | 82 | 72 | 529 | 596 | 537 | 575 |
| K07358 type 1 fimbriae regulatory protein FimE | 227 | 214 | 209 | 241 | 75 | 66 | 61 | 89 | 465 | 416 | 437 | 498 |
| K07443 methylated-DNA-protein-cysteine methyltransferase related protein | 1287 | 1315 | 1188 | 1313 | 1287 | 1216 | 1235 | 1229 | 1216 | 1113 | 1262 | 1080 |
| K07445 putative DNA methylase | 0 | 0 | 0 | 1 | 0 | 0 | 0 | 0 | 1 | 4 | 3 | 4 |
| K07449 similar to archaeal holliday junction resolvase and Mrr protein | 0 | 0 | 1 | 0 | 0 | 0 | 0 | 0 | 0 | 0 | 3 | 0 |
| K07453 putative restriction endonuclease | 1215 | 1177 | 1202 | 1184 | 694 | 661 | 601 | 673 | 387 | 370 | 394 | 401 |
| K07454 putative restriction endonuclease | 354 | 343 | 373 | 350 | 111 | 114 | 91 | 90 | 677 | 768 | 678 | 705 |
| K07457 endonuclease III related protein | 2 | 3 | 2 | 1 | 0 | 5 | 3 | 6 | 4 | 8 | 8 | 9 |
| K07459 putative ATP-dependent endonuclease of the OLD family | 1383 | 1211 | 1275 | 1304 | 694 | 761 | 765 | 799 | 1497 | 1557 | 1409 | 1602 |
| K07460 putative endonuclease | 1206 | 1125 | 1211 | 1149 | 1332 | 1438 | 1353 | 1419 | 1124 | 1133 | 1113 | 1116 |
| K07461 putative endonuclease | 843 | 785 | 809 | 835 | 449 | 476 | 459 | 444 | 594 | 574 | 572 | 611 |
| K07463 archaea-specific RecJ-like exonuclease | 2 | 3 | 3 | 3 | 0 | 2 | 0 | 0 | 13 | 19 | 3 | 15 |
| K07467 phage replication initiation protein | 242 | 220 | 232 | 278 | 382 | 373 | 409 | 335 | 531 | 554 | 499 | 477 |
| K07470 DNA gyrase inhibitor | 258 | 264 | 228 | 245 | 168 | 210 | 217 | 206 | 353 | 383 | 321 | 366 |
| K07474 phage terminase small subunit | 1043 | 1006 | 1024 | 1065 | 543 | 507 | 572 | 553 | 408 | 444 | 431 | 470 |
| K07478 putative ATPase | 4668 | 4590 | 4371 | 4582 | 4368 | 4351 | 4535 | 4295 | 4674 | 4577 | 4635 | 4600 |
| K07479 putative DNA topoisomerase | 334 | 354 | 329 | 401 | 262 | 268 | 271 | 261 | 507 | 506 | 517 | 475 |
| K07480 insertion element IS1 protein InsB | 487 | 520 | 510 | 484 | 713 | 762 | 730 | 666 | 63 | 88 | 83 | 95 |
| K07481 transposase, IS5 family | 788 | 886 | 856 | 843 | 941 | 965 | 997 | 999 | 213 | 243 | 228 | 203 |
| K07482 transposase, IS30 family | 0 | 3 | 1 | 1 | 2 | 3 | 3 | 2 | 1 | 0 | 0 | 4 |

| | | | | | | | | | | | | |
|---|---|---|---|---|---|---|---|---|---|---|---|---|
| K07483 transposase | 1577 | 1561 | 1677 | 1676 | 818 | 859 | 681 | 641 | 1312 | 1512 | 1355 | 1444 |
| K07484 transposase | 15019 | 16486 | 17586 | 16296 | 56980 | 57237 | 48110 | 48958 | 44299 | 42213 | 44864 | 41872 |
| K07486 transposase | 0 | 0 | 2 | 1 | 6 | 5 | 5 | 8 | 4 | 3 | 10 | 9 |
| K07487 transposase | 617 | 667 | 655 | 633 | 424 | 384 | 362 | 363 | 224 | 259 | 211 | 184 |
| K07488 transposase | 0 | 2 | 2 | 0 | 1 | 2 | 0 | 0 | 0 | 3 | 2 | 1 |
| K07489 transposase | 0 | 0 | 0 | 0 | 0 | 0 | 0 | 0 | 0 | 0 | 1 | 0 |
| K07491 putative transposase | 1064 | 1027 | 1017 | 1033 | 950 | 1033 | 943 | 1003 | 653 | 664 | 627 | 668 |
| K07492 putative transposase | 1 | 1 | 2 | 1 | 0 | 0 | 0 | 0 | 0 | 1 | 3 | 4 |
| K07493 putative transposase | 527 | 489 | 571 | 508 | 401 | 459 | 434 | 422 | 314 | 325 | 303 | 313 |
| K07495 putative transposase | 2 | 2 | 2 | 1 | 3 | 3 | 5 | 7 | 56 | 49 | 51 | 58 |
| K07496 putative transposase | 179 | 191 | 202 | 223 | 355 | 344 | 323 | 357 | 315 | 316 | 302 | 310 |
| K07497 putative transposase | 12498 | 12627 | 12716 | 12697 | 7720 | 8092 | 7462 | 7500 | 8336 | 8500 | 8214 | 8653 |
| K07498 putative transposase | 366 | 353 | 339 | 420 | 119 | 129 | 119 | 154 | 181 | 209 | 188 | 207 |
| K07499 putative transposase | 0 | 0 | 0 | 0 | 0 | 2 | 0 | 0 | 0 | 0 | 0 | 0 |
| K07501 3'-5' exonuclease | 0 | 2 | 0 | 1 | 2 | 0 | 1 | 0 | 1 | 0 | 0 | 0 |
| K07505 regulatory protein RepA | 1351 | 1385 | 1400 | 1341 | 783 | 816 | 764 | 754 | 458 | 483 | 506 | 506 |
| K09760 DNA recombination protein RmuC | 3554 | 3555 | 3754 | 3548 | 3958 | 3876 | 3718 | 3887 | 3297 | 3347 | 3239 | 3245 |
| K10762 putative replication protein | 0 | 0 | 0 | 1 | 0 | 2 | 1 | 0 | 2 | 0 | 5 | 0 |
| K14059 integrase | 1087 | 1011 | 1141 | 1152 | 904 | 911 | 932 | 907 | 275 | 302 | 305 | 278 |
| K14060 putative DNA-invertase from lambdoid prophage Rac | 182 | 202 | 211 | 235 | 284 | 339 | 277 | 280 | 112 | 129 | 128 | 124 |
| K18320 transposase, IS6 family | 379 | 331 | 371 | 334 | 540 | 546 | 465 | 491 | 46 | 49 | 27 | 37 |
| ko99998 Others | 526 | 539 | 513 | 562 | 402 | 436 | 438 | 419 | 684 | 774 | 691 | 747 |
| K03969 phage shock protein A | 335 | 356 | 357 | 411 | 241 | 280 | 262 | 221 | 450 | 570 | 501 | 533 |
| K03971 phage shock protein D | 135 | 112 | 103 | 110 | 75 | 76 | 76 | 86 | 176 | 158 | 156 | 184 |
| K03972 phage shock protein E | 53 | 69 | 51 | 41 | 80 | 66 | 89 | 98 | 30 | 23 | 11 | 10 |
| K07504 predicted type IV restriction endonuclease | 3 | 2 | 2 | 0 | 6 | 14 | 11 | 14 | 28 | 23 | 23 | 20 |
| Unclassified: signaling and cellular processes | 237972 | 237817 | 236931 | 239334 | 238163 | 240445 | 239553 | 240111 | 237762 | 238299 | 236919 | 238552 |
| ko99977 Transport | 97253 | 97592 | 97859 | 98594 | 107521 | 108423 | 108177 | 108675 | 107036 | 107130 | 106473 | 107028 |
| K03281 chloride channel protein, ClC family | 2389 | 2473 | 2551 | 2613 | 4587 | 4688 | 4631 | 4558 | 4831 | 4866 | 4806 | 4956 |
| K03285 general bacterial porin, GBP family | 31 | 28 | 34 | 33 | 46 | 56 | 43 | 44 | 46 | 66 | 64 | 54 |
| K03292 glycoside/pentoside/hexuronide:cation symporter, GPH family | 7894 | 8058 | 8396 | 8358 | 8592 | 9099 | 8797 | 8597 | 13208 | 13552 | 13427 | 13196 |
| K03293 amino acid transporter, AAT family | 3346 | 3279 | 3096 | 3410 | 1867 | 1934 | 1726 | 1963 | 2428 | 2552 | 2647 | 2497 |
| K03294 basic amino acid/polyamine antiporter, APA family | 2309 | 2495 | 2352 | 2391 | 1454 | 1515 | 1500 | 1503 | 1320 | 1367 | 1325 | 1432 |
| K03296 hydrophobic/amphiphilic exporter-1 (mainly G- bacteria), HAE1 family | 3315 | 3456 | 3722 | 3497 | 9140 | 9093 | 9294 | 9315 | 5694 | 5772 | 5707 | 5439 |
| K03300 citrate-Mg2+:H+ or citrate-Ca2+:H+ symporter, CitMHS family | 2168 | 2078 | 2159 | 2061 | 1151 | 1119 | 1029 | 1164 | 766 | 729 | 731 | 753 |
| K03305 proton-dependent oligopeptide transporter, POT family | 5195 | 5228 | 5336 | 5291 | 7097 | 7263 | 7464 | 7170 | 10519 | 10263 | 10203 | 10254 |
| K03306 inorganic phosphate transporter, PiT family | 5724 | 5602 | 5499 | 5631 | 4676 | 4763 | 4620 | 4842 | 3495 | 3168 | 3081 | 3184 |
| K03307 solute:Na+ symporter, SSS family | 9003 | 9079 | 8979 | 9136 | 12466 | 12498 | 12483 | 12230 | 10757 | 10607 | 10939 | 10700 |
| K03308 neurotransmitter:Na+ symporter, NSS family | 3965 | 3734 | 4017 | 3746 | 4022 | 4257 | 4481 | 4144 | 4913 | 5446 | 5236 | 5167 |
| K03309 dicarboxylate/amino acid:cation (Na+ or H+) symporter, DAACS family | 22 | 19 | 19 | 23 | 22 | 39 | 26 | 34 | 28 | 23 | 28 | 15 |
| K03310 alanine or glycine:cation symporter, AGCS family | 6660 | 6501 | 6504 | 6645 | 5871 | 5962 | 6067 | 5992 | 6675 | 6661 | 6599 | 6680 |
| K03311 branched-chain amino acid:cation transporter, LIVCS family | 3118 | 3088 | 2967 | 3142 | 1720 | 1764 | 1567 | 1789 | 2435 | 2448 | 2345 | 2580 |
| K03316 monovalent cation:H+ antiporter, CPA1 family | 5462 | 5420 | 5095 | 5649 | 3004 | 2861 | 2871 | 3195 | 4006 | 4030 | 3889 | 4015 |
| K03317 concentrative nucleoside transporter, CNT family | 1160 | 1091 | 1124 | 1086 | 564 | 623 | 576 | 602 | 1876 | 1760 | 1705 | 1983 |
| K03319 divalent anion:Na+ symporter, DASS family | 818 | 787 | 847 | 878 | 1053 | 1083 | 1074 | 1064 | 288 | 260 | 261 | 294 |
| K03328 polysaccharide transporter, PST family | 188 | 192 | 184 | 190 | 252 | 289 | 251 | 259 | 315 | 309 | 290 | 333 |
| K03451 betaine/carnitine transporter, BCCT family | 1198 | 1255 | 1185 | 1210 | 359 | 332 | 326 | 289 | 1405 | 1490 | 1489 | 1495 |
| K03453 bile acid:Na+ symporter, BASS family | 4057 | 4076 | 4073 | 4018 | 3819 | 3688 | 3608 | 3655 | 3078 | 2922 | 2950 | 2960 |
| K03455 monovalent cation:H+ antiporter-2, CPA2 family | 9837 | 9785 | 9947 | 9857 | 12870 | 12586 | 13185 | 13218 | 9703 | 9674 | 9497 | 9565 |
| K03457 nucleobase:cation symporter-1, NCS1 family | 8710 | 9204 | 8630 | 8750 | 5076 | 4901 | 4541 | 4979 | 4747 | 4393 | 4340 | 4614 |
| K03458 nucleobase:cation symporter-2, NCS2 family | 2371 | 2164 | 2202 | 2181 | 1162 | 1156 | 1102 | 1189 | 751 | 792 | 812 | 818 |
| K07011 rhamnosyltransferase | 0 | 0 | 0 | 0 | 1 | 5 | 0 | 0 | 0 | 0 | 0 | 1 |
| K07085 putative transport protein | 6241 | 6432 | 6970 | 6679 | 15436 | 15624 | 15869 | 15653 | 12419 | 12503 | 12752 | 12518 |
| K21990 formate-nitrite transporter family protein | 1883 | 1889 | 1790 | 1941 | 1115 | 1084 | 909 | 1103 | 1081 | 1171 | 1123 | 1249 |
| K23243 nickel/cobalt homeostasis protein | 189 | 179 | 181 | 178 | 99 | 141 | 137 | 124 | 252 | 306 | 227 | 276 |
| ko99978 Cell growth | 16189 | 16041 | 15938 | 16131 | 8820 | 9372 | 8971 | 8371 | 14171 | 15347 | 14813 | 15262 |
| K06190 intracellular septation protein | 1044 | 945 | 951 | 1038 | 600 | 718 | 579 | 580 | 560 | 559 | 601 | 578 |
| K06287 septum formation protein | 2035 | 2116 | 2163 | 2066 | 2157 | 2254 | 2200 | 2104 | 2116 | 2134 | 2020 | 2137 |
| K06295 spore germination protein KA | 32 | 19 | 15 | 37 | 9 | 3 | 10 | 4 | 21 | 24 | 25 | 27 |
| K06297 spore germination protein KC | 5 | 3 | 2 | 5 | 2 | 0 | 4 | 2 | 3 | 9 | 3 | 4 |
| K06298 germination protein M | 66 | 94 | 76 | 84 | 40 | 40 | 40 | 42 | 85 | 155 | 100 | 128 |
| K06305 spore germination protein Q | 0 | 0 | 0 | 0 | 0 | 0 | 0 | 0 | 0 | 1 | 0 | 0 |
| K06306 spore germination protein | 667 | 677 | 671 | 673 | 144 | 157 | 139 | 135 | 408 | 510 | 532 | 520 |
| K06308 spore germination protein | 3 | 2 | 0 | 0 | 0 | 0 | 0 | 0 | 0 | 0 | 0 | 0 |
| K06310 spore germination protein | 70 | 58 | 58 | 66 | 3 | 6 | 2 | 0 | 204 | 187 | 157 | 211 |
| K06313 spore germination protein | 2 | 0 | 0 | 0 | 0 | 0 | 0 | 0 | 0 | 0 | 0 | 0 |
| K06317 inhibitor of the pro-sigma K processing machinery | 18 | 12 | 15 | 14 | 8 | 16 | 13 | 5 | 26 | 18 | 43 | 27 |
| K06320 spore maturation protein CgeB | 6 | 16 | 8 | 11 | 1 | 5 | 6 | 3 | 15 | 41 | 29 | 21 |
| K06327 spore coat protein D | 0 | 0 | 0 | 0 | 0 | 0 | 0 | 1 | 0 | 0 | 0 | 0 |
| K06330 spore coat protein H | 8 | 6 | 7 | 12 | 8 | 10 | 3 | 3 | 5 | 5 | 8 | 6 |
| K06333 spore coat protein JB | 282 | 276 | 237 | 261 | 38 | 87 | 95 | 49 | 230 | 251 | 238 | 291 |
| K06334 spore coat protein JC | 527 | 502 | 495 | 530 | 132 | 142 | 158 | 117 | 557 | 602 | 570 | 555 |
| K06338 spore coat protein SA | 19 | 20 | 25 | 22 | 57 | 76 | 91 | 77 | 45 | 51 | 52 | 71 |
| K06344 spore coat protein Z | 3 | 0 | 0 | 0 | 0 | 0 | 0 | 2 | 0 | 0 | 0 | 0 |
| K06346 spoIIIJ-associated protein | 499 | 536 | 572 | 518 | 122 | 183 | 175 | 146 | 443 | 512 | 500 | 530 |
| K06348 sporulation inhibitor KapD | 11 | 13 | 13 | 19 | 0 | 0 | 0 | 0 | 20 | 28 | 38 | 26 |
| K06349 KinB signaling pathway activation protein | 0 | 0 | 0 | 0 | 0 | 0 | 0 | 2 | 0 | 0 | 0 | 0 |
| K06350 antagonist of KipI | 1 | 2 | 0 | 1 | 8 | 9 | 7 | 10 | 2 | 2 | 0 | 1 |
| K06351 inhibitor of KinA | 0 | 2 | 0 | 0 | 0 | 0 | 1 | 0 | 0 | 1 | 0 | 1 |
| K06373 spore maturation protein A | 342 | 333 | 346 | 310 | 106 | 111 | 82 | 98 | 276 | 312 | 301 | 308 |
| K06374 spore maturation protein B | 359 | 279 | 283 | 308 | 95 | 106 | 108 | 82 | 221 | 273 | 253 | 270 |
| K06378 stage II sporulation protein AA (anti-sigma F factor antagonist) | 153 | 129 | 144 | 142 | 46 | 59 | 50 | 39 | 124 | 97 | 93 | 138 |
| K06381 stage II sporulation protein D | 628 | 586 | 554 | 574 | 162 | 196 | 209 | 168 | 563 | 560 | 526 | 574 |
| K06384 stage II sporulation protein M | 140 | 108 | 111 | 128 | 68 | 72 | 57 | 33 | 109 | 113 | 141 | 132 |
| K06385 stage II sporulation protein P | 373 | 386 | 347 | 372 | 109 | 138 | 130 | 131 | 255 | 357 | 356 | 293 |
| K06386 stage II sporulation protein Q | 0 | 1 | 0 | 1 | 0 | 0 | 0 | 0 | 2 | 0 | 0 | 1 |
| K06387 stage II sporulation protein R | 306 | 296 | 262 | 308 | 62 | 64 | 81 | 71 | 263 | 276 | 254 | 251 |
| K06390 stage III sporulation protein AA | 526 | 480 | 456 | 519 | 134 | 140 | 124 | 122 | 347 | 346 | 377 | 369 |
| K06391 stage III sporulation protein AB | 200 | 193 | 171 | 168 | 48 | 40 | 48 | 48 | 129 | 162 | 151 | 179 |
| K06392 stage III sporulation protein AC | 86 | 87 | 90 | 90 | 28 | 24 | 34 | 22 | 71 | 80 | 52 | 81 |
| K06393 stage III sporulation protein AD | 188 | 165 | 177 | 177 | 46 | 67 | 46 | 30 | 152 | 167 | 140 | 147 |
| K06394 stage III sporulation protein AE | 444 | 410 | 443 | 392 | 118 | 111 | 101 | 122 | 299 | 316 | 348 | 353 |
| K06395 stage III sporulation protein AF | 162 | 132 | 174 | 183 | 42 | 41 | 56 | 38 | 100 | 107 | 121 | 125 |
| K06396 stage III sporulation protein AG | 220 | 199 | 199 | 214 | 44 | 71 | 67 | 54 | 159 | 153 | 129 | 133 |
| K06397 stage III sporulation protein AH | 343 | 317 | 306 | 312 | 81 | 91 | 84 | 64 | 264 | 237 | 255 | 261 |
| K06398 stage IV sporulation protein A | 642 | 667 | 741 | 704 | 217 | 219 | 231 | 203 | 565 | 756 | 579 | 630 |
| K06400 site-specific DNA recombinase | 83 | 93 | 83 | 91 | 113 | 123 | 129 | 113 | 35 | 68 | 63 | 71 |
| K06403 stage V sporulation protein AA | 238 | 249 | 259 | 276 | 28 | 65 | 57 | 42 | 239 | 215 | 246 | 270 |
| K06404 stage V sporulation protein AB | 146 | 169 | 161 | 127 | 35 | 46 | 29 | 29 | 116 | 135 | 126 | 141 |
| K06405 stage V sporulation protein AC | 213 | 226 | 201 | 200 | 91 | 81 | 69 | 74 | 177 | 219 | 199 | 198 |
| K06406 stage V sporulation protein AD | 431 | 493 | 480 | 463 | 176 | 149 | 149 | 135 | 470 | 519 | 480 | 478 |
| K06407 stage V sporulation protein AE | 193 | 169 | 204 | 170 | 71 | 47 | 69 | 53 | 152 | 156 | 148 | 107 |
| K06408 stage V sporulation protein AF | 597 | 648 | 619 | 595 | 172 | 185 | 212 | 152 | 344 | 444 | 355 | 353 |
| K06410 dipicolinate synthase subunit A | 258 | 263 | 248 | 269 | 50 | 41 | 38 | 39 | 206 | 230 | 220 | 230 |
| K06411 dipicolinate synthase subunit B | 197 | 204 | 205 | 238 | 49 | 59 | 41 | 49 | 158 | 159 | 226 | 194 |
| K06412 stage V sporulation protein G | 233 | 256 | 237 | 262 | 110 | 96 | 105 | 81 | 144 | 167 | 218 | 178 |
| K06413 stage V sporulation protein K | 0 | 0 | 0 | 0 | 0 | 0 | 0 | 0 | 0 | 1 | 0 | 0 |
| K06415 stage V sporulation protein R | 30 | 30 | 26 | 26 | 23 | 33 | 29 | 29 | 41 | 55 | 50 | 44 |
| K06416 stage V sporulation protein S | 6 | 5 | 5 | 4 | 2 | 2 | 3 | 0 | 8 | 8 | 4 | 7 |
| K06421 small acid-soluble spore protein D (minor alpha/beta-type SASP) | 0 | 0 | 0 | 0 | 0 | 1 | 0 | 0 | 0 | 0 | 0 | 0 |
| K06436 spore coat assemly protein | 3 | 5 | 9 | 0 | 0 | 0 | 2 | 0 | 4 | 8 | 5 | 11 |
| K06438 similar to stage IV sporulation protein | 416 | 362 | 395 | 404 | 156 | 144 | 146 | 123 | 331 | 374 | 389 | 417 |
| K07257 spore coat polysaccharide biosynthesis protein SpsF | 8 | 14 | 5 | 13 | 6 | 6 | 5 | 0 | 11 | 17 | 14 | 22 |
| K07321 CO dehydrogenase maturation factor | 125 | 143 | 115 | 88 | 85 | 48 | 88 | 77 | 283 | 269 | 284 | 307 |
| K07322 regulator of cell morphogenesis and NO signaling | 977 | 1009 | 1051 | 978 | 1856 | 1867 | 1798 | 1835 | 1518 | 1506 | 1484 | 1528 |
| K15722 cell division activator | 132 | 159 | 112 | 128 | 102 | 98 | 119 | 102 | 178 | 226 | 199 | 208 |
| K20073 mid-cell-anchored protein Z | 18 | 22 | 19 | 15 | 6 | 2 | 3 | 7 | 14 | 22 | 19 | 13 |
| K22304 cell division inhibition protein DicB | 1 | 0 | 1 | 0 | 1 | 3 | 3 | 3 | 10 | 7 | 7 | 6 |
| K22718 divisome-associated membrane protein | 64 | 65 | 59 | 54 | 41 | 63 | 70 | 57 | 82 | 84 | 97 | 88 |
| K22719 murein hydrolase activator | 1410 | 1390 | 1332 | 1471 | 912 | 957 | 776 | 764 | 1011 | 1056 | 988 | 1012 |
| ko99993 Cell motility | 4395 | 4287 | 4261 | 4313 | 2734 | 2849 | 2734 | 2733 | 3810 | 3953 | 3754 | 3855 |

| | | | | | | | | | | | | |
|---|---|---|---|---|---|---|---|---|---|---|---|---|
| K07335 basic membrane protein A and related proteins | 1860 | 1729 | 1839 | 1820 | 709 | 733 | 644 | 635 | 1445 | 1601 | 1582 | 1516 |
| K07337 penicillin-binding protein activator | 304 | 290 | 303 | 316 | 197 | 238 | 200 | 267 | 406 | 418 | 358 | 423 |
| K07343 DNA transformation protein and related proteins | 454 | 433 | 435 | 462 | 282 | 296 | 302 | 269 | 589 | 611 | 550 | 544 |
| K08086 pilus assembly protein FimV | 474 | 488 | 435 | 432 | 263 | 274 | 215 | 253 | 193 | 170 | 177 | 173 |
| K08087 fimbrial protein FimW | 0 | 0 | 0 | 0 | 0 | 2 | 2 | 2 | 1 | 0 | 0 | 0 |
| K15540 chaperone protein EcpD | 882 | 901 | 868 | 859 | 951 | 990 | 994 | 984 | 650 | 644 | 639 | 667 |
| K15723 SecY interacting protein Syd | 312 | 295 | 260 | 289 | 230 | 211 | 261 | 231 | 348 | 332 | 321 | 370 |
| K22720 surface composition regulator | 109 | 151 | 121 | 135 | 102 | 105 | 116 | 92 | 178 | 177 | 127 | 162 |
| ko99995 Signaling proteins | 38993 | 38110 | 38088 | 38920 | 30353 | 30602 | 30246 | 30102 | 32205 | 32468 | 32201 | 33127 |
| K03607 ProP effector | 513 | 461 | 466 | 521 | 259 | 342 | 284 | 282 | 439 | 469 | 440 | 488 |
| K03807 AmpE protein | 1961 | 1924 | 1860 | 1857 | 1043 | 1095 | 1120 | 1090 | 1138 | 1089 | 1140 | 1254 |
| K05791 tellurium resistance protein TerZ | 9 | 5 | 4 | 7 | 3 | 5 | 1 | 3 | 0 | 0 | 0 | 1 |
| K05792 tellurite resistance protein TerA | 0 | 0 | 0 | 0 | 0 | 1 | 3 | 3 | 2 | 0 | 0 | 1 |
| K05795 tellurium resistance protein TerD | 4 | 4 | 12 | 6 | 47 | 35 | 51 | 35 | 1 | 1 | 2 | 0 |
| K06149 universal stress protein A | 923 | 826 | 961 | 954 | 527 | 656 | 568 | 539 | 574 | 548 | 606 | 559 |
| K06200 carbon starvation protein | 6063 | 5852 | 5753 | 5863 | 3291 | 3241 | 3277 | 3396 | 3986 | 3996 | 4005 | 4130 |
| K06207 GTP-binding protein | 6026 | 6126 | 6272 | 6245 | 7201 | 7211 | 7165 | 6793 | 6744 | 6871 | 7093 | 6948 |
| K06217 phosphate starvation-inducible protein PhoH and related proteins | 4141 | 4035 | 4046 | 4045 | 3983 | 4084 | 3992 | 3952 | 4164 | 4152 | 4159 | 4200 |
| K06875 programmed cell death protein 5 | 0 | 0 | 0 | 1 | 1 | 0 | 0 | 0 | 0 | 0 | 1 | 1 |
| K07166 ACT domain-containing protein | 184 | 131 | 163 | 145 | 47 | 60 | 42 | 43 | 142 | 217 | 166 | 180 |
| K07167 putative transcriptional regulator | 0 | 1 | 0 | 0 | 0 | 0 | 0 | 0 | 0 | 2 | 4 | 0 |
| K07168 CBS domain-containing membrane protein | 1851 | 1822 | 1713 | 1821 | 910 | 936 | 883 | 903 | 1087 | 974 | 922 | 1088 |
| K07169 FHA domain-containing protein | 345 | 316 | 292 | 335 | 484 | 555 | 526 | 470 | 0 | 2 | 0 | 0 |
| K07175 PhoH-like ATPase | 2388 | 2385 | 2391 | 2450 | 3515 | 3369 | 3430 | 3512 | 2674 | 2624 | 2659 | 2642 |
| K07180 serine protein kinase | 3751 | 3493 | 3811 | 3952 | 2267 | 2278 | 2234 | 2294 | 2407 | 2413 | 2358 | 2366 |
| K07182 CBS domain-containing protein | 2864 | 2895 | 2684 | 2751 | 1547 | 1377 | 1467 | 1485 | 873 | 823 | 823 | 864 |
| K07184 SH3 domain protein | 1234 | 1128 | 1195 | 1117 | 728 | 719 | 721 | 751 | 696 | 723 | 649 | 706 |
| K07186 membrane protein | 366 | 342 | 369 | 390 | 284 | 286 | 278 | 263 | 466 | 542 | 529 | 647 |
| K07212 GGDEF domain | 7 | 6 | 6 | 9 | 0 | 0 | 0 | 0 | 4 | 4 | 6 | 5 |
| K09767 cyclic-di-GMP-binding protein | 1093 | 1175 | 1105 | 1148 | 661 | 602 | 623 | 621 | 744 | 778 | 730 | 827 |
| K11932 universal stress protein G | 539 | 512 | 481 | 536 | 288 | 330 | 261 | 303 | 862 | 960 | 891 | 911 |
| K13630 multiple antibiotic resistance protein MarB | 122 | 104 | 90 | 118 | 67 | 80 | 72 | 72 | 156 | 181 | 115 | 150 |
| K14055 universal stress protein E | 1593 | 1492 | 1460 | 1486 | 948 | 965 | 881 | 936 | 1019 | 976 | 941 | 974 |
| K14061 universal stress protein F | 253 | 238 | 190 | 245 | 159 | 152 | 171 | 157 | 300 | 289 | 281 | 292 |
| K14064 universal stress protein C | 233 | 243 | 264 | 285 | 167 | 181 | 158 | 181 | 407 | 402 | 440 | 378 |
| K14065 universal stress protein D | 2 | 1 | 2 | 2 | 3 | 2 | 9 | 4 | 31 | 28 | 26 | 39 |
| K19336 cyclic-di-GMP-binding biofilm dispersal mediator protein | 271 | 279 | 224 | 270 | 68 | 94 | 70 | 75 | 529 | 569 | 541 | 549 |
| K21637 anti-adapter protein IraD | 1 | 1 | 2 | 0 | 3 | 12 | 6 | 2 | 27 | 18 | 15 | 29 |
| K21638 anti-adapter protein IraM | 111 | 127 | 96 | 128 | 43 | 32 | 40 | 36 | 206 | 247 | 241 | 257 |
| K21973 blue light- and temperature-responsive anti-repressor | 1281 | 1373 | 1297 | 1309 | 913 | 989 | 978 | 996 | 1703 | 1827 | 1708 | 1882 |
| K21974 probable RcsB/C two-component-system connector | 260 | 221 | 246 | 223 | 206 | 226 | 229 | 254 | 264 | 223 | 232 | 224 |
| K21975 probable RcsB/C two-component-system connector | 302 | 301 | 330 | 349 | 358 | 372 | 342 | 337 | 318 | 293 | 283 | 314 |
| K21976 probable RcsB/C two-component-system connector, global regulator of biofilm formation and acid-resistance | 302 | 291 | 303 | 352 | 332 | 315 | 364 | 314 | 242 | 227 | 195 | 221 |
| ko99992 Structural proteins | 50112 | 50621 | 50511 | 50748 | 65251 | 65764 | 65890 | 66423 | 57775 | 56668 | 57542 | 56604 |
| K01993 HlyD family secretion protein | 4068 | 4114 | 4110 | 4006 | 6672 | 6846 | 7012 | 6995 | 6564 | 6174 | 6150 | 6292 |
| K02005 HlyD family secretion protein | 3239 | 3547 | 3669 | 3415 | 7825 | 7915 | 7704 | 7896 | 5876 | 5582 | 6107 | 5677 |
| K02022 HlyD family secretion protein | 7 | 12 | 4 | 19 | 0 | 2 | 2 | 3 | 7 | 8 | 3 | 4 |
| K02197 cytochrome c-type biogenesis protein CcmE | 826 | 842 | 800 | 913 | 393 | 420 | 436 | 448 | 540 | 563 | 579 | 587 |
| K02200 cytochrome c-type biogenesis protein CcmH | 2638 | 2614 | 2337 | 2404 | 1348 | 1314 | 1337 | 1683 | 1264 | 1186 | 1175 | 1201 |
| K02494 outer membrane lipoprotein LolB | 1471 | 1396 | 1301 | 1438 | 842 | 849 | 794 | 836 | 758 | 792 | 781 | 832 |
| K02497 HemX protein | 1 | 0 | 0 | 0 | 0 | 0 | 0 | 0 | 0 | 0 | 0 | 0 |
| K03634 outer membrane lipoprotein carrier protein | 1189 | 1205 | 1162 | 1172 | 649 | 665 | 596 | 625 | 697 | 769 | 696 | 787 |
| K03642 rare lipoprotein A | 1956 | 1882 | 1878 | 1997 | 1173 | 1198 | 1085 | 1188 | 1195 | 1146 | 1129 | 1193 |
| K04062 osmotically inducible lipoprotein OsmB | 100 | 85 | 86 | 96 | 73 | 87 | 65 | 95 | 144 | 130 | 114 | 128 |
| K04064 osmotically inducible lipoprotein OsmE | 744 | 768 | 713 | 734 | 439 | 454 | 425 | 465 | 444 | 476 | 464 | 450 |
| K04754 phospholipid-binding lipoprotein MlaA | 2299 | 2256 | 2364 | 2307 | 1442 | 1349 | 1401 | 1296 | 1286 | 1313 | 1267 | 1386 |
| K05790 lipopolysaccharide biosynthesis protein WzzE | 729 | 634 | 640 | 733 | 514 | 560 | 490 | 530 | 990 | 942 | 915 | 885 |
| K05803 lipoprotein NlpI | 555 | 539 | 498 | 570 | 408 | 458 | 416 | 405 | 728 | 744 | 758 | 732 |
| K06077 outer membrane lipoprotein SlyB | 723 | 688 | 631 | 675 | 384 | 370 | 331 | 411 | 394 | 381 | 378 | 359 |
| K06142 outer membrane protein | 2185 | 2253 | 2349 | 2266 | 4500 | 4412 | 4234 | 4280 | 3277 | 3374 | 3235 | 3211 |
| K06143 inner membrane protein | 3610 | 3693 | 3747 | 3633 | 4036 | 3886 | 4180 | 4077 | 3438 | 3348 | 3473 | 3370 |
| K06194 lipoprotein NlpD | 1730 | 1645 | 1652 | 1723 | 1014 | 1160 | 1127 | 1217 | 1155 | 1196 | 1112 | 1114 |
| K07265 capsular polysaccharide export protein | 240 | 252 | 266 | 278 | 436 | 516 | 345 | 414 | 71 | 57 | 41 | 52 |
| K07266 capsular polysaccharide export protein | 459 | 456 | 481 | 465 | 617 | 765 | 708 | 622 | 68 | 81 | 75 | 90 |
| K07273 lysozyme | 704 | 727 | 736 | 747 | 1930 | 1920 | 2065 | 1927 | 1585 | 1527 | 1580 | 1609 |
| K07275 outer membrane protein | 616 | 542 | 519 | 595 | 402 | 370 | 355 | 349 | 604 | 550 | 524 | 568 |
| K07282 gamma-polyglutamate biosynthesis protein CapA | 0 | 0 | 0 | 0 | 0 | 0 | 0 | 0 | 1 | 0 | 0 | 0 |
| K07283 putative salt-induced outer membrane protein | 351 | 406 | 457 | 438 | 337 | 319 | 303 | 273 | 530 | 553 | 487 | 589 |
| K07285 outer membrane lipoprotein | 293 | 359 | 357 | 321 | 224 | 286 | 246 | 254 | 460 | 508 | 495 | 558 |
| K07286 uncharacterized lipoprotein | 384 | 410 | 424 | 388 | 265 | 301 | 296 | 274 | 573 | 512 | 566 | 556 |
| K07289 AsmA protein | 4345 | 4371 | 4273 | 4353 | 2682 | 2649 | 2653 | 2824 | 2617 | 2651 | 2578 | 2651 |
| K07290 AsmA family protein | 4582 | 4537 | 4297 | 4701 | 2768 | 2800 | 2764 | 2952 | 2818 | 2772 | 2781 | 2816 |
| K07340 inner membrane protein | 1051 | 1037 | 876 | 987 | 643 | 581 | 588 | 567 | 512 | 534 | 571 | 514 |
| K11744 AI-2 transport protein TqsA | 357 | 372 | 361 | 353 | 132 | 116 | 118 | 137 | 778 | 769 | 760 | 829 |
| K12943 lipoprotein YgeR | 976 | 975 | 884 | 955 | 581 | 651 | 543 | 603 | 804 | 705 | 769 | 774 |
| K13665 pyruvyltransferase | 0 | 0 | 3 | 0 | 0 | 0 | 0 | 0 | 0 | 0 | 0 | 0 |
| K15539 cytoskeleton protein RodZ | 1522 | 1413 | 1424 | 1601 | 920 | 903 | 1009 | 1011 | 870 | 904 | 774 | 933 |
| K19061 macrolide resistance protein | 0 | 0 | 0 | 0 | 3 | 1 | 0 | 3 | 1 | 0 | 3 | 0 |
| K19525 vacuolar protein sorting-associated protein 13A/C | 0 | 0 | 0 | 0 | 0 | 0 | 0 | 0 | 0 | 2 | 1 | 0 |
| K21571 starch-binding outer membrane protein SusE/F | 5804 | 6228 | 6844 | 6136 | 21425 | 21440 | 22049 | 21602 | 16075 | 15701 | 16460 | 15149 |
| K21695 protein AaeX | 134 | 134 | 151 | 124 | 101 | 122 | 123 | 102 | 176 | 256 | 239 | 200 |
| K21908 membrane protein HdeD | 221 | 227 | 216 | 203 | 73 | 79 | 90 | 58 | 475 | 462 | 502 | 508 |
| K22116 gamma-polyglutamate biosynthesis protein CapC | 3 | 2 | 1 | 2 | 0 | 0 | 0 | 1 | 0 | 0 | 0 | 0 |
| ko99994 Others | 26899 | 26847 | 26169 | 26270 | 18478 | 18344 | 18317 | 18616 | 17515 | 17444 | 17123 | 17624 |
| K02039 phosphate transport system protein | 2262 | 2331 | 2315 | 2307 | 2363 | 2348 | 2459 | 2373 | 2513 | 2456 | 2518 | 2494 |
| K02192 bacterioferritin-associated ferredoxin | 478 | 450 | 473 | 467 | 293 | 276 | 268 | 270 | 297 | 304 | 268 | 276 |
| K02255 ferritin-like protein 2 | 280 | 259 | 274 | 241 | 197 | 177 | 197 | 180 | 410 | 464 | 412 | 415 |
| K03670 periplasmic glucans biosynthesis protein | 858 | 822 | 853 | 857 | 600 | 679 | 602 | 559 | 1157 | 1266 | 1137 | 1298 |
| K03926 periplasmic divalent cation tolerance protein | 185 | 174 | 197 | 224 | 144 | 120 | 141 | 138 | 266 | 245 | 256 | 241 |
| K05781 putative phosphonate transport system ATP-binding protein | 1039 | 1026 | 915 | 1040 | 420 | 386 | 431 | 431 | 1053 | 1039 | 935 | 978 |
| K05794 tellurite resistance protein TerC | 1400 | 1384 | 1395 | 1287 | 818 | 787 | 748 | 786 | 430 | 433 | 382 | 406 |
| K06039 uncharacterized protein involved in oxidation of intracellular sulfur | 161 | 196 | 172 | 187 | 139 | 124 | 136 | 112 | 248 | 268 | 220 | 242 |
| K06195 ApaG protein | 823 | 777 | 796 | 815 | 482 | 551 | 517 | 517 | 469 | 523 | 493 | 461 |
| K06201 copper homeostasis protein | 1104 | 1116 | 998 | 1071 | 1671 | 1698 | 1760 | 1718 | 1698 | 1611 | 1865 | 1662 |
| K06202 CyaY protein | 600 | 695 | 600 | 590 | 409 | 418 | 421 | 431 | 420 | 428 | 450 | 503 |
| K07214 enterochelin esterase and related enzymes | 1910 | 1905 | 1857 | 1960 | 1709 | 1746 | 1687 | 1770 | 2175 | 2287 | 2276 | 2434 |
| K07216 hemerythrin | 626 | 642 | 676 | 606 | 338 | 308 | 337 | 340 | 262 | 289 | 283 | 238 |
| K07217 Mn-containing catalase | 463 | 520 | 545 | 510 | 354 | 372 | 395 | 340 | 744 | 780 | 737 | 729 |
| K07218 nitrous oxidase accessory protein | 1827 | 1714 | 1758 | 1564 | 961 | 865 | 773 | 911 | 624 | 543 | 537 | 478 |
| K07219 putative molybdopterin biosynthesis protein | 0 | 0 | 2 | 0 | 3 | 2 | 5 | 2 | 0 | 4 | 0 | 4 |
| K07222 putative flavoprotein involved in K+ transport | 2279 | 2366 | 2180 | 2135 | 1466 | 1474 | 1353 | 1514 | 709 | 694 | 607 | 692 |
| K07225 putative hemin transport protein | 2322 | 2358 | 2153 | 2288 | 1378 | 1279 | 1301 | 1293 | 1427 | 1395 | 1317 | 1476 |
| K07226 heme iron utilization protein | 1199 | 1151 | 1215 | 1165 | 648 | 714 | 652 | 678 | 358 | 338 | 372 | 409 |
| K07227 heme iron utilization protein | 8 | 5 | 5 | 7 | 14 | 7 | 10 | 11 | 42 | 55 | 45 | 43 |
| K07231 putative iron-regulated protein | 1967 | 2007 | 1901 | 1973 | 1091 | 1110 | 1206 | 1155 | 683 | 654 | 589 | 633 |
| K07233 copper resistance protein B | 2608 | 2645 | 2695 | 2725 | 1736 | 1688 | 1613 | 1659 | 842 | 751 | 757 | 821 |
| K07234 uncharacterized protein involved in response to NO | 1631 | 1487 | 1484 | 1488 | 810 | 804 | 904 | 977 | 486 | 413 | 447 | 461 |
| K19342 copper chaperone NosL | 869 | 817 | 710 | 763 | 434 | 411 | 401 | 451 | 202 | 204 | 220 | 230 |
| ko99979 Unclassified viral proteins | 4131 | 4319 | 4105 | 4358 | 5006 | 5091 | 5218 | 5191 | 5250 | 5289 | 5013 | 5052 |
| K22014 terminase small subunit | 1 | 0 | 0 | 0 | 4 | 1 | 3 | 0 | 0 | 0 | 2 | 0 |
| Poorly characterized | 394582 | 394437 | 389152 | 395148 | 359198 | 360214 | 355286 | 358810 | 343574 | 343691 | 341481 | 343943 |
| ko99996 General function prediction only | 93819 | 93815 | 91925 | 93860 | 80950 | 80897 | 78999 | 79586 | 79302 | 80043 | 78278 | 79660 |
| K00389 putative membrane protein | 378 | 285 | 338 | 348 | 230 | 210 | 225 | 189 | 666 | 599 | 599 | 602 |
| K00479 glycine betaine catabolism A | 3533 | 3594 | 3602 | 3485 | 2233 | 2074 | 1985 | 2096 | 1139 | 1107 | 1096 | 1198 |
| K01421 putative membrane protein | 912 | 839 | 887 | 909 | 321 | 344 | 298 | 324 | 1037 | 1233 | 1009 | 1170 |
| K02348 ElaA protein | 1113 | 1091 | 1113 | 1129 | 656 | 600 | 576 | 661 | 660 | 682 | 685 | 693 |
| K02351 putative membrane protein | 3 | 0 | 0 | 0 | 0 | 0 | 0 | 0 | 0 | 0 | 0 | 0 |
| K02426 cysteine desulfuration protein SufE | 1331 | 1257 | 1285 | 1319 | 1701 | 1683 | 1654 | 1692 | 1632 | 1540 | 1631 | 1618 |
| K02441 GlpG protein | 1897 | 1839 | 1691 | 1816 | 1111 | 1101 | 981 | 1124 | 1127 | 1124 | 1029 | 1101 |
| K02442 membrane protein GlpM | 438 | 389 | 338 | 344 | 213 | 184 | 194 | 210 | 146 | 100 | 93 | 123 |
| K02617 phenylacetic acid degradation protein | 202 | 216 | 227 | 188 | 59 | 46 | 44 | 48 | 404 | 391 | 373 | 422 |

| | | | | | | | | | | | | |
|---|---|---|---|---|---|---|---|---|---|---|---|---|
| K02742 SprT protein | 1159 | 1083 | 1038 | 1093 | 657 | 666 | 653 | 607 | 594 | 695 | 728 | 696 |
| K03095 SprT-like protein | 23 | 16 | 34 | 27 | 6 | 3 | 11 | 5 | 31 | 28 | 38 | 30 |
| K03112 DamX protein | 2064 | 2084 | 1929 | 2085 | 1264 | 1245 | 1323 | 1379 | 991 | 935 | 994 | 1017 |
| K03558 membrane protein required for colicin V production | 1241 | 1357 | 1209 | 1261 | 1629 | 1792 | 1354 | 1346 | 1262 | 1240 | 1155 | 1183 |
| K03618 hydrogenase-1 operon protein HyaF | 21 | 30 | 38 | 32 | 39 | 45 | 64 | 63 | 108 | 87 | 108 | 78 |
| K03619 hydrogenase-1 operon protein HyaE | 8 | 9 | 21 | 12 | 24 | 36 | 20 | 11 | 40 | 42 | 34 | 26 |
| K03744 LemA protein | 1475 | 1498 | 1458 | 1544 | 2529 | 2508 | 2683 | 2529 | 2197 | 2080 | 2309 | 2106 |
| K03745 SlyX protein | 436 | 444 | 410 | 453 | 258 | 234 | 273 | 257 | 247 | 287 | 272 | 325 |
| K03747 Smg protein | 298 | 274 | 284 | 277 | 178 | 220 | 185 | 239 | 360 | 401 | 413 | 412 |
| K03748 SanA protein | 356 | 385 | 326 | 423 | 278 | 255 | 241 | 258 | 509 | 654 | 536 | 554 |
| K03749 DedD protein | 783 | 787 | 782 | 734 | 429 | 484 | 518 | 543 | 593 | 604 | 533 | 565 |
| K03791 putative chitinase | 1389 | 1443 | 1487 | 1491 | 927 | 853 | 916 | 876 | 462 | 393 | 441 | 407 |
| K03796 Bax protein | 492 | 516 | 449 | 517 | 403 | 404 | 366 | 378 | 700 | 804 | 664 | 715 |
| K03808 paraquat-inducible protein A | 3258 | 3194 | 3157 | 3409 | 2140 | 2151 | 1888 | 2090 | 2472 | 2610 | 2456 | 2615 |
| K03810 virulence factor | 549 | 576 | 519 | 567 | 373 | 323 | 389 | 330 | 741 | 789 | 716 | 778 |
| K03923 modulator of drug activity B | 1244 | 1125 | 1124 | 1214 | 709 | 729 | 784 | 741 | 735 | 707 | 691 | 710 |
| K03925 MraZ protein | 1698 | 1663 | 1665 | 1643 | 1808 | 1878 | 1803 | 1779 | 1782 | 1740 | 1793 | 1710 |
| K03931 putative isomerase | 637 | 646 | 654 | 664 | 767 | 783 | 840 | 796 | 360 | 486 | 437 | 415 |
| K03933 chitin-binding protein | 1810 | 1668 | 1657 | 1658 | 992 | 935 | 899 | 935 | 610 | 540 | 471 | 546 |
| K03975 membrane-associated protein | 1920 | 1748 | 1879 | 1779 | 2261 | 2225 | 2164 | 2265 | 1820 | 1764 | 1872 | 1879 |
| K04065 hyperosmotically inducible periplasmic protein | 900 | 930 | 868 | 855 | 542 | 595 | 567 | 572 | 678 | 691 | 742 | 678 |
| K04477 putative hydrolase | 991 | 943 | 1037 | 1049 | 570 | 618 | 640 | 569 | 1056 | 1194 | 1013 | 993 |
| K04750 PhnB protein | 1675 | 1709 | 1601 | 1577 | 715 | 620 | 618 | 736 | 1050 | 1171 | 1103 | 1185 |
| K04756 alkyl hydroperoxide reductase subunit D | 0 | 2 | 0 | 0 | 0 | 0 | 0 | 0 | 0 | 2 | 0 | 2 |
| K05594 ElaB protein | 46 | 74 | 56 | 56 | 81 | 98 | 78 | 96 | 14 | 21 | 16 | 14 |
| K05775 maltose operon periplasmic protein | 452 | 531 | 544 | 560 | 413 | 388 | 374 | 355 | 725 | 711 | 662 | 697 |
| K05805 CreA protein | 718 | 660 | 696 | 695 | 424 | 534 | 433 | 465 | 597 | 585 | 617 | 593 |
| K05811 putative lipoprotein | 799 | 698 | 702 | 732 | 470 | 444 | 375 | 444 | 433 | 471 | 399 | 473 |
| K05812 DTW domain-containing protein | 2110 | 2001 | 1878 | 2008 | 1126 | 1163 | 1053 | 1229 | 1072 | 1092 | 1081 | 1060 |
| K05997 Fe-S cluster assembly protein SufA | 197 | 197 | 169 | 201 | 121 | 156 | 140 | 118 | 306 | 301 | 276 | 281 |
| K06135 pyrroloquinoline quinone biosynthesis protein A | 244 | 210 | 195 | 185 | 90 | 102 | 116 | 119 | 107 | 129 | 104 | 108 |
| K06136 pyrroloquinoline quinone biosynthesis protein B | 1697 | 1637 | 1504 | 1541 | 824 | 741 | 725 | 815 | 1046 | 980 | 990 | 1162 |
| K06138 pyrroloquinoline quinone biosynthesis protein D | 255 | 272 | 266 | 239 | 106 | 114 | 110 | 109 | 244 | 258 | 214 | 231 |
| K06192 paraquat-inducible protein B | 4639 | 4636 | 4686 | 4693 | 2797 | 2775 | 2701 | 2685 | 2472 | 2549 | 2346 | 2320 |
| K06193 protein PhnA | 864 | 799 | 823 | 907 | 575 | 573 | 538 | 477 | 530 | 589 | 533 | 614 |
| K06197 cation transport regulator | 132 | 127 | 111 | 146 | 78 | 85 | 82 | 100 | 206 | 179 | 199 | 224 |
| K06198 competence protein CoiA | 53 | 68 | 72 | 66 | 8 | 16 | 12 | 22 | 83 | 71 | 77 | 93 |
| K06865 ATPase | 0 | 0 | 0 | 1 | 3 | 5 | 1 | 2 | 7 | 22 | 7 | 12 |
| K06866 autonomous glycyl radical cofactor | 227 | 231 | 204 | 212 | 204 | 178 | 172 | 163 | 308 | 327 | 317 | 312 |
| K06874 zinc finger protein | 0 | 0 | 0 | 1 | 0 | 0 | 0 | 0 | 4 | 4 | 4 | 5 |
| K06877 DEAD/DEAH box helicase domain-containing protein | 886 | 931 | 883 | 858 | 160 | 162 | 193 | 155 | 902 | 1019 | 918 | 955 |
| K06878 tRNA-binding protein | 555 | 538 | 533 | 487 | 289 | 380 | 242 | 255 | 226 | 223 | 245 | 179 |
| K06886 hemoglobin | 6 | 11 | 10 | 10 | 20 | 18 | 13 | 13 | 8 | 13 | 14 | 14 |
| K06994 putative drug exporter of the RND superfamily | 8 | 2 | 1 | 9 | 6 | 6 | 2 | 5 | 4 | 7 | 6 | 8 |
| K07001 NTE family protein | 6584 | 6813 | 6893 | 6772 | 12020 | 12028 | 12229 | 11579 | 9088 | 8913 | 9215 | 9016 |
| K07023 putative hydrolases of HD superfamily | 1226 | 1136 | 1101 | 1146 | 546 | 534 | 456 | 460 | 736 | 720 | 783 | 760 |
| K07025 putative hydrolase of the HAD superfamily | 1324 | 1493 | 1579 | 1549 | 3336 | 3424 | 3383 | 3384 | 2852 | 2734 | 2790 | 2778 |
| K07038 inner membrane protein | 1179 | 1228 | 1176 | 1224 | 817 | 872 | 840 | 877 | 730 | 732 | 839 | 753 |
| K07048 phosphotriesterase-related protein | 351 | 342 | 326 | 330 | 114 | 113 | 145 | 139 | 888 | 1020 | 982 | 1068 |
| K07049 TatD-related deoxyribonuclease | 0 | 0 | 2 | 0 | 0 | 2 | 0 | 0 | 0 | 0 | 0 | 0 |
| K07058 membrane protein | 5215 | 5483 | 5440 | 5539 | 5226 | 5470 | 5380 | 5224 | 4915 | 4694 | 4739 | 4707 |
| K07082 UPF0755 protein | 3131 | 3188 | 3112 | 3198 | 3383 | 3299 | 3366 | 3320 | 3068 | 3022 | 3131 | 3120 |
| K07100 putative phosphoribosyl transferase | 2 | 0 | 0 | 0 | 0 | 1 | 0 | 0 | 0 | 0 | 6 | 0 |
| K07113 UPF0716 protein FxsA | 1176 | 1083 | 1088 | 1127 | 735 | 776 | 699 | 648 | 606 | 638 | 566 | 621 |
| K07146 UPF0176 protein | 1961 | 2001 | 1976 | 2029 | 1178 | 1194 | 1090 | 1031 | 1305 | 1335 | 1217 | 1318 |
| K07153 high frequency lysogenization protein | 1266 | 1250 | 1145 | 1312 | 782 | 707 | 786 | 735 | 820 | 787 | 795 | 755 |
| K08276 ecotin | 404 | 400 | 376 | 424 | 189 | 179 | 167 | 171 | 440 | 471 | 405 | 481 |
| K08279 carnitine operon protein CaiE | 118 | 116 | 125 | 144 | 192 | 181 | 153 | 152 | 30 | 47 | 32 | 49 |
| K08971 putative membrane protein | 0 | 0 | 0 | 0 | 0 | 3 | 0 | 1 | 1 | 11 | 3 | 9 |
| K08972 putative membrane protein | 16 | 11 | 13 | 17 | 6 | 7 | 7 | 13 | 17 | 33 | 17 | 26 |
| K08973 putative membrane protein | 662 | 692 | 747 | 724 | 412 | 362 | 413 | 416 | 256 | 216 | 196 | 239 |
| K08974 putative membrane protein | 443 | 424 | 459 | 353 | 120 | 124 | 162 | 134 | 400 | 452 | 362 | 456 |
| K08976 putative membrane protein | 2 | 0 | 0 | 0 | 0 | 0 | 0 | 0 | 0 | 0 | 0 | 0 |
| K08979 putative membrane protein | 0 | 0 | 0 | 1 | 0 | 2 | 0 | 2 | 13 | 12 | 7 | 0 |
| K08981 putative membrane protein | 18 | 19 | 9 | 14 | 18 | 7 | 8 | 10 | 21 | 25 | 29 | 28 |
| K08982 putative membrane protein | 0 | 0 | 1 | 1 | 0 | 1 | 0 | 0 | 0 | 0 | 1 | 3 |
| K08983 putative membrane protein | 688 | 694 | 586 | 653 | 397 | 398 | 362 | 403 | 205 | 211 | 171 | 221 |
| K08984 putative membrane protein | 3 | 3 | 3 | 1 | 14 | 8 | 8 | 9 | 40 | 50 | 31 | 39 |
| K08985 putative lipoprotein | 676 | 669 | 705 | 696 | 378 | 367 | 304 | 385 | 234 | 194 | 187 | 201 |
| K08986 putative membrane protein | 313 | 311 | 305 | 288 | 51 | 49 | 60 | 31 | 195 | 202 | 248 | 253 |
| K08987 putative membrane protein | 162 | 160 | 174 | 169 | 101 | 120 | 107 | 126 | 165 | 171 | 177 | 177 |
| K08988 putative membrane protein | 0 | 0 | 0 | 0 | 3 | 0 | 0 | 0 | 1 | 4 | 3 | 1 |
| K08989 putative membrane protein | 95 | 73 | 89 | 101 | 120 | 124 | 105 | 128 | 24 | 37 | 36 | 16 |
| K08990 putative membrane protein | 605 | 570 | 491 | 594 | 328 | 395 | 389 | 384 | 838 | 827 | 828 | 901 |
| K08993 putative membrane protein | 606 | 590 | 519 | 567 | 338 | 318 | 276 | 349 | 400 | 337 | 378 | 409 |
| K08995 putative membrane protein | 79 | 96 | 102 | 106 | 55 | 59 | 61 | 54 | 49 | 21 | 25 | 42 |
| K08996 putative membrane protein | 4 | 3 | 0 | 3 | 9 | 16 | 14 | 3 | 29 | 30 | 41 | 35 |
| K09014 Fe-S cluster assembly protein SufB | 1995 | 2364 | 2212 | 2199 | 4223 | 4345 | 4187 | 4201 | 3971 | 4212 | 4037 | 3922 |
| K09015 Fe-S cluster assembly protein SufD | 1659 | 1815 | 1875 | 1744 | 3837 | 3678 | 3737 | 3703 | 3412 | 3262 | 3336 | 3216 |
| K09858 SEC-C motif domain protein | 1021 | 970 | 960 | 968 | 568 | 586 | 551 | 582 | 597 | 633 | 526 | 633 |
| K09914 putative lipoprotein | 928 | 855 | 868 | 883 | 529 | 505 | 514 | 579 | 628 | 628 | 559 | 630 |
| K11312 cupin 2 domain-containing protein | 0 | 1 | 1 | 3 | 0 | 3 | 6 | 4 | 1 | 0 | 0 | 0 |
| K11477 glc operon protein GlcG | 530 | 537 | 581 | 556 | 318 | 271 | 282 | 335 | 235 | 206 | 214 | 203 |
| K11739 bacteriophage N4 adsorption protein A | 7 | 8 | 17 | 10 | 41 | 36 | 33 | 31 | 174 | 163 | 167 | 157 |
| K11740 bacteriophage N4 adsorption protein B | 17 | 18 | 14 | 23 | 46 | 27 | 20 | 23 | 157 | 165 | 153 | 125 |
| K11811 arsenical resistance protein ArsH | 1093 | 1123 | 1004 | 1099 | 743 | 725 | 682 | 769 | 301 | 269 | 278 | 298 |
| K12147 acidic protein MsyB | 225 | 245 | 200 | 217 | 164 | 167 | 166 | 135 | 323 | 418 | 306 | 298 |
| K13255 ferric iron reductase protein FhuF | 502 | 516 | 510 | 538 | 340 | 330 | 354 | 366 | 692 | 709 | 659 | 704 |
| K13256 protein PsiE | 253 | 289 | 276 | 276 | 227 | 215 | 177 | 219 | 374 | 412 | 364 | 361 |
| K13918 glucarate dehydratase-related protein | 860 | 859 | 792 | 847 | 566 | 614 | 554 | 590 | 1091 | 1137 | 1163 | 1232 |
| K14591 protein AroM | 508 | 462 | 429 | 520 | 297 | 368 | 368 | 267 | 650 | 731 | 680 | 733 |
| K15640 uncharacterized phosphatase | 8 | 5 | 6 | 5 | 2 | 0 | 0 | 2 | 61 | 73 | 56 | 74 |
| K15724 iron-sulfur cluster insertion protein | 764 | 855 | 739 | 764 | 443 | 517 | 469 | 455 | 481 | 515 | 455 | 543 |
| K21740 reactive chlorine resistance protein B | 1 | 0 | 0 | 0 | 1 | 2 | 3 | 2 | 11 | 10 | 11 | 14 |
| K21741 reactive chlorine resistance protein C | 5 | 2 | 0 | 6 | 10 | 16 | 3 | 8 | 32 | 34 | 37 | 32 |
| K21832 glycine betaine catabolism B | 1727 | 1682 | 1579 | 1531 | 973 | 814 | 860 | 912 | 537 | 571 | 495 | 501 |
| K21833 dimethylglycine catabolism A | 1775 | 1832 | 1682 | 1813 | 931 | 935 | 885 | 1001 | 615 | 555 | 528 | 545 |
| K21834 dimethylglycine catabolism B | 3510 | 3266 | 3198 | 3267 | 2006 | 1821 | 1662 | 1959 | 1090 | 1123 | 955 | 1050 |
| ko99997 Function unknown | 300763 | 300622 | 297227 | 301288 | 278248 | 279317 | 276287 | 279224 | 264272 | 263648 | 263203 | 264283 |
| K00243 uncharacterized protein | 1483 | 1516 | 1448 | 1528 | 2142 | 2226 | 2340 | 2258 | 1980 | 1965 | 1912 | 1853 |
| K01163 uncharacterized protein | 1441 | 1417 | 1491 | 1448 | 2236 | 2242 | 2396 | 2175 | 2108 | 2064 | 2123 | 1951 |
| K05937 uncharacterized protein | 3 | 4 | 3 | 7 | 2 | 2 | 1 | 2 | 4 | 18 | 2 | 4 |
| K05952 uncharacterized protein | 390 | 424 | 386 | 389 | 224 | 189 | 209 | 180 | 226 | 214 | 254 | 241 |
| K05967 uncharacterized protein | 1 | 1 | 0 | 1 | 0 | 0 | 0 | 0 | 0 | 0 | 0 | 0 |
| K06867 uncharacterized protein | 2371 | 2528 | 2373 | 2547 | 1431 | 1361 | 867 | 932 | 873 | 884 | 815 | 849 |
| K06871 uncharacterized protein | 3560 | 3440 | 3696 | 3687 | 3519 | 3558 | 3735 | 3478 | 6026 | 6306 | 6190 | 6409 |
| K06872 uncharacterized protein | 3978 | 3724 | 3738 | 3857 | 3261 | 3292 | 3076 | 3355 | 2593 | 2499 | 2479 | 2454 |
| K06883 uncharacterized protein | 0 | 0 | 0 | 3 | 0 | 0 | 0 | 0 | 0 | 0 | 0 | 2 |
| K06884 uncharacterized protein | 485 | 516 | 531 | 538 | 266 | 347 | 105 | 115 | 426 | 452 | 402 | 431 |
| K06885 uncharacterized protein | 1212 | 1430 | 1375 | 1334 | 2670 | 2778 | 2699 | 2749 | 2995 | 2984 | 2966 | 2866 |
| K06887 uncharacterized protein | 169 | 215 | 174 | 154 | 257 | 288 | 257 | 279 | 96 | 95 | 82 | 114 |
| K06888 uncharacterized protein | 683 | 665 | 729 | 702 | 158 | 161 | 154 | 166 | 568 | 659 | 638 | 666 |
| K06889 uncharacterized protein | 10587 | 10667 | 10374 | 10888 | 14616 | 14609 | 14657 | 14464 | 12566 | 12376 | 12847 | 11990 |
| K06890 uncharacterized protein | 703 | 845 | 814 | 824 | 545 | 532 | 580 | 534 | 806 | 849 | 781 | 791 |
| K06893 uncharacterized protein | 1033 | 1119 | 940 | 1031 | 523 | 529 | 527 | 560 | 316 | 322 | 264 | 321 |
| K06894 uncharacterized protein | 12152 | 12310 | 11708 | 12126 | 15286 | 15604 | 15473 | 15980 | 13858 | 13634 | 13518 | 13588 |
| K06903 uncharacterized protein | 590 | 611 | 618 | 617 | 303 | 319 | 313 | 323 | 188 | 215 | 184 | 187 |
| K06904 uncharacterized protein | 1362 | 1362 | 1504 | 1533 | 1186 | 1180 | 1180 | 1100 | 483 | 442 | 510 | 414 |
| K06905 uncharacterized protein | 1817 | 1828 | 1787 | 1852 | 926 | 1000 | 974 | 1043 | 1623 | 1884 | 1815 | 1688 |
| K06906 uncharacterized protein | 1584 | 1544 | 1462 | 1493 | 840 | 807 | 810 | 824 | 454 | 472 | 423 | 513 |
| K06907 uncharacterized protein | 2297 | 2290 | 2382 | 2347 | 1428 | 1429 | 1312 | 1394 | 2322 | 2620 | 2540 | 2481 |

| | | | | | | | | | | | | |
|---|---|---|---|---|---|---|---|---|---|---|---|---|
| K06908 uncharacterized protein | 511 | 509 | 476 | 490 | 269 | 228 | 243 | 270 | 736 | 725 | 731 | 735 |
| K06910 uncharacterized protein | 850 | 867 | 786 | 836 | 625 | 640 | 617 | 642 | 1028 | 1079 | 1072 | 1178 |
| K06911 uncharacterized protein | 5847 | 5767 | 5670 | 5838 | 4365 | 4418 | 4279 | 4440 | 4522 | 4534 | 4368 | 4584 |
| K06913 uncharacterized protein | 0 | 2 | 0 | 0 | 0 | 2 | 0 | 2 | 8 | 8 | 1 | 11 |
| K06915 uncharacterized protein | 3344 | 3307 | 3187 | 3348 | 2009 | 1963 | 1911 | 1900 | 2261 | 2307 | 2248 | 2375 |
| K06918 uncharacterized protein | 567 | 480 | 481 | 526 | 500 | 551 | 407 | 432 | 643 | 637 | 597 | 603 |
| K06921 uncharacterized protein | 1733 | 1838 | 2036 | 1892 | 4446 | 4323 | 4594 | 4431 | 3663 | 3659 | 3855 | 3772 |
| K06923 uncharacterized protein | 1928 | 1927 | 1873 | 1917 | 797 | 869 | 700 | 828 | 957 | 1026 | 1047 | 959 |
| K06926 uncharacterized protein | 452 | 525 | 529 | 519 | 1338 | 1338 | 1423 | 1347 | 1173 | 1045 | 1144 | 1205 |
| K06929 uncharacterized protein | 254 | 249 | 238 | 227 | 150 | 201 | 163 | 146 | 295 | 384 | 343 | 344 |
| K06933 uncharacterized protein | 1 | 1 | 1 | 0 | 0 | 0 | 0 | 0 | 5 | 4 | 1 | 5 |
| K06934 uncharacterized protein | 87 | 72 | 60 | 56 | 13 | 16 | 10 | 20 | 200 | 177 | 162 | 235 |
| K06936 uncharacterized protein | 2 | 2 | 2 | 1 | 0 | 0 | 3 | 0 | 5 | 4 | 2 | 1 |
| K06938 uncharacterized protein | 1185 | 1132 | 1051 | 1104 | 629 | 628 | 602 | 629 | 500 | 509 | 484 | 489 |
| K06940 uncharacterized protein | 2704 | 2566 | 2445 | 2458 | 1359 | 1397 | 1297 | 1569 | 1475 | 1485 | 1411 | 1543 |
| K06944 uncharacterized protein | 0 | 0 | 0 | 0 | 2 | 1 | 0 | 3 | 0 | 7 | 7 | 10 |
| K06946 uncharacterized protein | 6 | 5 | 4 | 6 | 22 | 25 | 19 | 30 | 76 | 88 | 103 | 108 |
| K06950 uncharacterized protein | 1887 | 1934 | 2096 | 2064 | 2969 | 3024 | 3148 | 3049 | 2740 | 2931 | 2952 | 2906 |
| K06951 uncharacterized protein | 1 | 4 | 5 | 9 | 3 | 2 | 1 | 1 | 4 | 4 | 4 | 6 |
| K06952 uncharacterized protein | 3 | 0 | 0 | 3 | 3 | 7 | 5 | 0 | 20 | 31 | 39 | 49 |
| K06953 uncharacterized protein | 0 | 0 | 2 | 0 | 0 | 0 | 0 | 2 | 3 | 0 | 3 | 1 |
| K06954 uncharacterized protein | 477 | 435 | 386 | 423 | 158 | 155 | 184 | 188 | 741 | 774 | 739 | 775 |
| K06955 uncharacterized protein | 1047 | 1029 | 900 | 975 | 580 | 578 | 510 | 639 | 338 | 288 | 295 | 281 |
| K06956 uncharacterized protein | 937 | 901 | 902 | 936 | 602 | 606 | 635 | 568 | 1120 | 1276 | 1238 | 1283 |
| K06960 uncharacterized protein | 148 | 160 | 147 | 123 | 51 | 51 | 52 | 43 | 146 | 181 | 142 | 188 |
| K06962 uncharacterized protein | 71 | 46 | 62 | 69 | 14 | 14 | 42 | 24 | 116 | 127 | 107 | 136 |
| K06971 uncharacterized protein | 763 | 798 | 762 | 745 | 425 | 416 | 447 | 420 | 818 | 916 | 917 | 935 |
| K06973 uncharacterized protein | 790 | 854 | 911 | 894 | 1513 | 1432 | 1685 | 1535 | 1359 | 1432 | 1433 | 1431 |
| K06975 uncharacterized protein | 1069 | 1058 | 1107 | 1110 | 1616 | 1594 | 1558 | 1644 | 1353 | 1328 | 1447 | 1276 |
| K06976 uncharacterized protein | 98 | 103 | 80 | 106 | 23 | 13 | 33 | 20 | 137 | 120 | 111 | 133 |
| K06977 uncharacterized protein | 0 | 0 | 1 | 0 | 0 | 0 | 0 | 0 | 0 | 0 | 0 | 0 |
| K06978 uncharacterized protein | 326 | 357 | 385 | 363 | 330 | 394 | 375 | 350 | 131 | 100 | 146 | 128 |
| K06986 uncharacterized protein | 706 | 700 | 749 | 728 | 481 | 379 | 404 | 423 | 287 | 246 | 243 | 227 |
| K06987 uncharacterized protein | 2541 | 2451 | 2387 | 2302 | 1178 | 1129 | 1059 | 1150 | 1299 | 1327 | 1316 | 1406 |
| K06991 uncharacterized protein | 687 | 740 | 646 | 706 | 383 | 412 | 349 | 383 | 206 | 201 | 183 | 232 |
| K06992 uncharacterized protein | 1687 | 1612 | 1542 | 1706 | 934 | 893 | 869 | 972 | 538 | 513 | 488 | 504 |
| K06995 uncharacterized protein | 2798 | 2641 | 2593 | 2526 | 1465 | 1348 | 1349 | 1473 | 931 | 969 | 885 | 987 |
| K06996 uncharacterized protein | 955 | 920 | 968 | 1051 | 1179 | 1195 | 1226 | 1171 | 1036 | 942 | 962 | 963 |
| K07000 uncharacterized protein | 1940 | 1845 | 1809 | 1923 | 1110 | 1102 | 1116 | 1029 | 1311 | 1396 | 1383 | 1384 |
| K07002 uncharacterized protein | 881 | 850 | 776 | 798 | 489 | 599 | 554 | 602 | 225 | 210 | 181 | 208 |
| K07003 uncharacterized protein | 7973 | 7894 | 7822 | 8010 | 4085 | 4156 | 3903 | 4126 | 3108 | 2915 | 2845 | 2815 |
| K07004 uncharacterized protein | 3116 | 3113 | 2998 | 3019 | 1747 | 1750 | 1465 | 1767 | 1007 | 910 | 895 | 976 |
| K07005 uncharacterized protein | 496 | 558 | 623 | 615 | 1198 | 1139 | 1204 | 1133 | 1081 | 1011 | 951 | 1003 |
| K07006 uncharacterized protein | 1736 | 1648 | 1607 | 1599 | 867 | 816 | 791 | 871 | 966 | 952 | 941 | 1014 |
| K07013 uncharacterized protein | 8 | 6 | 8 | 9 | 0 | 1 | 2 | 3 | 8 | 10 | 9 | 15 |
| K07014 uncharacterized protein | 3808 | 3593 | 3561 | 3795 | 2189 | 2222 | 2069 | 2202 | 2255 | 2178 | 2194 | 2201 |
| K07015 uncharacterized protein | 297 | 245 | 282 | 277 | 98 | 92 | 109 | 89 | 341 | 350 | 298 | 337 |
| K07017 uncharacterized protein | 1307 | 1334 | 1139 | 1309 | 661 | 674 | 551 | 773 | 951 | 950 | 938 | 1041 |
| K07018 uncharacterized protein | 431 | 445 | 431 | 425 | 255 | 229 | 260 | 243 | 120 | 135 | 141 | 108 |
| K07019 uncharacterized protein | 1958 | 1857 | 1790 | 1844 | 1117 | 1279 | 1191 | 1190 | 1245 | 1237 | 1204 | 1256 |
| K07020 uncharacterized protein | 841 | 737 | 821 | 827 | 460 | 373 | 400 | 456 | 285 | 223 | 225 | 231 |
| K07028 uncharacterized protein | 2353 | 2265 | 2318 | 2368 | 1391 | 1203 | 1312 | 1469 | 782 | 679 | 815 | 764 |
| K07030 uncharacterized protein | 826 | 832 | 819 | 777 | 269 | 287 | 291 | 301 | 722 | 893 | 872 | 835 |
| K07032 uncharacterized protein | 687 | 716 | 677 | 675 | 391 | 401 | 360 | 375 | 267 | 229 | 235 | 227 |
| K07033 uncharacterized protein | 36 | 44 | 38 | 41 | 14 | 14 | 11 | 8 | 69 | 89 | 87 | 74 |
| K07034 uncharacterized protein | 397 | 407 | 371 | 381 | 248 | 284 | 305 | 276 | 555 | 508 | 516 | 541 |
| K07035 uncharacterized protein | 648 | 639 | 657 | 673 | 188 | 197 | 233 | 188 | 686 | 760 | 652 | 718 |
| K07039 uncharacterized protein | 1159 | 1241 | 1135 | 1249 | 736 | 708 | 764 | 776 | 834 | 815 | 827 | 850 |
| K07040 uncharacterized protein | 1315 | 1409 | 1336 | 1428 | 748 | 760 | 738 | 762 | 820 | 960 | 928 | 885 |
| K07041 uncharacterized protein | 2 | 2 | 2 | 1 | 3 | 2 | 0 | 3 | 7 | 8 | 10 | 10 |
| K07043 uncharacterized protein | 1561 | 1647 | 1732 | 1588 | 2274 | 2304 | 2311 | 2288 | 2102 | 2119 | 2124 | 2021 |
| K07044 uncharacterized protein | 3255 | 3246 | 3188 | 3165 | 1827 | 1726 | 1740 | 1865 | 1058 | 1021 | 1020 | 974 |
| K07045 uncharacterized protein | 811 | 759 | 816 | 804 | 196 | 222 | 211 | 175 | 730 | 707 | 698 | 767 |
| K07051 uncharacterized protein | 0 | 0 | 0 | 0 | 0 | 2 | 2 | 0 | 6 | 4 | 3 | 8 |
| K07052 uncharacterized protein | 3292 | 3395 | 3413 | 3393 | 3982 | 4016 | 4095 | 4156 | 3248 | 3091 | 3206 | 3182 |
| K07054 uncharacterized protein | 571 | 534 | 518 | 571 | 358 | 395 | 468 | 485 | 786 | 690 | 720 | 815 |
| K07063 uncharacterized protein | 0 | 1 | 0 | 0 | 0 | 0 | 0 | 0 | 0 | 0 | 1 | 1 |
| K07068 uncharacterized protein | 590 | 639 | 595 | 630 | 321 | 325 | 313 | 321 | 191 | 214 | 156 | 200 |
| K07069 uncharacterized protein | 53 | 45 | 63 | 60 | 17 | 19 | 16 | 20 | 83 | 110 | 105 | 85 |
| K07070 uncharacterized protein | 592 | 622 | 558 | 586 | 370 | 384 | 328 | 369 | 324 | 301 | 284 | 343 |
| K07071 uncharacterized protein | 2763 | 2663 | 2723 | 2707 | 3612 | 3519 | 3570 | 3654 | 3074 | 2946 | 3041 | 2970 |
| K07074 uncharacterized protein | 1896 | 1962 | 1939 | 1911 | 1185 | 1218 | 1062 | 1168 | 660 | 671 | 648 | 705 |
| K07075 uncharacterized protein | 765 | 876 | 869 | 821 | 2671 | 2545 | 2762 | 2725 | 1919 | 1626 | 1912 | 1785 |
| K07076 uncharacterized protein | 1165 | 1103 | 1119 | 1020 | 575 | 537 | 509 | 569 | 344 | 298 | 264 | 332 |
| K07077 uncharacterized protein | 0 | 0 | 2 | 3 | 3 | 2 | 0 | 0 | 1 | 1 | 2 | 11 |
| K07078 uncharacterized protein | 1602 | 1448 | 1606 | 1527 | 2045 | 2198 | 2134 | 2041 | 1433 | 1441 | 1456 | 1351 |
| K07079 uncharacterized protein | 2023 | 2069 | 2072 | 2219 | 3016 | 3101 | 3096 | 2971 | 3658 | 3984 | 3827 | 3840 |
| K07080 uncharacterized protein | 1249 | 1186 | 1150 | 1204 | 878 | 871 | 888 | 890 | 426 | 437 | 372 | 442 |
| K07088 uncharacterized protein | 3769 | 3890 | 3932 | 3869 | 2273 | 2405 | 2225 | 2317 | 2808 | 2804 | 2700 | 2945 |
| K07089 uncharacterized protein | 1765 | 1656 | 1733 | 1728 | 1084 | 1208 | 1173 | 1242 | 1720 | 1742 | 1848 | 1965 |
| K07090 uncharacterized protein | 11714 | 11633 | 11092 | 11548 | 5844 | 5696 | 5631 | 6211 | 5590 | 5571 | 5584 | 5720 |
| K07092 uncharacterized protein | 556 | 521 | 470 | 454 | 272 | 261 | 260 | 309 | 156 | 142 | 149 | 146 |
| K07093 uncharacterized protein | 228 | 234 | 243 | 245 | 114 | 126 | 133 | 131 | 91 | 75 | 62 | 67 |
| K07095 uncharacterized protein | 2032 | 2117 | 1982 | 2062 | 1898 | 1897 | 1826 | 1912 | 2001 | 2113 | 2106 | 1970 |
| K07096 uncharacterized protein | 23 | 19 | 23 | 27 | 4 | 3 | 8 | 3 | 14 | 14 | 4 | 17 |
| K07097 uncharacterized protein | 2 | 0 | 0 | 1 | 0 | 0 | 0 | 0 | 0 | 0 | 0 | 1 |
| K07098 uncharacterized protein | 1825 | 1845 | 1871 | 1834 | 2937 | 2991 | 3085 | 2911 | 2353 | 2319 | 2404 | 2393 |
| K07099 uncharacterized protein | 61 | 83 | 70 | 70 | 69 | 50 | 54 | 43 | 90 | 78 | 82 | 111 |
| K07101 uncharacterized protein | 16 | 12 | 19 | 19 | 38 | 39 | 50 | 39 | 53 | 40 | 31 | 30 |
| K07103 uncharacterized protein | 0 | 0 | 0 | 1 | 0 | 0 | 0 | 0 | 4 | 4 | 0 | 2 |
| K07105 uncharacterized protein | 152 | 148 | 146 | 170 | 63 | 70 | 56 | 51 | 202 | 210 | 179 | 197 |
| K07108 uncharacterized protein | 2 | 0 | 2 | 0 | 1 | 0 | 0 | 2 | 0 | 1 | 3 | 1 |
| K07109 uncharacterized protein | 317 | 330 | 329 | 340 | 234 | 267 | 257 | 253 | 494 | 458 | 505 | 481 |
| K07112 uncharacterized protein | 4577 | 4539 | 4150 | 4447 | 2601 | 2659 | 2574 | 2743 | 2243 | 2249 | 2066 | 2317 |
| K07118 uncharacterized protein | 1423 | 1509 | 1433 | 1444 | 1924 | 1904 | 1883 | 1848 | 1432 | 1330 | 1402 | 1329 |
| K07120 uncharacterized protein | 1933 | 1878 | 1817 | 1858 | 918 | 958 | 841 | 921 | 1199 | 1283 | 1304 | 1268 |
| K07121 uncharacterized protein | 4163 | 4147 | 4027 | 4157 | 2503 | 2481 | 2475 | 2652 | 2498 | 2481 | 2603 | 2626 |
| K07123 uncharacterized protein | 0 | 0 | 0 | 1 | 0 | 0 | 0 | 0 | 2 | 12 | 8 | 12 |
| K07124 uncharacterized protein | 3631 | 3926 | 3571 | 3821 | 2209 | 2298 | 2176 | 2231 | 2013 | 1816 | 1887 | 1992 |
| K07126 uncharacterized protein | 7306 | 7349 | 7465 | 7358 | 6094 | 6059 | 4548 | 4553 | 4966 | 4559 | 4904 | 4715 |
| K07129 uncharacterized protein | 33 | 41 | 30 | 31 | 2 | 0 | 3 | 2 | 46 | 61 | 57 | 60 |
| K07133 uncharacterized protein | 9046 | 9618 | 10346 | 9509 | 25822 | 26048 | 26838 | 26319 | 20872 | 20053 | 21045 | 20721 |
| K07134 uncharacterized protein | 0 | 0 | 2 | 0 | 0 | 3 | 0 | 0 | 1 | 5 | 2 | 8 |
| K07135 uncharacterized protein | 0 | 0 | 0 | 0 | 0 | 0 | 0 | 1 | 0 | 2 | 1 | 5 |
| K07136 uncharacterized protein | 279 | 228 | 215 | 264 | 181 | 179 | 176 | 154 | 365 | 402 | 382 | 380 |
| K07137 uncharacterized protein | 2626 | 2652 | 2631 | 2723 | 3939 | 3854 | 3926 | 3597 | 4038 | 4113 | 4163 | 3974 |
| K07138 uncharacterized protein | 1019 | 945 | 959 | 949 | 607 | 542 | 668 | 615 | 987 | 1018 | 965 | 1073 |
| K07139 uncharacterized protein | 1616 | 1569 | 1622 | 1605 | 2582 | 2610 | 2760 | 2486 | 2677 | 2656 | 2640 | 2715 |
| K07140 uncharacterized protein | 1735 | 1832 | 1727 | 1731 | 953 | 1010 | 993 | 959 | 1240 | 1258 | 1230 | 1300 |
| K07148 uncharacterized protein | 1999 | 2244 | 2401 | 2238 | 5524 | 5498 | 5405 | 5481 | 4422 | 4278 | 4508 | 4277 |
| K07149 uncharacterized protein | 25 | 36 | 30 | 29 | 12 | 11 | 10 | 10 | 17 | 39 | 21 | 39 |
| K07150 uncharacterized protein | 914 | 830 | 907 | 873 | 658 | 657 | 611 | 610 | 1031 | 1163 | 1095 | 1192 |
| K07157 uncharacterized protein | 241 | 261 | 217 | 223 | 115 | 117 | 103 | 138 | 65 | 68 | 91 | 79 |
| K07158 uncharacterized protein | 0 | 0 | 2 | 0 | 0 | 0 | 0 | 0 | 2 | 1 | 3 | 2 |
| K07159 uncharacterized protein | 1 | 0 | 0 | 0 | 0 | 0 | 2 | 2 | 5 | 2 | 5 | 3 |
| K07161 uncharacterized protein | 0 | 2 | 1 | 2 | 0 | 0 | 0 | 0 | 6 | 2 | 2 | 7 |
| K07164 uncharacterized protein | 476 | 540 | 544 | 477 | 1556 | 1491 | 1540 | 1496 | 1110 | 1155 | 1140 | 1111 |
| K07220 uncharacterized protein | 1412 | 1403 | 1419 | 1507 | 1931 | 1798 | 1884 | 1752 | 1508 | 1499 | 1476 | 1367 |
| K07269 uncharacterized protein | 337 | 347 | 376 | 381 | 239 | 257 | 239 | 221 | 516 | 489 | 488 | 500 |
| K07276 uncharacterized protein | 0 | 0 | 0 | 1 | 0 | 0 | 0 | 0 | 0 | 2 | 2 | 0 |
| K07338 uncharacterized protein | 1762 | 1715 | 1875 | 1664 | 943 | 951 | 910 | 1009 | 538 | 584 | 574 | 561 |

| | | | | | | | | | | | | |
|---|---|---|---|---|---|---|---|---|---|---|---|---|
| K07502 uncharacterized protein | 372 | 340 | 354 | 321 | 60 | 80 | 71 | 53 | 287 | 315 | 309 | 304 |
| K07584 uncharacterized protein | 181 | 168 | 166 | 174 | 62 | 57 | 47 | 47 | 161 | 165 | 140 | 146 |
| K07586 uncharacterized protein | 36 | 37 | 32 | 45 | 12 | 14 | 4 | 4 | 39 | 43 | 24 | 24 |
| K07742 uncharacterized protein | 185 | 184 | 185 | 203 | 57 | 73 | 64 | 48 | 141 | 184 | 147 | 167 |
| K07807 uncharacterized protein | 384 | 412 | 469 | 424 | 377 | 319 | 294 | 306 | 99 | 111 | 89 | 117 |
| K08997 uncharacterized protein | 884 | 854 | 797 | 857 | 640 | 647 | 590 | 546 | 1117 | 1178 | 1086 | 1191 |
| K08998 uncharacterized protein | 906 | 886 | 864 | 901 | 995 | 983 | 1002 | 1044 | 1027 | 1058 | 1013 | 1014 |
| K08999 uncharacterized protein | 380 | 358 | 423 | 396 | 1257 | 1338 | 1236 | 1251 | 873 | 864 | 862 | 875 |
| K09003 uncharacterized protein | 2 | 0 | 2 | 3 | 6 | 0 | 0 | 2 | 0 | 8 | 3 | 4 |
| K09004 uncharacterized protein | 70 | 69 | 69 | 53 | 9 | 16 | 9 | 18 | 48 | 62 | 50 | 48 |
| K09005 uncharacterized protein | 3 | 2 | 4 | 1 | 2 | 4 | 0 | 0 | 13 | 2 | 7 | 7 |
| K09009 uncharacterized protein | 2 | 3 | 0 | 1 | 0 | 2 | 1 | 6 | 8 | 2 | 1 | 4 |
| K09116 uncharacterized protein | 61 | 62 | 47 | 58 | 25 | 8 | 13 | 23 | 115 | 158 | 131 | 166 |
| K09117 uncharacterized protein | 934 | 902 | 965 | 935 | 1283 | 1375 | 1451 | 1383 | 1172 | 1264 | 1175 | 1169 |
| K09118 uncharacterized protein | 56 | 86 | 60 | 61 | 60 | 53 | 45 | 49 | 77 | 112 | 108 | 114 |
| K09122 uncharacterized protein | 0 | 0 | 0 | 0 | 0 | 0 | 0 | 0 | 2 | 0 | 0 | 0 |
| K09124 uncharacterized protein | 71 | 98 | 104 | 83 | 310 | 319 | 321 | 287 | 36 | 69 | 49 | 51 |
| K09125 uncharacterized protein | 1815 | 1888 | 1923 | 1901 | 2182 | 2134 | 2151 | 2276 | 1599 | 1607 | 1611 | 1614 |
| K09126 uncharacterized protein | 0 | 0 | 1 | 0 | 3 | 2 | 4 | 4 | 4 | 11 | 3 | 2 |
| K09128 uncharacterized protein | 88 | 98 | 99 | 118 | 39 | 35 | 29 | 38 | 169 | 179 | 169 | 186 |
| K09129 uncharacterized protein | 0 | 0 | 0 | 0 | 0 | 0 | 0 | 0 | 4 | 0 | 0 | 0 |
| K09131 uncharacterized protein | 536 | 560 | 547 | 588 | 336 | 362 | 346 | 347 | 362 | 325 | 294 | 329 |
| K09133 uncharacterized protein | 121 | 157 | 132 | 147 | 62 | 61 | 51 | 72 | 162 | 182 | 153 | 176 |
| K09137 uncharacterized protein | 14 | 8 | 15 | 10 | 0 | 0 | 0 | 0 | 27 | 26 | 35 | 29 |
| K09138 uncharacterized protein | 135 | 128 | 125 | 131 | 28 | 22 | 24 | 20 | 91 | 92 | 83 | 99 |
| K09139 uncharacterized protein | 2 | 0 | 0 | 0 | 3 | 0 | 0 | 0 | 0 | 5 | 2 | 4 |
| K09141 uncharacterized protein | 5 | 6 | 8 | 5 | 5 | 1 | 3 | 2 | 4 | 22 | 18 | 17 |
| K09142 uncharacterized protein | 0 | 0 | 0 | 0 | 0 | 0 | 0 | 0 | 0 | 2 | 4 | 3 |
| K09144 uncharacterized protein | 0 | 0 | 2 | 0 | 0 | 0 | 3 | 2 | 6 | 7 | 2 | 7 |
| K09145 uncharacterized protein | 23 | 41 | 31 | 35 | 3 | 4 | 3 | 10 | 82 | 68 | 66 | 80 |
| K09154 uncharacterized protein | 0 | 0 | 0 | 0 | 0 | 0 | 0 | 0 | 3 | 7 | 3 | 4 |
| K09155 uncharacterized protein | 155 | 172 | 145 | 159 | 108 | 109 | 114 | 112 | 61 | 93 | 74 | 81 |
| K09157 uncharacterized protein | 848 | 881 | 860 | 874 | 378 | 419 | 388 | 397 | 956 | 1033 | 994 | 992 |
| K09158 uncharacterized protein | 576 | 582 | 600 | 610 | 372 | 376 | 343 | 408 | 336 | 428 | 373 | 379 |
| K09160 uncharacterized protein | 1005 | 938 | 925 | 1004 | 572 | 587 | 558 | 512 | 655 | 609 | 565 | 657 |
| K09161 uncharacterized protein | 309 | 269 | 287 | 313 | 176 | 227 | 233 | 183 | 415 | 461 | 416 | 466 |
| K09163 uncharacterized protein | 48 | 44 | 42 | 54 | 7 | 9 | 11 | 17 | 75 | 89 | 83 | 116 |
| K09164 uncharacterized protein | 2 | 2 | 2 | 0 | 0 | 2 | 0 | 0 | 0 | 1 | 0 | 1 |
| K09167 uncharacterized protein | 2 | 6 | 2 | 7 | 3 | 2 | 4 | 6 | 13 | 14 | 4 | 12 |
| K09190 uncharacterized protein | 0 | 0 | 0 | 0 | 0 | 0 | 0 | 2 | 0 | 0 | 0 | 0 |
| K09384 uncharacterized protein | 43 | 40 | 40 | 47 | 22 | 13 | 17 | 10 | 60 | 89 | 77 | 109 |
| K09386 uncharacterized protein | 0 | 0 | 0 | 0 | 0 | 0 | 0 | 2 | 3 | 0 | 0 | 6 |
| K09701 uncharacterized protein | 467 | 424 | 431 | 464 | 284 | 308 | 287 | 310 | 577 | 602 | 578 | 603 |
| K09702 uncharacterized protein | 307 | 333 | 324 | 316 | 294 | 285 | 252 | 238 | 348 | 402 | 343 | 418 |
| K09703 uncharacterized protein | 197 | 175 | 219 | 204 | 35 | 32 | 41 | 28 | 225 | 275 | 257 | 257 |
| K09704 uncharacterized protein | 3013 | 3094 | 3329 | 3077 | 9166 | 9174 | 9448 | 8972 | 8143 | 7748 | 7916 | 7290 |
| K09705 uncharacterized protein | 115 | 139 | 104 | 136 | 30 | 31 | 28 | 22 | 88 | 99 | 82 | 96 |
| K09706 uncharacterized protein | 1 | 3 | 0 | 1 | 0 | 3 | 0 | 0 | 1 | 0 | 6 | 4 |
| K09707 uncharacterized protein | 202 | 188 | 152 | 175 | 37 | 18 | 20 | 24 | 200 | 192 | 174 | 179 |
| K09711 uncharacterized protein | 1 | 0 | 1 | 1 | 0 | 1 | 0 | 1 | 0 | 0 | 2 | 7 |
| K09712 uncharacterized protein | 137 | 143 | 125 | 135 | 74 | 73 | 54 | 84 | 189 | 221 | 216 | 193 |
| K09713 uncharacterized protein | 0 | 0 | 0 | 0 | 0 | 0 | 0 | 0 | 2 | 0 | 0 | 12 |
| K09717 uncharacterized protein | 0 | 4 | 0 | 0 | 0 | 3 | 2 | 0 | 8 | 7 | 2 | 3 |
| K09721 uncharacterized protein | 0 | 0 | 0 | 1 | 0 | 0 | 0 | 0 | 0 | 2 | 0 | 1 |
| K09724 uncharacterized protein | 0 | 2 | 0 | 2 | 2 | 4 | 2 | 2 | 5 | 13 | 4 | 11 |
| K09726 uncharacterized protein | 0 | 0 | 0 | 0 | 0 | 0 | 0 | 0 | 5 | 1 | 6 | 12 |
| K09727 uncharacterized protein | 0 | 2 | 2 | 0 | 0 | 0 | 0 | 0 | 1 | 0 | 0 | 4 |
| K09728 uncharacterized protein | 0 | 0 | 0 | 0 | 0 | 1 | 0 | 0 | 0 | 0 | 0 | 0 |
| K09729 uncharacterized protein | 26 | 44 | 37 | 39 | 3 | 2 | 7 | 2 | 54 | 51 | 48 | 58 |
| K09730 uncharacterized protein | 0 | 0 | 0 | 0 | 0 | 0 | 5 | 0 | 1 | 5 | 2 | 3 |
| K09735 uncharacterized protein | 0 | 0 | 0 | 0 | 1 | 0 | 2 | 0 | 2 | 5 | 2 | 5 |
| K09737 uncharacterized protein | 0 | 0 | 0 | 0 | 0 | 0 | 0 | 0 | 1 | 0 | 0 | 0 |
| K09738 uncharacterized protein | 0 | 0 | 2 | 0 | 0 | 0 | 0 | 0 | 0 | 2 | 0 | 3 |
| K09740 uncharacterized protein | 9 | 4 | 16 | 13 | 15 | 13 | 19 | 10 | 17 | 32 | 31 | 22 |
| K09747 uncharacterized protein | 942 | 918 | 882 | 931 | 489 | 512 | 416 | 510 | 757 | 751 | 778 | 745 |
| K09749 uncharacterized protein | 345 | 347 | 379 | 372 | 84 | 113 | 115 | 78 | 308 | 340 | 361 | 391 |
| K09762 uncharacterized protein | 622 | 526 | 564 | 502 | 160 | 165 | 203 | 166 | 432 | 599 | 576 | 482 |
| K09763 uncharacterized protein | 0 | 0 | 0 | 0 | 0 | 0 | 0 | 1 | 0 | 0 | 0 | 0 |
| K09764 uncharacterized protein | 10 | 14 | 12 | 10 | 13 | 18 | 19 | 18 | 18 | 24 | 24 | 28 |
| K09766 uncharacterized protein | 1 | 2 | 0 | 1 | 0 | 0 | 0 | 0 | 0 | 1 | 2 | 2 |
| K09768 uncharacterized protein | 1317 | 1268 | 1344 | 1296 | 691 | 733 | 690 | 775 | 904 | 858 | 912 | 953 |
| K09769 uncharacterized protein | 126 | 89 | 92 | 98 | 57 | 90 | 76 | 72 | 90 | 122 | 78 | 132 |
| K09770 uncharacterized protein | 0 | 0 | 0 | 3 | 0 | 0 | 0 | 0 | 2 | 0 | 0 | 2 |
| K09775 uncharacterized protein | 261 | 192 | 216 | 257 | 58 | 64 | 73 | 81 | 182 | 170 | 192 | 183 |
| K09776 uncharacterized protein | 0 | 0 | 0 | 0 | 0 | 0 | 0 | 0 | 0 | 0 | 2 | 0 |
| K09777 uncharacterized protein | 39 | 37 | 51 | 37 | 24 | 28 | 40 | 18 | 53 | 64 | 69 | 66 |
| K09779 uncharacterized protein | 139 | 122 | 164 | 145 | 50 | 49 | 45 | 37 | 122 | 123 | 145 | 176 |
| K09780 uncharacterized protein | 631 | 614 | 571 | 598 | 295 | 355 | 363 | 387 | 384 | 339 | 405 | 425 |
| K09781 uncharacterized protein | 1105 | 1072 | 1012 | 1048 | 635 | 656 | 634 | 698 | 834 | 829 | 725 | 765 |
| K09786 uncharacterized protein | 2519 | 2526 | 2440 | 2578 | 1503 | 1506 | 1463 | 1563 | 1569 | 1587 | 1621 | 1609 |
| K09787 uncharacterized protein | 186 | 191 | 224 | 201 | 60 | 70 | 57 | 63 | 163 | 183 | 177 | 203 |
| K09790 uncharacterized protein | 1022 | 1001 | 998 | 918 | 1124 | 1167 | 1180 | 1119 | 1085 | 978 | 1049 | 986 |
| K09791 uncharacterized protein | 368 | 403 | 335 | 329 | 261 | 211 | 183 | 214 | 269 | 287 | 251 | 224 |
| K09792 uncharacterized protein | 326 | 334 | 315 | 312 | 146 | 184 | 84 | 101 | 97 | 83 | 92 | 82 |
| K09793 uncharacterized protein | 230 | 224 | 217 | 243 | 656 | 627 | 935 | 885 | 576 | 431 | 571 | 514 |
| K09794 uncharacterized protein | 380 | 347 | 323 | 351 | 213 | 178 | 183 | 188 | 116 | 103 | 94 | 126 |
| K09795 uncharacterized protein | 2 | 1 | 2 | 0 | 2 | 2 | 3 | 1 | 1 | 0 | 3 | 2 |
| K09797 uncharacterized protein | 206 | 227 | 219 | 249 | 144 | 128 | 149 | 163 | 227 | 256 | 246 | 233 |
| K09798 uncharacterized protein | 1872 | 1868 | 1783 | 1786 | 1085 | 1143 | 1053 | 1087 | 1306 | 1269 | 1162 | 1250 |
| K09799 uncharacterized protein | 0 | 0 | 0 | 0 | 0 | 5 | 0 | 2 | 2 | 2 | 6 | 5 |
| K09801 uncharacterized protein | 678 | 575 | 588 | 586 | 401 | 364 | 414 | 411 | 314 | 328 | 315 | 352 |
| K09802 uncharacterized protein | 232 | 200 | 184 | 213 | 158 | 147 | 155 | 173 | 251 | 301 | 232 | 235 |
| K09803 uncharacterized protein | 28 | 33 | 21 | 31 | 5 | 4 | 3 | 10 | 7 | 12 | 26 | 15 |
| K09804 uncharacterized protein | 3 | 0 | 0 | 0 | 0 | 0 | 0 | 0 | 0 | 0 | 0 | 0 |
| K09805 uncharacterized protein | 8 | 14 | 9 | 9 | 12 | 5 | 8 | 4 | 33 | 48 | 26 | 34 |
| K09806 uncharacterized protein | 729 | 653 | 692 | 692 | 424 | 419 | 468 | 444 | 452 | 403 | 399 | 391 |
| K09807 uncharacterized protein | 1120 | 990 | 1052 | 1081 | 1796 | 1725 | 1784 | 1732 | 1815 | 1814 | 1759 | 1776 |
| K09824 uncharacterized protein | 671 | 693 | 719 | 755 | 660 | 634 | 628 | 628 | 763 | 817 | 739 | 864 |
| K09857 uncharacterized protein | 1608 | 1568 | 1444 | 1529 | 922 | 891 | 868 | 904 | 790 | 799 | 851 | 886 |
| K09861 uncharacterized protein | 2472 | 2540 | 2529 | 2480 | 2935 | 2941 | 2906 | 2865 | 2681 | 2641 | 2838 | 2772 |
| K09862 uncharacterized protein | 441 | 414 | 444 | 421 | 269 | 310 | 298 | 271 | 223 | 278 | 267 | 264 |
| K09891 uncharacterized protein | 379 | 340 | 375 | 389 | 263 | 291 | 273 | 233 | 537 | 506 | 522 | 499 |
| K09894 uncharacterized protein | 218 | 231 | 248 | 270 | 197 | 209 | 136 | 138 | 343 | 350 | 322 | 369 |
| K09895 uncharacterized protein | 508 | 540 | 517 | 538 | 329 | 397 | 337 | 331 | 483 | 606 | 539 | 608 |
| K09896 uncharacterized protein | 151 | 182 | 206 | 185 | 139 | 149 | 178 | 110 | 254 | 248 | 236 | 257 |
| K09897 uncharacterized protein | 282 | 230 | 238 | 274 | 183 | 216 | 187 | 164 | 313 | 338 | 325 | 355 |
| K09898 uncharacterized protein | 523 | 507 | 451 | 518 | 285 | 299 | 260 | 286 | 302 | 310 | 296 | 302 |
| K09899 uncharacterized protein | 235 | 211 | 202 | 248 | 175 | 177 | 183 | 177 | 323 | 415 | 373 | 392 |
| K09900 uncharacterized protein | 169 | 177 | 166 | 181 | 127 | 136 | 155 | 144 | 278 | 245 | 209 | 281 |
| K09901 uncharacterized protein | 172 | 186 | 160 | 191 | 105 | 135 | 115 | 129 | 274 | 250 | 198 | 294 |
| K09902 uncharacterized protein | 569 | 598 | 571 | 601 | 322 | 341 | 343 | 358 | 375 | 379 | 363 | 415 |
| K09904 uncharacterized protein | 180 | 185 | 152 | 195 | 120 | 121 | 115 | 102 | 269 | 291 | 267 | 270 |
| K09907 uncharacterized protein | 328 | 332 | 361 | 334 | 213 | 272 | 249 | 207 | 453 | 453 | 500 | 468 |
| K09908 uncharacterized protein | 1013 | 977 | 938 | 894 | 548 | 595 | 565 | 570 | 694 | 636 | 561 | 590 |
| K09909 uncharacterized protein | 133 | 145 | 136 | 134 | 197 | 231 | 231 | 187 | 18 | 37 | 40 | 27 |
| K09910 uncharacterized protein | 210 | 192 | 207 | 238 | 192 | 168 | 189 | 181 | 286 | 334 | 337 | 334 |
| K09911 uncharacterized protein | 242 | 238 | 247 | 276 | 198 | 192 | 158 | 140 | 322 | 325 | 371 | 351 |
| K09912 uncharacterized protein | 997 | 943 | 1030 | 964 | 564 | 546 | 527 | 507 | 651 | 640 | 641 | 684 |
| K09915 uncharacterized protein | 1001 | 995 | 859 | 988 | 591 | 584 | 585 | 636 | 710 | 755 | 747 | 738 |
| K09916 uncharacterized protein | 575 | 555 | 569 | 583 | 342 | 344 | 311 | 327 | 421 | 415 | 357 | 357 |
| K09917 uncharacterized protein | 98 | 92 | 101 | 114 | 63 | 87 | 88 | 62 | 151 | 183 | 150 | 136 |

| | | | | | | | | | | | | |
|---|---|---|---|---|---|---|---|---|---|---|---|---|
| K09918 uncharacterized protein | 425 | 413 | 434 | 368 | 289 | 254 | 289 | 322 | 158 | 147 | 129 | 123 |
| K09919 uncharacterized protein | 1789 | 1696 | 1686 | 1757 | 963 | 929 | 923 | 949 | 553 | 504 | 513 | 554 |
| K09920 uncharacterized protein | 973 | 972 | 959 | 931 | 582 | 621 | 591 | 635 | 708 | 638 | 605 | 635 |
| K09921 uncharacterized protein | 1384 | 1332 | 1248 | 1366 | 796 | 822 | 827 | 856 | 970 | 927 | 828 | 944 |
| K09922 uncharacterized protein | 645 | 738 | 777 | 699 | 1003 | 957 | 1044 | 1010 | 662 | 746 | 577 | 602 |
| K09923 uncharacterized protein | 709 | 686 | 725 | 731 | 462 | 449 | 358 | 348 | 451 | 472 | 485 | 465 |
| K09924 uncharacterized protein | 875 | 887 | 772 | 793 | 515 | 498 | 448 | 500 | 302 | 263 | 263 | 278 |
| K09925 uncharacterized protein | 859 | 844 | 904 | 835 | 462 | 460 | 429 | 454 | 353 | 328 | 320 | 322 |
| K09926 uncharacterized protein | 744 | 733 | 715 | 704 | 417 | 368 | 384 | 442 | 229 | 213 | 235 | 227 |
| K09927 uncharacterized protein | 3202 | 3095 | 3065 | 3110 | 1426 | 1498 | 1397 | 1434 | 2130 | 2158 | 2226 | 2147 |
| K09928 uncharacterized protein | 396 | 431 | 414 | 441 | 211 | 256 | 209 | 219 | 100 | 119 | 119 | 136 |
| K09929 uncharacterized protein | 760 | 814 | 772 | 744 | 397 | 411 | 328 | 375 | 273 | 219 | 242 | 224 |
| K09930 uncharacterized protein | 1300 | 1410 | 1432 | 1323 | 795 | 784 | 686 | 659 | 385 | 347 | 443 | 426 |
| K09931 uncharacterized protein | 849 | 785 | 718 | 708 | 404 | 367 | 327 | 390 | 264 | 274 | 232 | 211 |
| K09932 uncharacterized protein | 834 | 803 | 766 | 750 | 374 | 417 | 331 | 414 | 597 | 598 | 595 | 620 |
| K09934 uncharacterized protein | 861 | 878 | 892 | 846 | 537 | 497 | 506 | 541 | 374 | 270 | 316 | 301 |
| K09935 uncharacterized protein | 1114 | 1059 | 1050 | 1074 | 588 | 527 | 535 | 573 | 498 | 517 | 462 | 503 |
| K09937 uncharacterized protein | 503 | 543 | 509 | 510 | 306 | 292 | 278 | 268 | 289 | 336 | 352 | 367 |
| K09938 uncharacterized protein | 1629 | 1615 | 1522 | 1608 | 910 | 801 | 824 | 912 | 546 | 466 | 521 | 476 |
| K09939 uncharacterized protein | 58 | 68 | 79 | 67 | 97 | 107 | 87 | 98 | 2 | 19 | 9 | 12 |
| K09940 uncharacterized protein | 669 | 672 | 724 | 733 | 351 | 358 | 348 | 406 | 217 | 224 | 201 | 206 |
| K09945 uncharacterized protein | 1104 | 1051 | 1073 | 1092 | 603 | 640 | 721 | 850 | 704 | 662 | 585 | 664 |
| K09946 uncharacterized protein | 620 | 583 | 575 | 641 | 351 | 310 | 348 | 367 | 249 | 192 | 203 | 249 |
| K09947 uncharacterized protein | 491 | 459 | 492 | 477 | 280 | 304 | 266 | 268 | 124 | 132 | 138 | 148 |
| K09948 uncharacterized protein | 477 | 448 | 377 | 429 | 195 | 232 | 240 | 238 | 168 | 116 | 122 | 140 |
| K09949 uncharacterized protein | 36 | 26 | 45 | 47 | 4 | 11 | 3 | 4 | 41 | 59 | 49 | 58 |
| K09950 uncharacterized protein | 1764 | 1737 | 1569 | 1729 | 905 | 885 | 928 | 970 | 601 | 564 | 452 | 548 |
| K09954 uncharacterized protein | 491 | 489 | 459 | 488 | 270 | 312 | 263 | 284 | 315 | 340 | 299 | 308 |
| K09955 uncharacterized protein | 7153 | 7561 | 7758 | 7547 | 12477 | 12339 | 12973 | 12959 | 9589 | 9705 | 9638 | 9490 |
| K09956 uncharacterized protein | 517 | 510 | 443 | 411 | 257 | 268 | 215 | 266 | 154 | 153 | 151 | 154 |
| K09957 uncharacterized protein | 491 | 459 | 510 | 471 | 269 | 239 | 230 | 285 | 149 | 155 | 145 | 142 |
| K09958 uncharacterized protein | 1139 | 1158 | 1030 | 1081 | 638 | 734 | 580 | 648 | 650 | 631 | 628 | 648 |
| K09959 uncharacterized protein | 0 | 3 | 0 | 3 | 0 | 0 | 2 | 0 | 2 | 0 | 0 | 0 |
| K09960 uncharacterized protein | 227 | 212 | 238 | 249 | 16 | 19 | 12 | 14 | 13 | 9 | 18 | 15 |
| K09961 uncharacterized protein | 941 | 1003 | 1069 | 970 | 278 | 300 | 260 | 273 | 39 | 41 | 40 | 34 |
| K09962 uncharacterized protein | 889 | 878 | 823 | 841 | 475 | 477 | 467 | 487 | 274 | 255 | 207 | 246 |
| K09963 uncharacterized protein | 550 | 616 | 605 | 537 | 104 | 114 | 128 | 115 | 452 | 496 | 509 | 449 |
| K09964 uncharacterized protein | 861 | 784 | 766 | 788 | 1062 | 1038 | 1031 | 1077 | 743 | 670 | 728 | 638 |
| K09965 uncharacterized protein | 807 | 765 | 655 | 733 | 466 | 381 | 354 | 382 | 196 | 213 | 220 | 248 |
| K09967 uncharacterized protein | 474 | 452 | 393 | 450 | 605 | 671 | 570 | 596 | 36 | 38 | 48 | 38 |
| K09968 uncharacterized protein | 210 | 198 | 203 | 217 | 89 | 89 | 68 | 78 | 670 | 581 | 568 | 646 |
| K09973 uncharacterized protein | 946 | 866 | 954 | 952 | 2136 | 2029 | 2159 | 1976 | 2037 | 1924 | 1894 | 2021 |
| K09974 uncharacterized protein | 865 | 839 | 734 | 730 | 392 | 428 | 366 | 480 | 240 | 302 | 273 | 273 |
| K09975 uncharacterized protein | 1502 | 1534 | 1409 | 1644 | 944 | 1054 | 936 | 986 | 785 | 816 | 757 | 767 |
| K09976 uncharacterized protein | 16 | 12 | 10 | 5 | 2 | 4 | 3 | 3 | 6 | 7 | 6 | 11 |
| K09977 uncharacterized protein | 528 | 461 | 427 | 477 | 252 | 279 | 209 | 264 | 174 | 138 | 130 | 161 |
| K09978 uncharacterized protein | 685 | 597 | 645 | 647 | 370 | 322 | 314 | 348 | 442 | 484 | 443 | 426 |
| K09979 uncharacterized protein | 279 | 258 | 282 | 285 | 198 | 208 | 201 | 218 | 347 | 402 | 374 | 377 |
| K09980 uncharacterized protein | 980 | 997 | 1013 | 1053 | 629 | 518 | 517 | 599 | 341 | 364 | 377 | 313 |
| K09982 uncharacterized protein | 251 | 287 | 264 | 322 | 180 | 201 | 179 | 186 | 227 | 255 | 223 | 280 |
| K09983 uncharacterized protein | 610 | 623 | 570 | 605 | 342 | 330 | 303 | 351 | 223 | 183 | 176 | 182 |
| K09984 uncharacterized protein | 525 | 489 | 442 | 542 | 295 | 253 | 277 | 339 | 245 | 289 | 267 | 278 |
| K09985 uncharacterized protein | 0 | 0 | 0 | 0 | 2 | 2 | 0 | 2 | 0 | 0 | 2 | 2 |
| K09986 uncharacterized protein | 918 | 917 | 866 | 907 | 477 | 464 | 502 | 446 | 323 | 276 | 265 | 263 |
| K09987 uncharacterized protein | 0 | 0 | 0 | 0 | 2 | 0 | 0 | 0 | 0 | 0 | 5 | 0 |
| K09989 uncharacterized protein | 1626 | 1675 | 1607 | 1613 | 918 | 870 | 887 | 977 | 604 | 505 | 538 | 492 |
| K09990 uncharacterized protein | 2 | 0 | 0 | 1 | 0 | 0 | 0 | 0 | 3 | 0 | 3 | 0 |
| K09991 uncharacterized protein | 1 | 0 | 0 | 1 | 1 | 0 | 0 | 0 | 2 | 0 | 2 | 3 |
| K09992 uncharacterized protein | 76 | 84 | 65 | 69 | 11 | 19 | 27 | 27 | 147 | 180 | 130 | 120 |
